# Genomics of sable (*Martes zibellina)* × pine marten (*Martes martes*) hybridization

**DOI:** 10.1101/2025.08.01.668050

**Authors:** Andrey A. Tomarovsky, Azamat A. Totikov, Tatiana M. Bulyonkova, Polina L. Perelman, Alexei V. Abramov, Natalia A. Serdyukova, Aliya R. Yakupova, Dmitry Prokopov, Violetta R. Beklemisheva, Mikkel-Holger S. Sinding, Guzel Davletshina, Maria Pobedintseva, Ksenia Krasheninnikova, Daniel W. Foerster, Anna S. Mukhacheva, Alexandra Mironova, Michail Sidorov, Wenhui Nie, Jinhuan Wang, Svetlana A. Romanenko, Anastasiya A. Proskuryakova, Malcolm Ferguson-Smith, Fengtang Yang, Nikolay Cherkasov, Elena Balanovskaya, M. Thomas P. Gilbert, Innokentiy M. Okhlopkov, Anna Zhuk, Alexander S. Graphodatsky, Roger Powell, Klaus-Peter Koepfli, Sergei Kliver

**Affiliations:** Laboratory of Diversity and Evolution of Genomes, Institute of Molecular and Cellular Biology SB RAS, 8/2 Acad. Lavrentiev ave., Novosibirsk, 630090, Russia; Department of Natural Sciences, Novosibirsk State University, 1 Pirogova str., Novosibirsk, 630090, Russia; Youth Laboratory of Molecular Genetics, Yugra State University, 16 Ulitsa Chekhova, Khanty-Mansiysk, 628011, Russia; Laboratory for Theriology, Zoological Institute RAS, 1 Universitetskaya emb., St. Petersburg, 199034, Russia; Division of Evolutionary Biology, Ludwig-Maximilians-Universität, 2, Großhaderner str, Planegg, 82152, Germany; Microevolution and Biodiversity, Max Planck Institute for Biological Intelligence, Eberhard-Gwinner-Straße, Seewiesen, 82319, Germany; Centre for Haemato-Oncology, Barts Cancer Institute, Queen Mary University of London, London, UK; QMUL Centre for Epigenetics, Queen Mary University of London, London, UK; Center for Evolutionary Hologenomics, The Globe Institute, The University of Copenhagen, Copenhagen, Denmark; Department of Biology, The University of Copenhagen, Copenhagen, Denmark; Independent researcher, Wellcome Trust Genome Campus, Hinxton, Saffron Walden CB10 1RQ, United Kingdom; Leibniz Institute for Zoo and Wildlife Research (IZW), Alfred Kowalke Straße 17, 10315 Berlin, Germany; Sikhote-Alin Biosphere Zapovednik, 44 Partizanskaya str., Ternei, 692150, Russia; Laboratoire de Physiologie Cellulaire and Végétale, Univ. Grenoble Alpes/CNRS/CEA/INRA/IRIG, Grenoble, France; Institute of Biological Problems of Cryolithozone SB RAS, 41 Lenina ave., Yakutsk, 677000, Russia; State Key Laboratory of Genetic Resources and Evolution, Kunming Institute of Zoology, Chinese Academy of Sciences, Kunming 650223, China; Cambridge Resource Centre for Comparative Genomics, Department of Veterinary Medicine, University of Cambridge, Cambridge CB3 OES, UK; School of Life Sciences and Medicine, Shandong University of Technology, Zibo, China; Vavilov Institute of General Genetics, Moscow, Russia; Laboratory of human population genetics, Research Centre for Medical Genetics, Moscow 115522, Russia; Center for Evolutionary Hologenomics, The Globe Institute, The University of Copenhagen, 5A, Oester Farimagsgade, Copenhagen, 1353, Denmark; University Museum, NTNU, Trondheim, Norway; Institute of Applied Computer Science, ITMO University, 197101 St. Petersburg, Russia; Laboratory of Amyloid Biology, St. Petersburg State University, 199034 St. Petersburg, Russia; North Carolina State University); Smithsonian-Mason School of Conservation, 1500 Remount Road, Front Royal, VA 22630, USA

**Author notes:** equal contribution (Azamat A. Totikov, Tatiana M. Bulyonkova).

**Keywords:** sable, pine marten, interspecific hybridization, species divergence, conservation genomics

## Abstract

The sable (*Martes zibellina*) and pine marten (*Martes martes*) are two Palearctic mustelids with long-recognized hybrids (kidases), whose fertility was controversial for years. Early genetic studies confirmed hybrids beyond F1, but details remained unclear due to low-resolution methods. Both species were hunted for centuries, but anthropogenic pressures during the 20th-century caused severe bottlenecks in the sable followed by hunting bans and large-scale reintroduction programs across much of its range, including the sympatric zone, potentially affecting hybridization. We resequenced 30 individuals from most of the sables’ range and Eastern part of pine marten’s. Among samples, we found a broad spectrum of hybrid types with mosaic recombinant chromosomes that confirm hybrid fertility and indicate crossover is not suppressed in kidases. This necessitates re-evaluation of previous research, as we detected notable discrepancies between STR-based ancestry and whole-genome analysis. In pine martens, we revealed mitochondrial DNA introgression from sables, indicating displacement of native pine marten mitochondrial sequences. Pine marten heterozygosity is relatively low (∼0.5-0.6 hetSNPs/kbp) while sable diversity (∼1.5-1.8 hetSNPs/kbp) is unexpectedly high for a species with its demographic history, likely reflecting successful reintroduction programs. We dated species divergence at 1.52 (CI: 1.05-2.06) Mya and identified candidate genes associated with ecological, morphological, and dietary differences, as well as hybrid fertility issues. This study is the first to elucidate marten hybridization at the whole-genome level, opening new research directions for understanding hybridization among Holarctic martens, the genetic consequences of reintroduction programs, and comparative adaptomics.

## Introduction

Interspecific hybridization has played a significant role in the evolution of many mammalian lineages, including the Mustelidae family – a diverse group that encompasses ferrets, weasels, otters, badgers, and martens (Rozhnov et al. 2013; Colella et al. 2018; Kinoshita et al. 2019; Colella et al. 2021; Etherington et al. 2022). The sable (*Martes zibellina*) and pine marten (*Martes martes*) are two closely related but distinct species within genus *Martes* whose ranges partially overlap across Eurasia. In the sympatric zone, hybrids known as kidas (or kidus, Russian кидас/кидус) naturally occur, with mentions of them dating back to the 18th-19th centuries (Pavlinin 1963). According to data from the International Union for Conservation of Nature’s Red List of Threatened Species, this zone forms a wide band stretching from northwest to southeast across several regions of Russia: from the Komi Republic in the northwestern Urals through Khanty-Mansi Autonomous Okrug – Yugra, Tyumen Oblast, Omsk Oblast, Tomsk Oblast, and Novosibirsk Oblast (Herrero et al. 2015; Monakhov 2015a).

Although kidases have long been recognized in the fur industry, their official taxonomic and hybrid status has remained controversial for years. Morphologically, they exhibit a broad range of phenotypes, combining traits of both parental species. Yurgenson (1947) considered kidases to be interspecific hybrids, whereas Pavlinin (1963) challenged this view, suggesting that most individuals identified as kidases were actually highly variable sables or martens, with their morphological deviations arising within each species rather than as a result of hybridization (Yurgenson 1947; Pavlinin 1963). Grakov (1976) showed that in captivity, kidases were produced from male sables and female martens, whereas male martens and female sables yielded no offspring in the experiments (Grakov 1976). The backcrossing and intercrossing of kidases had limited success, causing some authors to claim that male F1 hybrids were infertile, thus limiting the introgression in the wild (Portnova 1941; Grakov 1974; Grakov 1981; Grakov 1993). However, these studies were based on a small number of crossing attempts, which involved few individuals (e.g., ∼10 in the Grakov, 1976 study), of unknown or poorly described ancestry. It is unclear whether the origin or features of the particular individuals affected the results or not. Nonetheless, based on these reports, some researchers hypothesized that the hybrid breakdown between the sable and the pine marten acts as a reproductive barrier, limiting its further expansion within each species’ range (Kassal and Sidorov 2013).

Generally, kidases possess mixed morphological traits inherited from both parent species (Figure 1). Traditionally, identification of kidases relies on tail length and coat coloration (Yurgenson 1947). Compared to pine martens, hybrid individuals often exhibit sable-like features, such as a shorter tail with fewer caudal vertebrae (typically 15–16, compared to 15–22 in pine martens), which barely extends beyond the hind feet (Monakhov 2011; Monakhov 2022). Their coat is more variably colored and often darker, with high-contrast banding of down hair, and the upper neck, ears and face tend to be paler. The throat patch is often reduced or completely absent, unlike pine martens, where it is normally prominent (Yurgenson 1947; Pavlinin 1963). Additional features inherited from sables may include shorter and darker guard hairs at the tail tip, paler face, and shorter guard hair on the rump and hind limbs. In winter, like the sable, kidases may develop elongated bristle hairs on their feet that facilitate movement in deep snow, as well as dense fur covering the footpads and toe pads, which provides protection against cold.

**Figure 1.**
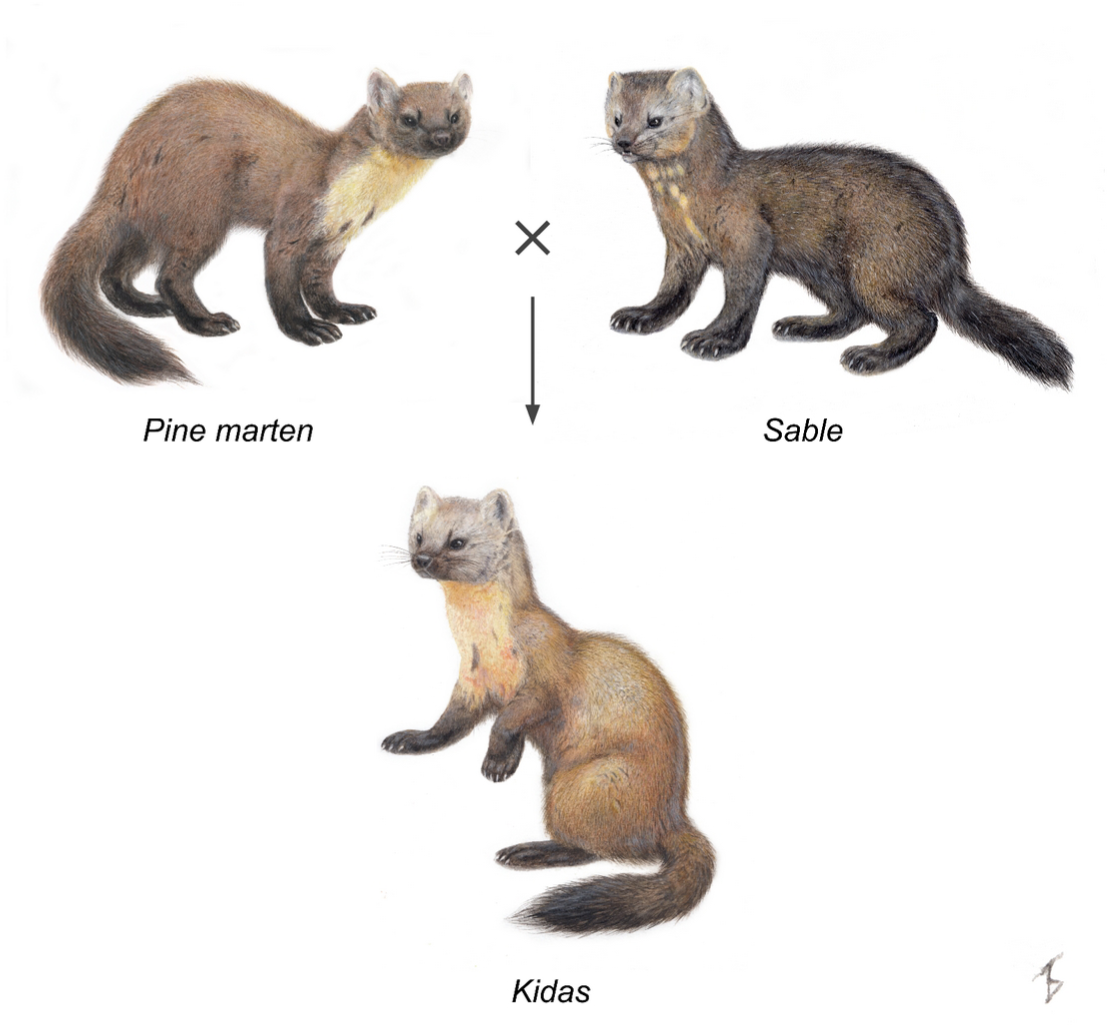
Sable (*M. zibellina*), pine marten (*M. martes*) and kidas (*M. martes* × *M. zibellina*). Illustrations by T. M. Bulyonkova.

While sable and pine marten show roughly similar head-body lengths in both males and females (Monakhov 2011; Monakhov 2022), a morphometric study showed that kidases occupy an intermediate position between the parental species, consistent with first-generation hybrids (F1), although the classification accuracy was limited to 62.5% (Monakhov and Uspenskaya 2013). Notably, the sable was considered vulnerable to genetic swamping due to this introgression. To improve identification, Monakhov (2021) introduced a craniometric index Δ (delta), defined as the distance from the postorbital constriction to a line connecting the postorbital processes in the sagittal plane (Monakhov 2021a). This index allowed for highly accurate discrimination between sables and pine martens, with over 97% classification accuracy. However, it remains unclear whether this index can reliably identify hybrids or assess the extent of introgression, prompting the need for molecular genetic analyses.

Human activities have significantly influenced the interaction between sables and pine martens.. Historically, both sable and pine marten furs were considered luxury goods and served as high-value natural currency for centuries. At the turn of the 20th century, the value of sable fur peaked due to a combination of declining harvest numbers and intense demand from the fashion industries in both Europe and North America. By 1925, prices soared to as much as $6,000 (12,000 rubles) per pelt – an equivalent of 8 kg/18 lbs of gold at the time (Shadiul and Vashukevich 2020). Overhunting for several centuries, beginning as early as the 17th century (Timofeev and Nadeev 1955), resulted in severe population declines, range reduction, and fragmentation, ultimately leading to a complete harvesting ban imposed in 1935-1940 (Mikhel 1938; Romanov 1941). Finally, a large-scale reintroduction program was implemented 1939-1941 and 1945-1970, with more than 18,000 individuals released throughout the original range of the sable (Timofeev and Nadeev 1955; Bobrov et al. 2008; Monakhov 2015b). However, harvesting for fur was not the only factor that affected the ranges and population dynamics of the two species. Most of the pine marten’s range overlaps with areas of relatively high human population density, where agricultural development and degradation of core forested habitats may have exerted significant pressure on populations. (Twining et al. 2020). Therefore, the diversity and population structures of both species were affected by anthropogenic activities, which must be taken into account while studying the hybridization between them.

Sable and pine marten have similar karyotypes (2n = 38), as do other closely related species from the subgenus *Martes* (Koepfli et al. 2008; Law et al. 2018; Hassanin et al. 2021): the Pacific marten (*M. caurina*), American marten (*M. americana*), Japanese marten (*M. melampus*) and the stone marten (*M. foina*). In particular, their chromosomes share similar morphologies, G-banding patterns and low heterochromatin content (Hsu and Benirschke 1971; Graphodatsky et al. 2020; Beklemisheva et al. 2023). A recent comparative chromosome painting study confirmed a one-to-one synteny and absence of large rearrangements among chromosomes of the sable, pine marten and stone marten, but highlighted differences in macrosatellites that form heterochromatin blocks (Beklemisheva et al. 2023). These findings suggest that no cytogenetic barriers prevent hybridization between these species.

Early genetic studies of sable-pine marten hybridization based on mitochondrial DNA (mtDNA) fragments revealed bidirectional introgression between *M. zibellina* and *M. martes* in sympatric populations of the Northern Urals (Rozhnov et al. 2010). Importantly, species-specific mtDNA haplotypes often do not correspond to phenotypic species assignments, even when external morphological traits appear unambiguous (Rozhnov et al. 2010). The distinctiveness of sable and pine marten mitotypes has been clearly established (Rozhnov et al. 2010; Kassal and Sidorov 2013), and the detection of pine marten individuals carrying sable mtDNA – despite earlier assumptions that such introgression was impossible based on results from captive breeding experiments – provided compelling evidence of natural hybridization (Rozhnov et al. 2010). These findings were supported by further reports of mtDNA introgression between the two species (Davison et al. 2001; Pishchulina 2013), suggesting the fertility of at least female hybrids (Pishchulina 2013). Later, the classification of hybrid individuals using microsatellite loci was conducted in several studies, confirming the persistence of hybridization within the zone of sympatry (Pishchulina 2013; Rozhnov et al. 2013; Modorov et al. 2020; Ranyuk et al. 2025). However, only a small number (up to 11) of microsatellites (STRs) was used, and their distribution on chromosomes was unknown.

To date, despite numerous studies based on morphological traits and molecular markers, the hybridization between sables and pine martens has not been studied at the whole-genome level. The key reason lies in the absence of high-quality genome assemblies not only for the genus *Martes*, but also for the Guloninae lineage within the family Mustelidae. Until 2021, only a highly fragmented assembly of the wolverine (*Gulo gulo*) was available (Ekblom et al. 2018). In recent years, assemblies of several marten species have appeared (O’Brien and Januszczak 2024), including those generated with long read and Hi-C sequencing (Liu et al. 2020; Tomarovsky et al. in prep).

In this work, we present the first detailed analysis of interspecific hybridization between sables and pine martens. We resequenced 30 individuals of non-hybrids, as well as putative hybrids from the zone of sympatry. We performed detailed analyses of ancestry using multiple approaches, assessed genetic diversity and its spatial distribution, and integrated our results with previous studies.

We also revealed the genomic legacy of the sable reintroduction program and assessed its results. Finally, we dated the divergence between sable and pine marten and revealed previously unknown genomic features in the two species.

## Results

### Chromosome-length genome assemblies, resequencing data, and macrosynteny

We resequenced twelve sables, six pine martens, and twelve putative hybrids from the sable-pine marten sympatric zone (Figure 2, Supplementary Table ST1). Combined with data from previously sequenced sable (sample CHN) (Liu et al. 2020) and two reference individuals (Tomarovsky et al. in prep), our final dataset reached 33 samples. The analysis of 23-mer distributions (Supplementary Figure SF1) showed no significant differences in genome size among the pine martens, sables, and putative hybrids. We observed the smallest genome sizes for the samples T72 (2.39 Gbp), 10xmzib, and T18 (2.40 Gbp), and the largest for S50 (2.54 Gbp), respectively (Supplementary Table ST2). Sample T87 showed an additional k-mer peak (at a half coverage) on the distribution, indicating a high heterozygosity level.

**Figure 2.**
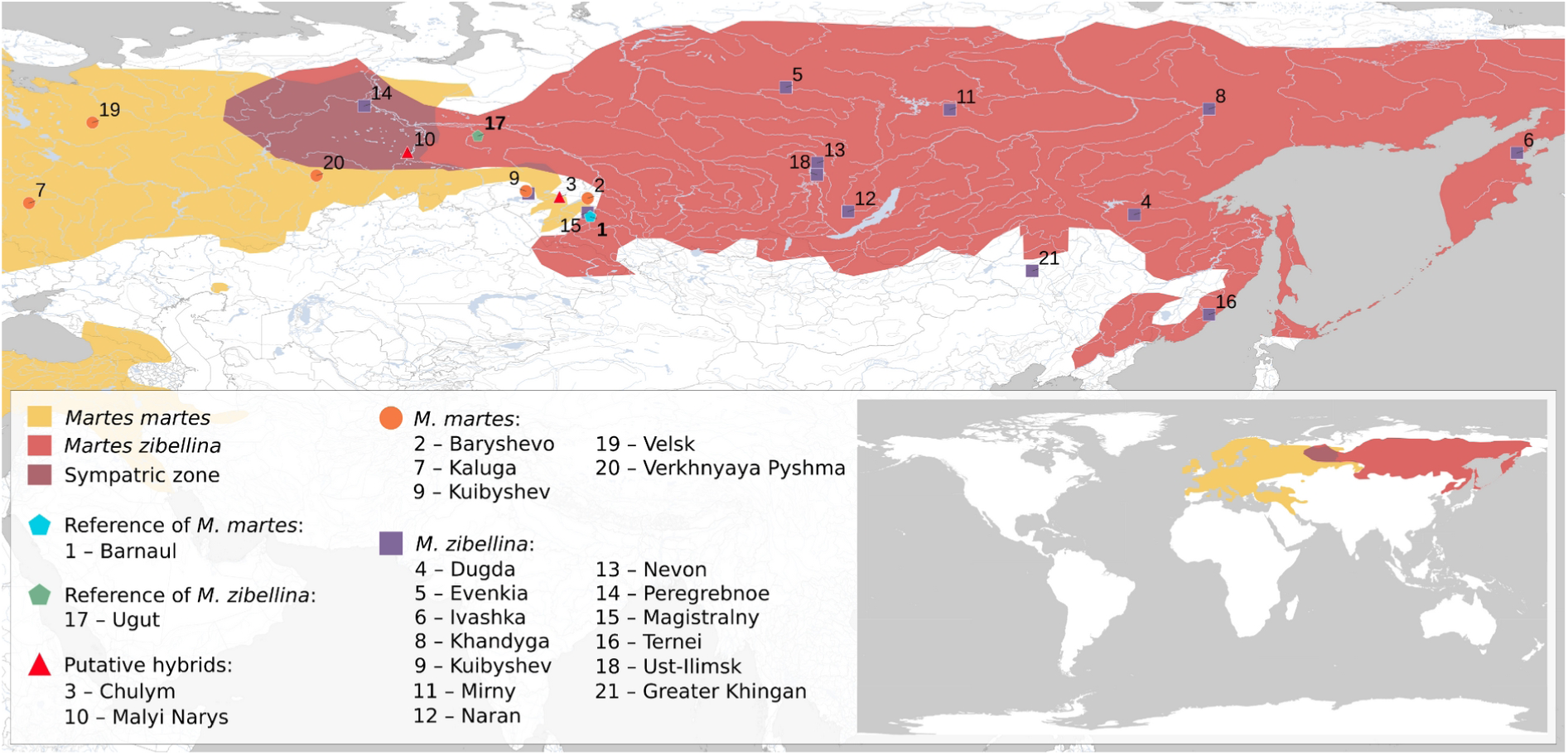
Geographic ranges of *M. zibellina* and *M. martes* along with the locations of samples used in this study. The pine marten range is shown in yellow, the range of the sable – in red, the sympatric zone – in maroon. Markers show sampling locations. Location codes correspond to the column “Point on map” in Supplementary Table ST1. Small map in the right bottom corner shows full (uncut) areas from the IUCN Red List of Threatened Species (2024-2) (Herrero et al. 2015; Monakhov 2015a)

### Divergence times

To estimate the divergence time between *M. martes* and *M. zibellina*, we used clockwise sites extracted from a codon alignment (1,384,523 bp) generated with 5,989 single-copy orthologs and fossil-based calibration priors for five nodes to reconstruct a chronogram that included six other mustelid species (Supplementary File <u>SF1</u>). We tested three types of molecular clocks: global, independent and correlated (Supplementary Table ST3). The global clock provided the youngest dating for all nodes except the root. For example, the Most Recent Common Ancestor (MRCA) of the sable and pine marten was dated at 1.35 (Confidence Interval (CI): 1.31 - 1.39) Mya versus 2.02 (CI: 1.53 - 2.58) Mya and 1.52 (CI: 1.05 - 2.06) Mya for the correlated and independent clocks, respectively. We also observed very narrow confidence intervals provided by the global model (Supplementary Figure SF2A) and decided to discard it. For the correlated (Supplementary Figure SF2B) and independent (Figure 3) clocks we obtained very similar (∼5% younger for the independent clock) datings for nodes 5, 6 and 7, whereas the discrepancy was significantly higher for the Guloninae lineage (up to 33%). Based on the distribution of the fossil calibrations and heterogeneity of substitution rates, we decided to rely on the estimates from the independent rate model (Figure 3).

**Figure 3.**
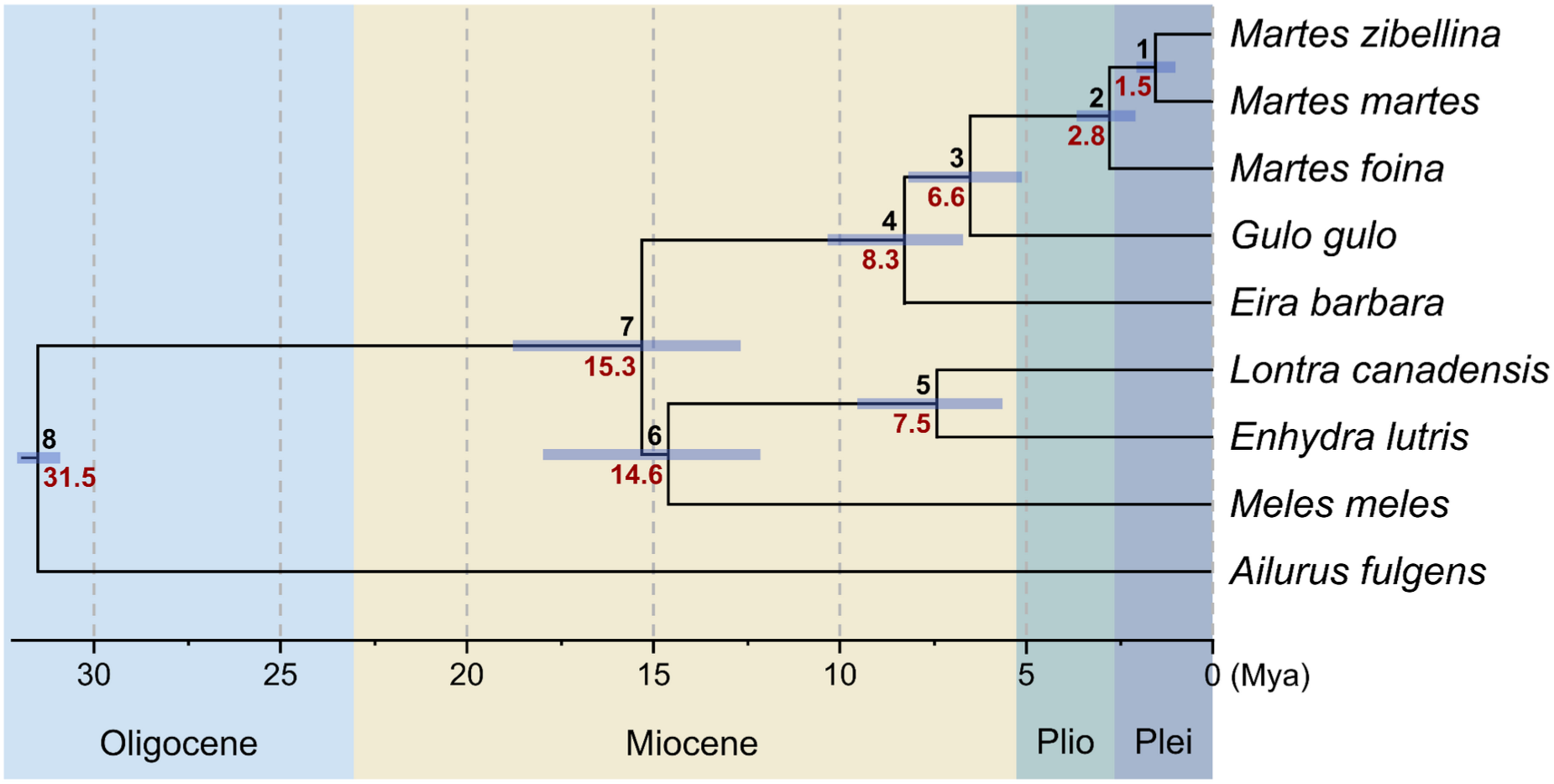
Time-calibrated phylogenetic tree based on the independent molecular clock model. Branch lengths are proportional to time, with divergence dates shown in millions of years. For each node, the node ID is displayed above the branch, and the estimated divergence time (Mya) is shown below. Node bars indicate the 95% Confidence Intervals (CIs) for node age uncertainty. Geological epochs: Plio – Pliocene, Plei – Pleistocene

### Genetic structure and ancestry

To explore genetic structure and ancestry proportions, we conducted principal component analysis (PCA) and global (whole genome) and local (independent admixture runs in 1 Mbp sliding windows with 100 kbp step) admixture analyses using the resequencing data. We further analyzed ancestry using known STR loci, mitochondrial genomes, and compared the morphology of putative hybrids with parental species, building on previous studies that applied these markers or characters. All analyses showed no reference-related biases, with results remaining consistent regardless of which reference genome assembly (sable or pine marten) was used (see Supplementary File <u>SF2</u> for those based on the pine marten reference).

### PCA and admixture of WGS data

PCA revealed two distinct clusters distinguishing sable (right) and pine marten (left) samples, with samples T87 and T84 located between these clusters, elucidating them as potential F1-like hybrids (Figure 4A, Supplementary Figure SF3). We also note two samples with a specific location on the plot. The first, T18, is located between the two hybrid samples and sable cluster, whereas the second, T72, is part of the sable cluster, but is an outlier located at the top right corner of the plot. We consider Principal Component 1 (PC1, 30.27 % of the variance) to be associated with interspecies differences, whereas PC2 (5.59 %), is associated with variation within the sable. Interestingly, the distribution of the sables along PC2 partly follows a longitudinal pattern: the further east the sample was gathered, the higher it is located on the figure. This pattern is consistent for samples collected from Eastern Yakutia to Kamchatka, as well as the CHN sample from China, (PC2 > 0), but the remaining samples (PC2 < 0), from Central Siberia and Western Yakutia, are mixed.

**Figure 4.**
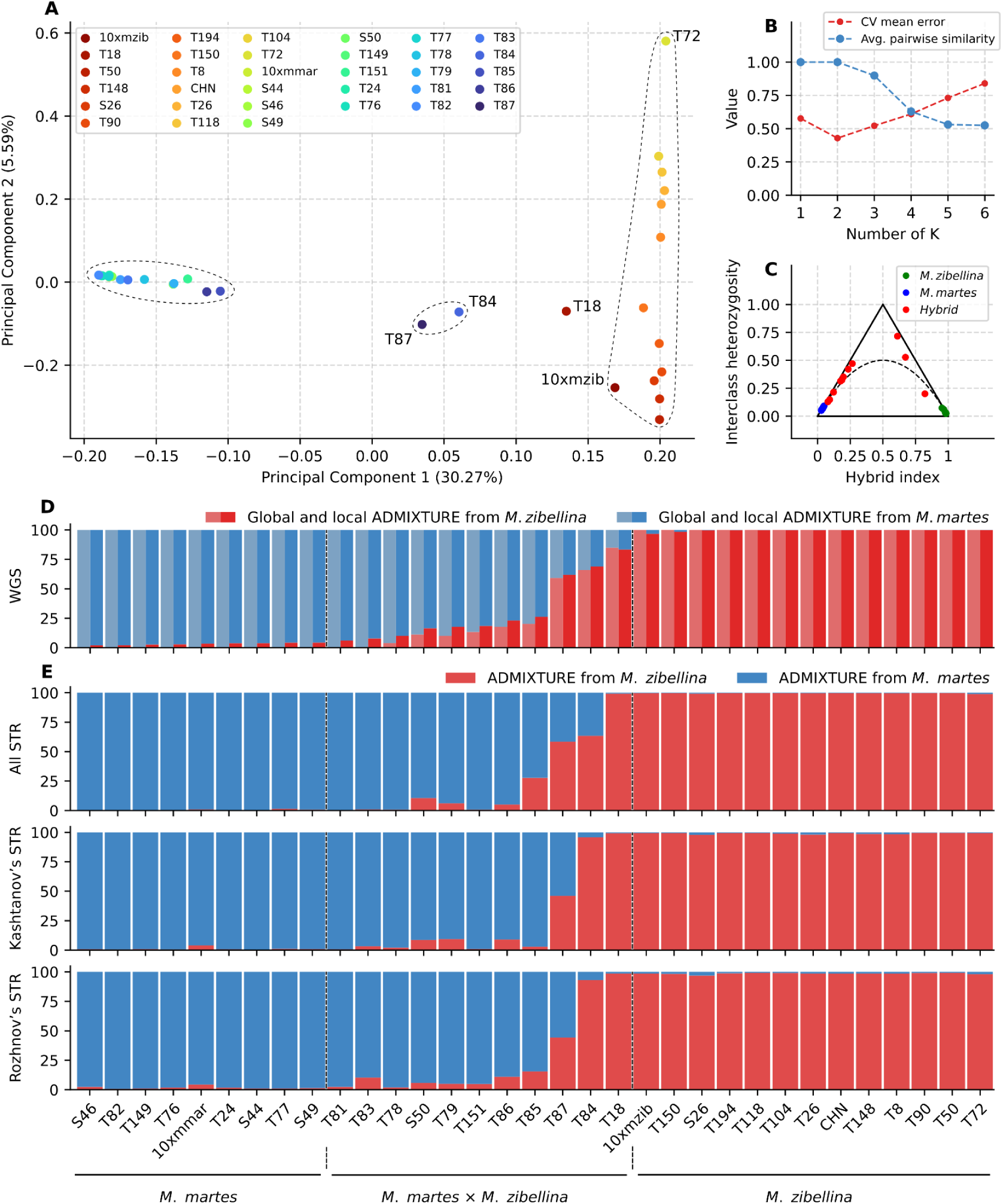
Principal component analysis and ancestry proportions of pine martens, sables, and putative hybrids. A – PCA plot based on 6,750,722 SNPs obtained from autosomes and the pseudoautosomal region (PAR). The dashed ellipses indicate clusters of pure pine martens and hybrids with a small fraction of sable (left), pure sables (right) and hybrids close to F1 (middle). Sample T18 was located outside of the clusters. The percentage of variance explained by the corresponding component is indicated in parentheses; B – cross-validation error (blue) and average pairwise similarity (red) based on global admixture. C – Triangle plot with allele frequency difference threshold (δ) of 0.75. D – Global admixture (left half) and local admixture (right half) based on the WGS data. Samples are labeled along the X-axis while the Y-axis represents the mean probability of ancestry. E – Admixture analysis for three sets of STR loci. All STRs – combined set of all mapped STRs loci, Rozhnov’s STRs – mapped markers from (Rozhnov et al. 2013), Kashtanov’s STRs – markers from (Kashtanov et al. 2022)

Global and local admixture analyses (Figure 4D) supported two clusters (K = 2) based on cross-validation error assessment (Figure 4B). Global analyses revealed admixture in 9 samples (including those initially classified as pure species): T18, S50, T151, T78, T79, T84, T85, T86 and T87, with admixture levels ranging from 3.9% (T78) to 40.8% (T87), while remaining samples showed no signs of introgression. Local analysis detected admixture in two additional samples (T81 and T83) at levels of 6.17% and 7.86%, respectively (Supplementary Table ST4), with accuracy confirmed by notable hybrid and sable components in the heterozygosity analysis (Supplementary Figure SF4). We found admixture signals between 1% and 5% in several other samples, including the reference genome individuals of both species: 10xmzib, 10xmmar, T150, S44, S46, S49, T149, T24, T76, T77, and T82, while remaining samples showed only small introgression traces (<1%). Using a 5% threshold to distinguish pure species ancestry from hybrid/introgressed individuals, we divided samples into three groups: pure sables (13 individuals: 10xmzib, S26, T8, T26, T50, T72, T90, T104, T118, T148, T150, T194, and CHN), pure pine martens (9 individuals: 10xmmar, S44, S46, S49, T149, T24, T76, T77, and T82), and hybrid or introgressed individuals with ≥5% introgression (11 individuals). The third group was further subdivided into F1-like (T87, 35–50% introgression), backcross-like (S50, T79, T151, T86, T85, T84, T18, 15–35% introgression), and atypical sables or martens (T81, T83, T78, 5–15% introgression).

We complemented PCA and admixture analyses with HyDe, F3, D and F4 statistics using the three-group classification of resequenced individuals based on the local admixture results (Supplementary File <u>SF3</u>). HyDe revealed a significant level of the hybridization with γ = 0.19, (Z = 224.47, p-value << 0.01). Individual tests of the samples from the hybrid group resulted in a broad range of γ values: from 0.02 (T81) to 0.74 (T18). Samples T84 and T87 were the closest to the 50/50 ratio, with γ of 0.53 and 0.47 (Z-scores of 289.06 and 304.55). The F3 metric calculated for the hybrid group (−0.16, Z = -90.9) showed a pattern similar to the HyDe results. Individual F3 tests showed statistically significant admixture (|Z| > 3) in all hybrid samples, with four samples exhibiting particularly extreme F3 values: T87 (−0.21), T85 (−0.18), T84 (−0.17) and T86 (−0.17). Values from D and F4 statistics were concordant and confirmed gene flow (|Z| > 3) into hybrid individuals from both sable and pine marten.

### STR markers: localization and admixture

Using an *in silico* PCR approach, we identified the locations of previously described STR (Short Tandem Repeat) loci for various Mustelidae species (Davis and Strobeck 1998; Fleming et al. 1999; Domingo-Roura 2002; Vincent et al. 2003; Basto et al. 2010; Natali et al. 2010), and checked if it was possible to 1) genotype them in our samples and 2) if so, to evaluate their performance in detecting admixture. Out of a total of 79 markers, only 36 and 44 loci were suitable for STR-genotyping from our reads (150 bp) using the sable and pine marten assemblies, respectively (Supplementary File <u>SF4</u>). In addition to the full set of STR markers, we also used two subsets called the Kashtanov’s (Kashtanov et al. 2022) subset and the Rozhnov’s (Rozhnov et al. 2013) subset (see Methods and Supplementary Method <u>SM1</u> for details). The full set of markers map to all chromosomes except chr11, chr16, and chrX (Supplementary Figure SF5A) with a mean density of 1.9 STRs/chr (STR markers per chromosome). Rozhnov’s and Kashtanov’s subsets have significantly lower coverage: the first maps to only 6 out of 19 chromosomes (0.4 STRs/chr, Supplementary Figure SF5B), while the second maps to 11 out of 19 (0.7 STRs/chr, Supplementary Figure SF5C), respectively.

Admixture analysis showed a notable difference between sets for classification of some hybrid martens (Figure 4E, Supplementary File <u>SF5</u>). For example, sample T85 was shown to have 27.8% of sable according to the full set of STRs, but for the Kashtanov and Rozhnov subsets, the values are significantly lower – 2.9% and 15.5%, respectively. Moreover, sample T84 showed an ∼30% difference between the full set (63.4% of the sable) and the two subsets (95.8% and 93.1%). For the rest of the hybrid martens, the difference is not as dramatic, but higher than that for other samples. Compared to whole-genome sequencing analysis, STR-based results for hybrid samples revealed significant underestimation of the extent of admixture.

### Mitochondrial genome sequences

A previous mtDNA-based phylogeographic study of martens (Li et al. 2021) was focused on sables (∼100 individuals) and included only a few pine martens as an outgroup. We expanded this dataset by adding our 32 samples and mtDNA sequences assembled from published raw reads (Liu et al. 2020; Manakhov et al. 2021), resulting in 140 total samples (Supplementary File <u>SF6</u>). The addition of new data improved coverage of previously under-sampled lineages within the C1 and A1 clades (Supplementary Figure SF6, Figure 5). We reconstructed a median-joining haplotype network (Figure 5) and maximum likelihood phylogenetic tree (Supplementary Figure SF6) using this dataset. The obtained topologies of both the network and tree include three major clades containing sables (A, B, and C), along with an outgroup consisting of pine martens, which agrees with previously reported results (Li et al. 2021). However, some of the minor clades (A1, B1 and C1) are non-monophyletic, and we propose to divide them into smaller subgroups. Clade A1 was split into three subgroups (A3, A4, A5) while clades B1 and C1 were each split into two subgroups (Figure 5, Supplementary Figure SF6): B3, B4, C3 and C4, respectively. We found no clear correlation between sable haplogroups and designated subspecies (inferred from the sampling location). Among our samples, only T72 clustered with its formal subspecies *M. z. kamtschadalica*. Finally, we note that only three of our samples (two pine martens S44, S46 and a sympatric marten T78) had pine marten mtDNA. The remaining samples, even those identified as pine martens based on location and morphology, have sable mtDNA representing different haplogroups.

**Figure 5.**
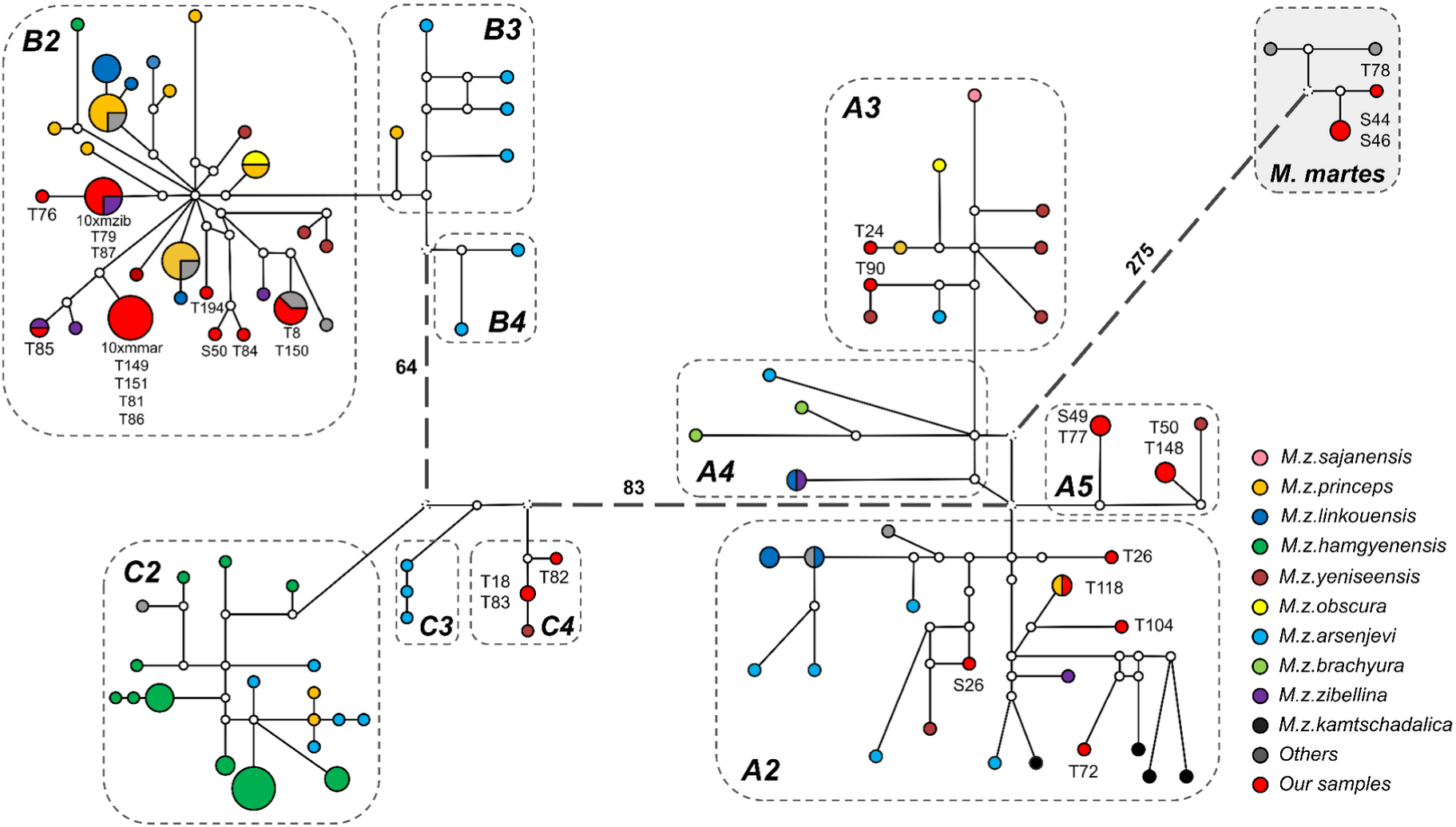
Median-joining network of mitogenomes haplotypes. The sizes of the circles are proportional to the haplotype frequency, and the distances between haplotypes are relative to the number of the substitutions between haplotypes. *Martes zibellina* subspecies are denoted by different colors (see legend). Substitutions between three main clades (A, B and C) are shown as a number. *M. foina* (NC_020643.1) was used as an outgroup (not shown). See Supplementary File <u>SF7</u> for a raw haplotype network with all distances labeled.

### Morphology of putative hybrids compared with parental species

All samples collected from putative hybrids (except T86, which was not analyzed) came from animals that had long tails protruding far beyond the hind legs. The throat patches were large, yellowish-red or white. These features are characteristic of the pine marten. Only in specimen T87 was the throat patch smaller and divided into several blotches. The heads of the examined putative hybrids were concolorous with the body, a feature typical for the pine marten, while the sable usually has a paler upper neck, ears and face. Multivariate analyses of craniometric characteristics failed to identify the species status of putative hybrids. They were morphologically closer to *M. martes* than to *M. zibellina*. According to Monakhov (2020), the average morphometric character Δ for sable is 4.67 ± 0.07, whereas that for pine marten is 7.81 ± 0.09. Our putative hybrids have a Δ = 6.82 (5.76 - 9.37), while specimen T87 has a Δ = 7.1.

### Heterozygosity

The total number of heterozygous SNPs (hetSNPs) varied greatly among our samples: from 2.06 million (10xmmar) to 9.09 million (T87) hetSNPs (Supplementary Table ST5). By analyzing the distributions of hetSNPs (counted in 1 Mbp windows with 100 kbp steps), we detected four major modes of heterozygosity, present in various combinations in different individuals (Figure 6A, Supplementary Figure SF7). By fitting a linear combination of negative binomial distributions to our data (see Supplementary Method <u>SM2</u> and Supplementary Results and Discussion (<u>SRD</u>) for the details), we resolved components of heterozygosity for all individuals and estimated corresponding mean values (Supplementary Figure SF4, <u>SRD</u> Figure 2, and Supplementary File <u>SF8</u>). The first component (R) at 0 hetSNPs/kbp corresponds to regions with low heterozygosity (potential segments of runs of homozygosity, RoH) and is found in multiple samples (Figure 6A). The second component (P), with a median value 0.57 hetSNPs/kbp is a characteristic feature of samples initially classified as pine martens, but is also present in sympatric martens. The third component (S, median value 1.70 hetSNPs/kbp) was detected in all sable samples and the putative hybrid martens T84 (comparable level to a priori sables) and T87 (significantly lower level). Finally, the last component (H, median value 4.51 hetSNPs/kbp) is mostly associated with the putative hybrids (T78, T79, T85, T86, T84, T87) and shows a dramatic expression in samples T84 and T87. However, H is also present in several sables (T18) and pine martens (S50 and T151). Densities of the hetSNPs along the chromosomes indicate that windows belonging to a particular component form long continuous runs (Figure 6, Supplementary File <u>SF9</u>). The relationship between the hybrid index and interclass heterozygosity (Figure 4C) showed a pattern consistent with the PCA analysis. At the selected allele frequency difference threshold (δ) of 0.75, a total of 502,893 Ancestry-Informative Markers (AIMs) were identified. The arrangement of the samples, forming a characteristic “arc”, rules out the isolation-by-distance model and supports the neutral diffusion model (Hewitt 1988). The dramatic difference in heterozygosity between the sable and pine marten is statistically significant. We confirmed it for both the median heterozygosity (0.64 vs 1.73 hetSNPs/kbp, one-sided Mann-Whitney p-value = 0.00005) and the corresponding heterozygosity components (mean P=0.57 vs mean S=1.9 hetSNPs/kbp, Mann-Whitney p-value = 0.00005).

**Figure 6.**
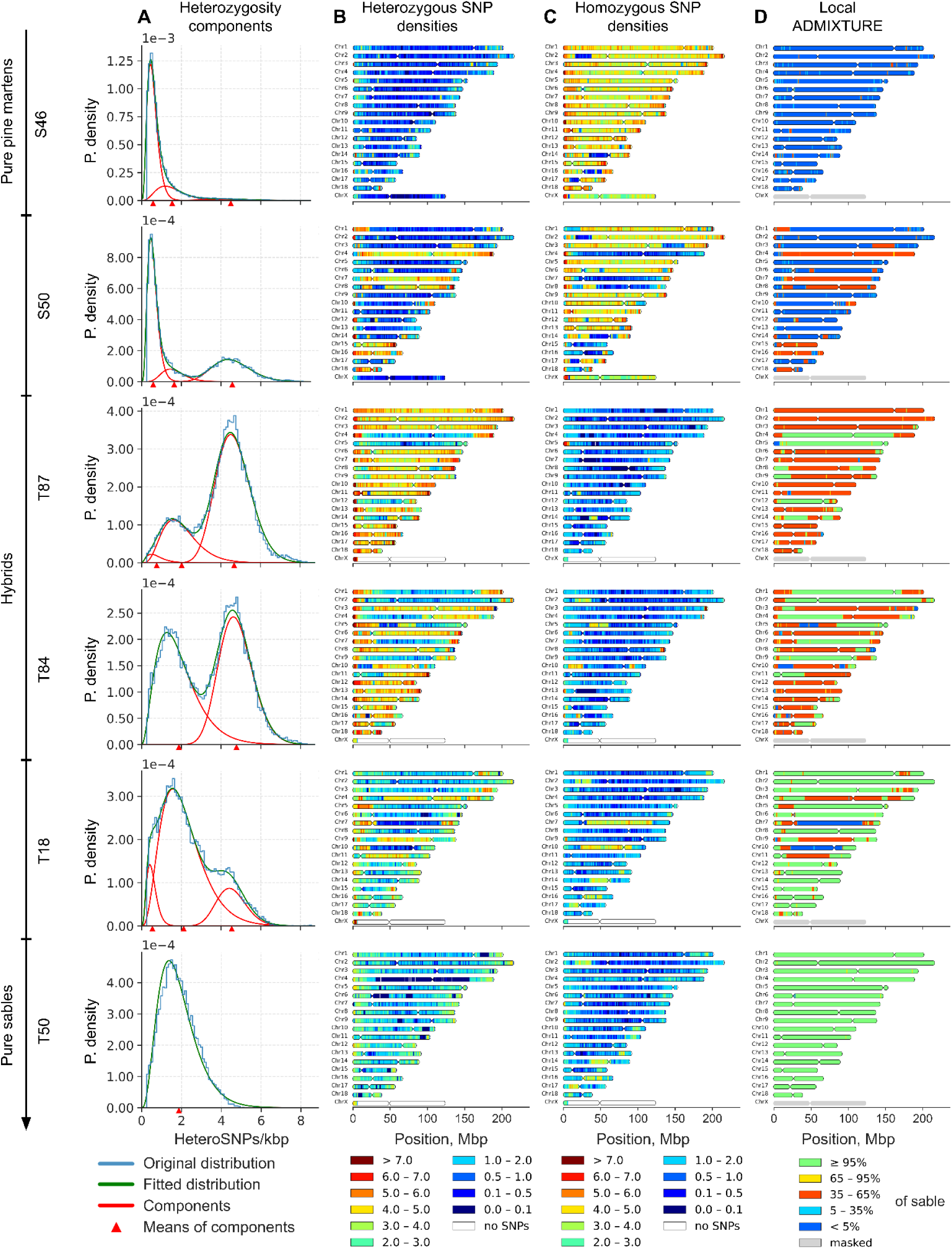
Distribution of heterozygous SNPs in representative samples showing the transition among pine martens, sables, and putative hybrids from the sympatric zone. A – Probability density of heterozygous SNPs/kbp and their fitted components. Blue – original distribution, green – fitted distribution, red – distribution of components with their means (red triangles); B, C – Distribution of heterozygous and homozygous SNP density along chromosomes, respectively. hetSNPs were counted in 1 Mbp windows with a 100 kbp step size and scaled to hetSNPs/kbp (represented by the color scale on the bottom, ranging from dark blue (extremely low heterozygosity) to brown (very high heterozygosity), with 0 to 0.1 and over 7 hetSNPs per 1 kbp respectively). Distributions of the remaining samples are presented in Supplementary File <u>SF9</u>. Reference: *M. zibellina*; D – local admixture (% of *M. zibellina*) on chromosomes (green – *M. zibellina*, blue – *M. martes*, red – hybrid) of 1 Mbp sliding windows with 100 kbp step. Local admixture for the other samples is presented in the Supplementary File <u>SF10</u>

### Runs of Homozygosity (RoH)

Among the samples, we observed significant variation in both the distribution and size of RoH, which we categorized as ultra long (L ≥ 10 Mbp), long (10 Mbp > L ≥ 1 Mbp), and short (L < 1 Mbp) (Supplementary Figure SF8, Supplementary Table ST6). Two obvious outliers emerged based on the fraction of the genome occupied by ultra long RoH: samples 10xmzib (24.1%) and 10xmmar (11.8%), with some of 10xmzib’s ultra long RoH exceeding 100 Mbp in length and nearly completely covering several chromosomes (chr3, chr6, chr11, and chr12), while in 10xmmar only chromosome 3 exhibited a similar pattern (Supplementary Figure SF8E-F). Cumulative distributions of RoHs (Figure 7B-D) followed different trajectories between species, with pine martens showing higher, convex or linear patterns, while sables displayed lower, concave distributions, indicating that short and long RoH (< 10 Mbp) occupy smaller genomic fractions in sables than pine martens (two-sided Mann-Whitney test, p-value = 0.0011), though no significant differences were detected for ultra long RoH (p-value = 0.12) or all RoH combined (p-value = 0.07). Sample T72 from the Kamchatka Peninsula showed a pine marten-like trajectory despite being a pure sable, while hybrid samples exhibited lower RoH numbers compared to pure species with varied trajectories depending on admixture rates, particularly samples T87 and T84 which had significantly fewer RoH (124 and 154) with shorter total lengths (34.06 and 53.88 Mbp).

**Figure 7.**
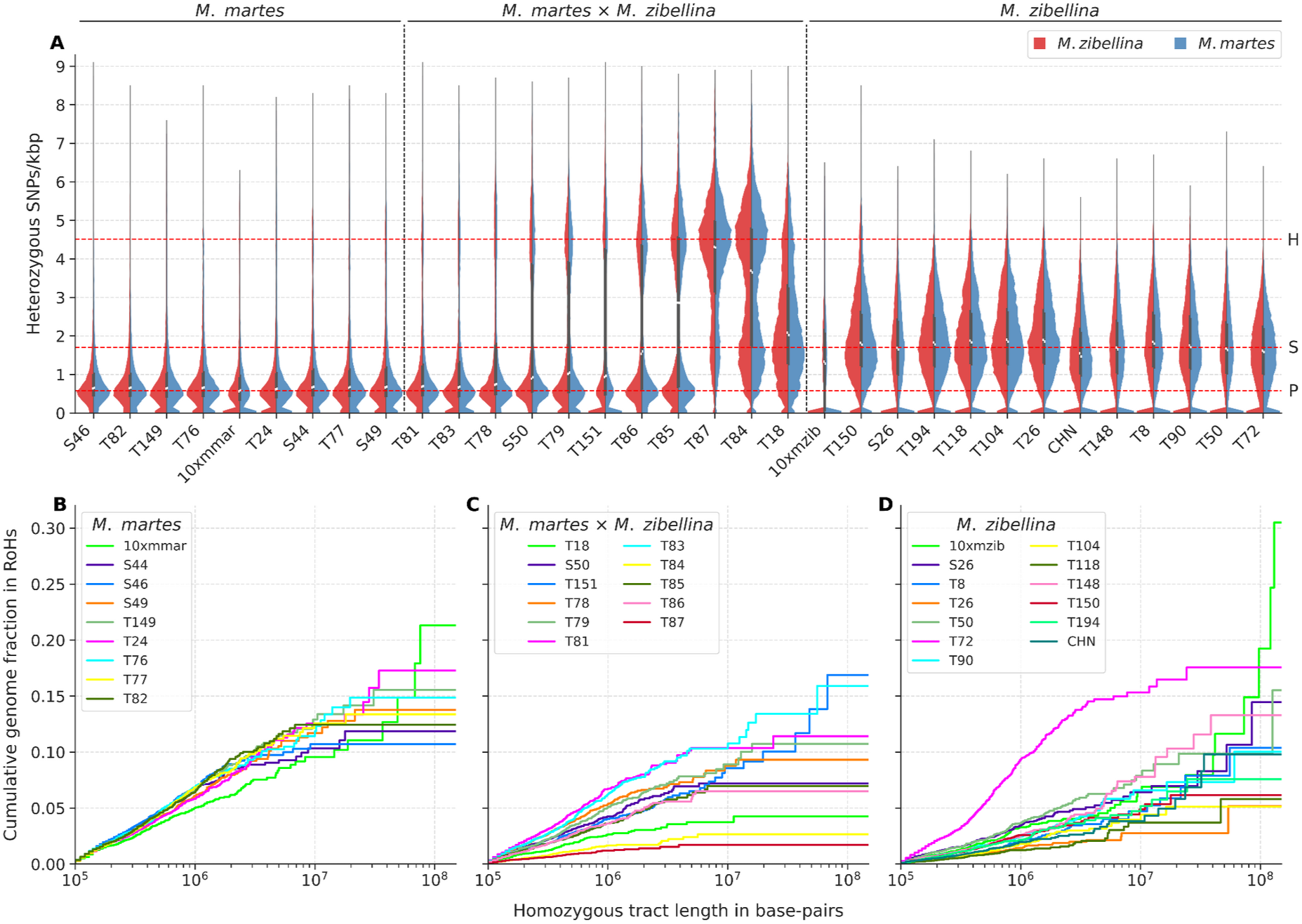
Heterozygosity and cumulative distributions of runs of homozygosity. A – Distribution of mean heterozygosity (SNP only) of the individuals counted in sliding windows of 1 Mbp with 100 kbp step size for both *M. zibellina* (left half in red) and *M. martes* assemblies (right half in blue). Each distribution was scaled with respect to the x-axis. Horizontal red lines indicate three levels of heterozygosity (0.57, 1.7 and 4.5 hetSNPs/kbp) corresponding to the median values (among the samples) of means for pine marten (P), sable (S) and hybrid (H) components. B, C and D – Cumulative distribution plots of RoH for *M. martes* (B), hybrids (C) and *M. zibellina* (D). Homozygous tract lengths and cumulative genome fraction in RoH are represented on X and Y axes, respectively. Tracts are ordered from shortest to longest. X chromosomes were excluded from all samples. RoH on chromosomes is presented in the Supplementary File <u>SF11</u>

### Demographic history

Pure sables and pine martens as well as hybrids showed distinct demographic trajectories (Figure 8). Sables (Figure 8A) showed an oscillating curve with three peaks at ∼2.5 (CI: 1.57 – 3.94), ∼0.5 (CI: 0.31 – 0.79), and ∼0.1 (CI: 0.06 – 0.15) Mya. In contrast, pine martens (Figure 8A) showed completely different curves: starting around 5 (3.15; 7.89) Mya their numbers increased and peaked at ∼1.3 (CI: 0.82 – 2.05) Mya. This was followed by a severe bottleneck overlapping with the Mid-Pleistocene Transition (MPT), which dropped N_e_ five-fold at ∼0.7 (CI: 0.44 – 1.1) Mya. Since then, the effective population size of pine martens has been slowly decreasing, with a small peak at ∼125 (CI: 79 – 197) kya. The hybrids (Figure 8B) revealed an interesting pattern: at ∼1.5 (CI: 0.94 – 2.37) Mya their trajectories curve steeply upward. We found that the strength (height of the peak) of this artificial “population explosion” strongly correlates (Kendall’s τ = 0.67, p-value = 0.003; Spearman’s ρ = 0.76, p-value = 0.006; Pearson’s r = 0.778, p-value = 0.005) with the introgression level in the hybrids (Supplementary Figure SF9). Finally, we note that until ∼3.5 (CI: 2.20 - 5.52) Mya all samples share nearly the same and growing trajectory (Figure 8A, zoomed on Supplementary Figure SF10).

**Figure 8.**
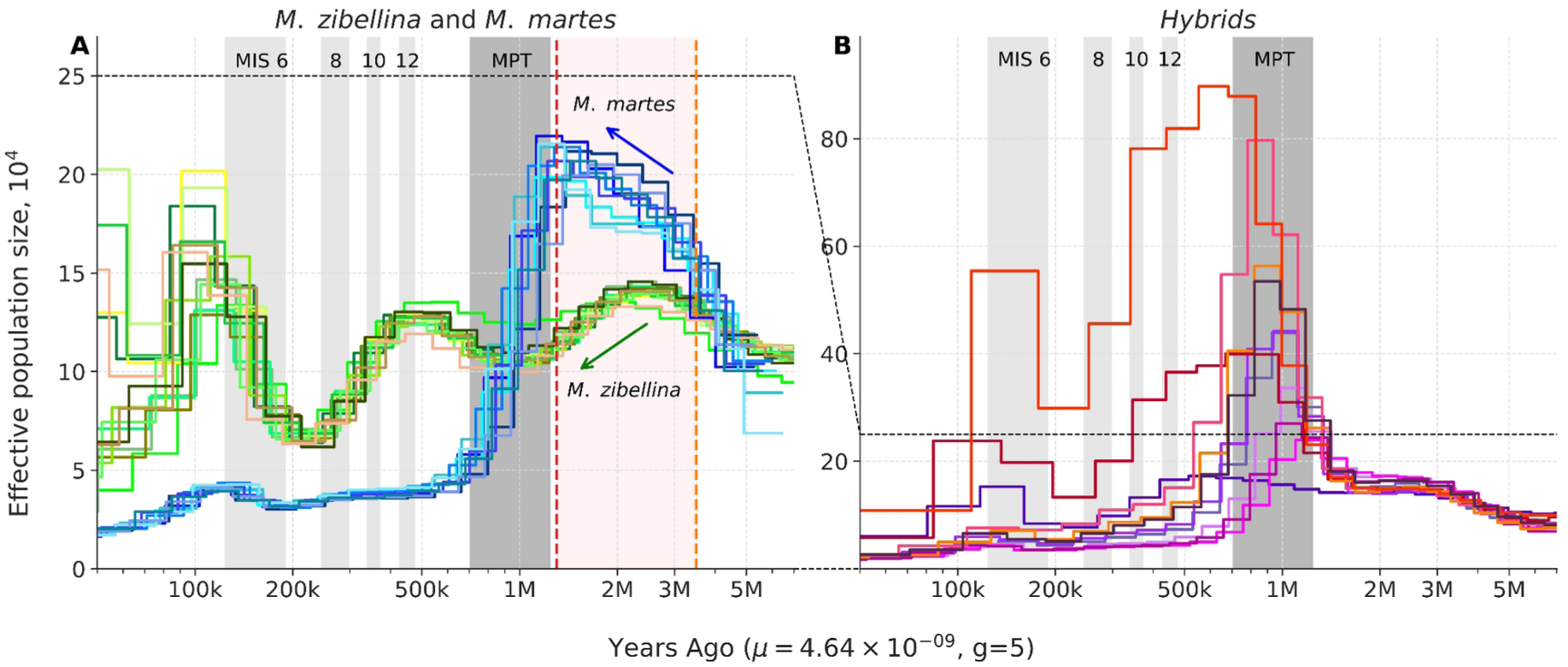
Demographic histories of *Martes zibellina*, *M. martes*, and hybrids. A – *M. zibellina* (green) and *M. martes* (blue), B – hybrids. Mutation rate (*μ*) = 4.64 × 10^-9^. Generation time (g) = 5. Abbreviations: MIS – Marine Isotope Stages (only four even, i.e. cold, stages are highlighted): 6 (130-191 kya), 8 (243-300 kya), 10 (337-374 kya), 12 (424-478 kya); MPT – Mid-Pleistocene Transition (0.7-1.25 Mya). Red and orange lines denote the period preceding species divergence (1.3-3.5 Mya). The blue and green arrows indicate the directions of demographic trajectory changes during this pre-divergence period for *M. martes* and *M. zibellina*, respectively. The X chromosome was excluded from the analysis

The demographic trajectories also provided some clues for dating the divergence between sable and pine marten. It should occur between the start of the N_e_ decline in the pine marten and the end of the shared with the sable trajectory. Using the mutation rate *μ* = 4.64 × 10⁻⁹ (Bergeron et al. 2023), the divergence event is estimated to have occurred between 1.3 and 3.5 Mya. Taking a more conservative approach and incorporating the confidence interval boundaries of the mutation rate (2.94 × 10⁻⁹ – 7.37 × 10⁻⁹) expands this estimate considerably to between 0.82 and 5.52 Mya. However, the divergence of demographic trajectories is expected to precede the actual speciation event by a substantial margin. We previously observed similar demographic trajectory differences within the least weasel (comparable to the 1.3-3.5 Mya divergence between sable and pine marten), yet no speciation occurred in that case (Totikov et al. 2025). This suggests that the upper boundary estimates of 3.5 Mya, and particularly 5.52 Mya, represent substantial overestimates. Alternatively, if we assume that isolation during the Mid-Pleistocene Transition was the primary driver of speciation and apply the mutation rate confidence interval boundaries to the 1.3 Mya estimate, we obtain a more reasonable confidence interval of 0.82-1.97 Mya.

### Protein-coding genes, inversions, Tajima’s D and Fst

Within the previously reported inverted regions between the sable and pine marten (Tomarovsky et al. in prep), we found relatively few genes: 31 on chr11 and 44 on chr12 (Supplementary File <u>SF12</u>), with Gene Ontology (GO) analysis showing no statistically significant enrichment of GO terms for these rearrangements, though manual examination revealed two reproduction-related genes (*SPMIP7* and *ZPBP*) within the chr11 inverted region. Both genes are located ∼1.5 Mbp from the inversion breakpoint, and the chromosomal rearrangement shifted them from a peritelomeric position in the pine marten and stone marten (representing the ancestral state) to a pericentromeric position in the sable. We found a higher differentiation for the p-arm of chr11 (mean Fst 0.71) than for p-arms of chromosomes of similar size (mean Fst values 0.50-0.56, all p-values << 0.001, see <u>SRD</u>). These results suggest that the inverted region has already accumulated a notable amount of substitutions.

Comparison of pure sables and pine martens revealed 12 regions with high values (>=0.9) of the weighted Fst metric (Supplementary Table ST7). The most interesting of them, FST11 (chr9: 86,600,000–88,100,000), showed enrichment in four GO categories: keratinization (GO:0031424), intermediate filament organization (GO:0045109), embryonic skeletal system development (GO:0048706) and anterior/posterior pattern specification (GO:0009952). Examination of the gene content showed that it includes a cluster of HOX-C genes (homeobox C cluster) and seven cytokeratin genes: *KRT76*, *KRT3*, *KRT4*, *KRT79*, *KRT78*, *KRT8* (type II) and *KRT18* (type I). Tajima’s D metric calculated for all samples together (<u>SRD</u> Figure 5A and D) revealed only a single region (TJD1) with a significantly high value (D >2). It is located on the chr1, encompasses 1.3 Mbp and is nested in the high Fst region FST4 (Supplementary Table ST7). No GO enrichment was detected for TJD1, but among the 20 genes located within it (Supplementary File <u>SF12</u>), we found the *FGFR3* gene (fibroblast growth factor receptor 3), which plays a key role in bone development and maintenance. The independent analysis of the pure individuals (<u>SRD</u> Figure 5B, E, C, F) detected 35 regions (TJD2-TJD36) with high negative values of Tajima’s D (D > -2) for the pine marten, and a single region (TJD37) with high positive value for the sable, respectively. Seven pine marten regions (TJD2, 5, 7, 14, 25, 30, and 34) showed enrichment in GO terms, one of them, TJD25, is enriched by GO:0045796 (negative regulation of intestinal cholesterol absorption) and GO:0010949 (negative regulation of intestinal phytosterol absorption), because it contains the *ABCG5* and *ABCG8* genes, which encode sterolin-1 and sterolin-2 proteins (Supplementary File <u>SF12</u>).

## Discussion

### Mosaic recombinant chromosomes in hybrids

Our analyses identified 11 hybrid individuals using a 5% introgression threshold to distinguish individuals with pure species ancestry from those with hybrid or introgressed genomes. In our hybrid samples, we observed multiple cases of mosaic recombinant chromosomes (Figure 9). If crossing over were suppressed, we would expect sable and pine marten chromosomes to remain intact and separate in descendants (Figure 10B), but our findings support a different model involving active recombination (Figure 10A). Because of the nature of our dataset (short read paired end Illumina resequencing only), we lack a long range information to phase the genetic variants. However, we can take into account distribution of the ancestry along chromosomes and propose possible variants of the large-scale phasing (Figure 9). For example, the ancestry along chr 10 of sample T18 suggests only a single variant of the phasing (phased I), whereas for chr 5 and chr10 of sample T84 there are two (phased I and II) and six (phased I-VI) possible options, respectively. Possible phasing patterns for selected chromosomes indicate that both homologs in some chromosome pairs are recombinant and consist of multiple segments, indicating that multiple crossover events have occurred during hybrid formation (Figure 9). Such patterns can only arise through multiple successive crosses involving various combinations of pure species and different hybrid types, and only if crossover is not completely suppressed. Our results suggest that the sympatric zone functions as a hybridization hotspot with introgression events radiating outward in all geographic directions (Figure 10C, Supplementary Figure SF11). We observed a continuum border between the species (Harrison and Larson 2014), characterized by a gradual transition from pine marten in the west to sable in the east (Figure 6). Due to the limited number of samples (for species with such large ranges) we cannot accurately define the borders of the introgression zone. The eastern border is more clearly delimited due to the documented ongoing historical expansion of the pine marten range eastwards, which passes roughly along the eastern edge of the Ob River basin; in the west, it possibly expands past the known sympatry zone in the pre-Urals, i.e. to the west of the Komi Republic, Permskiy Kray, and Sverdlovsk Oblast.

**Figure 9.**
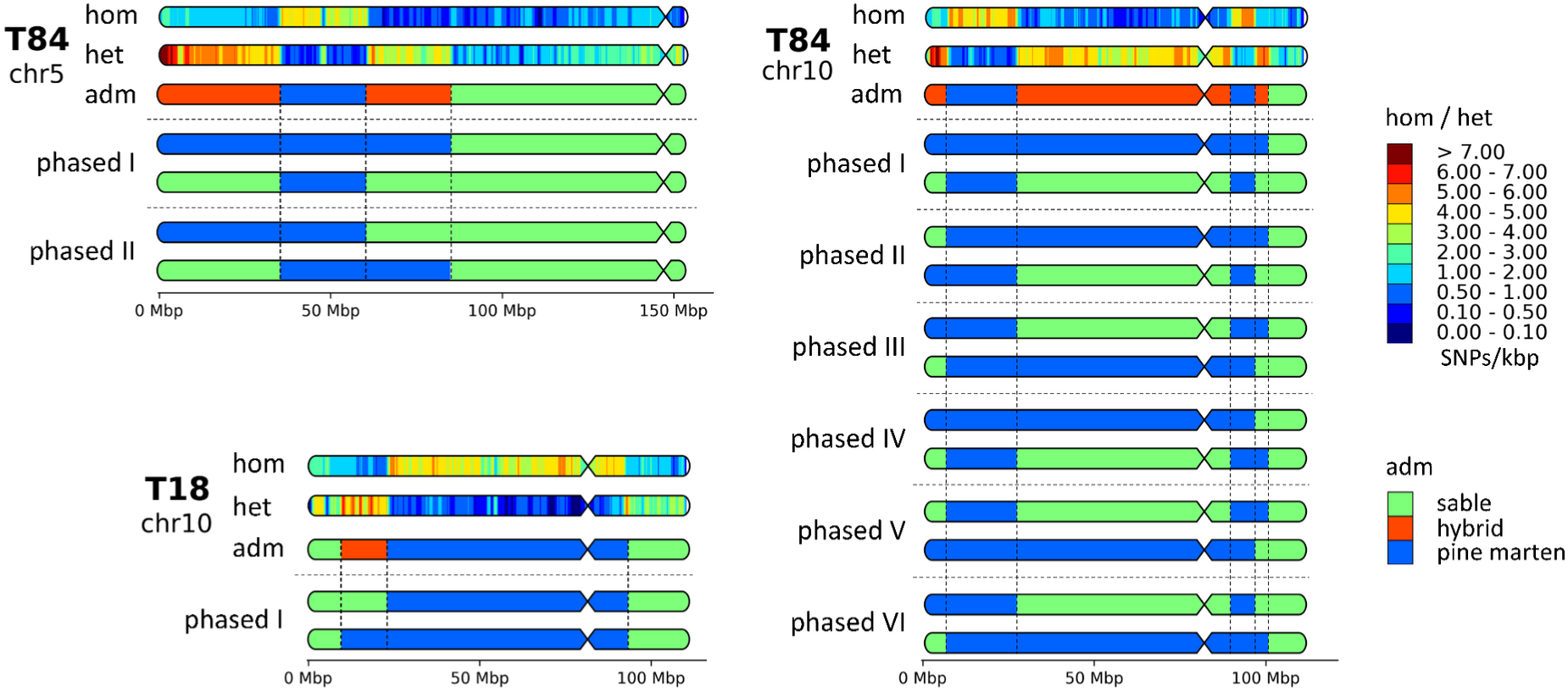
Mosaic recombinant chromosomes and possible variants of their phasing (scheme). Bold T** indicates the sample ID. Abbreviations: hom / het – density of homozygous and heterozygous SNPs, respectively. SNPs were counted in 1 Mbp windows with 100 kbp step size and scaled to SNPs/kbp (represented by the color scale on the bottom, ranging from dark blue (extremely low heterozygosity) to brown (very high heterozygosity), with 0 to 0.1 and over 7 heterozygous SNPs per 1 kbp, respectively; adm – local admixture (% of *M. zibellina*) on chromosomes of 1 Mbp sliding windows with 100 kbp step; phased ** – possible variants of chromosome phasing.

**Figure 10.**
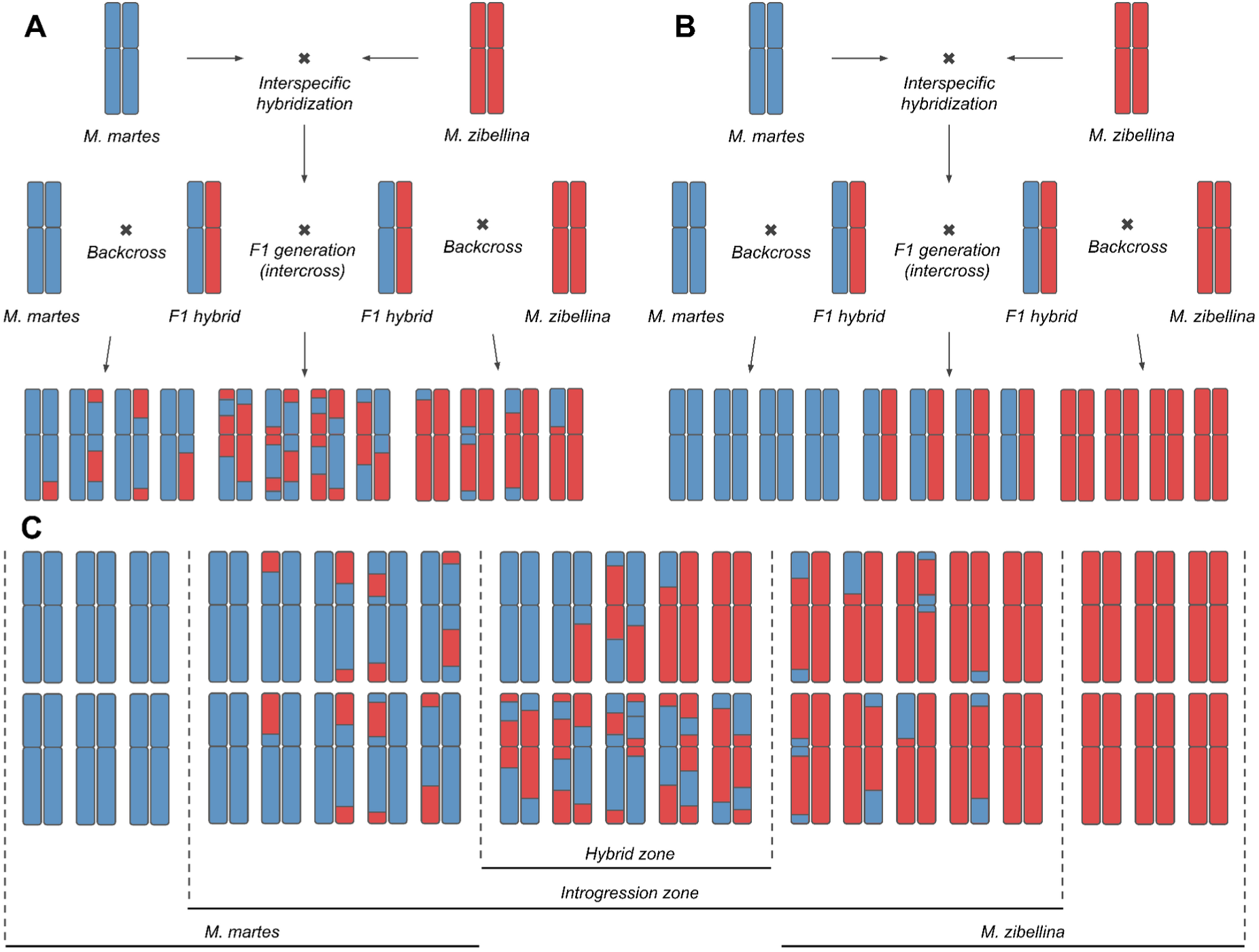
Hybridization, crossover and introgression. A – case of unsuppressed crossover between homologous chromosomes of different origin; B – hypothetical case of completely suppressed crossover; C – gradient of admixture (continuum border between species) between zone of sympatry and regions of pure species in case of unsuppressed crossover

### Genetic ancestry and discordance between different datatypes

We identified the ancestry of the sampled individuals using global and local admixture analyses. The latter analysis successfully detected low (<7%) rates of admixture (confirmed by the heterozygosity components plus F3-, D-, and F4-statistics), which the global analysis failed to detect. Our results showed that the classification of putative hybrids based on phenotypic traits was notably imprecise (Figure 11). Moreover, even the morphometric character Δ (see *Introduction*), previously proposed as a reliable metric for distinguishing sables and pine martens (Monakhov 2021a), was heavily biased toward pine marten values in putative kidases and therefore proved uninformative for hybrid verification.

**Figure 11.**
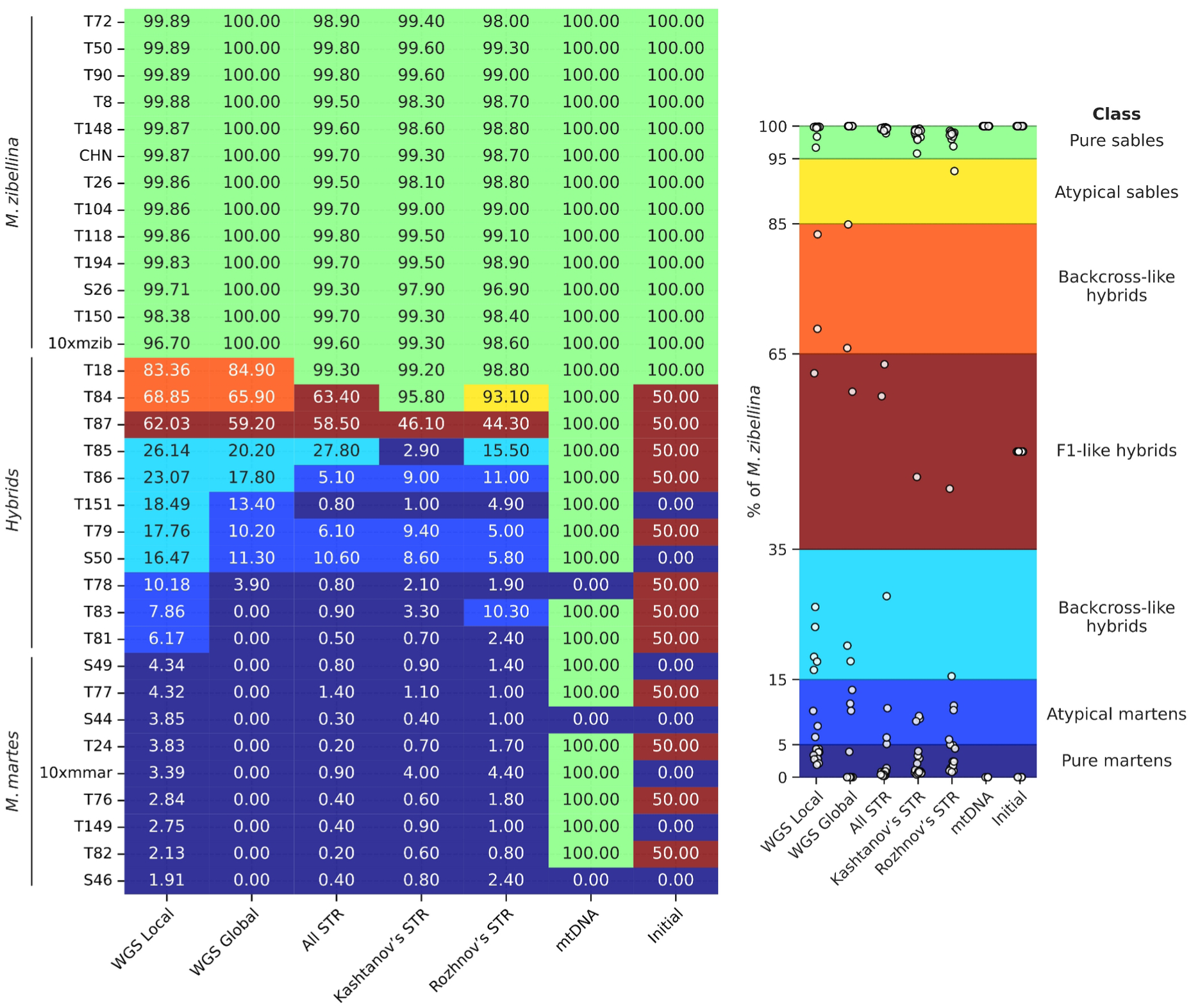
Discordance in ancestry estimates of sable × pine marten hybrids between different analytical methods and datasets. Individuals are coded as a percentage of sable ancestry, i.e., 100% indicates a pure sable, whereas 0% indicates a pure pine marten. Abbreviations of methods and data used to calculate ancestry: WGS Local – WGS data and the local approach, WGS Global – WGS data and the global approach, All STR – complete set of all mapped STR markers, Kashtanov’s STR – STR markers from (Kashtanov et al. 2022), Rozhnov’s STR – STR markers from (Rozhnov et al. 2013), mtDNA – the origin of mitochondrial DNA, and Initial – simple phenotypic traits: tail length, fur length and quality, throat patch size, respectively

The unsuppressed crossover in sable × pine marten hybrids revealed in this study suggests that accurate ancestry identification requires, if not WGS data, then at least thousands of markers uniformly distributed over all chromosomes. However, there are multiple studies of hybridization between sables and pine martens, as well as on the intraspecific diversity of sables, which were performed using various sets of STRs and mitochondrial markers (Rozhnov et al. 2010; Andrianov et al. 2012; Rozhnov et al. 2013; Kashtanov et al. 2015; Li et al. 2021; Kashtanov et al. 2022). Moreover, such studies continue to be published (Ranyuk et al. 2025). Application of known mustelid STR markers to classify samples in this study revealed significant discordance with WGS-based analysis for hybrid samples, primarily underestimating admixture levels (Figure 11). This result is expected given the limited number of STR markers, which fail to provide complete chromosomal coverage even when combined (Supplementary Figure SF5). STR markers can only be considered reliable when crossover is completely suppressed and proven coverage of all chromosomes is achieved, suggesting that previous assessments require reevaluation (Rozhnov et al. 2010; Rozhnov et al. 2013; Ranyuk et al. 2025).

Mitochondrial DNA obviously can’t be used for classification of hybrids, but in the case of our samples, it doesn’t even correlate with the nuclear genome for pure martens (Figure 11). Therefore, mtDNA of the sable and marten, and, probably, of other actively hybridizing mustelid species, should be used with precautions even in the phylogenetic studies.

### Legacy of the sable reintroduction program

We found that the sable exhibits unexpectedly high heterozygosity (1.5 – 1.8 hetSNPs/kbp) given its documented history of population declines due to fur harvesting. It is comparable to the heterozygosity (1.78 hetSNPs/kbp) of the tayra (*Eira barbara*), a neotropical gulonine that experiences some pressure from human, but never was extensively hunted . Moreover, it is approximately three times higher than in the pine marten (Figure 12). The concave сumulative RoH trajectories of all sable samples, except one, are also very different from that of the pine martens (Figure 7B and D). We suspect that interbreeding with reintroduced sable individuals, mostly from the Barguzin (*M. z. princeps*) and Vitim (*M. z. vitimensis*) subspecies (Timofeev and Nadeev 1955; Monakhov 2011), may have introduced new genetic diversity, increasing genetic variation, thereby reducing the probability of consanguinity, which in turn could explain the low content of RoH.

**Figure 12.**
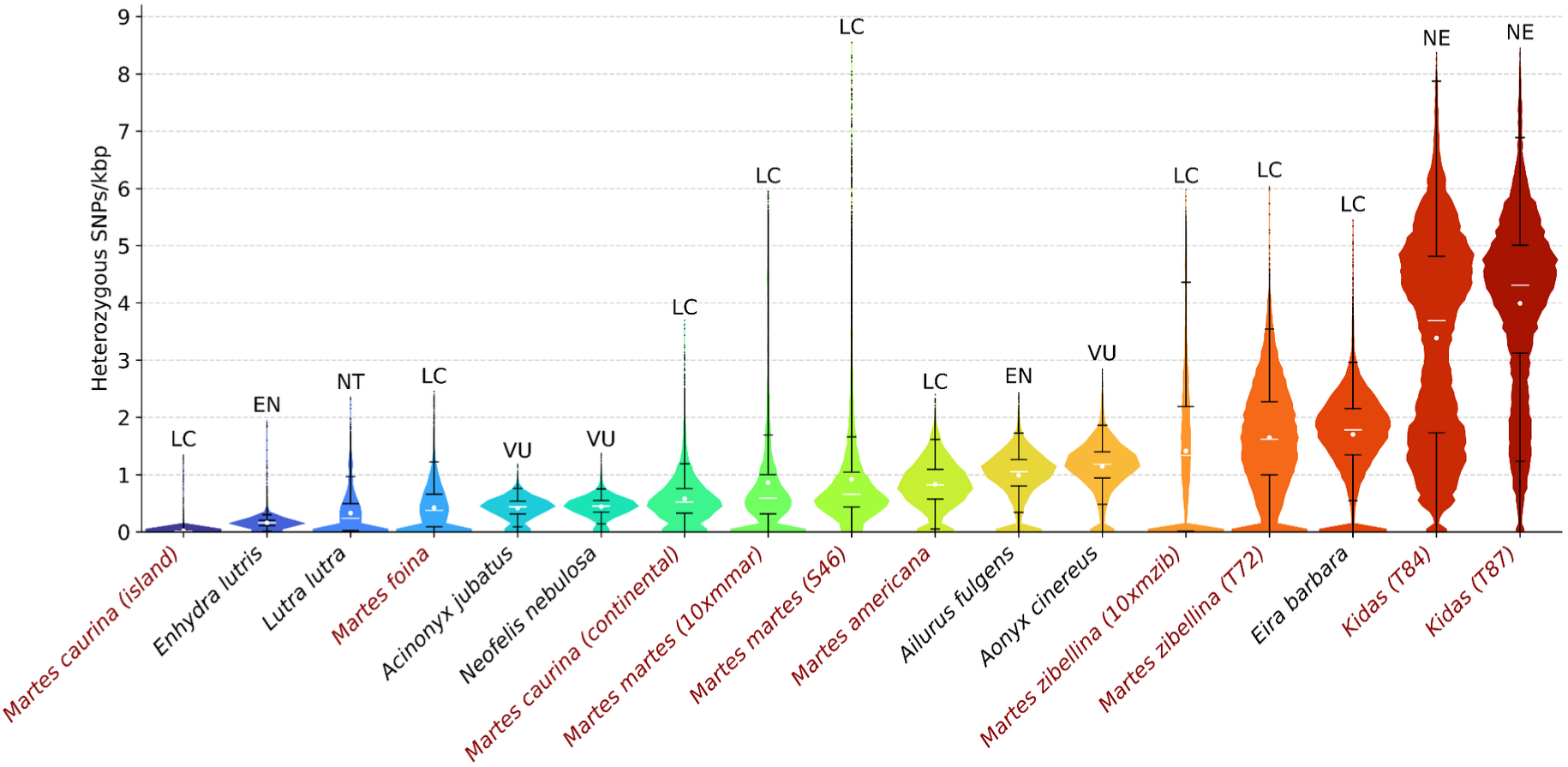
Distribution of the heterozygous SNP density in mustelids and three well-known in conservation genetics species: cheetah (*Acinonyx jubatus*), clouded leopard (*Neofelis nebulosa*), and red panda (*Ailurus fulgens*). Distributions were counted in 1Mbp sliding windows with 100 kbp step. Mean values are marked with white dots, medians are represented by white lines. Color gradient indicates level of heterozygosity, ranging from lower (blue) to higher (red) values. Marten species (including hybrids) are highlighted in red. American marten (*Martes americana*; SRR11575352) and pacific marten (*Martes caurina*; SRR11575348 (island) and SRR11575351 (continental)) samples were obtained from (Colella et al. 2021); *M. zibellina* genome assembly was used as reference. Distributions for other species were obtained from (Totikov et al. 2021; Derežanin et al. 2022; Tomarovsky et al. 2025). Abbreviations for Global conservation status according to IUCN Red List data: LC – Least Concern, NT – Near Threatened, VU – Vulnerable, EN – Endangered, NE – Not Evaluated

Among the sables we examined, sample T72 (∼1.6 hetSNPs/kbp) from Kamchatka is worth highlighting, as it has a cumulative RoH trajectory very different from that of other sables (Figure 7D), but similar to that of the pine martens (Figure 7B). The Kamchatkan population remained relatively stable throughout the bottlenecks and was used to restock some areas in Eastern Siberia (Magadan Oblast, parts of Yakutia, etc.) where the sable had nearly disappeared (Sablina 1958). This supports the hypothesis that Kamchatka may still harbor an autochthonous populations not affected by admixture with other subspecies, consistent with previous studies that have highlighted distinctive features of Kamchatka sables (Malyarchuk et al. 2010; Malyarchuk et al. 2014; Kinoshita et al. 2015; Li et al. 2021). Following this hypothesis, the similar heterozygosity of T72 and sables from the Western and Central Siberia, regions where sables experienced severe population declines due to overharvesting, suggests that these results reflect the success of the reintroduction programs. However, our hypothesis should be interpreted with caution, as we have sequenced a limited number of sables, and only one of them belongs to the Kamchatka population.

### Displacement of the pine marten mtDNA

Surprisingly, only two of the pure pine marten samples (S44 and S46) and only one hybrid (T78) have pine marten mtDNA (Figure 5). Moreover, our pure pine martens have mtDNA from four different sable clades: A3, A5, B2, and C4 (Figure 5, Supplementary Figure SF6), which implies that interspecific mtDNA introgression has occurred multiple times. Such a phenomenon might be explained by the biological and behavioral features of these species. Sable is hypothesized to be better adapted to life in deep snow, and several morphological traits, such as its shorter tail, thicker fur, thicker-furred soles, and better-developed strips of bristly fur on the feet (“snowshoes”) are indirect evidence of this (Timofeev and Nadeev 1955). In regions with harsh, snowy winters such as the sympatric zone, pine marten may be vulnerable in direct confrontations with sables, particularly smaller females. Some sources claim sables are more aggressive to conspecifics than pine marten, with cases of intraspecific killings being exceptionally rare in pine martens but common in sables. In fact, sables cannibalizing trapped conspecifics is a common occurrence and has been attributed to territorial behavior rather than hunger-motivated carnivory (Buskirk et al. 1994). Therefore, female pine martens might be casualties of interspecific competition in the sympatric zone. Also, it implies that sable mtDNA might have spread far West of the sympatric zone and that pine marten mtDNA should only be occasionally found in the East. Despite the limited sample size of our dataset, we see such a trend in our results: two martens (the pine marten S49 and hybrid S50) collected in the far West have sable mtDNA, and no specimen collected in the East has pine marten mtDNA.

### Ancient roots of low heterozygosity in the pine marten

Surprisingly, we found that the pine marten is significantly less heterozygous than the sable (Figure 7A, <u>SRD</u> Figure 2). Its median heterozygosity of ∼0.5-0.6 hetSNPs/kbp (Figure 7A, Supplementary Figure SF12), is smaller than in some endangered species, for example, the red panda (1.05 hetSNPs/kbp) and Asian small-clawed otter (1.18 hetSNPs/kbp) (Figure 12). It is only 40% higher than the heterozygosity of such iconic species of conservation biology and genomics as the cheetah (0.44 hetSNPs/kbp) and clouded leopard (0.45 hetSNPs/kbp). Our demographic reconstruction (Figure 8A) suggests that it is a consequence of the severe population decline during the Mid-Pleistocene Transition (0.7-1.25 Mya) and, therefore, is an ancient trait of the species rather than a modern feature. Given its range, census numbers and no reports of inbreeding depression, we consider it a successful species, which probably purged most of the recessive lethal alleles from its genome, similar to the Channel Island fox (*Urocyon littoralis*) (Robinson et al. 2016; Robinson et al. 2018). An expanded dataset and further study are necessary to confirm this hypothesis. Moreover, all our conclusions and interpretations apply specifically to Eastern pine marten populations, while the dynamics of Western populations remain poorly understood.

### Demography and associated migration routes

The trajectories of effective population size of sables and pine martens are markedly different (Figure 8A), although they show similar curves before 3.5 Mya (Supplementary Figure SF10). We hypothesize that the significant increase in N_e_ observed in pine martens (1.3 - 3.5 Mya, Figure 8A) was associated with the colonization of new territories by ancestral pine martens, supported by the strong unidirectional gene flow from the “mainland” area inhabited by the shared ancestors with the sable. The subsequent decrease of the pine marten’s N_e_ during the Mid-Pleistocene Transition (0.7-1.25 Mya) can be considered not only as a potential result of climatic changes, but also indicative of isolation from ancestral sables, as the N_e_ of sables drops negligibly around the same time period (Figure 8A). Isolation in different refugia during the Pleistocene glaciation(s) (Ruiz-González et al. 2013) may have served as a driver of speciation that severed the gene flow between the two diverging species.

Currently, martens *sensu stricto* (subgenus *Martes*) are distributed across the Northern Hemisphere. The sable is an Asian species whereas the pine marten is mostly European (but it penetrates into Asia). The American marten (*M. americana*) and the Pacific marten (*M. caurina*) range in Northern America, whereas the Japanese sable (*M. melampus*) inhabits the Japanese archipelago. Given the distribution and relatively stable (but oscillating) trajectory of the sable, we assume that the MRCA of pine marten and sable inhabited an area in Northern Asia close or partly overlapping with the current range of the sable. Therefore, the expansion of pine marten ancestors should have been towards Europe. However, the evolutionary history of sable-pine marten interactions is likely more complex than simple ancient divergence followed by recent hybridization. The Mid-Pleistocene Transition changed not only the period of glacial-interglacial cycles, but also significantly increased the amplitude of temperature changes (Pisias and Moore 1981; Clark et al. 2006; Bajo et al. 2020). The high vagility of martens (Zalewski et al. 2004) suggests that under such conditions, species ranges should be highly dynamic, fluctuating from small refugia to vast areas, with cold-adapted species gaining substantial advantages. The evolutionary processes after divergence from the MRCA have shaped the pine marten and the sable differently. The modern sable is considered to have greater adaptability to cold habitats, but the cost was a reduced tolerance to hot summers (Monakhov 2021b). Some researchers even suggest that during the cold (but not glacial) periods of the Pleistocene, sable temporarily expanded westwards, even reaching Fennoscandia, and had retracted back to the Siberian forests during warmer times. In contrast, the pine marten was hypothesized to replace the sable during warm interglacial periods (Davison et al. 2001; Stojak and Jędrzejewska 2022). However, both species share a broad temperature range optimal for both, suggesting that multiple hybridization events likely occurred during periods of favorable climatic conditions. Thus, the current ranges of the sable and pine marten most likely are the result of multiple recolonization processes shaped by Pleistocene climatic fluctuations, which led to the formation of ancient and modern zones of sympatry and hybridization after the two lineages had diverged.

### Divergence of the species

The fossil calibrated tree (Figure 3) estimated the split between the sable and pine marten at 1.52 Mya (CI: 1.05 – 2.06 Mya), which substantially overlaps with the demographic trajectory-based estimate of 1.3 Mya (CI: 0.82 – 1.97 Mya). This divergence has been estimated in at least four previous genetic studies (Supplementary Table ST8) (Koepfli et al. 2008; Li, Wolsan, et al. 2014; Law et al. 2018; Hassanin et al. 2021). Hassanin et al. (2021) and Li et al. (2014) used complete mitochondrial genomes (∼16 kbp), Koepfli et al. (2008) analyzed 21 nuclear gene segments and 1 mitochondrial gene (∼12 kbp), while Law et al. (2018) employed 4 mitochondrial genes and 42 nuclear gene segments (∼35 kbp). Most of these studies agree that the MRCA of sable and pine marten existed ∼1.1 Mya (CI: ∼0.6 – 1.6 Mya), with the exception of Li et al. (2014), who estimated substantially younger divergence of 0.68 Mya (CI: 0.54 – 0.84 Mya). Interestingly, for the MRCA of all three marten species (including the stone marten, *Martes foina*), the outlier dating estimate belongs to Hassanin et al. (2021) (5.1 Mya, CI unavailable), whereas the other studies estimated it no older than 2.93 Mya (Li et al., CI: 2.35 – 3.54 Mya) or even younger, at 2.8 Mya (Koepfli et al., CI: 1.9 - 3.8 Mya) or 2.56 Mya (Law et al., CI: 1.93 – 3.29 Mya). Our mean estimate of 2.83 Mya and the associated confidence intervals (CI: 2.11 – 3.66 Mya) are consistent with these values. In contrast to the earlier studies, our phylogenetic analyses employed significantly more sites: ∼100x more sites (1.38 Mbp) than the mtDNA genome-based works and ∼40x more than the study by Law et al. (2018). Moreover, we utilized only clockwise sites – fourfold-degenerate sites from the third codon positions of single-copy genes. Another potential source of discrepancy may be incomplete lineage sorting or ancient hybridization with the lineages leading to American and/or Japanese martens, which could have affected mtDNA and the small set of nuclear genes used by Law et al. (2018).

### Candidate genes related to the fertility issues, phenotypic and diet differences

We found that two fertility-related genes, *SPMIP7* and *ZPBP*, are located within previously cytogenetically confirmed 11.5 Mbp inversion between the sable and pine marten on the p-arm of chr 11 (Tomarovsky et al. in prep). *SPMIP7* (Spermatozoon Microtubule Inner Protein 7) is essential for normal spermatogenesis (Leung et al. 2023), while *ZPBP* (Zona Pellucida Binding Protein) functions in both spermatogenesis and sperm-egg binding (Lin et al. 2007). In humans, mutations in *ZPBP* are associated with spermatogenic failure-66 (SPGF66) disorder and male infertility due to globozoospermia (Oud et al. 2020). Inversions of such size are also known to completely suppress recombination, as their length is insufficient to form the characteristic inverted loop during meiosis (Anton et al. 2005). The notably higher Fst values on the p-arm of chr 11 relative to that on other chromosomes (see Results) support this. We suspect that recombination in this region was likely suppressed in the MRCA of sable and pine marten, which should have resulted in the accumulation of multiple substitutions (Wellenreuther and Bernatchez 2018). Therefore, the chromosomal environment of these genes differs between these species and their regulatory mechanisms may have evolved divergently as well. The presence of alleles of different origin in hybrids can lead to the disrupted regulation of the related processes (Runemark et al. 2025), and potentially can explain the reported fertility issues in kidasses (see *Introduction*).

The same inversion also contains *GLI3*, a gene involved in developmental processes (Supplementary File <u>SF12</u>). This gene is known to participate in the regulation of limb and skeletal development (Kalff-Suske et al. 1999; Wang et al. 2024). In humans, the associated disorders include acrocallosal syndrome, characterized by distinct craniofacial features such as hypertelorism (widely spaced eyes) and a prominent forehead (Elson et al. 2002), which are similar to the skull differences between the sable and the pine marten (Monakhov 2021a). This suggests that *GLI3* may be a promising candidate to study the genetics of morphological evolution of marten species. Other candidate genes we identified may also be associated with other phenotypic features. For example, the *FGFR3* gene, associated with craniosynostosis and multiple types of skeletal dysplasia (Schibler et al. 2009), might be linked to the increased vertebral number in the pine marten (Monakhov 2011; Monakhov 2022). Similarly, the *HoxC* cluster, known for its roles in limb development (Okamoto et al. 2011), could relate to the differences in paw size.

The two species are also known to exhibit significant dietary differences. The sable shows a flexible feeding pattern, shifting from a diet primarily composed of plant matter (nuts and berries) during years of abundant harvest to one predominantly consisting of animal prey (small vertebrates) (Monakhov 2016; Cheprasov and Mordosov 2019). In areas with high yields of nut-producing five-needle conifers, sables rely heavily on nuts produced through windfall or cached by birds and squirrels throughout the fall and early winter (Zakharov et al. 2016). In contrast, the pine marten does not feed on nuts in appreciable quantities, if at all (Helldin 2000; Posłuszny et al. 2007; Twining et al. 2019). Plant matter (mostly berries) contributes up to 50% or more of its diet in southern areas, but in most regions it is reported to be a hypercarnivore (>70% animal prey) during the year (Zalewski 2005). We found two genes, *ABCG5* and *ABCG8*, directly involved in the transport of dietary sterols (both cholesterol and plant sitosterols) (Hazard and Patel 2007) overlap with regions having high negative Tajima’s D in the pure pine martens, but not in the pure sables. Previously, it was shown that simultaneous knockout of both genes in mice decreased the concentration of the cholesterol in the bile, resulting in its accumulation in the liver, as well as increased the plasma level of sitosterols (Yu, Hammer, et al. 2002). In contrast, the overexpression of these genes increased the bile secretion of the cholesterol and halved its fractional absorption from the digestive tract (Yu, Li-Hawkins, et al. 2002). Low Tajima’s D are typically interpreted as indicators of either negative selection or population expansion (Carlson et al. 2005). Therefore, *ABCG5* and *ABCG8* are considered prime candidates for future genetic studies focusing on the dietary differences between sable and pine marten, specifically regarding the latter’s adaptation to hypercarnivory.

## Conclusion

We conducted a thorough genomic investigation into the hybridization between the sable and pine marten; however many of the patterns revealed warrant further study. The first set of new questions is directly related to the hybrids. We observed bidirectional introgression, including the substitution of marten mtDNA by sable mtDNA in many samples extending far beyond the sympatric zone. This suggests that current species boundaries may be obsolete and require reevaluation. The notable discrepancy between our results and previous mtDNA-based divergence dating necessitates an investigation into potential ancient gene flow from other species. Such introgression, for example, from a closely related *M. americana* and *M. caurina* hybrid system, may bias estimates. However, such a complex interspecies study should be studied using a pangenome approach instead of usual linear reference(s). In our study we performed all the analyses twice, using both sable and pine marten genome assemblies as references, to confirm absence of the reference-related biases. The pangenome methodology is free from such issues. Second, the lower heterozygosity in the pine marten is surprising, as from the history of both species we expected the opposite pattern. Our samples cover only the eastern part of the range, so the diversity of western populations remains unclear. Our dataset, while spanning much of both species’ ranges, included only 33 individuals; thus, for such widely distributed species, this study should be considered a pilot investigation. Moreover, it is still unclear how the population declines and one of the largest reintroduction programs in history have affected the subspecies and population structure, and whether the diversity was indeed restored or not. Our findings provide reasons for cautious optimism, but the heterozygosity of sables before the population declines of the 19th and 20th centuries remains unknown. To address these questions, a broader sampling and sequencing of hundreds of individuals, including museum and ancient samples, is required. Finally, we have taken the initial step in investigating the adaptomics of martens, identifying a set of candidate genes potentially associated with the morphological and dietary differences between the sable and the pine marten. Future investigations will necessitate fieldwork, detailed gene sequence analysis, transcriptomic and metagenomic studies, and biochemical experiments. Our work thus establishes a foundation for at least four research directions and highlights the appropriate methodologies that can be used to pursue them.

## Materials and methods

### Samples, DNA extraction and sequencing

For whole genome resequencing, we used previously collected tissue samples from Novosibirsk Cell Line Collection located at the Institute of Molecular and Cellular Biology, Siberian Branch of the Russian Academy of Sciences (IMCB SB RAS). The initial classification (pure/putative hybrid) of samples (Supplementary Table ST1) was based on the descriptions of simple morphological traits distinguishing sables, pine martens, and kidases: tail length, fur length and quality, and throat patch size (Kassal and Sidorov 2013).

DNA extraction was performed using the standard phenol-chloroform protocol (Sambrook and Russell 2006). The libraries for resequencing were constructed using the TruSeq DNA PCR-Free kit (Illumina, Inc, San Diego, CA, USA). All prepared libraries were sequenced with paired-end 150 bp reads on the Illumina NovaSeq 6000 or Illumina HiSeq X Ten platforms. All manipulations with the samples were performed according to the permission of IMCB Ethical Committee № 01/21 issued on 26 January 2021.

### Data quality control, filtration and genome size estimation

Our dataset consisted of 33 samples, including three previously sequenced martens (Tomarovsky et al. in prep). Two of published samples (10xmmar (SRR22412800) and 10xmzib (SRR22412799)) were sequenced using 10X Genomics linked reads, and therefore included barcoded at 5’ ends of the forward reads, which we trimmed using EMerAld (EMA) v0.6.2 (Shajii et al. 2017) prior processing. Initial quality control of raw and filtered data was performed using FastQC v0.11.9 (Andrews 2010). Trimming of Illumina adapters and filtering based on read quality was performed in two stages with initial 23 kmer-based trimming of large adapter fragments using Cookiecutter v1 (Starostina et al. 2015), followed by trimming of small fragments and quality filtering using Trimmomatic v0.36 (Bolger et al. 2014) with the following parameters: “ILLUMINACLIP:TruSeq2-PE.fa:2:30:10:1 SLIDINGWINDOW:8:20 MINLEN:50”. Distributions of 23-mers after read filtration were counted using Jellyfish v2.3.0 (Marçais and Kingsford 2011) with the parameters “-m 23 -s 30G” for *jellyfish count* and “-l 1 -h 100000000” for *jellyfish histo*. Then, we visualized the distributions using KrATER v2.6.1 (Kliver et al. 2017) with the parameters “-m 23 -u 1”. Genome size estimation and coverage based on paired-end reads was performed using KrATER, incorporating GenomeScope2 v2.0.1 (Ranallo-Benavidez et al. 2020) in diploid mode with a k-mer length of 23. To achieve equal coverage and maximize similarity among all samples, we downsampled them to 22x coverage using the *reformat.sh* script with the “samplerate=” parameter from BBmap v38.96 (Bushnell 2014). The downsampling fraction was calculated as the ratio of the desired coverage to the initial coverage of the sample.

### Phylogenetic analyses

The phylogenetic tree was reconstructed using the BuscoClade pipeline v1.7 (https://github.com/tomarovsky/BuscoClade), based on a multiple codon alignment of conserved single-copy orthologous gene coding sequences (BUSCOs). We included genome assemblies from a total of 21 species (Supplementary Table ST9) (Peng et al. 2014; Foote et al. 2015; Hu et al. 2017; Jones et al. 2017; Dudchenko et al. 2017; Dudchenko et al. 2018; Taylor et al. 2018; Miranda et al. 2021; Peng et al. 2021; Newman et al. 2022; Karimi et al. 2022; Lok et al. 2022; Mohr et al. 2022; Derežanin et al. 2022; Kliver et al. 2023; Tomarovsky et al. 2025; Tomarovsky et al. in prep). BUSCO sequences were identified with BUSCO v5.4.2 (Manni et al. 2021) using the database Mammalia_odb v10, 2021-02-19. Only single-copy sequences common for all species were used in the alignment. The codon alignments were performed separately for each sequence using PRANK v170427 (Löytynoja 2014), followed by a filtration of hypervariable and poorly aligned regions using GBlocks v0.91b (Castresana 2000). The resulting filtered alignments were concatenated, yielding a final alignment of 9,707,517 bp. Phylogenetic inference was performed using the maximum likelihood (ML) method implemented in IQ-TREE v2.2.0 (Minh et al. 2020) with automatic selection of the best-fitting substitution model using ModelFinder (Kalyaanamoorthy et al. 2017). Node support was assessed with 1,000 bootstrap replicates. Bayesian phylogeny inference was conducted using MrBayes v3.2.6 with the GTR substitution model and 2,000,000 generations of Markov chain Monte Carlo (MCMC). The first 15% of generations were discarded as burn-in before generation of the posterior probability majority consensus tree. An alternative phylogenetic reconstruction was performed using ASTRAL-III v5.7.1 (Zhang et al. 2018), based on a set of separately reconstructed gene trees using IQ-TREE v2.2.0. To reduce noise, we applied a filtering step by collapsing nodes with bootstrap support below 70% prior to species tree inference. A summary of support metrics for each internal branch in the ASTRAL species tree is presented in Supplementary Table ST10. Visualization of phylogenetic trees was performed using the ETE Toolkit v3.1.2 (Huerta-Cepas et al. 2016). For the final analysis and time calibration we used a reduced tree of nine species (see <u>SRD</u>).

Divergence times were dated using the MCMCTree tool from PAML v4.10.7 (Yang 2007) and the fossil-based calibrations listed in Supplementary File <u>SF1</u>. (Stach 1959; Wolsan 1990; Wang et al. 2005; Salesa et al. 2013; Samuels and Cavin 2013; Li, Wolsan, et al. 2014; Law et al. 2018; Marciszak et al. 2024). First, we extracted 4-fold degenerate sites (1,384,523 bp) from the concatenated alignment used to reconstruct the tree. Next, we performed analyses twice (according to the recommendations from authors of the MCMCTree tool) for each of the three molecular clock types (global, correlated and independent) with 2,200,000 MCMC generations. The first 200,000 generations were discarded as burn-in. We found no discordance between the two runs for each of the clock model analyses. We verified satisfactory convergence of parameters using Tracer v1.7 (Rambaut et al. 2018) for each of the runs. Visualization of the dated phylogenetic tree was performed using FigTree v1.4.4 (Rambaut 2018). All phylogenetic trees generated in this study are included in Supplementary File <u>SF13</u>.

### Reference genome assemblies, alignments, coverage and PAR identification

Filtered reads from the 33 samples were aligned to the reference assemblies of both sable (10xmzib, GCA_040938815.1) and pine marten (10xmmar, GCA_040938825.1) (Tomarovsky et al. in prep) using BWA v0.7.11 (Li and Durbin 2009) with default parameters. All the downstream analyses were performed twice for each of the references and compared. Read processing (pairing, sorting, quality filtering, marking duplicates, and indexing the alignments) was performed using Samtools v1.15.1 (Li et al. 2009). Per-base genome coverage was calculated using Mosdepth v0.3.0 (Pedersen and Quinlan 2018) with the parameter “--mapq 10”. Individual genome coverage masks for each sample were obtained based on Mosdepth assessments using the *generate_mask_from_coverage_bed.py* script from the MAVR v0.97 package (https://github.com/mahajrod/MAVR) with the parameters “-x 2.5 -n 0.33”, corresponding to coverage thresholds of >250% and <33% of the median genome coverage.

To set the correct ploidy of the X chromosome during variant calling we identified the coordinates of the pseudoautosomal region (PAR) in male samples based on individual coverage masks using the *pseudoautosomal_region.py* script from the Biocrutch software package (https://github.com/tomarovsky/Biocrutch). The algorithm is described in Supplementary Methods <u>SM3</u>.

### Variant calling and runs of homozygosity

Variant calling for all samples was performed using Bcftools v1.15.1 (Li 2011) with parameters: “-d 250 -q 30 -Q 30 --adjust-MQ 50 -a AD,INFO/AD,ADF,INFO/ADF,ADR,INFO/ADR,DP,SP,SCR,INFO/SCR -O u” for *bcftools mpileup* and “--group-samples - -m -O u -v -f GQ,GP” for *bcftools call*. Low-quality genetic variants were removed using the *bcftools filter* with the following thresholds (’*QUAL < 20.0 || (FORMAT/SP > 60.0 | FORMAT/DP < 5.0 | FORMAT/GQ < 20.0)*’). The PAR coordinates were specified for variant calling via the “--ploidy-file” parameter. The genetic variants in regions with too high and too low coverage were removed using individual per-sample masks (described in the previous section) and BEDTools v2.31.0 (Quinlan and Hall 2010) with the default parameters. The filtered and masked genetic variants for each sample were divided into heterozygous and homozygous single nucleotide polymorphisms (SNPs) and insertions/deletions (indels) using the bcftools filter with the parameters “- i ‘TYPE=“snp”’”, “-i ‘TYPE=“indel”’”, “-i ‘FMT/GT=“het”’” and “-i ‘FMT/GT=“hom”’”. All subsequent analyses were performed using only filtered and masked SNPs, which were counted in overlapping 1 Mbp sliding windows with a 100 kbp step size. As some homologous scaffolds in the genome assemblies had different orientations, we inverted corresponding pine marten chromosomes. In such cases, recalculation of SNP coordinates was performed using Picard v.2.27.4 LiftoverVcf (Anon 2019). The visualization of SNP density heatmaps was performed using the *draw_variant_window_densities.py* from the MACE v1.1.32 package (Kliver 2024).

Runs of homozygosity (RoH) were identified based on the previously calculated heterozygous SNP densities in overlapping 100 kbp sliding windows with a 10 kbp step size. The algorithm was based on filtering of the windows by level of heterozygosity followed by merging into a final RoH file. A window was considered to be an RoH if its heterozygosity didn’t exceed 0.05. Windows with intermediate heterozygosity (between 0.05 and 0.1) were also treated as RoH windows, but only if there were no more than five such windows in a row. Windows with higher heterozygosity (greater than 0.1) were excluded. Next, the filtered windows were merged together if the distance between them (between the end of the previous window and the start of the next one) was less than half of the window size. The X chromosome was excluded from the RoH analysis. Visualization of RoH distribution along the chromosomal scaffolds was performed using the *draw_features.py* script from the MACE v1.1.32 package (Kliver 2024).

### Classification of individuals and introgression analysis

As input for population structure and admixture analysis we used the filtered SNPs as described in the previous section. However, instead of individual per-sample masks, we generated a unified mask based on pairwise intersections of individual per-sample coverage masks (see *Alignments, coverage and PAR identification*) using BEDOPS v2.4.40 (Neph et al. 2012) followed by merging of all intersections. The resulting mask included genomic regions with excessively high (>250%) or excessively low (<33%) coverage in at least two samples. Filtered and masked SNP data were further filtered using PLINK v1.9 (Purcell et al. 2007). This involved filtering by a minimum minor allele frequency of 0.03 (“--maf 0.03”) and retaining only variants with a 100% genotyping rate across all samples. (“--geno 0”). Next, we pruned SNPs with a high level of pairwise linkage disequilibrium (LD) in sliding windows of 50 SNPs with a step size of 10 SNPs using an r ^2^ threshold of 0.7 (“--indep-pairwise 50 10 0.7”). To analyze introgression and hybridization, we performed multiple types of analyses and tests. Principal component analysis was performed using PLINK (“-- pca” option) (Purcell et al. 2007). Genome-wide admixture (global admixture) was performed using ADMIXTURE v1.3.0 (Alexander et al. 2009) for K values (number of populations) ranging from 2 to 6 with three replicates each. We chose the K with the minimal CV mean error (calculated by ADMIXTURE) and maximal average pairwise similarity. For K=2, we performed a local admixture analysis to identify introgressed regions in the putative hybrid samples. The reference genome was split into 1 Mbp sliding windows with a 100 kbp step size, and ADMIXTURE analysis was performed independently for each window. The visualization of introgressed regions along the chromosomal scaffolds was performed using the *draw_features.py* script from the MACE v1.1.32 package (Kliver 2024).

To statistically assess hybridization and gene flow, we repeated the variant calling, this time including *Martes foina* (SRR22412409), which was used as an outgroup in subsequent analyses. SNP filtering, masking, and pruning were performed following the same procedure described in the *Variant calling and runs of homozygosity* section. Hybridization detection was performed using HyDe v.0.4.3 (Blischak et al. 2018) with default parameters. To detect introgression and gene flow, we calculated F3-, D- and F4-statistics using AdmixTools v.7.0.2 (Patterson et al. 2012) with default parameters. To identify Ancestry-Informative Markers (AIMs), calculate interclass heterozygosity, and generate triangle plots, we used the triangulaR package v1.0 (Wiens et al. 2025).

### In silico PCR

Primers from 79 previously identified STR (Short Tandem Repeats) loci (Supplementary File <u>SF14</u>) were compiled from the literature for eight distinct mustelid species – European pine marten (*M. martes*), stone marten (*M. foina*), American marten (*M. americana*), wolverine (*Gulo gulo*), American badger (*Taxidea taxus*), European badger (*Meles meles*), American mink (*Neogale vison*), and ermine (*Mustela erminea*) (Davis and Strobeck 1998; Fleming et al. 1999; Domingo-Roura 2002; Vincent et al. 2003; Basto et al. 2010; Natali et al. 2010). Next, an *in silico* PCR with selected primers was performed for the reference assemblies of the pine marten and sable using Simulate_PCR 1.2 (Gardner and Slezak 2014). No more than four mismatches between the target sequence and each primer were allowed. The amplicon length was limited to 50–1000 bp. Further filtration of the amplicons was performed as previously described (Totikov et al. 2021). Finally, remaining markers were manually investigated in both genome assemblies for repeat motif, repeat length, and number of repeats. The loci with STRs longer than 100 bp, with 2 or more STRs, or without STRs were excluded from all the downstream analyses as well as STRs that were X-linked (Supplementary Methods <u>SM1</u>).

### STR-typing and STR-based admixture analysis

The STR genotyping methodology is described in detail in Supplementary Methods <u>SM4</u>. Initially, for each sample and STR locus, we extracted reads aligned to it, including 1000 bp flanking regions, using Samtools v1.19.2. We then verified pairing with Bazam v1.0.1, remapped them to the reference genome using BWA v0.7.17, and performed indel-aware realignment using IndelRealigner from GATK v3.7. (McKenna et al. 2010; Sadedin and Oshlack 2019). Next, we performed STR genotyping using hipSTR v0.6.2 (Willems et al. 2017). In the next stage, in addition to the full set of STR loci, we created two subsets with intersections between the full STR set and the STR sets used in Rozhnov et al. (Rozhnov et al. 2013) and Kashtanov et al. (Kashtanov et al. 2022) (Supplementary File <u>SF4</u>). Downstream analyses were performed for all three STR sets. Finally, we performed the STR-based admixture analysis using STRUCTURE v2.3.4 and Clumpp v1.1.2 (Pritchard et al. 2000; Jakobsson and Rosenberg 2007). All stages were integrated in the SnakeSTR pipeline (https://github.com/mahajrod/snakeSTR).

### Demographic reconstruction

To infer the historical dynamics of the effective population size (N_e_) for each species, we used the Pairwise Sequentially Markovian Coalescent software package (PSMC) v0.6.5 with the parameters “-N25 -t15 -r5 -p “4+25*2+4+6”” (Li and Durbin 2011). The consensus diploid sequences (input for PSMC) were created using Samtools v0.1.19 (Li et al. 2009) with the alignment quality parameter “-C 50” for *samtools mpileup* and the variant calling parameter “-c” for *bcftools view*. Diploid consensus sequences were generated using *vcfutils.pl vcf2fq* from Samtools, with minimum (“-d”) and maximum (“-D”) coverage thresholds for each sample. These thresholds were defined as one-third and 2.5 times the median genome coverage of a sample, respectively. Positions with coverage values outside this range were excluded. Fasta-like input files were generated using *fq2psmcfa* with a minimum base quality threshold set to “-q20”. PSMC bootstrap replicates for each sample were created by segmenting the sequences using *splitfa* and performing 100 bootstrap replicates (“-b 100”). Generation time (g) was set to 5 years (Colella et al. 2021), and mutation rate (*μ*) to 4.64 × 10^-9^ (Bergeron et al. 2023). Values of 2.94 × 10^-9^ and 7.37 × 10^-9^ were used as the lower and upper limits of the confidence interval (CI) for *μ*, respectively (Bergeron et al. 2023).

### Mitochondrial genome assemblies and analysis

Complete mitochondrial DNA (mtDNA) genomes were assembled using MitoZ v2.3 (Meng et al. 2019) with the parameters “all --genetic_code 2 --clade Chordata --insert_size 350 --requiring_taxa ’Mammalia’”. We also extracted and assembled four mtDNA genomes of *M. zibellina* from reads previously published by Liu et al. (PRJNA495455) (Liu et al. 2020) and Manakhov et al. (SRR13213810, SRR13213811, SRR13213812) (Manakhov et al. 2021). Prior to assembly, all samples were downsampled to 20,000,000 reads using the *reformat.sh* script with the “samplereadstarget=” parameter from BBmap v38.96 (Bushnell 2014). All publicly available mtDNA genomes for *M. zibellina* and *M. martes* from the NCBI database were included in the analysis using the query “(‘Martes zibellina’[Orgn] OR ‘Martes martes’[Orgn]) AND (mitochondrial OR mitochondrion) AND 10000:20000[SLEN]” (Yu et al. 2011; Xu et al. 2012; Li, Wu, et al. 2014; Malyarchuk et al. 2014; Hua et al. 2017; Li et al. 2021). To ensure that all the sequences begin at the tRNA-Phe start codon (nucleotide position 1) and have the same orientation, a manual preprocessing was performed: sequences were reverse-complemented and/or adjusted as necessary. A total of 140 mtDNA genomes were used in the analysis (Supplementary File <u>SF6</u>).

The mtDNA genome sequences were aligned using MAFFT v7.490 (Katoh et al. 2002) with default parameters followed by the filtration of the hypervariable regions of the control region and other poorly aligned regions using TrimAl v1.2rev59 (Capella-Gutiérrez et al. 2009) with the parameters “-automated1 -nogaps”. The multiple sequence alignment length prior to and after filtering was 17,326 bp and 15,701 bp, respectively. Next, we reconstructed a maximum likelihood tree using IQ-TREE v2.2.0 (automatic model selection, 1000 ultra-fast bootstrap replicates). *M. foina* (NC_020643.1) was used as an outgroup to root the tree. The filtered mtDNA sequences were clustered using CD-HIT v4.8.1 (Fu et al. 2012) with options “-c 1.0 -s 1.0 -S -aL 1.0 -aS 1.0” to count identical sequences and assign haplotype ids. Next, a haplotype network was constructed and visualized using the median joining approach implemented in PopArt v1.7 (Leigh and Bryant 2015). The input file for PopArt is provided in Supplementary File <u>SF15</u>.

### Morphological analysis

Skulls and skins of ten martens from the sympatric zone (specimens T76-T79, T81-T85, T87) were investigated for differences in morphology. The skulls were compared with specimens of *M. martes* (n=32) and *M. zibellina* (n=44) from allopatric populations that represent putatively pure ancestries for each species, kept in the collections of the Zoological Institute, Russian Academy of Sciences (Saint Petersburg, Russia). Twenty four measurements were taken on each skull using a sliding caliper with an accuracy of 0.1 mm. See Abramov & Tumanov (2003) and Monakhov (2020) for the scheme of measurements (Abramov and Tumanov 2003; Monakhov 2021a). Multivariate analyses (cluster analysis and discriminant analysis) were carried out to evaluate differences among the samples. Statistical analyses were performed in Statistica 6.0 (StatSoft 2001).

### Fst and Tajima’s D

Fst and Tajima’s D statistics were calculated using Vcftools v0.1.16 and VCF-kit v0.2.9 (Danecek et al. 2011; Cook and Andersen 2017), respectively, in 1 Mbp sliding windows with a 100 kbp step size, using the sable genome assembly as a reference. The comparison was performed between pure sable and pure pine marten samples (based on their classification – see section *PCA and admixture of WGS data*). Genome windows with more than 50,000 Ns or containing more than 607,592 bp (see Supplementary Methods <u>SM5</u> for detailed description of the threshold selection) repetitive sequences were excluded from the downstream analyses.

## Supporting information

SRD

## Supplementary

Supplementary Results And Discussion (SRD): link

Supplementary Figures (SF): link

Supplementary Tables (ST): link

Supplementary Methods (SM): link

## Acknowledgements

We thank Sergei Pisarev and Pavel Reznichenko from the Barnaul Zoo “Lesnayia skazka” (eng. “Forest tale”, Barnaul, Russia) for providing samples used to generate the *Martes martes* cell line. We acknowledge Alexander M. Migura, Valerii A. Zelepukhin, Victor D. Hazagaev, Valeriy V. Seredkin, Valentin S. Slobodenuk, Sergei D. Krekhnov, Irina A. Shuvalova, Svetlana V. Zubakina, Aleksey A. Lakhtin, Sergei M. Tsiplenkov and Valeriy T. Zhdanuk for their excellent assistance in the collection of samples from *M. martes* and *M. zibellina* specimens.

The unpublished assemblies of the Ailurus fulgens, Canis lupus familiaris, Enhydra lutris, Erignathus barbatus, Lontra canadensis, Mustela putorius furo, Odobenus rosmarus and Ursus arctos were used with permission from the DNA Zoo Consortium (https://www.dnazoo.org/).

The research reported in this study was partially completed using equipment (materials) belonging to the large-scale research facilities of the “Cryobank of cell cultures” at the Institute of Molecular and Cellular Biology SB RAS (Novosibirsk, Russia). We thank the Ministry of Science and Higher Education of the Russian Federation for granting access to the equipment. This research was supported in part through computational resources of the HPC facilities provided by the collaborative center «Bioinformatics» ICG SB RAS (Novosibirsk, Russia), as well as the computing cluster of ITMO University (Saint Petersburg, Russia).

## Data availability

Resequencing data is available from BioProject: PRJNA1102534. Mitochondrial genomes generated in this study are available at the following accession numbers: PP934007-PP934038. Mitochondrial genomes assembled from publicly available resequencing data are available in the GenBank Third Party Annotation database under the following accession numbers: PP960514-PP960517.

## Abbreviations

CI: Confidence Interval
SRD: Supplementary Results and Discussion
RoH: Run of Homozygosity
WGA: Whole Genome Alignment
PAR: PseudoAutosomal Region
PCA: Principal Component Analysis
STR: Short Tandem Repeat
MRCA: Most Recent Common Ancestor
MPT: Mid-Pleistocene Transition

