## Supplementary material for "Genomics of sable (*Martes zibellina)* × pine marten (*Martes martes*) hybridization": SRD

* corresponding author

[x] equal contribution

### List of abbreviations

MT – Main Text

CI – Confidence Interval

SRD – Supplementary Results and Discussion

HCA – Heterozygosity Component Analysis

*Samples and ranges*

Our dataset included 33 marten samples (Supplementary Table ST1), 30 of which were sequenced as a part of this study. After classification 9 have been assigned as pure pine martens, 13 as pure sables and 11 as hybrid/introgressed individuals, i.e. the final groups of pure species were of similar size. For the sable, we sequenced at least one individual per sampled population, and covered most of its geographic range from Khanty-Mansi Autonomous Okrug – Yugra in the west to the Kamchatka peninsula in the east. For the pine marten, fewer samples were available, which limited our study to the Eastern part of its range. The hybrid samples mostly (8 of 11, except samples T18, T151, S50) were gathered from a single location (Tyumen Oblast, Malyi Narys) in the sympatric zone. T18 originated from another point within the zone of sympatry (Khanty-Mansi Autonomous Okrug–Yugra, Peregrebnoe). T151 was collected in the very Eastern part of the pine marten range, but outside of the overlap with sable area according to the IUCN (2024-02) (Herrero et al. 2015; Monakhov 2015). The ranges for both species were assessed by IUCN in 2015 (Herrero et al. 2015; Monakhov 2015) and, probably, are already outdated as it has been reported that sable has increased in abundance and is migrating at least in the northern (Kashtanov et al. 2022) direction out of its range. The status of the western border of the range is unclear.

The origin of the last hybrid sample (S50) is mysterious. It is the most Western among all the samples, and was gathered deep inside the area of the pine marten (Kaluga Oblast, Kaluga). Given the high fraction of the pine marten (only 16.47% of the sable), we consider this individual as a descendant (at least second or higher generation or backcross) of an escapee from a fur farm. Such fugitives were previously reported to even reach Finland, and are considered there as invasive aliens (Partanen et al. 2020).

*Coverage and pseudoautosomal region (PAR)*

The initial coverage of our samples varied greatly (21x-76x), and we downsampled them to reach a 22x (±10%) coverage (Supplementary Tables ST2, ST11) to avoid coverage-related biases. After generation of per-sample masking tracks (Supplementary Table ST12) we detected some differences depending on what assembly was used as a reference for alignment (Supplementary Figure SF13). This, along with usage of two distinct species as well as potential hybrids from the zone of sympatry, made the choice of the reference (sable or pine marten assembly) for downstream analysis quite challenging. To mitigate these issues, we performed all analyses twice using each of the assemblies, and compared the results.

We checked the size and coordinates of the pseudoautosomal region (PAR) in male samples using a coverage-based approach. We found no discrepancy among them either in length (6.45 Mbp in all samples) nor location (SRD Figure 1, Supplementary Table ST13).

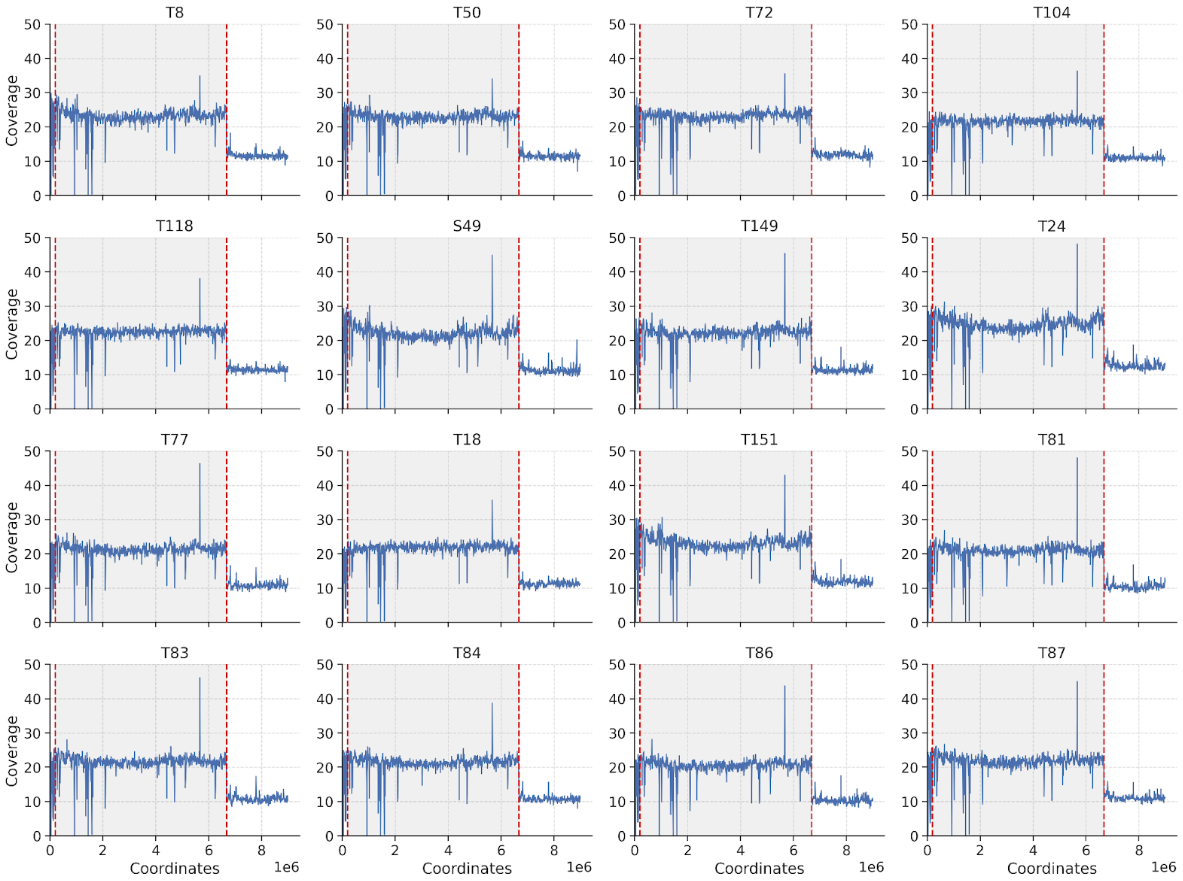


**SRD Figure 1**. Median coverage in non-overlapping 10 kbp windows for males across the pseudoautosomal region in *M. zibellina* genome assembly.

*Concept of the heterozygosity component analysis (HCA)*

Mean and median heterozygosity are significantly affected by RoH and introgression and, therefore, provide only a “summary” estimate. A common way to step over this issue is to use a window-based approach, which provides a distribution of estimates in each of the windows instead of a single value (MT Figure 6 and 7A). It allows to study distribution of heterozygosity along the chromosomes (MT Figure 6B, Supplementary File SF9), identify regions outlying by derivative metrics like Fst and Tajima’s D, etc. We decided to step further and introduce a concept of a new method - heterozygosity component analysis or HCA. An inspiration for it were heterozygosity distributions of our hybrid samples and Genomescope2 tool (Ranallo-Benavidez et al. 2020), which uses a similar approach, but for a different purpose (genome size estimation). Briefly (see Supplementary Methods SM2 for details), the basis of our method is a fitting of a linear combination of negative binomial distributions to the empirical distribution of the heterozygosity. We successfully decomposed distributions into the combination of pure pine marten (P), pure sable (S) and hybrid (H) components for all our samples (SRD Figure 2, MT Figure 6A). Mean of the P-component was very similar among all samples containing it (0.536 - 0.763, σ = 0.047) and for pure pine martens (0.559 - 0.576, σ = 0.008) was close to the global median values (0.56 - 0.67, orange bars on SRD Figure 2).

However, the notable difference with the global mean values (0.84 - 1.07, light orange bars on SRD Figure 2) is easy to explain by a small introgression from the sable to our pure pine marten samples. Because of high values of H-component (4.17 - 4.51, $\sigma = 0.11$) and low values of P-component (green dashes on SRD Figure 2), global mean values were notably biased upwards by a small number of outlying hybrid windows. Presence of the small H-component (red vertical dashes on SRD Figure 2) in pure pine martens was confirmed by the local (but not the global) admixture analysis (MT Figure 4D). Therefore, we have confirmed the robustness of our method and one more time highlighted that a single global mean value is a bad metric even in a case of low introgression.

After some additions (automatic detection of the starting parameters) and optimizations (better fitting procedure), our method can be used as a test for hybrid origin of the sample if no samples of pure species are available, or their number is not enough for a reliable Admixture/HyDe/D-statistic analysis. Principal limitation for such an application is a presence of a notable number of windows with significantly different heterozygosity, so it, for example, will not work for F1-hybrids, but should work for F1-like, as our sample T87 (MT Figure 6A). As a weak point of our method we have to mention that it is yet unclear at what degree imprecision of distribution fitting affects the values and if its contribution is comparable (or not) to biological variation among the samples.

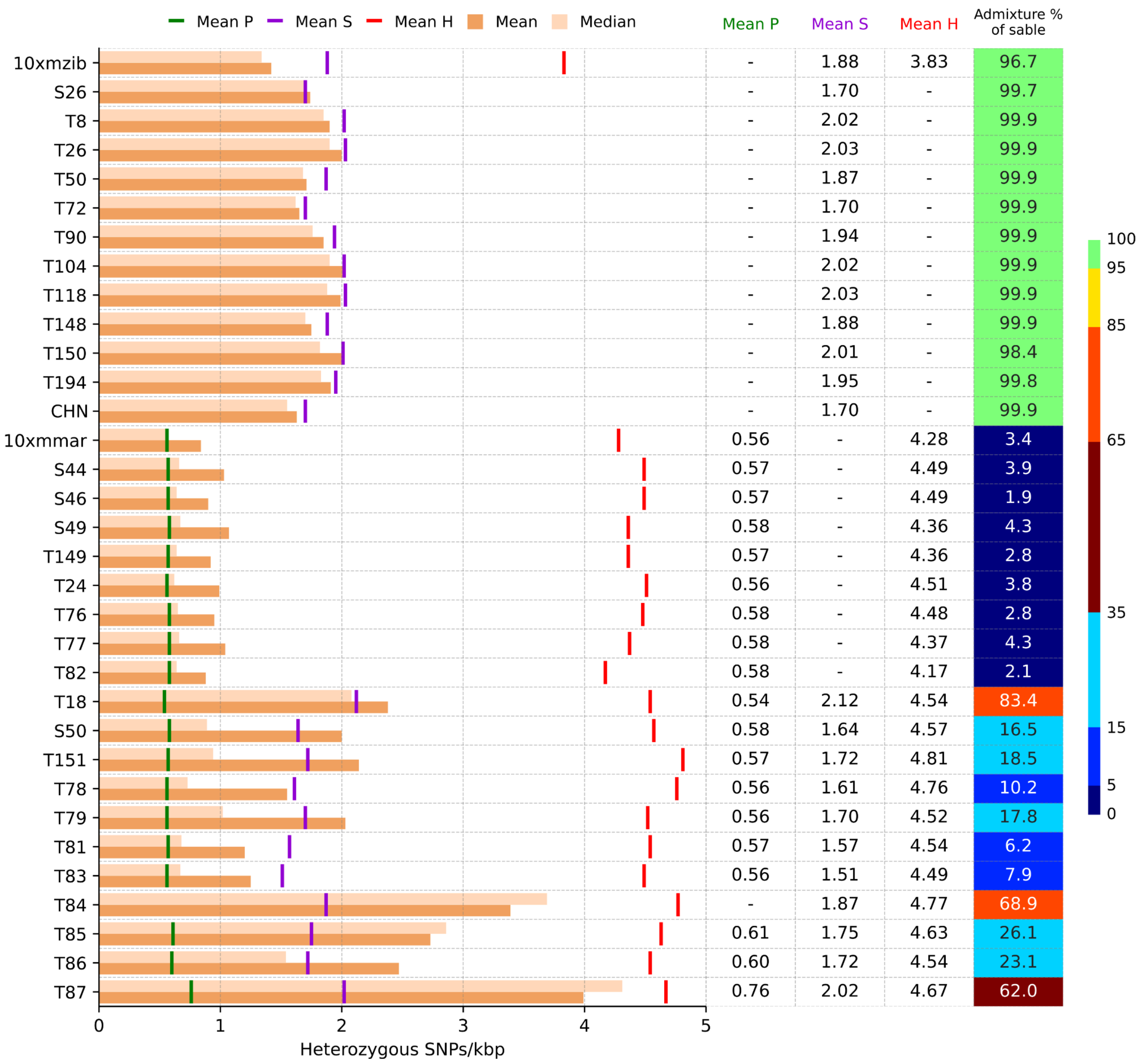


| Sample | Mean P | Mean S | Mean H | Admixture % of sable |
|---|---|---|---|---|
| 10xmzib | - | 1.88 | 3.83 | 96.7 |
| S26 | - | 1.70 | - | 99.7 |
| T8 | - | 2.02 | - | 99.9 |
| T26 | - | 2.03 | - | 99.9 |
| T50 | - | 1.87 | - | 99.9 |
| T72 | - | 1.70 | - | 99.9 |
| T90 | - | 1.94 | - | 99.9 |
| T104 | - | 2.02 | - | 99.9 |
| T118 | - | 2.03 | - | 99.9 |
| T148 | - | 1.88 | - | 99.9 |
| T150 | - | 2.01 | - | 98.4 |
| T194 | - | 1.95 | - | 99.8 |
| CHN | - | 1.70 | - | 99.9 |
| 10xmmar | 0.56 | - | 4.28 | 3.4 |
| S44 | 0.57 | - | 4.49 | 3.9 |
| S46 | 0.57 | - | 4.49 | 1.9 |
| S49 | 0.58 | - | 4.36 | 4.3 |
| T149 | 0.57 | - | 4.36 | 2.8 |
| T24 | 0.56 | - | 4.51 | 3.8 |
| T76 | 0.58 | - | 4.48 | 2.8 |
| T77 | 0.58 | - | 4.37 | 4.3 |
| T82 | 0.58 | - | 4.17 | 2.1 |
| T18 | 0.54 | 2.12 | 4.54 | 83.4 |
| S50 | 0.58 | 1.64 | 4.57 | 16.5 |
| T151 | 0.57 | 1.72 | 4.81 | 18.5 |
| T78 | 0.56 | 1.61 | 4.76 | 10.2 |
| T79 | 0.56 | 1.70 | 4.52 | 17.8 |
| T81 | 0.57 | 1.57 | 4.54 | 6.2 |
| T83 | 0.56 | 1.51 | 4.49 | 7.9 |
| T84 | - | 1.87 | 4.77 | 68.9 |
| T85 | 0.61 | 1.75 | 4.63 | 26.1 |
| T86 | 0.60 | 1.72 | 4.54 | 23.1 |
| T87 | 0.76 | 2.02 | 4.67 | 62.0 |

**SRD Figure 2**. Heterozygosity Component Analysis (HCA)

The histogram displays mean (orange) and median (light orange) global heterozygosity across samples. Vertical dashes indicate the mean component values: green (pine marten, P), purple (sable, S), and red (hybrid, H). Absolute values of the P, S, and H components, along with admixture (%) from sable, are shown on the right

*Classification of individuals by morphology*

It is believed that the kidases (hybrids *M. martes* × *M. zibellina*) can be identified by the fur coloration and the relative length of the tail (Pavlinin 1963). In most cases, the coloration of kidas is similar to the color of the pine marten: the head is darker than the back (in sable, it is lighter), the belly is evenly colored, the throat spot is usually absent or represented by separate small spots. The pine marten has a long tail, more than half the length of the body. Usually it protrudes more than 1/4 of its length beyond the ends of the hind limbs extended backwards. The tail of the sable is relatively shorter; it protrudes slightly beyond the ends of hind legs. The tail of the kidas is of medium length.

Based on the exterior features, all the studied specimens can be attributed to Martes martes, except for specimen T87, which can be identified as a hybrid (according to the color of the throat).

Many characters were used to distinguish the skulls of *M. zibellina* and *M. martes* – position and size of auditory bullae, various dental characters, mandibular characters, location of the carotid fossae, etc., however all of them are not certain enough (Pavlinin 1963; Heptner et al. 1967). Recently, Monakhov (2020) proposed an additional craniological character for the identification of sable and pine marten (Monakhov 2020). According to his data, sable and pine marten differ in the distance from the postorbital constriction to the line between the postorbital processes in the sagittal plane (character Δ) with the average value of Δ for sable being much smaller than that for pine marten. This character showed a high level (more than 97%) of correct species identifications (Monakhov 2020). Based on this character our putative hybrids were closer to *M. martes* than to *M. zibellina*. However, skull morphology of hybrids in Mustelidae is poorly studied. Usually, F1 hybrids look craniologically similar to one of the parental species. Comparative analysis of the skulls of European mink *Mustela lutreola*, polecat *Mustela putorius* and their hybrids from the north-western part of Russia showed that most of the hybrid specimens were craniologically very similar to *M. putorius* (Abramov and Tumanov 2003). Skulls of hybrids between European and Asian badgers, *Meles meles* and *M. leucurus*, look more similar to *M. leucurus* (Abramov and Puzachenko 2007).

*Issues with datings of demographic trajectories*

Dating of the demographic trajectory is always a difficult task due to difficulties with reliable estimation of generation time to the mutation rate ratio ($g/\mu$). As a reference value for the $\mu$ used 4.64 $\times 10^{-9}$ substitution per site per generation, measured from the trio of American minks (Bergeron et al. 2023). Borders of its confidence interval (CI: $2.94 \times 10^{-9}$ - $7.37 \times 10^{-9}$) we used to calculate CI for our datings. We have to note that Bergeron et al have sequenced and analyzed only a single mink trio for the estimation, and CI was inferred from a negative binomial distribution. American mink belongs to a different subfamily (Mustelinae) and our initial phylogeny reconstruction showed significantly longer branches for Mustelinae than for Guloninae (Supplementary Figure SF14A and SF14B). Therefore, mink mutation rate might not be a good estimate for marten species, but no better estimates are available. For g we found no way to estimate a CI and used a fixed g=5. So the imprecision of the generation time is not included in the CIs of our datings.

In hybrid samples (MT Figure 8B) we found a specific pattern – a steeply and high peak, which can be used as an indicator of notable introgression. Moreover, we detected a strong and significant correlation (Kendall's $\tau = 0.67$, p-value = 0.003; Spearman's $\rho = 0.76$, p-value = 0.006; Pearson's $r = 0.778$, p-value = 0.005) between height of the peak (Supplementary Figure SF9) and admixture level. It suggests that the ratio of its heights in two or more samples can be used for comparative quantification of the admixture in terms more/less. The differences in trajectories observed in hybrids are likely related not only to the number of hybrid segments but also to their merging (Cahill et al. 2016).

*Phylogeny and dating*

We reconstructed and dated the phylogenetic tree of 21 Caniformia species using Maximum Likelihood (ML) (Supplementary Figure SF14A), Bayesian (Supplementary Figure SF14B) and coalescent-based (Supplementary Figures SF15, SF16; Supplementary Table ST10) methods and 5,989 single-copy genes (codon alignment 9,707,517 bp). We found no discrepancy in topology between ML and Bayesian trees, but on the coalescent tree the *Meles meles* was placed as an outgroup to all other included Mustelidae species instead of sister position to Lutrinae + Mustelinae. However, such a topology has a low quartet support (only 40%) close to the first alternative (36%), and the effective number (EN) of genes used to resolve the branches is low as well (2563.92 or 42.81%). On the ML and Bayesian tree, the related node has a slightly reduced bootstrap support (83) and posterior probability (95), respectively.

The fossil-based calibrated time trees provide more reliable estimates, but are highly dependent on reliable fossil calibrations and a proper choice of molecular clock model. Our phylogenetic reconstruction revealed a high heterogeneity in substitution rates within Arctoidea and even within Mustelidae. Such a pattern forces usage of the independent rate model and requires multiple fossil calibrations ideally covering most of the nodes, which is unrealistic. This forced us to remove all lineages with either too low (all except *Ailurus fulgens* and Mustelidae) or too high (Mustelinae) substitution rates. Two fossil records (see *Fossil calibrations*) dating the split between Ailuridae and Mustelidae from both sides were an additional argument to retain *Ailurus fulgens*. It resulted in a reduced tree of 9 species (MT Figure 3), which was used to date divergence times between *Martes* species.

*Fossil calibrations*

For our set of nine species we constrained five nodes. Nearly all of them were limited either only from the lower boundary, or the upper boundary got a very relaxed geostratigraphic restriction (Supplementary File SF1, SRD Figure 3). The only exception was the root node, which we have constrained from both sides.

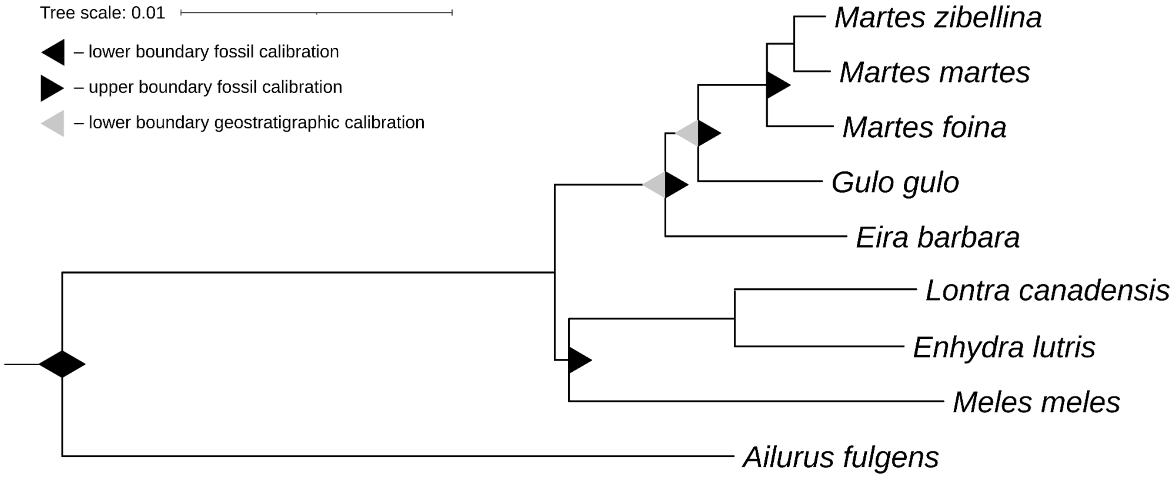


**SRD Figure 3**. Fossil calibrated nodes.

*Mustelidae – Ailuridae split*

The root node of our tree was constrained from both sides. As the elder restriction we have used *Amphicticeps dorog* (Wang et al. 2005), as the younger one (30.9 Mya) – *Mustelictis olivieri* (30.9-32.8 Mya) (Wang et al. 2005), respectively. The first calibration was considered as the stem Arcoidea in previous publications, the second one - as a sister lineage to the extant and extinct Mustelidae (Wang et al. 2005; Law et al. 2018).

*Meles – Neogale + Lutrinae split*

*Taxodon sansaniensis* (Salesa et al. 2013) was considered by Law et, 2018 as the oldest extinct species within Melinae. We used this fossil to constrain the split between *Meles meles* and Lutrinae-Neogale lineages.

*Eira – Martes + Gulo split*

Li et al, 2014 used *Pekania occulta* (Samuels and Cavin 2013) to set the lower boundary for split between *Pekania* and *Martes-Gulo* lineages. *Eira barbar*a (tayra) is considered to be a basal to all Mustelidae (Koepfli et al. 2008) species or a sister to Pekania pennanti (Law et al. 2018). Therefore, *Pekania occulta* can be used to restrict split between *Eira barbara* and *Martes-Gulo* lineages from the lower side. As upper boundary for this node we used a geostratigrafic constraint of 20.44 Mya proposed by Li et al, 2014 (Law et al. 2018).

*Calibrations within Martes genus (2 nodes)*

A paleontological evidence of undoubted extinct marten species is limited. Two main paleontological databases NOW (New and Old Worlds database of fossil mammals) (Žliobaitė et al.

2023) and Paleobiology Database (PBDB; https://paleobiodb.org) contain multiple records of unrealistically old *"Martes"* species, for example, *"Martes" sansaniensis* (14-16 Mya), *"Martes" sainjon*i (16-17 Mya), *"Martes" munki* (11-17 Mya), *"Martes" laevidens* (17-19 Mya) and others. Such datings make the assignment of these fossils to the *Martes* genus unreliable as their ages are comparable to the epoch of early radiation of Mustelidae lineages (Law et al. 2018; Hassanin et al. 2021). Tracing the related publication showed that even in the paleontological publications (Nagel et al. 2009; Kargopoulos et al. 2022) the genus names of these species are mentioned in quotes or/and with 'cf.' abbreviation (short for confer, i.e. "compare"), for example, *"Martes" cf. munki* (Kargopoulos et al. 2022) or *Martes cf. munki* (Nagel et al. 2009). Therefore, only verified fossils can be used as calibrations within the *Martes* genus. There are only three trusted records so far: *Martes wenzensis* (3.3 - 4 Mya, the European locality of Węże 1, Central Poland) (Stach 1959; Marciszak et al. 2024), *Martes crassidens* (Jinyuan Cave, Liaoning Province, Northern China) (Jiangzuo et al. 2021a), *Martes vetus* (1.75 - 2 Mya, European locality of Kielniki 3A, Southern Poland) (Wolsan 1990). Nonetheless the broader dating interval (mid-earlier pleistocene), *M. crassidens* is considered to be older than *M. vetus*. However, the conservative assignments place both of them closer to the Holarctic marten group (*M. martes*, *M. zibellina*, *M. melampus*, *M. caurina* and *M. americana*) than to *M. foina* (Jiangzuo et al. 2021a). *M. wenzensis* is treated as the most ancient verified *Martes* species, but the precise dating is unknown yet - the current estimation is based on layers and covers early-mid pleistocene interval, i.e. (1.25 - 2.5 Mya) (Jiangzuo et al. 2021b).

*Martes wenzensis* was previously used to calibrate the split between *Gulo-Martes* lineages (3.3 Mya, lower side) (Li et al. 2014). Given that recently found *M. crassidens* is considered to be younger, we did the same. Due to the relaxed lower boundary (1.25 Mya), *Martes crassidens* is the worse calibration for the split between *M. foina* and Holarctic martens than *M. vestus* (1.75 Mya), so we decided to use *M. vestus* to restrict this node from the lower side. The same constraint was used by Li et al, 2014 (Li et al. 2014).

*Tajima's D and Fst*

To detect the most differentiated loci between the pure sables and pure martens, we calculated Fst (pure sables vs pure pine martens) and Tajima's D (all samples as a single population) statistics in the sliding windows of 1 Mbp with a step of 100 kbp based *M. zibellina* genome assembly. The weighted Fst (SRD Figure 4) showed a high level of genetic differentiation between two species, with a mean value of 0.66. We detected 12 genome regions consisting of windows with high Fst values (>=0.9; Supplementary Table ST7), ranging from 1 to 3.1 Mbp in length and predominantly gene-rich. Among them we found three loci (FST3, FST6, FST11) enriched by proteins with specific GO terms (Supplementary File SF12). The most interesting region, FST11 (chr9: 86,600,000–88,100,000), containing 48 genes, revealed a statistically significant enrichment in four GO categories: keratinization (GO:0031424), intermediate filament organization (GO:0045109), embryonic skeletal system development (GO:0048706) and anterior/posterior pattern specification (GO:0009952). Further examination of the gene list showed that it includes a cluster of HOX-C genes (homeobox C cluster) and seven cytokeratin type I and II genes: KRT76, KRT3, KRT4, KRT79, KRT78, KRT8 (type II) and KRT18 (type I).

Tajima's D (all samples as a single population, SRD Figure 5A and D) metric revealed only a single region (TJD1) with significantly high value (>=2). It is located on the chr1, encompasses 1.3 Mbp and is nested in the high Fst loci FST4 (Supplementary Table ST7). Although no GO enrichment was found for this region, among its 20 genes is FGFR3, which plays a key role in bone development and maintenance (Supplementary File SF12).

We also calculated Tajima's D in 1 Mbp windows with a 100 kbp step separately for pure sables and pure pine martens (SRD Figure 5B, E, C, F). The mean values were -0.52 for sables and -0.41 for pine martens. This analysis revealed 35 regions (TJD2–TJD36) with significantly negative Tajima's D values (D < -2). Of these, seven (TJD2, 5, 7, 14, 25, 30, and 34) showed enrichment for proteins with specific GO terms (Supplementary File SF12).

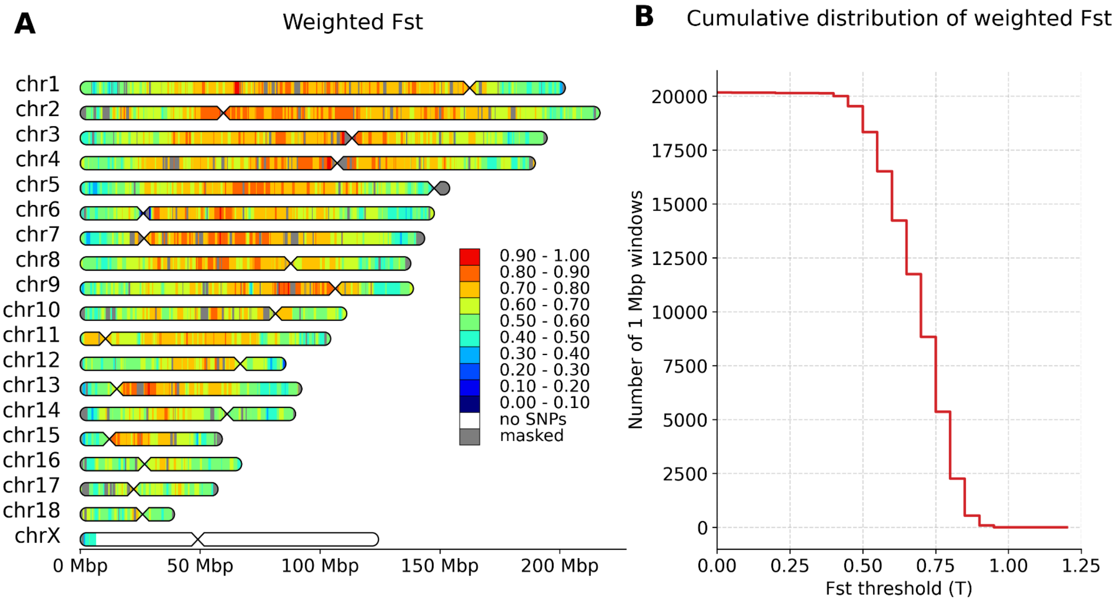


**SRD Figure 4**. Heatmap (A) and cumulative distribution (B) of weighted Fst (pure sables vs pure pine martens).

Estimates were calculated in the sliding windows of 1 Mbp with a step of 100 kbp

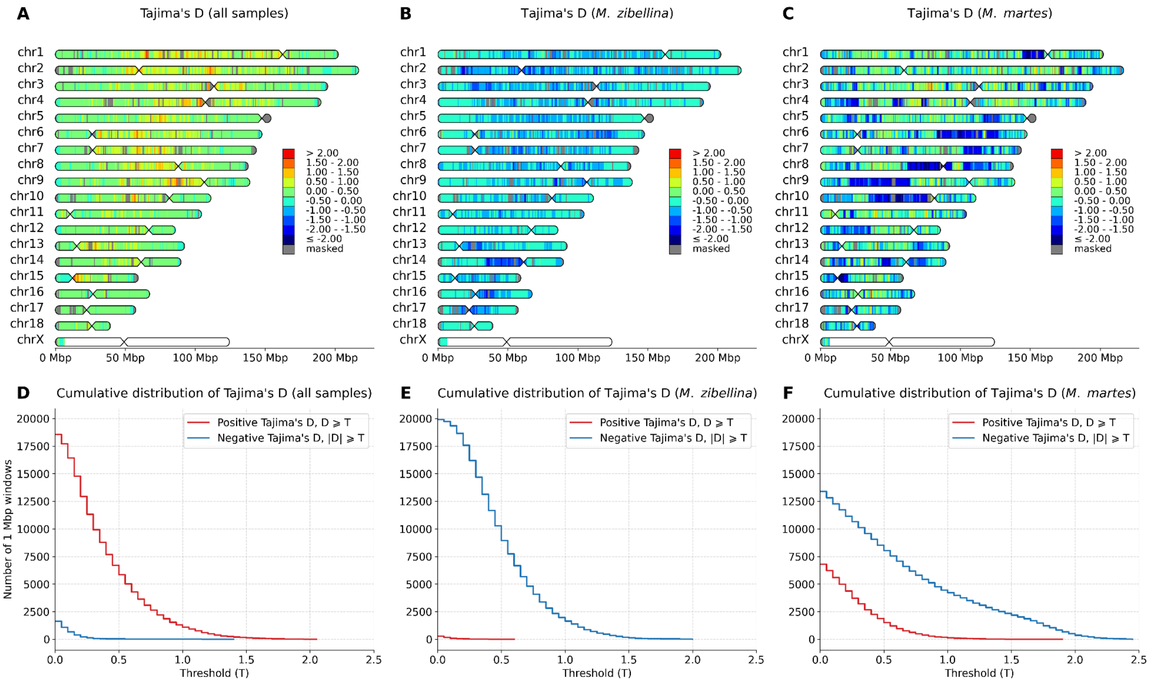


**SRD Figure 5**. Distribution of Tajima's D values on chromosomes.

A, B, C – Tajima's D for all samples, pure sables and pure pine martens, respectively; D, E, F – cumulative distributions of Tajima's D for the same groups. Tajima's D values were calculated in the sliding windows of 1 Mbp with a step of 100 kbp

Previously reported synteny analysis revealed an inversion on chr 11 (cytogenetically confirmed) between the sable and the pine marten (Tomarovsky et al. in prep). Chromosomal inversions in a heterozygous state are known to alter the functional regulation of gene expression and to suppress recombination within the inverted region (so-called "crossover suppression"), leading to the gradual accumulation of differences between species (Berdan et al. 2023). In the location of the inversion and its relatively small size (11.5 Mbp), we initially hypothesized that in a heterozygous state it would contribute to crossover suppression and promote the accumulation of species-specific differences within the inverted region. To test this hypothesis, we compared genetic differentiation in the p-arm of chr 11 (inversion) with the p-arms of other chromosomes with comparable size of p-arm (no more than twice as long). For this purpose, we analyzed the distributions of Fst values (SRD Figure 6) and found statistically significant differences (p-values << 0.001) between the p-arm of chr 11 and other chromosomes (SRD Table 1). After Bonferroni correction for multiple testing (adjusted for four comparisons), the p-values remained significant.

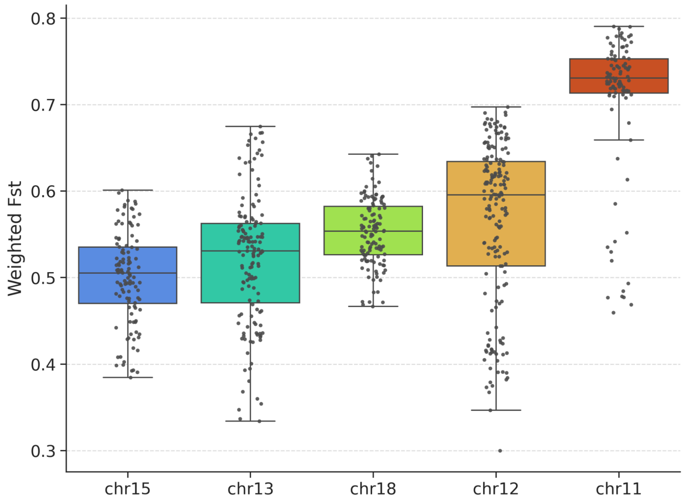


**SRD Figure 6**. Distribution of Fst estimates for p-arms of chr11, 12, 13, 15 and 18.

Distributions of weighted Fst (sables and pines) values were counted in sliding windows of 1 Mbp with 100 kbp step (Reference: *M. zibellina*)

**SRD Table 1**. Comparison of Fst values for the p-arm of chr 11 with estimates on the p-arms of other chromosomes.

| Chr | Coordinates of p-arm (*M. zibellina*) | Size of p-arm, bp | Number of windows | One-sided Mann–Whitney U test, p-value | |
|---|---|---|---|---|---|
| | | | | raw | corrected* |
| chr11 | 0 - 10553932 | 10’553’932 | 96 | NA | NA |
| chr12 | 67017660 - 85782648 | 18’764’988 | 177 | 4.38E-27 | 1.75E-26 |
| chr13 | 0 - 15232218 | 15’232’218 | 143 | 3.55E-28 | 1.42E-27 |
| chr15 | 0 - 12185599 | 12’185’599 | 112 | 1.45E-27 | 5.78E-27 |
| chr18 | 26068618 - 39344853 | 13’276’235 | 123 | 1.26E-23 | 5.04E-23 |

* corrected – Bonferroni-adjusted p-values.
