## Supplementary Figures for "Genomics of sable (*Martes zibellina)* × pine marten (*Martes martes*) hybridization"

* corresponding author

[x] equal contribution

**Supplementary Figure SF1.** Distribution of 23-mers for filtered data of sables (A), pine martens (B) and sympatric martens (C).

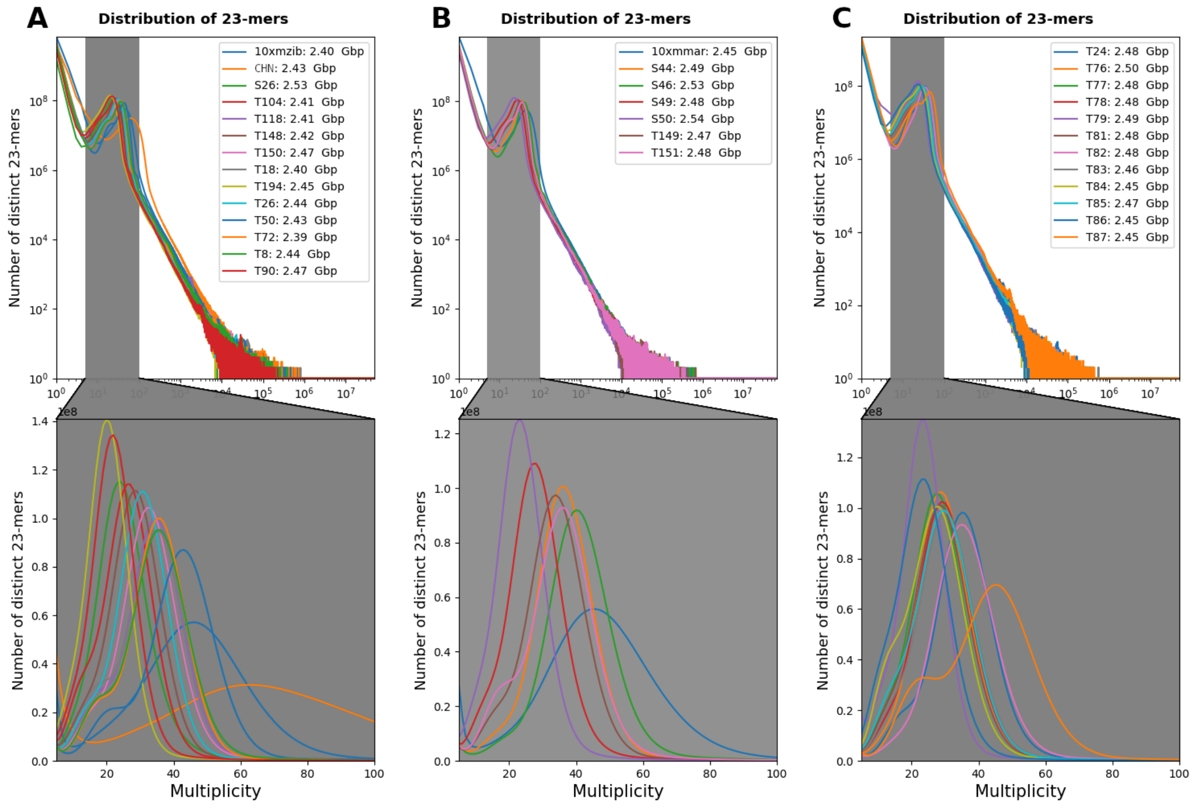


**Supplementary Figure SF2**. Time-calibrated phylogenetic trees for global and correlated clock models.

Types of molecular clocks: A – global; B – correlated. Branch lengths are proportional to time, with divergence dates shown in millions of years. For each node, the node ID is displayed above the branch, and the estimated divergence time (100 Mya) is shown below. Node bars indicate the 95% highest posterior density (HPD) intervals for node age uncertainty.

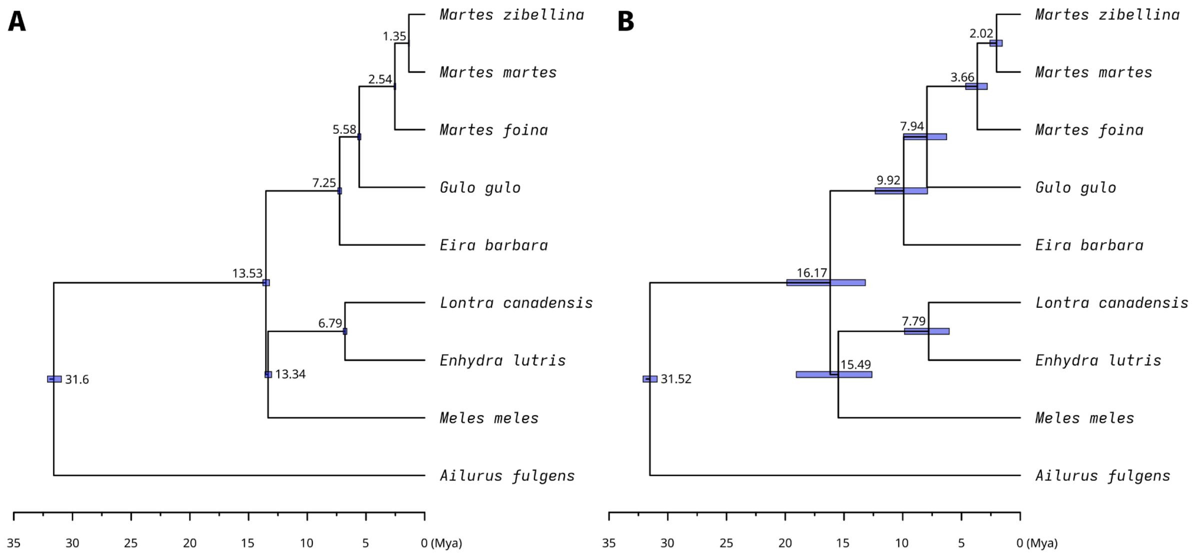

**Supplementary Figure SF3.** Principal component analysis (PCA) plot based on 6,750,722 SNPs obtained from autosomes and the pseudoautosomal region (PAR) of the *M. zibellina* assembly.

The first and second principal components are represented on the X and Y axes, respectively. The percentage of variance explained on each axis is indicated in parentheses. Plot with sample labels.

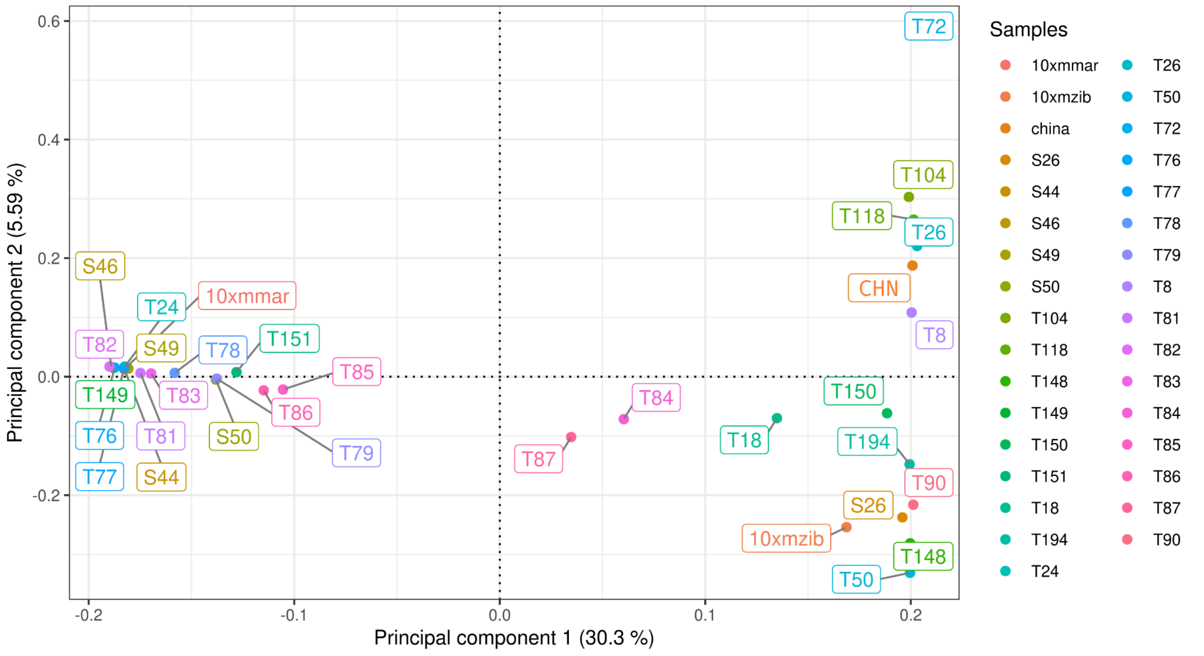

**Supplementary Figure SF4.** Fitting models for heterozygosity distributions.

Red triangles indicate mean values for the distribution of individual model components.

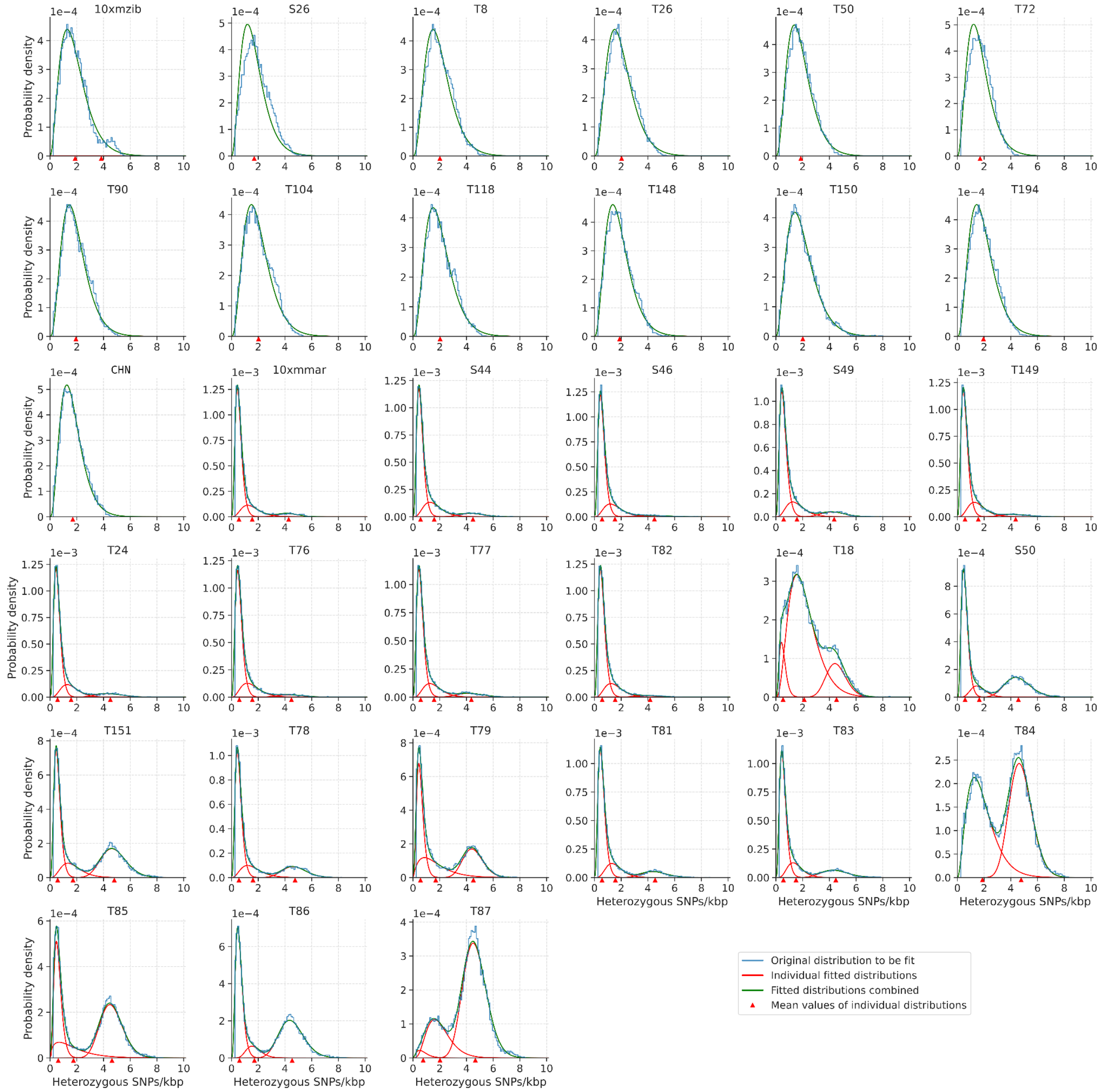

**Supplementary Figure SF5.** Localization of markers and admixture analysis for three sets of STR loci (All, Rozhnov's, Kashatonov's).

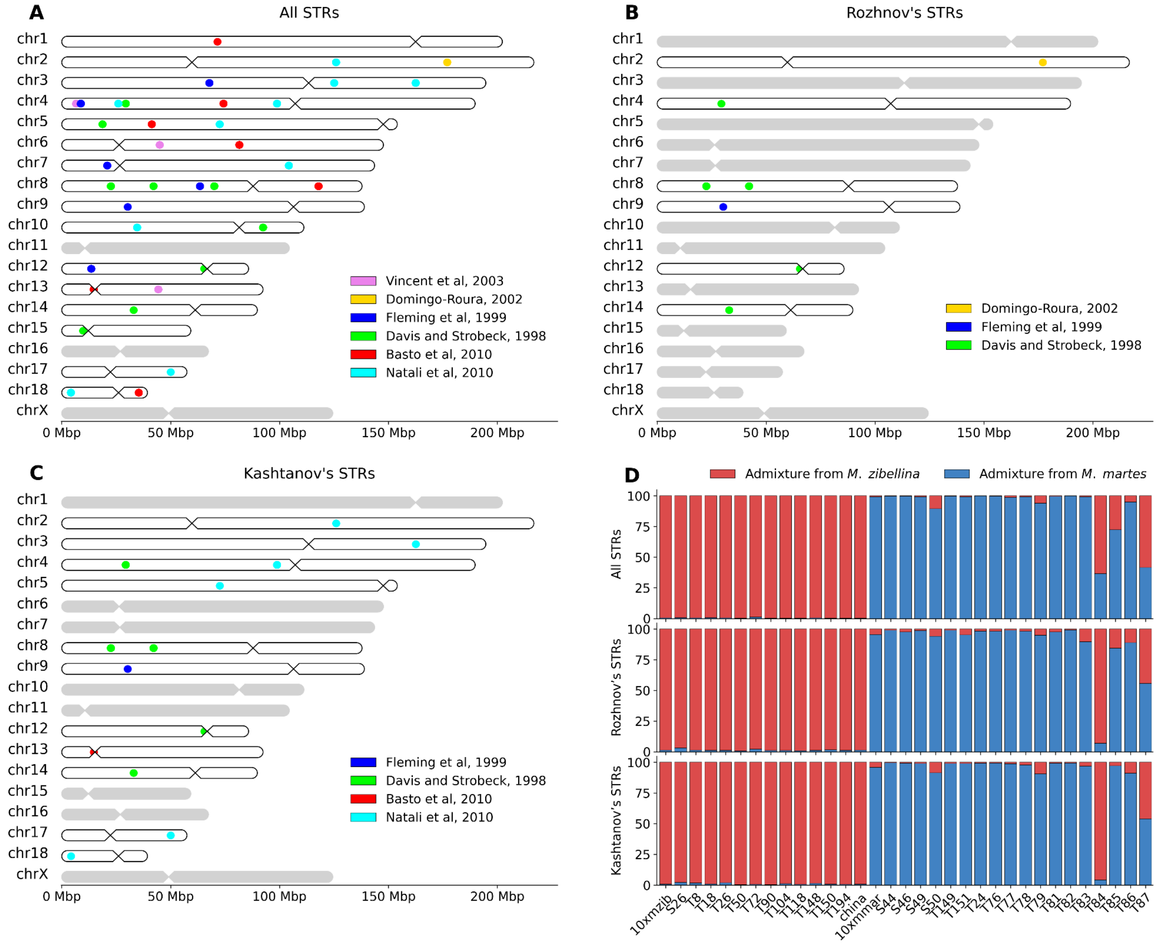

**Supplementary figure SF6.** Mitochondrial phylogenetic tree.

Nodes with more than 90 bootstrap support are marked in green, and nodes between 70 and 90 are marked in yellow. Nodes with support values less than 70 were deleted. Three main clade groups (A, B, and C) were identified, which included subclades (A1-5, B1-4 and C1-4). The studied samples are color-coded: *M. zibellina* samples with mtDNA from *M. zibellina* are shown in purple; *M. martes* samples with mtDNA from *M. martes* are shown in blue; *M. martes* samples with mtDNA from *M. zibellina* are shown in red; hybrids with mtDNA from *M. zibellina* are shown in green. *M. foina* (NC_020643.1) was used as an outgroup (not shown).

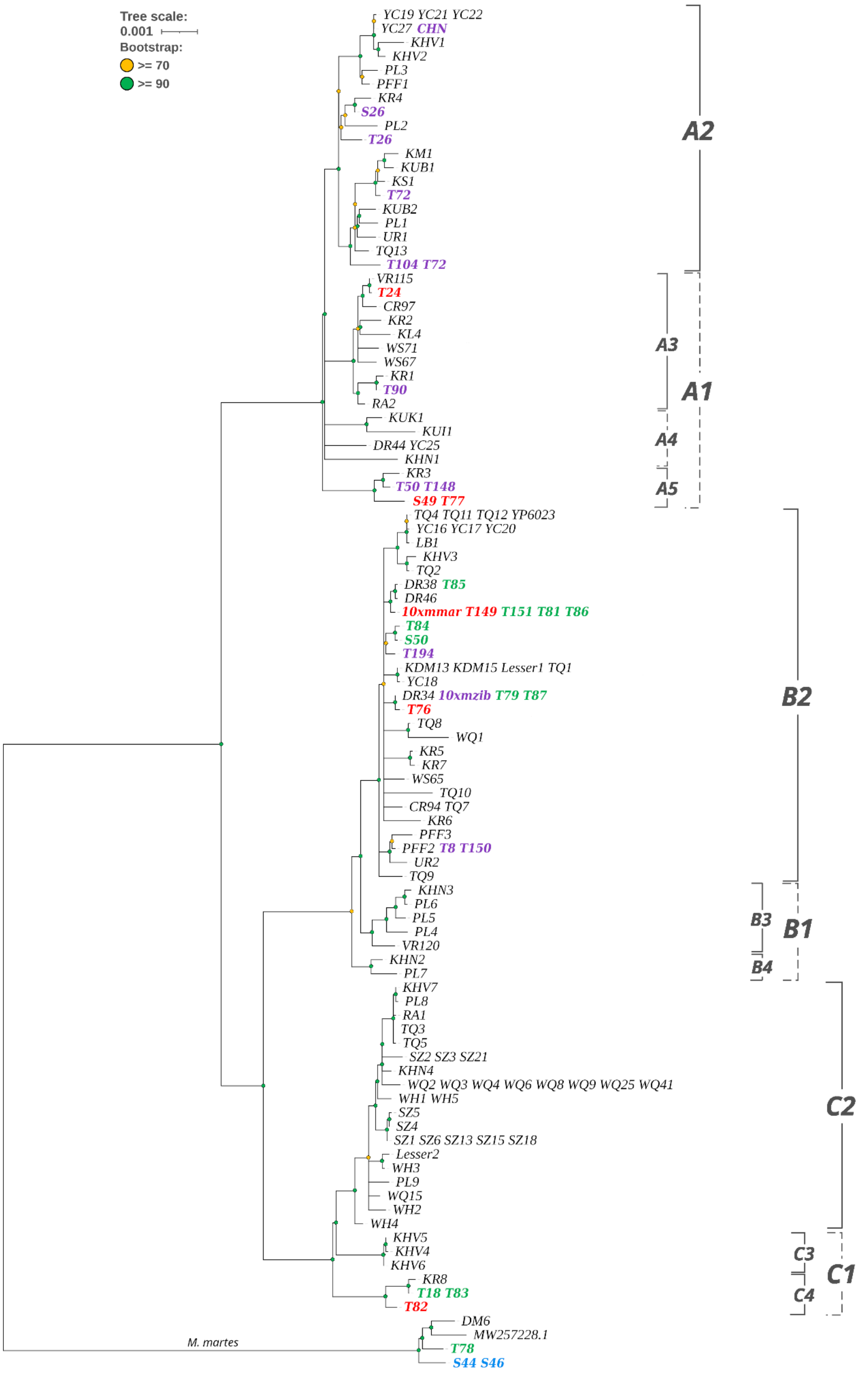

**Supplementary Figure SF7**. Distributions of mean heterozygosity (SNP only) counted in sliding windows of 1 Mb with 100 kbp step for both *M. zibellina* (red) and *M. martes* (blue) genome assemblies.

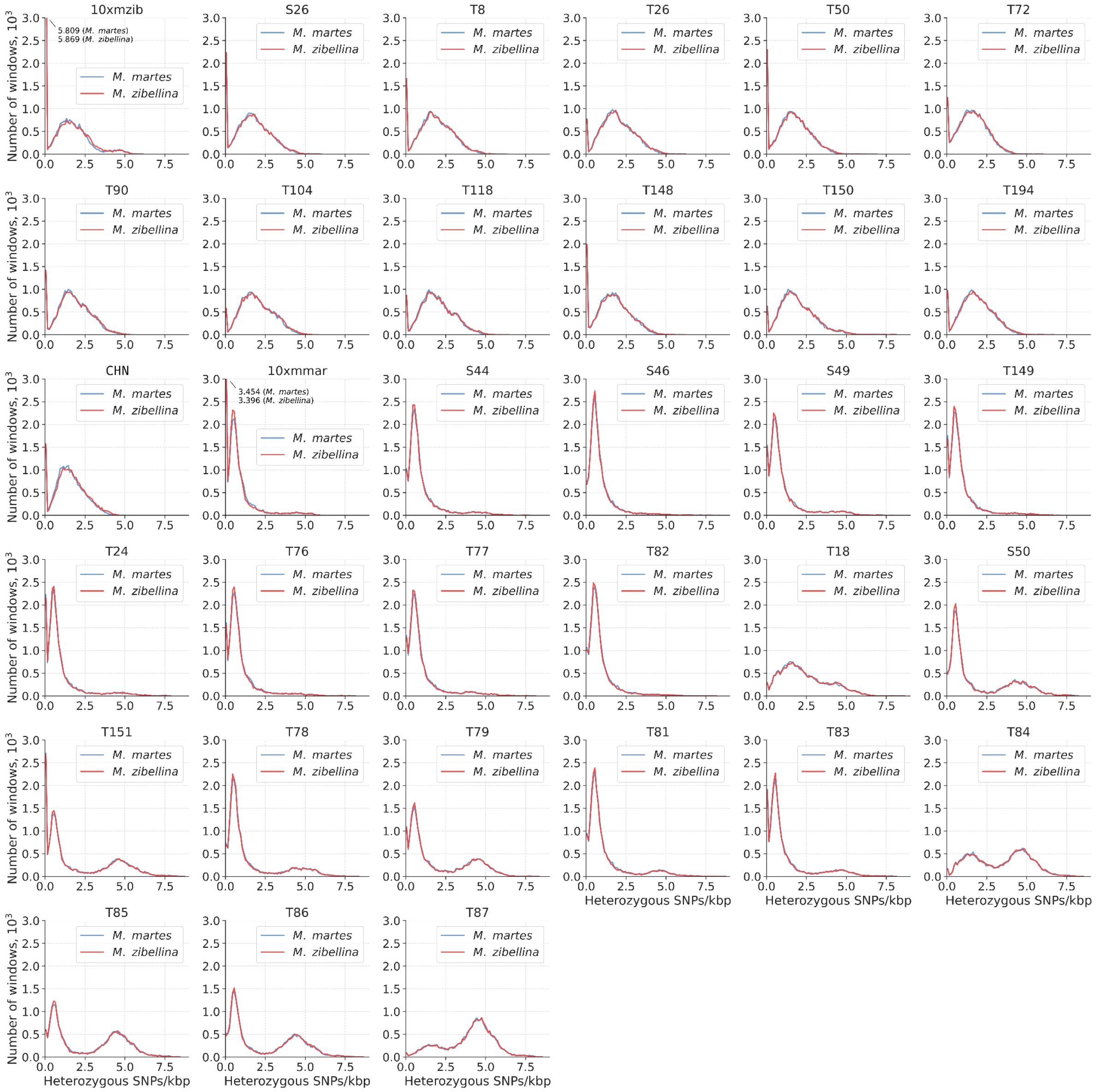

**Supplementary Figure SF8.** Runs of homozygosity (RoH).

A – Cumulative lengths (fraction of the genome) of the RoH of different categories: ultra long (L ≥ 10 Mbp), long (10 Mbp > L ≥ 1 Mbp) and short (L < 1 Mbp), in all samples , B-D – cumulative distribution of RoH length in the pure martens, pure sables and hybrids, E-J – localization of the RoH in the specific samples: 10xmmar, 10xmzib, T87, S46, T50 and T72. The X chromosome was excluded from all samples.

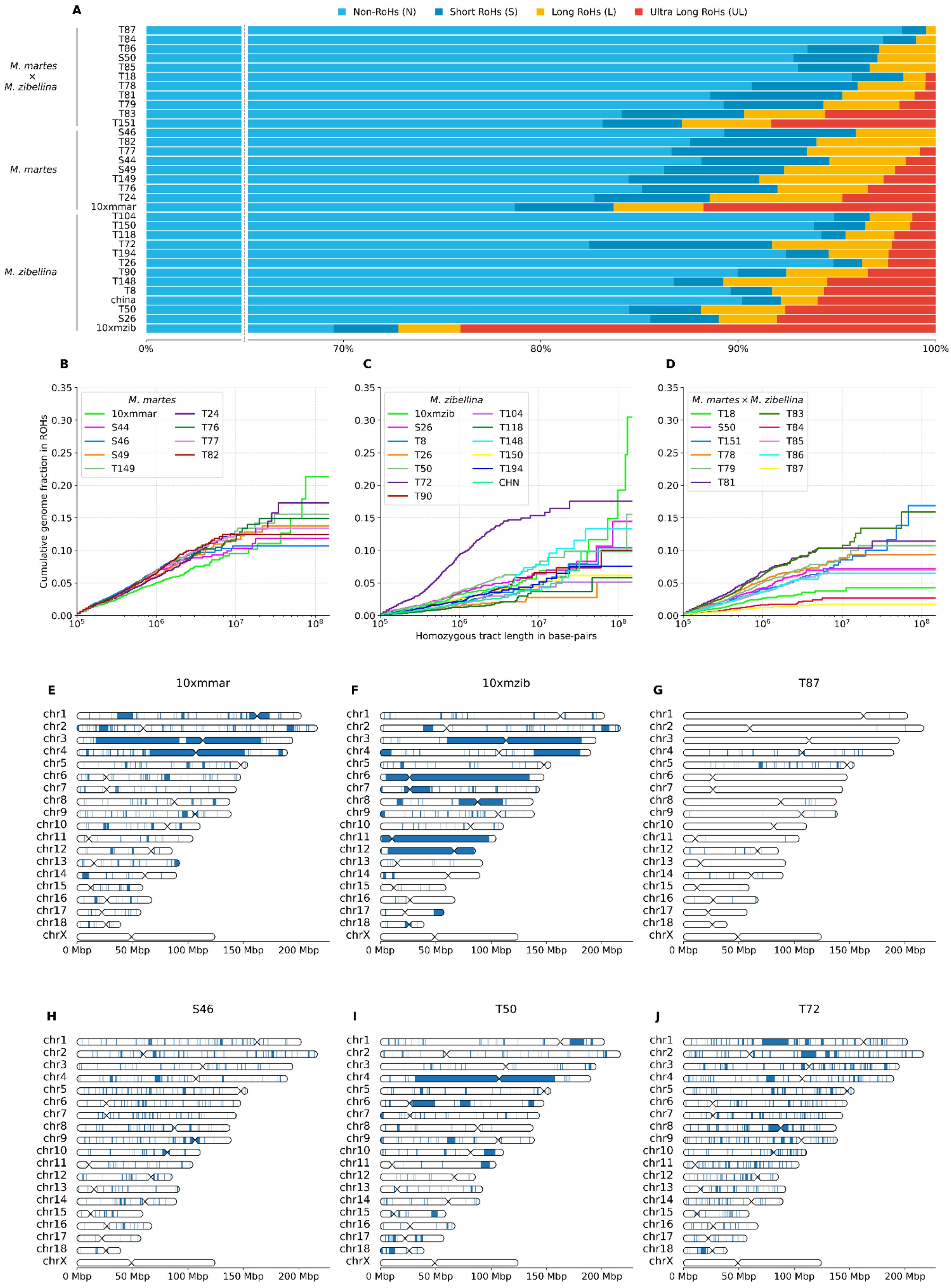

**Supplementary Figure SF9.** Correlation between the introgression level and height of the “population explosion” peak for the hybrids.

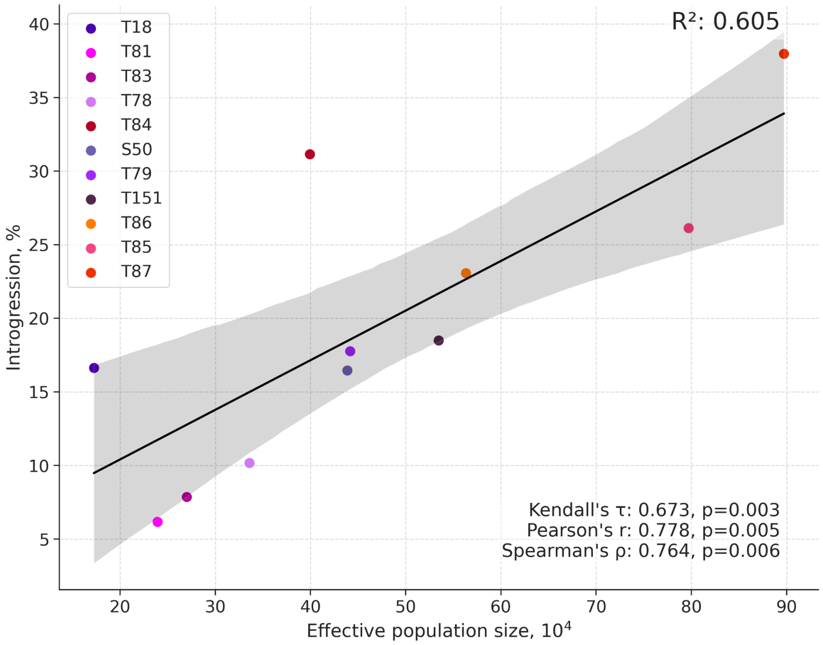


**Supplementary Figure SF10.** Scoped demographic history reconstruction based on *M. zibellina* genome assembly.

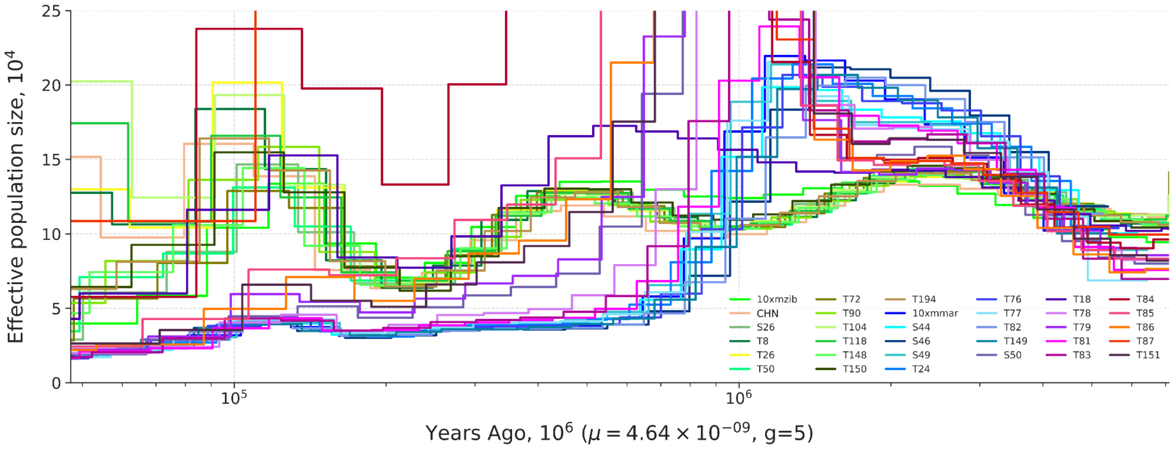

**Supplementary Figure SF11.** Species map with local ADMIXTURE results for each sample.

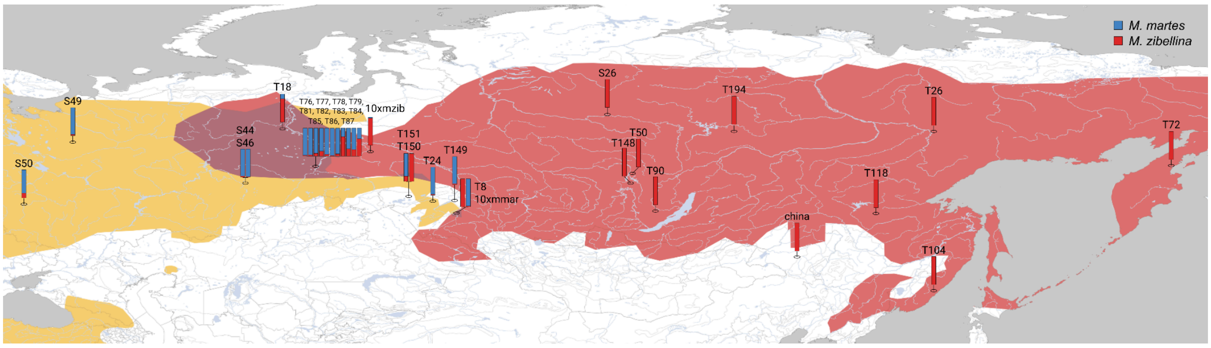


**Supplementary Figure SF12.** Histograms of mean heterozygosity (SNP only) for each species.

The X-axis represents the number of SNPs per kbp, calculated in overlapping 1 Mbp windows with a 100 kbp step, while the Y-axis indicates the corresponding number of windows

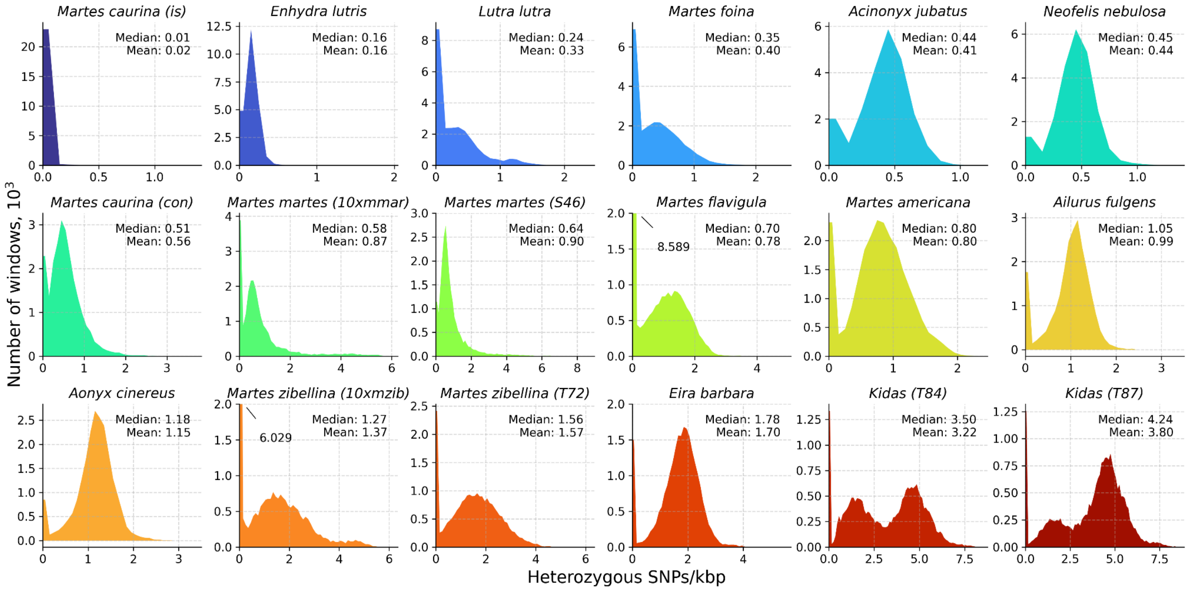

**Supplementary Figure SF13.** Comparison of coverage masking of sable and pine marten genome assemblies for classified groups of individuals.

Absolute values are provided in Supplementary Table ST11. Statistics in Supplementary Table ST14.

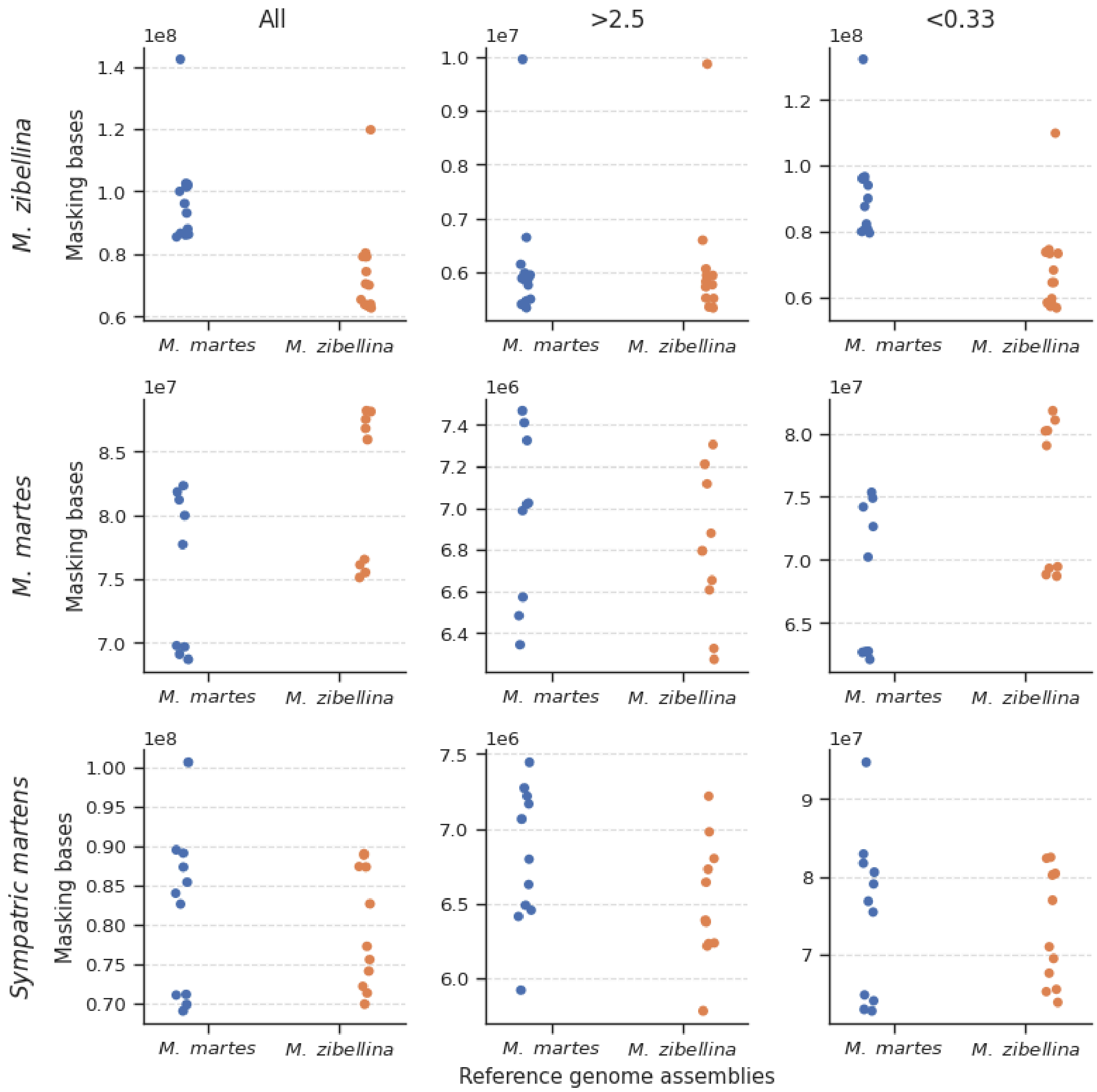

**Supplementary Figure SF14.** Phylogenetic trees based on data matrix of 5989 common single-copy BUSCOs.

A – IQtree (ML), B – MrBayes (BI).

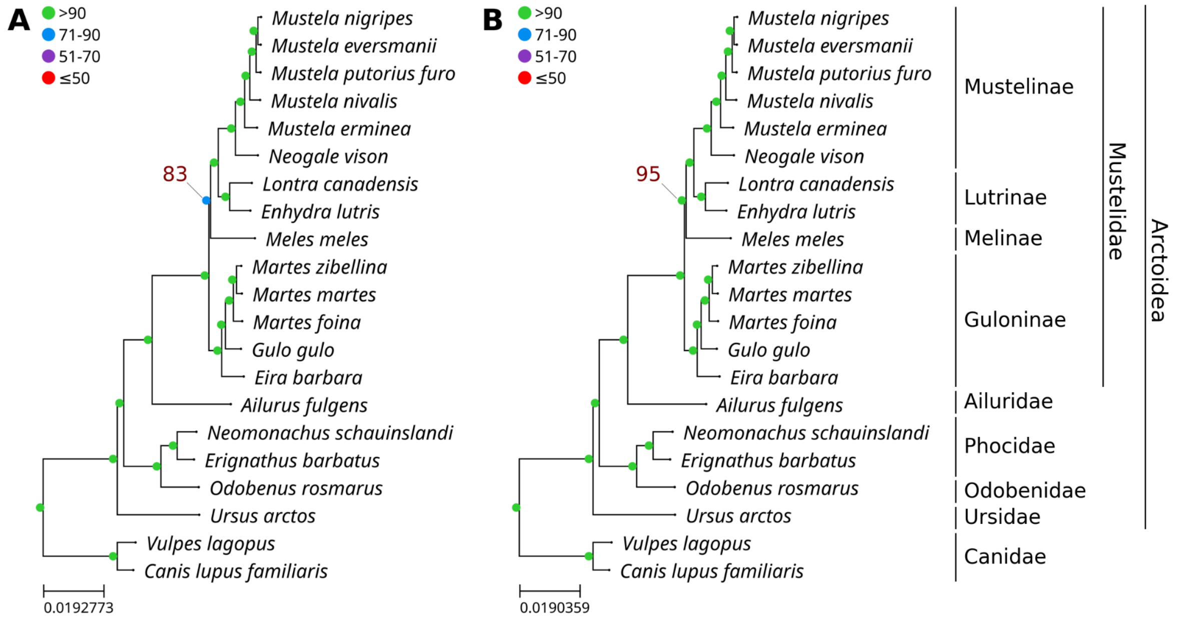


**Supplementary Figure SF15.** ASTRAL-III phylogenetic tree was reconstructed based on 5989 single-copy BUSCO gene alignments.

Individual gene trees were inferred using IQTree (1000 bootstraps). Nodes with bootstrap support values below 70 were excluded from the input trees. Numbers displayed at each node represent unique node identifiers, for which detailed statistical values are provided in Supplementary Table ST13. Colored dots indicate the main local posterior probabilities (pp1), expressed as percentages.

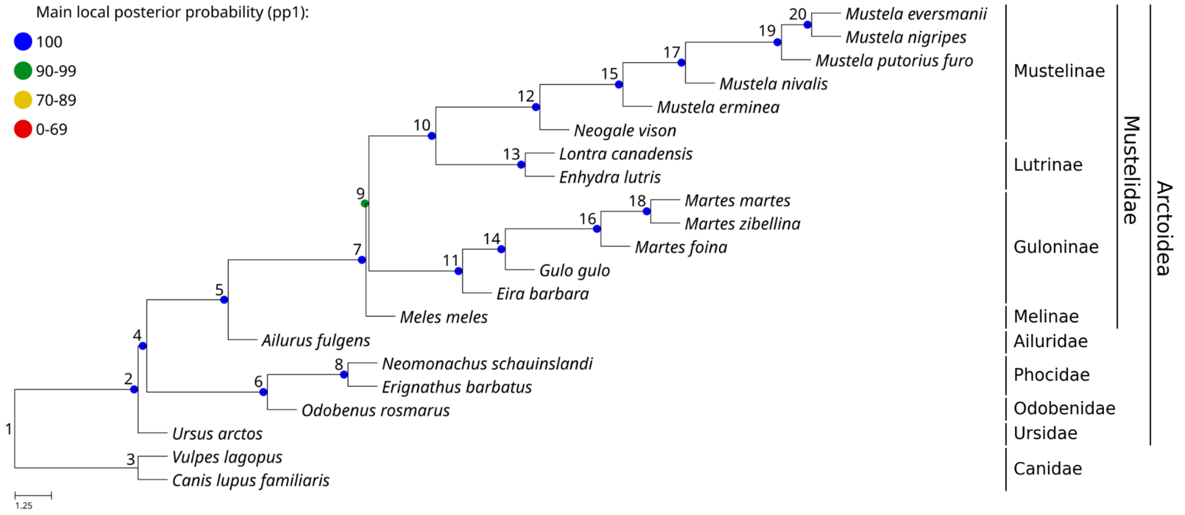

**Supplementary Figure SF16.** ASTRAL-III phylogenetic tree was reconstructed based on 5989 single-copy BUSCO gene alignments.

Individual gene trees were inferred using IQTree (1000 bootstraps). Nodes with bootstrap support values below 70 were excluded from the input trees. Detailed statistical values are provided in Supplementary Table ST13. Piecharts indicate the quartet supports.

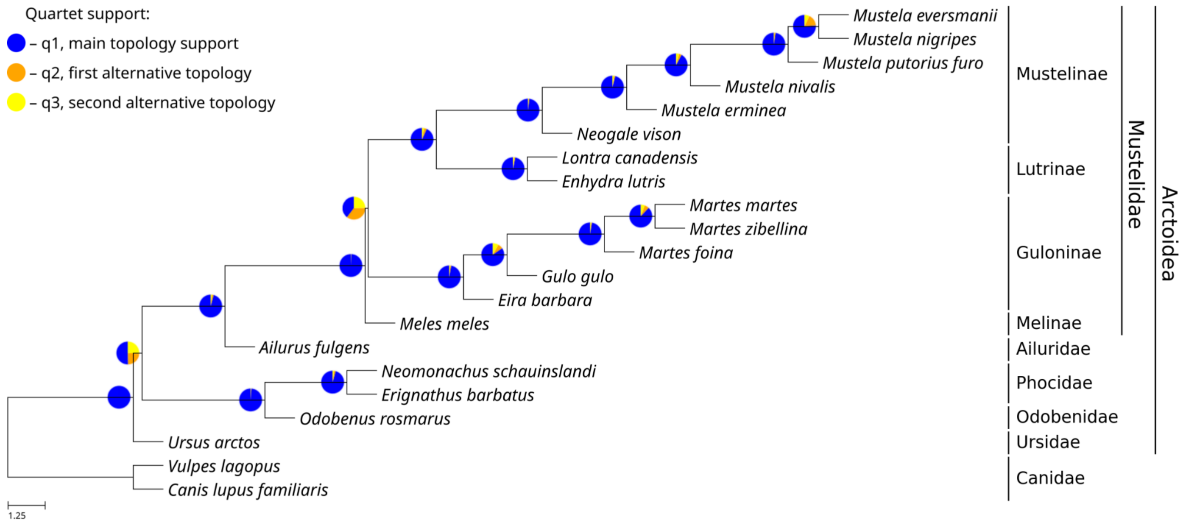
