## Supplementary Tables for "Genomics of sable (*Martes zibellina)* × pine marten (*Martes martes*) hybridization"

**Supplementary Table ST1. Sample information.**

| Species * | Data type ** | ID | SRA / BioProject ID | Origin | Point on map | Sex ***** |
|---|---|---|---|---|---|---|
| *M. zibellina* | 10X | 10xmzib*** | SRR22412799 | Khanty-Mansi Autonomous Okrug – Yugra, Ugut | 17 | F |
| *M. zibellina* | PE | S26 | SRR28749757 | Krasnoyarsk Krai, Evenkia Autonomous Okrug, Baikit | 5 | F |
| *M. zibellina* | PE | T8 | SRR28749756 | Altai Krai complex "Magistralny" | 15 | M |
| *M. zibellina* | PE | T18 | SRR28749745 | Khanty-Mansi Autonomous Okrug – Yugra, Peregrebnoe | 14 | M |
| *M. zibellina* | PE | T26 | SRR28749734 | Sakha – Yakutia, Okhotskiy Perevoz | 8 | F |
| *M. zibellina* | PE | T50 | SRR28749733 | Irkutsk Oblast, Nevon | 13 | M |
| *M. zibellina* | PE | T72 | SRR28749732 | Kamchatskiy Krai, Ivashka | 6 | M |
| *M. zibellina* | PE | T90 | SRR28749731 | Irkutsk Oblast, Naran | 12 | F |
| *M. zibellina* | PE | T104 | SRR28749730 | Primorskiy Krai, Terney | 16 | M |
| *M. zibellina* | PE | T118 | SRR28749729 | Amur Oblast, Dugda | 4 | M |
| *M. zibellina* | PE | T148 | SRR28749728 | Irkutsk region, Ust'-Ilimsk | 18 | F |
| *M. zibellina* | PE | T150 | SRR28749755 | Novosibirsk Oblast, Kuibyshev | 9 | F |
| *M. zibellina* | PE | T194 | SRR28749754 | Sakha – Yakutia, Mirnyi | 11 | F |
| *M. zibellina* | PE | CHN **** | PRJNA495455 | Greater Khingan Mountains, China | 21 | F |
| *M. martes* | 10X | 10xmmar *** | SRR22412800 | Altai Krai, Barnaul, "Lesnaya skazka" zoo | 1 | F |
| *M. martes* | PE | S44 | SRR28749753 | Sverdlovsk Oblast, Verkhnyaya Pyshma | 20 | F |
| *M. martes* | PE | S46 | SRR28749752 | Sverdlovsk Oblast, Verkhnyaya Pyshma | 20 | F |
| *M. martes* | PE | S49 | SRR28749751 | Arkhangelsk Oblast, Velsk | 19 | M |
| *M. martes* | PE | S50 | SRR28749750 | Kaluga Oblast, Kaluga | 7 | F |
| *M. martes* | PE | T149 | SRR28749749 | Novosibirsk Oblast, Baryshevo | 2 | M |
| *M. martes* | PE | T151 | SRR28749748 | Novosibirsk Oblast, Kuibyshev | 9 | M |
| *Putative hybrid* | PE | T24 | SRR28749747 | Novosibirsk Oblast, Chulym | 3 | M |
| *Putative hybrid* | PE | T76 | SRR28749746 | Tyumen Oblast, Malyi Narys | 10 | F |
| *Putative hybrid* | PE | T77 | SRR28749744 | Tyumen Oblast, Malyi Narys | 10 | M |
| *Putative hybrid* | PE | T78 | SRR28749743 | Tyumen Oblast, Malyi Narys | 10 | F |
| *Putative hybrid* | PE | T79 | SRR28749742 | Tyumen Oblast, Malyi Narys | 10 | F |
| *Putative hybrid* | PE | T81 | SRR28749741 | Tyumen Oblast, Malyi Narys | 10 | M |

| | | | | | | |
|---|---|---|---|---|---|---|
| *Putative hybrid* | PE | T82 | SRR28749740 | Tyumen Oblast, Malyi Narys | 10 | F |
| *Putative hybrid* | PE | T83 | SRR28749739 | Tyumen Oblast, Malyi Narys | 10 | M |
| *Putative hybrid* | PE | T84 | SRR28749738 | Tyumen Oblast, Malyi Narys | 10 | M |
| *Putative hybrid* | PE | T85 | SRR28749737 | Tyumen Oblast, Malyi Narys | 10 | F |
| *Putative hybrid* | PE | T86 | SRR28749736 | Tyumen Oblast, Malyi Narys | 10 | M |
| *Putative hybrid* | PE | T87 | SRR28749735 | Tyumen Oblast, Malyi Narys | 10 | M |

* The initial classification of the samples, based on simple morphological traits: tail length, fur length and quality and throat patch size.

** Data type: “PE” – paired end Illumina reads, “10X” – 10x Genomics linked reads;

*** Reference individuals, previously published in (Tomarovsky et al. in prep);

**** Previously published sample (Liu et al. 2020);

***** Sex: M – male, F – female;

| Supplementary Table ST2. Samples quality. Number of reads, k-mers coverage, genome sizes and downsampling fraction. | | | | | | | | | |
|---|---|---|---|---|---|---|---|---|---|
| **Sample** | **Read length** | **Number of raw reads, mln** | **Number of filtered reads, mln** | **Saved reads, %** | **Kmer multiplicity at first maximum** | **Estimated haplome coverage (genomescope2)** | **Coverage (genomescope2)** | **Genome size (genomescope2), Gbp** | **Downsampling fraction** |
| 10xmmar | 128+150 | 633.01 | 578.26 | 91.35 | 45 | 24.81 | 49.62 | 2.45 | 0.44 |
| S44 | 150+150 | 380.48 | 369.5 | 97.11 | 36 | 18.66 | 37.32 | 2.49 | 0.59 |
| S46 | 150+150 | 427.28 | 417.27 | 97.66 | 40 | 20.8 | 41.6 | 2.53 | 0.53 |
| S49 | 150+150 | 291.26 | 284.51 | 97.68 | 28 | 14.46 | 28.92 | 2.48 | 0.76 |
| S50 | 150+150 | 250.61 | 244.71 | 97.65 | 23 | 12.18 | 24.36 | 2.54 | 0.9 |
| T149 | 150+150 | 351.28 | 343.66 | 97.83 | 34 | 17.59 | 35.18 | 2.47 | 0.63 |
| T151 | 150+150 | 377.11 | 367.49 | 97.45 | 36 | 18.68 | 37.36 | 2.48 | 0.59 |
| 10xmzib | 128+150 | 619.13 | 566.65 | 91.52 | 46 | 24.85 | 49.7 | 2.4 | 0.44 |
| CHN | 120+120 | 977.18 | 955.63 | 97.79 | 63 | 38.39 | 76.78 | 2.43 | 0.29 |
| S26 | 150+150 | 269.04 | 261.32 | 97.13 | 24 | 12.66 | 25.32 | 2.53 | 0.87 |
| T104 | 150+150 | 275.03 | 266.11 | 96.76 | 27 | 13.87 | 27.74 | 2.41 | 0.79 |
| T118 | 150+150 | 352.81 | 342.09 | 96.96 | 35 | 17.82 | 35.64 | 2.41 | 0.62 |
| T148 | 150+150 | 294.84 | 286.42 | 97.15 | 29 | 14.84 | 29.68 | 2.42 | 0.74 |
| T150 | 150+150 | 341 | 331.36 | 97.17 | 33 | 16.88 | 33.76 | 2.47 | 0.65 |
| T18 | 150+150 | 356.65 | 345.64 | 96.91 | 35 | 18.07 | 36.14 | 2.4 | 0.61 |
| T194 | 150+150 | 214.47 | 207.9 | 96.93 | 20 | 10.62 | 21.24 | 2.45 | NA |
| T26 | 150+150 | 319.08 | 309.51 | 97 | 31 | 15.91 | 31.82 | 2.44 | 0.69 |
| T50 | 150+150 | 437.93 | 423.14 | 96.62 | 43 | 22.01 | 44.02 | 2.43 | 0.5 |
| T72 | 150+150 | 361.13 | 349.85 | 96.88 | 36 | 18.34 | 36.68 | 2.39 | 0.6 |
| T8 | 150+150 | 366.9 | 357.24 | 97.37 | 36 | 18.44 | 36.88 | 2.44 | 0.6 |
| T90 | 150+150 | 232.63 | 225.76 | 97.05 | 22 | 11.49 | 22.98 | 2.47 | 0.96 |
| T24 | 150+150 | 376.9 | 365.39 | 96.95 | 35 | 18.24 | 36.48 | 2.48 | 0.6 |
| T76 | 150+150 | 307.97 | 302.5 | 98.22 | 29 | 15.07 | 30.14 | 2.5 | 0.73 |
| T77 | 150+150 | 295.04 | 289.54 | 98.14 | 28 | 14.53 | 29.06 | 2.48 | 0.76 |
| T78 | 150+150 | 312.95 | 306.79 | 98.03 | 29 | 15.4 | 30.8 | 2.48 | 0.71 |
| T79 | 150+150 | 253.9 | 245.87 | 96.84 | 23 | 12.14 | 24.28 | 2.49 | 0.91 |
| T81 | 150+150 | 305.17 | 299.3 | 98.08 | 28 | 15.06 | 30.12 | 2.48 | 0.73 |
| T82 | 150+150 | 370.46 | 363.71 | 98.18 | 35 | 18.29 | 36.58 | 2.48 | 0.6 |
| T83 | 150+150 | 317.82 | 311.74 | 98.08 | 30 | 15.81 | 31.62 | 2.46 | 0.7 |
| T84 | 150+150 | 292.12 | 287.13 | 98.29 | 28 | 14.63 | 29.26 | 2.45 | 0.75 |
| T85 | 150+150 | 315.36 | 309.73 | 98.21 | 30 | 15.63 | 31.26 | 2.47 | 0.7 |
| T86 | 150+150 | 251 | 246.61 | 98.25 | 23 | 12.58 | 25.16 | 2.45 | 0.87 |
| T87 | 150+150 | 471.25 | 462.92 | 98.23 | 45 | 23.59 | 47.18 | 2.45 | 0.47 |

| |
|---|
| * NA – not applicable, i.e. downsampling was unnecessary. |

**Supplementary Table ST3. Dates of species divergence (in millions of years) for correlated, independent and global molecular clocks.**

| Node ID | Description | Correlated | | | Independent | | | Global | | |
|---|---|---|---|---|---|---|---|---|---|---|
| | | Date | HPD Lower | HPD Upper | Date | HPD Lower | HPD Upper | Date | HPD Lower | HPD Upper |
| 1 | MRCA of M. martes and M. zibellina | 2.02 | 1.53 | 2.58 | 1.52 | 1.05 | 2.06 | 1.35 | 1.31 | 1.39 |
| 2 | MRCA of M. foina, M. zibellina and M. martes | 3.66 | 2.81 | 4.65 | 2.83 | 2.11 | 3.66 | 2.54 | 2.47 | 2.6 |
| 3 | MRCA of G. gulo and Martes genus | 7.94 | 6.27 | 9.94 | 6.57 | 5.16 | 8.27 | 5.58 | 5.45 | 5.7 |
| 4 | MRCA of Guloninae | 9.92 | 7.89 | 12.35 | 8.32 | 6.7 | 10.33 | 7.25 | 7.08 | 7.4 |
| 5 | MRCA of E. lutris and L. canadensis | 7.79 | 6.04 | 9.86 | 7.45 | 5.64 | 9.59 | 6.79 | 6.63 | 6.93 |
| 6 | MRCA of Lutrinae and Melinae | 15.49 | 12.62 | 19.06 | 14.64 | 12.12 | 17.99 | 13.34 | 13.04 | 13.6 |
| 7 | MRCA of Guloninae, Lutrinae and Melinae | 16.17 | 13.18 | 19.87 | 15.34 | 12.68 | 18.8 | 13.53 | 13.22 | 13.78 |
| 8 | MRCA of Ailuridae and Mustelidae | 31.52 | 30.9 | 32.1 | 31.53 | 30.91 | 32.11 | 31.6 | 30.94 | 32.13 |

**Supplementary Table ST4. Global (whole genome) and local (in sliding 1 Mbp windows with 100 kbp step) ADMIXTURE (k = 2).**

| Sample | Global ADMIXTURE | | Local ADMIXTURE | |
|---|---|---|---|---|
| | **M. zibellina, %** | ***M. martes, %*** | **M. zibellina, %** | ***M. martes, %*** |
| 10xmzib | 100 | 0 | 96.7 | 3.3 |
| S26 | 100 | 0 | 99.71 | 0.29 |
| T8 | 100 | 0 | 99.88 | 0.12 |
| T18 | 84.9 | 15.1 | 83.36 | 16.64 |
| T26 | 100 | 0 | 99.86 | 0.14 |
| T50 | 100 | 0 | 99.89 | 0.11 |
| T72 | 100 | 0 | 99.89 | 0.11 |
| T90 | 100 | 0 | 99.89 | 0.11 |
| T104 | 100 | 0 | 99.86 | 0.14 |
| T118 | 100 | 0 | 99.86 | 0.14 |
| T148 | 100 | 0 | 99.87 | 0.13 |
| T150 | 100 | 0 | 98.38 | 1.62 |
| T194 | 100 | 0 | 99.83 | 0.17 |
| CHN | 100 | 0 | 99.87 | 0.13 |
| 10xmmar | 0 | 100 | 3.39 | 96.61 |
| S44 | 0 | 100 | 3.85 | 96.15 |
| S46 | 0 | 100 | 1.91 | 98.09 |
| S49 | 0 | 100 | 4.34 | 95.66 |
| S50 | 11.3 | 88.7 | 16.47 | 83.53 |
| T149 | 0 | 100 | 2.75 | 97.25 |
| T151 | 13.4 | 86.6 | 18.49 | 81.51 |
| T24 | 0 | 100 | 3.83 | 96.17 |
| T76 | 0 | 100 | 2.84 | 97.16 |
| T77 | 0 | 100 | 4.32 | 95.68 |
| T78 | 3.9 | 96.1 | 10.18 | 89.82 |
| T79 | 10.2 | 89.8 | 17.76 | 82.24 |

| | | | | |
|---|---|---|---|---|
| T81 | 0 | 100 | 6.17 | 93.83 |
| T82 | 0 | 100 | 2.13 | 97.87 |
| T83 | 0 | 100 | 7.86 | 92.14 |
| T84 | 65.9 | 34.1 | 68.85 | 31.15 |
| T85 | 20.2 | 79.8 | 26.14 | 73.86 |
| T86 | 17.8 | 82.2 | 23.07 | 76.93 |
| T87 | 59.2 | 40.8 | 62.03 | 37.97 |

**Supplementary Table ST5. Total number of heterozygous SNPs. Mean and median heterozygosity (SNPs/kbp) calculated in 1 Mbp windows with 100 kbp step.**

| Sample | Number of hetSNPs*, mln | Mean heterozygosity | Median heterozygosity |
|---|---|---|---|
| 10xmzib | 3.12 | 1.42 | 1.34 |
| S26 | 3.89 | 1.74 | 1.69 |
| T8 | 4.23 | 1.9 | 1.85 |
| T26 | 4.44 | 2 | 1.9 |
| T50 | 3.81 | 1.71 | 1.68 |
| T72 | 3.67 | 1.65 | 1.62 |
| T90 | 4.09 | 1.85 | 1.76 |
| T104 | 4.47 | 2.02 | 1.9 |
| T118 | 4.42 | 1.99 | 1.88 |
| T148 | 3.88 | 1.75 | 1.7 |
| T150 | 4.45 | 2 | 1.82 |
| T194 | 4.23 | 1.91 | 1.83 |
| CHN | 3.53 | 1.63 | 1.55 |
| 10xmmar | 2.01 | 0.84 | 0.56 |
| S44 | 2.42 | 1.03 | 0.66 |
| S46 | 2.14 | 0.9 | 0.64 |
| S49 | 2.51 | 1.07 | 0.67 |
| T149 | 2.17 | 0.92 | 0.64 |
| T24 | 2.33 | 0.99 | 0.62 |
| T76 | 2.24 | 0.95 | 0.65 |
| T77 | 2.45 | 1.04 | 0.66 |
| T82 | 2.09 | 0.88 | 0.64 |
| T18 | 5.31 | 2.38 | 2.08 |
| S50 | 4.6 | 2 | 0.89 |
| T151 | 4.9 | 2.14 | 0.94 |

| | | | |
|---|---|---|---|
| T78 | 3.58 | 1.55 | 0.73 |
| T79 | 4.64 | 2.03 | 1.02 |
| T81 | 2.81 | 1.2 | 0.68 |
| T83 | 2.92 | 1.25 | 0.67 |
| T84 | 7.62 | 3.39 | 3.69 |
| T85 | 6.23 | 2.73 | 2.86 |
| T86 | 5.66 | 2.47 | 1.54 |
| T87 | 9.01 | 3.99 | 4.31 |
| * heterozygous SNPs | | | |

| Supplementary Table ST6. RoHs content. | | | | | | | |
|---|---|---|---|---|---|---|---|
| Sample | Number of RoH | Length, Mbp | RoH, % | Non-RoH, % | % of RoH * | | |
| | | | | | Short | Long | Ultra Long |
| 10xmmar | 513 | 476.1 | 21.3 | 78.7 | 5 | 4.5 | 11.8 |
| S44 | 625 | 264.7 | 11.9 | 88.1 | 6.5 | 3.9 | 1.5 |
| S46 | 667 | 238.9 | 10.7 | 89.3 | 6.7 | 4 | 0 |
| S49 | 600 | 307.3 | 13.8 | 86.2 | 6.1 | 5.6 | 2.1 |
| T149 | 644 | 347.2 | 15.5 | 84.5 | 6.6 | 6.3 | 2.6 |
| T24 | 598 | 385.8 | 17.3 | 82.7 | 5.8 | 6.7 | 4.7 |
| T76 | 652 | 332.2 | 14.9 | 85.1 | 6.9 | 4.6 | 3.4 |
| T77 | 632 | 298.9 | 13.4 | 86.6 | 6.8 | 5.7 | 0.8 |
| T82 | 647 | 277.7 | 12.4 | 87.6 | 6.4 | 6 | 0 |
| 10xmzib | 294 | 681.3 | 30.5 | 69.5 | 3.3 | 3.1 | 24.1 |
| S26 | 345 | 322.7 | 14.5 | 85.5 | 3.5 | 3 | 8 |
| T8 | 196 | 232 | 10.4 | 89.6 | 2.1 | 2.6 | 5.7 |
| T26 | 140 | 115.5 | 5.2 | 94.8 | 1.5 | 1.3 | 2.4 |
| T50 | 351 | 346.7 | 15.5 | 84.5 | 3.6 | 4.3 | 7.6 |
| T72 | 769 | 392.1 | 17.6 | 82.4 | 9.3 | 6.1 | 2.2 |
| T90 | 257 | 224.2 | 10 | 90 | 2.5 | 4.1 | 3.5 |
| T104 | 162 | 114.7 | 5.1 | 94.9 | 1.8 | 2.2 | 1.2 |
| T118 | 125 | 129.3 | 5.8 | 94.2 | 1.2 | 2.5 | 2.1 |
| T148 | 267 | 296.7 | 13.3 | 86.7 | 2.5 | 5.3 | 5.5 |
| T150 | 231 | 137.6 | 6.2 | 93.8 | 2.6 | 2.3 | 1.3 |
| T194 | 227 | 169.1 | 7.6 | 92.4 | 2.2 | 3 | 2.4 |
| CHN | 187 | 219 | 9.8 | 90.2 | 2 | 1.9 | 6 |
| T18 | 256 | 95 | 4.3 | 95.7 | 2.6 | 1.1 | 0.5 |
| S50 | 436 | 160.7 | 7.2 | 92.8 | 4.3 | 2.9 | 0 |
| T151 | 400 | 376.9 | 16.9 | 83.1 | 4 | 4.5 | 8.3 |
| T78 | 524 | 207.8 | 9.3 | 90.7 | 5.3 | 3.4 | 0.5 |

| | | | | | | | |
|---|---|---|---|---|---|---|---|
| T79 | 463 | 239.8 | 10.7 | 89.3 | 5 | 3.9 | 1.8 |
| T81 | 629 | 255.1 | 11.4 | 88.6 | 6.7 | 3.7 | 1.1 |
| T83 | 582 | 355.2 | 15.9 | 84.1 | 6.2 | 4.1 | 5.6 |
| T84 | 155 | 59.5 | 2.7 | 97.3 | 1.7 | 1 | 0 |
| T85 | 371 | 155.7 | 7 | 93 | 3.6 | 3.3 | 0 |
| T86 | 366 | 145 | 6.5 | 93.5 | 3.6 | 2.9 | 0 |
| T87 | 125 | 38 | 1.7 | 98.3 | 1.2 | 0.5 | 0 |
| * Short RoH (< 1 Mbp), Long RoH (>= 1 Mbp < 10 Mbp) and Ultra Long RoH (>= 10 Mbp). | | | | | | | |

**Supplementary Table ST7. Candidate regions, associated with differences between the sable and the pine marten**

| Region ID | Description | Chr | Start | End | Length (Mbp) | Number of genes | Mean Fst | Mean Tajima's D |
|---|---|---|---|---|---|---|---|---|
| INV1 | Inversion | chr11 | 2079555 | 13548101 | 11.5 | 31 | 0.74 | 0.44 |
| INV2 | Inversion | chr12 | 78576795 | 85165887 | 6.6 | 44 | 0.48 | 0.1 |
| INV3 | Inversion | chr15 | 4271212 | 12359108 | 8.1 | 77 | 0.53 | -0.24 |
| **TJD1** | **High Tajima's D** | **chr1** | **65400000** | **66700000** | **1.3** | **20** | **0.93** | **2.05** |
| FST1 | High Fst | chr2 | 58200000 | 59300000 | 1.1 | 9 | 0.9 | 1.47 |
| FST2 | High Fst | chr2 | 60500000 | 61800000 | 1.3 | 6 | 0.9 | 1.61 |
| *FST3* | *High Fst* | *chr2* | *111100000* | *112100000* | *1* | *26* | *0.9* | *1.62* |
| **FST4** | **High Fst** | **chr1** | **64600000** | **67100000** | **2.5** | **39** | **0.92** | **1.9** |
| FST5 | High Fst | chr3 | 109200000 | 111100000 | 1.9 | 14 | 0.92 | 1.47 |
| *FST6* | *High Fst* | *chr3* | *112600000* | *115700000* | *3.1* | *27* | *0.91* | *1.26* |
| FST7 | High Fst | chr4 | 103000000 | 105400000 | 2.4 | 79 | 0.91 | 1.62 |
| FST8 | High Fst | chr6 | 58200000 | 59600000 | 1.4 | 32 | 0.91 | 1.72 |
| FST9 | High Fst | chr7 | 59800000 | 60900000 | 1.1 | 23 | 0.9 | 1.29 |
| FST10 | High Fst | chr9 | 82800000 | 84100000 | 1.3 | 34 | 0.91 | 1.41 |
| *FST11* | *High Fst* | *chr9* | *86600000* | *88100000* | *1.5* | *48* | *0.91* | *1.73* |
| FST12 | High Fst | chr13 | 28100000 | 29900000 | 1.8 | 53 | 0.9 | 1.46 |

**Bold** markers nested loci with both high Tajima’s D (TJD1) and high FST (FST1)

*Italic* highlights a loci with GO term enrichment.

| **Supplementary Table ST8. Estimated divergence times between Martes species (M. zibellina, M. martes and M. foina) across phylogenetic studies.** | | | | |
|---|---|---|---|---|
| **Node** | **Study** | **Divergence Time (Mya)** | **CI** | **Markers** |
| *M. zibellina* - *M. martes* | (Law et al. 2018) | 1.06 | 0.66 - 1.55 | 46 genes (4 mitochondrial and 42 nuclear genes) |
| | (Hassanin et al. 2021) | 1.1 | unavailable | whole mtDNA |
| | (Koepfli et al. 2008) | 1.1 | 0.6 - 1.6 | 22 mitochondrial genes |
| | (Li et al. 2014) | 0.68 | 0.54 - 0.84 | whole mtDNA |
| | This study (correlated) | 2.05 | 1.52 - 2.70 | 4-fold degenerated sites from 5989 genes |
| | This study (independent) | 1.49 | 1.05 - 1.98 | 4-fold degenerated sites from 5989 genes |
| *M. foina* - *M. zibellina + M. martes* | (Law et al. 2018) | 2.56 | 1.93 - 3.29 | 46 genes (4 mitochondrial and 42 nuclear genes) |
| | (Hassanin et al. 2021) | 5.1 | unavailable | whole mtDNA |
| | (Koepfli et al. 2008) | 2.8 | 1.9 - 3.8 | 22 mitochondrial genes |
| | (Li et al. 2014) | 2.93 | 2.35 - 3.54 | whole mtDNA |
| | This study (correlated) | 3.72 | 2.78 - 4.84 | 4-fold degenerated sites from 5989 genes |
| | This study (independent) | 2.77 | 2.14 - 3.55 | 4-fold degenerated sites from 5989 genes |

| **Supplementary Table ST9. Species used for whole-genome phylogenetic reconstruction and dating.** | | | |
|---|---|---|---|
| **Species** | **Level of assembly** | **ID** | **Source** |
| *Martes martes* | Chromosome | DNAzoo | (Tomarovsky et al. in prep) |
| *Martes zibellina* | Chromosome | DNAzoo | (Tomarovsky et al. in prep) |
| *Martes foina* | Chromosome | DNAzoo | (Tomarovsky et al. 2025) |
| *Mustela putorius furo* | Chromosome | DNAzoo | (Peng et al. 2014; Dudchenko et al. 2017; Dudchenko et al. 2018) |
| *Mustela nigripes* | Chromosome | DNAzoo | (Kliver et al. 2023) |
| *Mustela nivalis* | Scaffold | GCA_019141155.1 | (Miranda et al. 2021) |
| *Mustela erminea* | Chromosome | GCA_009829155.1 | (Rhie et al. 2021) |
| *Neogale vison* | Chromosome | GCA_020171115.1 | (Karimi et al. 2022) |
| *Gulo gulo* | PseudoChromosome | GCA_024510155.1 | (Lok et al. 2022) |
| *Meles meles* | Chromosome | GCA_922984935.1 | (Newman et al. 2022) |
| *Ailurus fulgens* | Chromosome | DNAzoo | (Dudchenko et al. 2017; Hu et al. 2017; Dudchenko et al. 2018) |
| *Enhydra lutris* | Chromosome | DNAzoo | (Dudchenko et al. 2017; Jones et al. 2017; Dudchenko et al. 2018) |
| *Erignathus barbatus* | Chromosome | DNAzoo | (Dudchenko et al. 2017) |
| *Lontra canadensis* | Chromosome | DNAzoo | (Dudchenko et al. 2017) |
| *Neomonachus schauinslandi* | Chromosome | GCA_002201575.2 | (Mohr et al. 2022) |
| *Odobenus rosmarus* | Chromosome | DNAzoo | (Foote et al. 2015; Dudchenko et al. 2017; Dudchenko et al. 2018) |
| *Ursus arctos* | Chromosome | DNAzoo | (Dudchenko et al. 2017; Dudchenko et al. 2018; Taylor et al. 2018) |
| *Canis lupus familiaris* | Chromosome | DNAzoo | (Dudchenko et al. 2017; Dudchenko et al. 2018) |
| *Mustela eversmanii* | Scaffold | GCA_963422785.1 | (Derežanin et al. in prep) |
| *Eira barbara* | Scaffold | GCA_020311275.1 | (Derežanin et al. 2022) |
| *Vulpes lagopus* | Chromosome | GCA_018345385.1 | (Peng et al. 2021) |

**Supplementary Table ST10. Summary of support metrics for each internal branch in the ASTRAL species tree.**

The table reports quartet support values (q1, q2, q3), representing the proportion of gene tree quartets supporting the main topology and two alternatives; the corresponding counts of supporting quartets across all gene trees (f1, f2, f3); local posterior probabilities for each topology (pp1, pp2, pp3); the total number of informative quartets (QC); and the effective number of genes contributing to each branch (EN).

| Node | q1 | q2 | q3 | f1 | f2 | f3 | pp1 | pp2 | pp3 | QC | EN |
|---|---|---|---|---|---|---|---|---|---|---|---|
| 1 | - | - | - | - | - | - | - | - | - | - | - |
| 2 | 0.999786 | 0.000093 | 0.000121 | 5,977.94 | 0.556 | 0.722 | 1 | 0 | 0 | 18 | 5,979.22 |
| 3 | - | - | - | - | - | - | - | - | - | - | - |
| 4 | 0.504337 | 0.238758 | 0.256905 | 1,800.62 | 852.433 | 917.222 | 1 | 0 | 0 | 90 | 3,570.28 |
| 5 | 0.959436 | 0.021889 | 0.018674 | 4,961.98 | 113.206 | 96.579 | 1 | 0 | 0 | 126 | 5,171.76 |
| 6 | 0.989571 | 0.005326 | 0.005103 | 5,617.54 | 30.233 | 28.967 | 1 | 0 | 0 | 90 | 5,676.74 |
| 7 | 0.994269 | 0.002648 | 0.003083 | 5,763.06 | 15.346 | 17.872 | 1 | 0 | 0 | 78 | 5,796.28 |
| 8 | 0.958165 | 0.021062 | 0.020773 | 5,327.61 | 117.111 | 115.5 | 1 | 0 | 0 | 18 | 5,560.22 |
| 9 | 0.397202 | 0.355236 | 0.247562 | 1,018.39 | 910.796 | 634.729 | 0.999999998 | 0.000000002 | 0 | 280 | 2,563.92 |
| 10 | 0.933255 | 0.036117 | 0.030628 | 3,699.09 | 143.154 | 121.4 | 1 | 0 | 0 | 480 | 3,963.64 |
| 11 | 0.973431 | 0.014125 | 0.012444 | 4,774.00 | 69.273 | 61.031 | 1 | 0 | 0 | 256 | 4,904.31 |
| 12 | 0.981458 | 0.009831 | 0.008711 | 5,221.82 | 52.308 | 46.346 | 1 | 0 | 0 | 130 | 5,320.48 |
| 13 | 0.969222 | 0.013683 | 0.017095 | 4,774.03 | 67.397 | 84.205 | 1 | 0 | 0 | 78 | 4,925.63 |
| 14 | 0.846536 | 0.071584 | 0.08188 | 3,030.33 | 256.25 | 293.104 | 1 | 0 | 0 | 48 | 3,579.69 |
| 15 | 0.961569 | 0.01665 | 0.021782 | 4,730.97 | 81.917 | 107.167 | 1 | 0 | 0 | 60 | 4,920.05 |
| 16 | 0.97518 | 0.010772 | 0.014048 | 4,649.03 | 51.353 | 66.971 | 1 | 0 | 0 | 34 | 4,767.35 |
| 17 | 0.922293 | 0.036949 | 0.040758 | 3,980.77 | 159.479 | 175.917 | 1 | 0 | 0 | 48 | 4,316.17 |
| 18 | 0.879455 | 0.064054 | 0.056491 | 3,165.50 | 230.556 | 203.333 | 1 | 0 | 0 | 18 | 3,599.39 |
| 19 | 0.975488 | 0.01443 | 0.010082 | 4,598.82 | 68.029 | 47.529 | 1 | 0 | 0 | 34 | 4,714.38 |
| 20 | 0.762988 | 0.16628 | 0.070733 | 2,227.50 | 485.444 | 206.5 | 1 | 0 | 0 | 18 | 2,919.44 |

| Supplementary Table ST11. Whole genome coverage statistics. | | | | |
|---|---|---|---|---|
| **Sample** | **Median** | **Mean** | **Max** | **Min** |
| 10xmmar | 23 | 23.91 | 78676 | 0 |
| S44 | 22 | 22.43 | 60435 | 0 |
| S46 | 22 | 22.51 | 64371 | 0 |
| S49 | 21 | 21.19 | 73018 | 0 |
| S50 | 21 | 21.42 | 59636 | 0 |
| T149 | 21 | 21.54 | 62018 | 0 |
| T151 | 22 | 22.59 | 68279 | 0 |
| 10xmzib | 23 | 23.34 | 70122 | 0 |
| CHN | 20 | 21.07 | 41207 | 0 |
| S26 | 23 | 23.71 | 59272 | 0 |
| T104 | 21 | 21.34 | 55593 | 0 |
| T118 | 22 | 22.22 | 49561 | 0 |
| T148 | 22 | 22.05 | 46180 | 0 |
| T150 | 22 | 22.56 | 49432 | 0 |
| T18 | 22 | 21.89 | 43685 | 0 |
| T194 | 21 | 21.02 | 64791 | 0 |
| T26 | 22 | 22.57 | 37577 | 0 |
| T50 | 22 | 22.32 | 55577 | 0 |
| T72 | 23 | 22.68 | 44591 | 0 |
| T8 | 22 | 22.44 | 59392 | 0 |
| T90 | 21 | 21.57 | 49280 | 0 |
| T24 | 23 | 23.51 | 50622 | 0 |
| T76 | 20 | 21.03 | 47816 | 0 |
| T77 | 20 | 20.68 | 53836 | 0 |
| T78 | 20 | 20.41 | 50296 | 0 |
| T79 | 24 | 24.25 | 60921 | 0 |
| T81 | 20 | 20.29 | 59904 | 0 |

| | | | | |
|---|---|---|---|---|
| T82 | 20 | 20.85 | 61273 | 0 |
| T83 | 20 | 20.54 | 71353 | 0 |
| T84 | 20 | 20.7 | 67450 | 0 |
| T85 | 20 | 20.29 | 47703 | 0 |
| T86 | 19 | 19.79 | 55570 | 0 |
| T87 | 21 | 21.19 | 52188 | 0 |

**Supplementary Table ST12. Coverage masked bases of all samples for both references.**

The table shows the total number of masked bases (“all”), bases with coverage <33% (“<0.33”) and >250% (“>2.5”) of the median genome coverage.

| Sample | *M. martes* | | | *M. zibellina* | | |
|---|---|---|---|---|---|---|
| | **all** | **>2.5** | **<0.33** | **all** | **>2.5** | **<0.33** |
| 10xmmar | 77690575 | 7467627 | 70222948 | 87535669 | 7305340 | 80230329 |
| S44 | 69089660 | 6343178 | 62746482 | 75107591 | 6273326 | 68834265 |
| S46 | 69657638 | 6987587 | 62670051 | 76100527 | 6653635 | 69446892 |
| S49 | 81211898 | 7019223 | 74192675 | 86805493 | 6606650 | 80198843 |
| S50 | 69897077 | 7064285 | 62832792 | 72228119 | 6799836 | 65428283 |
| T149 | 79969683 | 7324894 | 72644789 | 85938085 | 6879475 | 79058610 |
| T151 | 87339644 | 6795080 | 80544564 | 88869345 | 6386397 | 82482948 |
| 10xmzib | 100019837 | 5946170 | 94073667 | 70512427 | 5936822 | 64575605 |
| CHN | 142339413 | 9961222 | 132378191 | 119773932 | 9872441 | 109901491 |
| S26 | 86692485 | 6645479 | 80047006 | 63937509 | 6594431 | 57343078 |
| T104 | 96212242 | 6146189 | 90066053 | 74343770 | 6060894 | 68282876 |
| T118 | 102631121 | 5851772 | 96779349 | 80263274 | 5768144 | 74495130 |
| T148 | 93117777 | 5465243 | 87652534 | 70092723 | 5510204 | 64582519 |
| T150 | 86308923 | 5496513 | 80812410 | 63912325 | 5515943 | 58396382 |
| T18 | 100666826 | 5918099 | 94748727 | 82704420 | 5779188 | 76925232 |
| T194 | 86096441 | 5340166 | 80756275 | 63437198 | 5336734 | 58100464 |
| T26 | 88052151 | 5761283 | 82290868 | 65370993 | 5727879 | 59643114 |
| T50 | 102323097 | 5886465 | 96436632 | 79189229 | 5822456 | 73366773 |
| T72 | 101444464 | 5403974 | 96040490 | 79133305 | 5354574 | 73778731 |
| T8 | 101837884 | 5980124 | 95857760 | 79208271 | 5831246 | 73377025 |
| T90 | 85484068 | 5872449 | 79611619 | 62786786 | 5938766 | 56848020 |
| T24 | 81822420 | 6482883 | 75339537 | 88133795 | 6325230 | 81808565 |
| T76 | 69756766 | 7024754 | 62732012 | 76533331 | 7211061 | 69322270 |
| T77 | 82307204 | 7410651 | 74896553 | 88194290 | 7116188 | 81078102 |

| | | | | | | |
|---|---|---|---|---|---|---|
| T78 | 71096727 | 7165845 | 63930882 | 74136549 | 6640974 | 67495575 |
| T79 | 69094119 | 6453865 | 62640254 | 71360191 | 6233641 | 65126550 |
| T81 | 84016376 | 7218598 | 76797778 | 89078733 | 6727928 | 82350805 |
| T82 | 68684974 | 6571699 | 62113275 | 75503772 | 6794480 | 68709292 |
| T83 | 82673798 | 7271922 | 75401876 | 87357337 | 6978491 | 80378846 |
| T84 | 89544850 | 6625818 | 82919032 | 77274482 | 6373443 | 70901039 |
| T85 | 71181116 | 6487501 | 64693615 | 69953242 | 6229687 | 63723555 |
| T86 | 89144285 | 7444519 | 81699766 | 87406188 | 7218063 | 80188125 |
| T87 | 85447452 | 6411189 | 79036263 | 75624899 | 6212422 | 69412477 |

**Supplementary Table ST13. PAR coordinates for all male samples for both references.**

| Sample | *M. martes* | | | *M. zibellina* | | |
|---|---|---|---|---|---|---|
| | **start (bp)** | **stop (bp)** | **length (bp)** | **start (bp)** | **stop (bp)** | **length (bp)** |
| T8 | 118000000 | 124450000 | 6450000 | 200000 | 6680000 | 6480000 |
| T18 | 118000000 | 124450000 | 6450000 | 200000 | 6680000 | 6480000 |
| T50 | 118000000 | 124450000 | 6450000 | 200000 | 6680000 | 6480000 |
| T72 | 118000000 | 124450000 | 6450000 | 200000 | 6680000 | 6480000 |
| T104 | 118000000 | 124450000 | 6450000 | 200000 | 6680000 | 6480000 |
| T118 | 118000000 | 124450000 | 6450000 | 200000 | 6680000 | 6480000 |
| S49 | 118000000 | 124450000 | 6450000 | 200000 | 6680000 | 6480000 |
| T149 | 118000000 | 124450000 | 6450000 | 200000 | 6680000 | 6480000 |
| T151 | 118000000 | 124450000 | 6450000 | 200000 | 6680000 | 6480000 |
| T24 | 118000000 | 124450000 | 6450000 | 200000 | 6680000 | 6480000 |
| T77 | 118000000 | 124450000 | 6450000 | 200000 | 6680000 | 6480000 |
| T81 | 118000000 | 124450000 | 6450000 | 200000 | 6680000 | 6480000 |
| T83 | 118000000 | 124450000 | 6450000 | 200000 | 6680000 | 6480000 |
| T84 | 118000000 | 124450000 | 6450000 | 200000 | 6680000 | 6480000 |
| T86 | 118000000 | 124450000 | 6450000 | 200000 | 6680000 | 6480000 |
| T87 | 118000000 | 124450000 | 6450000 | 200000 | 6680000 | 6480000 |

**Supplementary Table ST14. P-values of two- and one-sided Mann-Whitney U tests conducted to compare the number of masked bases between M. zibellina and M. martes genome assemblies relative to classified groups of individuals.**

| Comparison | Statistic | Type * | Masked bases ** | | |
|---|---|---|---|---|---|
| | | | All | >2.5 | <0.33 |
| *M. zibellina - M. zibellina* | Two-sided U-test | p-value<br>p-value (corrected) | 0.00022<br>0.00200 | 0.71961<br>6.47653 | 0.00022<br>0.00200 |
| *M. martes - M. martes* | Two-sided U-test | p-value<br>p-value (corrected) | 0.07739<br>0.69650 | 0.37722<br>3.39502 | 0.07739<br>0.69650 |
| *kidases - kidases* | Two-sided U-test | p-value<br>p-value (corrected) | 0.94764<br>8.52880 | 0.10067<br>0.90601 | 1.000008<br>9.00000 |
| *M. zibellina - M. zibellina* | One-sided U-test | p-value<br>p-value (corrected) | 0.00011<br>0.00100 | 0.35981<br>3.23826 | 0.00011<br>0.00100 |
| *M. martes - M. martes* | One-sided U-test | p-value<br>p-value (corrected) | 0.03869<br>0.34825 | 0.18861<br>1.69751 | 0.03869<br>0.34825 |
| *kidases - kidases* | One-sided U-test | p-value<br>p-value (corrected) | 0.47382<br>4.26440 | 0.05033<br>0.45300 | 0.50000<br>4.50000 |

* P-value (corrected) – Bonferroni correction for 9 tests.

** All – all masked bases; >2.5 – those with coverage 2.5 times greater than the median of the whole genome; <0.33 – those with coverage 0.33 times less than the median of the whole genome.

Tomarovsky AA, Khan R, Dudchenko O, Beklemisheva VR, Perelman PL, Totikov AA, Serdyukova NA, Bulyonkova TM, Weisz D, Yakupova A, et al. in prep. Chromosome-length genome assemblies and macrosynteny of three marten species.
