## Supplementary Methods for "Genomics of sable (*Martes zibellina)* × pine marten (*Martes martes*) hybridization"

* corresponding author

[x] equal contribution

**Supplementary method SM1**. Localization and genotyping of previously known STR loci.

Using an *in silico* PCR approach, we identified the locations of STR loci previously described for various Mustelidae species (Davis and Strobeck 1998; Fleming et al. 1999; Domingo-Roura 2002; Vincent et al. 2003; Basto et al. 2010; Natali et al. 2010). Based on the results, the markers were divided into three categories: Localized (L), Declined (D) and Not Amplified (NA). Using sable reference assembly, we localized 44 STRs (Supplementary File SF14). An additional 16 markers had low-confidence mapping and were declined. The identification rate using the pine marten reference assembly was nearly the same: 48 high confidence mappings and 15 ambiguous. None of the remaining loci were amplified. All three categories have significant overlaps between the two reference assemblies: 87.7 % (43/49) for localized STRs, 72.2 % (13/18) for declined STRs, and 75 % (15/20) for not amplified STRs, respectively. The localized markers cover all chromosomes of both species, except chr16 (Supplementary Methods Figure SMF 1). The mean density of markers is 2.3 and 2.5 loci per chromosome for sable and pine marten, respectively.

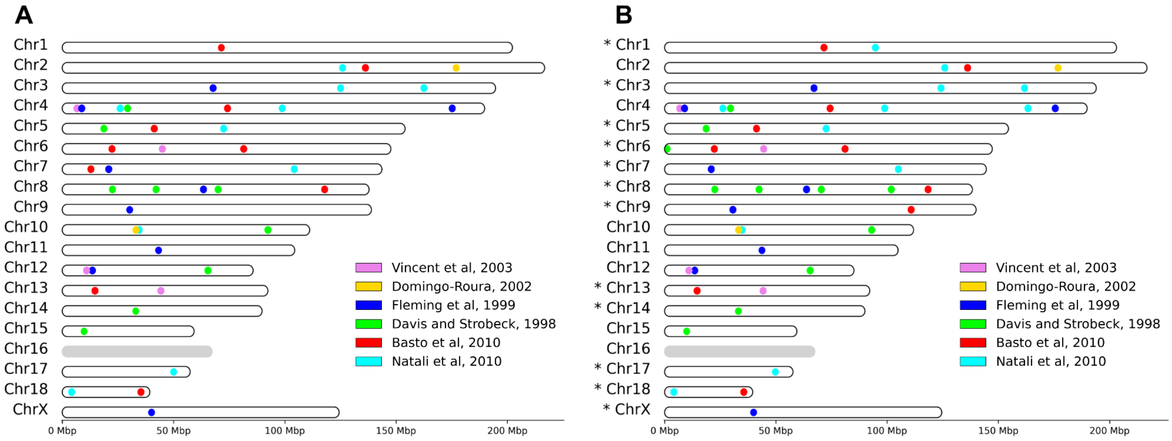


**Supplementary Methods Figure SMF 1.** Localization of previously identified STR loci from various species of Mustelidae.

A – Distribution of STR loci in the sable reference assembly, B – Distribution of STR loci in the pine marten reference assembly. Pine marten chromosomes labeled with an asterisk were inverted to match the orientation of the sable homologous chromosomes

Further investigation of repeat motifs in the two reference assemblies (described in methods Section "Whole genome alignments and connection between assembly and karyotype") reduced the number of the suitable STR loci for genotyping to 36 and 44 in the sable and pine marten assemblies, respectively (Supplementary File SF4). The reasons for discarding these loci were different. For example, Mf8.8, Mf4.10, Mf6.5 were too long for genotyping from 150 bp reads. The STR itself in each of these markers was longer than 100 bp in the reference sable genome. Moreover, the repeat in Mf8.8 was not completely assembled and contained a gap. Mer095 contained not one but two complementary STRs separated by a 26 bp insertion and also exceeded the 100 bp threshold. Mvis099 did not contain STR at all, and the sequences of repeats within Mel08 and Mvi_1273 were very different from the way they were originally described. Finally, Mvis020 was removed because it was X-linked.

Rozhnov et al. in their study of hybridization between the sable and the pine marten used only 9 STR markers: Mel10, Ma-1, Ma-3, Ma-8, Ma-15, Ma-18, Ma-19, Mvis072 and Mer041 (Rozhnov et al. 2013). Seven of them passed our filters and were genotyped in our resequencing data (except Ma-3 and Mer041). Kashtanov et al. investigated population structure of the sable in Central Siberia using 8 STR markers from the Rozhnov et al. set (Mel10 was excluded) and 8 additional STR loci (Mf3.7,

Mar08, Mar21, Mar36, Mar43, Mar53, Mar58 and Mar64) (Kashtanov et al. 2022). Among the additional STR markers, only one (Mar53) was not genotyped using the *M. zibellina* genome assembly in our analysis. To compare our WGS-based ADMIXTURE results with these studies, we performed STRUCTURE analysis using three sets of STR markers: (1) full set, including all loci, (2) Rozhnov's set and (3) Kashtanov's set, excluding non genotyped loci.

**Supplementary method SM2**. Components of heterozygosity distributions

To analyze the components of heterozygosity distributions, we fitted our data with a linear combination of negative binomial distributions. For each sample we tried models with different numbers of components or different starting values of the parameters. For each sample we selected one of six models (SMT 1). Comparing the distributions of our samples, we initially empirically identified 4 components that corresponded to modes of the groups of studied individuals: ROH component (0 SNPs/kbp), pine marten component (P) (0.5 SNPs/kbp), sable component (S) (1.5 SNPs/kbp) and hybrid component (H) (4.5 SNPs/kbp). Thus, starting values for mean (μ) and standard deviation (σ) of each component (SMT 2) were empirically determined based on heterozygosity distributions of the studied samples.

As input data we used counts of the heterozygous SNPs in sliding windows of 1 Mb with a 100 kbp step size. Counts from X chromosomes were removed prior to the analysis for all samples. Distributions of counts were constructed with a bin width of 100 counts (0.1 SNPs/kbp). The first two bins (corresponding to ROH) were removed to improve model fitting. Model fitting for heterozygosity distributions was performed using the mix() function with 'dist = "nbinom"' from the R package mixdist v.0.5-5 (Macdonald 2018).

**Supplementary Methods Table SMT 1.** Fitted models and their components.

| Model | Formula * | Components | | |
|---|---|---|---|---|
| | | **Pine marten** | **Sable** | **Hybrid** |
| General model | $a_p * Nb_p + a_s * Nb_s + a_h * Nb_h$ | + | + | + |
| Model without hybrid heterozygosity | $a_p * Nb_p + a_s * Nb_s$ | + | + | – |
| Pure pine model | $a_p * Nb_p$ | + | – | – |
| Pure sable model | $a_s * Nb_s$ | – | + | – |
| Pine + hybrid model | $a_r * Nb_r + a_p * Nb_p + a_h * Nb_h$ | + | – | + |
| Sable + hybrid model | $a_r * Nb_r + a_s * Nb_s + a_h * Nb_h$ | – | + | + |

* Formula coefficients:

- a – weight of an individual component (coefficient in linear combination).
- Nb – negative binomial distribution with mean value (μ) and standard deviation (σ).
- $_{p\ s\ h}$ – subscripts of individual components (pine marten, sable and hybrid).

**Supplementary Methods Table SMT 2.** Starting values for mean (μ) and standard deviation (σ) of each component.

| Component | μ | σ |
|---|---|---|
| Pine marten (P) | 700 | 200 |
| Sable (S) | 1700 | 500 |
| Hybrid (H) | 4500 | 700 |

We selected the best model for each sample in two steps. First, we filtered models based on the final mean (μ) of the fitted model; if any component's μ fell outside the range of the starting μ ± standard deviation (σ), we discarded the model. Next, we selected the best model based on the smallest quadratic deviation, calculated as the sum of squared differences between the original and fitted distributions, excluding the first bin.

After selecting the best model, we identified means and modes for each fitted distribution (Supplementary File SF8). We clustered all mean values of individual fitted distributions from the best-fit models using hierarchical agglomerative clustering. We used the *linkage()* function from Scipy v1.13.1 with the "average" method. We visualized the results of the clustering as a dendrogram.

We formed flat clusters using the *fcluster* function from Scipy, setting the optimal number of clusters to 3 (sable, pine marten, and hybrid clusters). We clustered the mean values of individual component distributions for each of the three groups using the maxclust criterion and calculated medians for each cluster. We performed all stages of the analysis based on both genome assemblies of *M. zibellina* and *M. martes*. Specifically, we calculated three median values for each of the three clusters relative to the *M. zibellina* reference, and similarly for the *M. martes* reference. Finally, we obtained and visualized the mean values of these medians for each cluster on the stripped histograms (Main text, Figure 7A).

**Supplementary method SM3.** Algorithm for identification of a pseudoautosomal region.

To set the correct ploidy of X chromosome segments during variant calling we identified coordinates of a pseudoautosomal region (PAR). First, median coverage was estimated in non-overlapping sliding windows of 10 kbp. Chains of at least 10 consecutive windows with median coverage (of each window) ≥70% of the whole genome value were merged. Next, a median coverage was calculated for windows between adjacent chains. If it exceeded 70% of the whole genome level adjacent chains were merged too to generate a set of candidate regions. Finally, we compared the candidates with per-base coverage and selected the correct one. The scheme of the procedure is provided on Figure SMF 2.

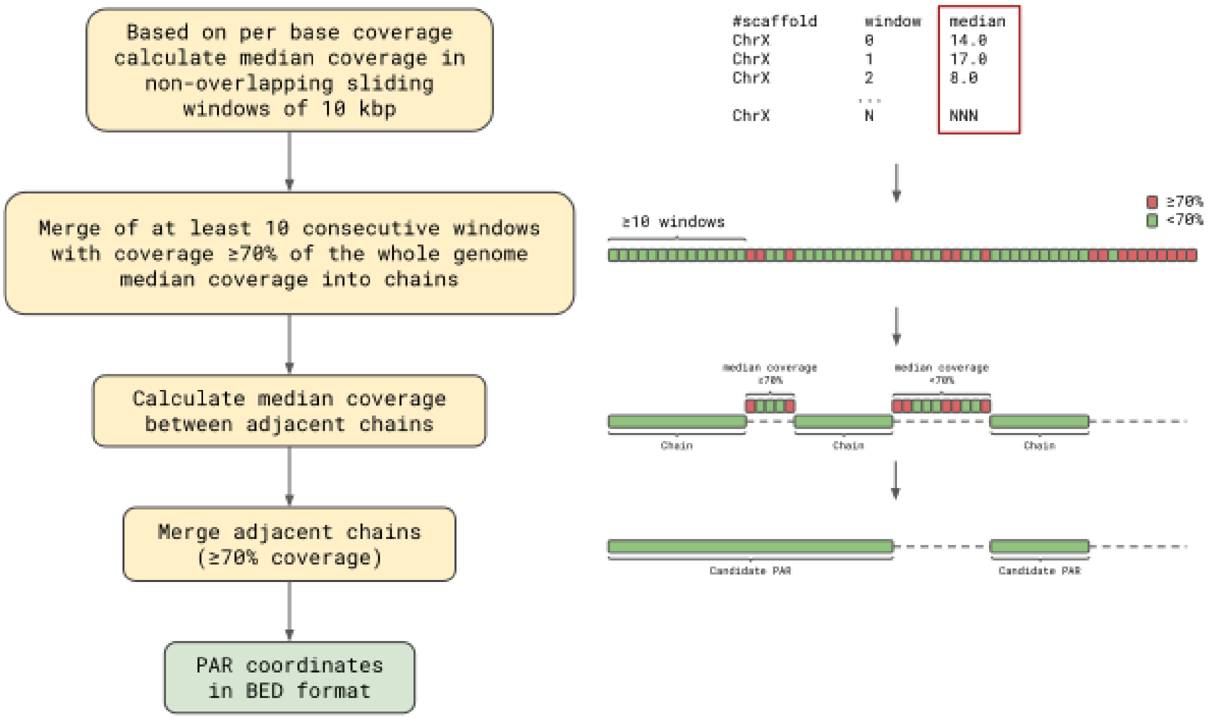


**Supplementary Methods Figure SMF 2**. Scheme of algorithm for PAR identification.

**Supplementary method SM4**. snakeSTR pipeline.

The snakeSTR pipeline (https://github.com/mahajrod/snakeSTR), built on Snakemake, enables STR-typing and STR-based admixture analysis (Figure SMF 3). Pipeline description:

**I - V. Preprocessing**. On the first stage for each sample and STR loci we extracted reads aligned to it, including 1000 bp flanks using Samtools v1.19.2 (Li et al. 2009) (I, II), verified pairing using Bazam v1.0.1 (Sadedin and Oshlack 2019) (III), mapped them back to the reference genome using BWA v0.7.17 (Li and Durbin 2009) (IV) and performed an indel-aware realignment using IndelRealigner from GATK v3.7 (McKenna et al. 2010) (V).

**VI - VII. STR-typing**. Next, we performed STR-typing using hipSTR v0.6.2 (Willems et al. 2017) (VI-VII). The following workflow includes both analysis of raw and filtered STR-loci sets. Analysis of raw STR-loci sets was performed only with a minimum read depth threshold of 5 (--min-reads). Analysis of filtered STR-loci sets was performed with additional filtering (VI a) according to the following criteria: also with a minimum read depth threshold of 5, a minimum call quality threshold of 0.9 (--min_call_qual), a maximum allowable fraction of reads with flank indels set to 0.15

(--max_call_flank_indel), a maximum allowable fraction of reads with stutter artifacts set to 0.15 (--max_call_stutter), a minimum call allele bias threshold of -2 (--min_call_allele_bias), and a minimum call strand bias threshold of -2 (--min_call_strand_bias). From this point, all downstream stages were performed for both raw and filtered STR-loci sets to assess the effect of the filtration on the final results.

**VIII - XIV. Admixture analysis**. STR-based admixture analysis performed using STRUCTURE v2.3.4 and Clumpp v1.1.2 (Pritchard et al. 2000; Jakobsson and Rosenberg 2007) (VIII-XIV). Admixture analysis was performed for values of parameter K (number of populations) from 2 to 6 in triplicate for each. The presented DAG includes only K = 2 for clarity. Regarding Clumpp, the following parameters were employed: M = 1 (indicating the FullSearch method), W = 0 (denoting no weighting by the number of individuals in each population), and S = 2 (representing the use of the pairwise matrix similarity statistic G'). Additionally, the Greedy algorithm was employed with the option set to 2, indicating random input orders, and 100 repeats were performed to ensure robustness of the results.

**XV - XVI. Visualization**. Creation input files needed to visualize the clustering results using Pong v1.5 (Behr et al. 2016) (XV). Generation a Bash script to automatically run an interactive visualization using Pong with default parameters (XVI).

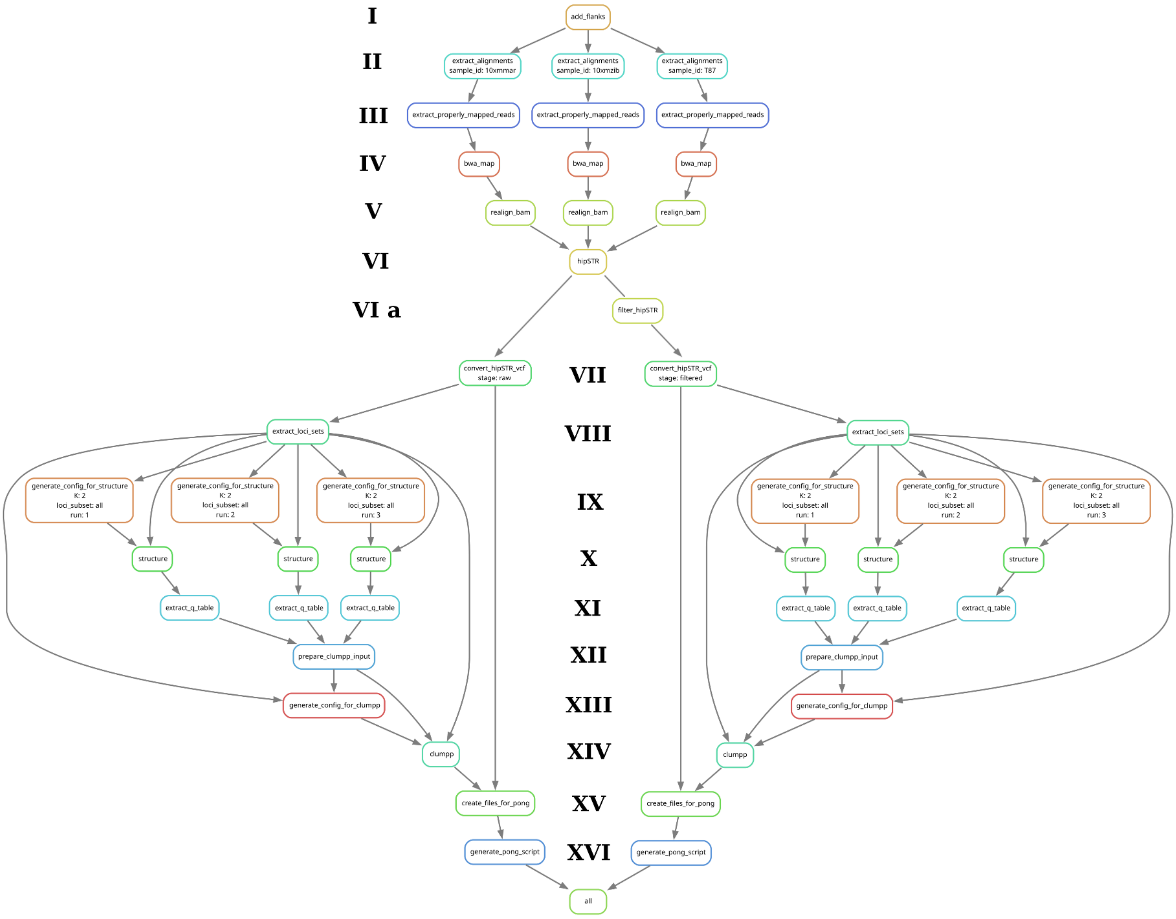


**Supplementary methods figure SMF 3.** Rule graph of the snakeSTR pipeline.

Roman numerals indicate the steps of workflow.

**Supplementary method SM5**. Fst and Tajima's D

Fst and Tajima's D statistics were calculated using Vcftools v.0.1.16 and VCF-kit v.0.2.9 (Danecek et al. 2011; Cook and Andersen 2017), respectively, in 1 Mbp sliding windows with a 100 kbp step, using the sable genome assembly as a reference. The comparison was performed between groups of pure sables and pure pine martens (based on local Admixture classification). Genomic windows with Fst and Tajima's D estimates were filtered in two stages. First, windows containing more than 50,000 Ns were discarded. Next, filtering was performed based on repeat content, which was identified using Tandem Repeats Finder v4.09.1 (Benson 1999), WindowMasker 1.0.0 (Morgulis et al. 2006), and RepeatMasker v.4.1.2.p1 (Smit et al. 2013). Conversion of GFF with repeats to BED format was performed using gff2bed from BEDOPS v.2.4.40 (Neph et al. 2012). Obtaining masked bases by windows was done by merging and then intersecting using Bcftools v1.15.1 (Danecek et al. 2021). For each window, we calculated the number of masked bases. The distributions of repeats were visualized as histograms for the whole assembly (Figure SMF 4) and for each scaffold (Figure SMF

5), as well as in the form of a heatmap (Figure SMF 6). Based on the visualization of these distributions, we applied a cutoff at the 95th percentile, meaning that windows containing more than 607,592 bp of repeats were removed. Finally, the coordinates of regions consisting of overlapping windows are obtained using Bedtools v.2.31.1 (Quinlan and Hall 2010).

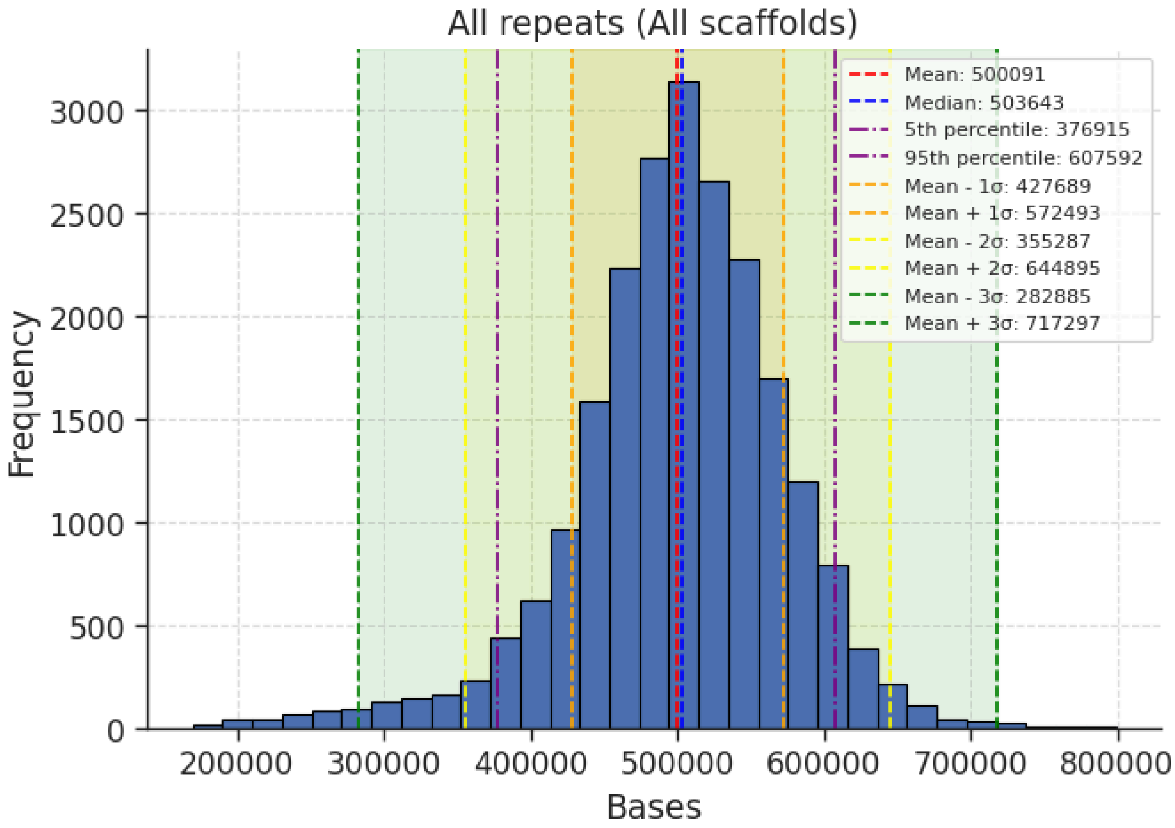


**Supplementary Methods Figure SMF 4.** Distribution of repeats across 1 Mbp sliding windows in all scaffolds of the *M. zibellina* genome assembly.

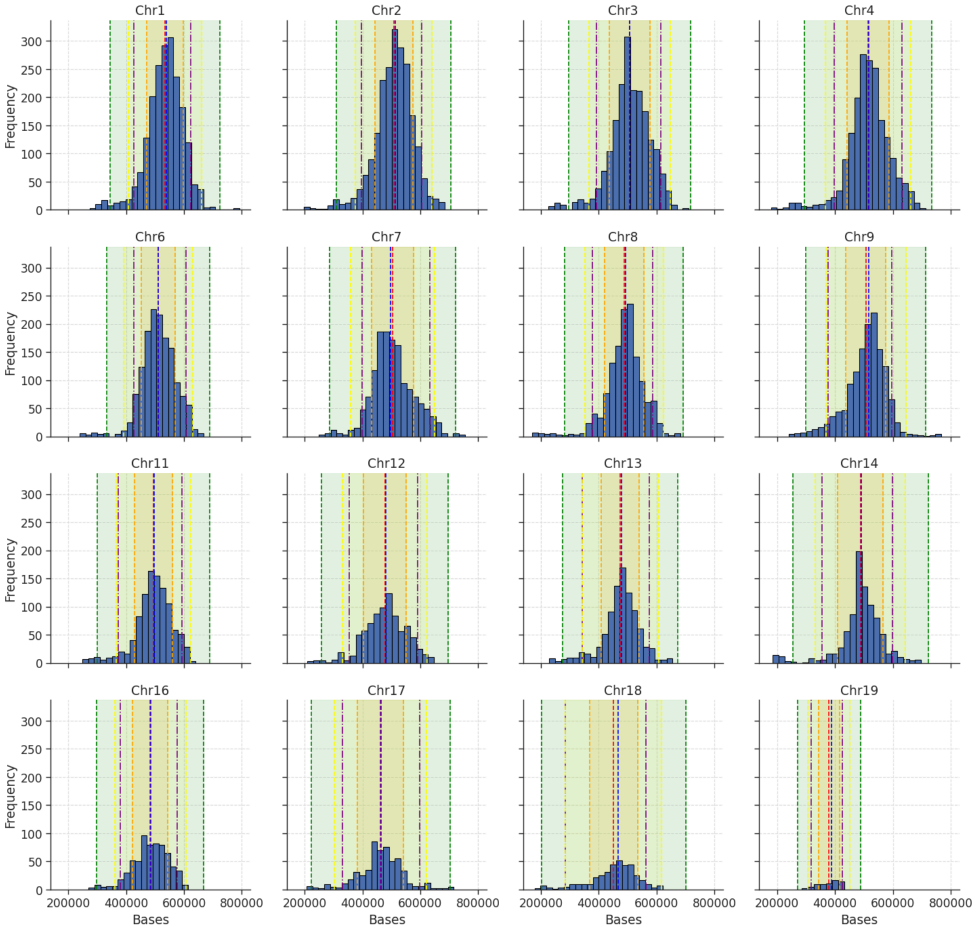


**Supplementary methods figure SMF5.** Distribution of repeats across 1 Mbp sliding windows for each scaffold in the *M. zibellina* genome assembly.

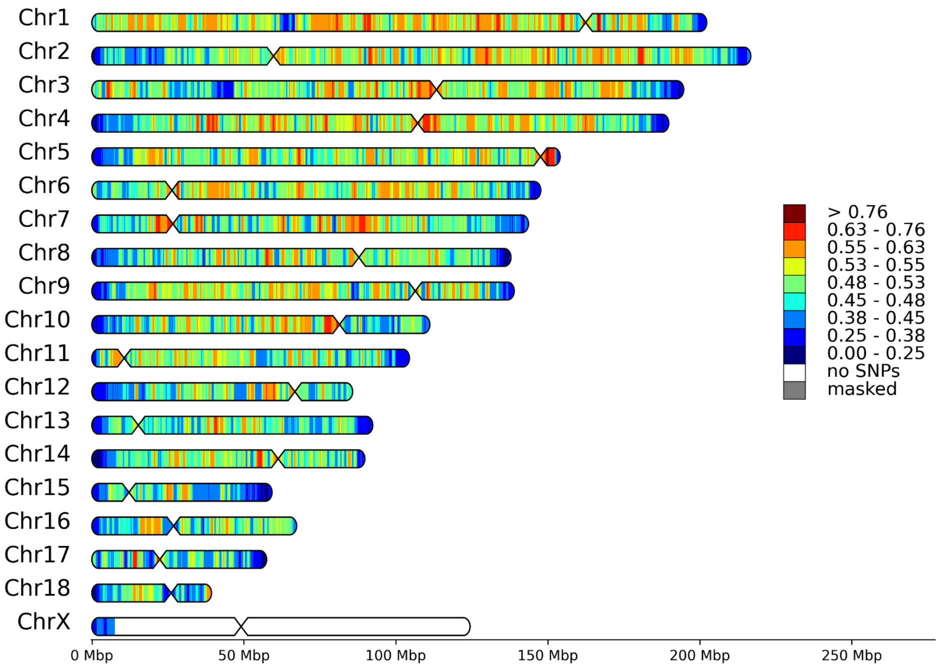


**Supplementary methods figure SMF6.** Density of the all repeats in the sliding windows of 1 Mbp with a step of 100 kbp (Reference: *M. zibellina*)

Region. *J. Mammal.* 83:907–912.

Fleming, Ostrander, Cook. 1999. Microsatellite markers for American mink (Mustela vison) and ermine (Mustela erminea). *Mol. Ecol.*:1352–1354.

Jakobsson M, Rosenberg NA. 2007. CLUMPP: a cluster matching and permutation program for dealing with label switching and multimodality in analysis of population structure. *Bioinformatics* 23:1801–1806.

Kashtanov S, Zakharov E, Begletsov O, Svishcheva G, Rychkov SY, Filimonov P, Onokhov A, Levenkova E, Meschersky I, Rozhnov V. 2022. Expansion of the sable (Martes zibellina L.) from the north of the Central Siberian Plateau into tundra ecosystems. *Russ. J. Genet.* 58:955–966.

Li H, Durbin R. 2009. Fast and accurate short read alignment with Burrows–Wheeler transform. *Bioinformatics* 25:1754–1760.

Li H, Handsaker B, Wysoker A, Fennell T, Ruan J, Homer N, Marth G, Abecasis G, Durbin R. 2009. The Sequence Alignment/Map format and SAMtools. *Bioinformatics* 25:2078–2079.

Macdonald P. 2018. mixdist: Finite Mixture Distribution Models. Available from: https://CRAN.R-project.org/package=mixdist

McKenna A, Hanna M, Banks E, Sivachenko A, Cibulskis K, Kernytsky A, Garimella K, Altshuler D, Gabriel S, Daly M, et al. 2010. The Genome Analysis Toolkit: A MapReduce framework for analyzing next-generation DNA sequencing data. *Genome Res.* 20:1297–1303.

Morgulis A, Gertz EM, Schäffer AA, Agarwala R. 2006. WindowMasker: window-based masker for sequenced genomes. *Bioinforma. Oxf. Engl.* 22:134–141.

Natali C, Banchi E, Ciofi C, Manzo E, Bartolommei P, Cozzolino R. 2010. Characterization of 13 polymorphic microsatellite loci in the European pine marten Martes martes. *Conserv. Genet. Resour.* 2:397–399.

Neph S, Kuehn MS, Reynolds AP, Haugen E, Thurman RE, Johnson AK, Rynes E, Maurano MT, Vierstra J, Thomas S, et al. 2012. BEDOPS: high-performance genomic feature operations. *Bioinformatics* 28:1919–1920.

Pritchard JK, Stephens M, Donnelly P. 2000. Inference of Population Structure Using Multilocus Genotype Data. *Genetics* 155:945–959.

Quinlan AR, Hall IM. 2010. BEDTools: a flexible suite of utilities for comparing genomic features. *Bioinformatics* 26:841–842.

Rozhnov VV, Pishchulina SL, Meschersky IG, Simakin LV. 2013. On the ratio of phenotype and genotype of sable and pine marten in sympatry zone in the Northern Urals. *Mosc. Univ. Biol. Sci. Bull.* 68:178–181.

Sadedin SP, Oshlack A. 2019. Bazam: a rapid method for read extraction and realignment of high-throughput sequencing data. *Genome Biol.* 20:78.

Smit A, Hubley R, Green P. 2013. 2015 RepeatMasker Open-4.0.

Vincent IR, Farid A, Otieno CJ. 2003. Variability of thirteen microsatellite markers in American mink ( *Mustela vison* ). *Can. J. Anim. Sci.* 83:597–599.

Willems T, Zielinski D, Yuan J, Gordon A, Gymrek M, Erlich Y. 2017. Genome-wide profiling of heritable and de novo STR variations. *Nat. Methods* 14:590–592.
