## Supplementary File SF2 for "Genomics of sable (*Martes zibellina)* × pine marten (*Martes martes*) hybridization"

* corresponding author

[x] equal contribution

**Figures:**

**Figure 1.** PCA plot based on 6,697,683 SNPs obtained from autosomes and the pseudoautosomal region (PAR) of the *M. martes* assembly.

The first and second principal components are represented on the X and Y axes, respectively. The percentage of variance explained on each axis is indicated in parentheses. Top: plot with sample labels. Bottom: plot without sample labels.

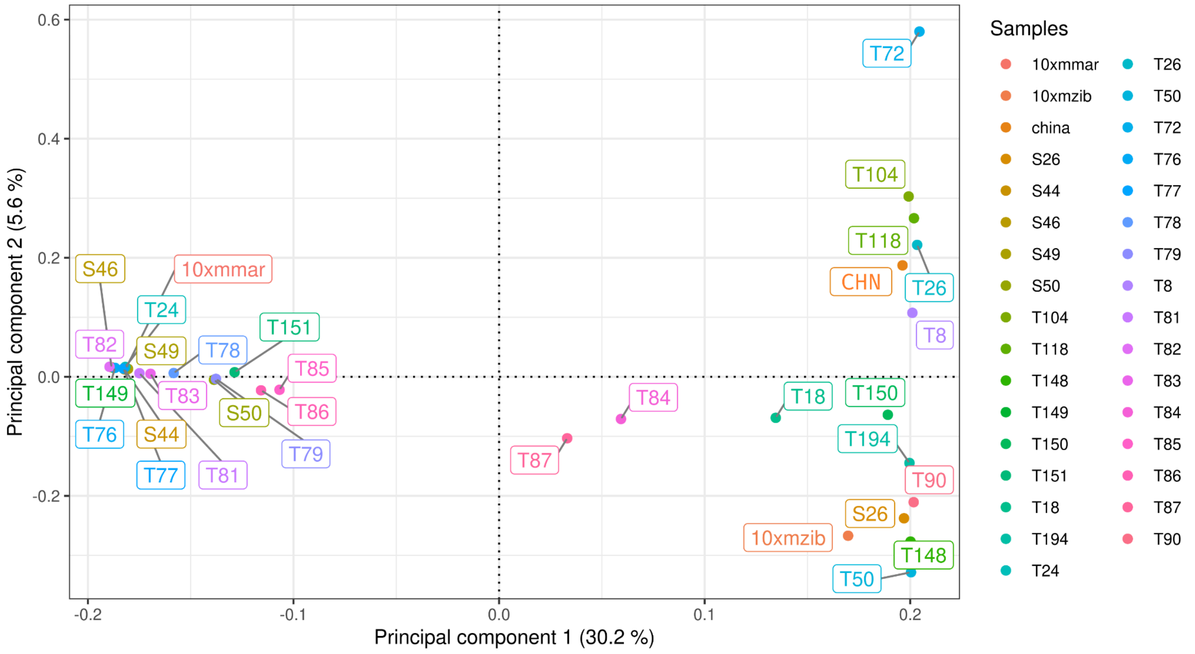


**Figure 2.** Global ADMIXTURE (left half) and local ADMIXTURE (right half) of 1 Mb sliding windows with 100 kbp step.

Samples are labeled along the X-axis while the Y-axis is the mean probability of ancestry.

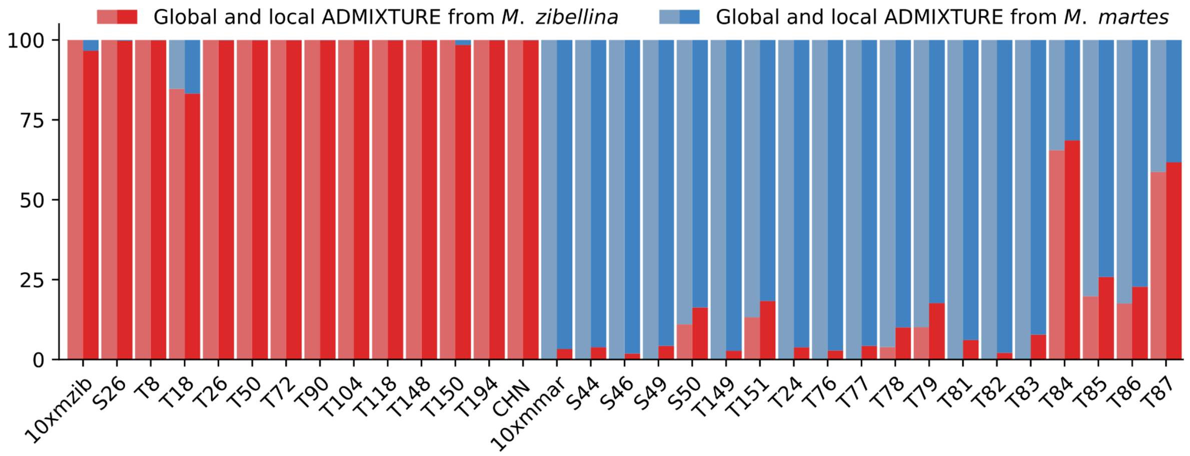

**Figure 3.** Fitting models for heterozygosity distributions.

Red triangles indicate mean values for the distribution of individual model components.

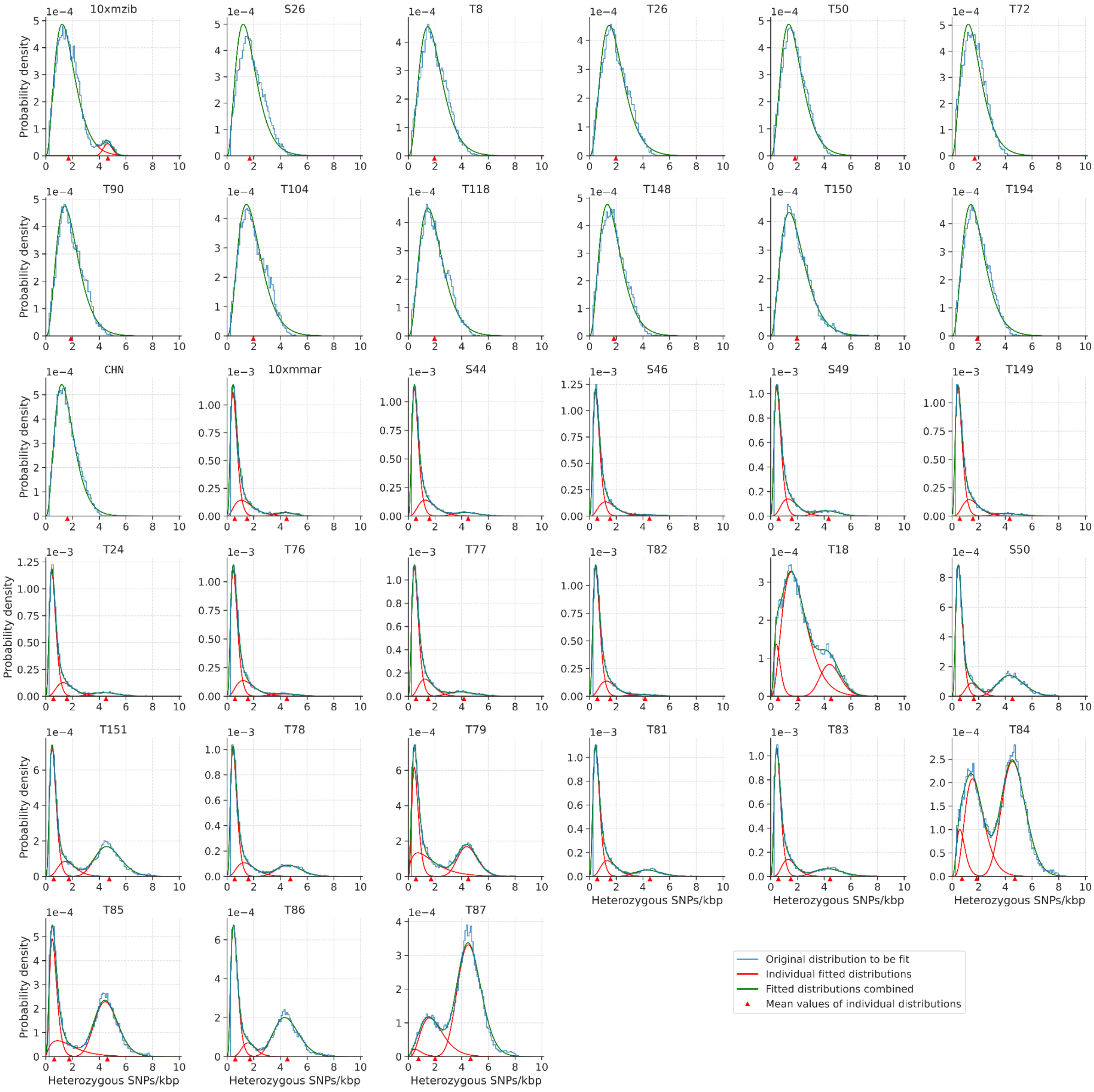

**Figure 4.** Localization of markers and admixture analysis for three sets of STR loci (All, Rozhnov's, Kashatonov's).

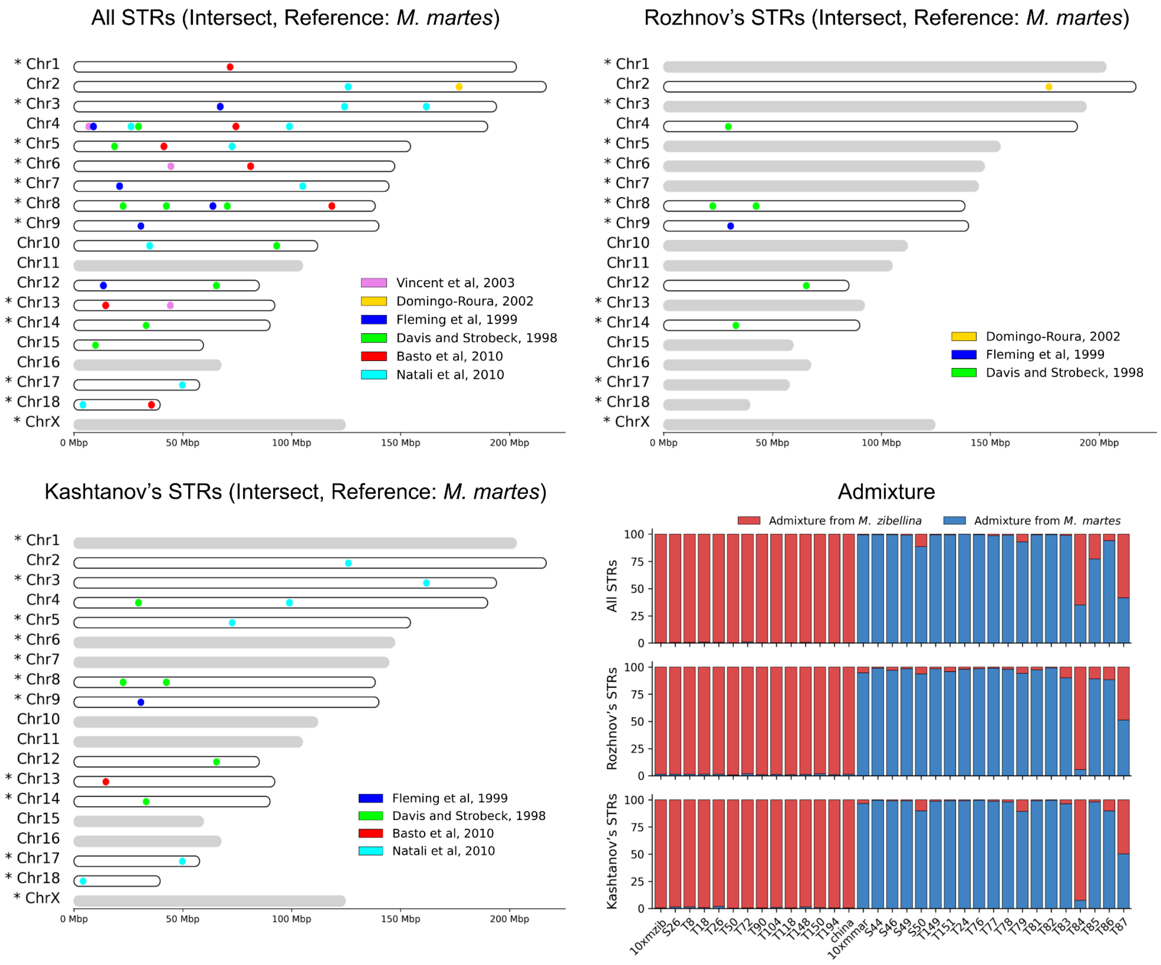

**Figure 5.** Runs of homozygosity.

A-C – cumulative distribution of RoH length in the pure martens, pure sables and hybrids; D – cumulative lengths (fraction of the genome) of the RoH of different categories: ultra long (L ≥ 10 Mbp), long (10 Mbp > L ≥ 1 Mbp) and short (L < 1 Mbp), in all samples. The X chromosome was excluded from all samples.

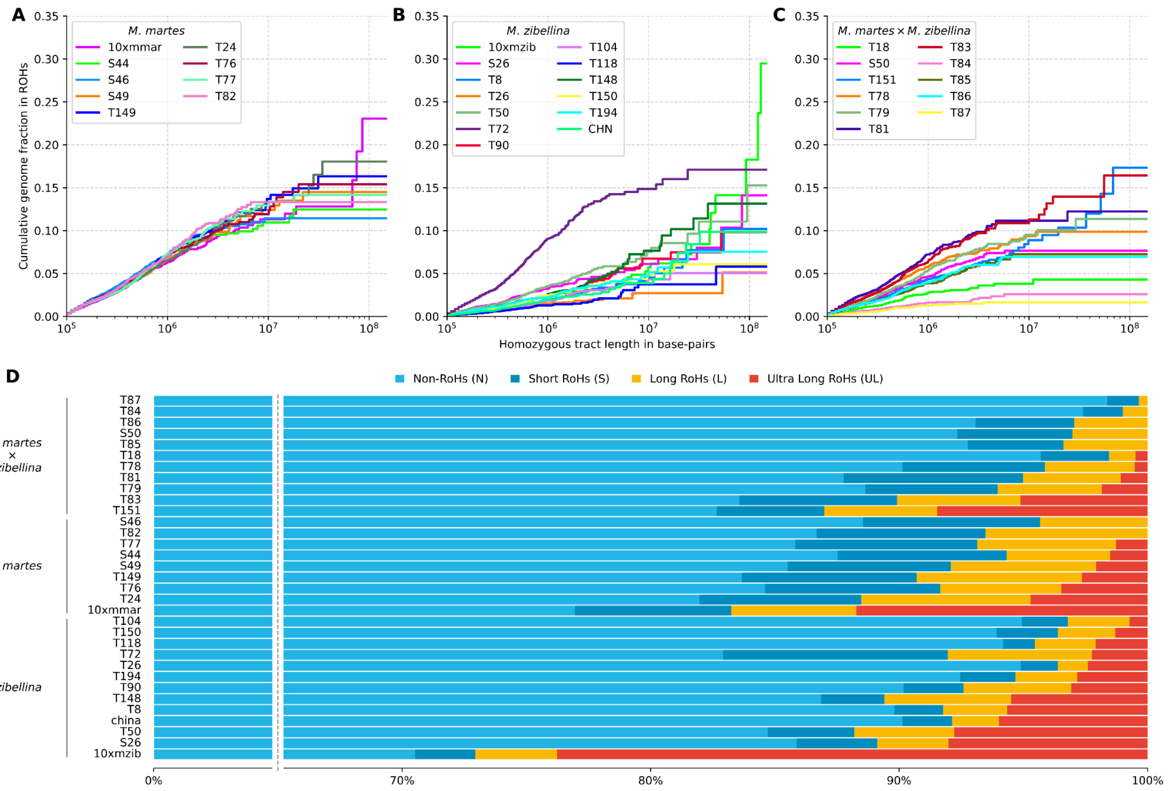

**Figure 6.** Demographic history reconstruction.

A – "pure" sables, B – "pure" martens, C – hybrids, D – all samples. Mutation rate ($\mu$) = $4.64 \times 10^{-9}$. Generation time (g) = 5 for all plots. The X chromosome was excluded from the analysis.

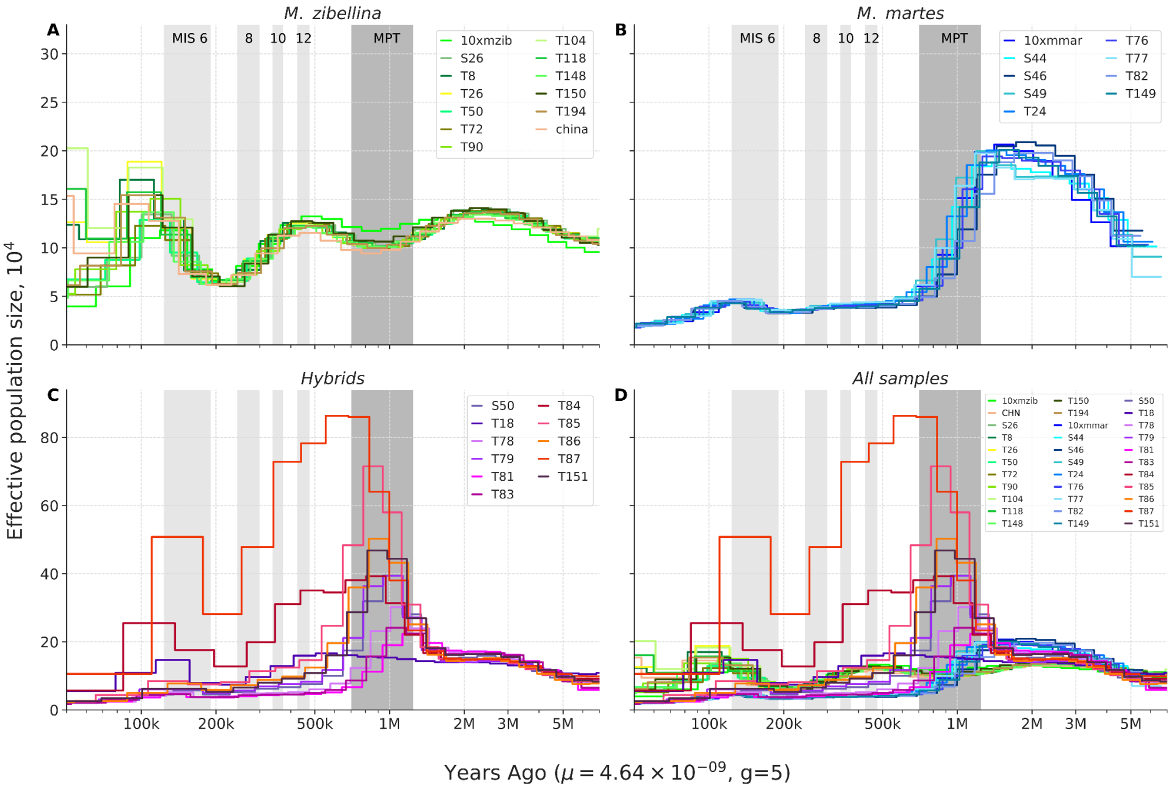

**Figure 7**. Median coverage in non-overlapping 10 kbp windows for males across the pseudoautosomal region in *M. martes* genome assembly.

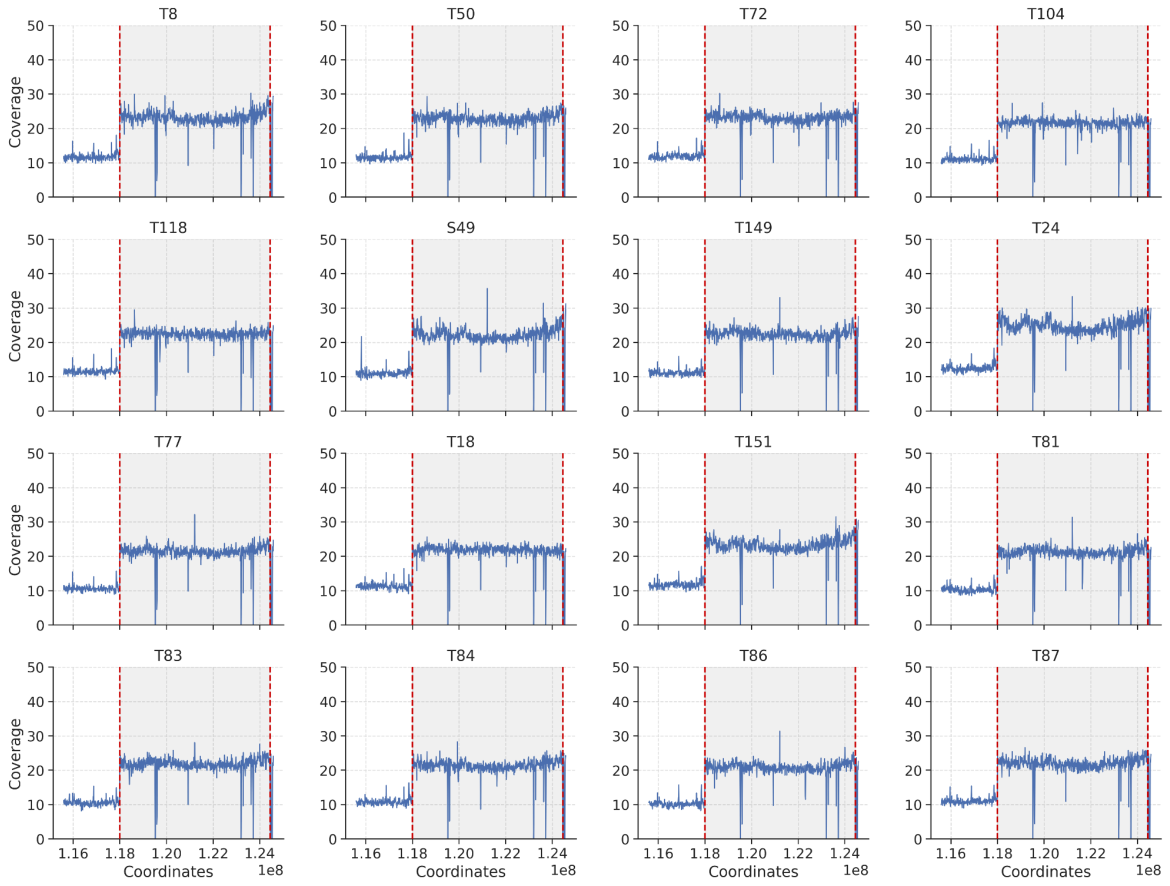


**Tables:**

**Table 1.** Global (whole genome) and local (in sliding 1 Mbp windows with 100 kbp step) ADMIXTURE (k = 2).

| Sample | Global ADMIXTURE | | Local ADMIXTURE | |
|---|---|---|---|---|
| | M. zibellina, % | *M. martes*, % | M. zibellina, % | *M. martes*, % |
| 10xmzib | 100 | 0 | 96.62 | 3.38 |
| S26 | 100 | 0 | 99.73 | 0.27 |
| T8 | 100 | 0 | 99.89 | 0.11 |
| T18 | 84.7 | 15.3 | 83.22 | 16.78 |
| T26 | 100 | 0 | 99.88 | 0.12 |

| Sample | Global ADMIXTURE | | Local ADMIXTURE | |
|---|---|---|---|---|
| | **M. zibellina, %** | ***M. martes*, %** | **M. zibellina, %** | ***M. martes*, %** |
| T50 | 100 | 0 | 99.89 | 0.11 |
| T72 | 100 | 0 | 99.91 | 0.09 |
| T90 | 100 | 0 | 99.90 | 0.10 |
| T104 | 100 | 0 | 99.86 | 0.14 |
| T118 | 100 | 0 | 99.88 | 0.12 |
| T148 | 100 | 0 | 99.88 | 0.12 |
| T150 | 100 | 0 | 98.40 | 1.60 |
| T194 | 100 | 0 | 99.85 | 0.15 |
| CHN | 100 | 0 | 99.87 | 0.13 |
| 10xmmar | 0 | 100 | 3.29 | 96.71 |
| S44 | 0 | 100 | 3.82 | 96.18 |
| S46 | 0 | 100 | 1.86 | 98.14 |
| S49 | 0 | 100 | 4.26 | 95.74 |
| S50 | 11 | 89 | 16.28 | 83.72 |
| T149 | 0 | 100 | 2.71 | 97.29 |
| T151 | 13.2 | 86.8 | 18.33 | 81.67 |
| T24 | 0 | 100 | 3.81 | 96.19 |
| T76 | 0 | 100 | 2.79 | 97.21 |
| T77 | 0 | 100 | 4.25 | 95.75 |
| T78 | 3.9 | 96.1 | 10.04 | 89.96 |
| T79 | 10.1 | 89.9 | 17.64 | 82.36 |
| T81 | 0 | 100 | 6.07 | 93.93 |
| T82 | 0 | 100 | 2.10 | 97.90 |
| T83 | 0 | 100 | 7.78 | 92.22 |
| T84 | 65.5 | 34.5 | 68.57 | 31.43 |
| T85 | 19.8 | 80.2 | 25.83 | 74.17 |

| Sample | Global ADMIXTURE | | Local ADMIXTURE | |
|---|---|---|---|---|
| | M. zibellina, % | *M. martes*, % | M. zibellina, % | *M. martes*, % |
| T86 | 17.5 | 82.5 | 22.78 | 77.22 |
| T87 | 58.7 | 41.3 | 61.68 | 38.32 |

**Table 2**. Total number of heterozygous SNPs.

Mean and median heterozygosity (SNPs/kbp) calculated in 1 Mbp windows with 100 kbp step. The X chromosome was excluded for all samples.

| Sample | Number of hetSNPs, mln | Mean heterozygosity | Median heterozygosity |
|---|---|---|---|
| 10xmzib | 3.24 | 1.37 | 1.28 |
| S26 | 3.99 | 1.7 | 1.65 |
| T8 | 4.35 | 1.85 | 1.8 |
| T26 | 4.58 | 1.95 | 1.85 |
| T50 | 3.91 | 1.67 | 1.63 |
| T72 | 3.77 | 1.6 | 1.58 |
| T90 | 4.21 | 1.79 | 1.71 |
| T104 | 4.60 | 1.96 | 1.85 |
| T118 | 4.55 | 1.94 | 1.83 |
| T148 | 3.99 | 1.7 | 1.66 |
| T150 | 4.57 | 1.95 | 1.78 |
| T194 | 4.36 | 1.85 | 1.78 |
| CHN | 3.72 | 1.54 | 1.46 |
| 10xmmar | 1.93 | 0.86 | 0.59 |
| S44 | 2.37 | 1.05 | 0.68 |
| S46 | 2.09 | 0.92 | 0.66 |
| S49 | 2.46 | 1.09 | 0.68 |
| T149 | 2.13 | 0.94 | 0.65 |
| T24 | 2.29 | 1.01 | 0.63 |

| Sample | Number of hetSNPs, mln | Mean heterozygosity | Median heterozygosity |
|---|---|---|---|
| T76 | 2.19 | 0.97 | 0.67 |
| T77 | 2.40 | 1.06 | 0.68 |
| T82 | 2.03 | 0.9 | 0.66 |
| T18 | 5.41 | 2.34 | 2.03 |
| S50 | 4.57 | 2.01 | 0.93 |
| T151 | 4.88 | 2.14 | 0.97 |
| T78 | 3.54 | 1.57 | 0.75 |
| T79 | 4.62 | 2.04 | 1.05 |
| T81 | 2.76 | 1.22 | 0.7 |
| T83 | 2.88 | 1.27 | 0.69 |
| T84 | 7.67 | 3.36 | 3.65 |
| T85 | 6.21 | 2.73 | 2.86 |
| T86 | 5.63 | 2.48 | 1.6 |
| T87 | 9.04 | 3.97 | 4.29 |

* heterozygous SNPs

**Table 3**. RoHs content.

| Sample | Number of RoH | Length, Mbp | RoH, % | Non-RoH, % | % of RoH * | | |
|---|---|---|---|---|---|---|---|
| | | | | | Short (S) | Long (L) | Ultra Long (UL) |
| 10xmmar | 603 | 515.8 | 23.1 | 76.9 | 6.3 | 5 | 11.7 |
| S44 | 668 | 279.1 | 12.5 | 87.5 | 6.8 | 4.2 | 1.5 |
| S46 | 700 | 256.3 | 11.5 | 88.5 | 7.1 | 4.3 | 0 |
| S49 | 634 | 324.3 | 14.5 | 85.5 | 6.6 | 5.8 | 2.1 |
| T149 | 673 | 365.5 | 16.4 | 83.6 | 7.1 | 6.7 | 2.7 |
| T24 | 641 | 403.8 | 18.1 | 81.9 | 6.5 | 6.8 | 4.7 |
| T76 | 670 | 344.5 | 15.4 | 84.6 | 7.1 | 4.9 | 3.5 |

| Sample | Number of RoH | Length, Mbp | RoH, % | Non-RoH, % | % of RoH * | | |
|---|---|---|---|---|---|---|---|
| | | | | | Short (S) | Long (L) | Ultra Long (UL) |
| T77 | 651 | 317.3 | 14.2 | 85.8 | 7.3 | 5.6 | 1.3 |
| T82 | 684 | 298.0 | 13.3 | 86.7 | 6.8 | 6.5 | 0 |
| 10xmzib | 255 | 659.7 | 29.5 | 70.5 | 2.4 | 3.3 | 23.8 |
| S26 | 329 | 316.1 | 14.2 | 85.8 | 3.3 | 2.9 | 8 |
| T8 | 192 | 228.2 | 10.2 | 89.8 | 2 | 2.6 | 5.7 |
| T26 | 142 | 114.4 | 5.1 | 94.9 | 1.5 | 1.2 | 2.4 |
| T50 | 335 | 342.0 | 15.3 | 84.7 | 3.5 | 4 | 7.8 |
| T72 | 776 | 382.5 | 17.1 | 82.9 | 9.1 | 5.8 | 2.2 |
| T90 | 253 | 219.6 | 9.8 | 90.2 | 2.4 | 4.3 | 3.1 |
| T104 | 169 | 113.1 | 5.1 | 94.9 | 1.8 | 2.5 | 0.7 |
| T118 | 130 | 130.2 | 5.8 | 94.2 | 1.3 | 2.5 | 2.1 |
| T148 | 266 | 294.2 | 13.2 | 86.8 | 2.6 | 5.1 | 5.5 |
| T150 | 233 | 136.0 | 6.1 | 93.9 | 2.5 | 2.3 | 1.3 |
| T194 | 247 | 168.9 | 7.6 | 92.4 | 2.2 | 2.5 | 2.8 |
| CHN | 187 | 220.7 | 9.9 | 90.1 | 2 | 1.9 | 6 |
| T18 | 257 | 96.1 | 4.3 | 95.7 | 2.7 | 1.1 | 0.5 |
| S50 | 452 | 171.5 | 7.7 | 92.3 | 4.6 | 3 | 0 |
| T151 | 418 | 388.0 | 17.4 | 82.6 | 4.3 | 4.6 | 8.5 |
| T78 | 565 | 221.0 | 9.9 | 90.1 | 5.8 | 3.6 | 0.5 |
| T79 | 491 | 254.3 | 11.4 | 88.6 | 5.3 | 4.2 | 1.8 |
| T81 | 678 | 273.6 | 12.3 | 87.7 | 7.2 | 3.9 | 1.1 |
| T83 | 612 | 367.6 | 16.5 | 83.5 | 6.4 | 5 | 5.1 |
| T84 | 161 | 58.2 | 2.6 | 97.4 | 1.6 | 1 | 0 |
| T85 | 379 | 162.2 | 7.3 | 92.7 | 3.9 | 3.4 | 0 |
| T86 | 396 | 155.2 | 7.0 | 93 | 4 | 3 | 0 |

| Sample | Number of RoH | Length, Mbp | RoH, % | Non-RoH, % | % of RoH * | | |
|---|---|---|---|---|---|---|---|
| | | | | | Short (S) | Long (L) | Ultra Long (UL) |
| T87 | 124 | 36.6 | 1.6 | 98.4 | 1.3 | 0.4 | 0 |

* Short RoH (< 1 Mbp), Long RoH (>= 1 Mbp < 10 Mbp) and  Ultra Long RoH (>= 10 Mbp).

**Table 4.** Whole genome coverage statistics.

| Sample | Median | Mean | Max | Min |
|---|---|---|---|---|
| 10xmmar | 23 | 24 | 38987 | 0 |
| S44 | 22 | 22.45 | 59070 | 0 |
| S46 | 22 | 22.54 | 68078 | 0 |
| S49 | 21 | 21.23 | 54065 | 0 |
| S50 | 21 | 21.39 | 54873 | 0 |
| T149 | 21 | 21.6 | 58932 | 0 |
| T151 | 22 | 22.6 | 78958 | 0 |
| 10xmzib | 23 | 22.99 | 36551 | 0 |
| CHN | 20 | 20.84 | 39693 | 0 |
| S26 | 23 | 23.46 | 78413 | 0 |
| T104 | 21 | 21.13 | 54566 | 0 |
| T118 | 22 | 21.99 | 58505 | 0 |
| T148 | 22 | 21.81 | 54344 | 0 |
| T150 | 22 | 22.33 | 64501 | 0 |
| T18 | 22 | 21.69 | 52075 | 0 |
| T194 | 21 | 20.8 | 66051 | 0 |
| T26 | 22 | 22.34 | 58942 | 0 |
| T50 | 22 | 22.08 | 65570 | 0 |
| T72 | 23 | 22.46 | 53159 | 0 |
| T8 | 22 | 22.21 | 64849 | 0 |

| Sample | Median | Mean | Max | Min |
|---|---|---|---|---|
| T90 | 21 | 21.36 | 55955 | 0 |
| T24 | 23 | 23.57 | 86673 | 0 |
| T76 | 21 | 21.04 | 69063 | 0 |
| T77 | 20 | 20.71 | 50848 | 0 |
| T78 | 20 | 20.4 | 48726 | 0 |
| T79 | 24 | 24.24 | 66740 | 0 |
| T81 | 20 | 20.32 | 57566 | 0 |
| T82 | 21 | 20.87 | 53307 | 0 |
| T83 | 20 | 20.57 | 59175 | 0 |
| T84 | 20 | 20.56 | 57272 | 0 |
| T85 | 20 | 20.24 | 45092 | 0 |
| T86 | 19 | 19.78 | 52040 | 0 |
| T87 | 21 | 21.06 | 50136 | 0 |
