## Supplementary figures and images for "Genomics of sable (*Martes zibellina)* × pine marten (*Martes martes*) hybridization"

### Supplementary File SF7

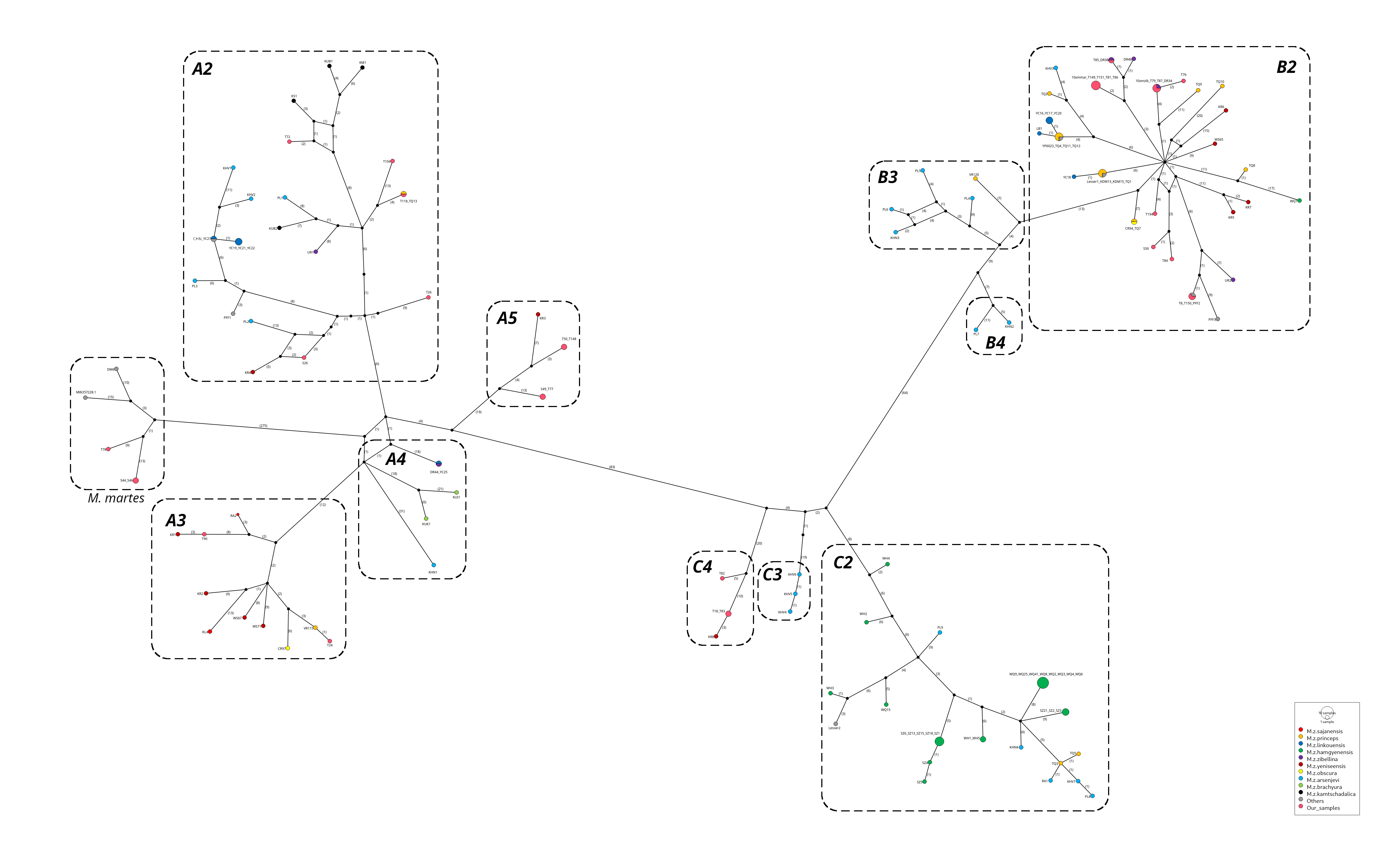
