## Supplementary File SF9 for "Genomics of sable (*Martes zibellina)* × pine marten (*Martes martes*) hybridization"

### 10xmmar

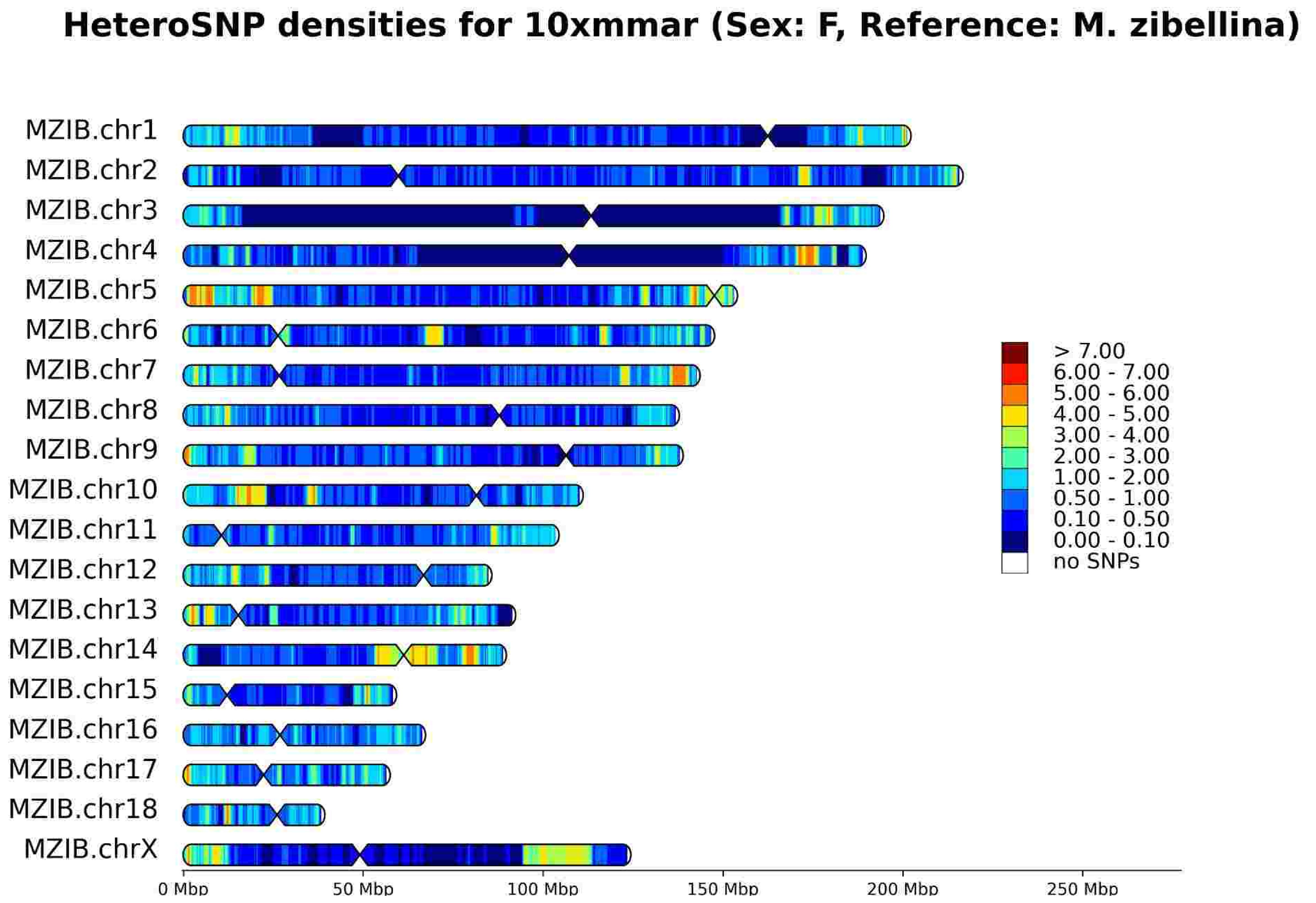
HeteroSNP densities for 10xmmar (Sex: F, Reference: M. zibellina)
MZIB.chr1
MZIB.chr2
MZIB.chr3
MZIB.chr4
MZIB.chr5
MZIB.chr6
MZIB.chr7
MZIB.chr8
MZIB.chr9
MZIB.chr10
MZIB.chr11
MZIB.chr12
MZIB.chr13
MZIB.chr14
MZIB.chr15
MZIB.chr16
MZIB.chr17
MZIB.chr18
MZIB.chrX
> 7.00
6.00 - 7.00
5.00 - 6.00
4.00 - 5.00
3.00 - 4.00
2.00 - 3.00
1.00 - 2.00
0.50 - 1.00
0.10 - 0.50
0.00 - 0.10
no SNPs
0 Mbp
50 Mbp
100 Mbp
150 Mbp
200 Mbp
250 Mbp

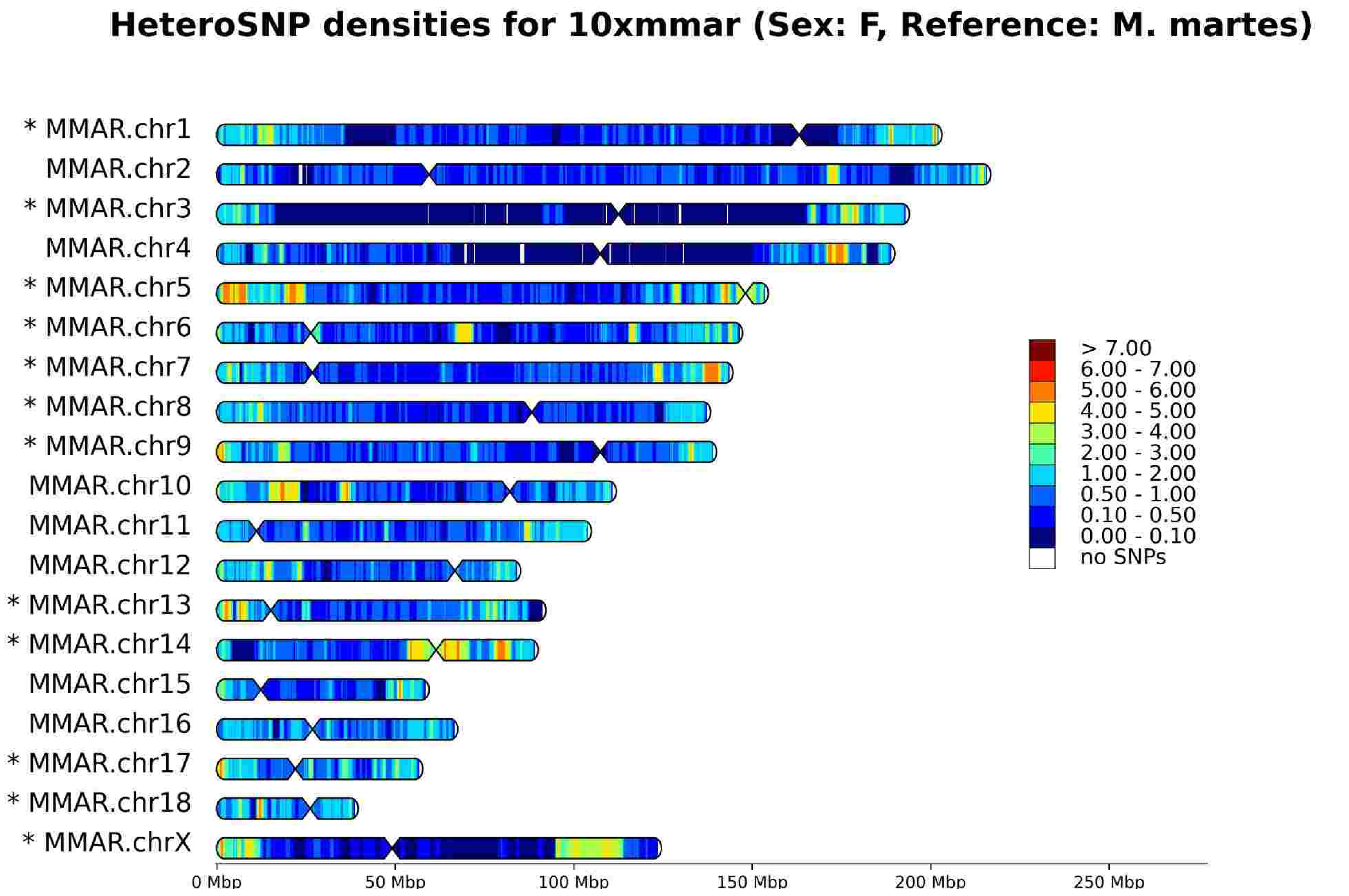
HeteroSNP densities for 10xmmar (Sex: F, Reference: M. martes)
* MMAR.chr1
MMAR.chr2
* MMAR.chr3
MMAR.chr4
* MMAR.chr5
* MMAR.chr6
* MMAR.chr7
* MMAR.chr8
* MMAR.chr9
MMAR.chr10
MMAR.chr11
MMAR.chr12
* MMAR.chr13
* MMAR.chr14
MMAR.chr15
MMAR.chr16
* MMAR.chr17
* MMAR.chr18
* MMAR.chrX
> 7.00
6.00 - 7.00
5.00 - 6.00
4.00 - 5.00
3.00 - 4.00
2.00 - 3.00
1.00 - 2.00
0.50 - 1.00
0.10 - 0.50
0.00 - 0.10
no SNPs
0 Mbp
50 Mbp
100 Mbp
150 Mbp
200 Mbp
250 Mbp

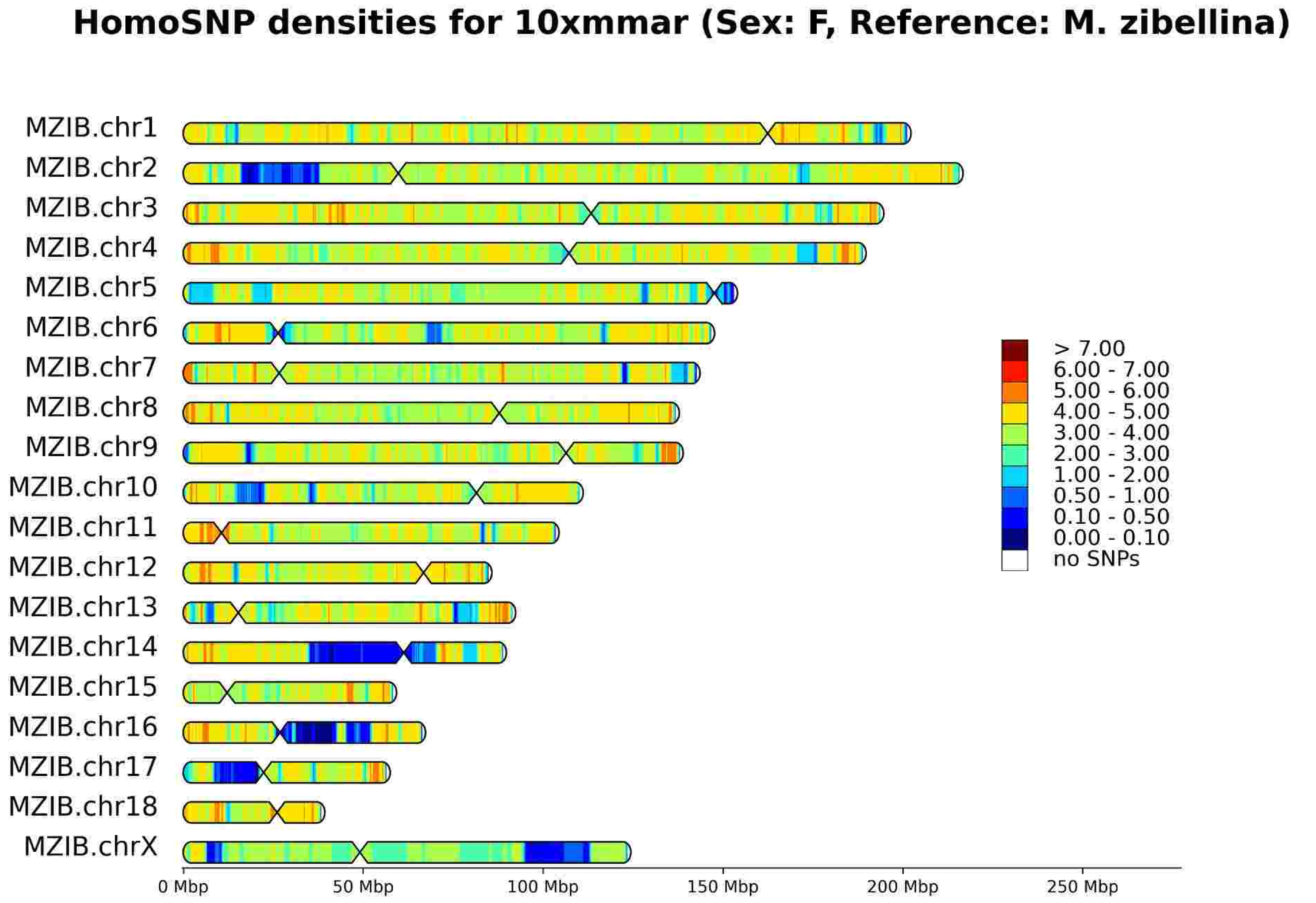
HomoSNP densities for 10xmmar (Sex: F, Reference: M. zibellina)
MZIB.chr1
MZIB.chr2
MZIB.chr3
MZIB.chr4
MZIB.chr5
MZIB.chr6
MZIB.chr7
MZIB.chr8
MZIB.chr9
MZIB.chr10
MZIB.chr11
MZIB.chr12
MZIB.chr13
MZIB.chr14
MZIB.chr15
MZIB.chr16
MZIB.chr17
MZIB.chr18
MZIB.chrX
> 7.00
6.00 - 7.00
5.00 - 6.00
4.00 - 5.00
3.00 - 4.00
2.00 - 3.00
1.00 - 2.00
0.50 - 1.00
0.10 - 0.50
0.00 - 0.10
no SNPs
0 Mbp
50 Mbp
100 Mbp
150 Mbp
200 Mbp
250 Mbp

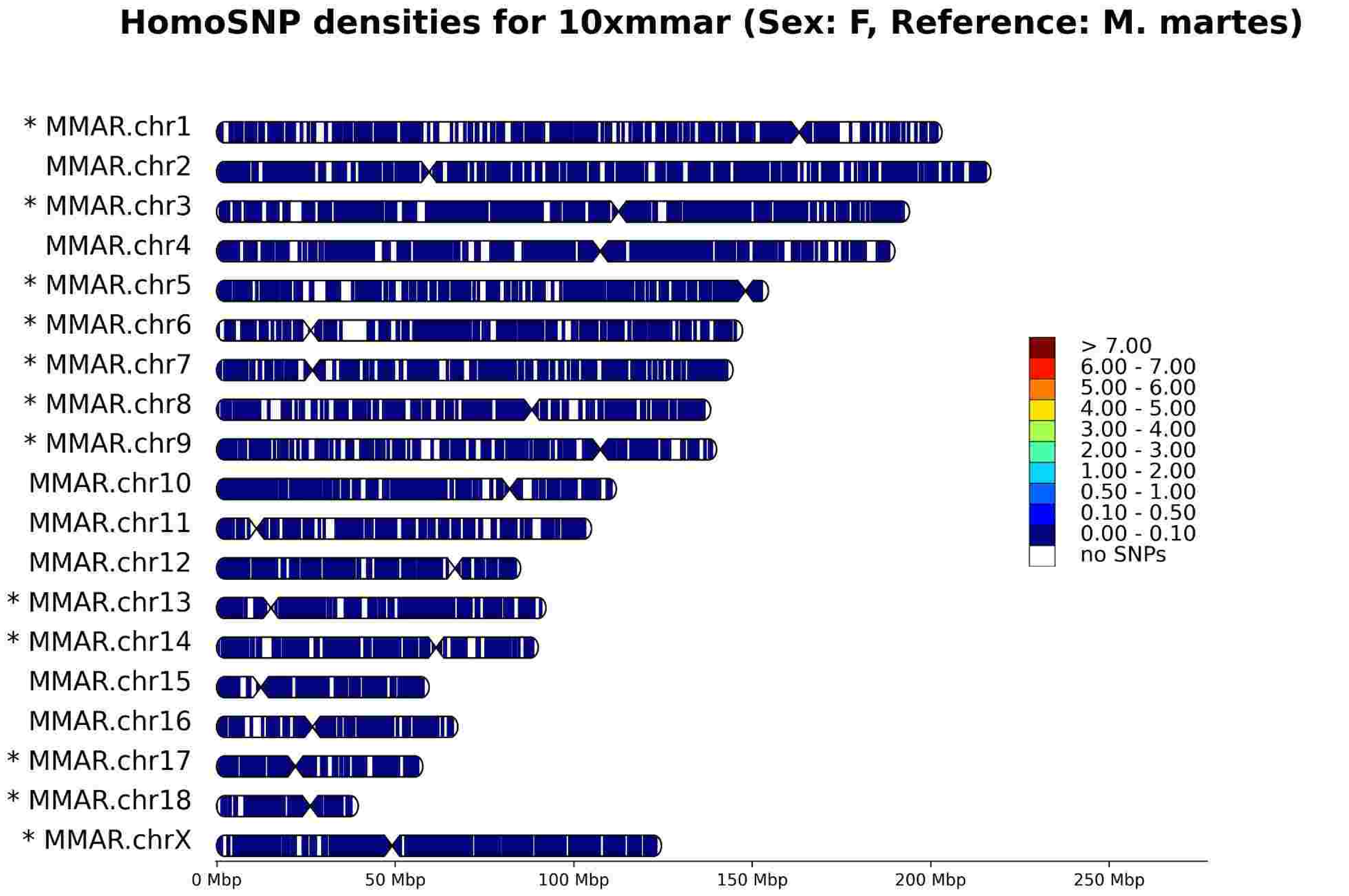
HomoSNP densities for 10xmmar (Sex: F, Reference: M. martes)
* MMAR.chr1
MMAR.chr2
* MMAR.chr3
MMAR.chr4
* MMAR.chr5
* MMAR.chr6
* MMAR.chr7
* MMAR.chr8
* MMAR.chr9
MMAR.chr10
MMAR.chr11
MMAR.chr12
* MMAR.chr13
* MMAR.chr14
MMAR.chr15
MMAR.chr16
* MMAR.chr17
* MMAR.chr18
* MMAR.chrX
> 7.00
6.00 - 7.00
5.00 - 6.00
4.00 - 5.00
3.00 - 4.00
2.00 - 3.00
1.00 - 2.00
0.50 - 1.00
0.10 - 0.50
0.00 - 0.10
no SNPs
0 Mbp
50 Mbp
100 Mbp
150 Mbp
200 Mbp
250 Mbp

### 10xmzib

HeteroSNP densities for 10xmzib (Sex: F, Reference: M. zibellina)

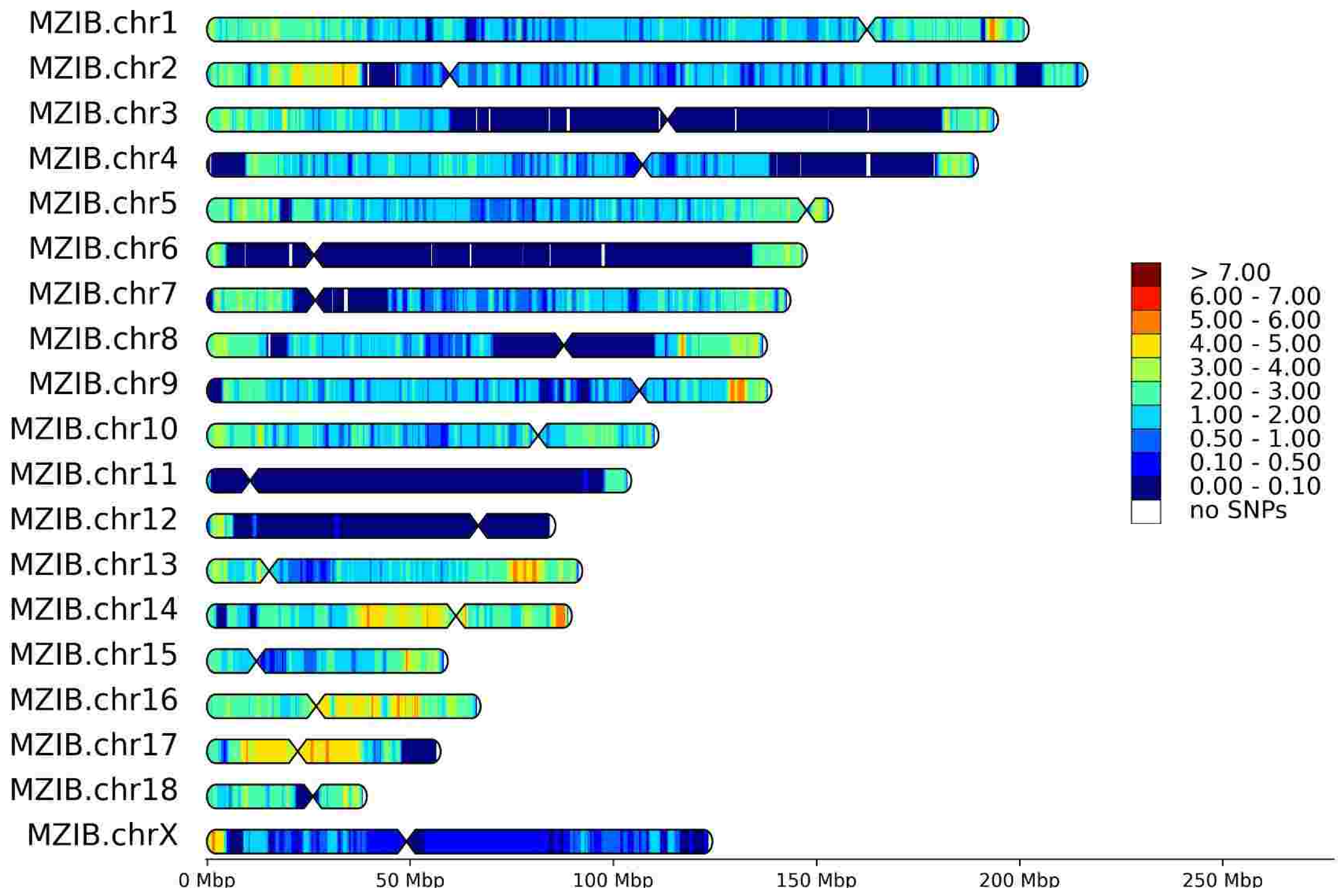


HeteroSNP densities for 10xmzib (Sex: F, Reference: M. martes)

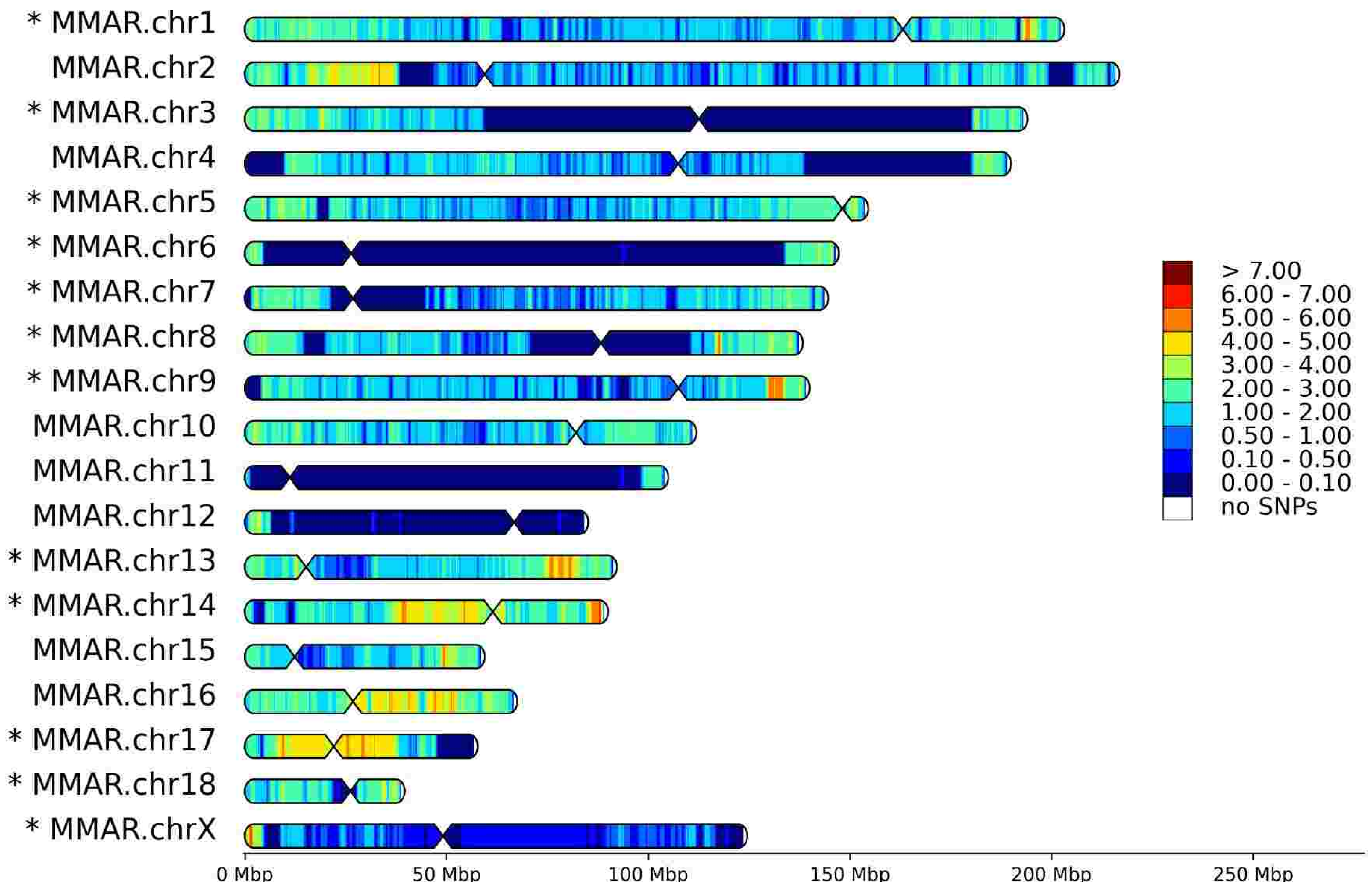


HomoSNP densities for 10xmzib (Sex: F, Reference: M. zibellina)

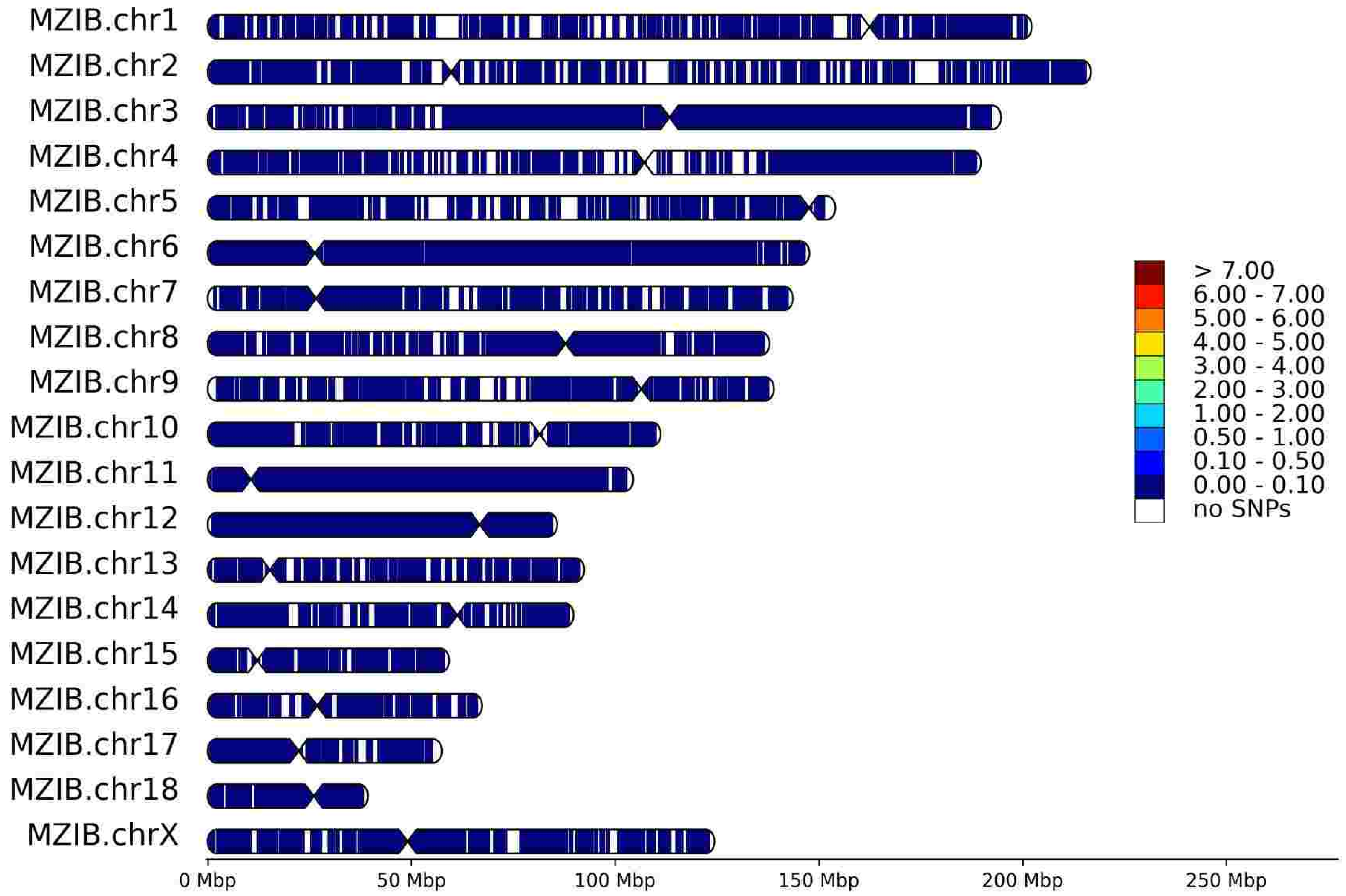


HomoSNP densities for 10xmzib (Sex: F, Reference: M. martes)

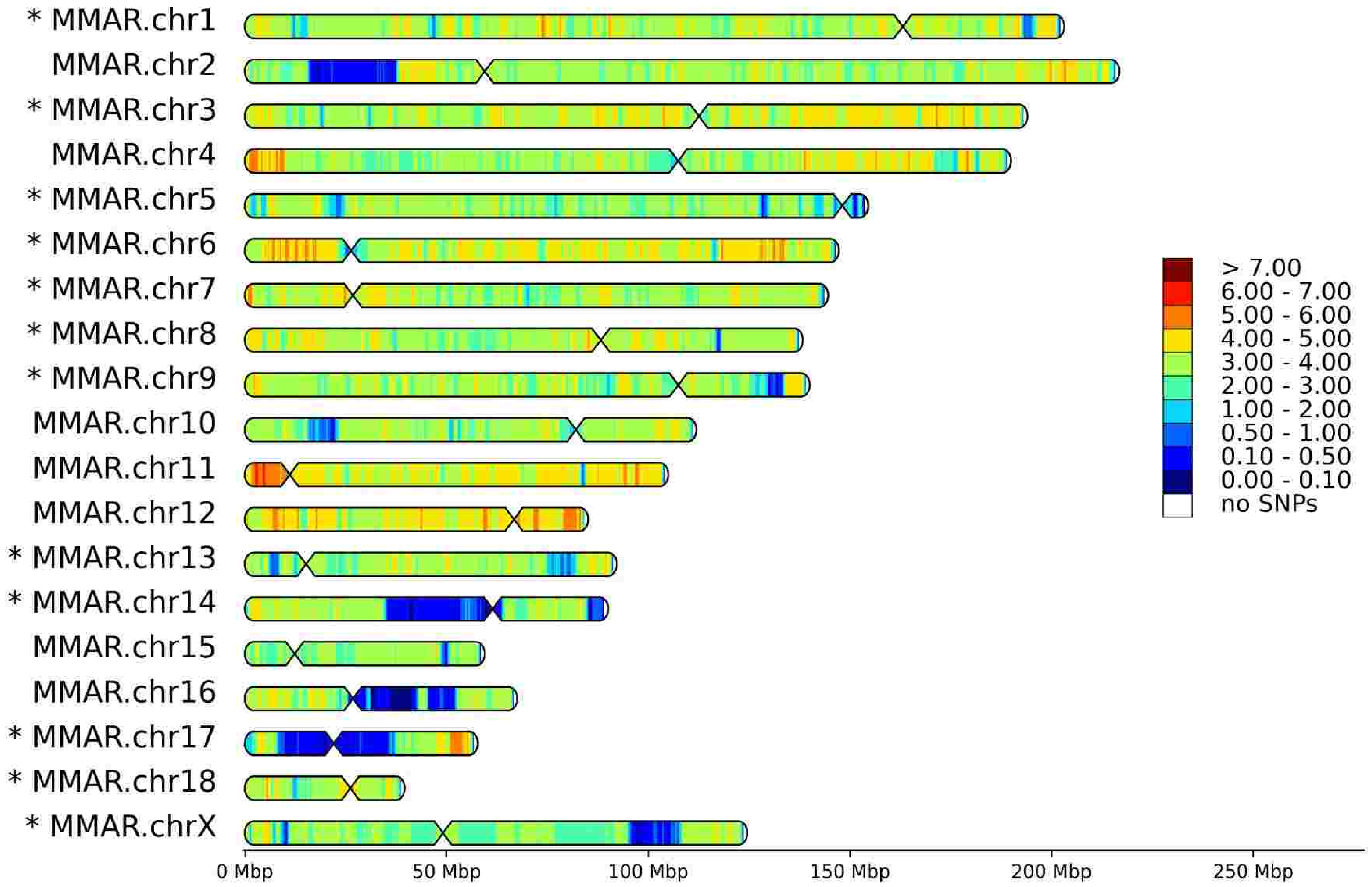

### CHN

HeteroSNP densities for CHN (Sex: F, Reference: M. zibellina)

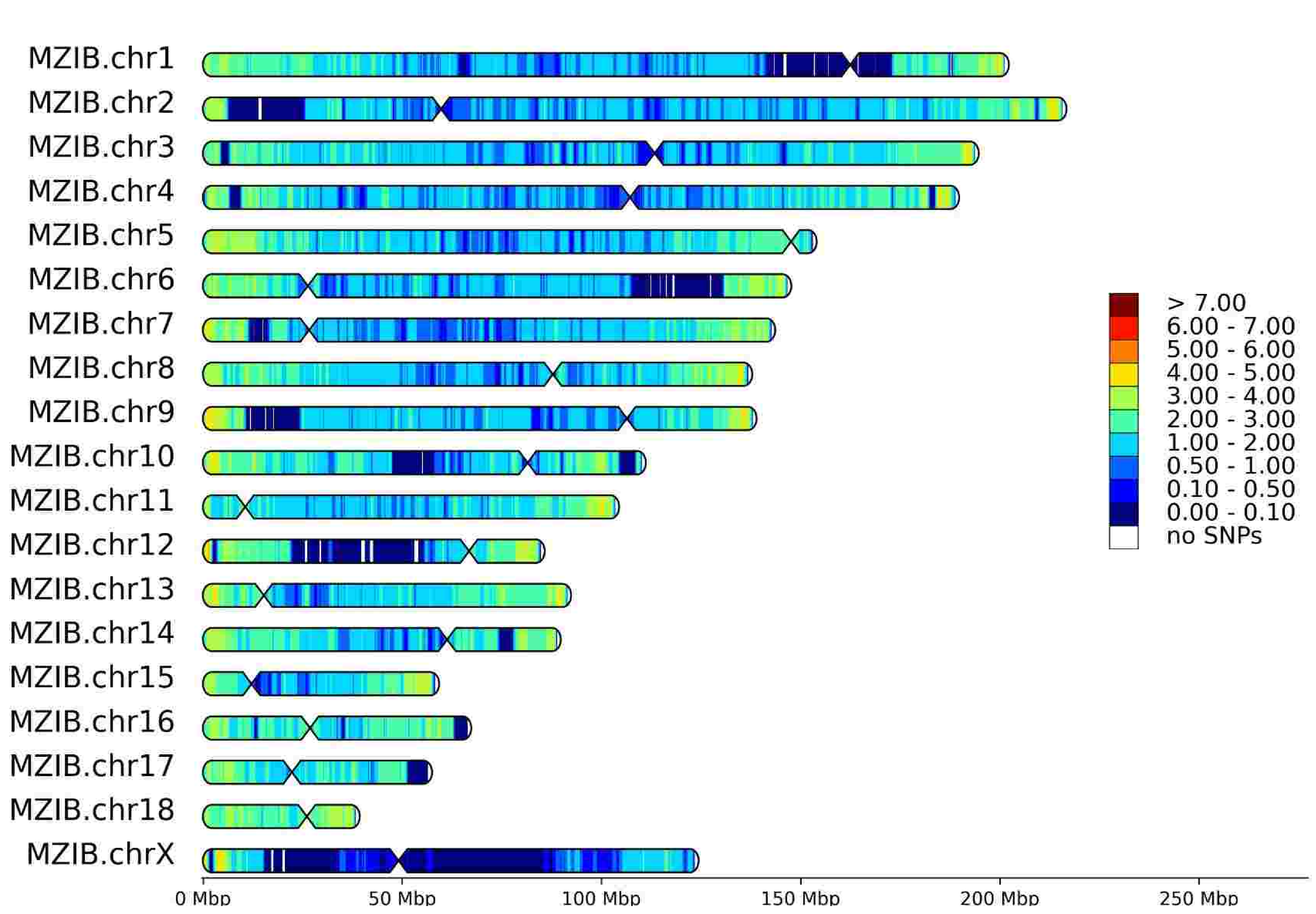


HeteroSNP densities for CHN (Sex: F, Reference: M. martes)

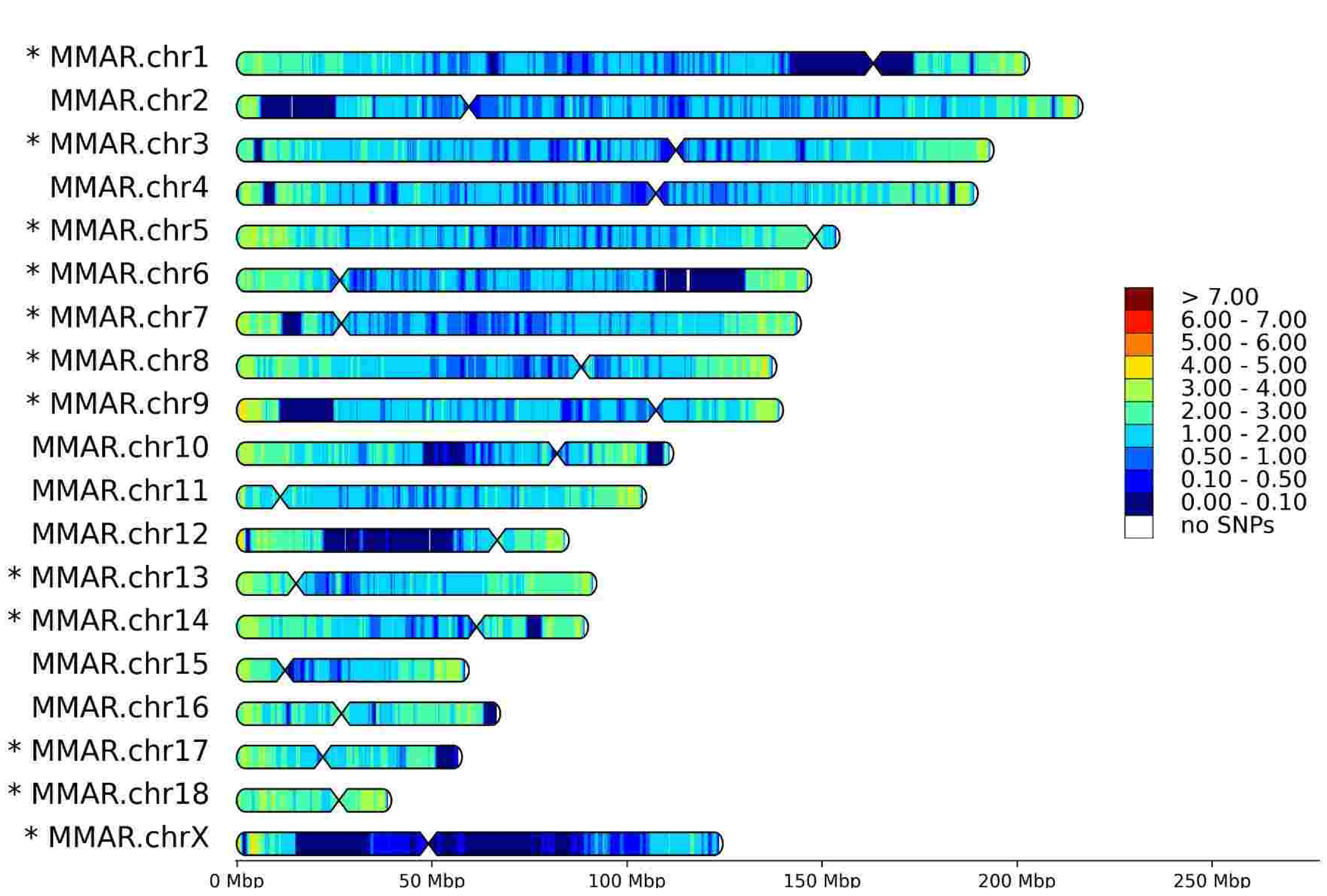


HomoSNP densities for CHN (Sex: F, Reference: M. zibellina)

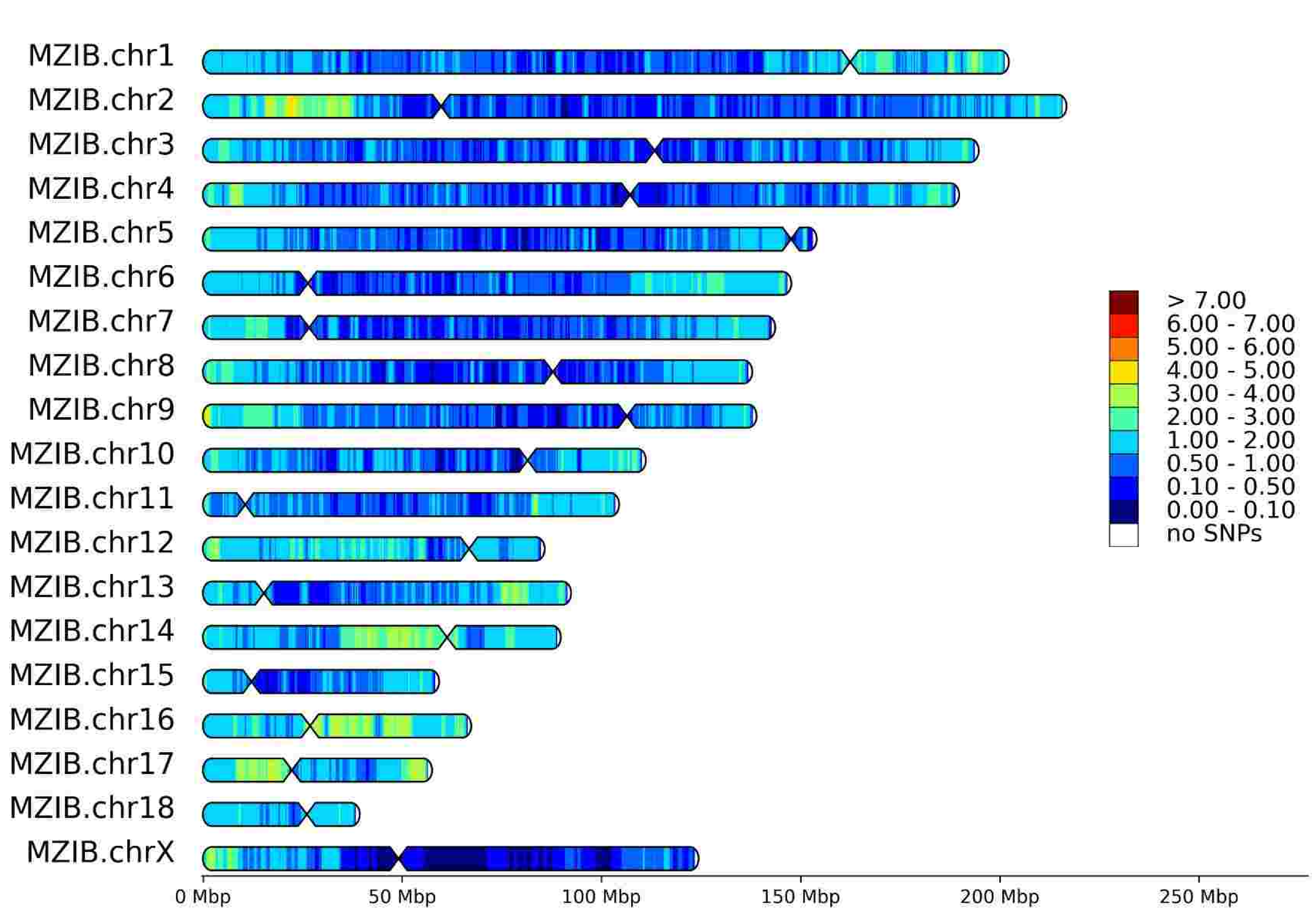


HomoSNP densities for CHN (Sex: F, Reference: M. martes)

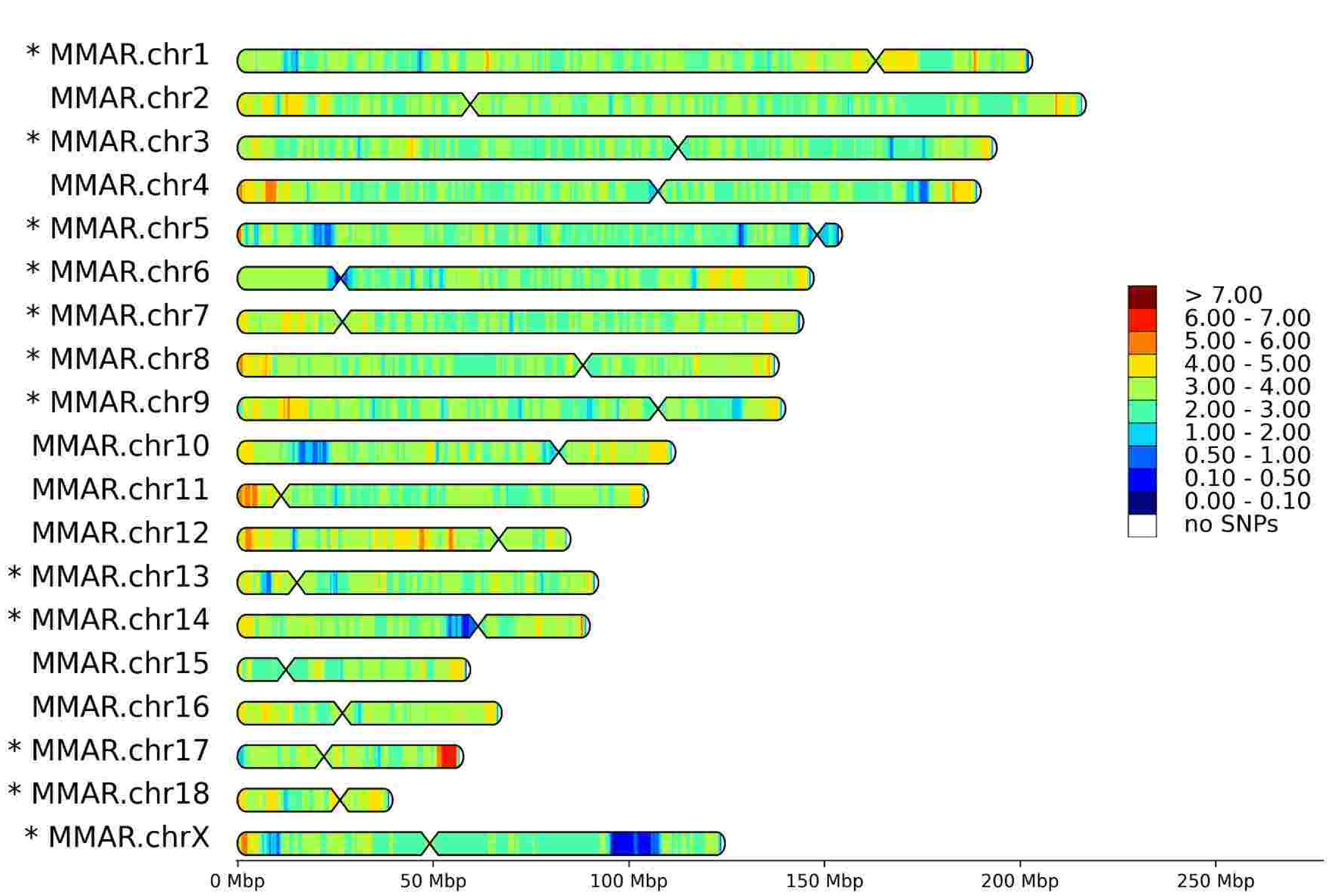

# S26

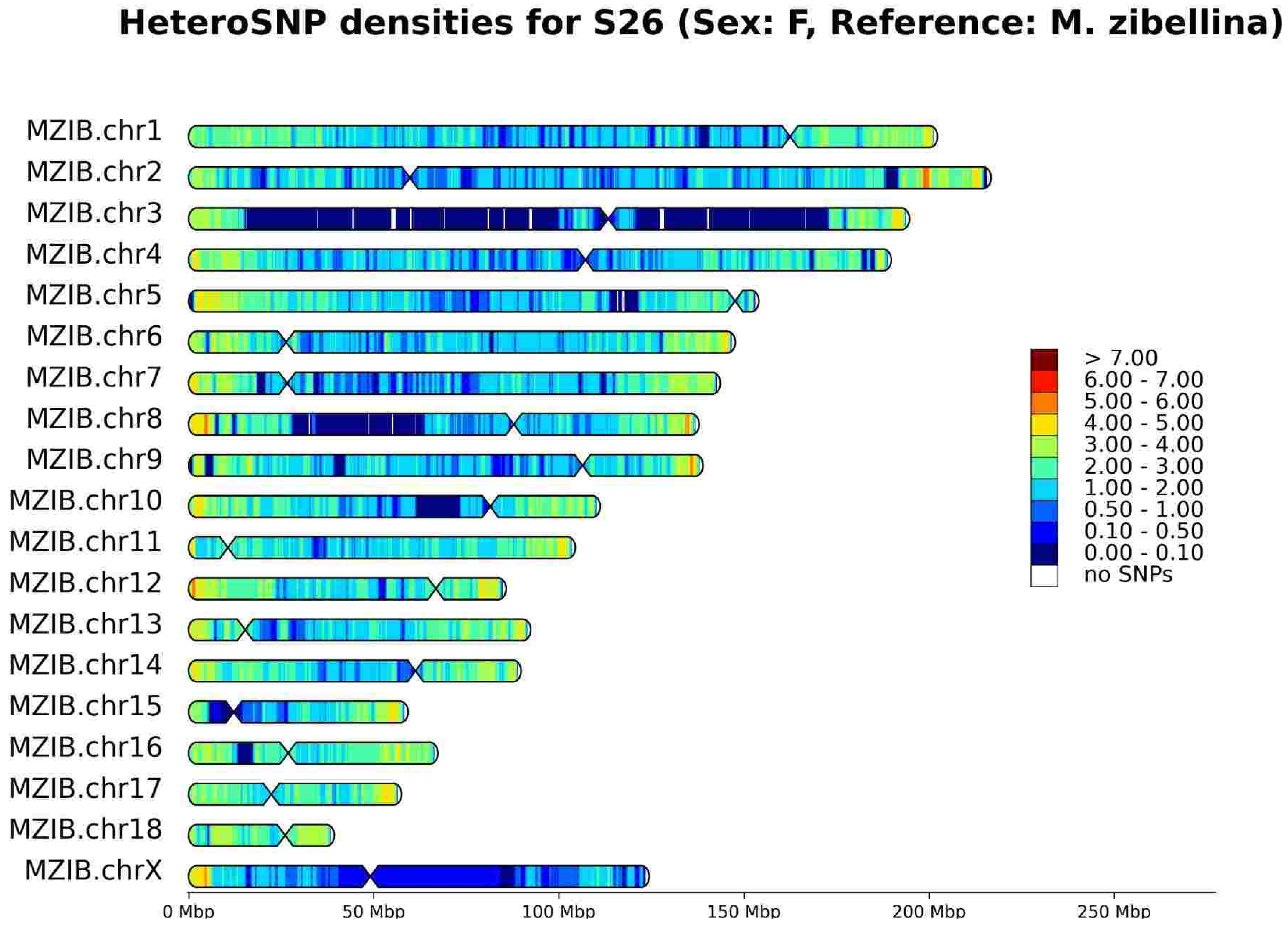
HeteroSNP densities for S26 (Sex: F, Reference: M. zibellina)
MZIB.chr1
MZIB.chr2
MZIB.chr3
MZIB.chr4
MZIB.chr5
MZIB.chr6
MZIB.chr7
MZIB.chr8
MZIB.chr9
MZIB.chr10
MZIB.chr11
MZIB.chr12
MZIB.chr13
MZIB.chr14
MZIB.chr15
MZIB.chr16
MZIB.chr17
MZIB.chr18
MZIB.chrX
> 7.00
6.00 - 7.00
5.00 - 6.00
4.00 - 5.00
3.00 - 4.00
2.00 - 3.00
1.00 - 2.00
0.50 - 1.00
0.10 - 0.50
0.00 - 0.10
no SNPs
0 Mbp
50 Mbp
100 Mbp
150 Mbp
200 Mbp
250 Mbp

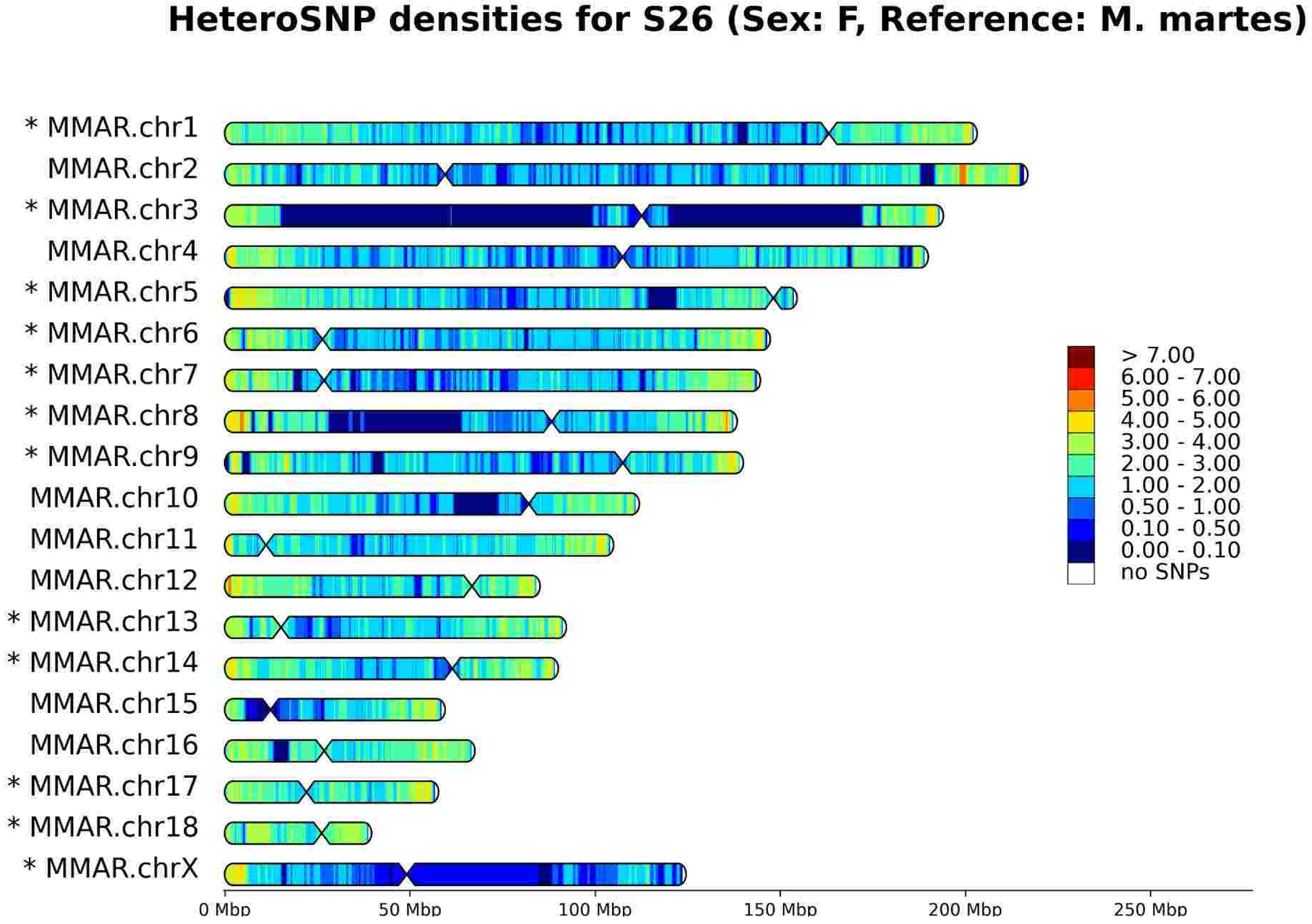
HeteroSNP densities for S26 (Sex: F, Reference: M. martes)
* MMAR.chr1
MMAR.chr2
* MMAR.chr3
MMAR.chr4
* MMAR.chr5
* MMAR.chr6
* MMAR.chr7
* MMAR.chr8
* MMAR.chr9
MMAR.chr10
MMAR.chr11
MMAR.chr12
* MMAR.chr13
* MMAR.chr14
MMAR.chr15
MMAR.chr16
* MMAR.chr17
* MMAR.chr18
* MMAR.chrX
> 7.00
6.00 - 7.00
5.00 - 6.00
4.00 - 5.00
3.00 - 4.00
2.00 - 3.00
1.00 - 2.00
0.50 - 1.00
0.10 - 0.50
0.00 - 0.10
no SNPs
0 Mbp
50 Mbp
100 Mbp
150 Mbp
200 Mbp
250 Mbp

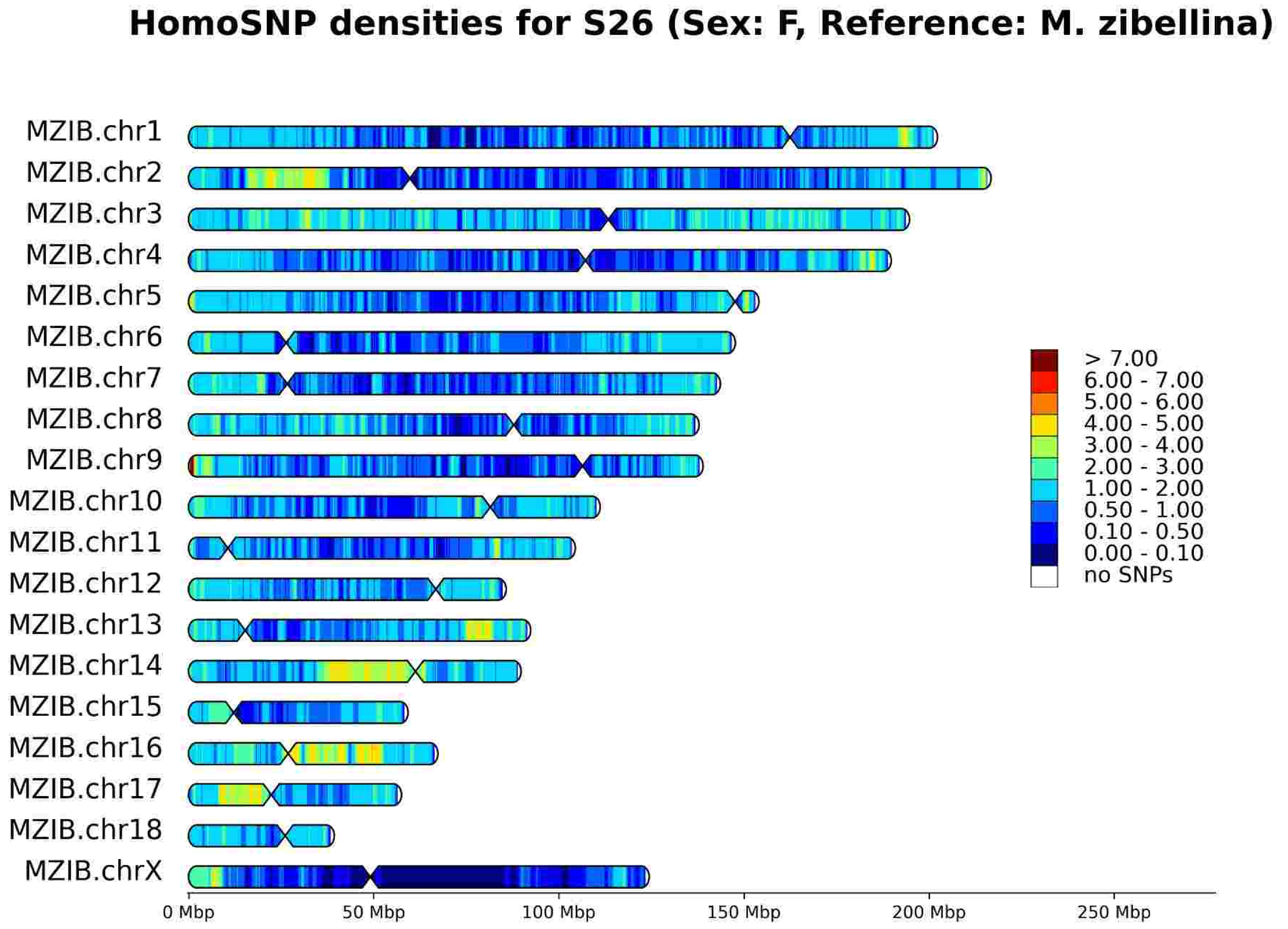
HomoSNP densities for S26 (Sex: F, Reference: M. zibellina)
MZIB.chr1
MZIB.chr2
MZIB.chr3
MZIB.chr4
MZIB.chr5
MZIB.chr6
MZIB.chr7
MZIB.chr8
MZIB.chr9
MZIB.chr10
MZIB.chr11
MZIB.chr12
MZIB.chr13
MZIB.chr14
MZIB.chr15
MZIB.chr16
MZIB.chr17
MZIB.chr18
MZIB.chrX
> 7.00
6.00 - 7.00
5.00 - 6.00
4.00 - 5.00
3.00 - 4.00
2.00 - 3.00
1.00 - 2.00
0.50 - 1.00
0.10 - 0.50
0.00 - 0.10
no SNPs
0 Mbp
50 Mbp
100 Mbp
150 Mbp
200 Mbp
250 Mbp

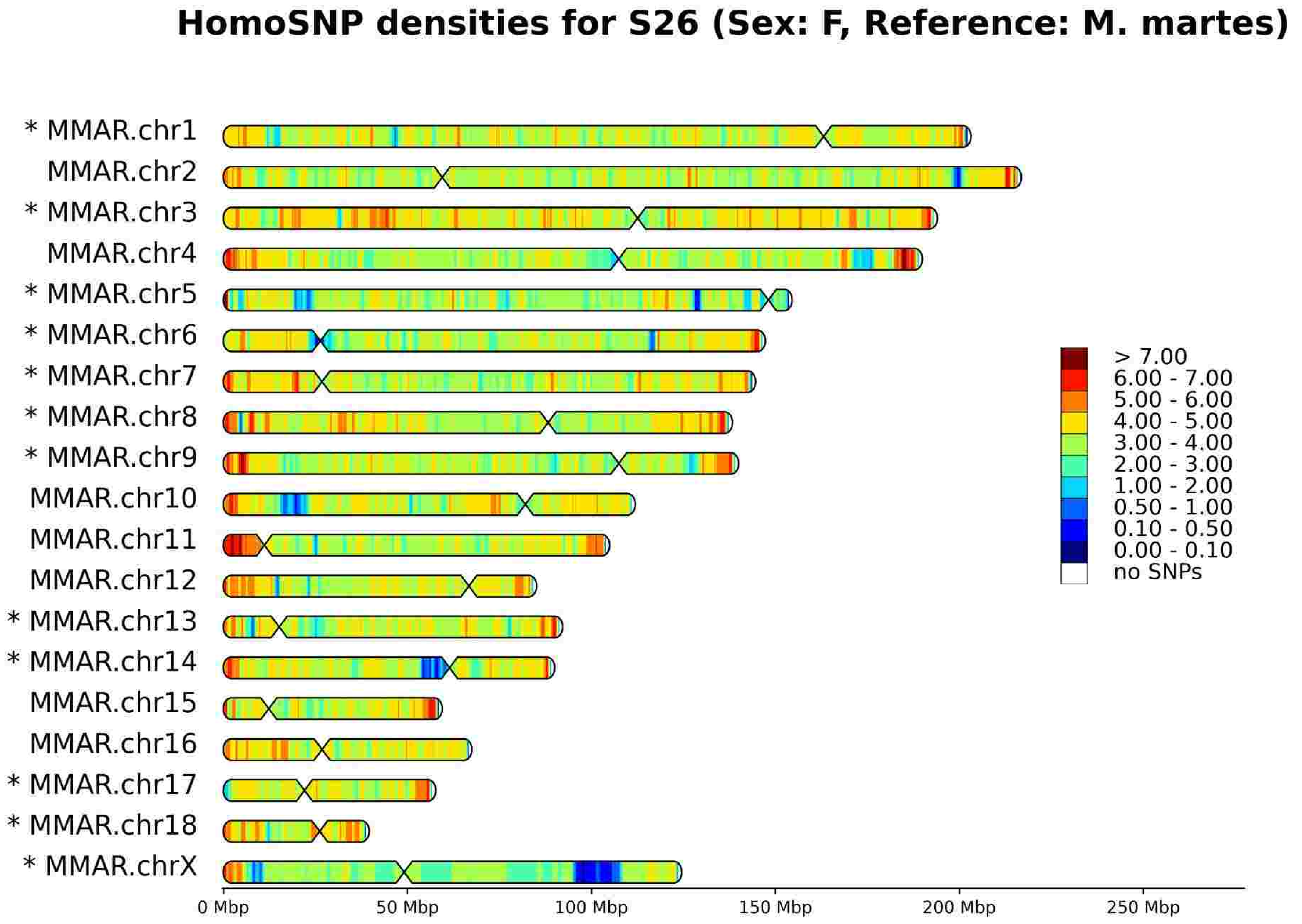
HomoSNP densities for S26 (Sex: F, Reference: M. martes)
* MMAR.chr1
MMAR.chr2
* MMAR.chr3
MMAR.chr4
* MMAR.chr5
* MMAR.chr6
* MMAR.chr7
* MMAR.chr8
* MMAR.chr9
MMAR.chr10
MMAR.chr11
MMAR.chr12
* MMAR.chr13
* MMAR.chr14
MMAR.chr15
MMAR.chr16
* MMAR.chr17
* MMAR.chr18
* MMAR.chrX
> 7.00
6.00 - 7.00
5.00 - 6.00
4.00 - 5.00
3.00 - 4.00
2.00 - 3.00
1.00 - 2.00
0.50 - 1.00
0.10 - 0.50
0.00 - 0.10
no SNPs
0 Mbp
50 Mbp
100 Mbp
150 Mbp
200 Mbp
250 Mbp

# S44

HeteroSNP densities for S44 (Sex: F, Reference: M. zibellina)

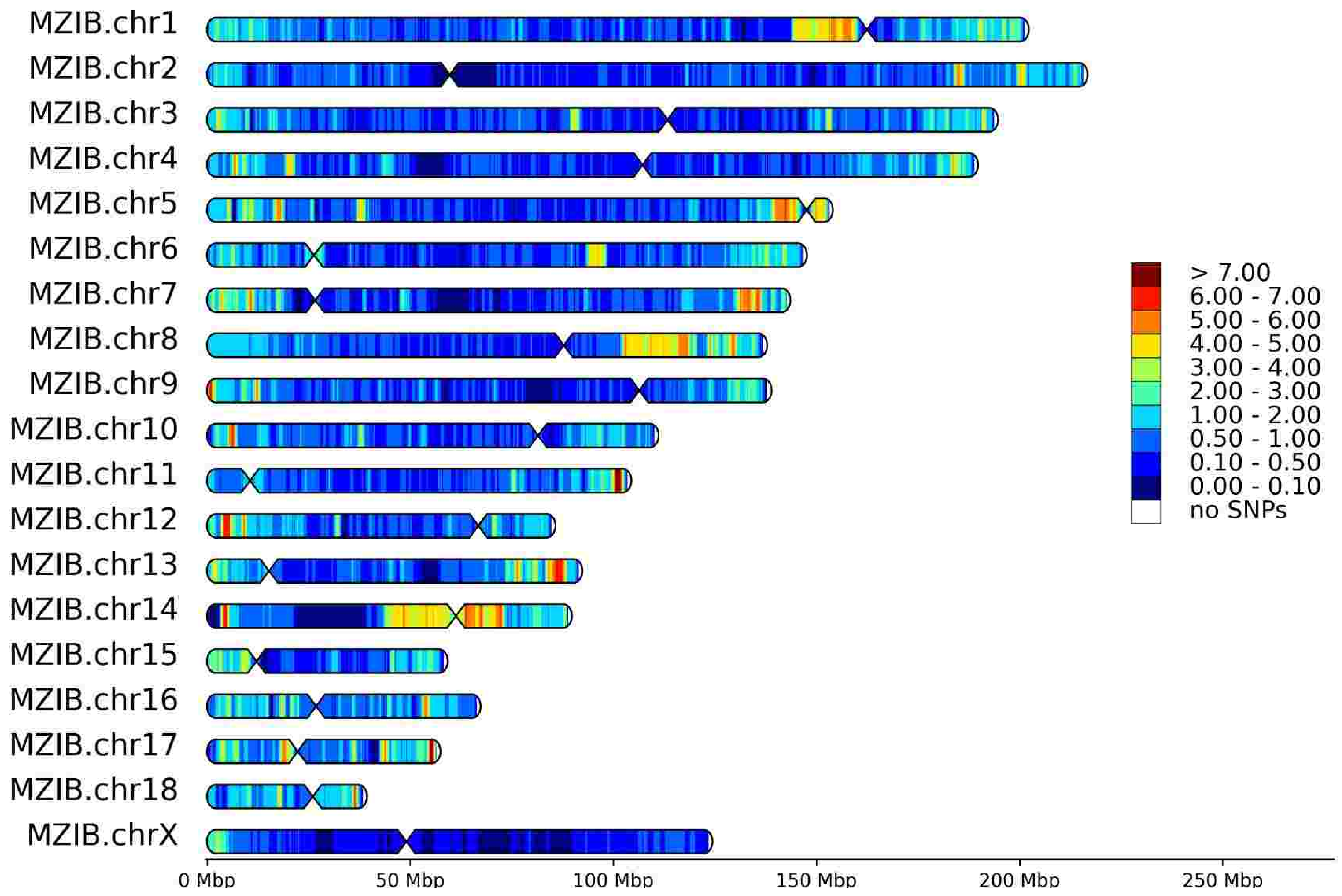


HeteroSNP densities for S44 (Sex: F, Reference: M. martes)

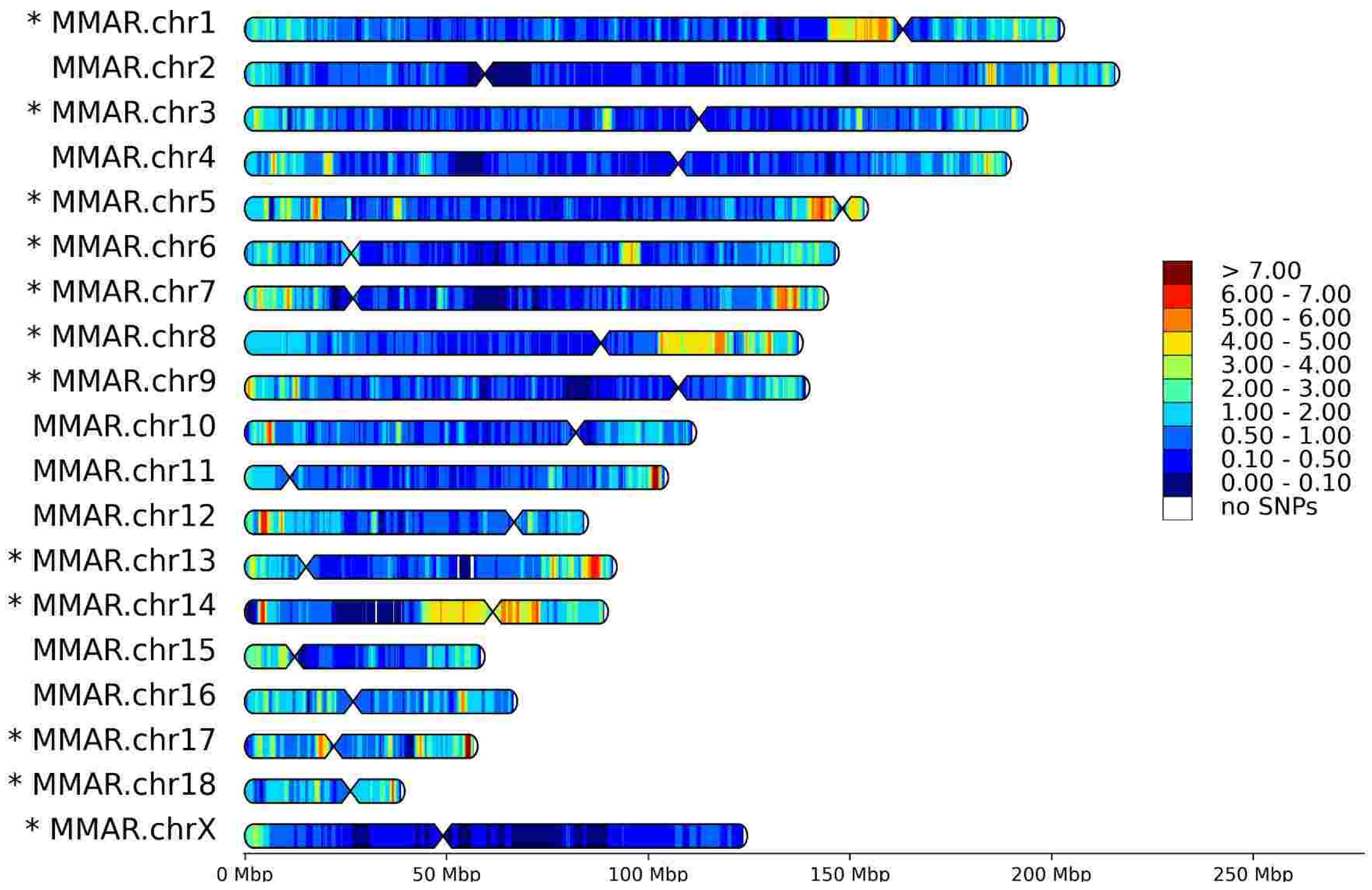


HomoSNP densities for S44 (Sex: F, Reference: M. zibellina)

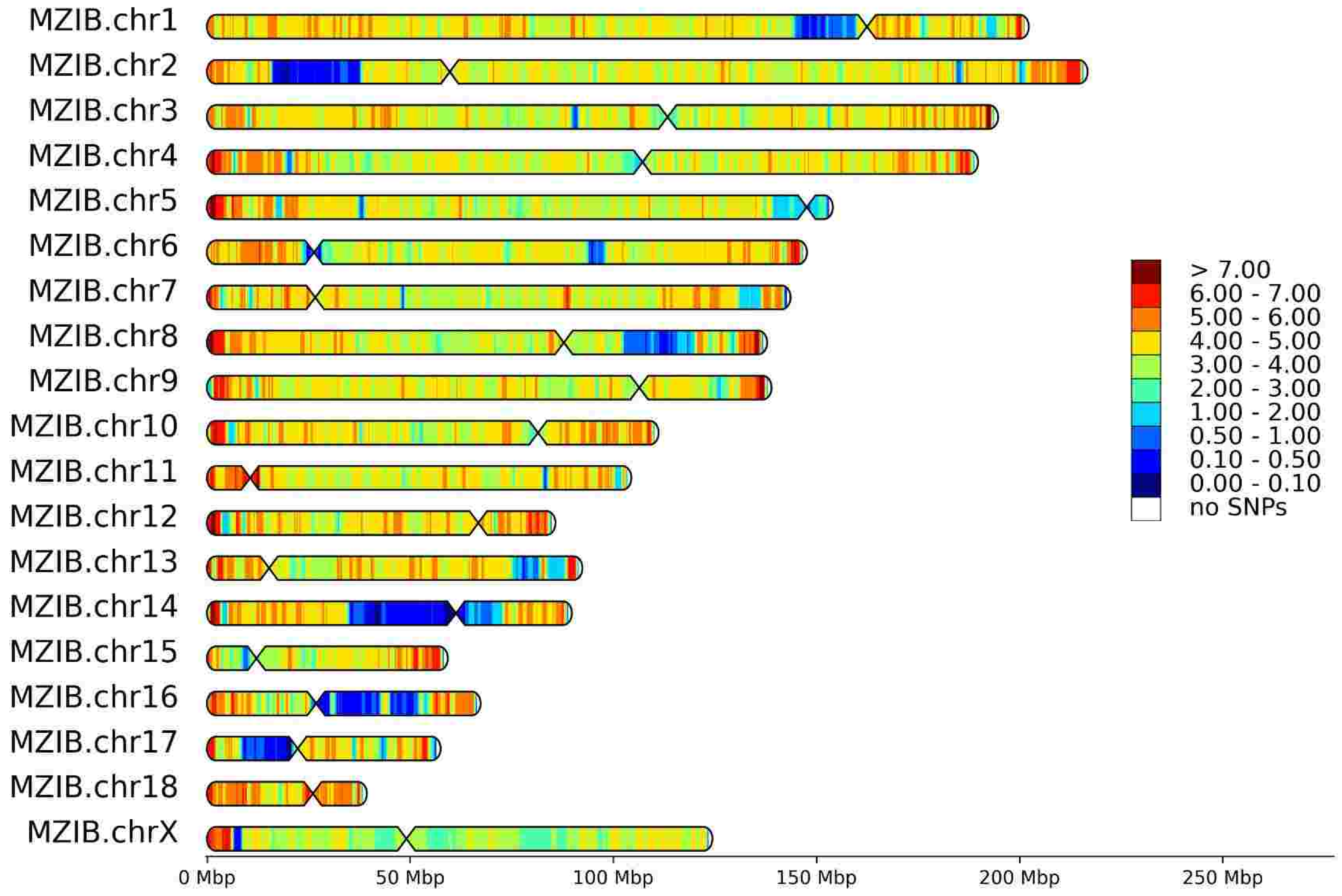


HomoSNP densities for S44 (Sex: F, Reference: M. martes)

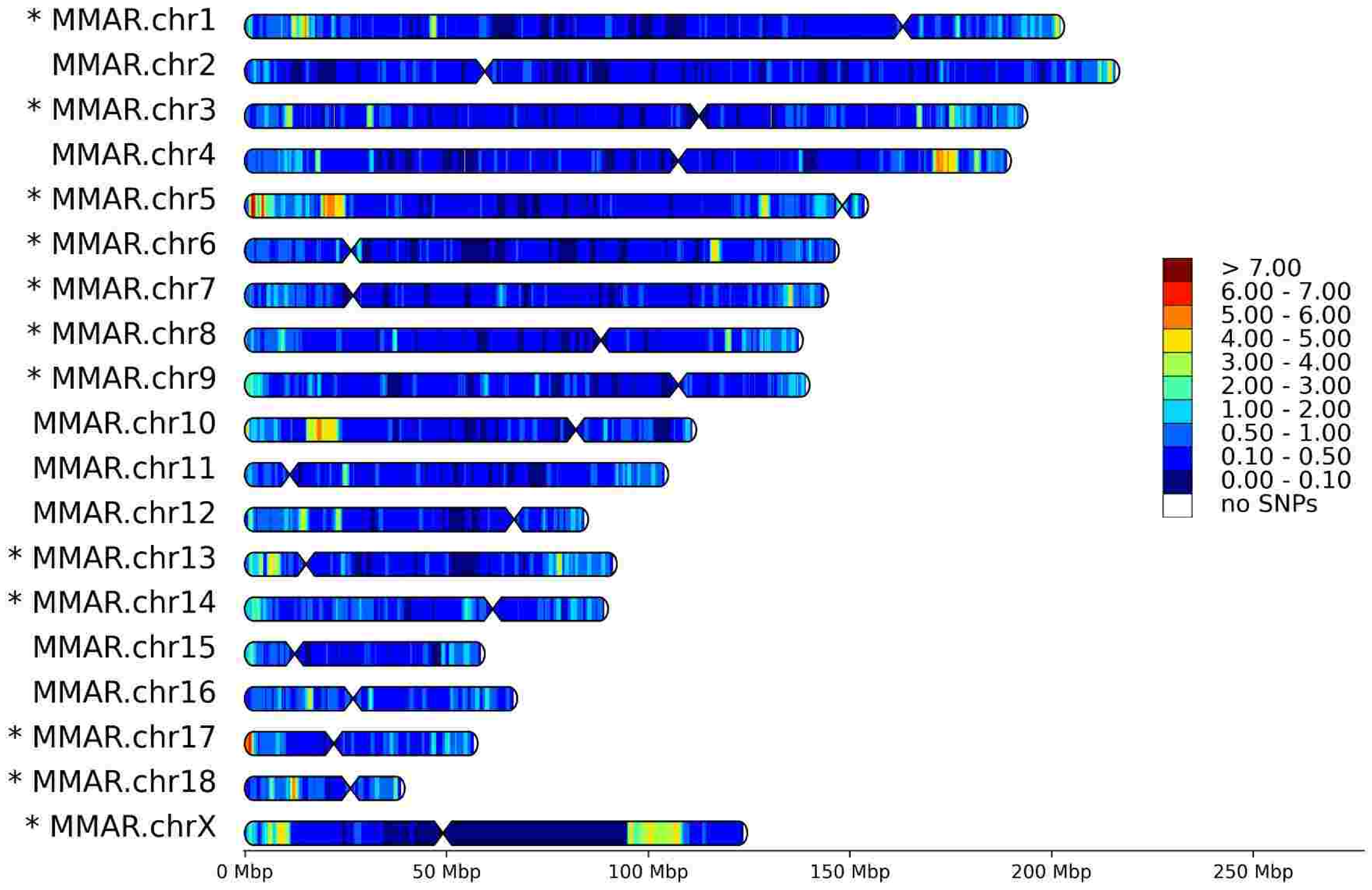

# S46

HeteroSNP densities for S46 (Sex: F, Reference: M. zibellina)

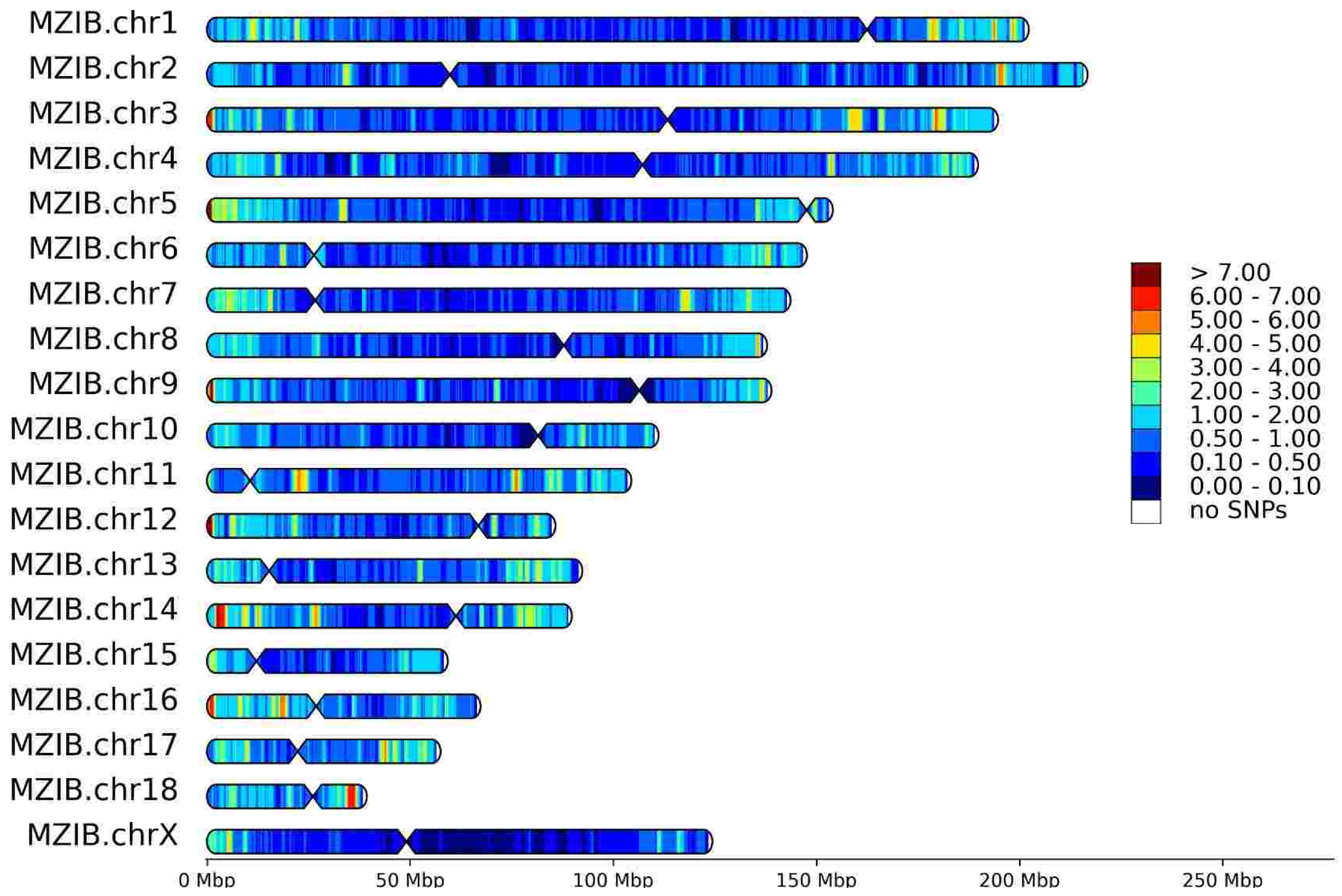


HeteroSNP densities for S46 (Sex: F, Reference: M. martes)

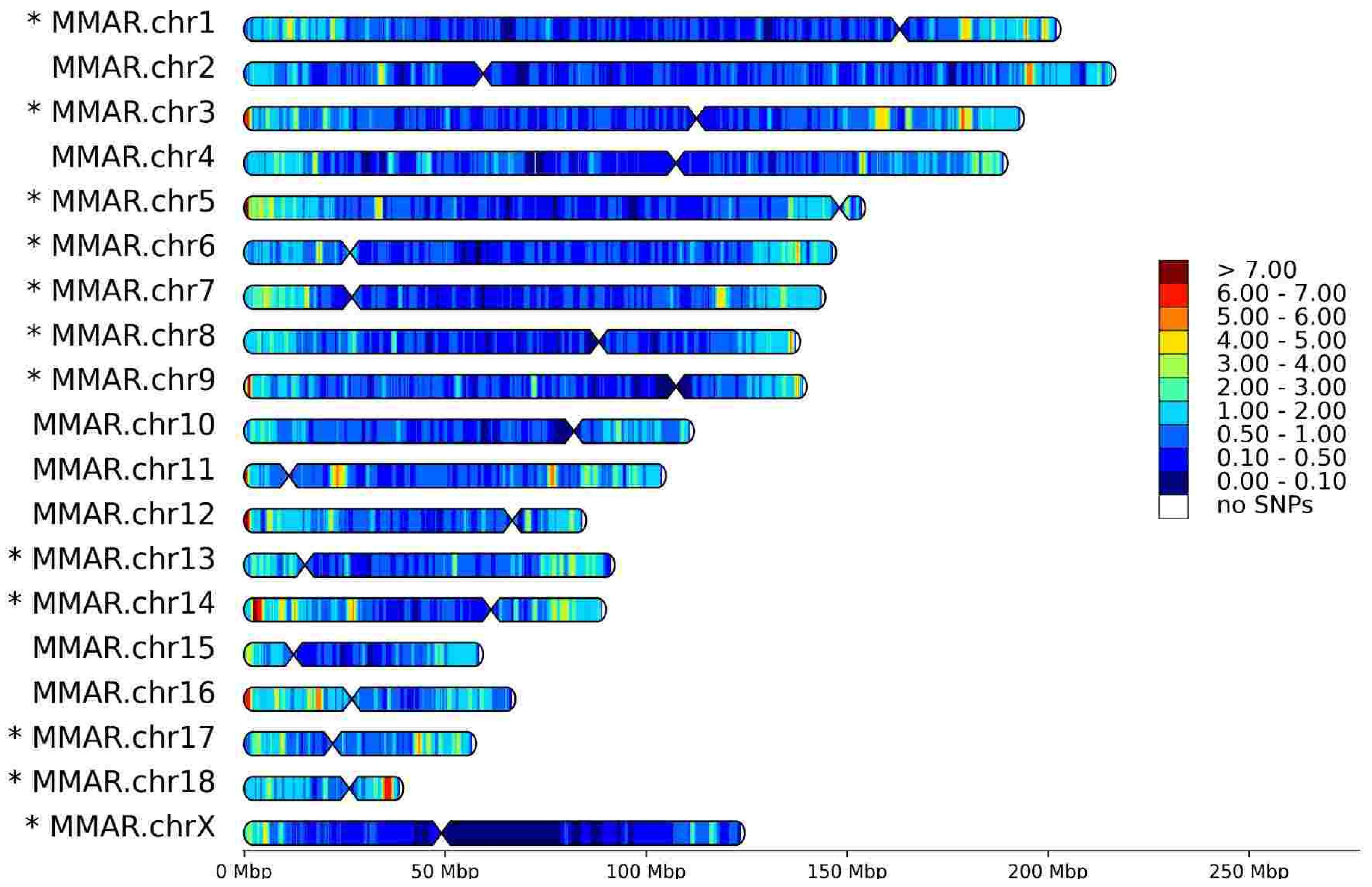


HomoSNP densities for S46 (Sex: F, Reference: M. zibellina)

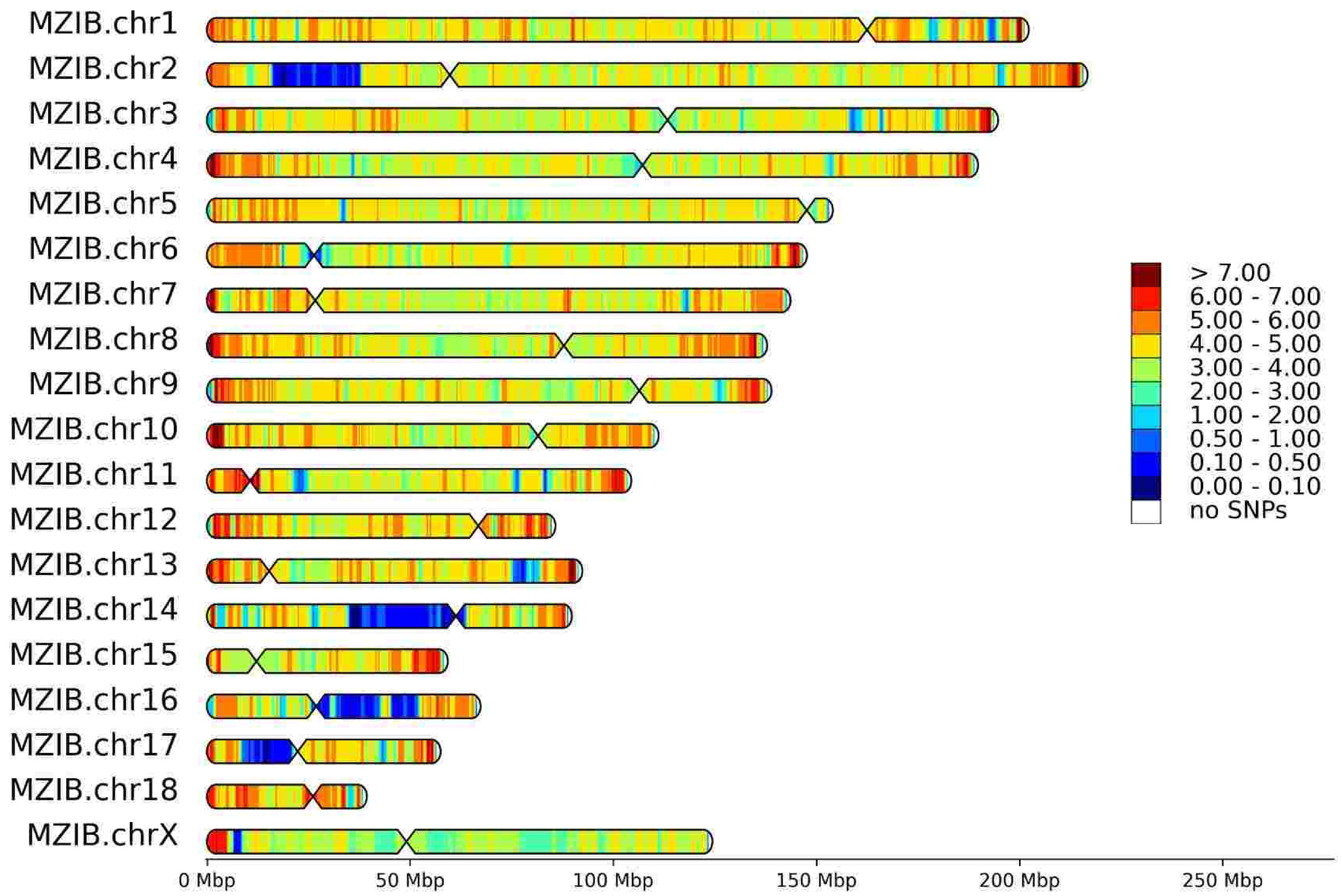


HomoSNP densities for S46 (Sex: F, Reference: M. martes)

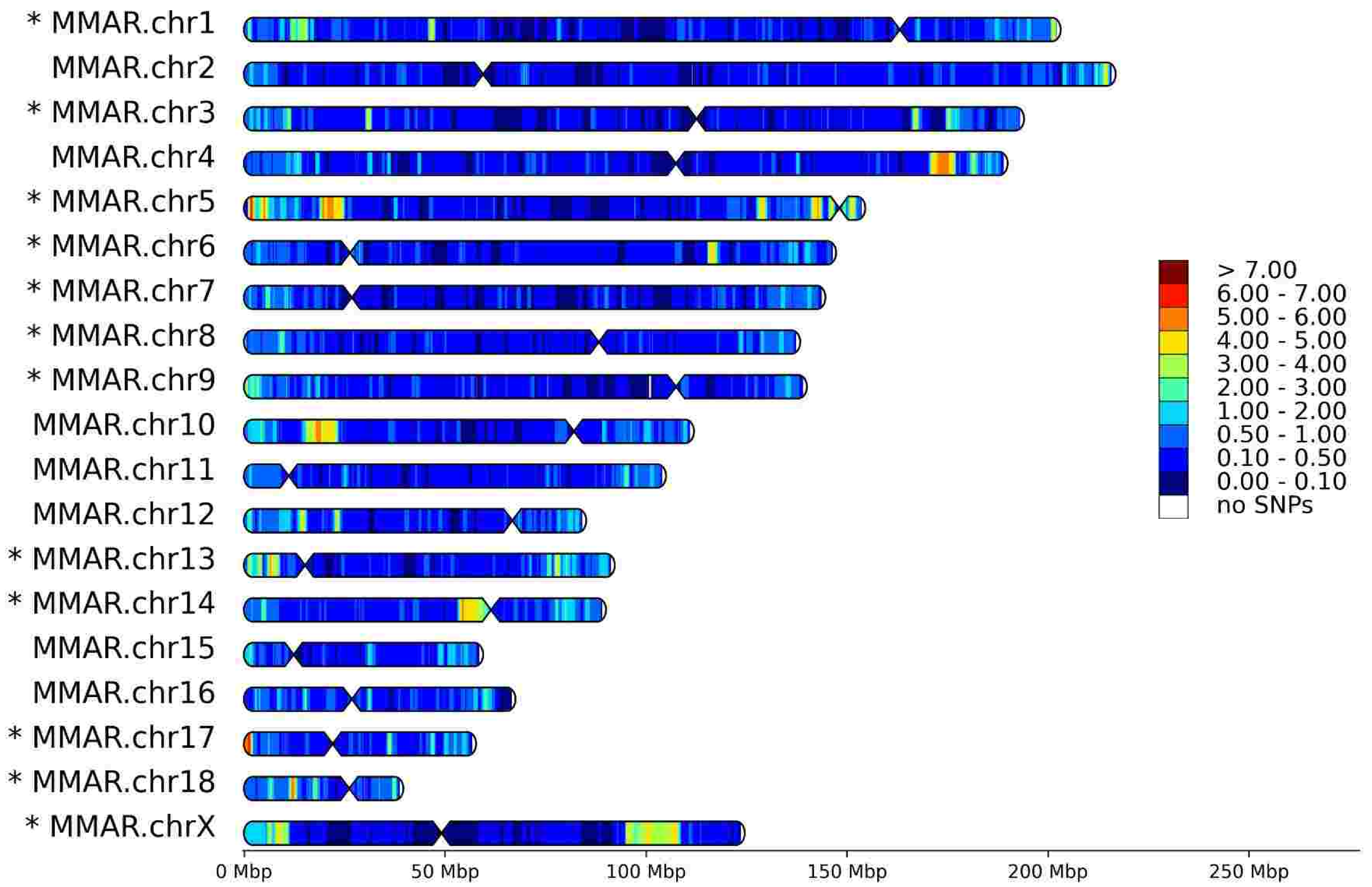

# S49

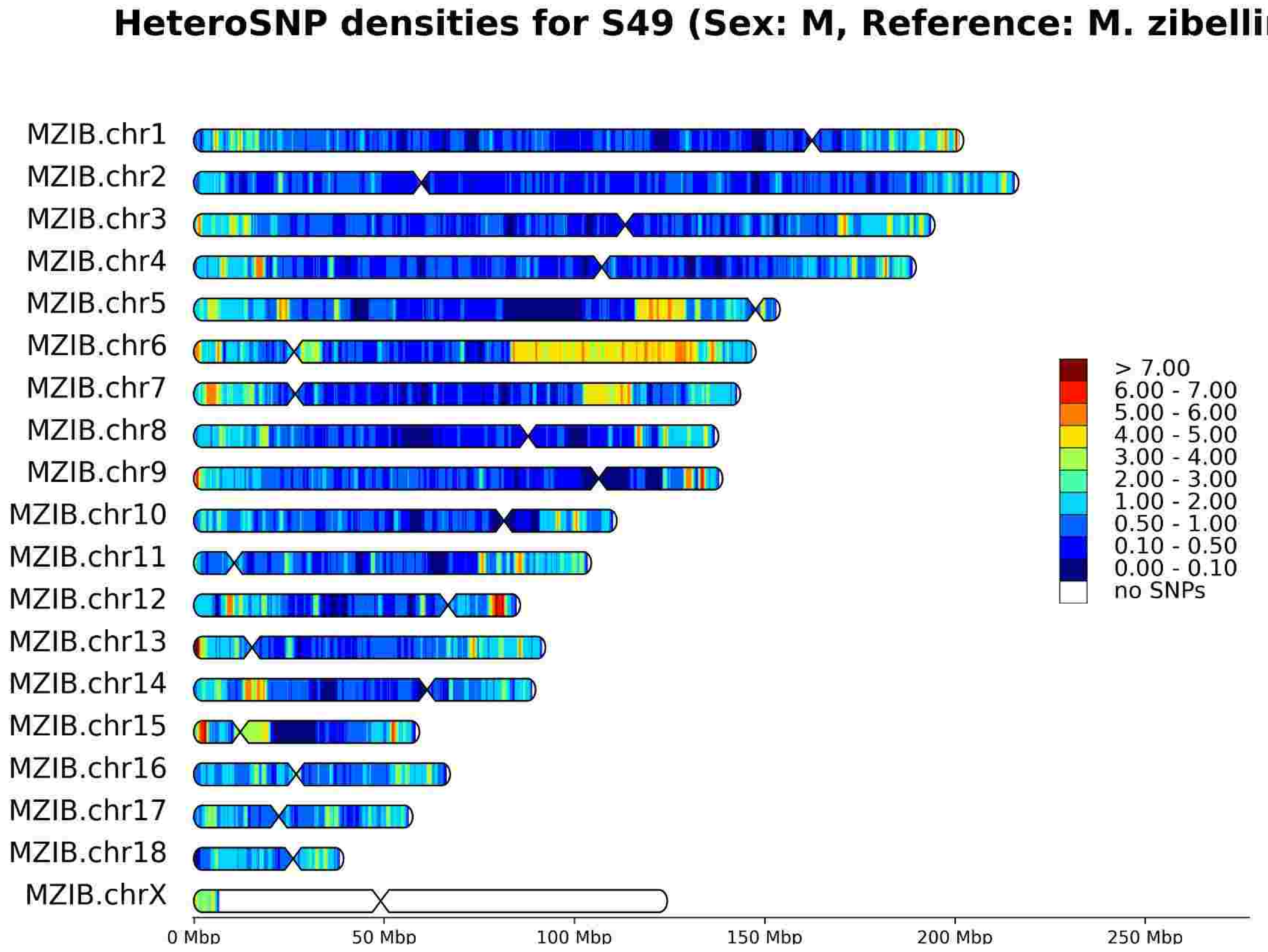
HeteroSNP densities for S49 (Sex: M, Reference: M. zibellina)
MZIB.chr1
MZIB.chr2
MZIB.chr3
MZIB.chr4
MZIB.chr5
MZIB.chr6
MZIB.chr7
MZIB.chr8
MZIB.chr9
MZIB.chr10
MZIB.chr11
MZIB.chr12
MZIB.chr13
MZIB.chr14
MZIB.chr15
MZIB.chr16
MZIB.chr17
MZIB.chr18
MZIB.chrX
> 7.00
6.00 - 7.00
5.00 - 6.00
4.00 - 5.00
3.00 - 4.00
2.00 - 3.00
1.00 - 2.00
0.50 - 1.00
0.10 - 0.50
0.00 - 0.10
no SNPs
0 Mbp
50 Mbp
100 Mbp
150 Mbp
200 Mbp
250 Mbp

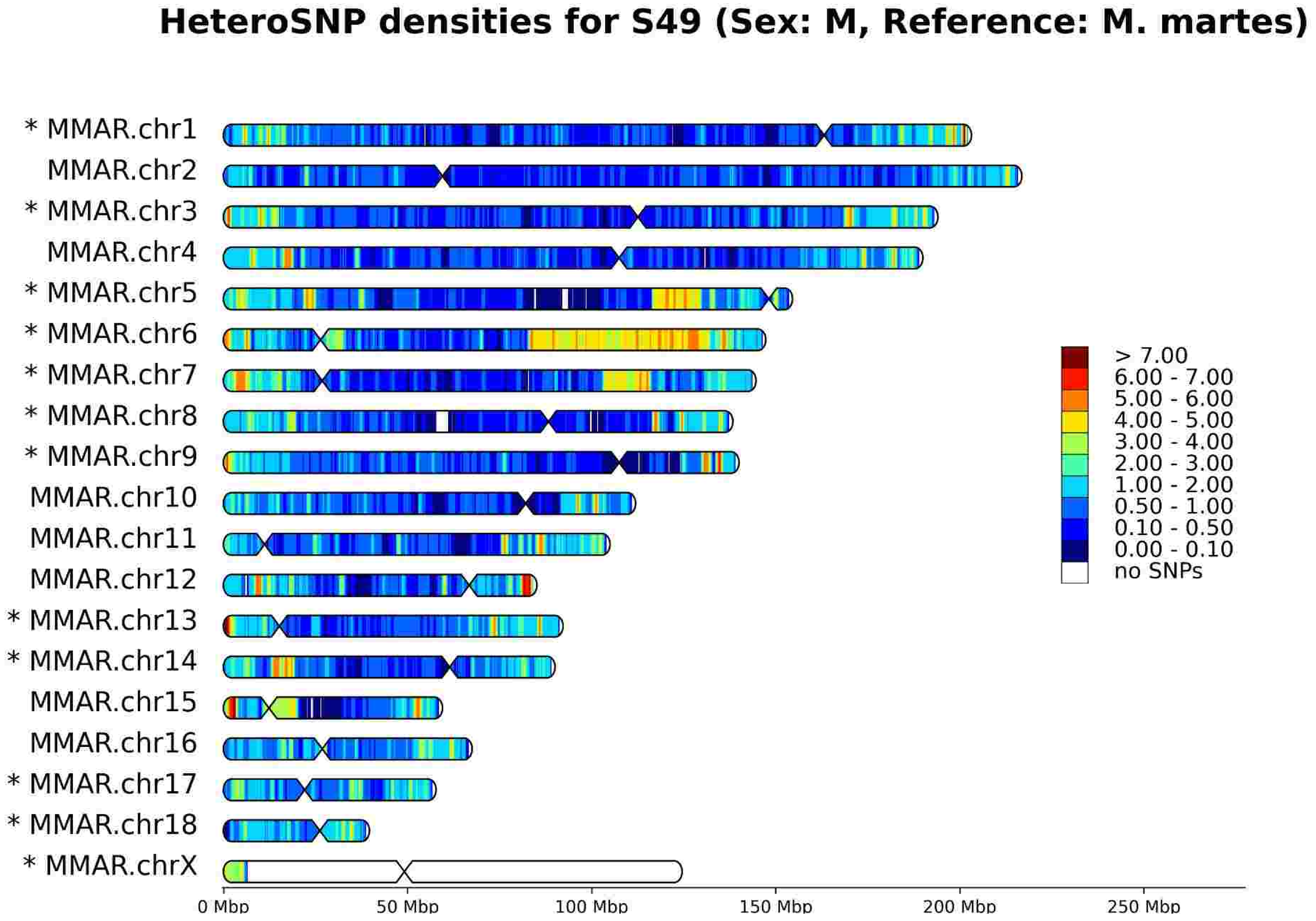
HeteroSNP densities for S49 (Sex: M, Reference: M. martes)
* MMAR.chr1
MMAR.chr2
* MMAR.chr3
MMAR.chr4
* MMAR.chr5
* MMAR.chr6
* MMAR.chr7
* MMAR.chr8
* MMAR.chr9
MMAR.chr10
MMAR.chr11
MMAR.chr12
* MMAR.chr13
* MMAR.chr14
MMAR.chr15
MMAR.chr16
* MMAR.chr17
* MMAR.chr18
* MMAR.chrX
> 7.00
6.00 - 7.00
5.00 - 6.00
4.00 - 5.00
3.00 - 4.00
2.00 - 3.00
1.00 - 2.00
0.50 - 1.00
0.10 - 0.50
0.00 - 0.10
no SNPs
0 Mbp
50 Mbp
100 Mbp
150 Mbp
200 Mbp
250 Mbp

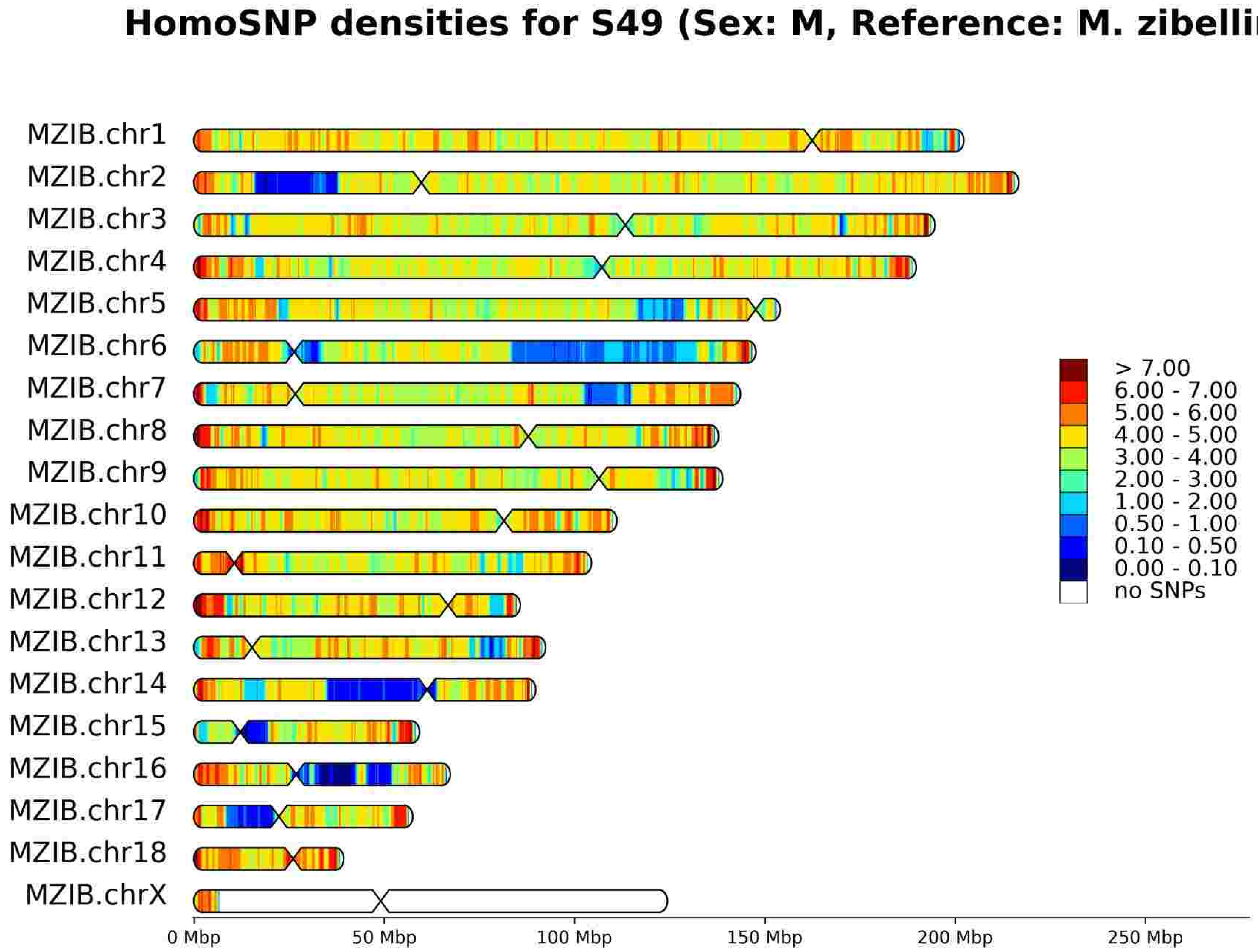
HomoSNP densities for S49 (Sex: M, Reference: M. zibellina)
MZIB.chr1
MZIB.chr2
MZIB.chr3
MZIB.chr4
MZIB.chr5
MZIB.chr6
MZIB.chr7
MZIB.chr8
MZIB.chr9
MZIB.chr10
MZIB.chr11
MZIB.chr12
MZIB.chr13
MZIB.chr14
MZIB.chr15
MZIB.chr16
MZIB.chr17
MZIB.chr18
MZIB.chrX
> 7.00
6.00 - 7.00
5.00 - 6.00
4.00 - 5.00
3.00 - 4.00
2.00 - 3.00
1.00 - 2.00
0.50 - 1.00
0.10 - 0.50
0.00 - 0.10
no SNPs
0 Mbp
50 Mbp
100 Mbp
150 Mbp
200 Mbp
250 Mbp

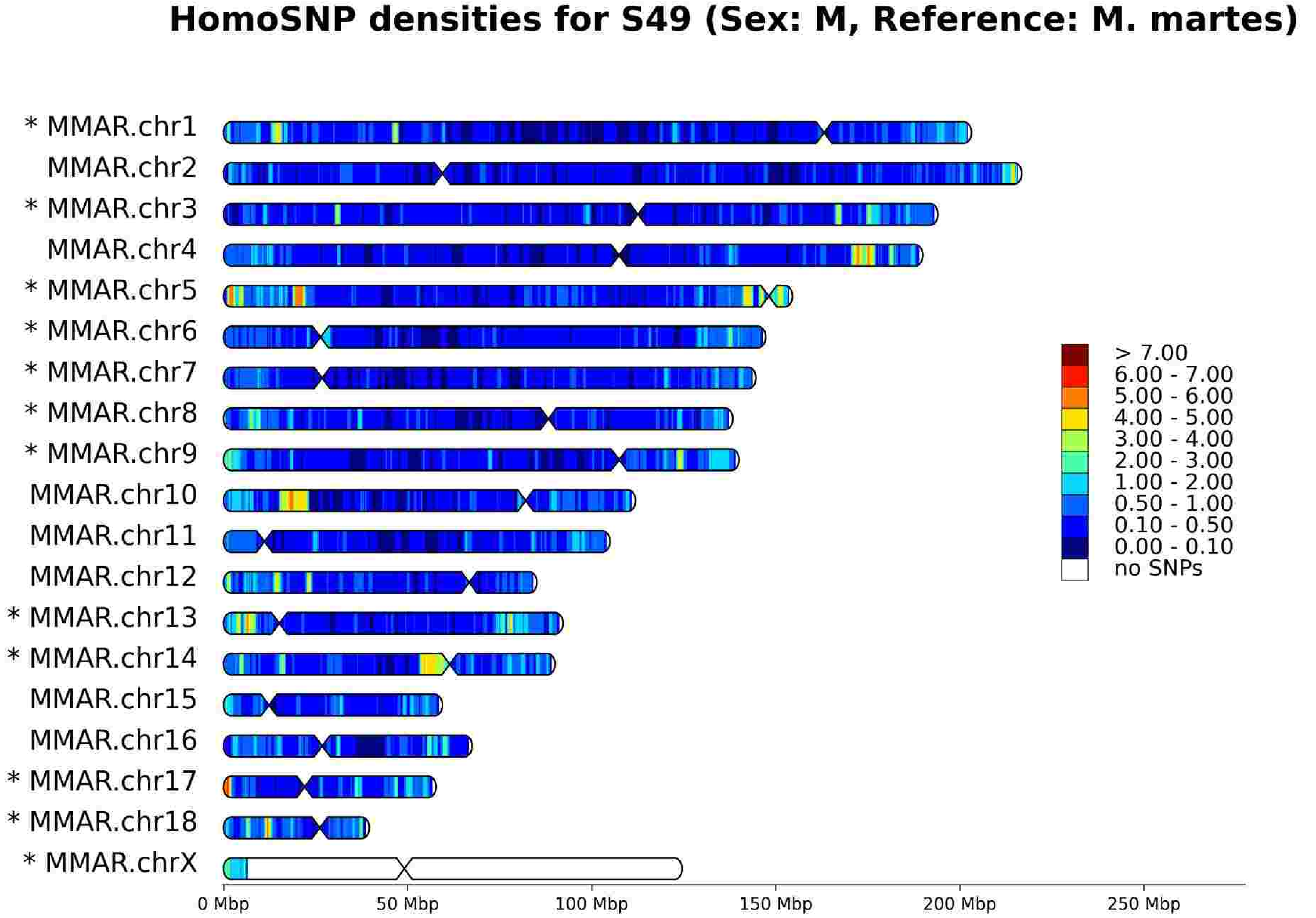
HomoSNP densities for S49 (Sex: M, Reference: M. martes)
* MMAR.chr1
MMAR.chr2
* MMAR.chr3
MMAR.chr4
* MMAR.chr5
* MMAR.chr6
* MMAR.chr7
* MMAR.chr8
* MMAR.chr9
MMAR.chr10
MMAR.chr11
MMAR.chr12
* MMAR.chr13
* MMAR.chr14
MMAR.chr15
MMAR.chr16
* MMAR.chr17
* MMAR.chr18
* MMAR.chrX
> 7.00
6.00 - 7.00
5.00 - 6.00
4.00 - 5.00
3.00 - 4.00
2.00 - 3.00
1.00 - 2.00
0.50 - 1.00
0.10 - 0.50
0.00 - 0.10
no SNPs
0 Mbp
50 Mbp
100 Mbp
150 Mbp
200 Mbp
250 Mbp

# S50

HeteroSNP densities for S50 (Sex: F, Reference: M. zibellina)

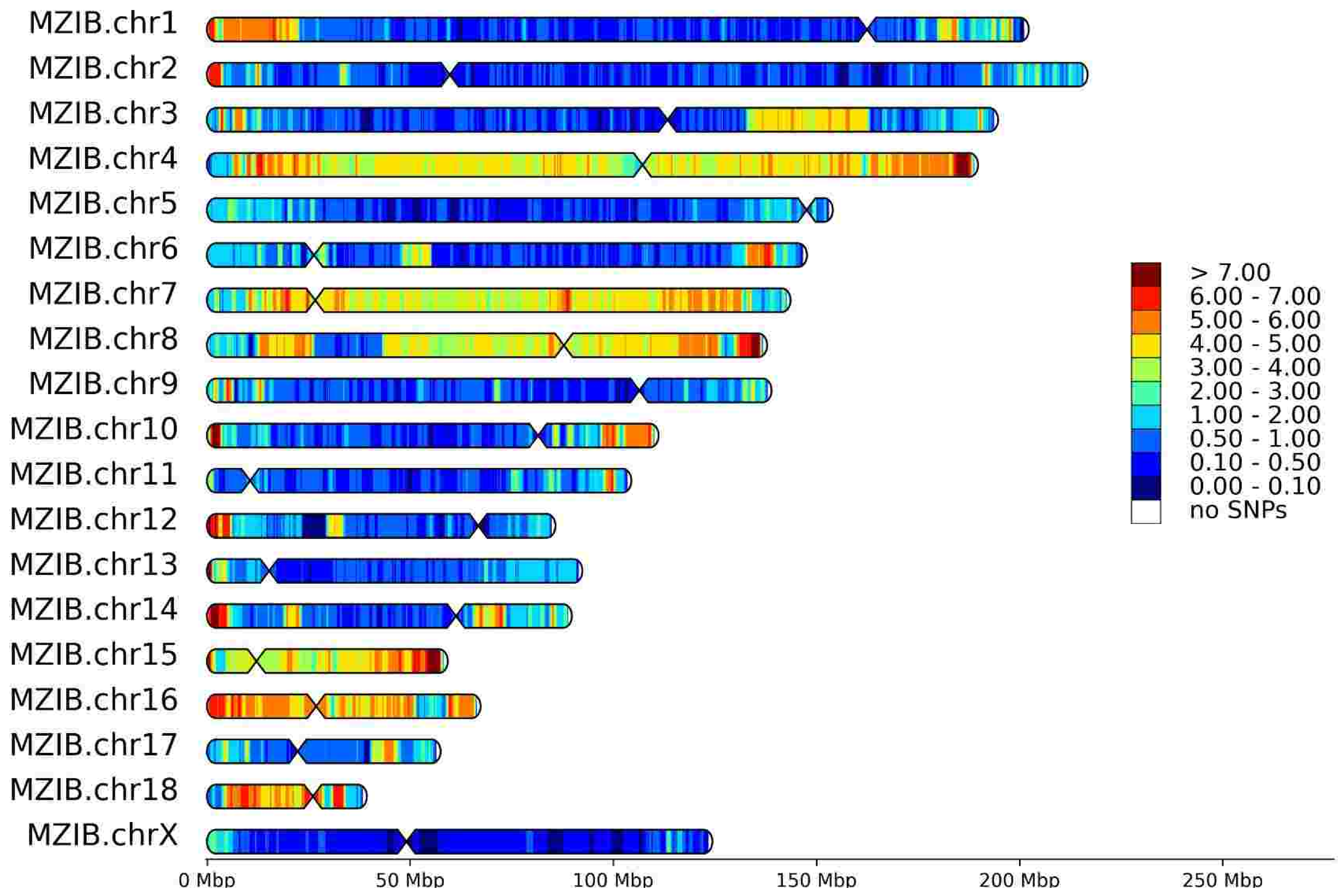


HeteroSNP densities for S50 (Sex: F, Reference: M. martes)

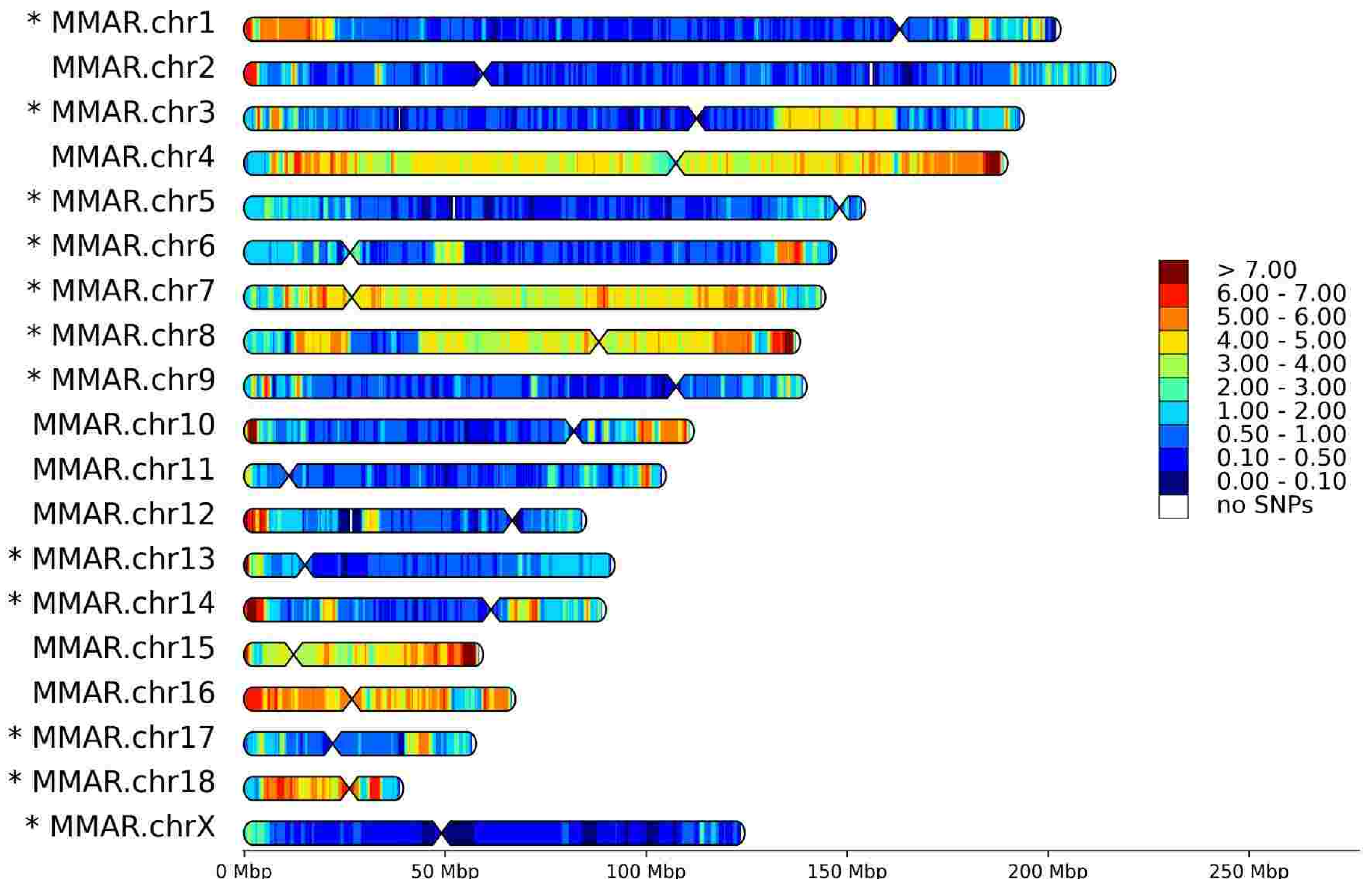


HomoSNP densities for S50 (Sex: F, Reference: M. zibellina)

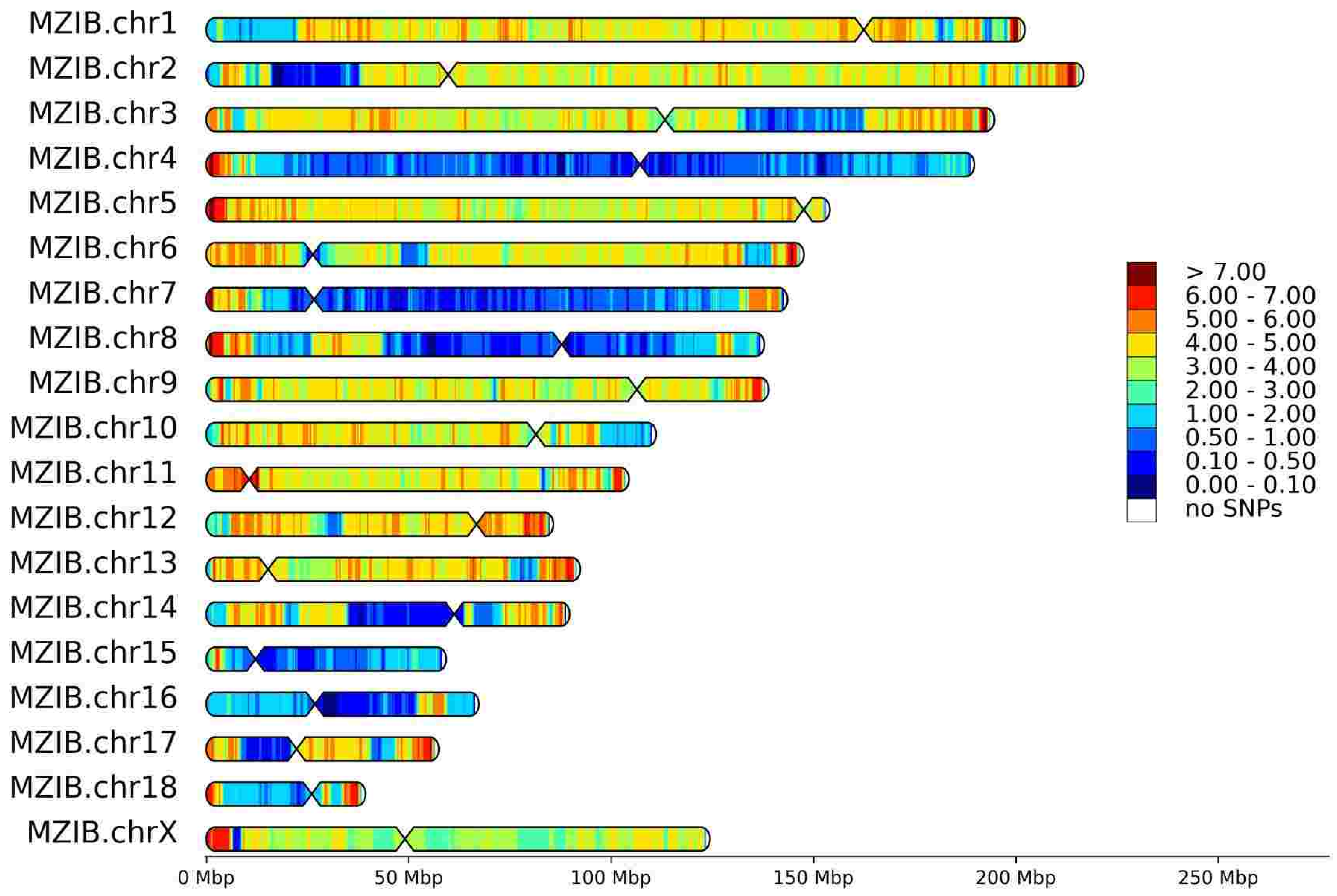


HomoSNP densities for S50 (Sex: F, Reference: M. martes)

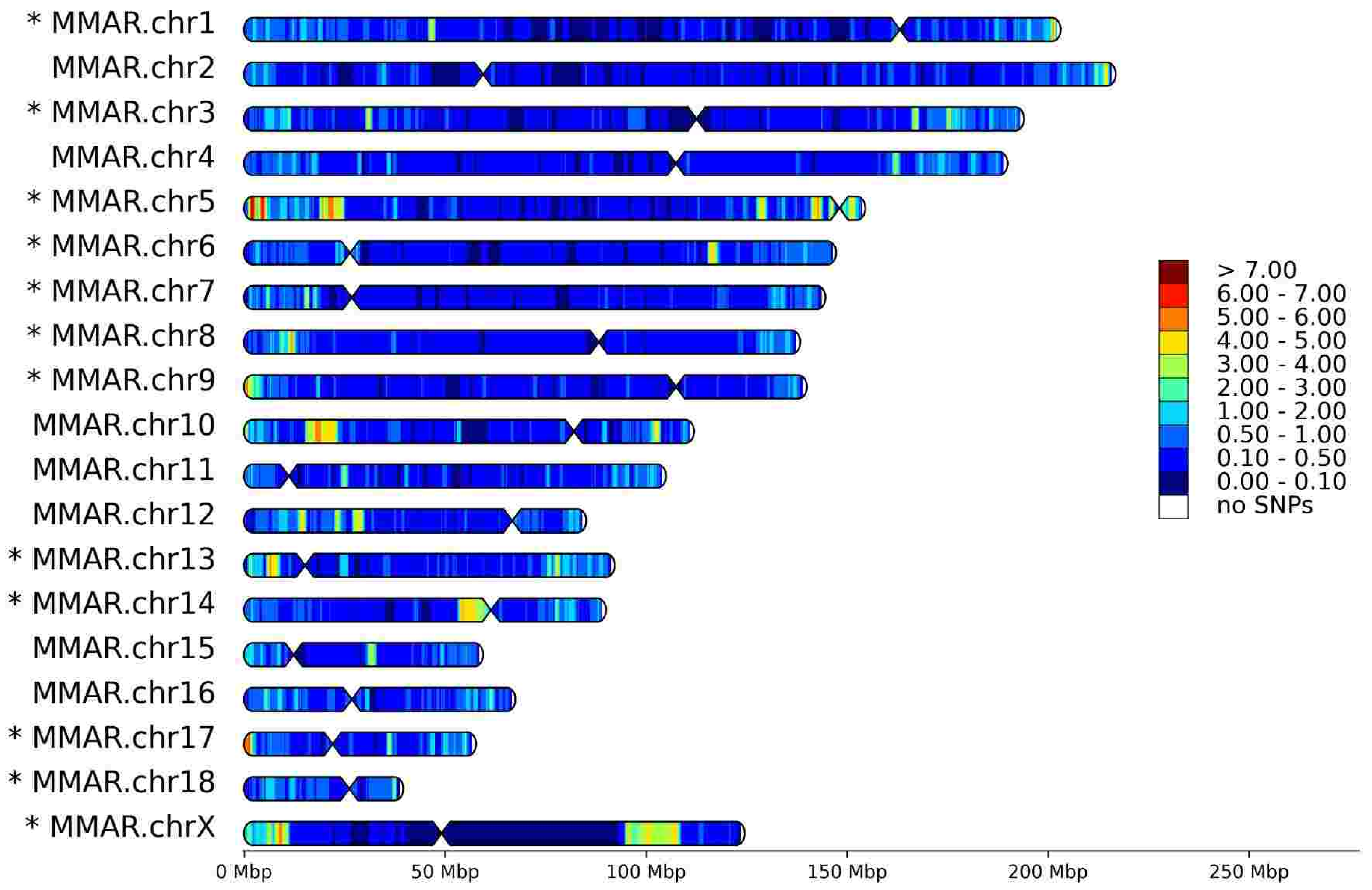

# T8

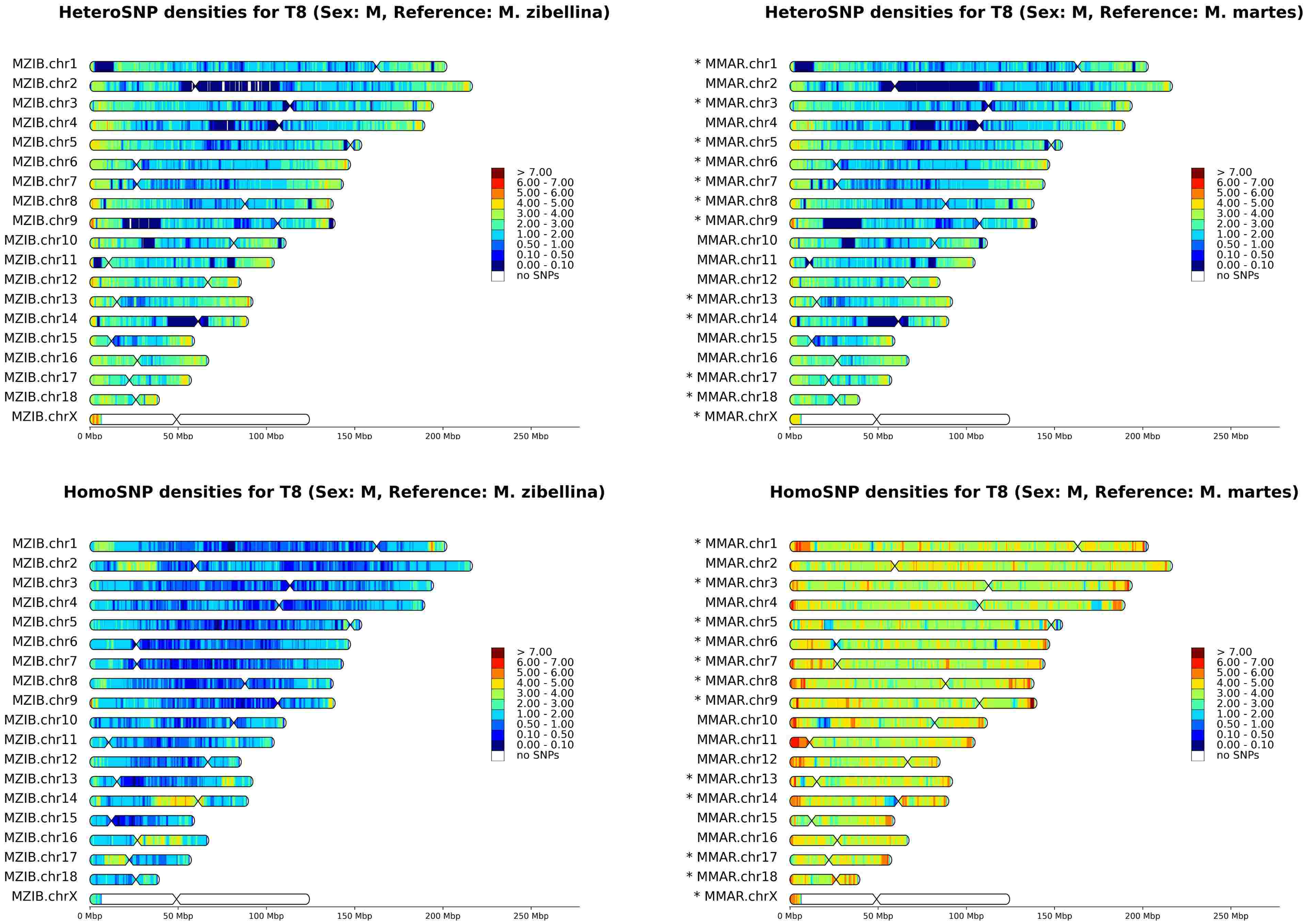

HeteroSNP densities for T8 (Sex: M, Reference: M. zibellina)
HeteroSNP densities for T8 (Sex: M, Reference: M. martes)
HomoSNP densities for T8 (Sex: M, Reference: M. zibellina)
HomoSNP densities for T8 (Sex: M, Reference: M. martes)
MZIB.chr1
MZIB.chr2
MZIB.chr3
MZIB.chr4
MZIB.chr5
MZIB.chr6
MZIB.chr7
MZIB.chr8
MZIB.chr9
MZIB.chr10
MZIB.chr11
MZIB.chr12
MZIB.chr13
MZIB.chr14
MZIB.chr15
MZIB.chr16
MZIB.chr17
MZIB.chr18
MZIB.chrX
* MMAR.chr1
MMAR.chr2
* MMAR.chr3
MMAR.chr4
* MMAR.chr5
* MMAR.chr6
* MMAR.chr7
* MMAR.chr8
* MMAR.chr9
MMAR.chr10
MMAR.chr11
MMAR.chr12
* MMAR.chr13
* MMAR.chr14
MMAR.chr15
MMAR.chr16
* MMAR.chr17
* MMAR.chr18
* MMAR.chrX
> 7.00
6.00 - 7.00
5.00 - 6.00
4.00 - 5.00
3.00 - 4.00
2.00 - 3.00
1.00 - 2.00
0.50 - 1.00
0.10 - 0.50
0.00 - 0.10
no SNPs
0 Mbp
50 Mbp
100 Mbp
150 Mbp
200 Mbp
250 Mbp

# T18

HeteroSNP densities for T18 (Sex: M, Reference: M. zibellina)

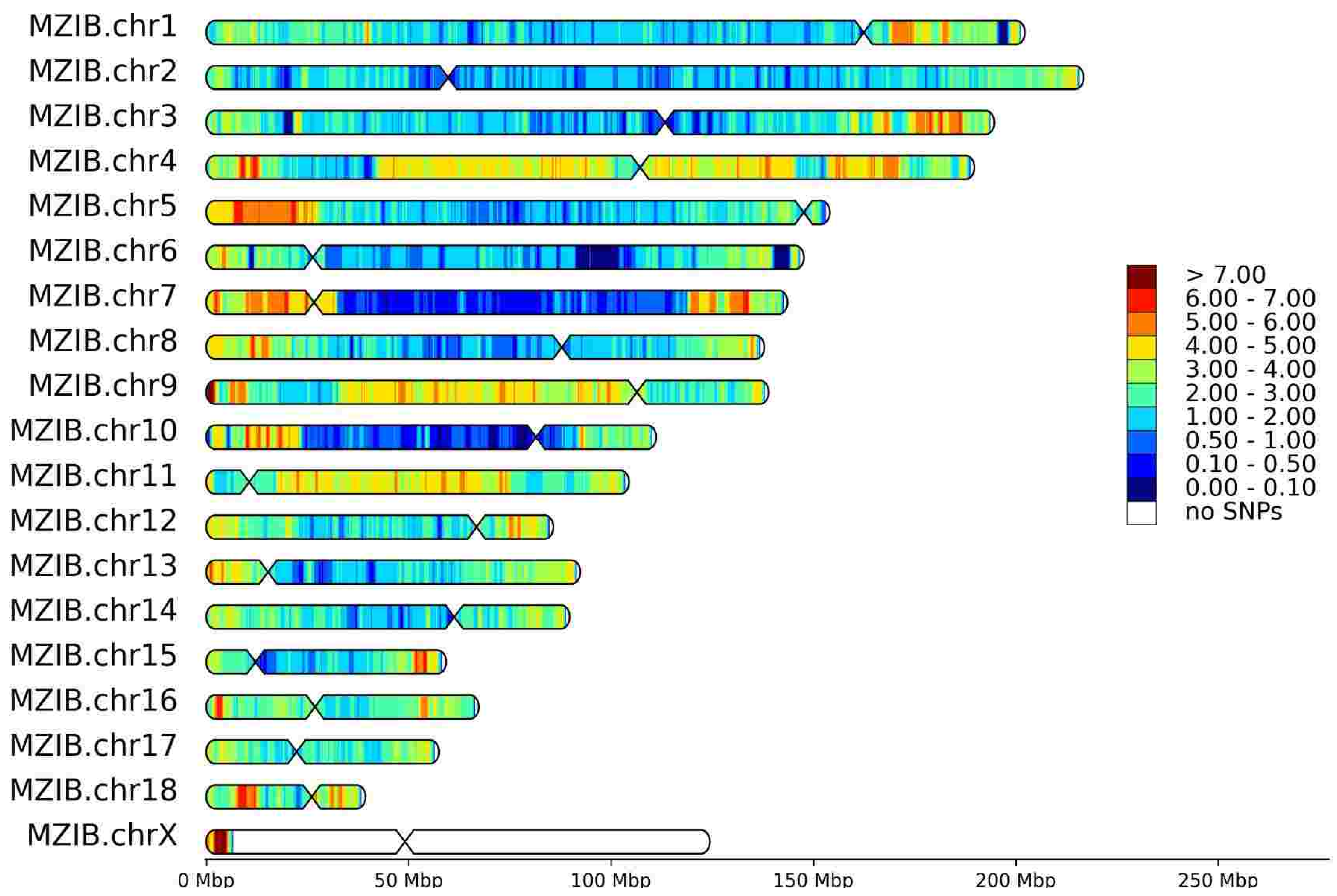


HeteroSNP densities for T18 (Sex: M, Reference: M. martes)

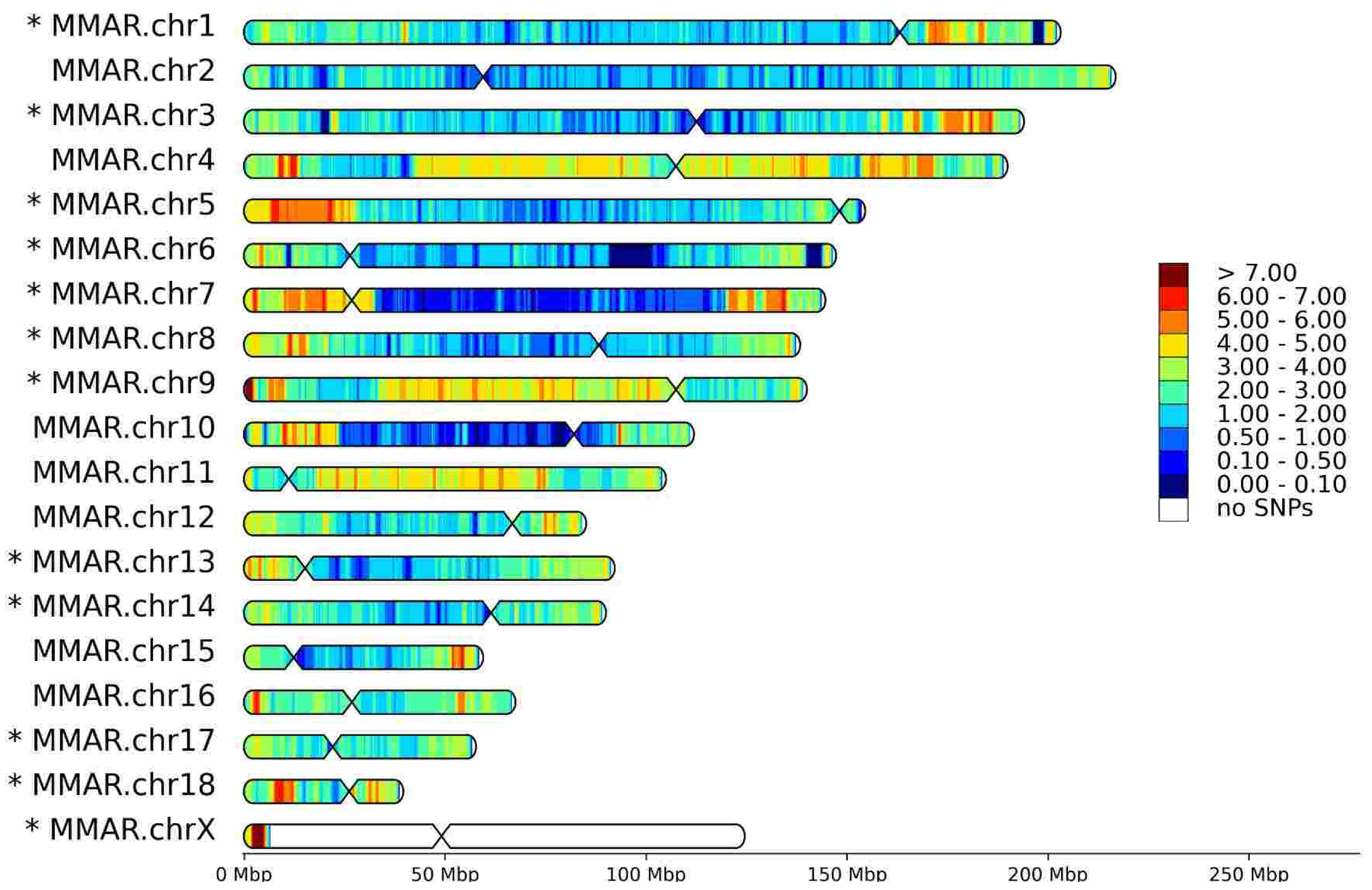


HomoSNP densities for T18 (Sex: M, Reference: M. zibellina)

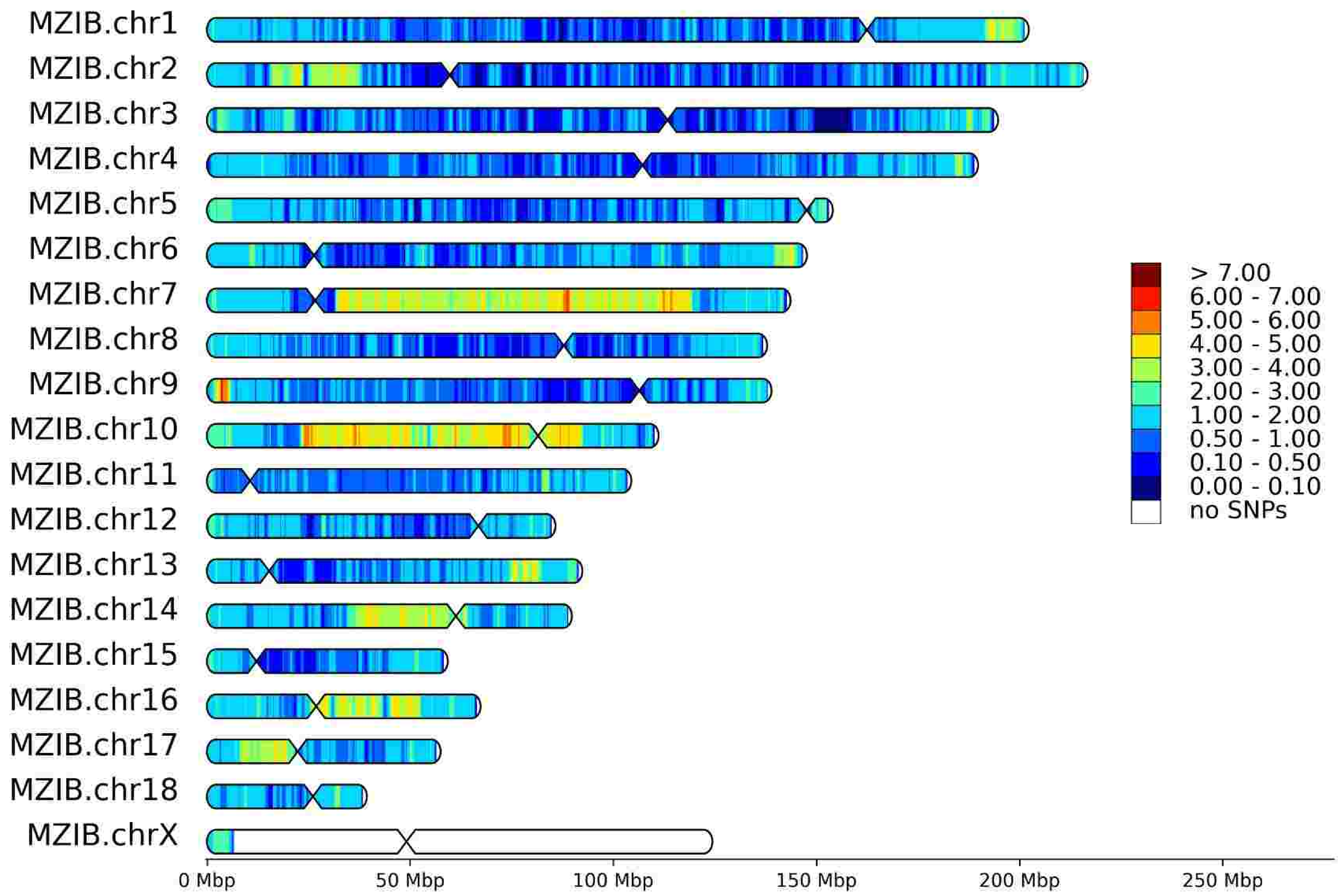


HomoSNP densities for T18 (Sex: M, Reference: M. martes)

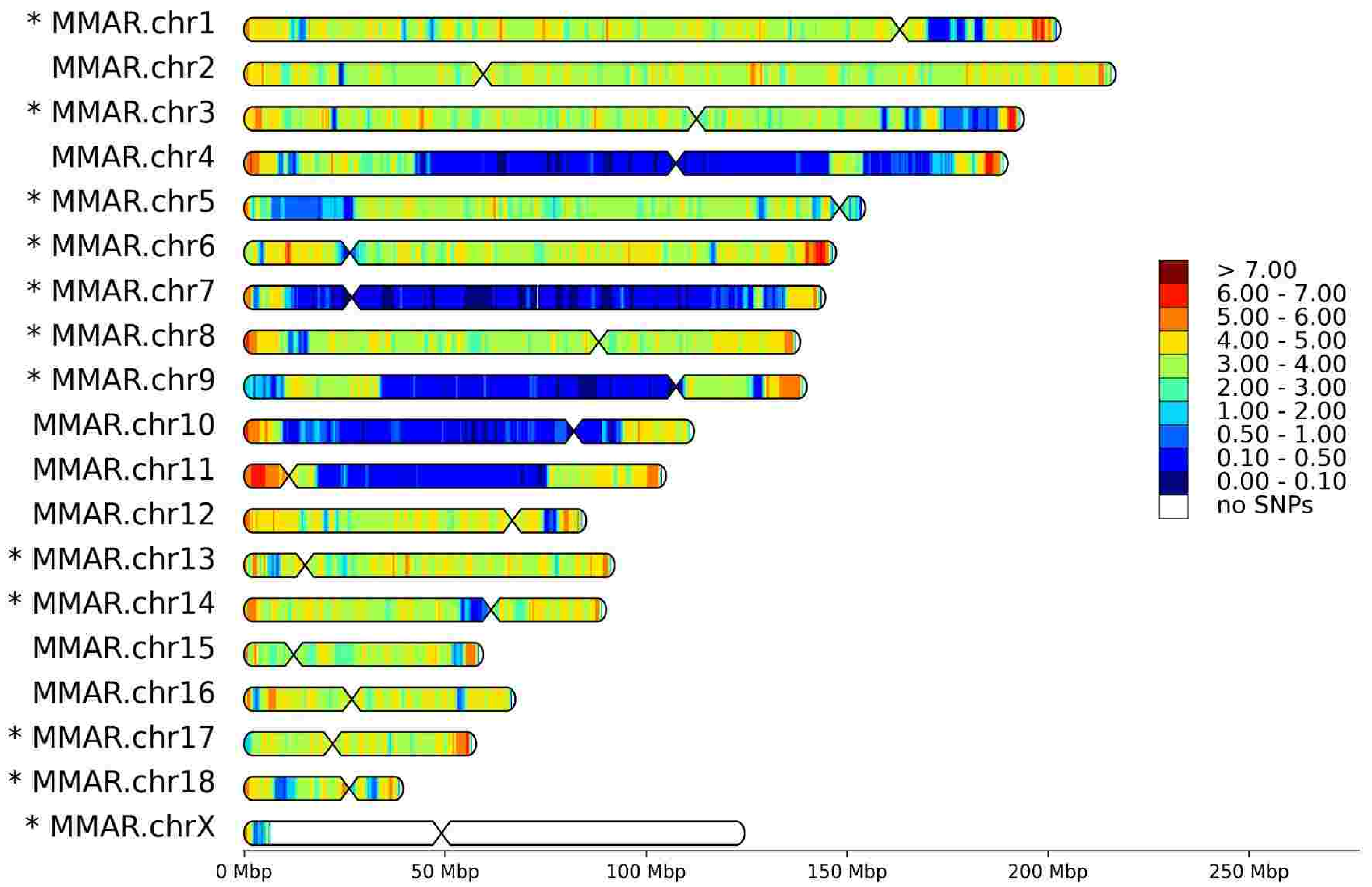

# T24

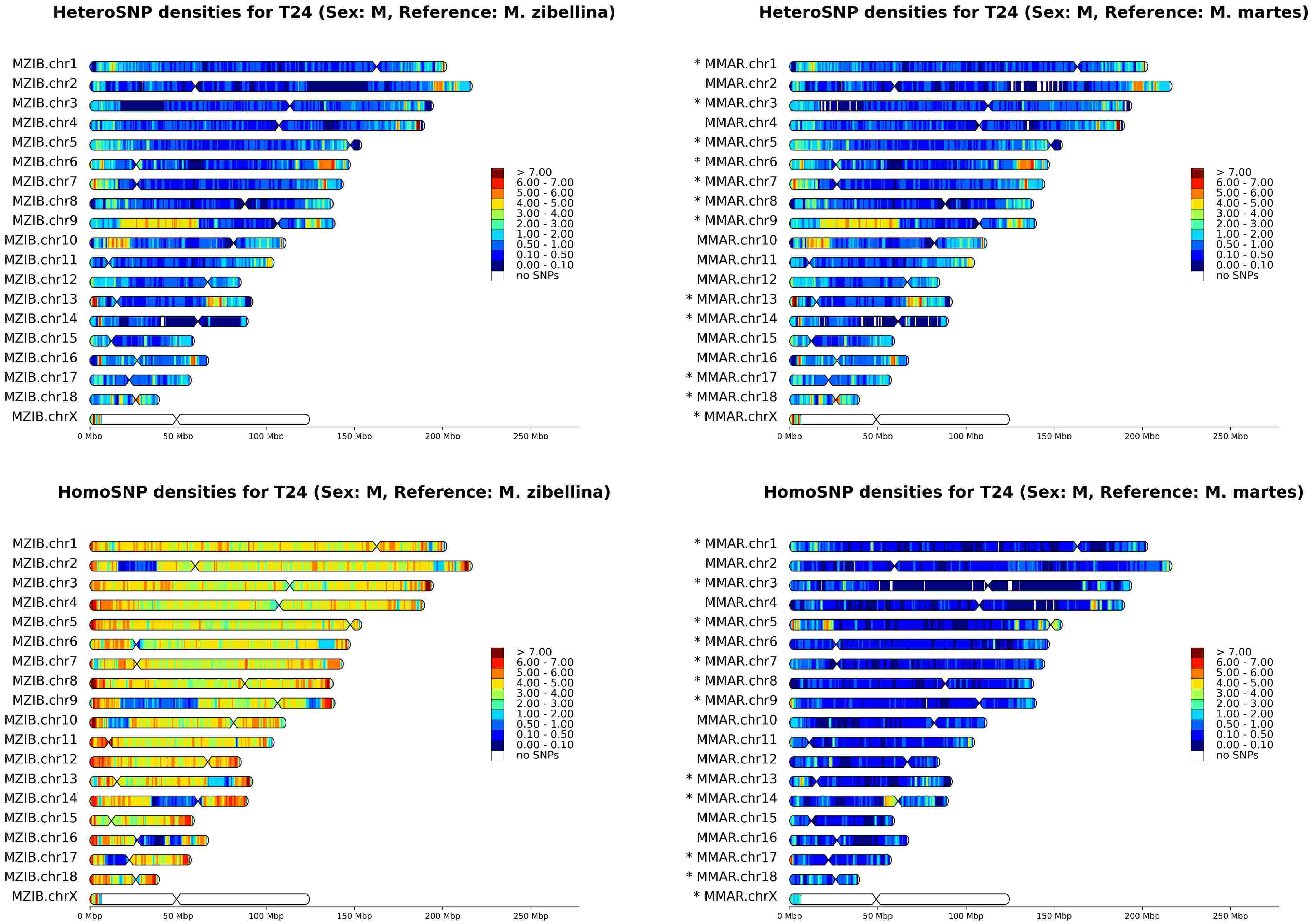
HeteroSNP densities for T24 (Sex: M, Reference: M. zibellina)
MZIB.chr1
MZIB.chr2
MZIB.chr3
MZIB.chr4
MZIB.chr5
MZIB.chr6
MZIB.chr7
MZIB.chr8
MZIB.chr9
MZIB.chr10
MZIB.chr11
MZIB.chr12
MZIB.chr13
MZIB.chr14
MZIB.chr15
MZIB.chr16
MZIB.chr17
MZIB.chr18
MZIB.chrX
0 Mbp
50 Mbp
100 Mbp
150 Mbp
200 Mbp
250 Mbp
> 7.00
6.00 - 7.00
5.00 - 6.00
4.00 - 5.00
3.00 - 4.00
2.00 - 3.00
1.00 - 2.00
0.50 - 1.00
0.10 - 0.50
0.00 - 0.10
no SNPs
HeteroSNP densities for T24 (Sex: M, Reference: M. martes)
* MMAR.chr1
MMAR.chr2
* MMAR.chr3
MMAR.chr4
* MMAR.chr5
* MMAR.chr6
* MMAR.chr7
* MMAR.chr8
* MMAR.chr9
MMAR.chr10
MMAR.chr11
MMAR.chr12
* MMAR.chr13
* MMAR.chr14
MMAR.chr15
MMAR.chr16
* MMAR.chr17
* MMAR.chr18
* MMAR.chrX
HomoSNP densities for T24 (Sex: M, Reference: M. zibellina)
HomoSNP densities for T24 (Sex: M, Reference: M. martes)

# T26

HeteroSNP densities for T26 (Sex: F, Reference: M. zibellina)

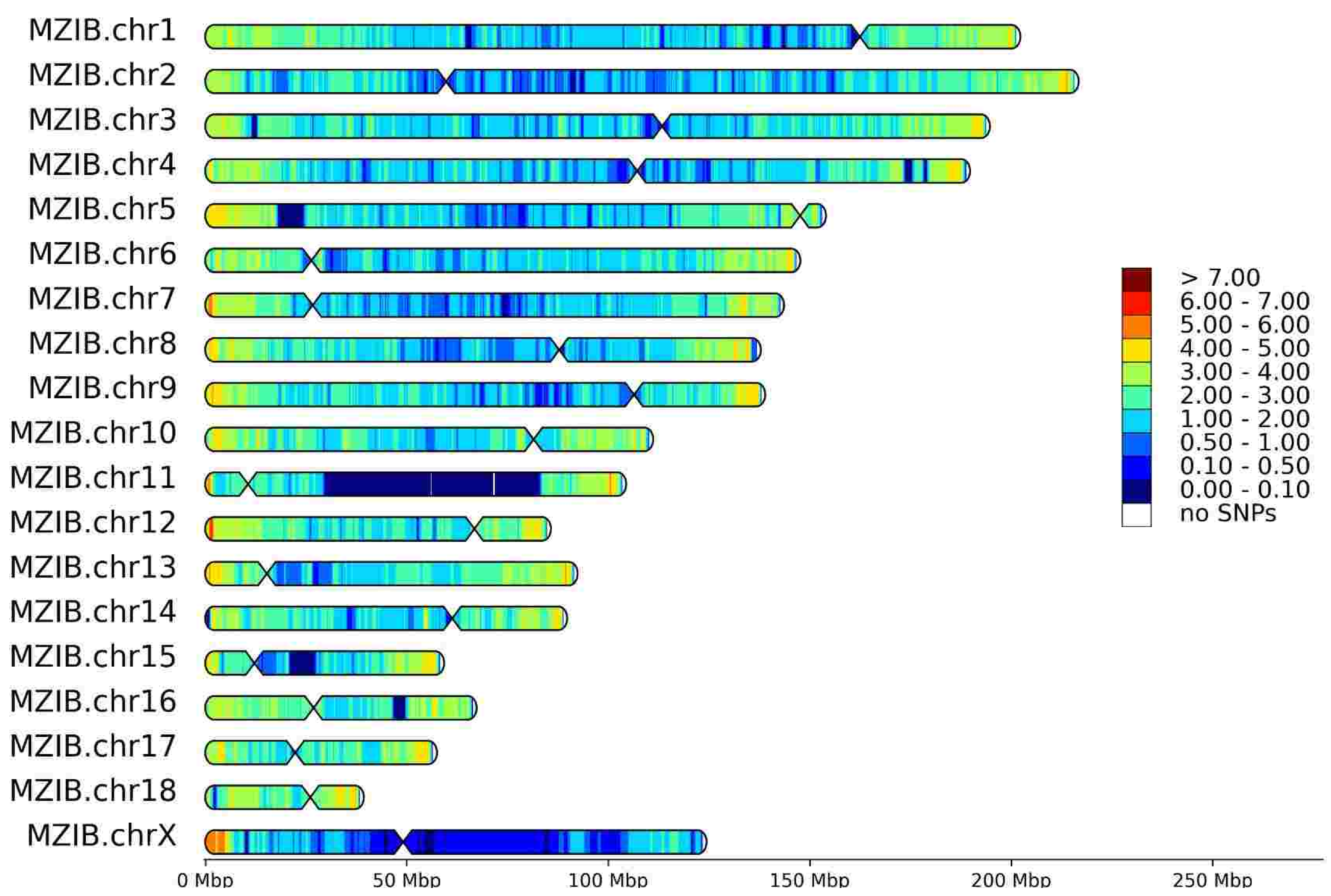


HeteroSNP densities for T26 (Sex: F, Reference: M. martes)

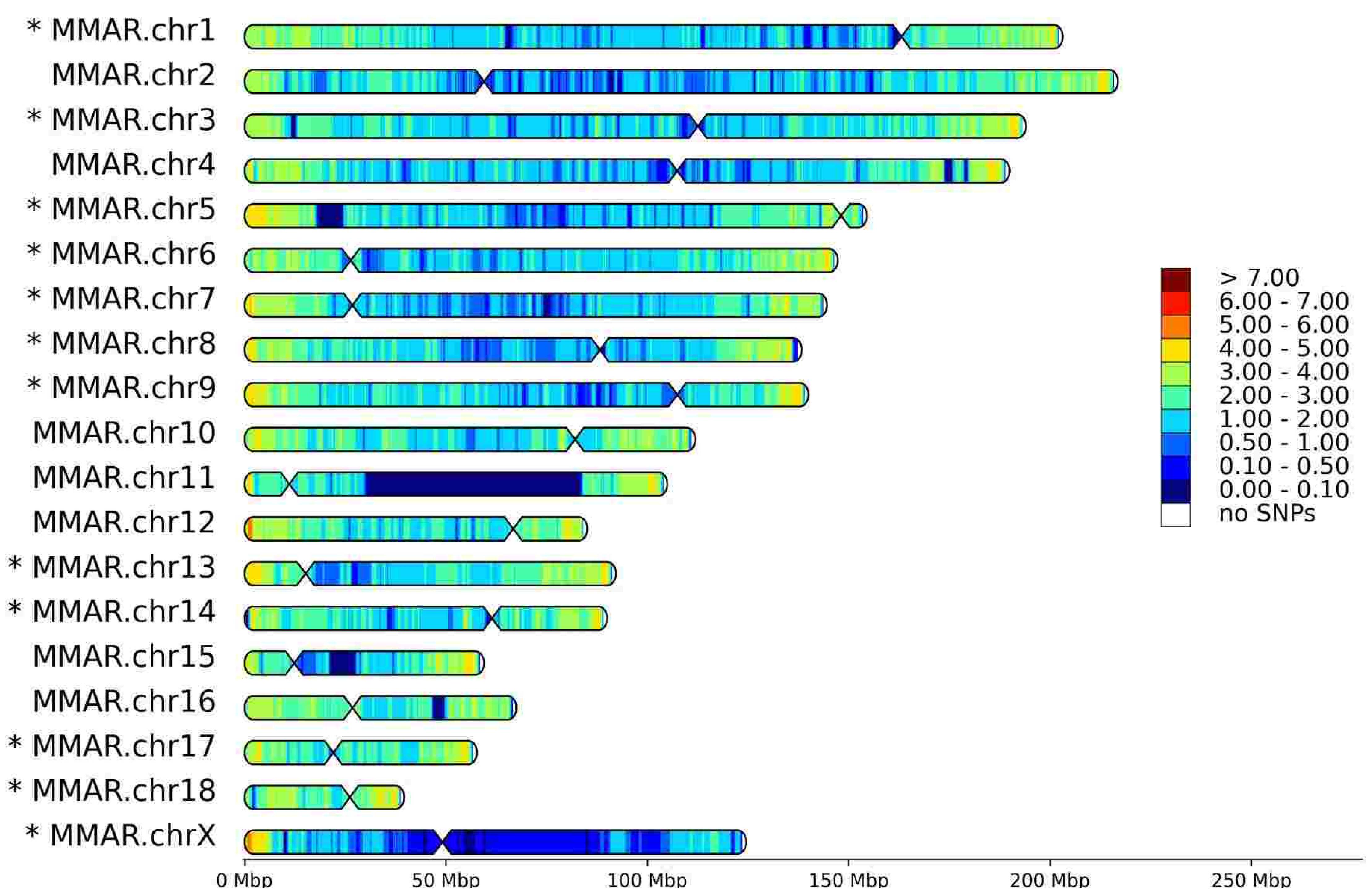


HomoSNP densities for T26 (Sex: F, Reference: M. zibellina)

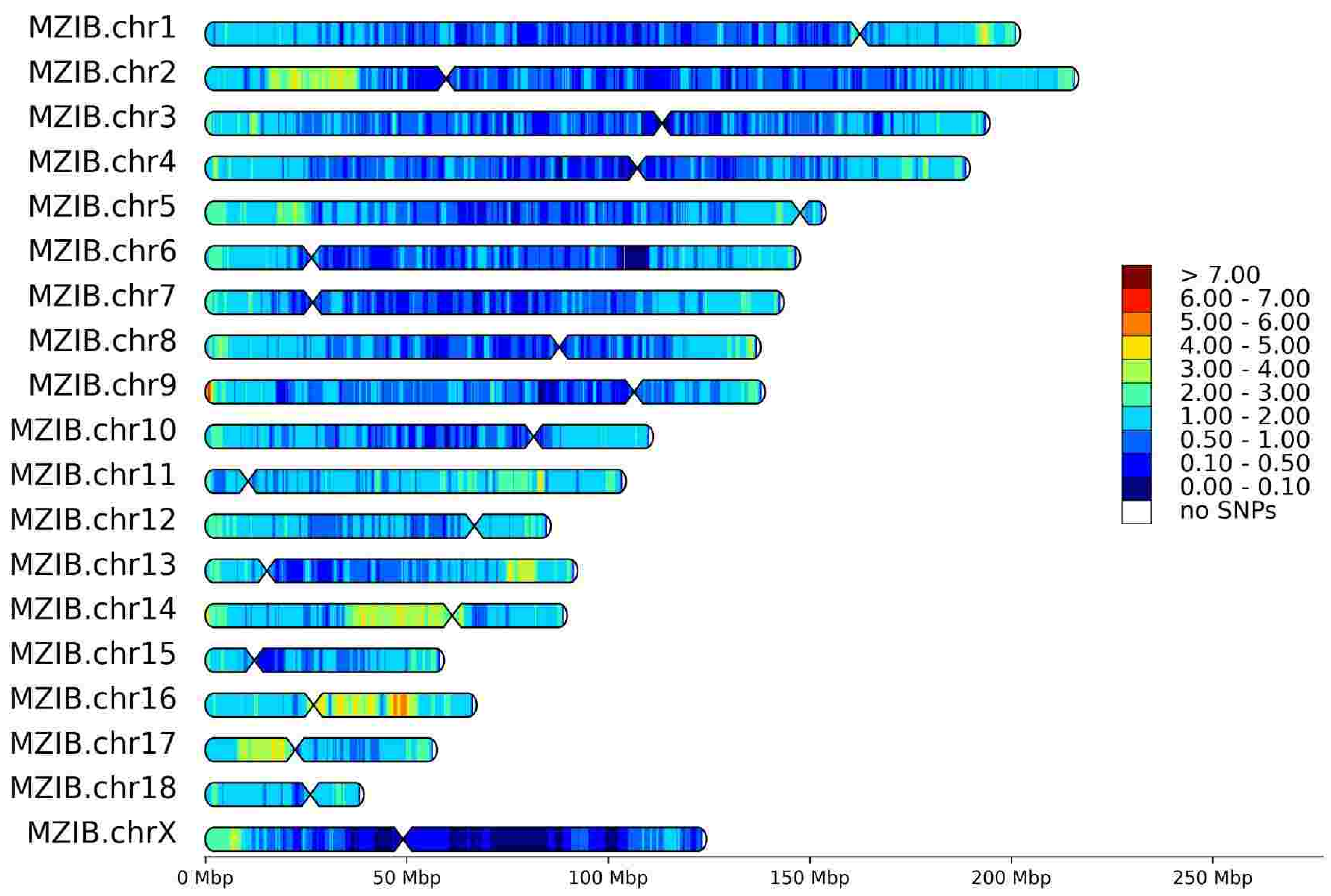


HomoSNP densities for T26 (Sex: F, Reference: M. martes)

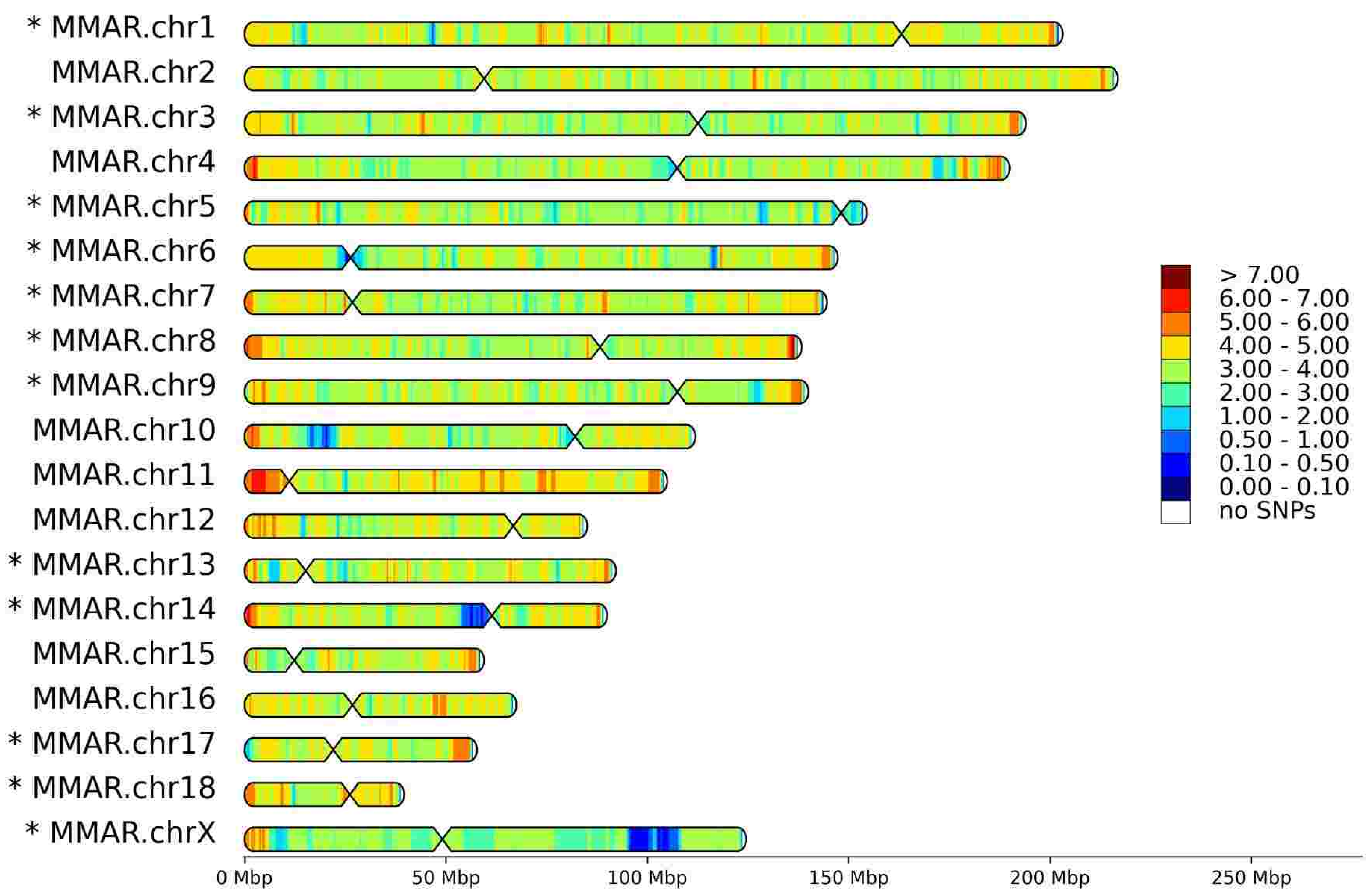

# T50

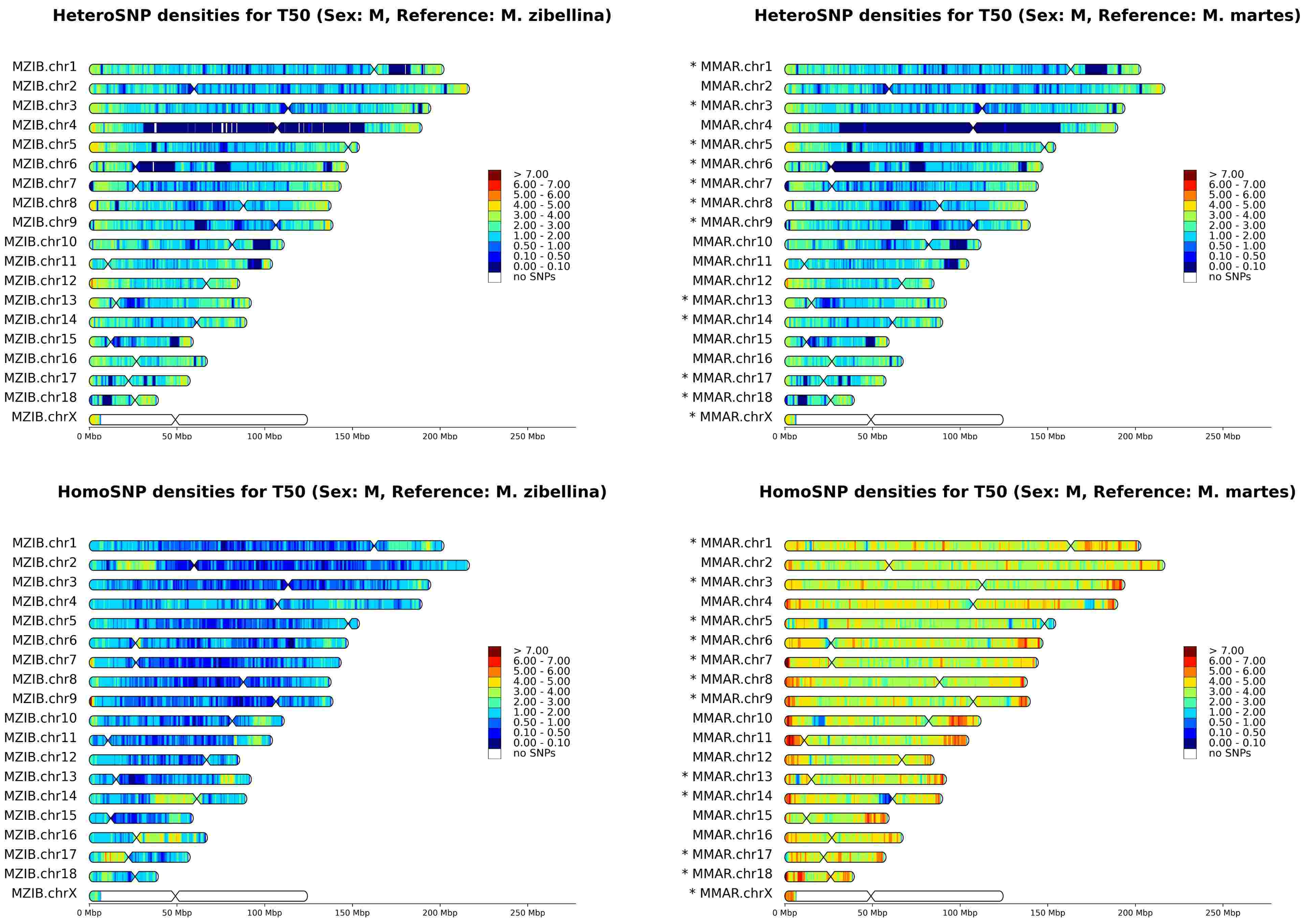
HeteroSNP densities for T50 (Sex: M, Reference: M. zibellina)
HeteroSNP densities for T50 (Sex: M, Reference: M. martes)
HomoSNP densities for T50 (Sex: M, Reference: M. zibellina)
HomoSNP densities for T50 (Sex: M, Reference: M. martes)
MZIB.chr1
MZIB.chr2
MZIB.chr3
MZIB.chr4
MZIB.chr5
MZIB.chr6
MZIB.chr7
MZIB.chr8
MZIB.chr9
MZIB.chr10
MZIB.chr11
MZIB.chr12
MZIB.chr13
MZIB.chr14
MZIB.chr15
MZIB.chr16
MZIB.chr17
MZIB.chr18
MZIB.chrX
* MMAR.chr1
MMAR.chr2
* MMAR.chr3
MMAR.chr4
* MMAR.chr5
* MMAR.chr6
* MMAR.chr7
* MMAR.chr8
* MMAR.chr9
MMAR.chr10
MMAR.chr11
MMAR.chr12
* MMAR.chr13
* MMAR.chr14
MMAR.chr15
MMAR.chr16
* MMAR.chr17
* MMAR.chr18
* MMAR.chrX
> 7.00
6.00 - 7.00
5.00 - 6.00
4.00 - 5.00
3.00 - 4.00
2.00 - 3.00
1.00 - 2.00
0.50 - 1.00
0.10 - 0.50
0.00 - 0.10
no SNPs
0 Mbp
50 Mbp
100 Mbp
150 Mbp
200 Mbp
250 Mbp

# T72

HeteroSNP densities for T72 (Sex: M, Reference: M. zibellina)

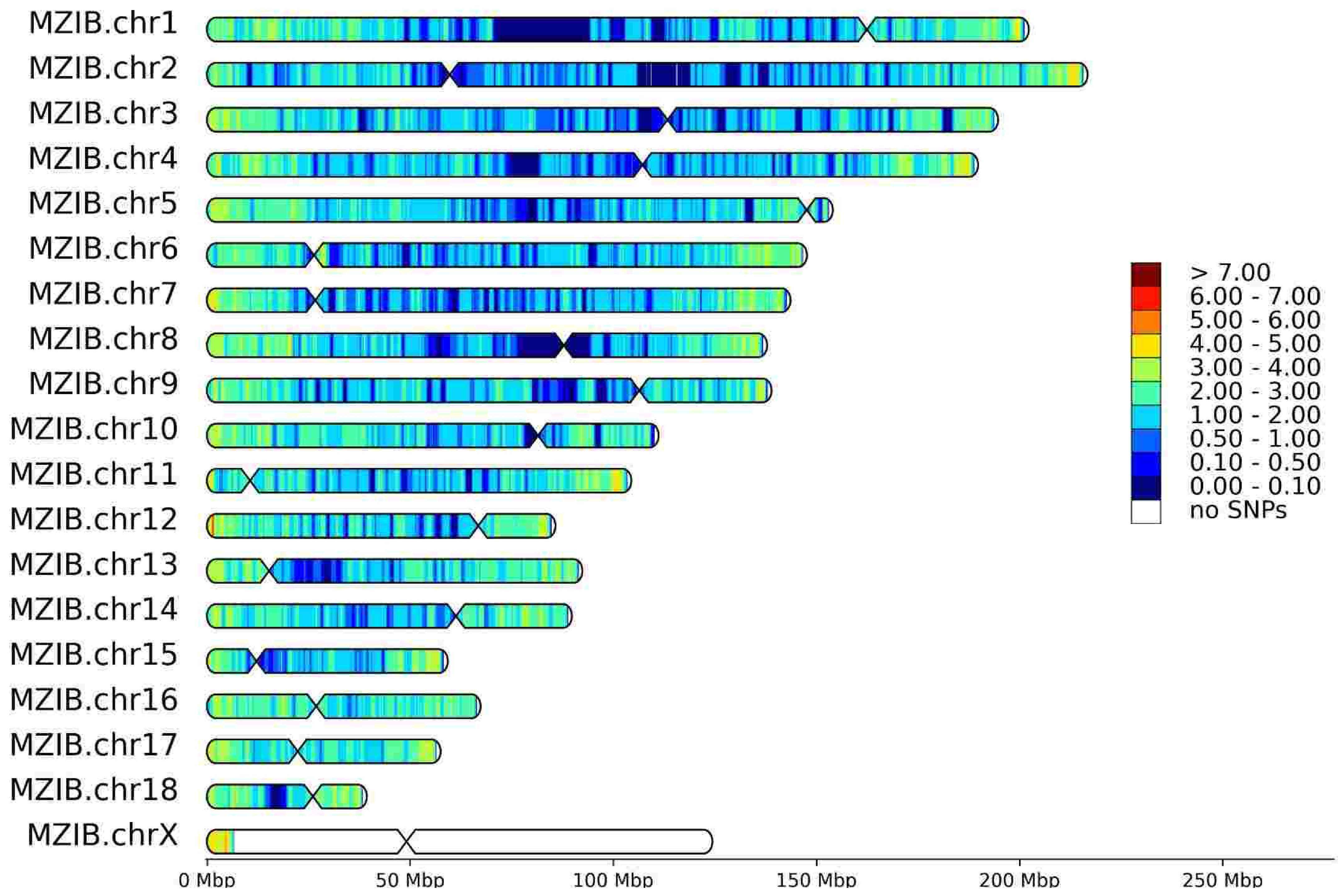


HeteroSNP densities for T72 (Sex: M, Reference: M. martes)

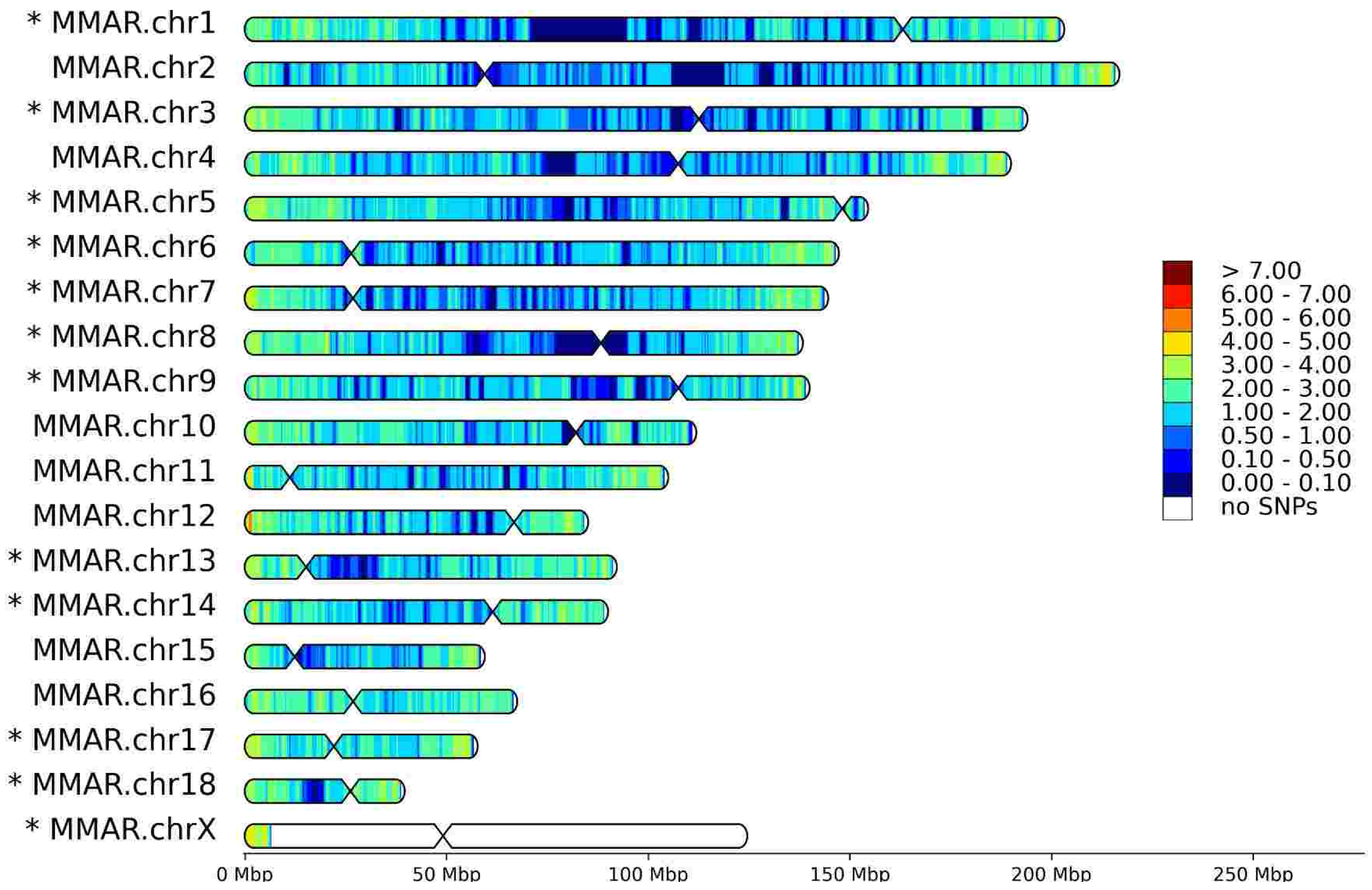


HomoSNP densities for T72 (Sex: M, Reference: M. zibellina)

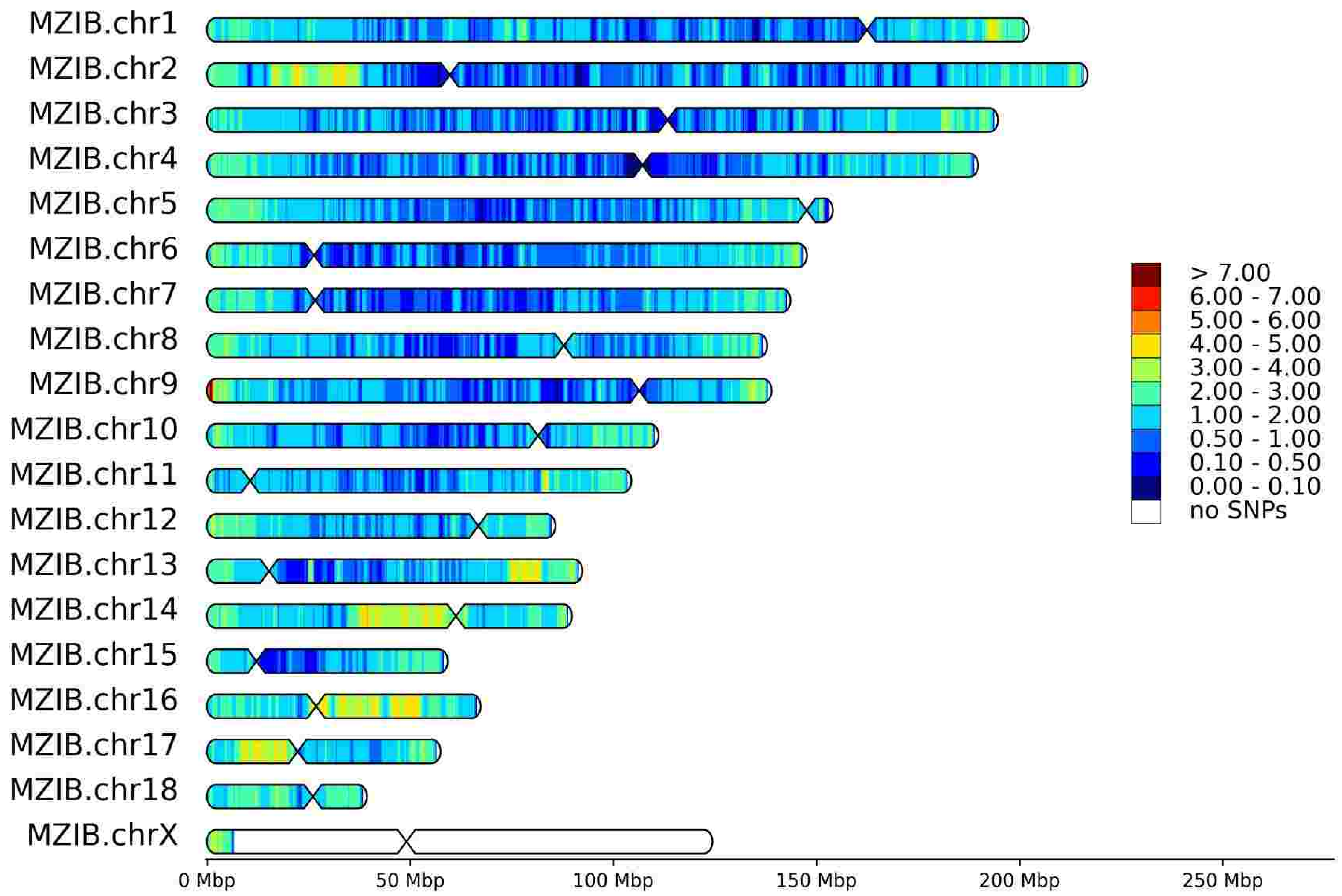


HomoSNP densities for T72 (Sex: M, Reference: M. martes)

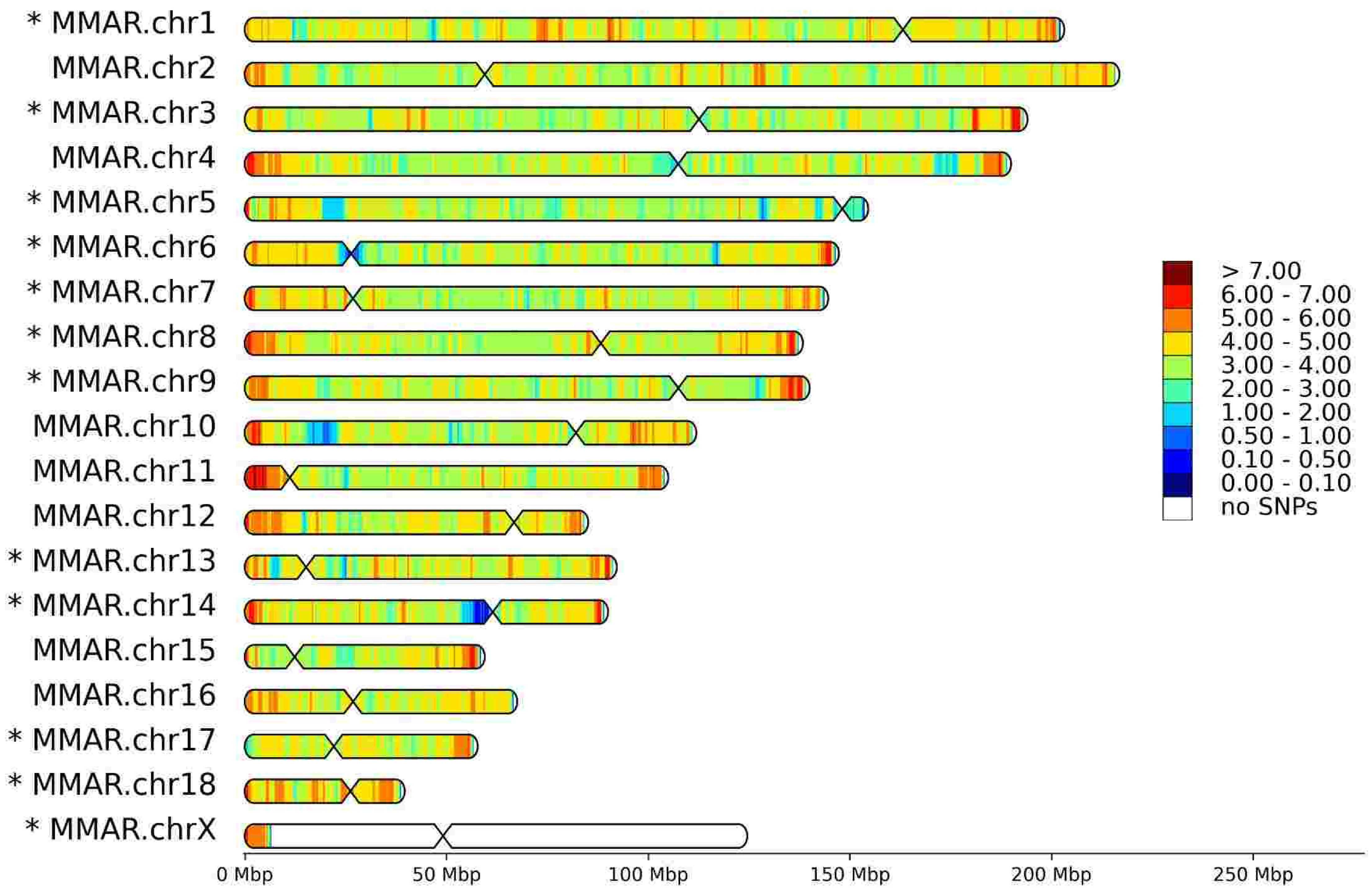

# T76

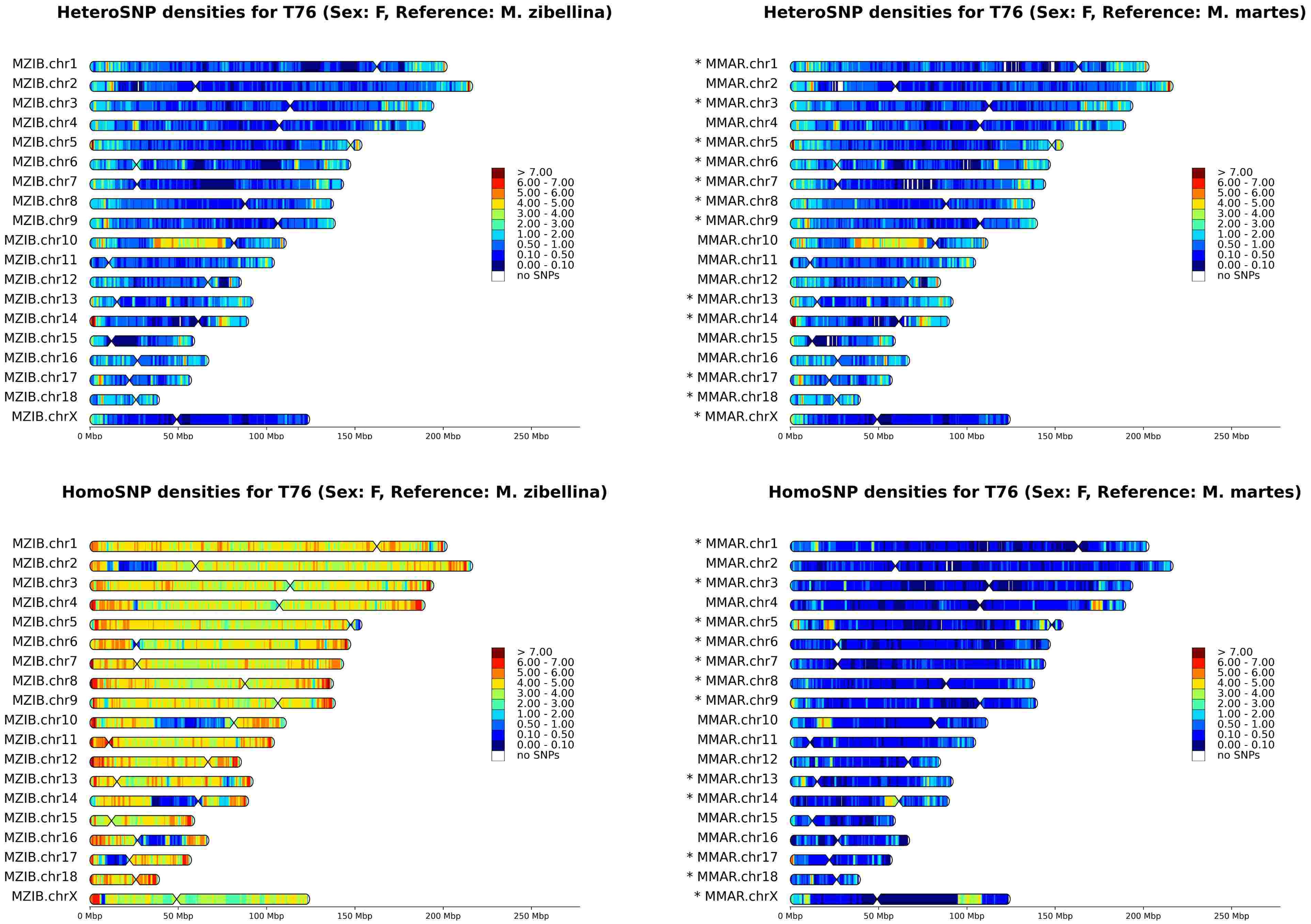
HeteroSNP densities for T76 (Sex: F, Reference: M. zibellina)
HeteroSNP densities for T76 (Sex: F, Reference: M. martes)
HomoSNP densities for T76 (Sex: F, Reference: M. zibellina)
HomoSNP densities for T76 (Sex: F, Reference: M. martes)
MZIB.chr1
MZIB.chr2
MZIB.chr3
MZIB.chr4
MZIB.chr5
MZIB.chr6
MZIB.chr7
MZIB.chr8
MZIB.chr9
MZIB.chr10
MZIB.chr11
MZIB.chr12
MZIB.chr13
MZIB.chr14
MZIB.chr15
MZIB.chr16
MZIB.chr17
MZIB.chr18
MZIB.chrX
* MMAR.chr1
MMAR.chr2
* MMAR.chr3
MMAR.chr4
* MMAR.chr5
* MMAR.chr6
* MMAR.chr7
* MMAR.chr8
* MMAR.chr9
MMAR.chr10
MMAR.chr11
MMAR.chr12
* MMAR.chr13
* MMAR.chr14
MMAR.chr15
MMAR.chr16
* MMAR.chr17
* MMAR.chr18
* MMAR.chrX
> 7.00
6.00 - 7.00
5.00 - 6.00
4.00 - 5.00
3.00 - 4.00
2.00 - 3.00
1.00 - 2.00
0.50 - 1.00
0.10 - 0.50
0.00 - 0.10
no SNPs
0 Mbp
50 Mbp
100 Mbp
150 Mbp
200 Mbp
250 Mbp

# T77

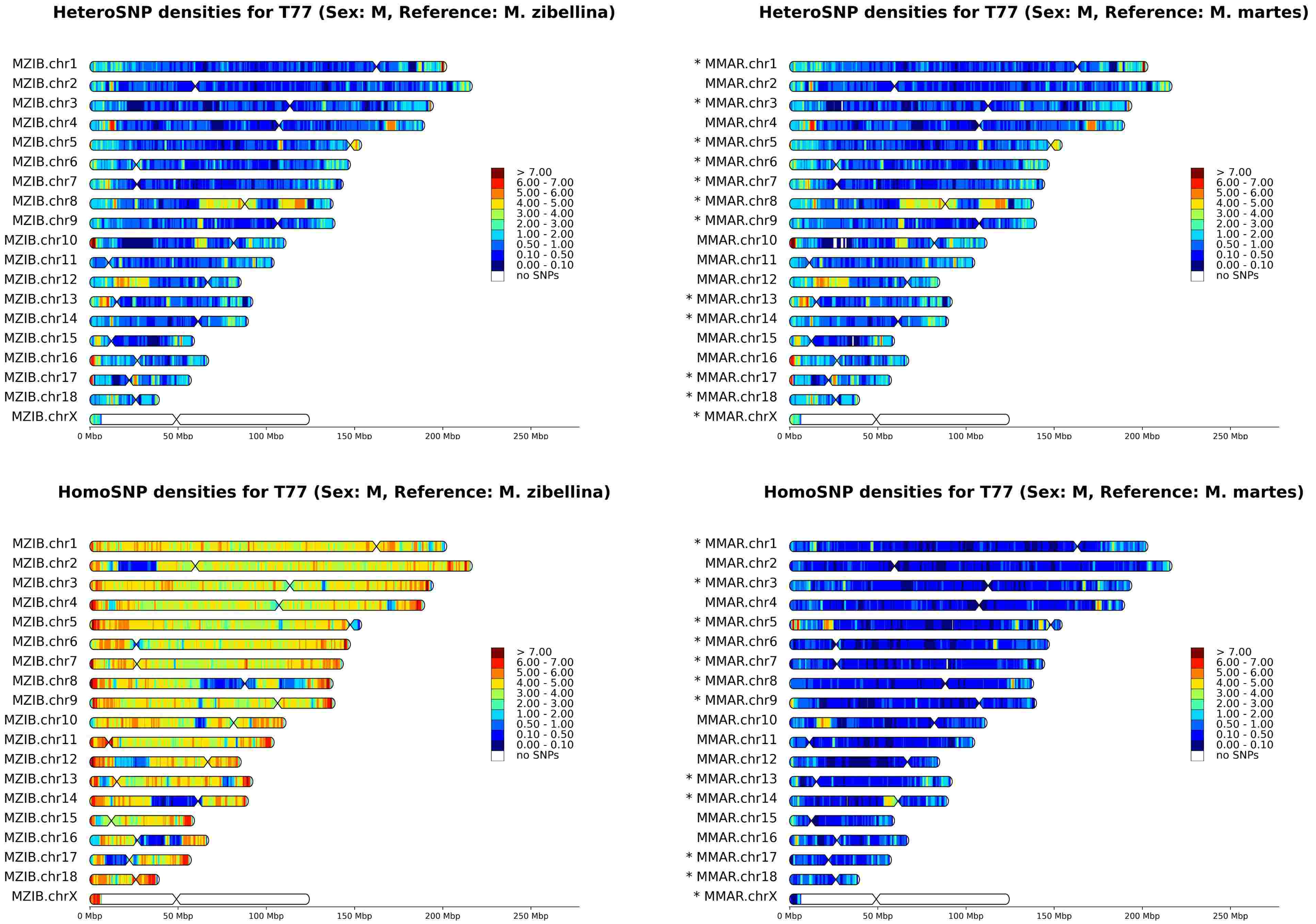
HeteroSNP densities for T77 (Sex: M, Reference: M. zibellina)
MZIB.chr1
MZIB.chr2
MZIB.chr3
MZIB.chr4
MZIB.chr5
MZIB.chr6
MZIB.chr7
MZIB.chr8
MZIB.chr9
MZIB.chr10
MZIB.chr11
MZIB.chr12
MZIB.chr13
MZIB.chr14
MZIB.chr15
MZIB.chr16
MZIB.chr17
MZIB.chr18
MZIB.chrX
0 Mbp
50 Mbp
100 Mbp
150 Mbp
200 Mbp
250 Mbp
> 7.00
6.00 - 7.00
5.00 - 6.00
4.00 - 5.00
3.00 - 4.00
2.00 - 3.00
1.00 - 2.00
0.50 - 1.00
0.10 - 0.50
0.00 - 0.10
no SNPs
HeteroSNP densities for T77 (Sex: M, Reference: M. martes)
* MMAR.chr1
MMAR.chr2
* MMAR.chr3
MMAR.chr4
* MMAR.chr5
* MMAR.chr6
* MMAR.chr7
* MMAR.chr8
* MMAR.chr9
MMAR.chr10
MMAR.chr11
MMAR.chr12
* MMAR.chr13
* MMAR.chr14
MMAR.chr15
MMAR.chr16
* MMAR.chr17
* MMAR.chr18
* MMAR.chrX
HomoSNP densities for T77 (Sex: M, Reference: M. zibellina)
HomoSNP densities for T77 (Sex: M, Reference: M. martes)

# T78

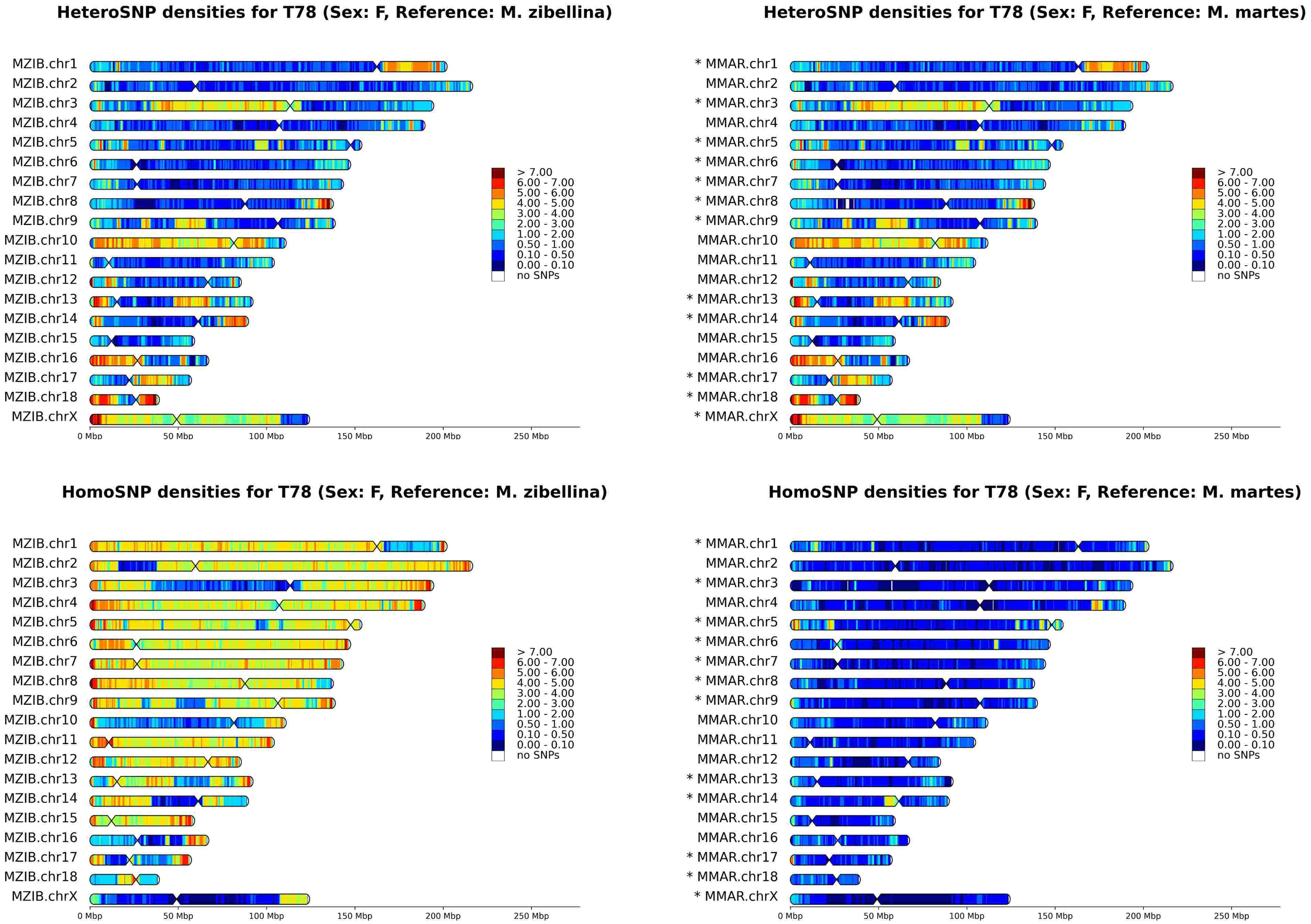
HeteroSNP densities for T78 (Sex: F, Reference: M. zibellina)
HeteroSNP densities for T78 (Sex: F, Reference: M. martes)
HomoSNP densities for T78 (Sex: F, Reference: M. zibellina)
HomoSNP densities for T78 (Sex: F, Reference: M. martes)
MZIB.chr1
MZIB.chr2
MZIB.chr3
MZIB.chr4
MZIB.chr5
MZIB.chr6
MZIB.chr7
MZIB.chr8
MZIB.chr9
MZIB.chr10
MZIB.chr11
MZIB.chr12
MZIB.chr13
MZIB.chr14
MZIB.chr15
MZIB.chr16
MZIB.chr17
MZIB.chr18
MZIB.chrX
* MMAR.chr1
MMAR.chr2
* MMAR.chr3
MMAR.chr4
* MMAR.chr5
* MMAR.chr6
* MMAR.chr7
* MMAR.chr8
* MMAR.chr9
MMAR.chr10
MMAR.chr11
MMAR.chr12
* MMAR.chr13
* MMAR.chr14
MMAR.chr15
MMAR.chr16
* MMAR.chr17
* MMAR.chr18
* MMAR.chrX
> 7.00
6.00 - 7.00
5.00 - 6.00
4.00 - 5.00
3.00 - 4.00
2.00 - 3.00
1.00 - 2.00
0.50 - 1.00
0.10 - 0.50
0.00 - 0.10
no SNPs
0 Mbp
50 Mbp
100 Mbp
150 Mbp
200 Mbp
250 Mbp

# T79

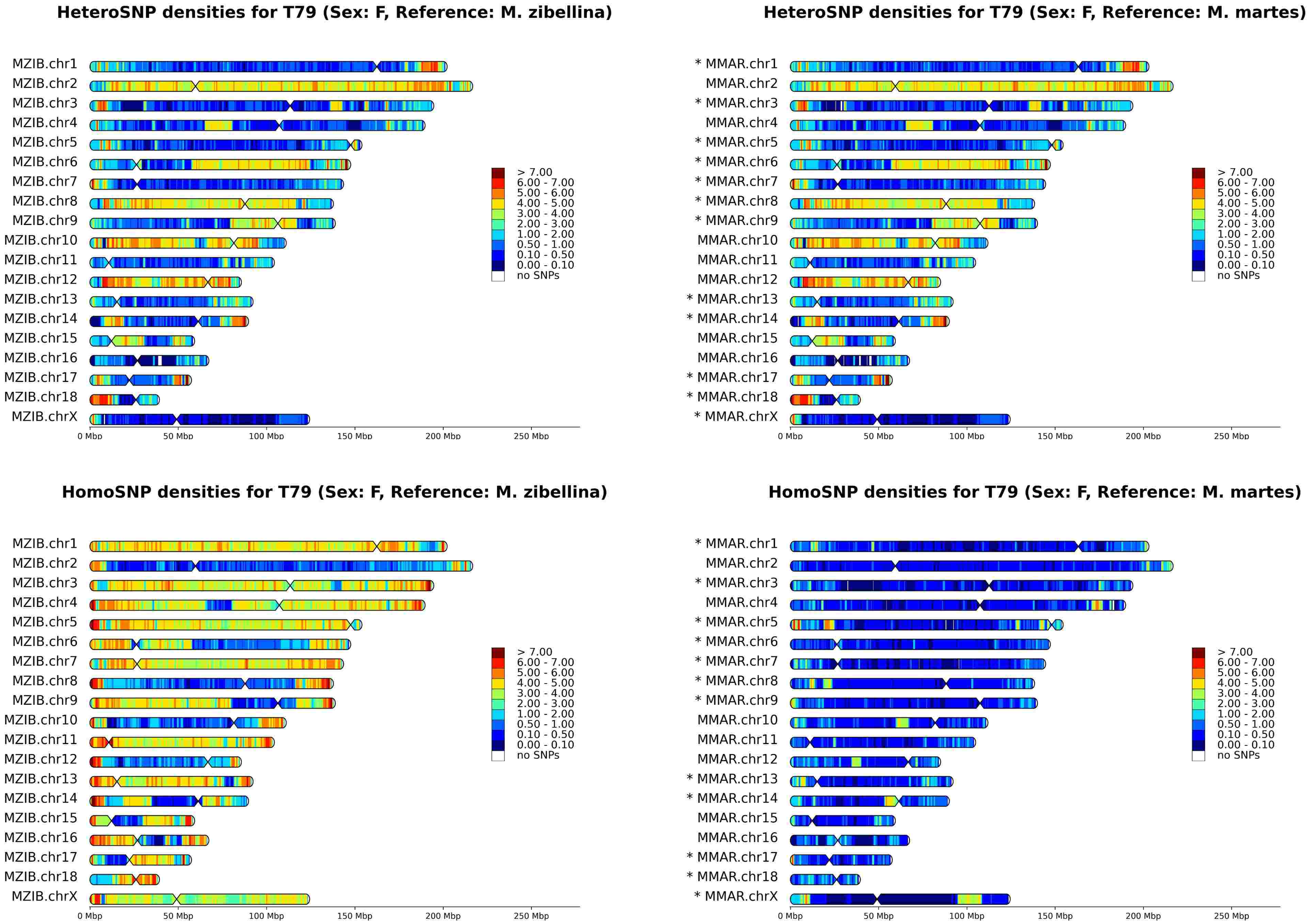
HeteroSNP densities for T79 (Sex: F, Reference: M. zibellina)
MZIB.chr1
MZIB.chr2
MZIB.chr3
MZIB.chr4
MZIB.chr5
MZIB.chr6
MZIB.chr7
MZIB.chr8
MZIB.chr9
MZIB.chr10
MZIB.chr11
MZIB.chr12
MZIB.chr13
MZIB.chr14
MZIB.chr15
MZIB.chr16
MZIB.chr17
MZIB.chr18
MZIB.chrX
0 Mbp
50 Mbp
100 Mbp
150 Mbp
200 Mbp
250 Mbp
> 7.00
6.00 - 7.00
5.00 - 6.00
4.00 - 5.00
3.00 - 4.00
2.00 - 3.00
1.00 - 2.00
0.50 - 1.00
0.10 - 0.50
0.00 - 0.10
no SNPs
HeteroSNP densities for T79 (Sex: F, Reference: M. martes)
* MMAR.chr1
MMAR.chr2
* MMAR.chr3
MMAR.chr4
* MMAR.chr5
* MMAR.chr6
* MMAR.chr7
* MMAR.chr8
* MMAR.chr9
MMAR.chr10
MMAR.chr11
MMAR.chr12
* MMAR.chr13
* MMAR.chr14
MMAR.chr15
MMAR.chr16
* MMAR.chr17
* MMAR.chr18
* MMAR.chrX
HomoSNP densities for T79 (Sex: F, Reference: M. zibellina)
HomoSNP densities for T79 (Sex: F, Reference: M. martes)

# T81

**HeteroSNP densities for T81 (Sex: M, Reference: M. zibellina)**

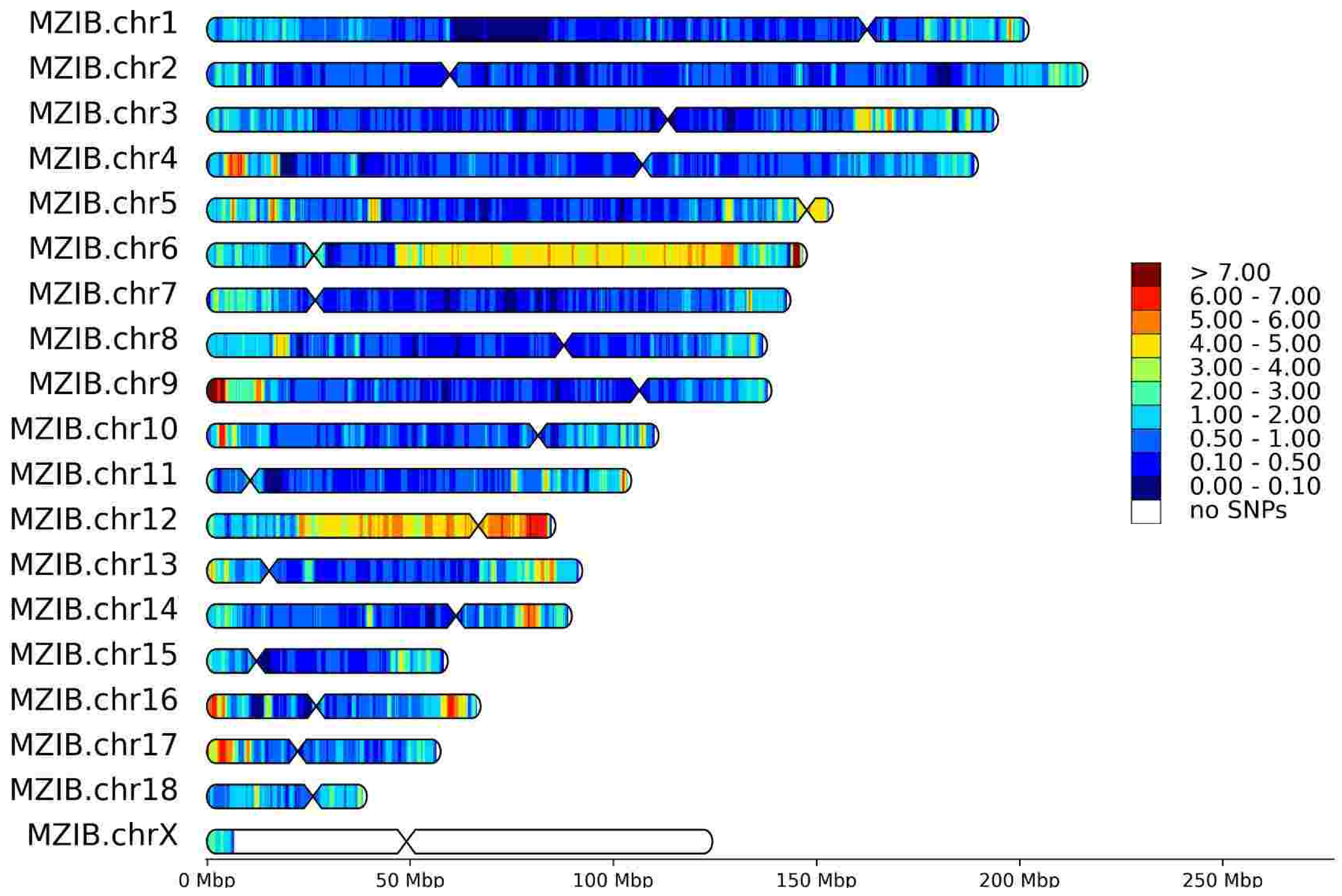


**HeteroSNP densities for T81 (Sex: M, Reference: M. martes)**

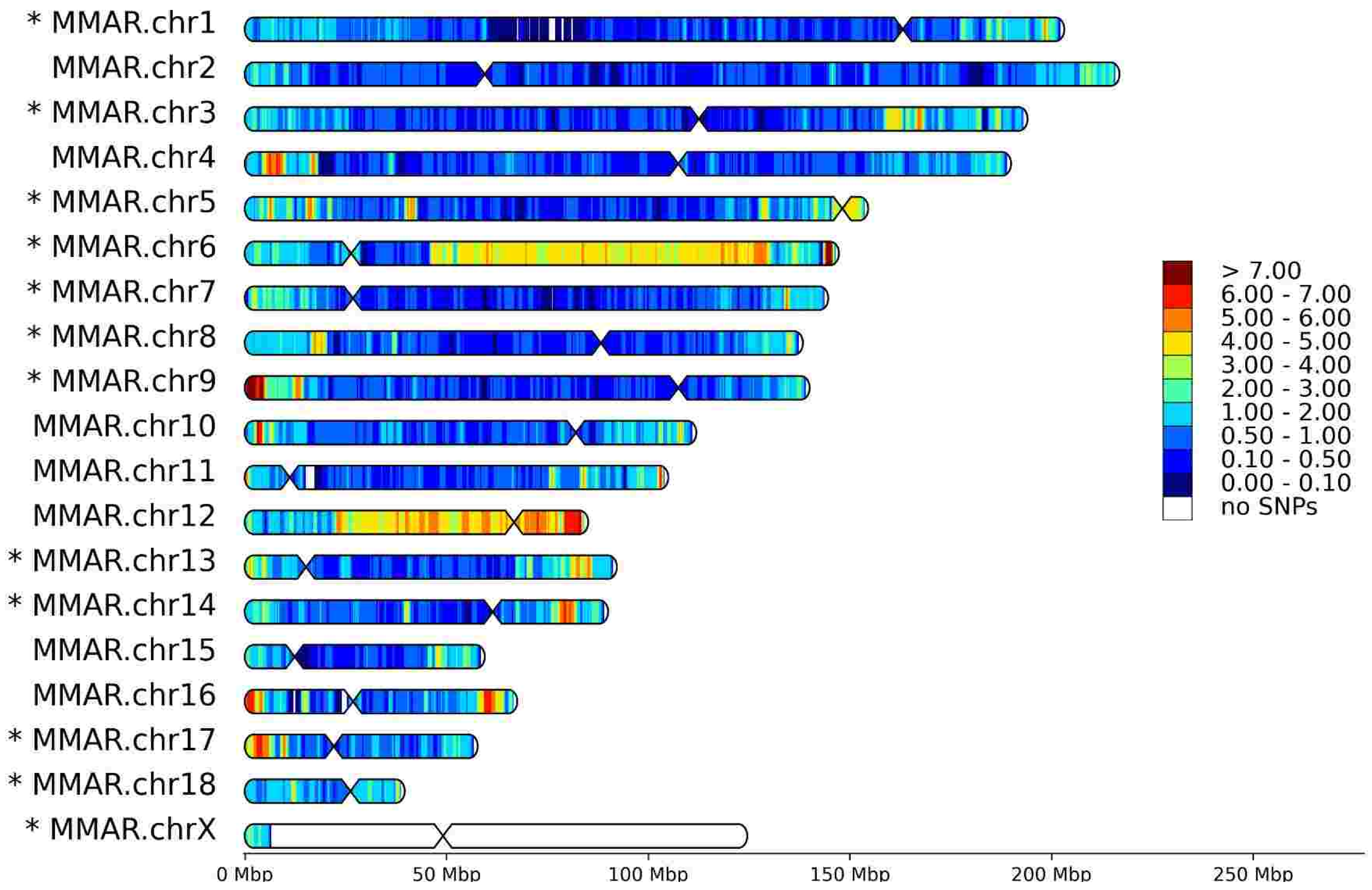


**HomoSNP densities for T81 (Sex: M, Reference: M. zibellina)**

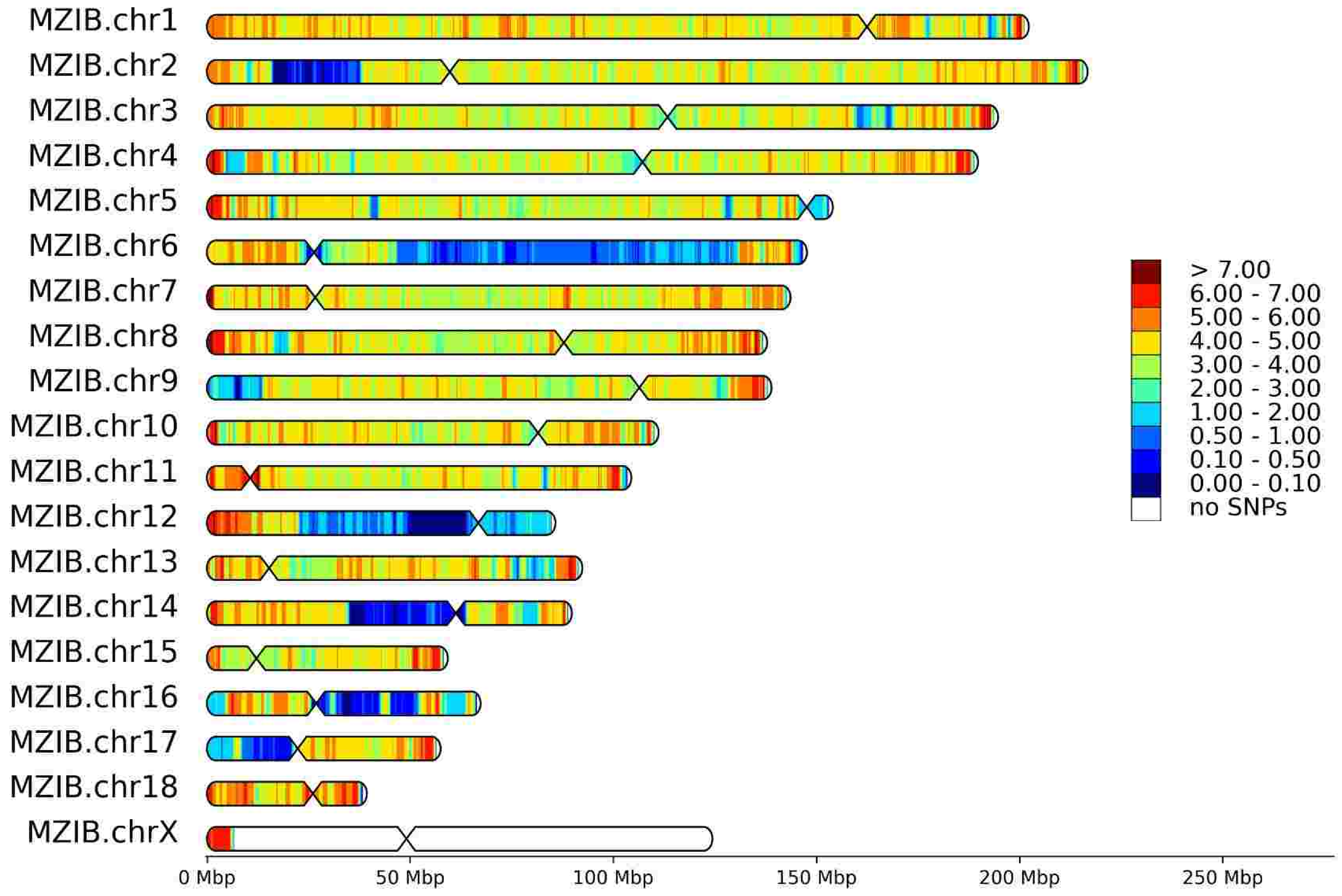


**HomoSNP densities for T81 (Sex: M, Reference: M. martes)**

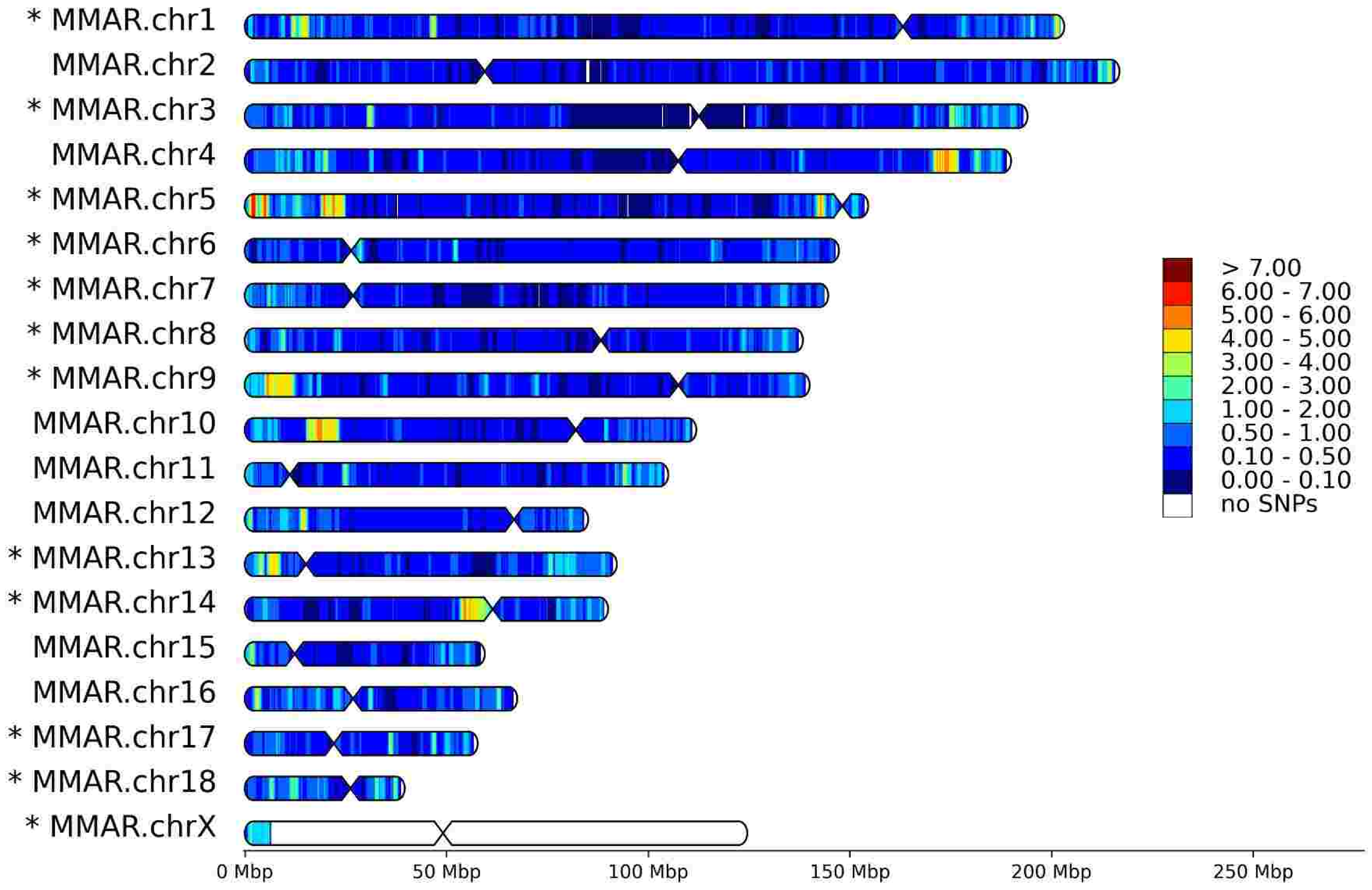

# T82

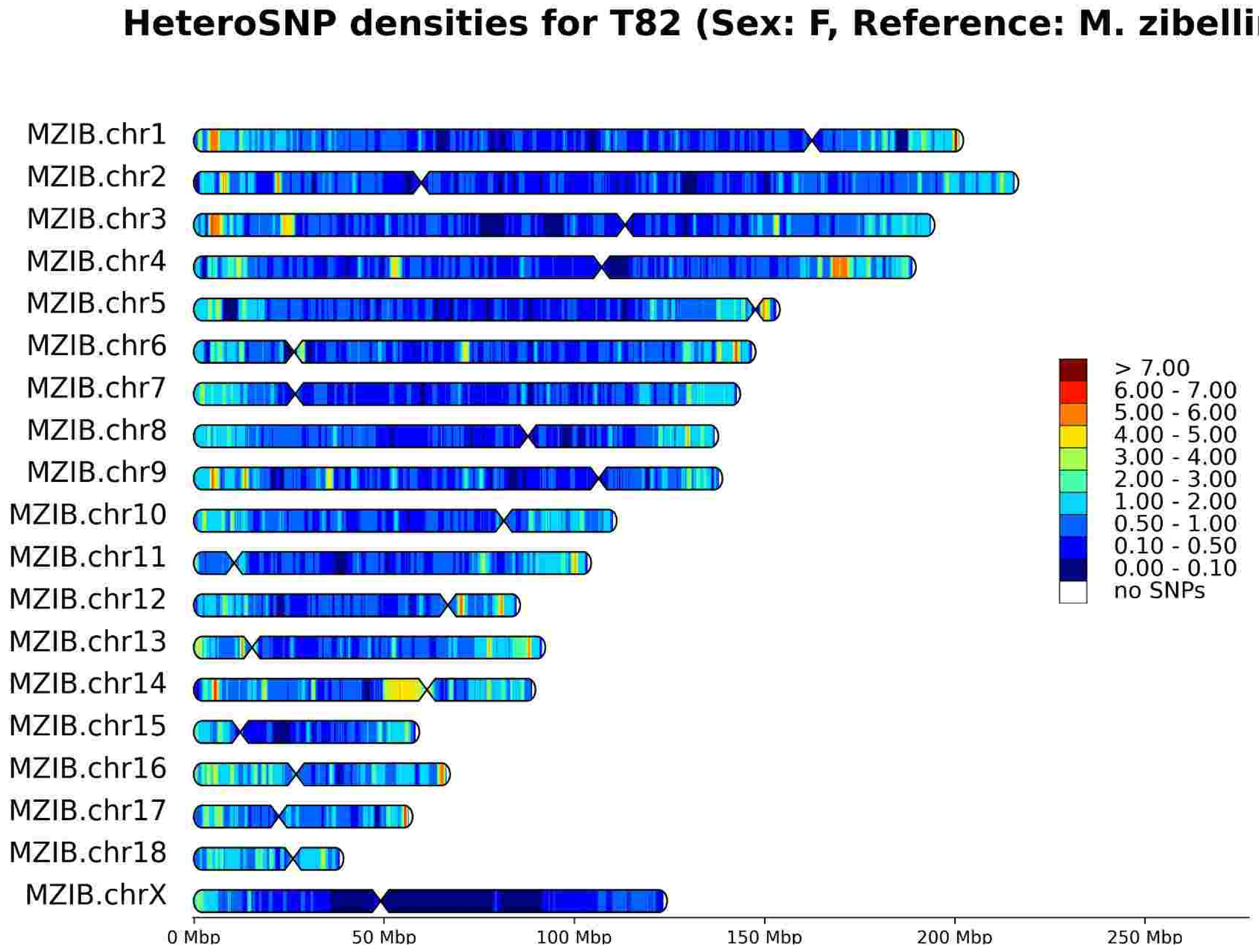
HeteroSNP densities for T82 (Sex: F, Reference: M. zibellina)
MZIB.chr1
MZIB.chr2
MZIB.chr3
MZIB.chr4
MZIB.chr5
MZIB.chr6
MZIB.chr7
MZIB.chr8
MZIB.chr9
MZIB.chr10
MZIB.chr11
MZIB.chr12
MZIB.chr13
MZIB.chr14
MZIB.chr15
MZIB.chr16
MZIB.chr17
MZIB.chr18
MZIB.chrX
> 7.00
6.00 - 7.00
5.00 - 6.00
4.00 - 5.00
3.00 - 4.00
2.00 - 3.00
1.00 - 2.00
0.50 - 1.00
0.10 - 0.50
0.00 - 0.10
no SNPs
0 Mbp
50 Mbp
100 Mbp
150 Mbp
200 Mbp
250 Mbp

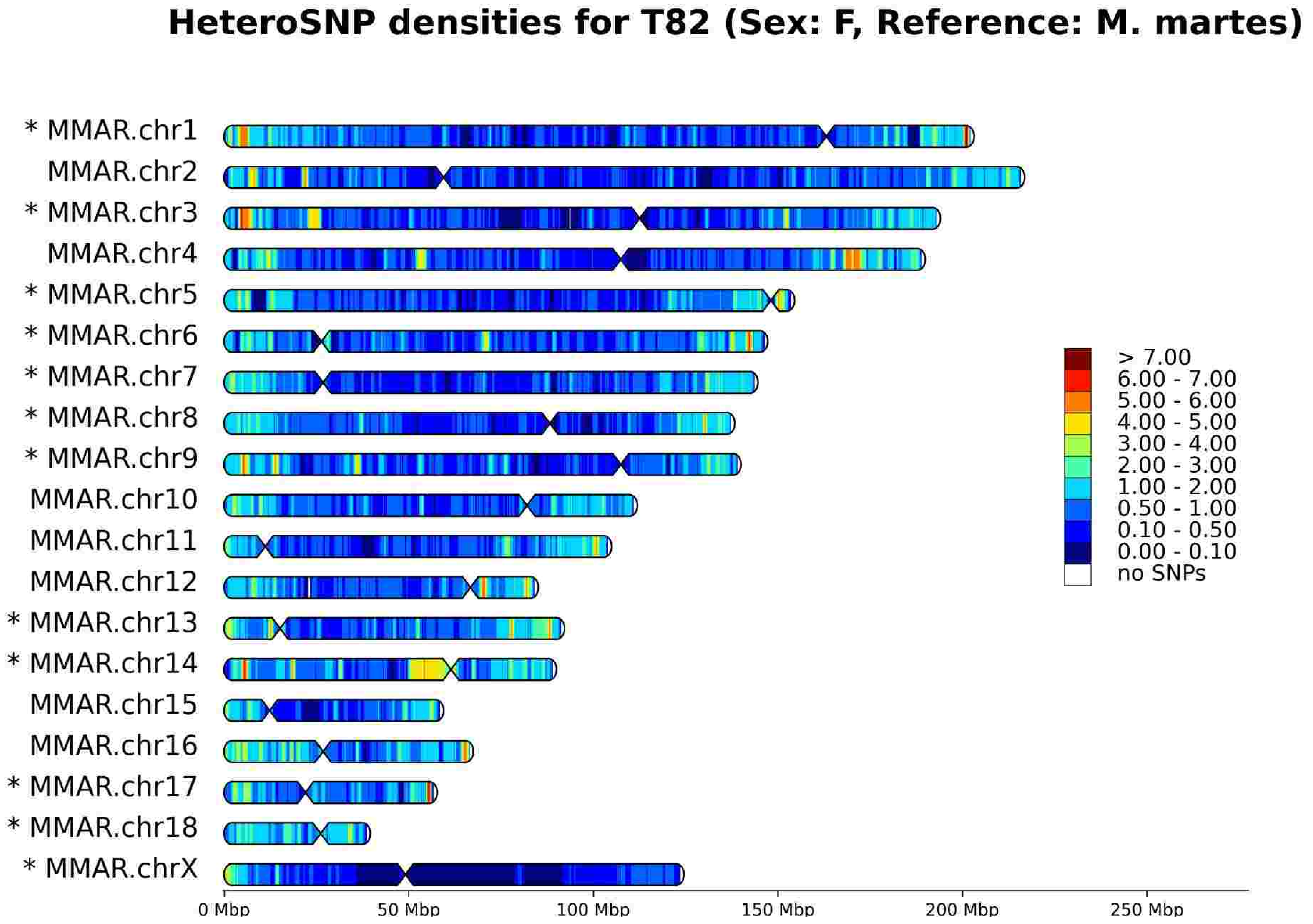
HeteroSNP densities for T82 (Sex: F, Reference: M. martes)
* MMAR.chr1
MMAR.chr2
* MMAR.chr3
MMAR.chr4
* MMAR.chr5
* MMAR.chr6
* MMAR.chr7
* MMAR.chr8
* MMAR.chr9
MMAR.chr10
MMAR.chr11
MMAR.chr12
* MMAR.chr13
* MMAR.chr14
MMAR.chr15
MMAR.chr16
* MMAR.chr17
* MMAR.chr18
* MMAR.chrX
> 7.00
6.00 - 7.00
5.00 - 6.00
4.00 - 5.00
3.00 - 4.00
2.00 - 3.00
1.00 - 2.00
0.50 - 1.00
0.10 - 0.50
0.00 - 0.10
no SNPs
0 Mbp
50 Mbp
100 Mbp
150 Mbp
200 Mbp
250 Mbp

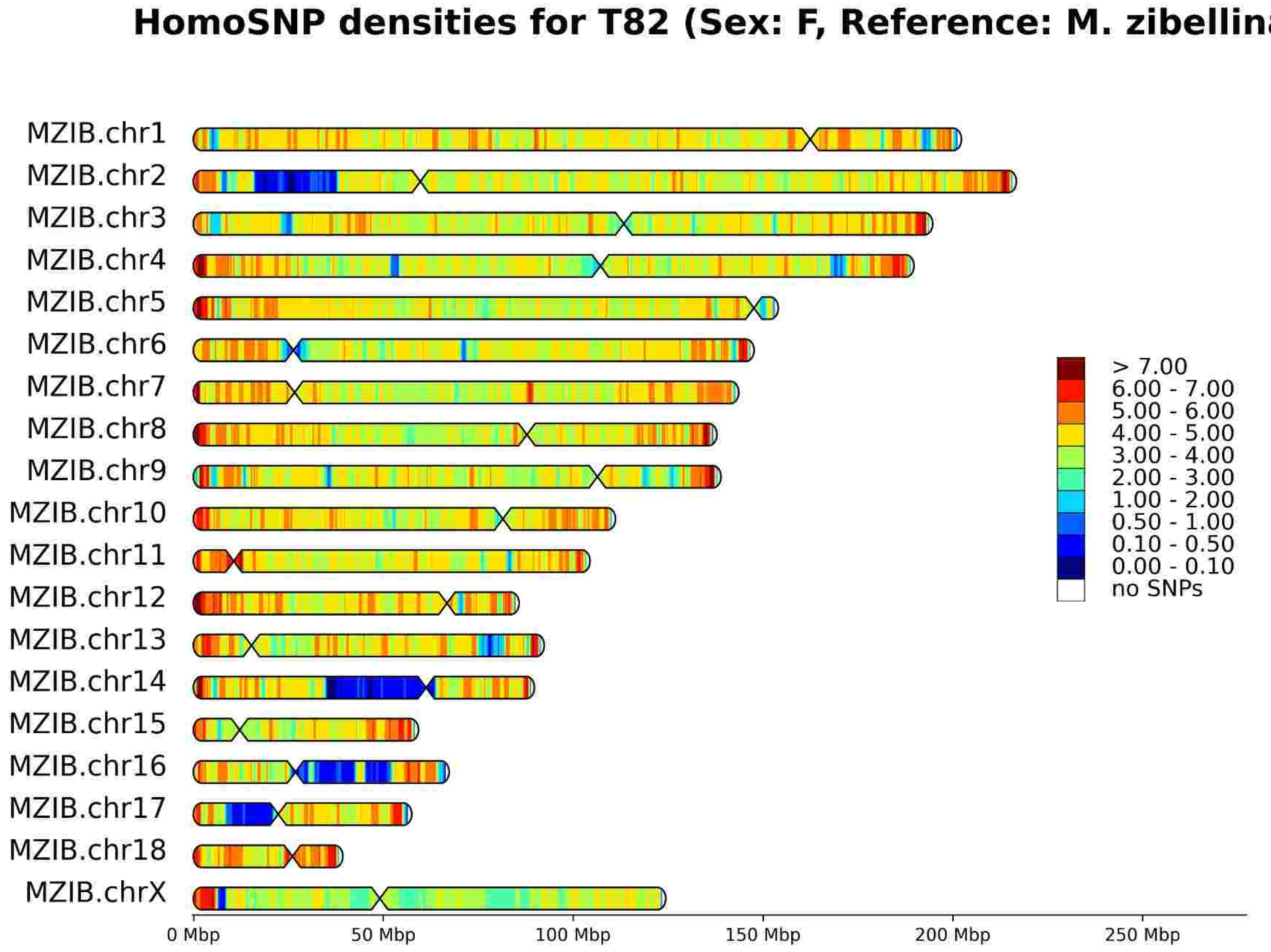
HomoSNP densities for T82 (Sex: F, Reference: M. zibellina)
MZIB.chr1
MZIB.chr2
MZIB.chr3
MZIB.chr4
MZIB.chr5
MZIB.chr6
MZIB.chr7
MZIB.chr8
MZIB.chr9
MZIB.chr10
MZIB.chr11
MZIB.chr12
MZIB.chr13
MZIB.chr14
MZIB.chr15
MZIB.chr16
MZIB.chr17
MZIB.chr18
MZIB.chrX
> 7.00
6.00 - 7.00
5.00 - 6.00
4.00 - 5.00
3.00 - 4.00
2.00 - 3.00
1.00 - 2.00
0.50 - 1.00
0.10 - 0.50
0.00 - 0.10
no SNPs
0 Mbp
50 Mbp
100 Mbp
150 Mbp
200 Mbp
250 Mbp

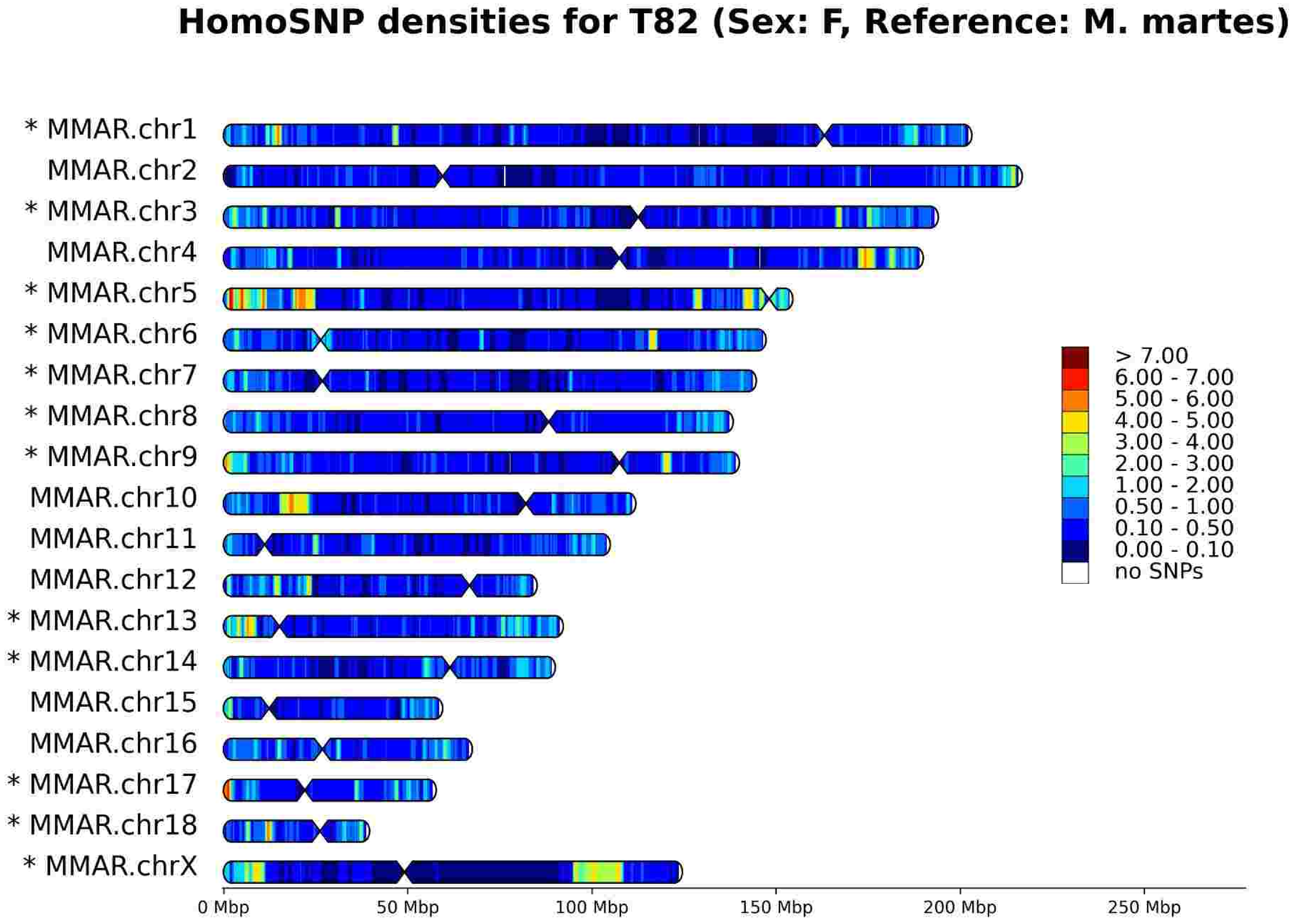
HomoSNP densities for T82 (Sex: F, Reference: M. martes)
* MMAR.chr1
MMAR.chr2
* MMAR.chr3
MMAR.chr4
* MMAR.chr5
* MMAR.chr6
* MMAR.chr7
* MMAR.chr8
* MMAR.chr9
MMAR.chr10
MMAR.chr11
MMAR.chr12
* MMAR.chr13
* MMAR.chr14
MMAR.chr15
MMAR.chr16
* MMAR.chr17
* MMAR.chr18
* MMAR.chrX
> 7.00
6.00 - 7.00
5.00 - 6.00
4.00 - 5.00
3.00 - 4.00
2.00 - 3.00
1.00 - 2.00
0.50 - 1.00
0.10 - 0.50
0.00 - 0.10
no SNPs
0 Mbp
50 Mbp
100 Mbp
150 Mbp
200 Mbp
250 Mbp

# T83

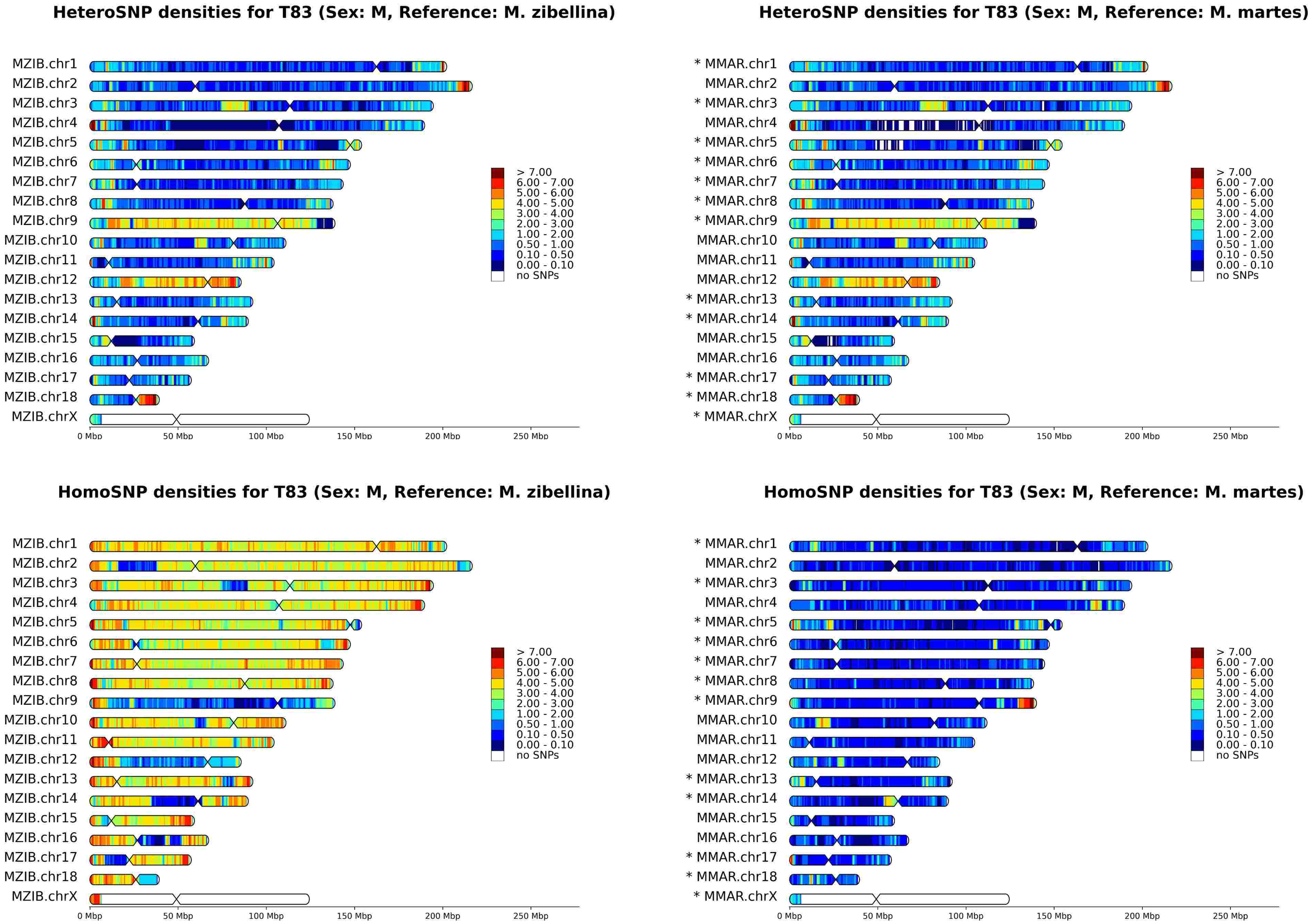
HeteroSNP densities for T83 (Sex: M, Reference: M. zibellina)
HeteroSNP densities for T83 (Sex: M, Reference: M. martes)
HomoSNP densities for T83 (Sex: M, Reference: M. zibellina)
HomoSNP densities for T83 (Sex: M, Reference: M. martes)
MZIB.chr1
MZIB.chr2
MZIB.chr3
MZIB.chr4
MZIB.chr5
MZIB.chr6
MZIB.chr7
MZIB.chr8
MZIB.chr9
MZIB.chr10
MZIB.chr11
MZIB.chr12
MZIB.chr13
MZIB.chr14
MZIB.chr15
MZIB.chr16
MZIB.chr17
MZIB.chr18
MZIB.chrX
* MMAR.chr1
MMAR.chr2
* MMAR.chr3
MMAR.chr4
* MMAR.chr5
* MMAR.chr6
* MMAR.chr7
* MMAR.chr8
* MMAR.chr9
MMAR.chr10
MMAR.chr11
MMAR.chr12
* MMAR.chr13
* MMAR.chr14
MMAR.chr15
MMAR.chr16
* MMAR.chr17
* MMAR.chr18
* MMAR.chrX
0 Mbp
50 Mbp
100 Mbp
150 Mbp
200 Mbp
250 Mbp
> 7.00
6.00 - 7.00
5.00 - 6.00
4.00 - 5.00
3.00 - 4.00
2.00 - 3.00
1.00 - 2.00
0.50 - 1.00
0.10 - 0.50
0.00 - 0.10
no SNPs

# T84

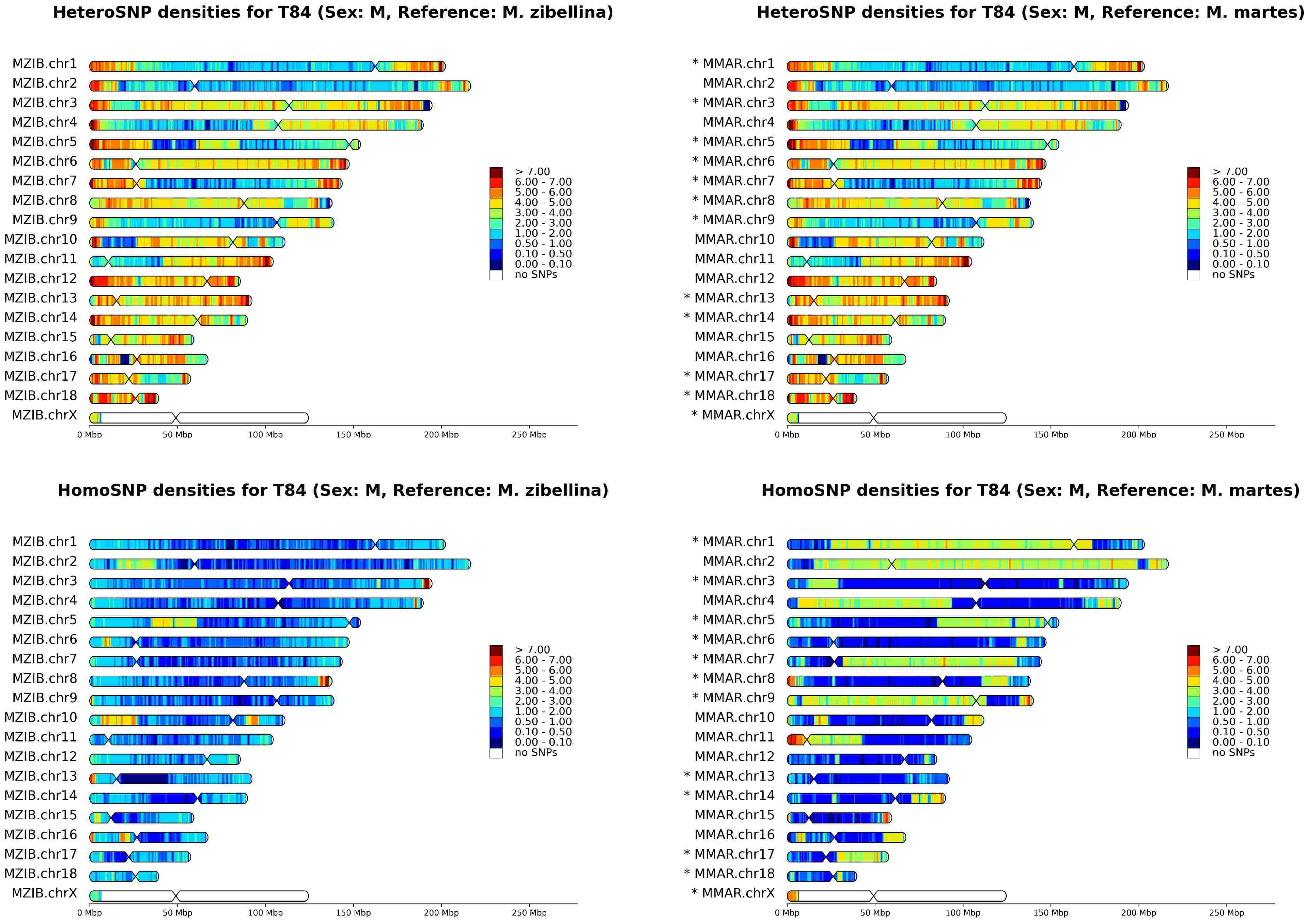

HeteroSNP densities for T84 (Sex: M, Reference: M. zibellina)
MZIB.chr1
MZIB.chr2
MZIB.chr3
MZIB.chr4
MZIB.chr5
MZIB.chr6
MZIB.chr7
MZIB.chr8
MZIB.chr9
MZIB.chr10
MZIB.chr11
MZIB.chr12
MZIB.chr13
MZIB.chr14
MZIB.chr15
MZIB.chr16
MZIB.chr17
MZIB.chr18
MZIB.chrX
0 Mbp
50 Mbp
100 Mbp
150 Mbp
200 Mbp
250 Mbp
> 7.00
6.00 - 7.00
5.00 - 6.00
4.00 - 5.00
3.00 - 4.00
2.00 - 3.00
1.00 - 2.00
0.50 - 1.00
0.10 - 0.50
0.00 - 0.10
no SNPs
HeteroSNP densities for T84 (Sex: M, Reference: M. martes)
* MMAR.chr1
MMAR.chr2
* MMAR.chr3
MMAR.chr4
* MMAR.chr5
* MMAR.chr6
* MMAR.chr7
* MMAR.chr8
* MMAR.chr9
MMAR.chr10
MMAR.chr11
MMAR.chr12
* MMAR.chr13
* MMAR.chr14
MMAR.chr15
MMAR.chr16
* MMAR.chr17
* MMAR.chr18
* MMAR.chrX
0 Mbp
50 Mbp
100 Mbp
150 Mbp
200 Mbp
250 Mbp
> 7.00
6.00 - 7.00
5.00 - 6.00
4.00 - 5.00
3.00 - 4.00
2.00 - 3.00
1.00 - 2.00
0.50 - 1.00
0.10 - 0.50
0.00 - 0.10
no SNPs
HomoSNP densities for T84 (Sex: M, Reference: M. zibellina)
MZIB.chr1
MZIB.chr2
MZIB.chr3
MZIB.chr4
MZIB.chr5
MZIB.chr6
MZIB.chr7
MZIB.chr8
MZIB.chr9
MZIB.chr10
MZIB.chr11
MZIB.chr12
MZIB.chr13
MZIB.chr14
MZIB.chr15
MZIB.chr16
MZIB.chr17
MZIB.chr18
MZIB.chrX
0 Mbp
50 Mbp
100 Mbp
150 Mbp
200 Mbp
250 Mbp
> 7.00
6.00 - 7.00
5.00 - 6.00
4.00 - 5.00
3.00 - 4.00
2.00 - 3.00
1.00 - 2.00
0.50 - 1.00
0.10 - 0.50
0.00 - 0.10
no SNPs
HomoSNP densities for T84 (Sex: M, Reference: M. martes)
* MMAR.chr1
MMAR.chr2
* MMAR.chr3
MMAR.chr4
* MMAR.chr5
* MMAR.chr6
* MMAR.chr7
* MMAR.chr8
* MMAR.chr9
MMAR.chr10
MMAR.chr11
MMAR.chr12
* MMAR.chr13
* MMAR.chr14
MMAR.chr15
MMAR.chr16
* MMAR.chr17
* MMAR.chr18
* MMAR.chrX
0 Mbp
50 Mbp
100 Mbp
150 Mbp
200 Mbp
250 Mbp
> 7.00
6.00 - 7.00
5.00 - 6.00
4.00 - 5.00
3.00 - 4.00
2.00 - 3.00
1.00 - 2.00
0.50 - 1.00
0.10 - 0.50
0.00 - 0.10
no SNPs

# T85

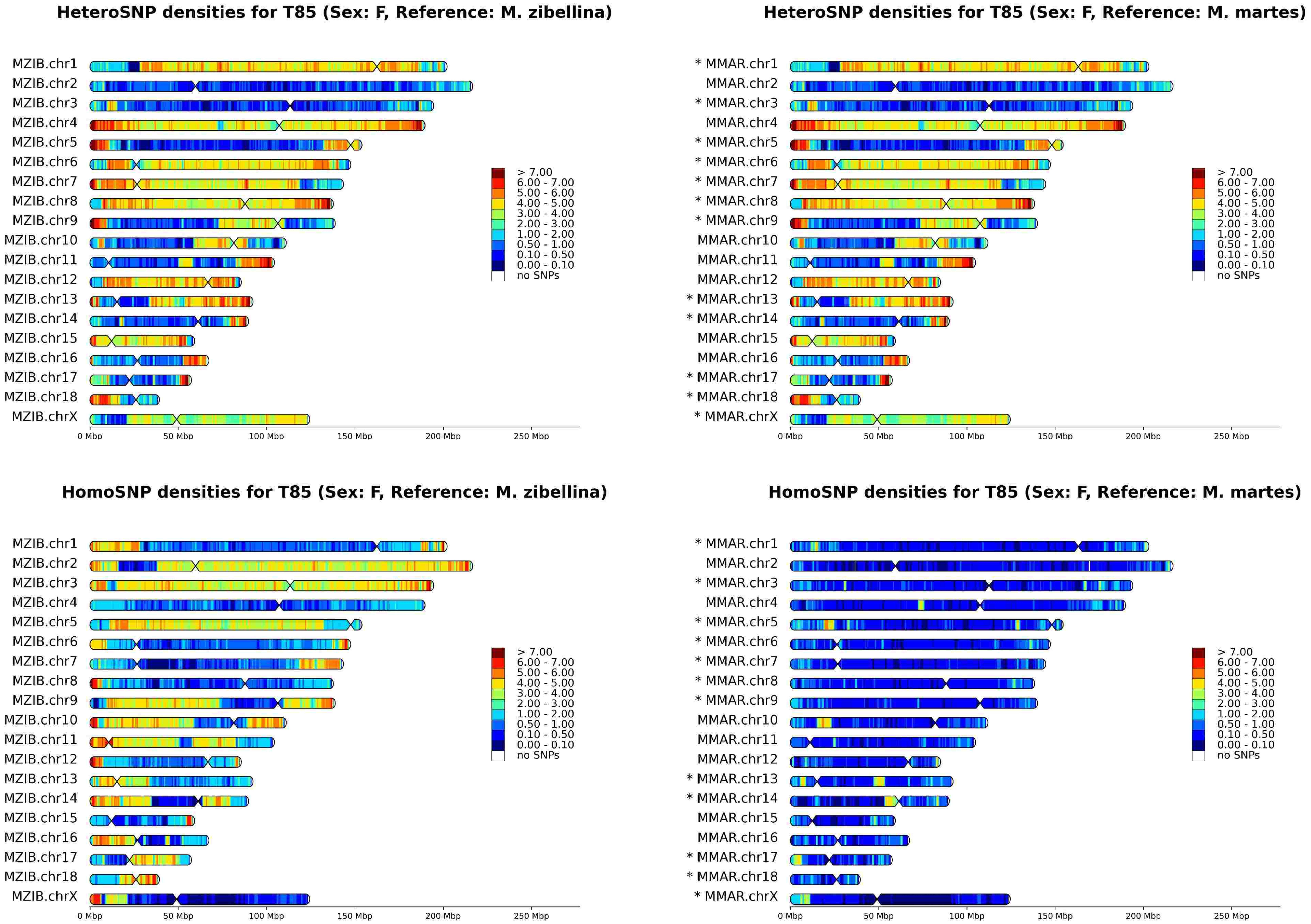

HeteroSNP densities for T85 (Sex: F, Reference: M. zibellina)
MZIB.chr1
MZIB.chr2
MZIB.chr3
MZIB.chr4
MZIB.chr5
MZIB.chr6
MZIB.chr7
MZIB.chr8
MZIB.chr9
MZIB.chr10
MZIB.chr11
MZIB.chr12
MZIB.chr13
MZIB.chr14
MZIB.chr15
MZIB.chr16
MZIB.chr17
MZIB.chr18
MZIB.chrX
0 Mbp
50 Mbp
100 Mbp
150 Mbp
200 Mbp
250 Mbp
> 7.00
6.00 - 7.00
5.00 - 6.00
4.00 - 5.00
3.00 - 4.00
2.00 - 3.00
1.00 - 2.00
0.50 - 1.00
0.10 - 0.50
0.00 - 0.10
no SNPs
HeteroSNP densities for T85 (Sex: F, Reference: M. martes)
* MMAR.chr1
MMAR.chr2
* MMAR.chr3
MMAR.chr4
* MMAR.chr5
* MMAR.chr6
* MMAR.chr7
* MMAR.chr8
* MMAR.chr9
MMAR.chr10
MMAR.chr11
MMAR.chr12
* MMAR.chr13
* MMAR.chr14
MMAR.chr15
MMAR.chr16
* MMAR.chr17
* MMAR.chr18
* MMAR.chrX
HomoSNP densities for T85 (Sex: F, Reference: M. zibellina)
HomoSNP densities for T85 (Sex: F, Reference: M. martes)

# T86

HeteroSNP densities for T86 (Sex: M, Reference: M. zibellina)

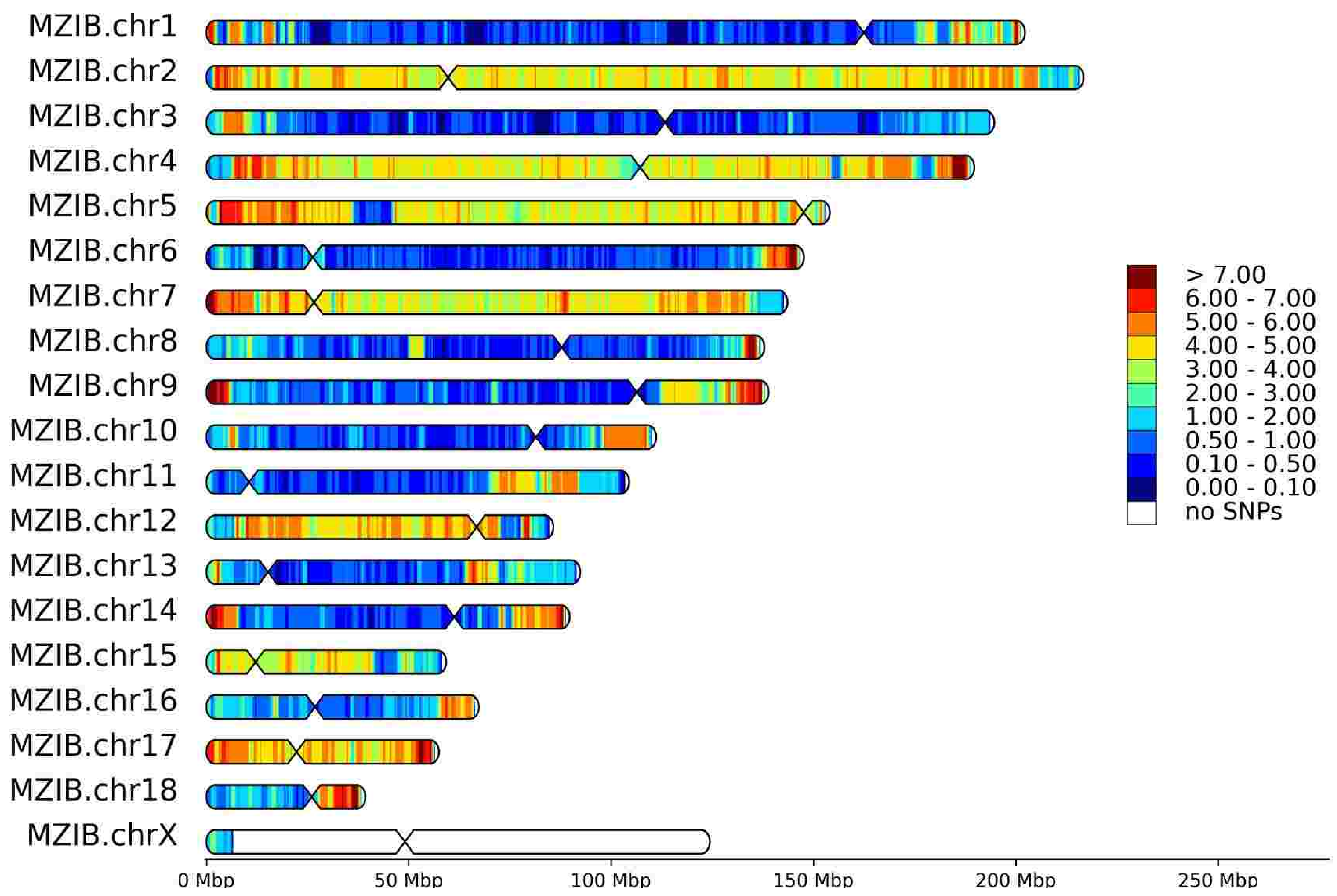


HeteroSNP densities for T86 (Sex: M, Reference: M. martes)

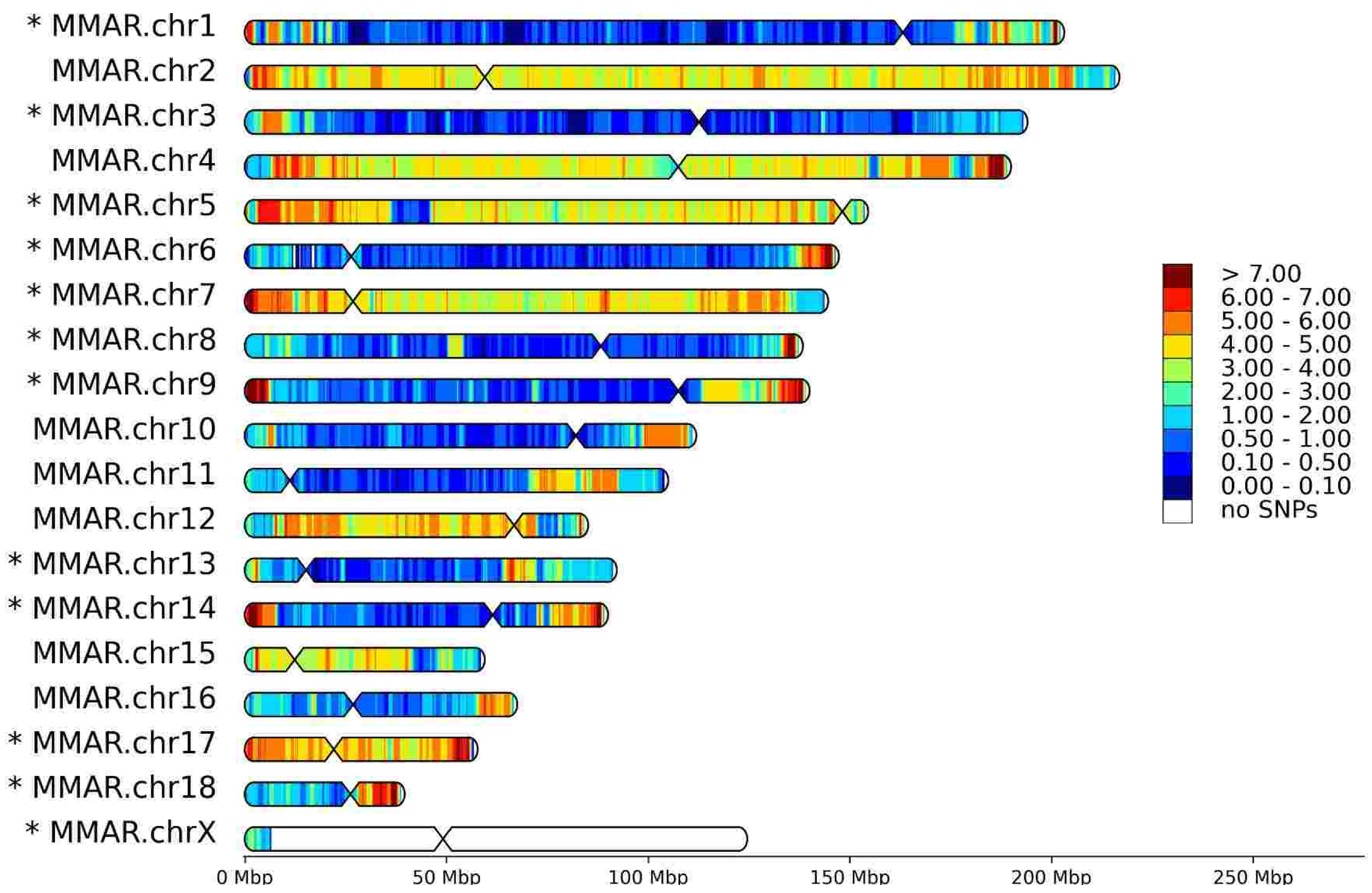


HomoSNP densities for T86 (Sex: M, Reference: M. zibellina)

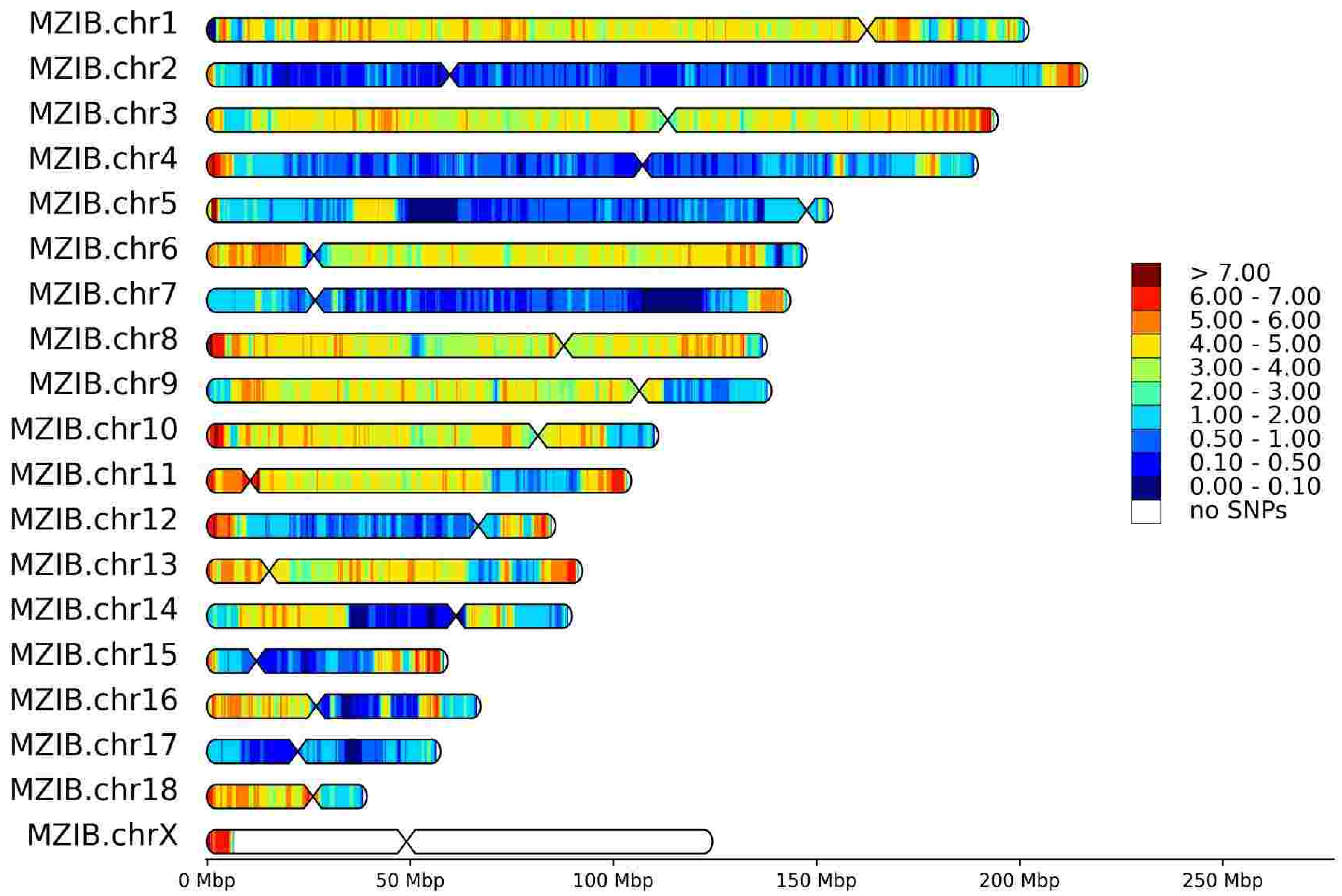


HomoSNP densities for T86 (Sex: M, Reference: M. martes)

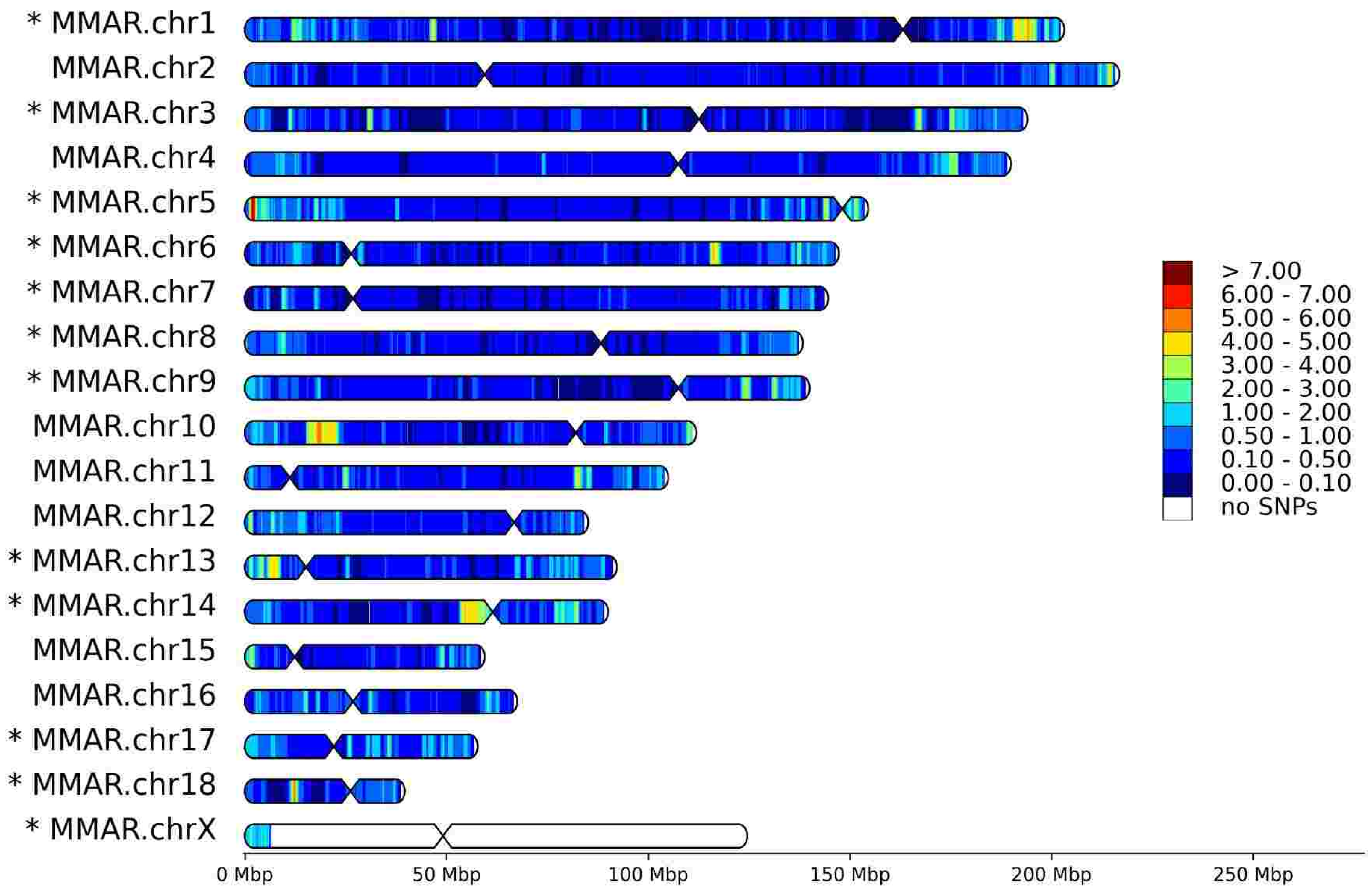

# T87

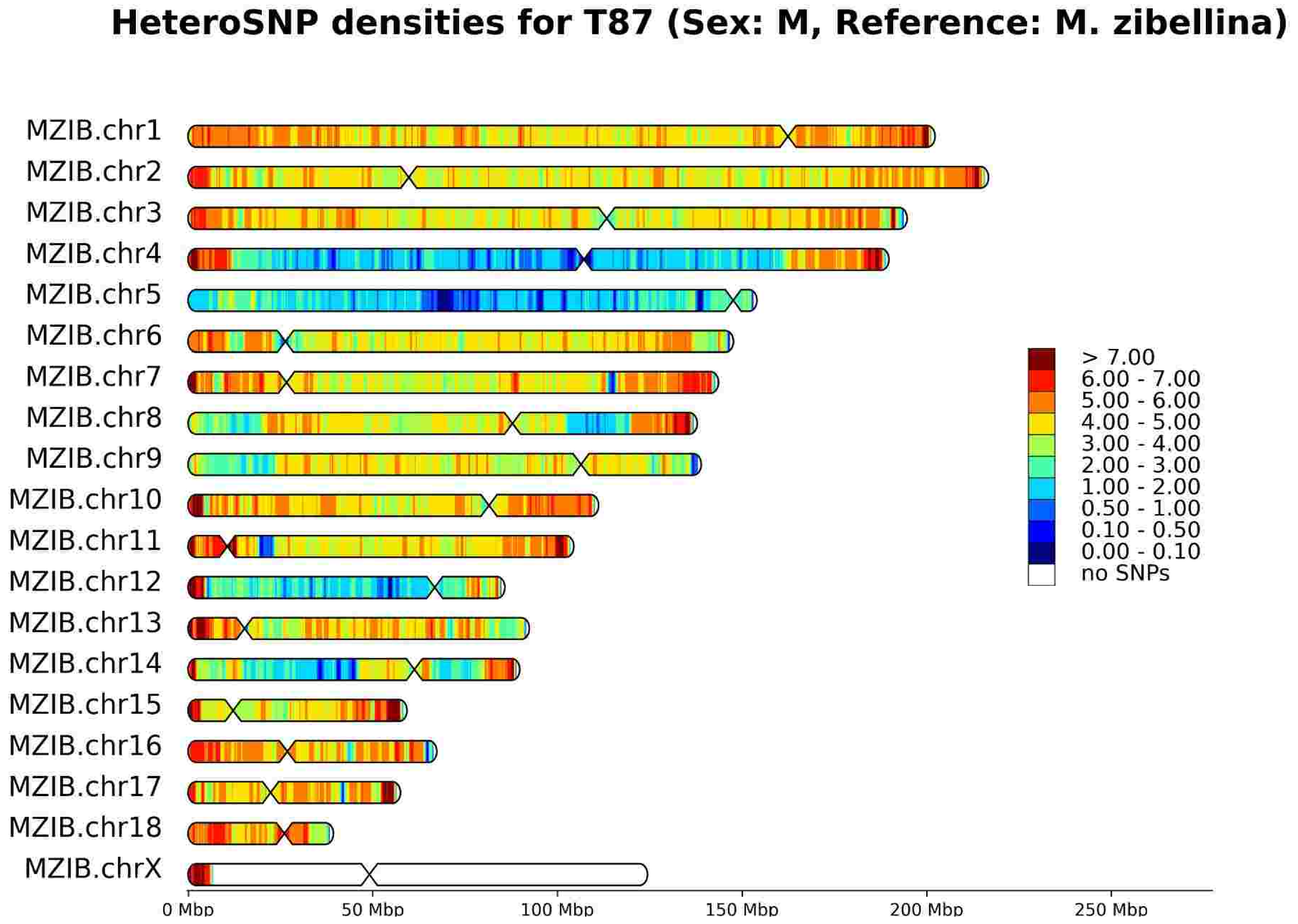

HeteroSNP densities for T87 (Sex: M, Reference: M. zibellina)
MZIB.chr1
MZIB.chr2
MZIB.chr3
MZIB.chr4
MZIB.chr5
MZIB.chr6
MZIB.chr7
MZIB.chr8
MZIB.chr9
MZIB.chr10
MZIB.chr11
MZIB.chr12
MZIB.chr13
MZIB.chr14
MZIB.chr15
MZIB.chr16
MZIB.chr17
MZIB.chr18
MZIB.chrX
> 7.00
6.00 - 7.00
5.00 - 6.00
4.00 - 5.00
3.00 - 4.00
2.00 - 3.00
1.00 - 2.00
0.50 - 1.00
0.10 - 0.50
0.00 - 0.10
no SNPs
0 Mbp
50 Mbp
100 Mbp
150 Mbp
200 Mbp
250 Mbp


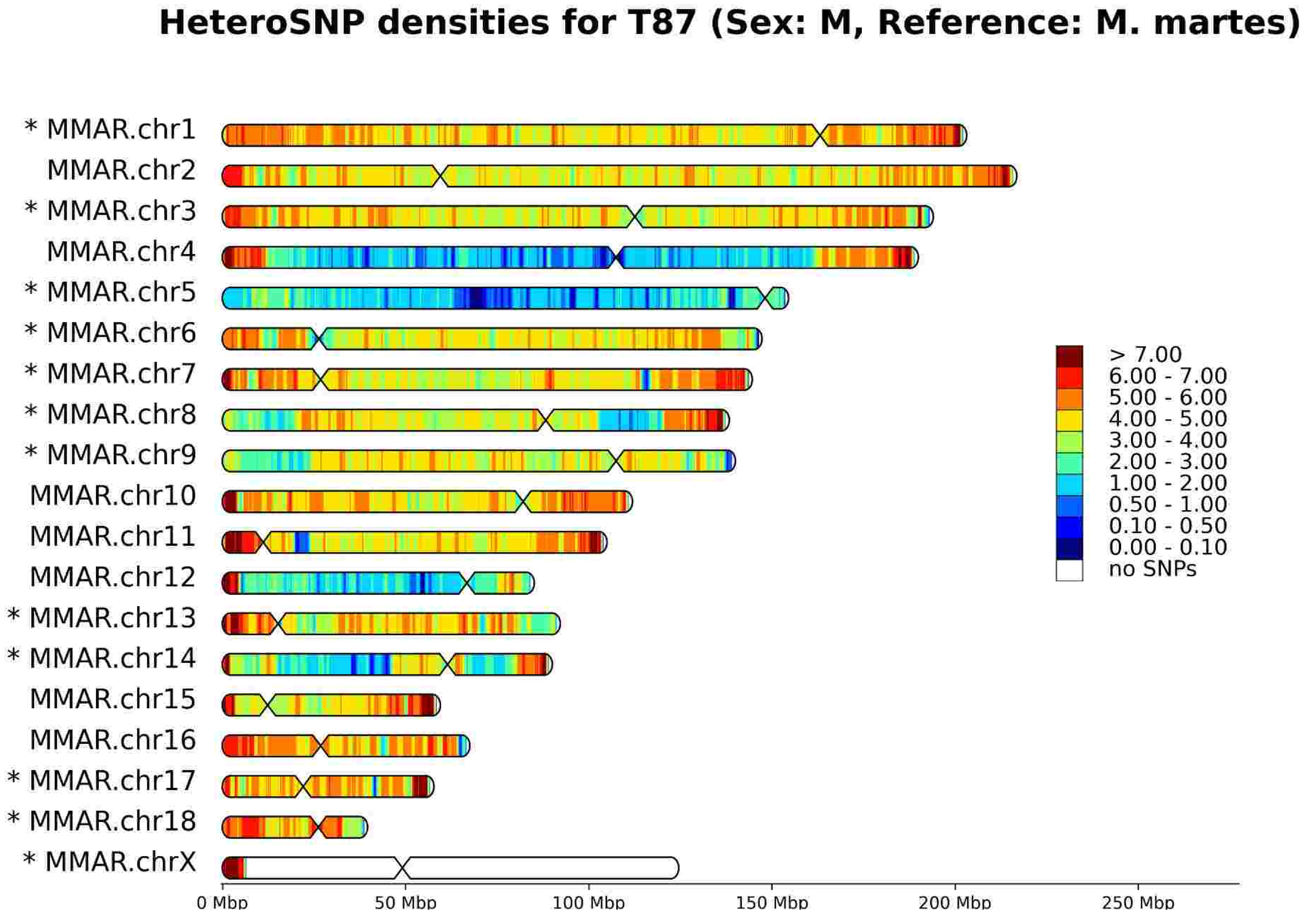

HeteroSNP densities for T87 (Sex: M, Reference: M. martes)
* MMAR.chr1
MMAR.chr2
* MMAR.chr3
MMAR.chr4
* MMAR.chr5
* MMAR.chr6
* MMAR.chr7
* MMAR.chr8
* MMAR.chr9
MMAR.chr10
MMAR.chr11
MMAR.chr12
* MMAR.chr13
* MMAR.chr14
MMAR.chr15
MMAR.chr16
* MMAR.chr17
* MMAR.chr18
* MMAR.chrX
> 7.00
6.00 - 7.00
5.00 - 6.00
4.00 - 5.00
3.00 - 4.00
2.00 - 3.00
1.00 - 2.00
0.50 - 1.00
0.10 - 0.50
0.00 - 0.10
no SNPs
0 Mbp
50 Mbp
100 Mbp
150 Mbp
200 Mbp
250 Mbp


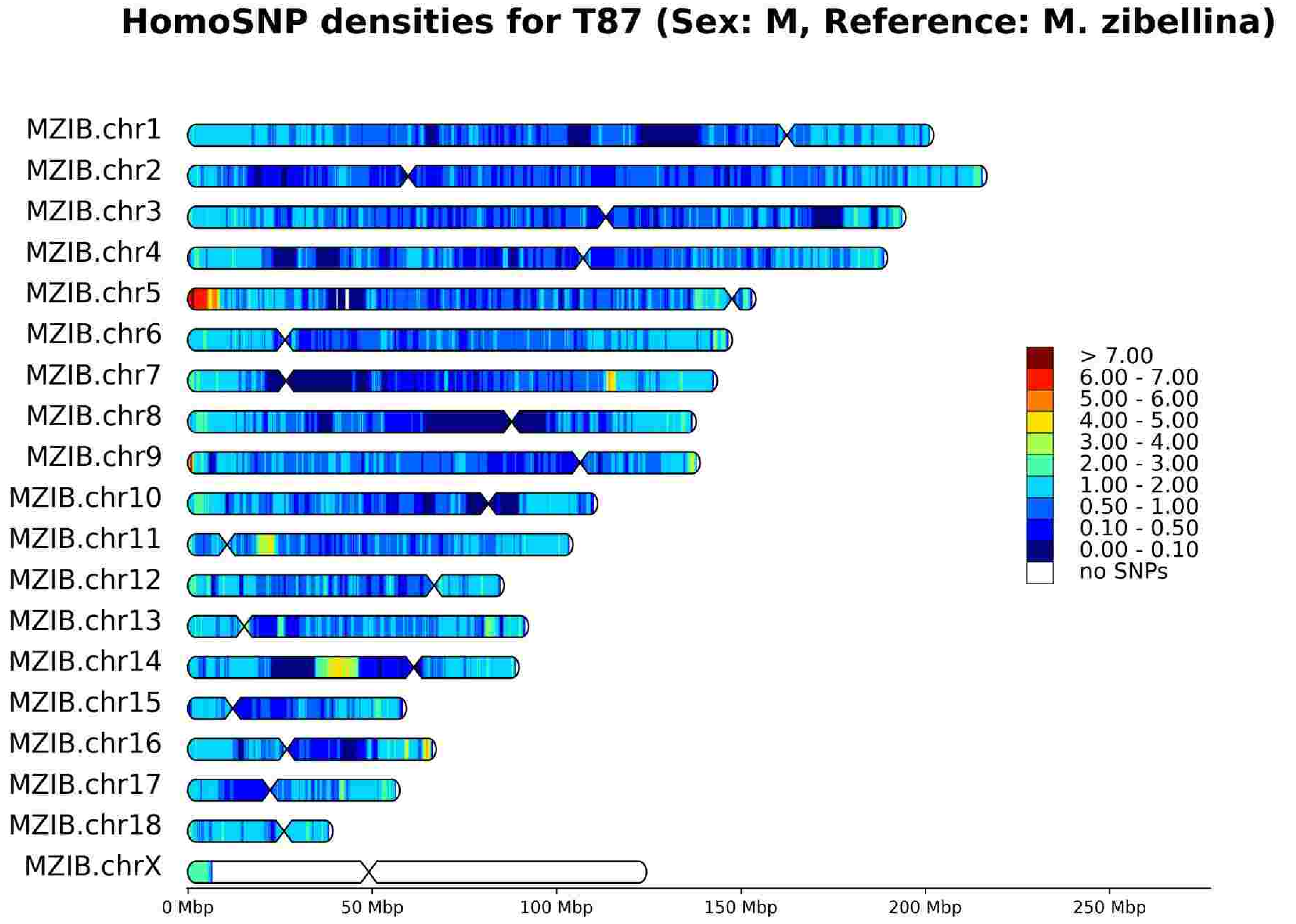

HomoSNP densities for T87 (Sex: M, Reference: M. zibellina)
MZIB.chr1
MZIB.chr2
MZIB.chr3
MZIB.chr4
MZIB.chr5
MZIB.chr6
MZIB.chr7
MZIB.chr8
MZIB.chr9
MZIB.chr10
MZIB.chr11
MZIB.chr12
MZIB.chr13
MZIB.chr14
MZIB.chr15
MZIB.chr16
MZIB.chr17
MZIB.chr18
MZIB.chrX
> 7.00
6.00 - 7.00
5.00 - 6.00
4.00 - 5.00
3.00 - 4.00
2.00 - 3.00
1.00 - 2.00
0.50 - 1.00
0.10 - 0.50
0.00 - 0.10
no SNPs
0 Mbp
50 Mbp
100 Mbp
150 Mbp
200 Mbp
250 Mbp


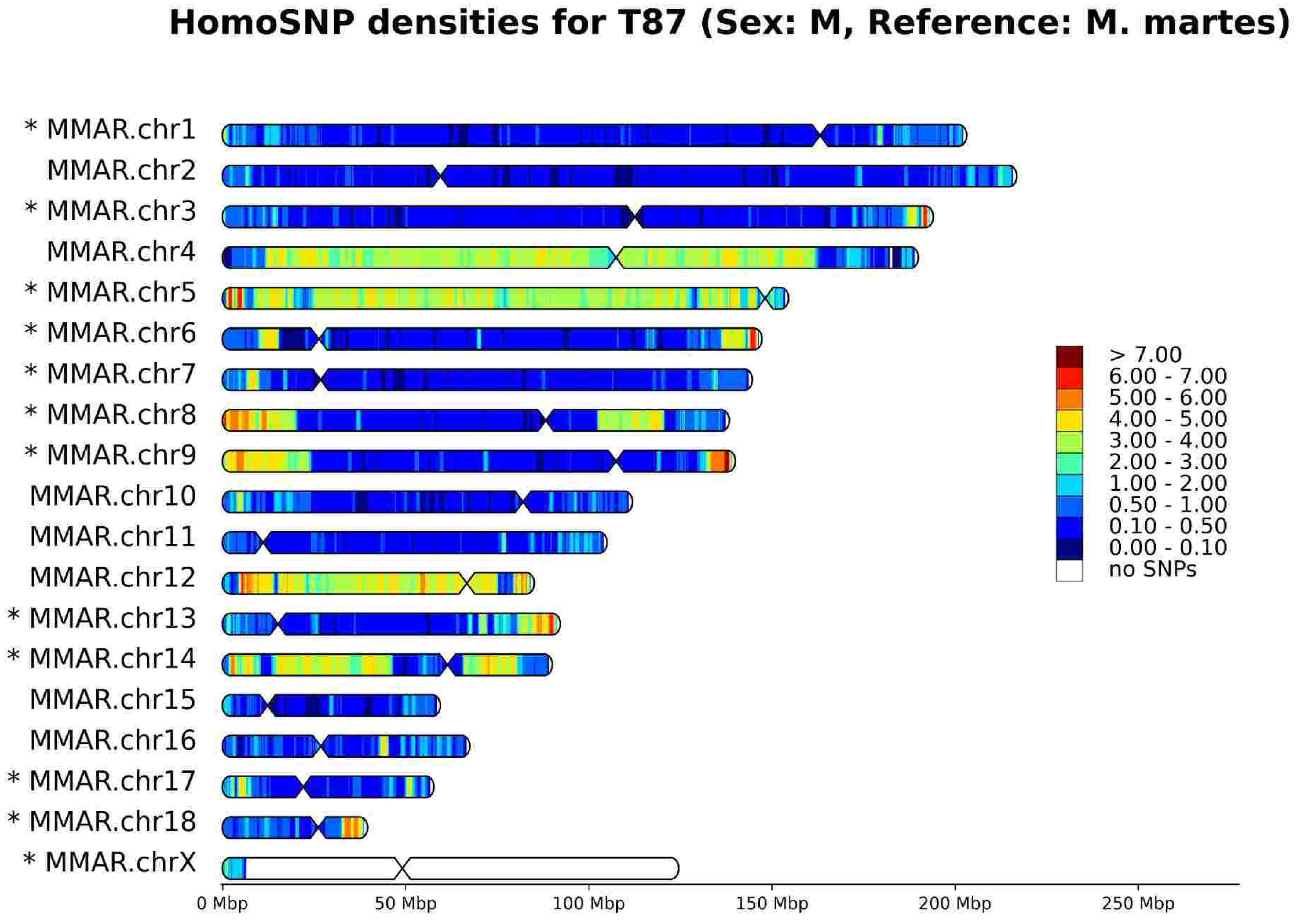

HomoSNP densities for T87 (Sex: M, Reference: M. martes)
* MMAR.chr1
MMAR.chr2
* MMAR.chr3
MMAR.chr4
* MMAR.chr5
* MMAR.chr6
* MMAR.chr7
* MMAR.chr8
* MMAR.chr9
MMAR.chr10
MMAR.chr11
MMAR.chr12
* MMAR.chr13
* MMAR.chr14
MMAR.chr15
MMAR.chr16
* MMAR.chr17
* MMAR.chr18
* MMAR.chrX
> 7.00
6.00 - 7.00
5.00 - 6.00
4.00 - 5.00
3.00 - 4.00
2.00 - 3.00
1.00 - 2.00
0.50 - 1.00
0.10 - 0.50
0.00 - 0.10
no SNPs
0 Mbp
50 Mbp
100 Mbp
150 Mbp
200 Mbp
250 Mbp

# T90

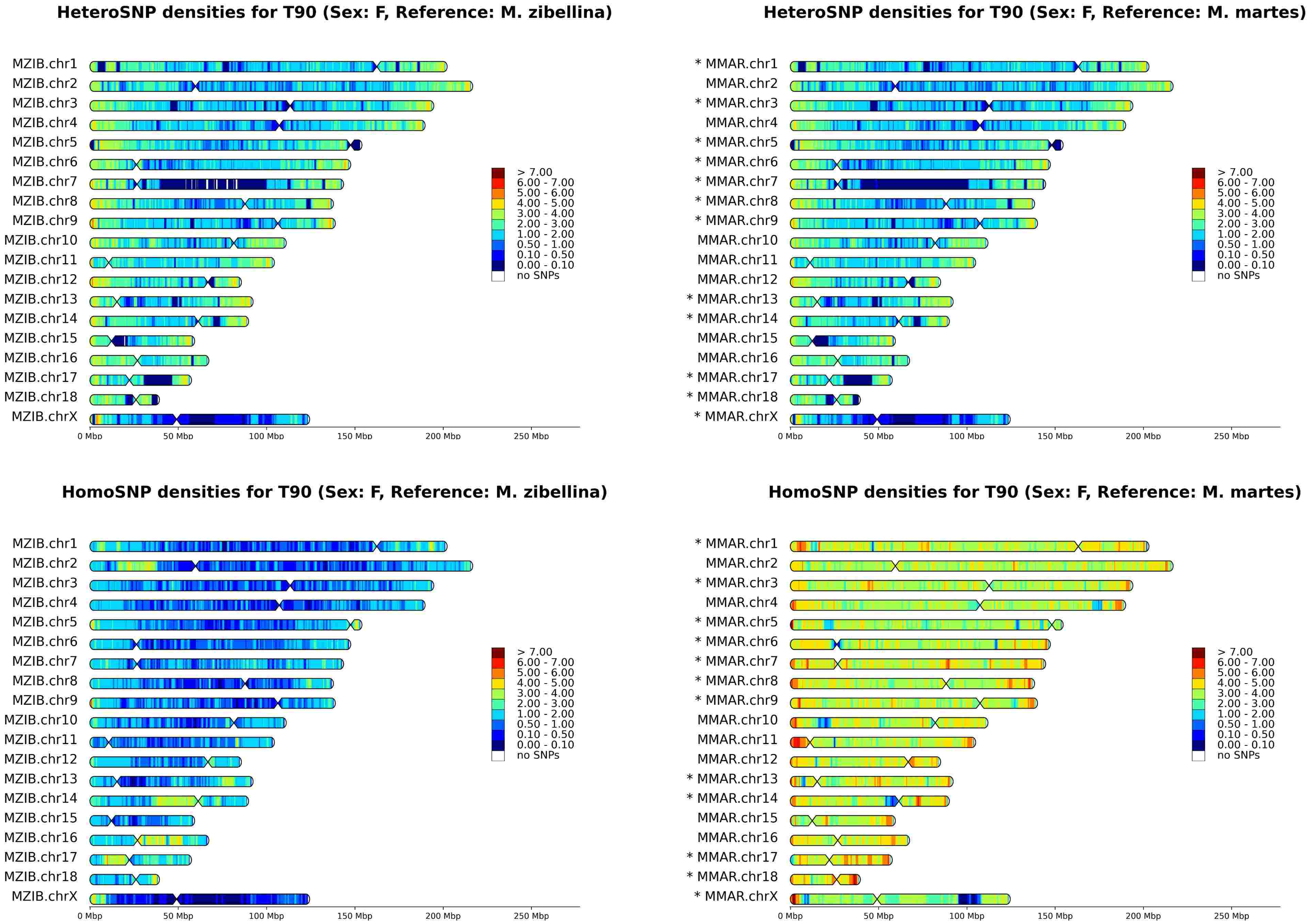
HeteroSNP densities for T90 (Sex: F, Reference: M. zibellina)
MZIB.chr1
MZIB.chr2
MZIB.chr3
MZIB.chr4
MZIB.chr5
MZIB.chr6
MZIB.chr7
MZIB.chr8
MZIB.chr9
MZIB.chr10
MZIB.chr11
MZIB.chr12
MZIB.chr13
MZIB.chr14
MZIB.chr15
MZIB.chr16
MZIB.chr17
MZIB.chr18
MZIB.chrX
0 Mbp
50 Mbp
100 Mbp
150 Mbp
200 Mbp
250 Mbp
> 7.00
6.00 - 7.00
5.00 - 6.00
4.00 - 5.00
3.00 - 4.00
2.00 - 3.00
1.00 - 2.00
0.50 - 1.00
0.10 - 0.50
0.00 - 0.10
no SNPs
HeteroSNP densities for T90 (Sex: F, Reference: M. martes)
* MMAR.chr1
MMAR.chr2
* MMAR.chr3
MMAR.chr4
* MMAR.chr5
* MMAR.chr6
* MMAR.chr7
* MMAR.chr8
* MMAR.chr9
MMAR.chr10
MMAR.chr11
MMAR.chr12
* MMAR.chr13
* MMAR.chr14
MMAR.chr15
MMAR.chr16
* MMAR.chr17
* MMAR.chr18
* MMAR.chrX
HomoSNP densities for T90 (Sex: F, Reference: M. zibellina)
HomoSNP densities for T90 (Sex: F, Reference: M. martes)

# T104

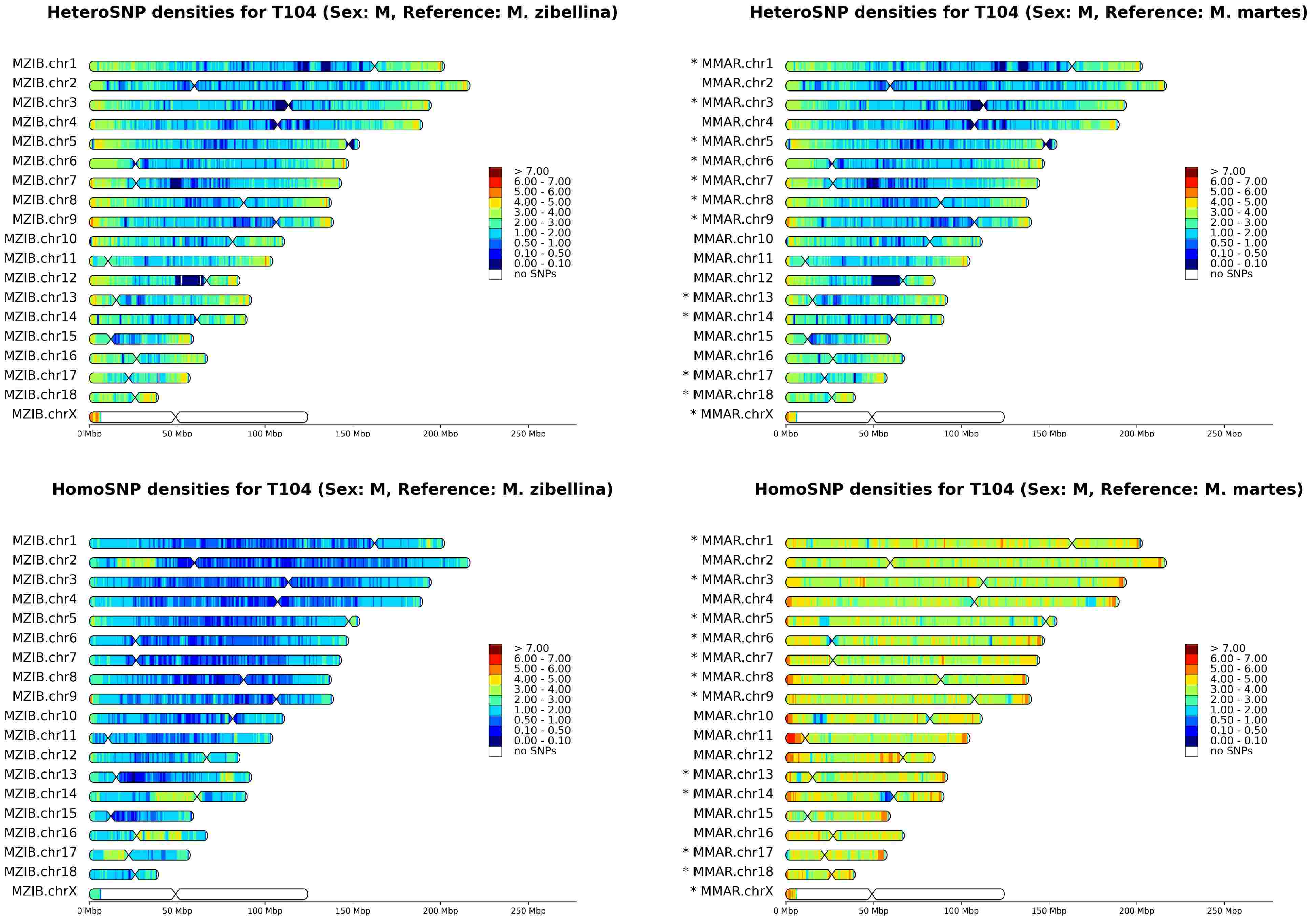
HeteroSNP densities for T104 (Sex: M, Reference: M. zibellina)
MZIB.chr1
MZIB.chr2
MZIB.chr3
MZIB.chr4
MZIB.chr5
MZIB.chr6
MZIB.chr7
MZIB.chr8
MZIB.chr9
MZIB.chr10
MZIB.chr11
MZIB.chr12
MZIB.chr13
MZIB.chr14
MZIB.chr15
MZIB.chr16
MZIB.chr17
MZIB.chr18
MZIB.chrX
0 Mbp
50 Mbp
100 Mbp
150 Mbp
200 Mbp
250 Mbp
> 7.00
6.00 - 7.00
5.00 - 6.00
4.00 - 5.00
3.00 - 4.00
2.00 - 3.00
1.00 - 2.00
0.50 - 1.00
0.10 - 0.50
0.00 - 0.10
no SNPs
HeteroSNP densities for T104 (Sex: M, Reference: M. martes)
* MMAR.chr1
MMAR.chr2
* MMAR.chr3
MMAR.chr4
* MMAR.chr5
* MMAR.chr6
* MMAR.chr7
* MMAR.chr8
* MMAR.chr9
MMAR.chr10
MMAR.chr11
MMAR.chr12
* MMAR.chr13
* MMAR.chr14
MMAR.chr15
MMAR.chr16
* MMAR.chr17
* MMAR.chr18
* MMAR.chrX
HomoSNP densities for T104 (Sex: M, Reference: M. zibellina)
HomoSNP densities for T104 (Sex: M, Reference: M. martes)

# T118

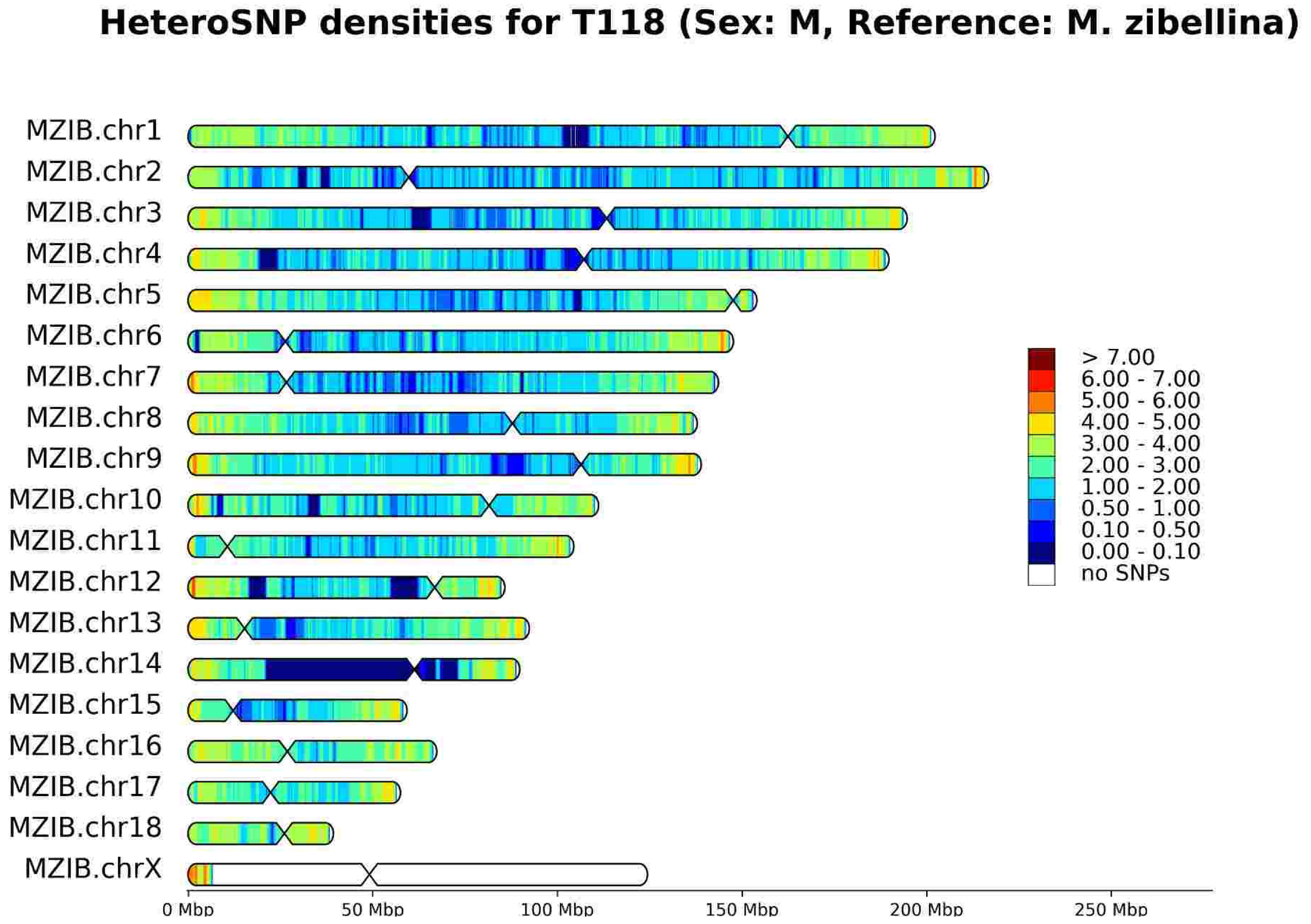
HeteroSNP densities for T118 (Sex: M, Reference: M. zibellina)
MZIB.chr1
MZIB.chr2
MZIB.chr3
MZIB.chr4
MZIB.chr5
MZIB.chr6
MZIB.chr7
MZIB.chr8
MZIB.chr9
MZIB.chr10
MZIB.chr11
MZIB.chr12
MZIB.chr13
MZIB.chr14
MZIB.chr15
MZIB.chr16
MZIB.chr17
MZIB.chr18
MZIB.chrX
0 Mbp
50 Mbp
100 Mbp
150 Mbp
200 Mbp
250 Mbp
> 7.00
6.00 - 7.00
5.00 - 6.00
4.00 - 5.00
3.00 - 4.00
2.00 - 3.00
1.00 - 2.00
0.50 - 1.00
0.10 - 0.50
0.00 - 0.10
no SNPs

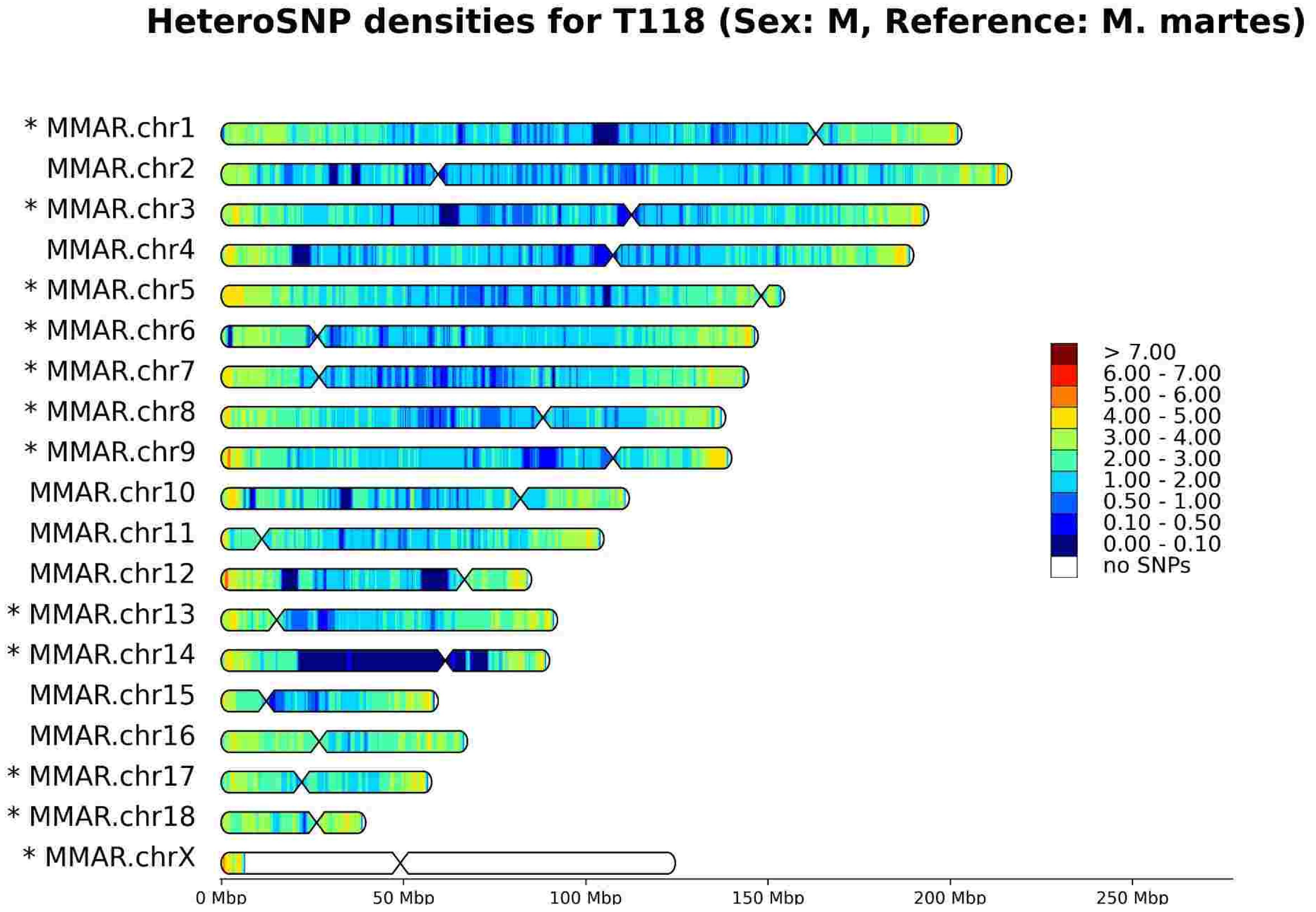
HeteroSNP densities for T118 (Sex: M, Reference: M. martes)
* MMAR.chr1
MMAR.chr2
* MMAR.chr3
MMAR.chr4
* MMAR.chr5
* MMAR.chr6
* MMAR.chr7
* MMAR.chr8
* MMAR.chr9
MMAR.chr10
MMAR.chr11
MMAR.chr12
* MMAR.chr13
* MMAR.chr14
MMAR.chr15
MMAR.chr16
* MMAR.chr17
* MMAR.chr18
* MMAR.chrX
0 Mbp
50 Mbp
100 Mbp
150 Mbp
200 Mbp
250 Mbp
> 7.00
6.00 - 7.00
5.00 - 6.00
4.00 - 5.00
3.00 - 4.00
2.00 - 3.00
1.00 - 2.00
0.50 - 1.00
0.10 - 0.50
0.00 - 0.10
no SNPs

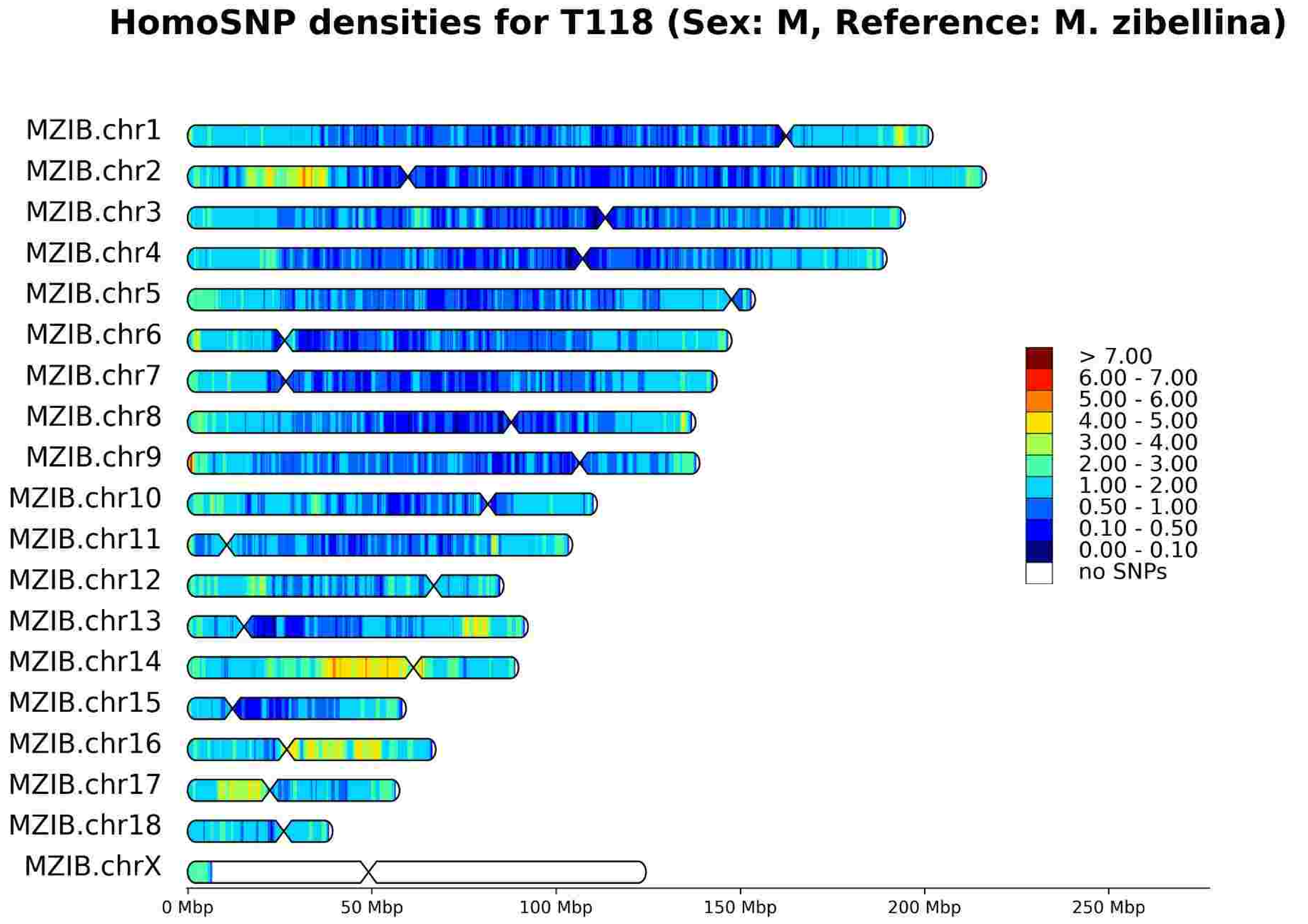
HomoSNP densities for T118 (Sex: M, Reference: M. zibellina)
MZIB.chr1
MZIB.chr2
MZIB.chr3
MZIB.chr4
MZIB.chr5
MZIB.chr6
MZIB.chr7
MZIB.chr8
MZIB.chr9
MZIB.chr10
MZIB.chr11
MZIB.chr12
MZIB.chr13
MZIB.chr14
MZIB.chr15
MZIB.chr16
MZIB.chr17
MZIB.chr18
MZIB.chrX
0 Mbp
50 Mbp
100 Mbp
150 Mbp
200 Mbp
250 Mbp
> 7.00
6.00 - 7.00
5.00 - 6.00
4.00 - 5.00
3.00 - 4.00
2.00 - 3.00
1.00 - 2.00
0.50 - 1.00
0.10 - 0.50
0.00 - 0.10
no SNPs

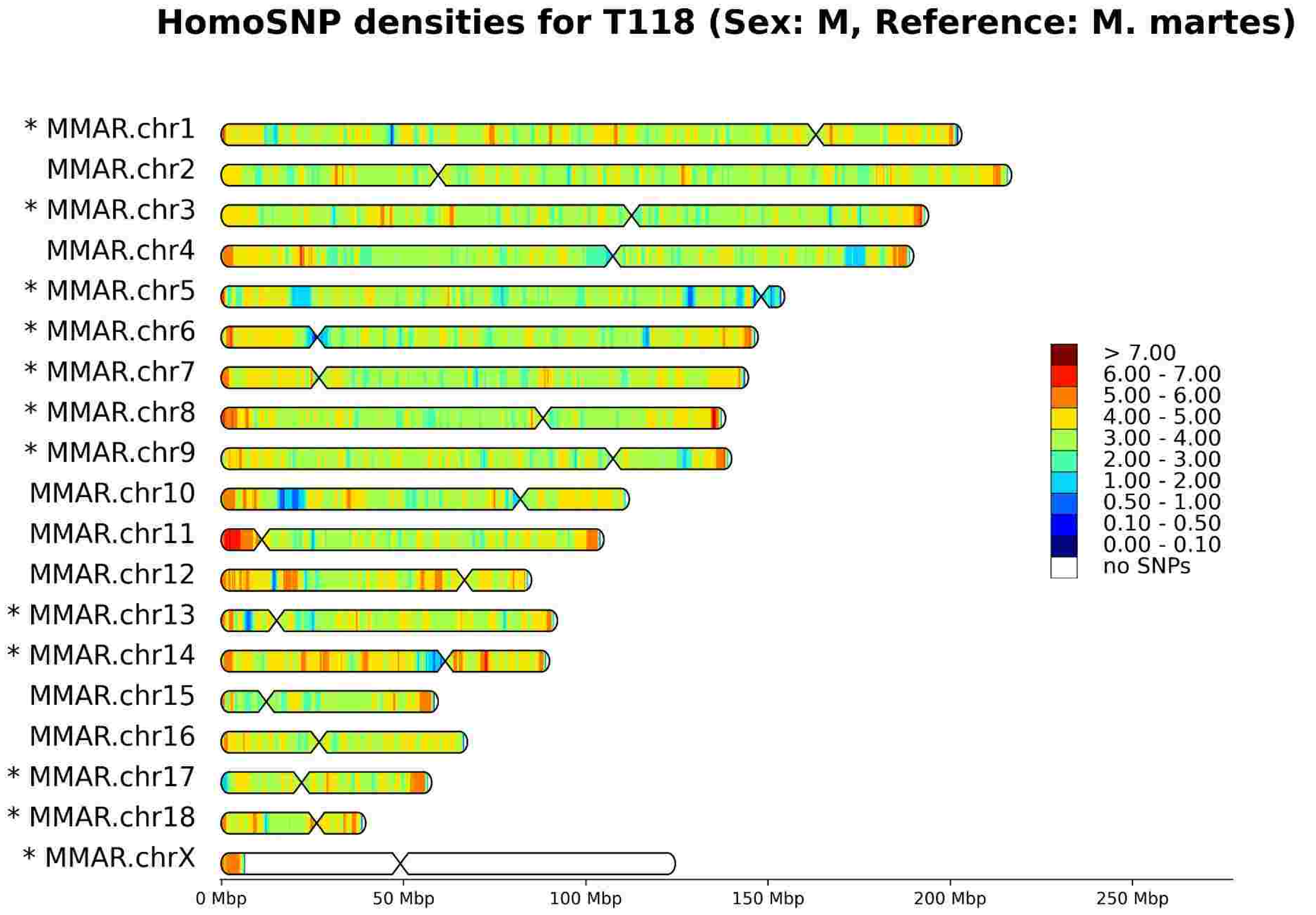
HomoSNP densities for T118 (Sex: M, Reference: M. martes)
* MMAR.chr1
MMAR.chr2
* MMAR.chr3
MMAR.chr4
* MMAR.chr5
* MMAR.chr6
* MMAR.chr7
* MMAR.chr8
* MMAR.chr9
MMAR.chr10
MMAR.chr11
MMAR.chr12
* MMAR.chr13
* MMAR.chr14
MMAR.chr15
MMAR.chr16
* MMAR.chr17
* MMAR.chr18
* MMAR.chrX
0 Mbp
50 Mbp
100 Mbp
150 Mbp
200 Mbp
250 Mbp
> 7.00
6.00 - 7.00
5.00 - 6.00
4.00 - 5.00
3.00 - 4.00
2.00 - 3.00
1.00 - 2.00
0.50 - 1.00
0.10 - 0.50
0.00 - 0.10
no SNPs

# T148

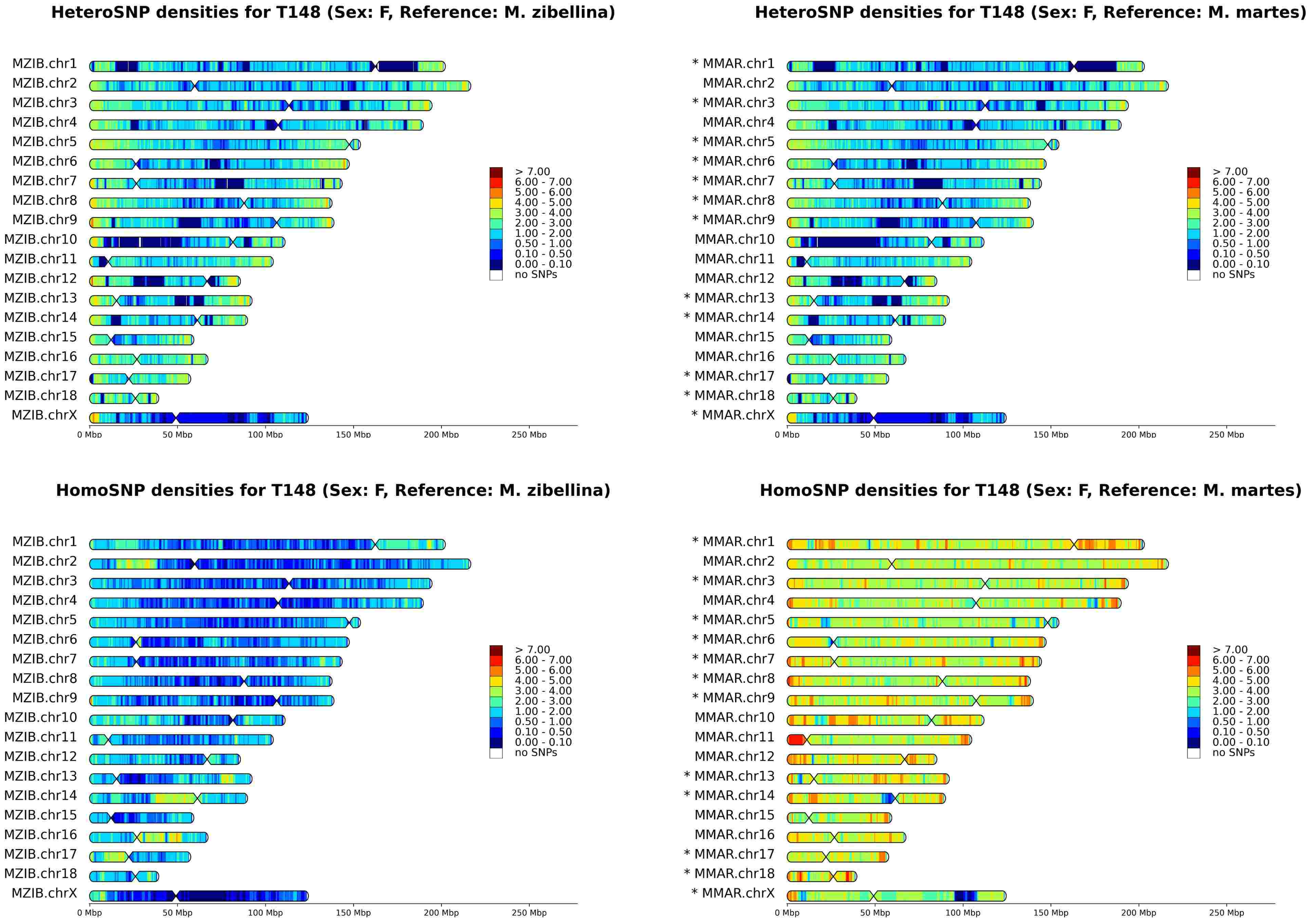
HeteroSNP densities for T148 (Sex: F, Reference: M. zibellina)
HeteroSNP densities for T148 (Sex: F, Reference: M. martes)
HomoSNP densities for T148 (Sex: F, Reference: M. zibellina)
HomoSNP densities for T148 (Sex: F, Reference: M. martes)
MZIB.chr1
MZIB.chr2
MZIB.chr3
MZIB.chr4
MZIB.chr5
MZIB.chr6
MZIB.chr7
MZIB.chr8
MZIB.chr9
MZIB.chr10
MZIB.chr11
MZIB.chr12
MZIB.chr13
MZIB.chr14
MZIB.chr15
MZIB.chr16
MZIB.chr17
MZIB.chr18
MZIB.chrX
* MMAR.chr1
MMAR.chr2
* MMAR.chr3
MMAR.chr4
* MMAR.chr5
* MMAR.chr6
* MMAR.chr7
* MMAR.chr8
* MMAR.chr9
MMAR.chr10
MMAR.chr11
MMAR.chr12
* MMAR.chr13
* MMAR.chr14
MMAR.chr15
MMAR.chr16
* MMAR.chr17
* MMAR.chr18
* MMAR.chrX
> 7.00
6.00 - 7.00
5.00 - 6.00
4.00 - 5.00
3.00 - 4.00
2.00 - 3.00
1.00 - 2.00
0.50 - 1.00
0.10 - 0.50
0.00 - 0.10
no SNPs
0 Mbp
50 Mbp
100 Mbp
150 Mbp
200 Mbp
250 Mbp

# T149

**HeteroSNP densities for T149 (Sex: M, Reference: M. zibellina)**

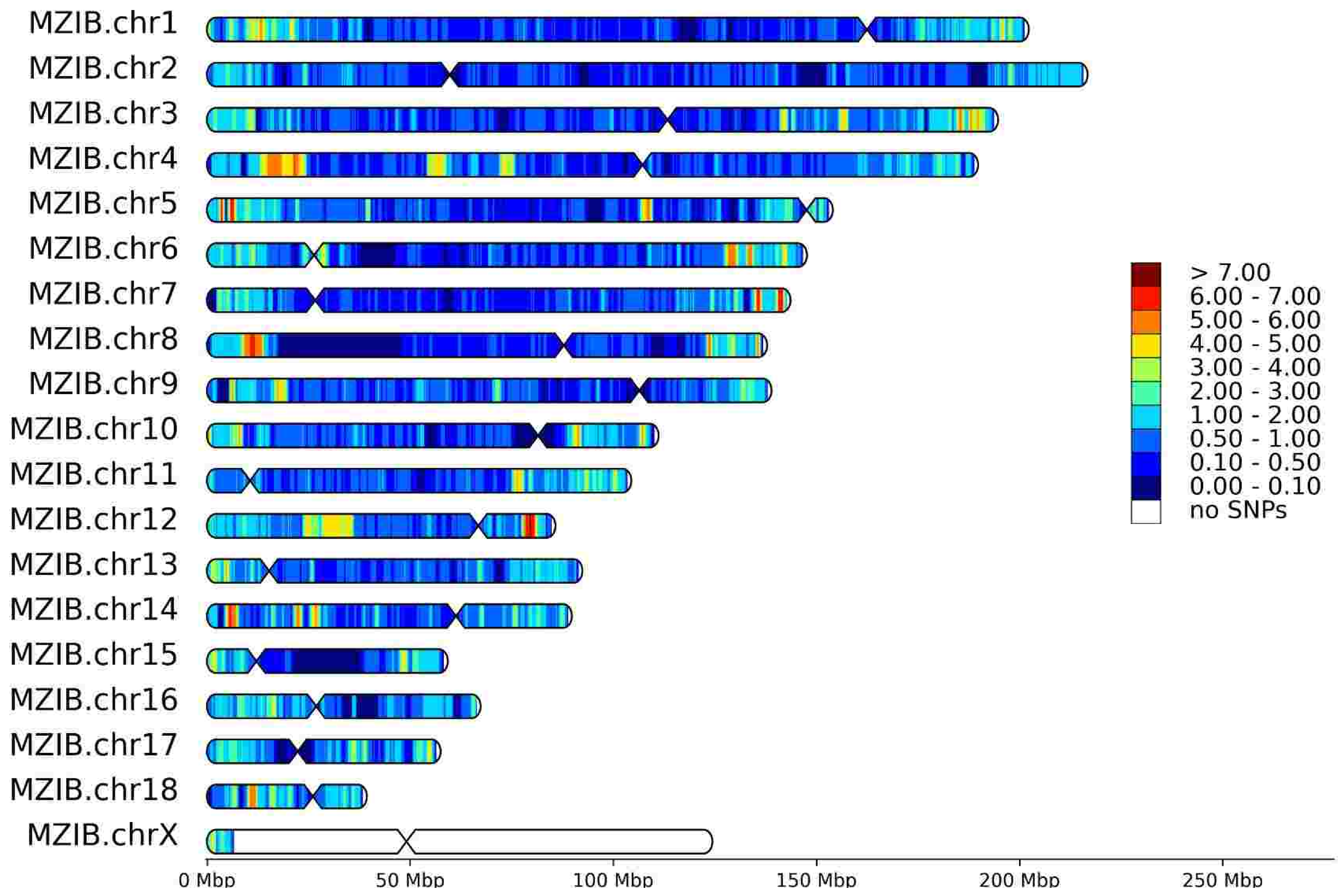


**HeteroSNP densities for T149 (Sex: M, Reference: M. martes)**

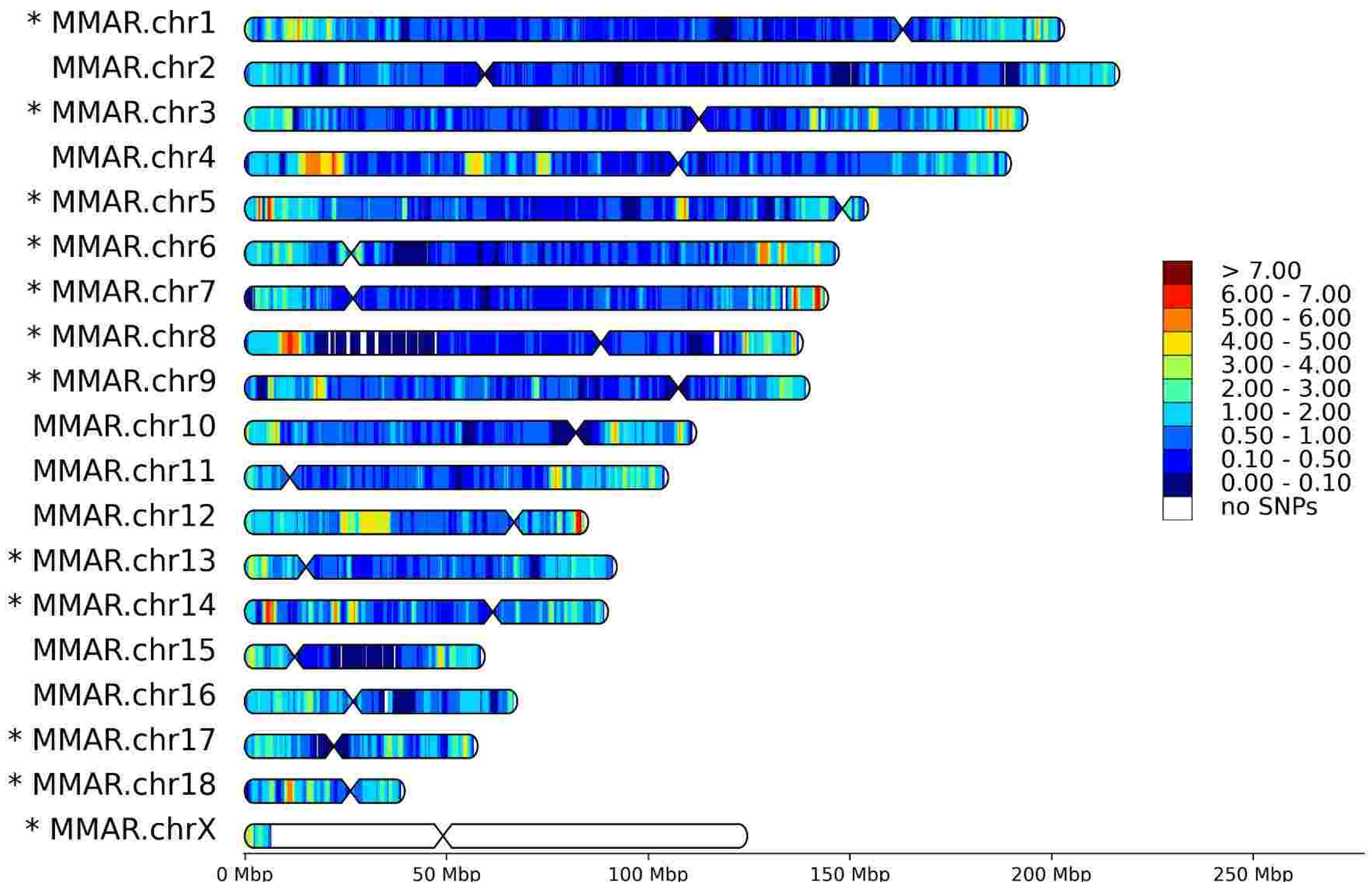


**HomoSNP densities for T149 (Sex: M, Reference: M. zibellina)**

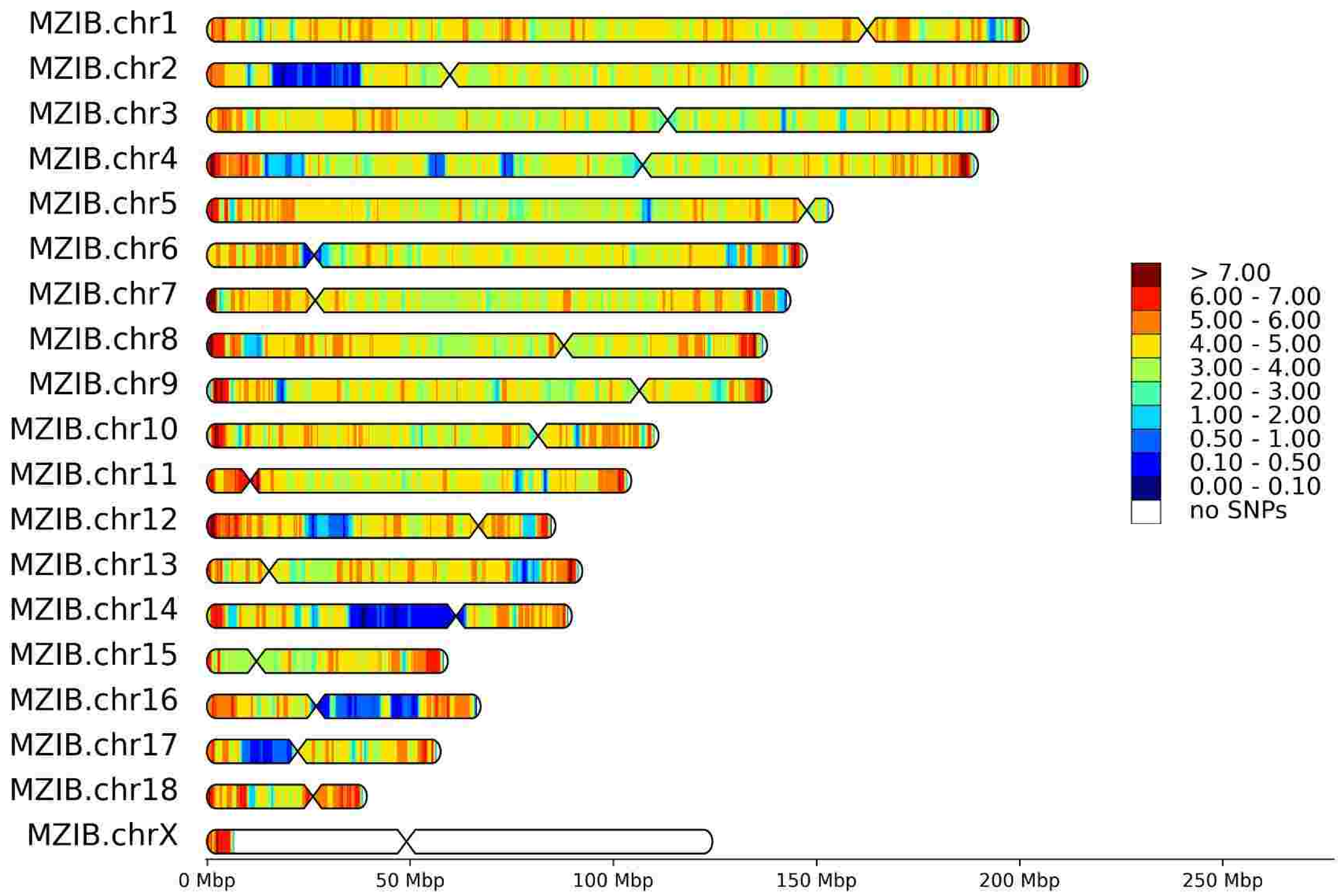


**HomoSNP densities for T149 (Sex: M, Reference: M. martes)**

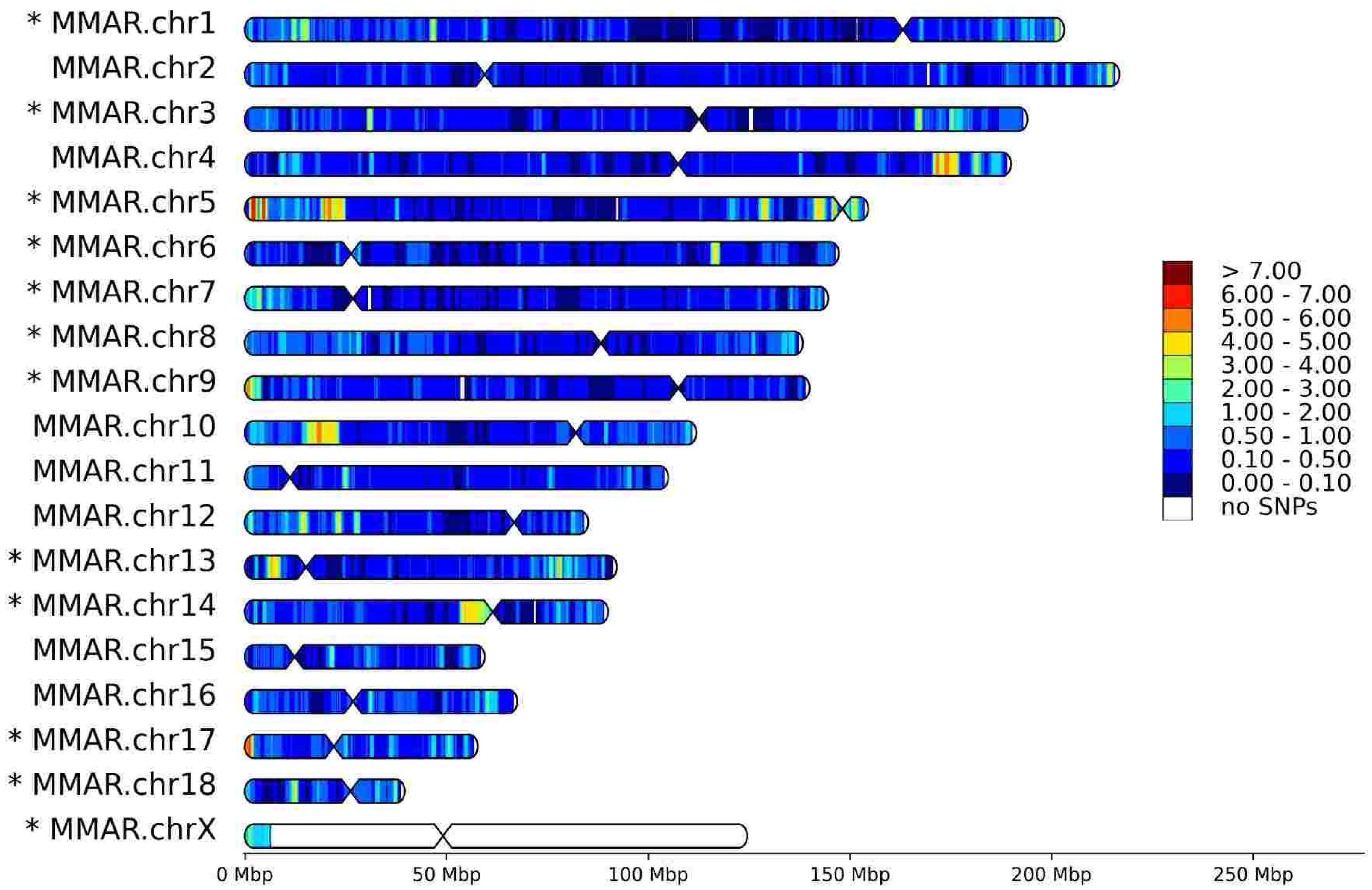

# T150

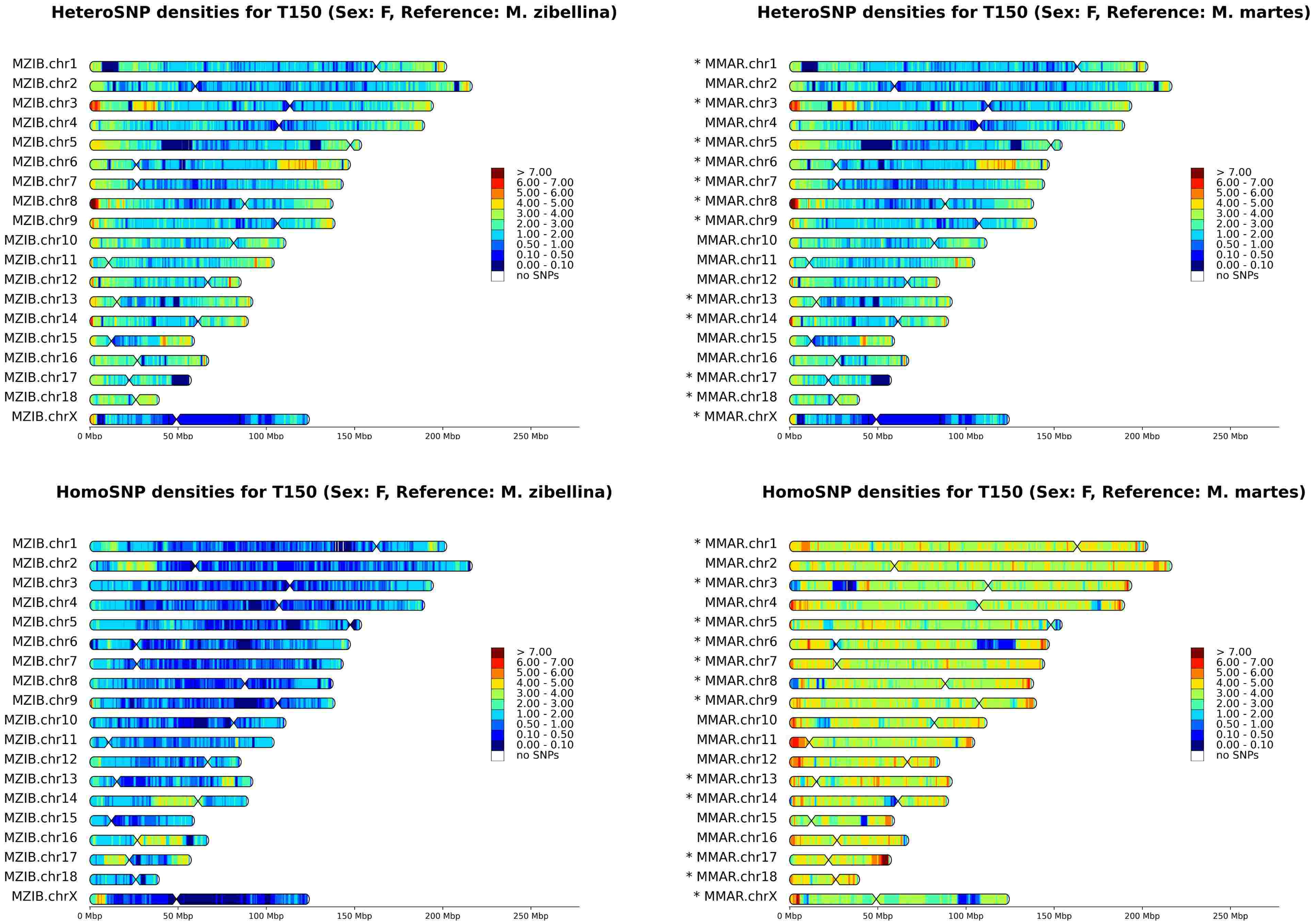
HeteroSNP densities for T150 (Sex: F, Reference: M. zibellina)
MZIB.chr1
MZIB.chr2
MZIB.chr3
MZIB.chr4
MZIB.chr5
MZIB.chr6
MZIB.chr7
MZIB.chr8
MZIB.chr9
MZIB.chr10
MZIB.chr11
MZIB.chr12
MZIB.chr13
MZIB.chr14
MZIB.chr15
MZIB.chr16
MZIB.chr17
MZIB.chr18
MZIB.chrX
0 Mbp
50 Mbp
100 Mbp
150 Mbp
200 Mbp
250 Mbp
> 7.00
6.00 - 7.00
5.00 - 6.00
4.00 - 5.00
3.00 - 4.00
2.00 - 3.00
1.00 - 2.00
0.50 - 1.00
0.10 - 0.50
0.00 - 0.10
no SNPs
HeteroSNP densities for T150 (Sex: F, Reference: M. martes)
* MMAR.chr1
MMAR.chr2
* MMAR.chr3
MMAR.chr4
* MMAR.chr5
* MMAR.chr6
* MMAR.chr7
* MMAR.chr8
* MMAR.chr9
MMAR.chr10
MMAR.chr11
MMAR.chr12
* MMAR.chr13
* MMAR.chr14
MMAR.chr15
MMAR.chr16
* MMAR.chr17
* MMAR.chr18
* MMAR.chrX
HomoSNP densities for T150 (Sex: F, Reference: M. zibellina)
HomoSNP densities for T150 (Sex: F, Reference: M. martes)

# T151

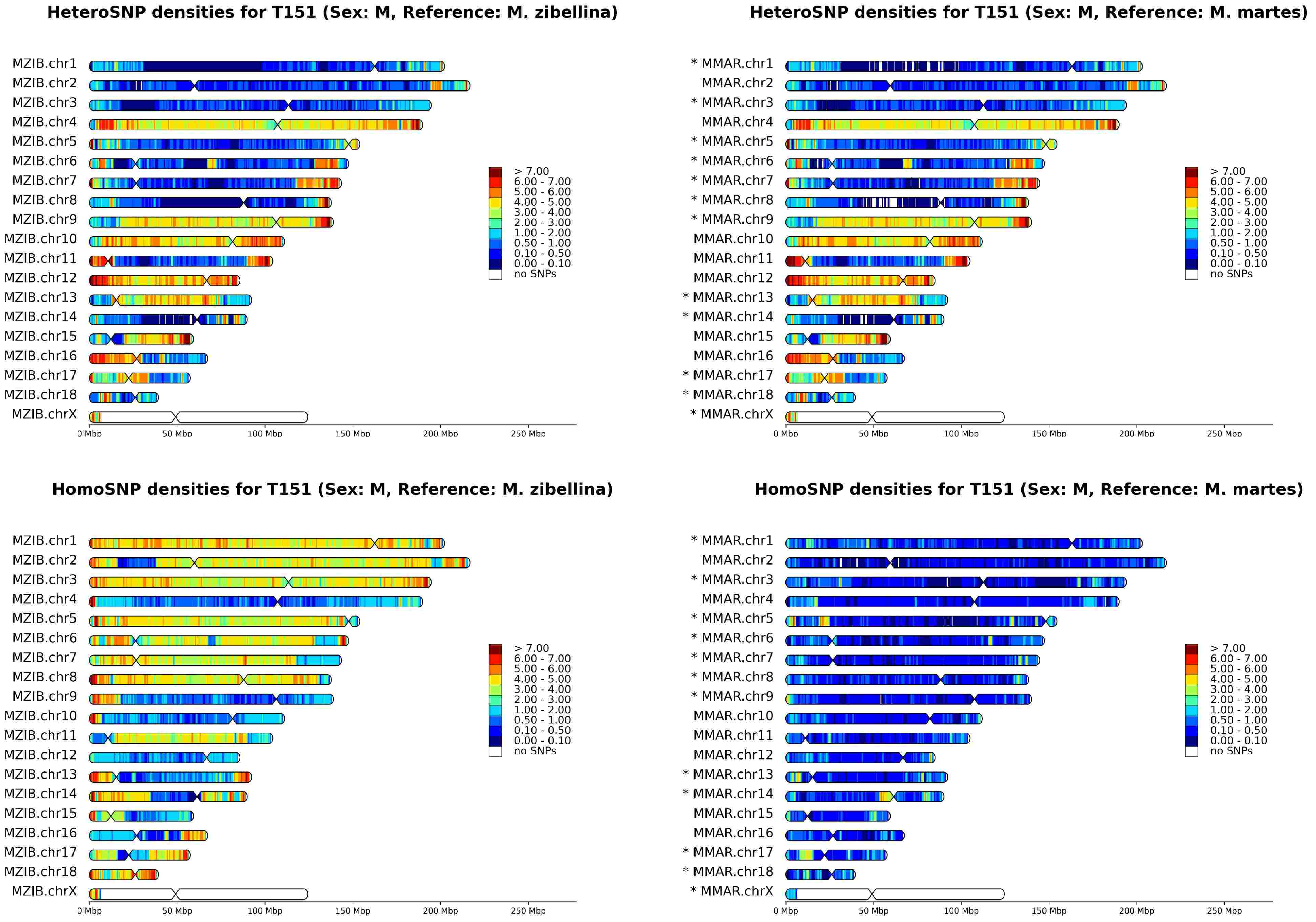
HeteroSNP densities for T151 (Sex: M, Reference: M. zibellina)
HeteroSNP densities for T151 (Sex: M, Reference: M. martes)
HomoSNP densities for T151 (Sex: M, Reference: M. zibellina)
HomoSNP densities for T151 (Sex: M, Reference: M. martes)
MZIB.chr1
MZIB.chr2
MZIB.chr3
MZIB.chr4
MZIB.chr5
MZIB.chr6
MZIB.chr7
MZIB.chr8
MZIB.chr9
MZIB.chr10
MZIB.chr11
MZIB.chr12
MZIB.chr13
MZIB.chr14
MZIB.chr15
MZIB.chr16
MZIB.chr17
MZIB.chr18
MZIB.chrX
* MMAR.chr1
MMAR.chr2
* MMAR.chr3
MMAR.chr4
* MMAR.chr5
* MMAR.chr6
* MMAR.chr7
* MMAR.chr8
* MMAR.chr9
MMAR.chr10
MMAR.chr11
MMAR.chr12
* MMAR.chr13
* MMAR.chr14
MMAR.chr15
MMAR.chr16
* MMAR.chr17
* MMAR.chr18
* MMAR.chrX
0 Mbp
50 Mbp
100 Mbp
150 Mbp
200 Mbp
250 Mbp
> 7.00
6.00 - 7.00
5.00 - 6.00
4.00 - 5.00
3.00 - 4.00
2.00 - 3.00
1.00 - 2.00
0.50 - 1.00
0.10 - 0.50
0.00 - 0.10
no SNPs

# T194

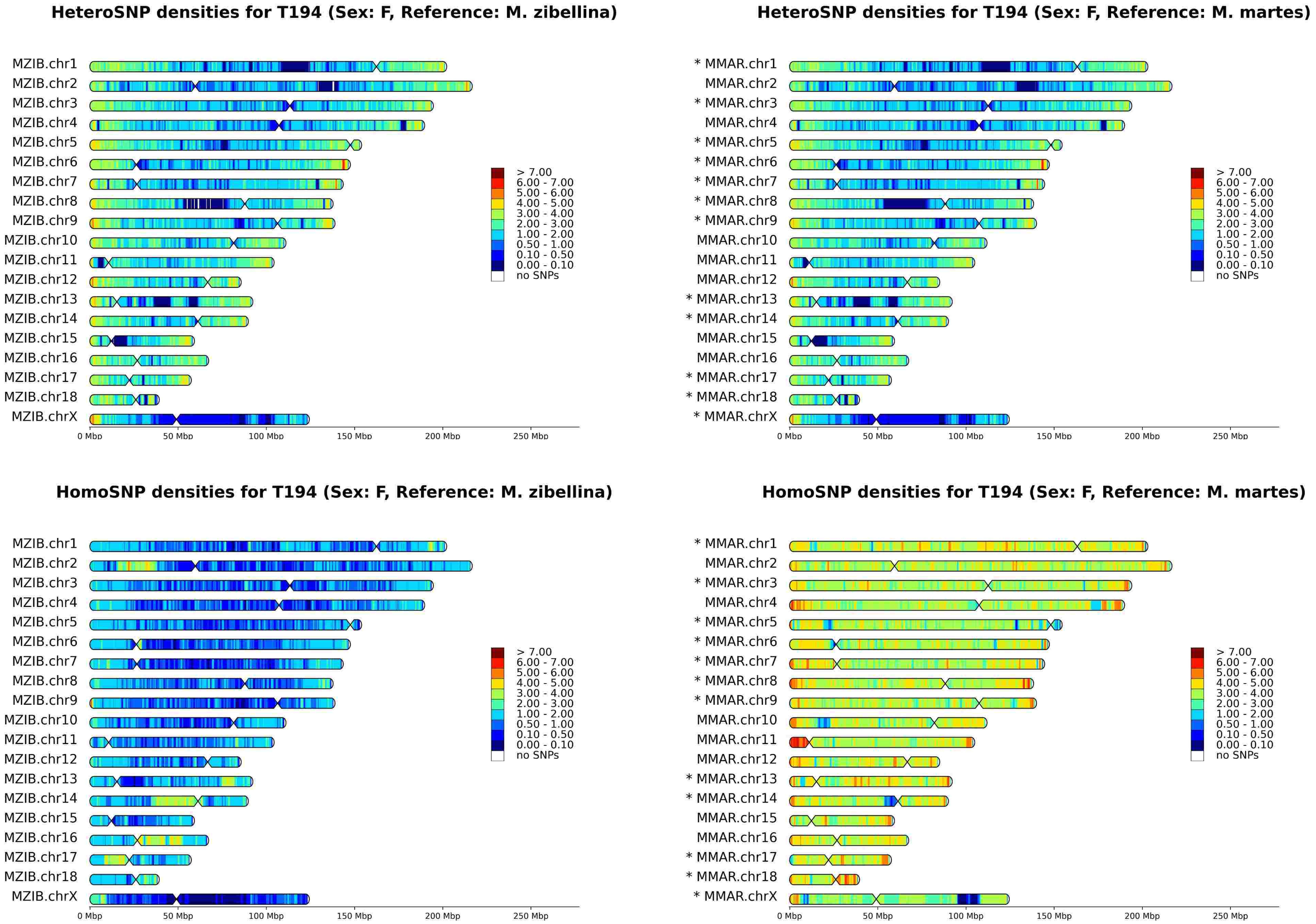
HeteroSNP densities for T194 (Sex: F, Reference: M. zibellina)
HeteroSNP densities for T194 (Sex: F, Reference: M. martes)
HomoSNP densities for T194 (Sex: F, Reference: M. zibellina)
HomoSNP densities for T194 (Sex: F, Reference: M. martes)
MZIB.chr1
MZIB.chr2
MZIB.chr3
MZIB.chr4
MZIB.chr5
MZIB.chr6
MZIB.chr7
MZIB.chr8
MZIB.chr9
MZIB.chr10
MZIB.chr11
MZIB.chr12
MZIB.chr13
MZIB.chr14
MZIB.chr15
MZIB.chr16
MZIB.chr17
MZIB.chr18
MZIB.chrX
* MMAR.chr1
MMAR.chr2
* MMAR.chr3
MMAR.chr4
* MMAR.chr5
* MMAR.chr6
* MMAR.chr7
* MMAR.chr8
* MMAR.chr9
MMAR.chr10
MMAR.chr11
MMAR.chr12
* MMAR.chr13
* MMAR.chr14
MMAR.chr15
MMAR.chr16
* MMAR.chr17
* MMAR.chr18
* MMAR.chrX
> 7.00
6.00 - 7.00
5.00 - 6.00
4.00 - 5.00
3.00 - 4.00
2.00 - 3.00
1.00 - 2.00
0.50 - 1.00
0.10 - 0.50
0.00 - 0.10
no SNPs
0 Mbp
50 Mbp
100 Mbp
150 Mbp
200 Mbp
250 Mbp
