## Supplementary File SF10 for "Genomics of sable (*Martes zibellina)* × pine marten (*Martes martes*) hybridization"

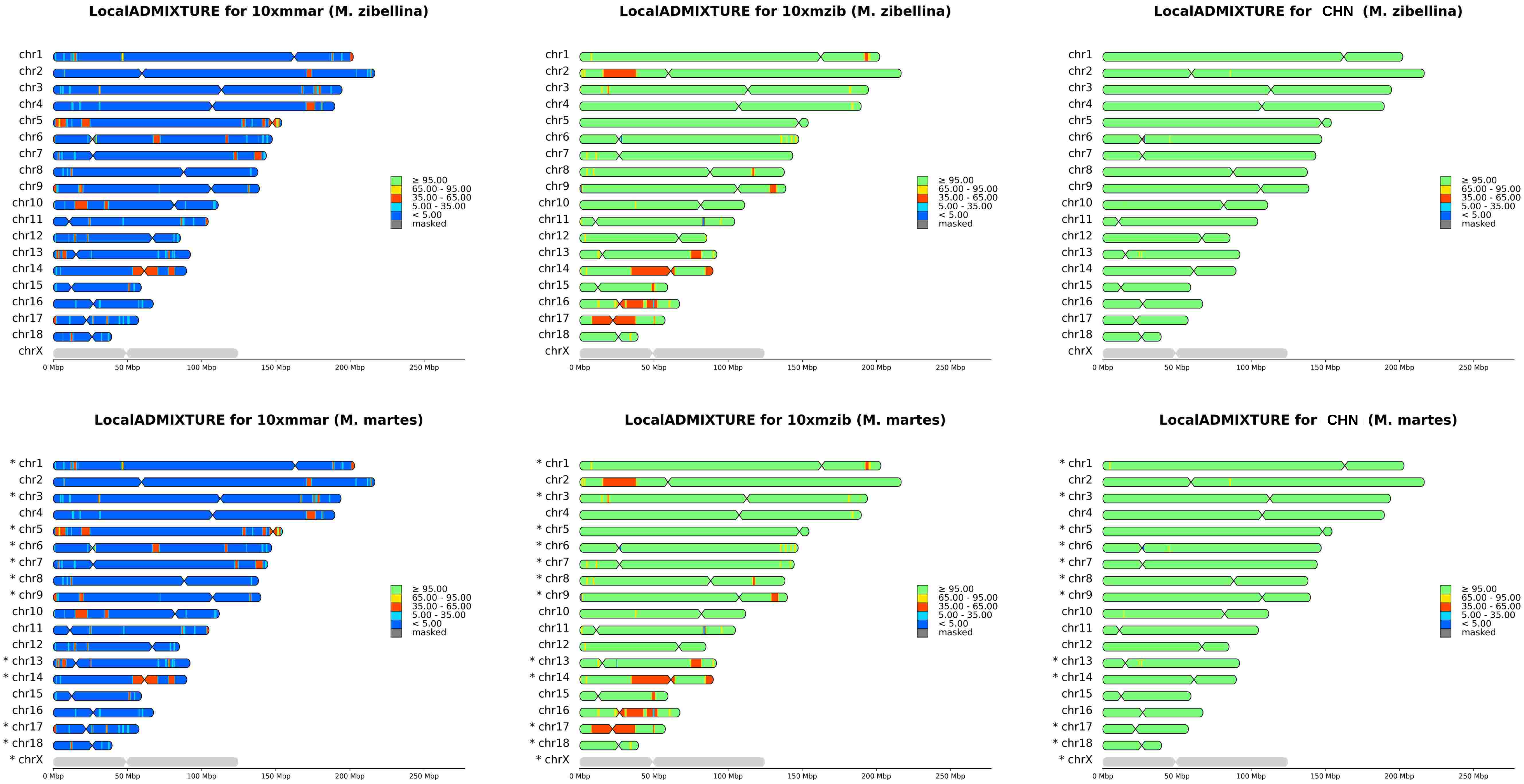
LocalADMIXTURE for 10xmmar (M. zibellina)
LocalADMIXTURE for 10xmzib (M. zibellina)
LocalADMIXTURE for CHN (M. zibellina)
LocalADMIXTURE for 10xmmar (M. martes)
LocalADMIXTURE for 10xmzib (M. martes)
LocalADMIXTURE for CHN (M. martes)
chr1
chr2
chr3
chr4
chr5
chr6
chr7
chr8
chr9
chr10
chr11
chr12
chr13
chr14
chr15
chr16
chr17
chr18
chrX
* chr1
* chr3
* chr5
* chr6
* chr7
* chr8
* chr9
* chr13
* chr14
* chr17
* chr18
* chrX
≥ 95.00
65.00 - 95.00
35.00 - 65.00
5.00 - 35.00
< 5.00
masked
0 Mbp
50 Mbp
100 Mbp
150 Mbp
200 Mbp
250 Mbp

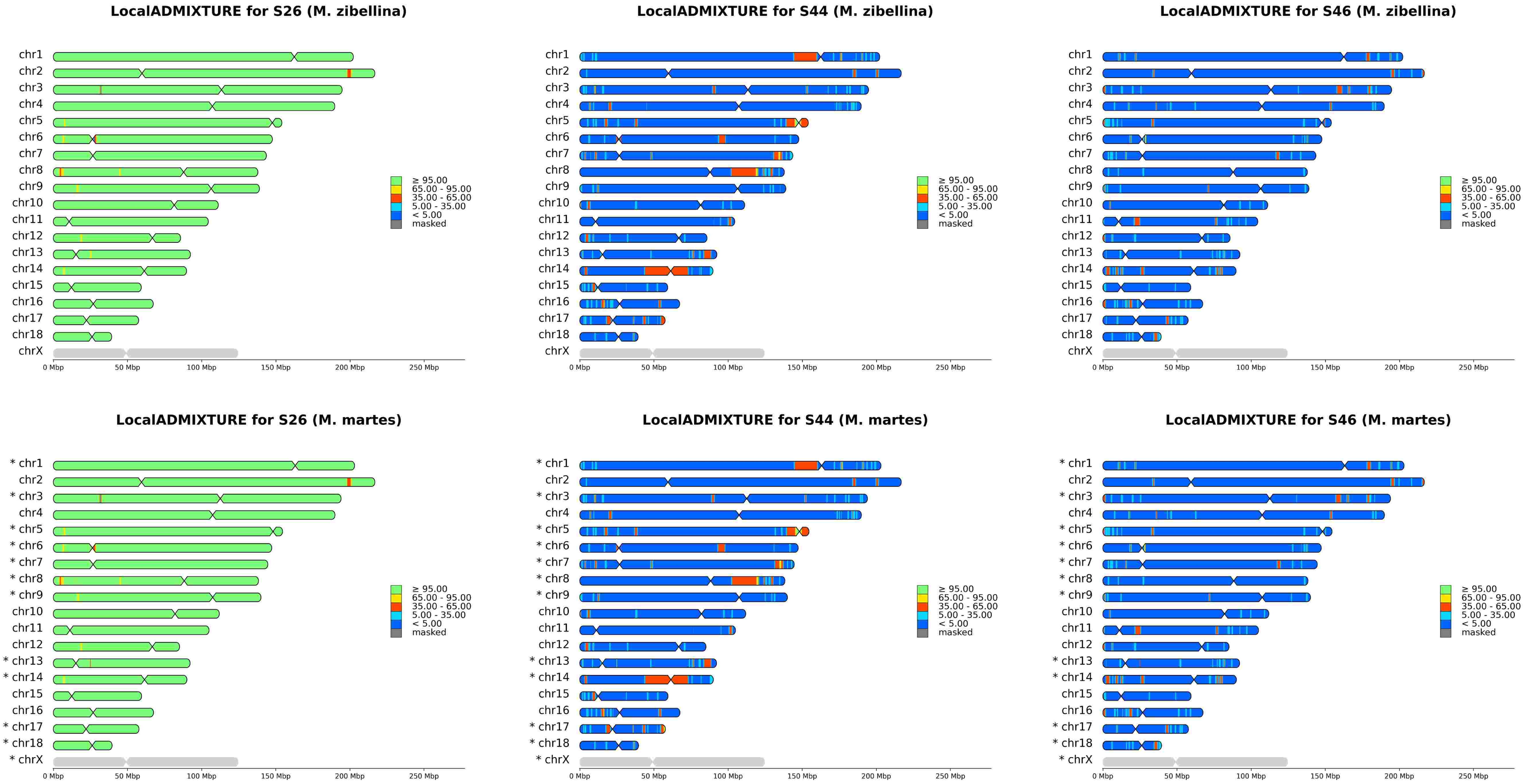
LocalADMIXTURE for S26 (M. zibellina)
LocalADMIXTURE for S44 (M. zibellina)
LocalADMIXTURE for S46 (M. zibellina)
chr1
chr2
chr3
chr4
chr5
chr6
chr7
chr8
chr9
chr10
chr11
chr12
chr13
chr14
chr15
chr16
chr17
chr18
chrX
≥ 95.00
65.00 - 95.00
35.00 - 65.00
5.00 - 35.00
< 5.00
masked
0 Mbp
50 Mbp
100 Mbp
150 Mbp
200 Mbp
250 Mbp
LocalADMIXTURE for S26 (M. martes)
LocalADMIXTURE for S44 (M. martes)
LocalADMIXTURE for S46 (M. martes)
* chr1
chr2
* chr3
chr4
* chr5
* chr6
* chr7
* chr8
* chr9
chr10
chr11
chr12
* chr13
* chr14
chr15
chr16
* chr17
* chr18
* chrX

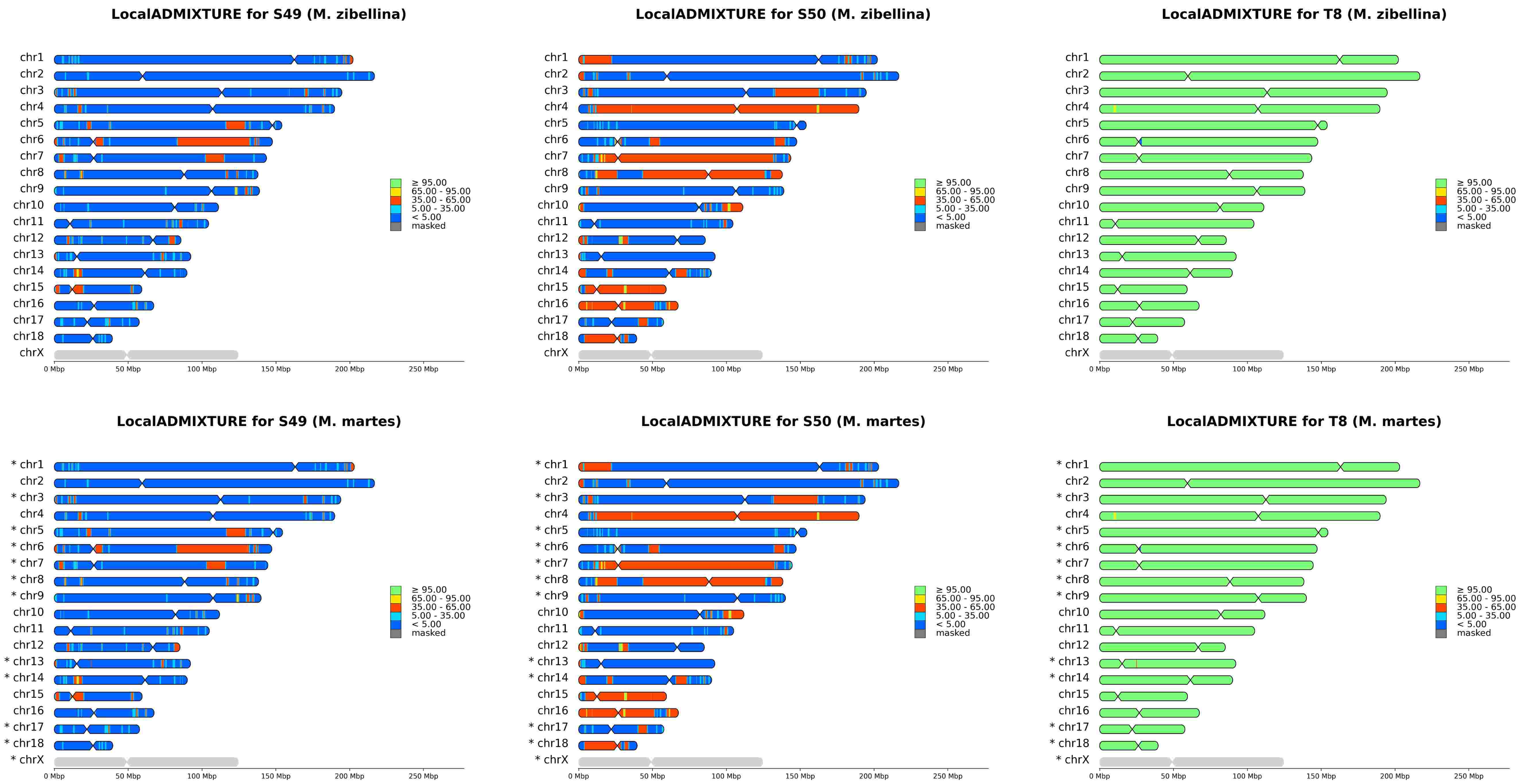

LocalADMIXTURE for S49 (M. zibellina)
LocalADMIXTURE for S50 (M. zibellina)
LocalADMIXTURE for T8 (M. zibellina)
chr1
chr2
chr3
chr4
chr5
chr6
chr7
chr8
chr9
chr10
chr11
chr12
chr13
chr14
chr15
chr16
chr17
chr18
chrX
≥ 95.00
65.00 - 95.00
35.00 - 65.00
5.00 - 35.00
< 5.00
masked
0 Mbp
50 Mbp
100 Mbp
150 Mbp
200 Mbp
250 Mbp
LocalADMIXTURE for S49 (M. martes)
LocalADMIXTURE for S50 (M. martes)
LocalADMIXTURE for T8 (M. martes)
* chr1
chr2
* chr3
chr4
* chr5
* chr6
* chr7
* chr8
* chr9
chr10
chr11
chr12
* chr13
* chr14
chr15
chr16
* chr17
* chr18
* chrX
≥ 95.00
65.00 - 95.00
35.00 - 65.00
5.00 - 35.00
< 5.00
masked
0 Mbp
50 Mbp
100 Mbp
150 Mbp
200 Mbp
250 Mbp

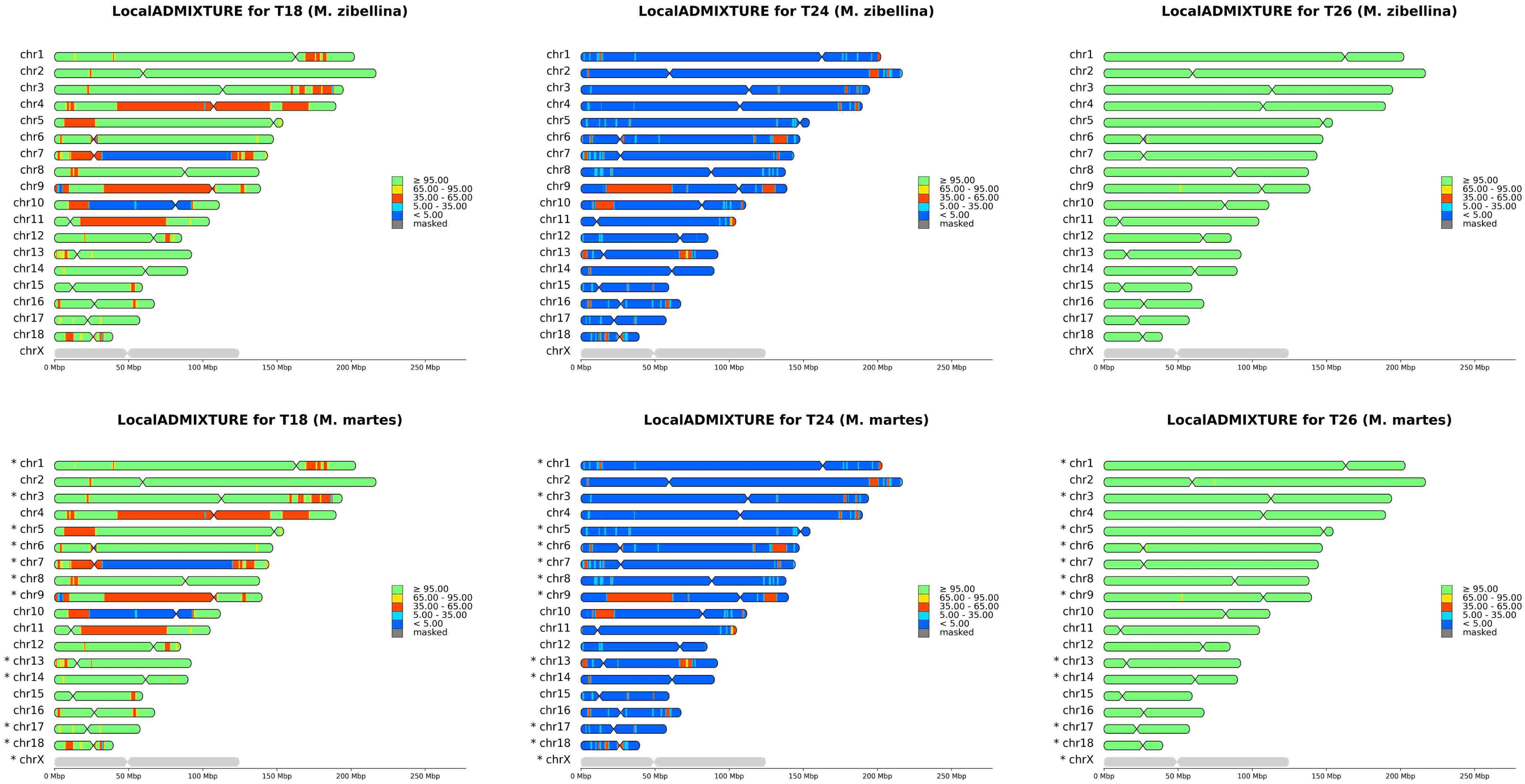
LocalADMIXTURE for T18 (M. zibellina)
LocalADMIXTURE for T24 (M. zibellina)
LocalADMIXTURE for T26 (M. zibellina)
chr1
chr2
chr3
chr4
chr5
chr6
chr7
chr8
chr9
chr10
chr11
chr12
chr13
chr14
chr15
chr16
chr17
chr18
chrX
≥ 95.00
65.00 - 95.00
35.00 - 65.00
5.00 - 35.00
< 5.00
masked
0 Mbp
50 Mbp
100 Mbp
150 Mbp
200 Mbp
250 Mbp
LocalADMIXTURE for T18 (M. martes)
LocalADMIXTURE for T24 (M. martes)
LocalADMIXTURE for T26 (M. martes)
* chr1
chr2
* chr3
chr4
* chr5
* chr6
* chr7
* chr8
* chr9
chr10
chr11
chr12
* chr13
* chr14
chr15
chr16
* chr17
* chr18
* chrX

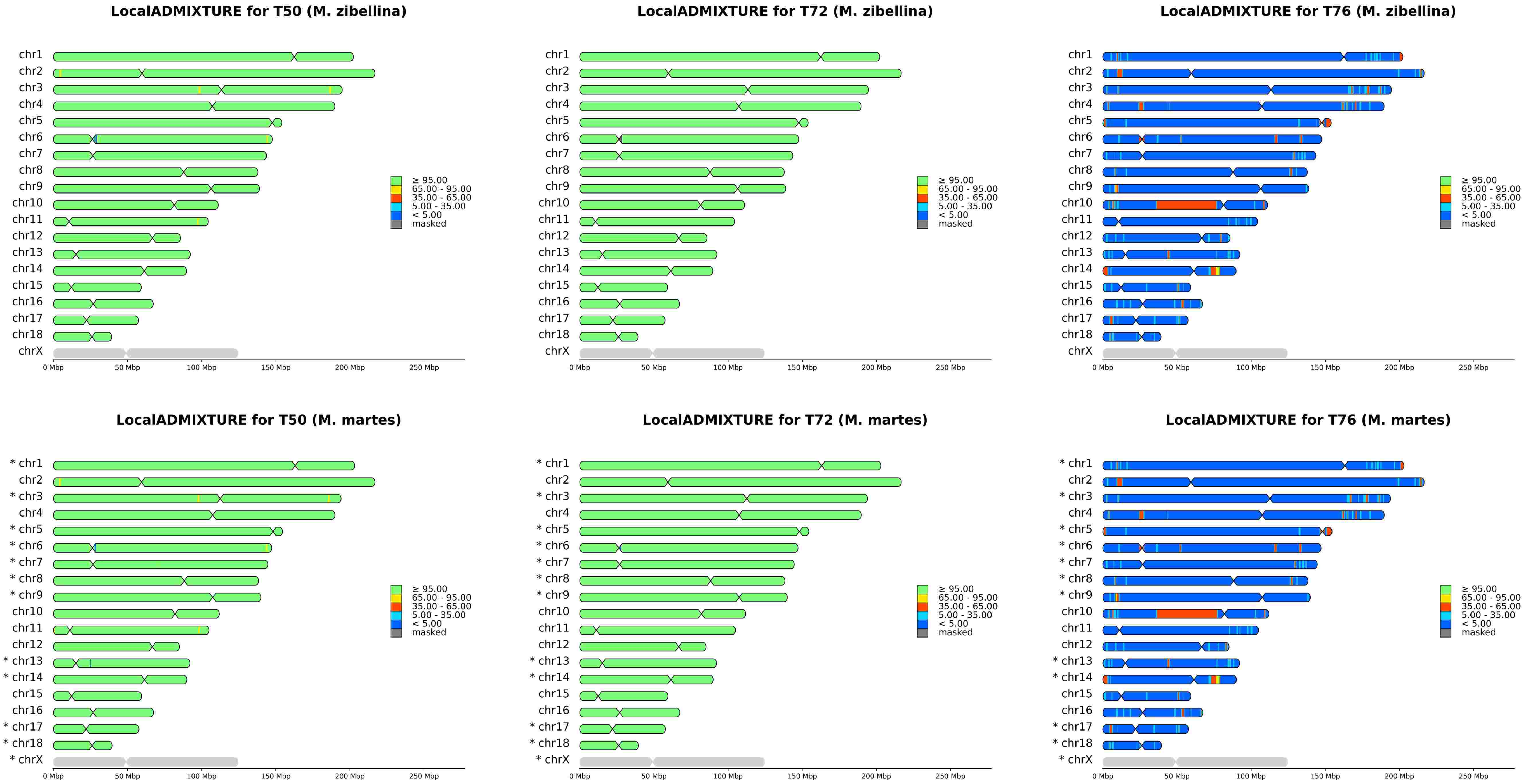
LocalADMIXTURE for T50 (M. zibellina)
chr1
chr2
chr3
chr4
chr5
chr6
chr7
chr8
chr9
chr10
chr11
chr12
chr13
chr14
chr15
chr16
chr17
chr18
chrX
≥ 95.00
65.00 - 95.00
35.00 - 65.00
5.00 - 35.00
< 5.00
masked
0 Mbp
50 Mbp
100 Mbp
150 Mbp
200 Mbp
250 Mbp
LocalADMIXTURE for T72 (M. zibellina)
chr1
chr2
chr3
chr4
chr5
chr6
chr7
chr8
chr9
chr10
chr11
chr12
chr13
chr14
chr15
chr16
chr17
chr18
chrX
≥ 95.00
65.00 - 95.00
35.00 - 65.00
5.00 - 35.00
< 5.00
masked
0 Mbp
50 Mbp
100 Mbp
150 Mbp
200 Mbp
250 Mbp
LocalADMIXTURE for T76 (M. zibellina)
chr1
chr2
chr3
chr4
chr5
chr6
chr7
chr8
chr9
chr10
chr11
chr12
chr13
chr14
chr15
chr16
chr17
chr18
chrX
≥ 95.00
65.00 - 95.00
35.00 - 65.00
5.00 - 35.00
< 5.00
masked
0 Mbp
50 Mbp
100 Mbp
150 Mbp
200 Mbp
250 Mbp
LocalADMIXTURE for T50 (M. martes)
* chr1
chr2
* chr3
chr4
* chr5
* chr6
* chr7
* chr8
* chr9
chr10
chr11
chr12
* chr13
* chr14
chr15
chr16
* chr17
* chr18
* chrX
≥ 95.00
65.00 - 95.00
35.00 - 65.00
5.00 - 35.00
< 5.00
masked
0 Mbp
50 Mbp
100 Mbp
150 Mbp
200 Mbp
250 Mbp
LocalADMIXTURE for T72 (M. martes)
* chr1
chr2
* chr3
chr4
* chr5
* chr6
* chr7
* chr8
* chr9
chr10
chr11
chr12
* chr13
* chr14
chr15
chr16
* chr17
* chr18
* chrX
≥ 95.00
65.00 - 95.00
35.00 - 65.00
5.00 - 35.00
< 5.00
masked
0 Mbp
50 Mbp
100 Mbp
150 Mbp
200 Mbp
250 Mbp
LocalADMIXTURE for T76 (M. martes)
* chr1
chr2
* chr3
chr4
* chr5
* chr6
* chr7
* chr8
* chr9
chr10
chr11
chr12
* chr13
* chr14
chr15
chr16
* chr17
* chr18
* chrX
≥ 95.00
65.00 - 95.00
35.00 - 65.00
5.00 - 35.00
< 5.00
masked
0 Mbp
50 Mbp
100 Mbp
150 Mbp
200 Mbp
250 Mbp

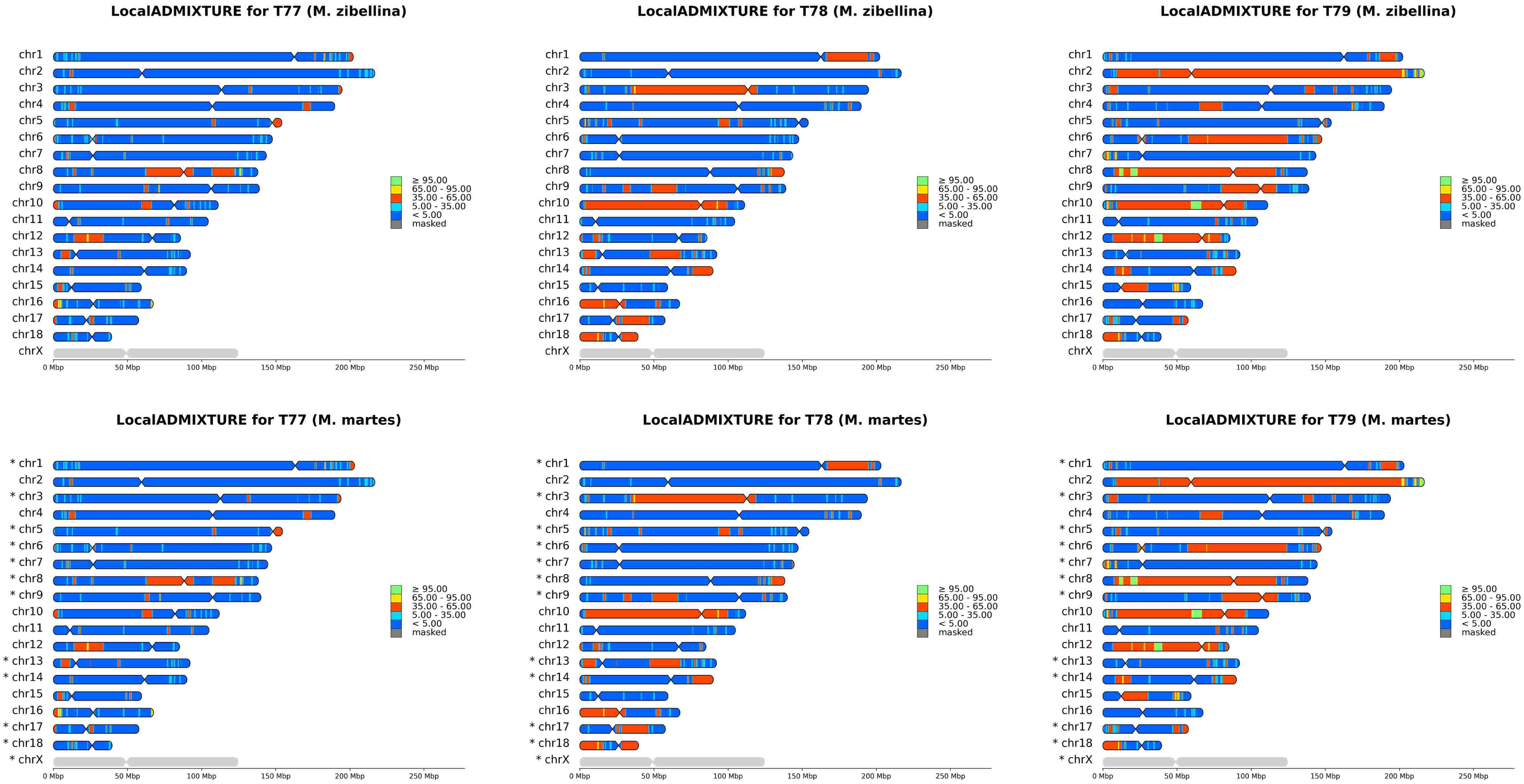
LocalADMIXTURE for T77 (M. zibellina)
LocalADMIXTURE for T78 (M. zibellina)
LocalADMIXTURE for T79 (M. zibellina)
chr1
chr2
chr3
chr4
chr5
chr6
chr7
chr8
chr9
chr10
chr11
chr12
chr13
chr14
chr15
chr16
chr17
chr18
chrX
≥ 95.00
65.00 - 95.00
35.00 - 65.00
5.00 - 35.00
< 5.00
masked
0 Mbp
50 Mbp
100 Mbp
150 Mbp
200 Mbp
250 Mbp
LocalADMIXTURE for T77 (M. martes)
LocalADMIXTURE for T78 (M. martes)
LocalADMIXTURE for T79 (M. martes)
* chr1
chr2
* chr3
chr4
* chr5
* chr6
* chr7
* chr8
* chr9
chr10
chr11
chr12
* chr13
* chr14
chr15
chr16
* chr17
* chr18
* chrX

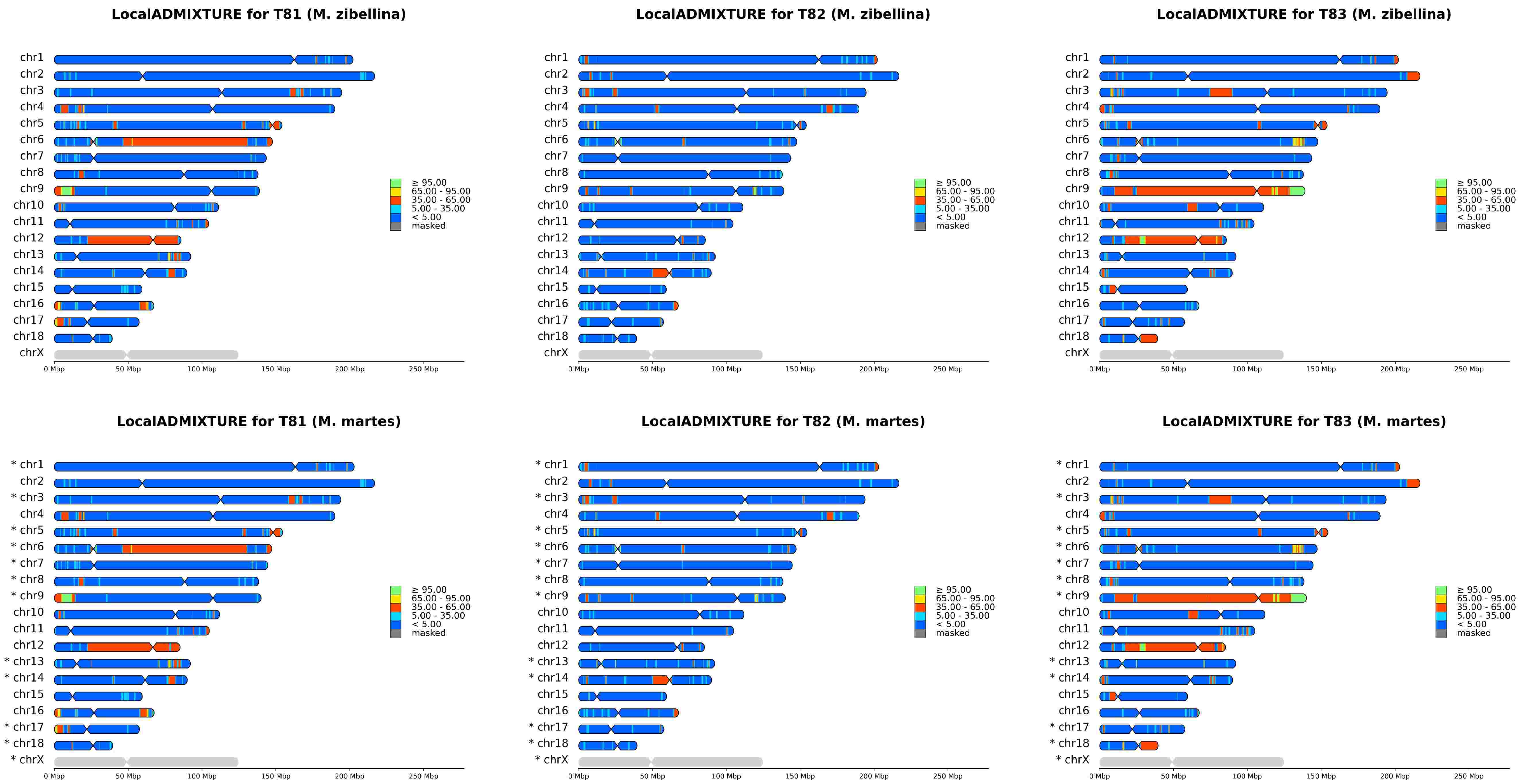
LocalADMIXTURE for T81 (M. zibellina)
LocalADMIXTURE for T82 (M. zibellina)
LocalADMIXTURE for T83 (M. zibellina)
chr1
chr2
chr3
chr4
chr5
chr6
chr7
chr8
chr9
chr10
chr11
chr12
chr13
chr14
chr15
chr16
chr17
chr18
chrX
≥ 95.00
65.00 - 95.00
35.00 - 65.00
5.00 - 35.00
< 5.00
masked
0 Mbp
50 Mbp
100 Mbp
150 Mbp
200 Mbp
250 Mbp
LocalADMIXTURE for T81 (M. martes)
LocalADMIXTURE for T82 (M. martes)
LocalADMIXTURE for T83 (M. martes)
* chr1
chr2
* chr3
chr4
* chr5
* chr6
* chr7
* chr8
* chr9
chr10
chr11
chr12
* chr13
* chr14
chr15
chr16
* chr17
* chr18
* chrX

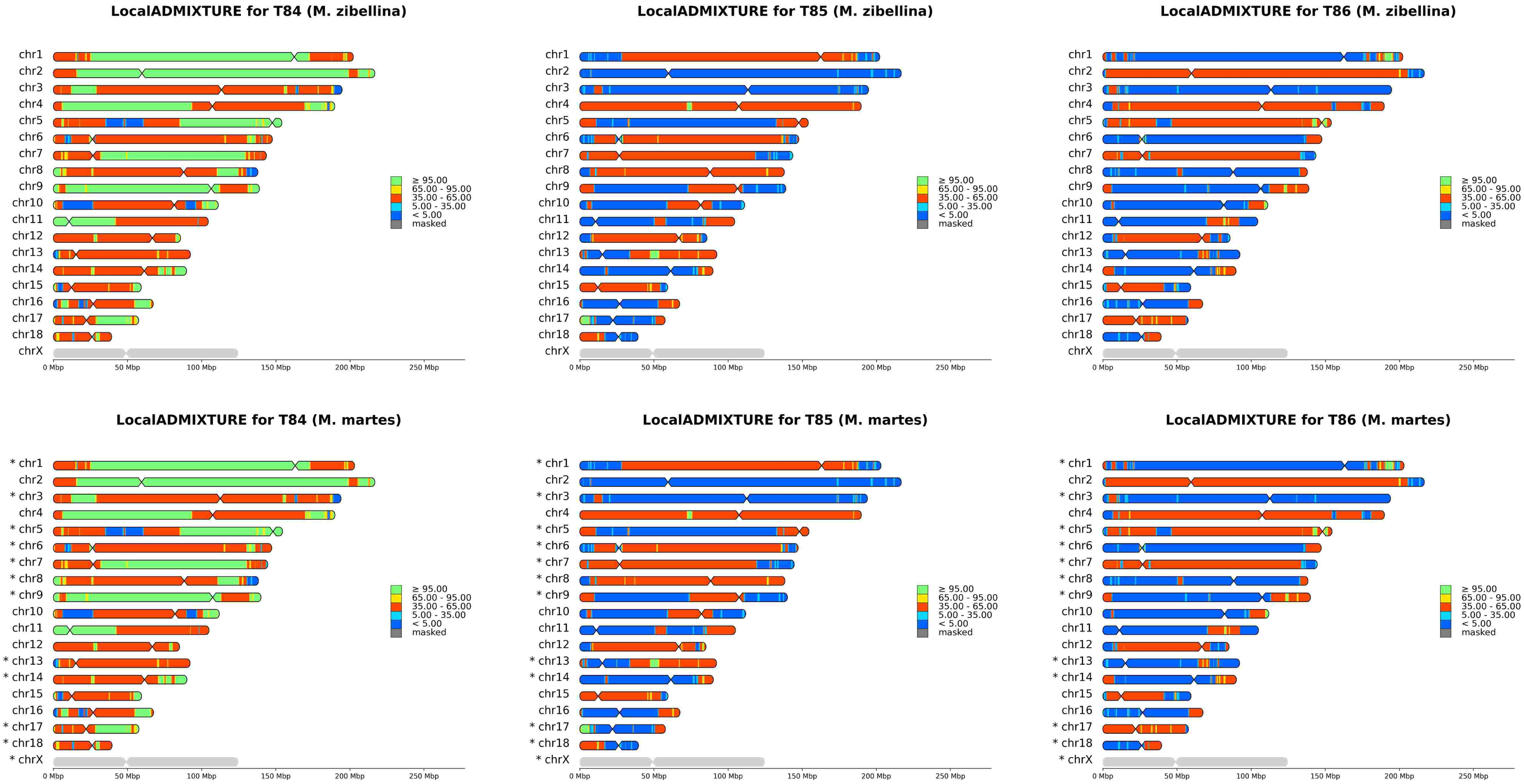
LocalADMIXTURE for T84 (M. zibellina)
LocalADMIXTURE for T85 (M. zibellina)
LocalADMIXTURE for T86 (M. zibellina)
chr1
chr2
chr3
chr4
chr5
chr6
chr7
chr8
chr9
chr10
chr11
chr12
chr13
chr14
chr15
chr16
chr17
chr18
chrX
≥ 95.00
65.00 - 95.00
35.00 - 65.00
5.00 - 35.00
< 5.00
masked
0 Mbp
50 Mbp
100 Mbp
150 Mbp
200 Mbp
250 Mbp
LocalADMIXTURE for T84 (M. martes)
LocalADMIXTURE for T85 (M. martes)
LocalADMIXTURE for T86 (M. martes)
* chr1
chr2
* chr3
chr4
* chr5
* chr6
* chr7
* chr8
* chr9
chr10
chr11
chr12
* chr13
* chr14
chr15
chr16
* chr17
* chr18
* chrX

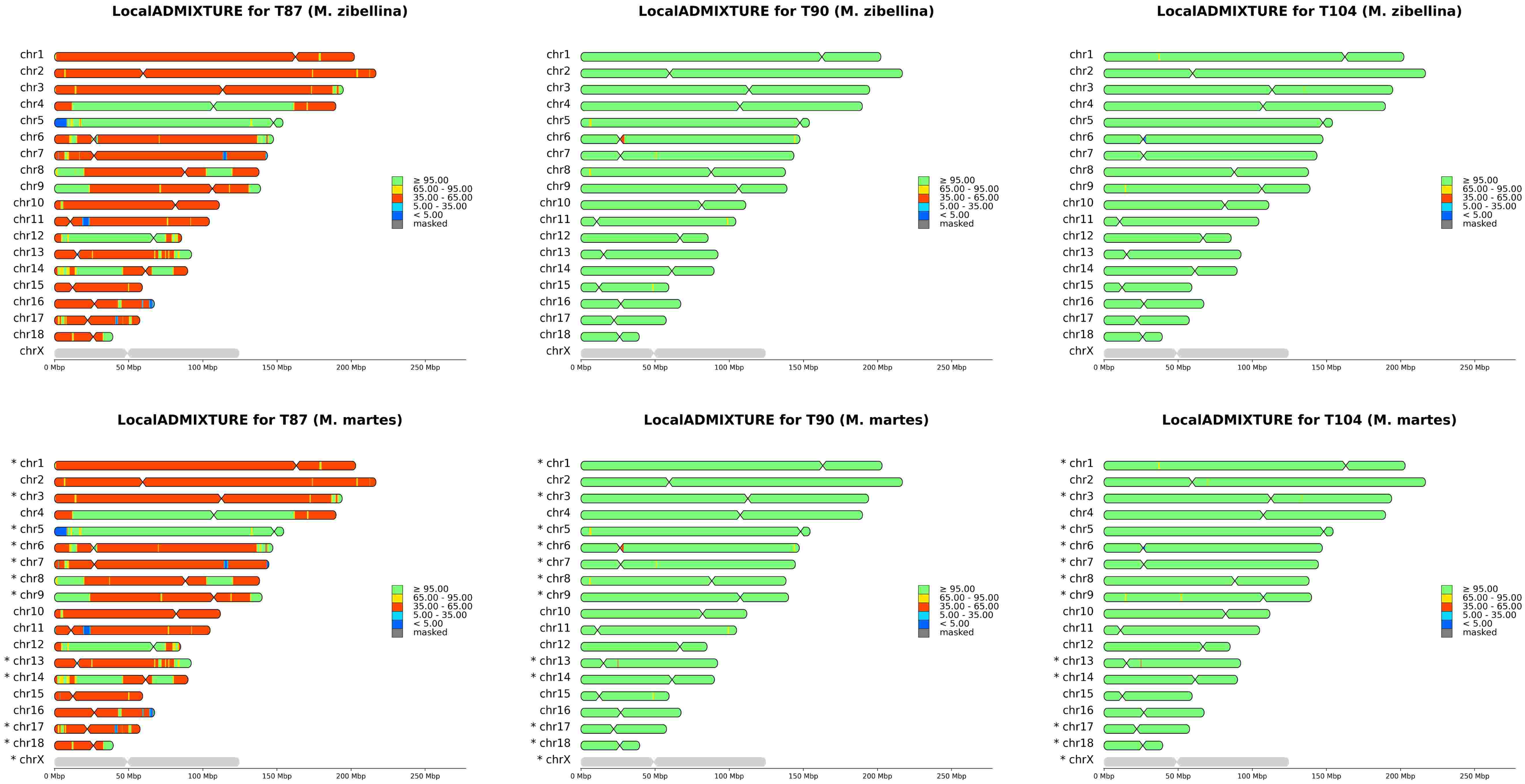
LocalADMIXTURE for T87 (M. zibellina)
LocalADMIXTURE for T90 (M. zibellina)
LocalADMIXTURE for T104 (M. zibellina)
chr1
chr2
chr3
chr4
chr5
chr6
chr7
chr8
chr9
chr10
chr11
chr12
chr13
chr14
chr15
chr16
chr17
chr18
chrX
≥ 95.00
65.00 - 95.00
35.00 - 65.00
5.00 - 35.00
< 5.00
masked
0 Mbp
50 Mbp
100 Mbp
150 Mbp
200 Mbp
250 Mbp
LocalADMIXTURE for T87 (M. martes)
LocalADMIXTURE for T90 (M. martes)
LocalADMIXTURE for T104 (M. martes)
* chr1
chr2
* chr3
chr4
* chr5
* chr6
* chr7
* chr8
* chr9
chr10
chr11
chr12
* chr13
* chr14
chr15
chr16
* chr17
* chr18
* chrX

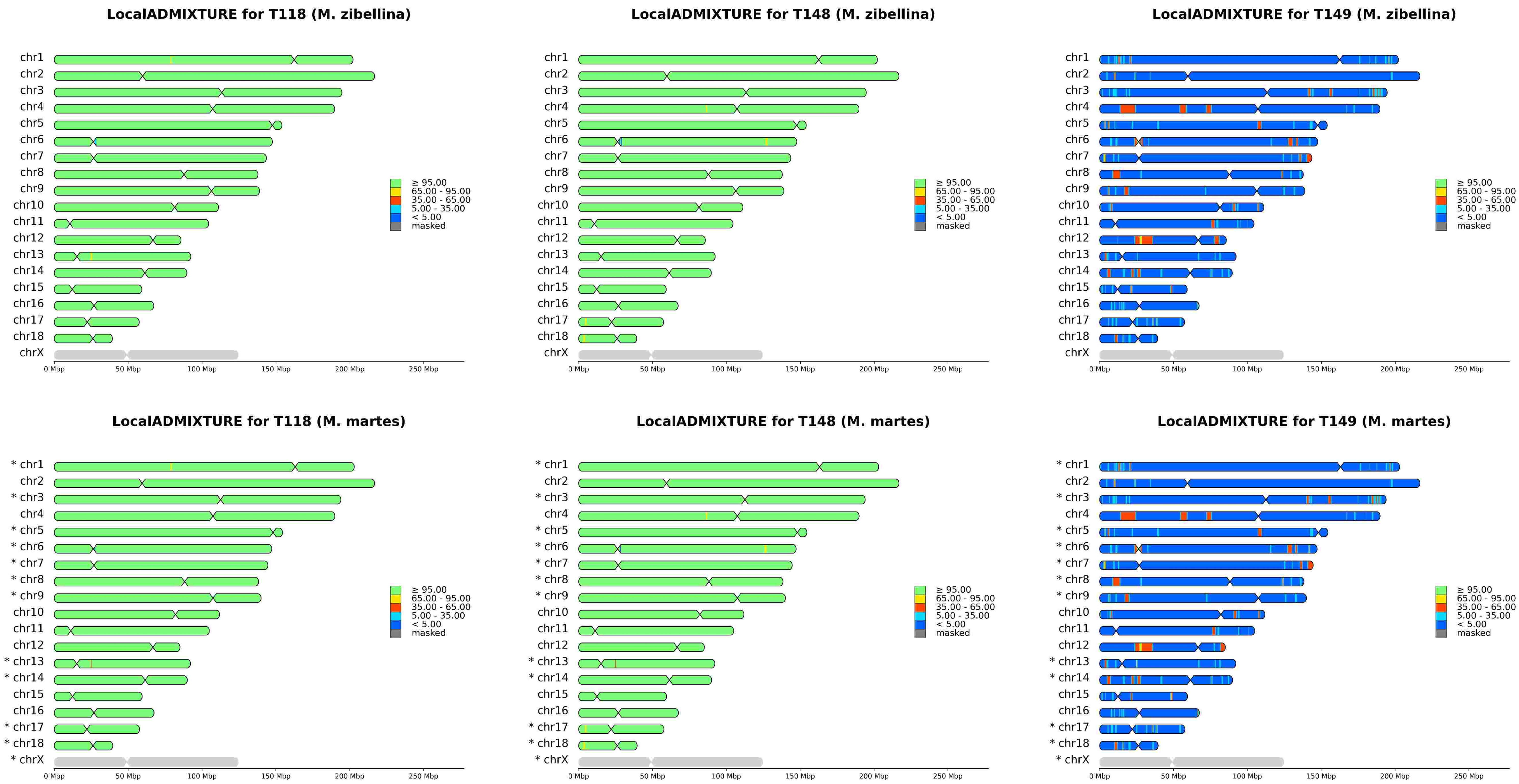
LocalADMIXTURE for T118 (M. zibellina)
chr1
chr2
chr3
chr4
chr5
chr6
chr7
chr8
chr9
chr10
chr11
chr12
chr13
chr14
chr15
chr16
chr17
chr18
chrX
≥ 95.00
65.00 - 95.00
35.00 - 65.00
5.00 - 35.00
< 5.00
masked
0 Mbp
50 Mbp
100 Mbp
150 Mbp
200 Mbp
250 Mbp
LocalADMIXTURE for T148 (M. zibellina)
chr1
chr2
chr3
chr4
chr5
chr6
chr7
chr8
chr9
chr10
chr11
chr12
chr13
chr14
chr15
chr16
chr17
chr18
chrX
≥ 95.00
65.00 - 95.00
35.00 - 65.00
5.00 - 35.00
< 5.00
masked
0 Mbp
50 Mbp
100 Mbp
150 Mbp
200 Mbp
250 Mbp
LocalADMIXTURE for T149 (M. zibellina)
chr1
chr2
chr3
chr4
chr5
chr6
chr7
chr8
chr9
chr10
chr11
chr12
chr13
chr14
chr15
chr16
chr17
chr18
chrX
≥ 95.00
65.00 - 95.00
35.00 - 65.00
5.00 - 35.00
< 5.00
masked
0 Mbp
50 Mbp
100 Mbp
150 Mbp
200 Mbp
250 Mbp
LocalADMIXTURE for T118 (M. martes)
* chr1
chr2
* chr3
chr4
* chr5
* chr6
* chr7
* chr8
* chr9
chr10
chr11
chr12
* chr13
* chr14
chr15
chr16
* chr17
* chr18
* chrX
≥ 95.00
65.00 - 95.00
35.00 - 65.00
5.00 - 35.00
< 5.00
masked
0 Mbp
50 Mbp
100 Mbp
150 Mbp
200 Mbp
250 Mbp
LocalADMIXTURE for T148 (M. martes)
* chr1
chr2
* chr3
chr4
* chr5
* chr6
* chr7
* chr8
* chr9
chr10
chr11
chr12
* chr13
* chr14
chr15
chr16
* chr17
* chr18
* chrX
≥ 95.00
65.00 - 95.00
35.00 - 65.00
5.00 - 35.00
< 5.00
masked
0 Mbp
50 Mbp
100 Mbp
150 Mbp
200 Mbp
250 Mbp
LocalADMIXTURE for T149 (M. martes)
* chr1
chr2
* chr3
chr4
* chr5
* chr6
* chr7
* chr8
* chr9
chr10
chr11
chr12
* chr13
* chr14
chr15
chr16
* chr17
* chr18
* chrX
≥ 95.00
65.00 - 95.00
35.00 - 65.00
5.00 - 35.00
< 5.00
masked
0 Mbp
50 Mbp
100 Mbp
150 Mbp
200 Mbp
250 Mbp

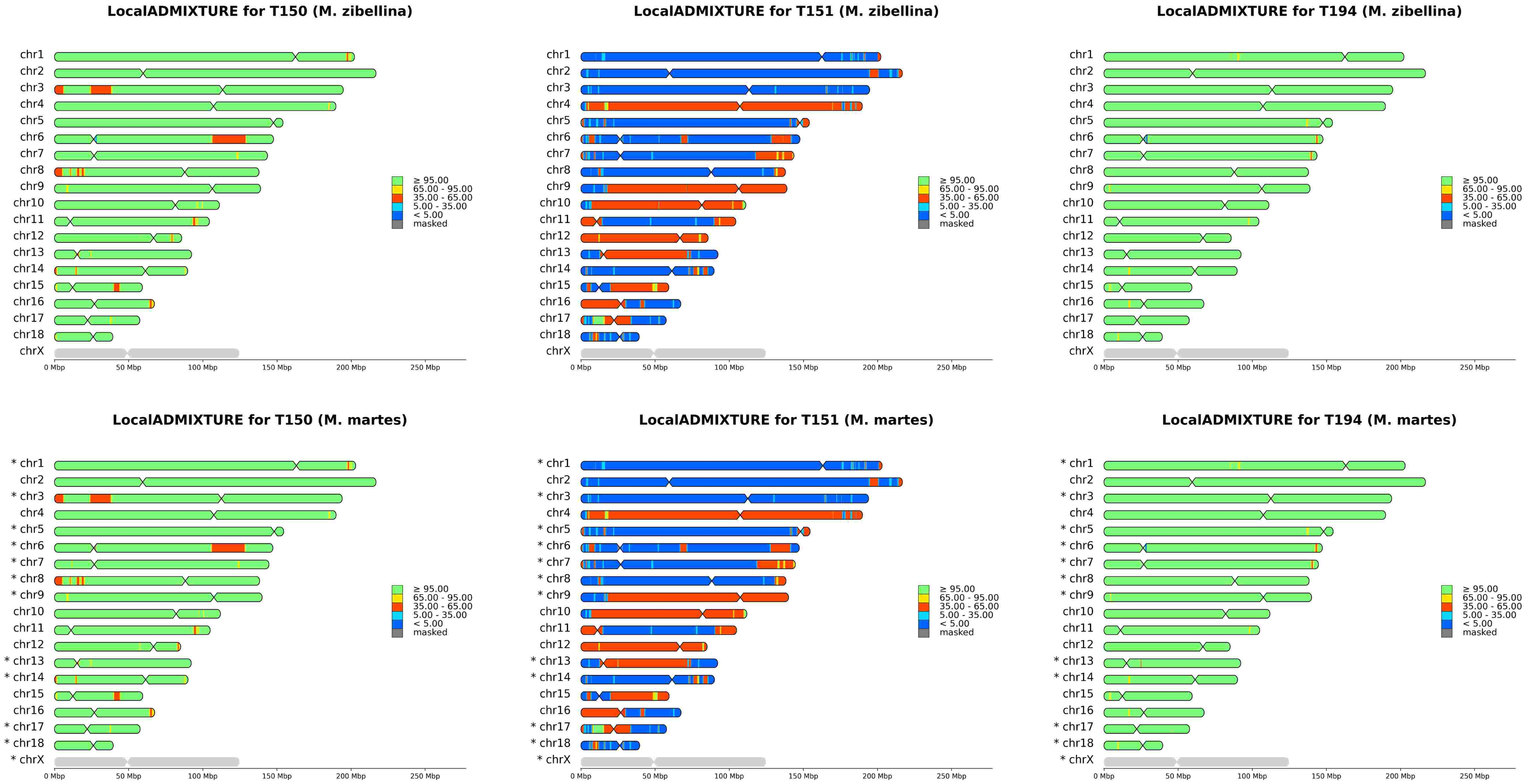
LocalADMIXTURE for T150 (M. zibellina)
LocalADMIXTURE for T151 (M. zibellina)
LocalADMIXTURE for T194 (M. zibellina)
LocalADMIXTURE for T150 (M. martes)
LocalADMIXTURE for T151 (M. martes)
LocalADMIXTURE for T194 (M. martes)
chr1
chr2
chr3
chr4
chr5
chr6
chr7
chr8
chr9
chr10
chr11
chr12
chr13
chr14
chr15
chr16
chr17
chr18
chrX
* chr1
* chr3
* chr5
* chr6
* chr7
* chr8
* chr9
* chr13
* chr14
* chr17
* chr18
* chrX
≥ 95.00
65.00 - 95.00
35.00 - 65.00
5.00 - 35.00
< 5.00
masked
0 Mbp
50 Mbp
100 Mbp
150 Mbp
200 Mbp
250 Mbp
