## Supplementary File SF11 for "Genomics of sable (*Martes zibellina)* × pine marten (*Martes martes*) hybridization"

### 10xmmar

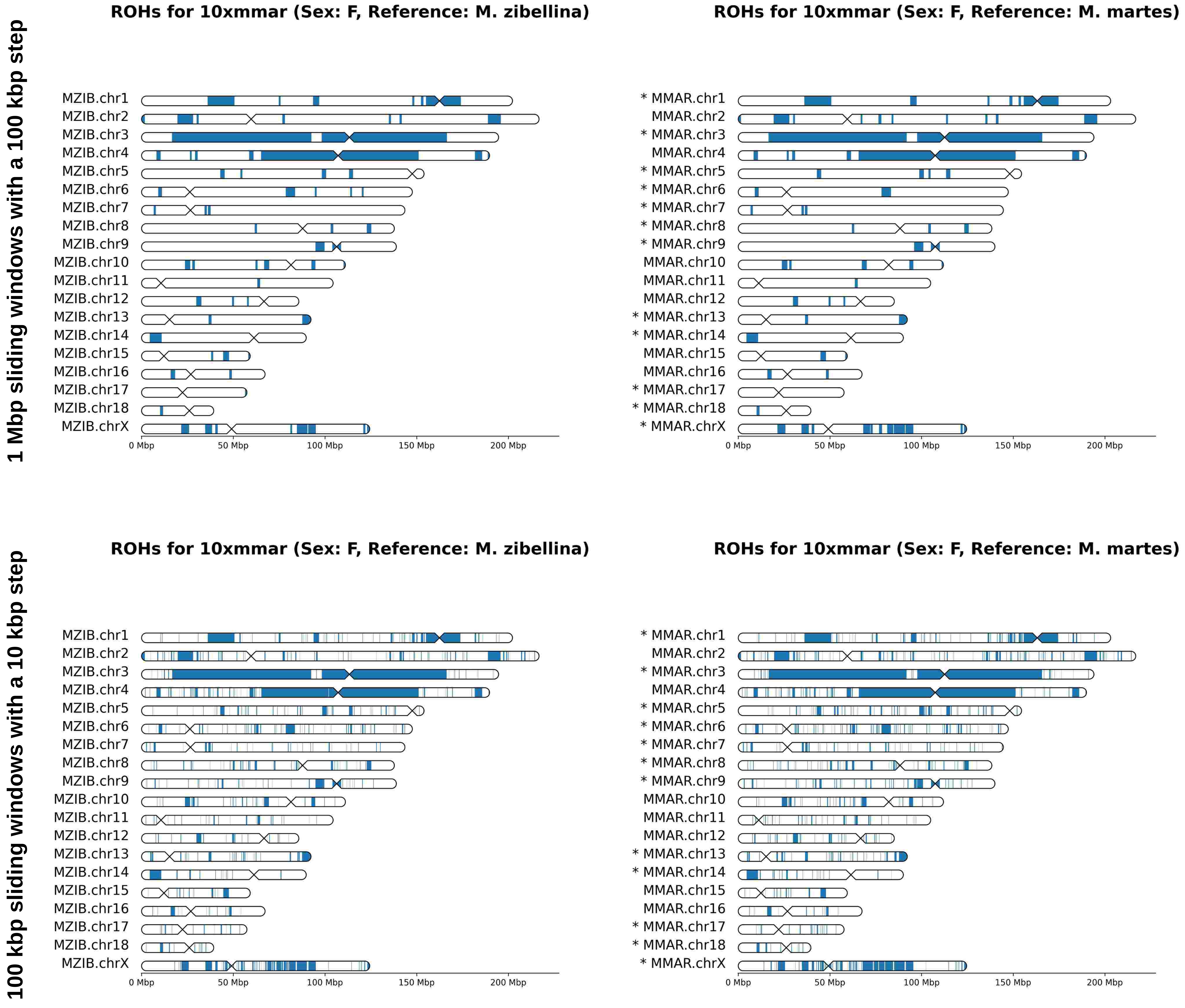

1 Mbp sliding windows with a 100 kbp step
ROHs for 10xmmar (Sex: F, Reference: M. zibellina)
MZIB.chr1
MZIB.chr2
MZIB.chr3
MZIB.chr4
MZIB.chr5
MZIB.chr6
MZIB.chr7
MZIB.chr8
MZIB.chr9
MZIB.chr10
MZIB.chr11
MZIB.chr12
MZIB.chr13
MZIB.chr14
MZIB.chr15
MZIB.chr16
MZIB.chr17
MZIB.chr18
MZIB.chrX
0 Mbp
50 Mbp
100 Mbp
150 Mbp
200 Mbp
ROHs for 10xmmar (Sex: F, Reference: M. martes)
* MMAR.chr1
MMAR.chr2
* MMAR.chr3
MMAR.chr4
* MMAR.chr5
* MMAR.chr6
* MMAR.chr7
* MMAR.chr8
* MMAR.chr9
MMAR.chr10
MMAR.chr11
MMAR.chr12
* MMAR.chr13
* MMAR.chr14
MMAR.chr15
MMAR.chr16
* MMAR.chr17
* MMAR.chr18
* MMAR.chrX
0 Mbp
50 Mbp
100 Mbp
150 Mbp
200 Mbp
100 kbp sliding windows with a 10 kbp step
ROHs for 10xmmar (Sex: F, Reference: M. zibellina)
MZIB.chr1
MZIB.chr2
MZIB.chr3
MZIB.chr4
MZIB.chr5
MZIB.chr6
MZIB.chr7
MZIB.chr8
MZIB.chr9
MZIB.chr10
MZIB.chr11
MZIB.chr12
MZIB.chr13
MZIB.chr14
MZIB.chr15
MZIB.chr16
MZIB.chr17
MZIB.chr18
MZIB.chrX
0 Mbp
50 Mbp
100 Mbp
150 Mbp
200 Mbp
ROHs for 10xmmar (Sex: F, Reference: M. martes)
* MMAR.chr1
MMAR.chr2
* MMAR.chr3
MMAR.chr4
* MMAR.chr5
* MMAR.chr6
* MMAR.chr7
* MMAR.chr8
* MMAR.chr9
MMAR.chr10
MMAR.chr11
MMAR.chr12
* MMAR.chr13
* MMAR.chr14
MMAR.chr15
MMAR.chr16
* MMAR.chr17
* MMAR.chr18
* MMAR.chrX
0 Mbp
50 Mbp
100 Mbp
150 Mbp
200 Mbp

### 10xmzib

1 Mbp sliding windows with a 100 kbp step

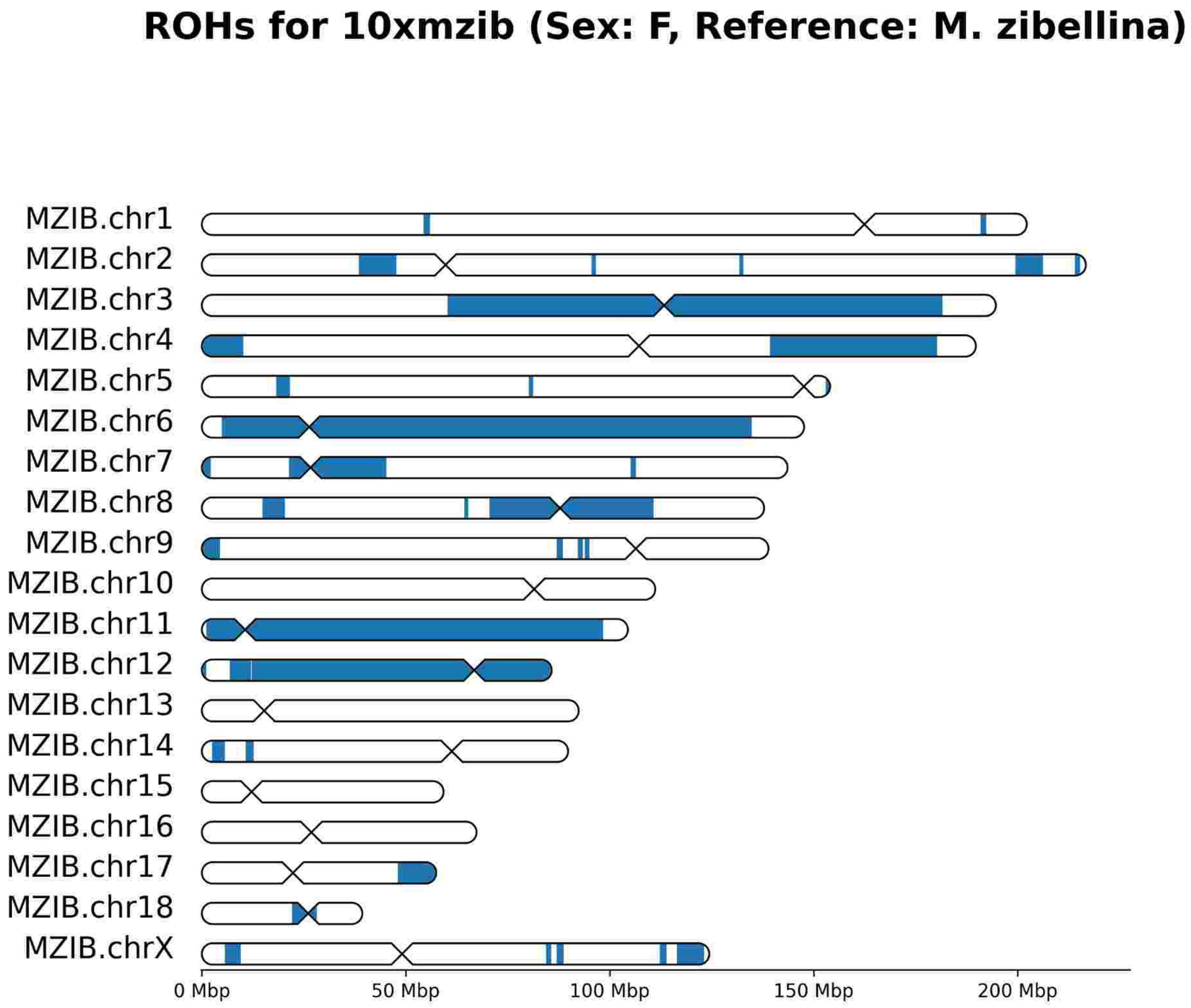


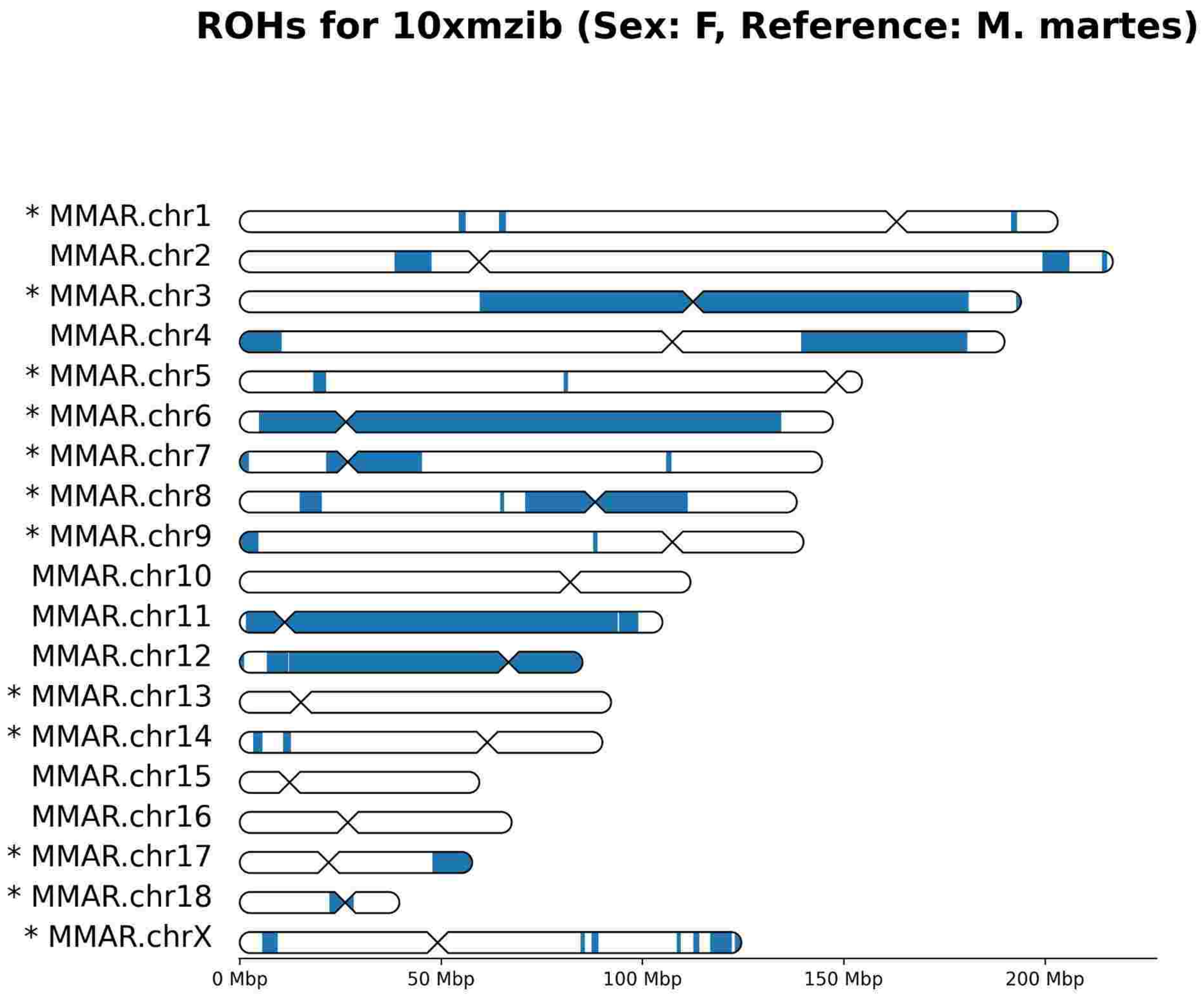


100 kbp sliding windows with a 10 kbp step

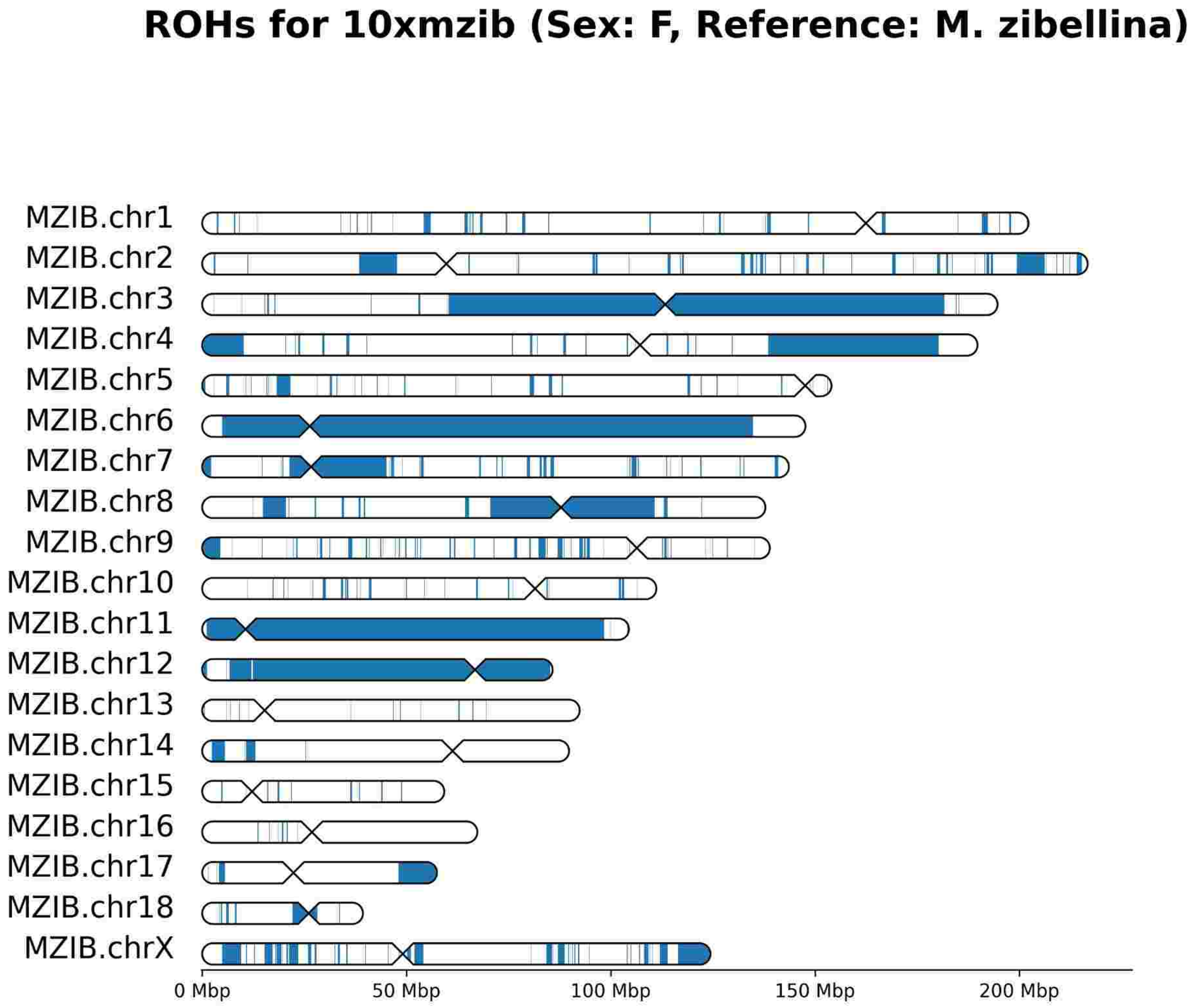


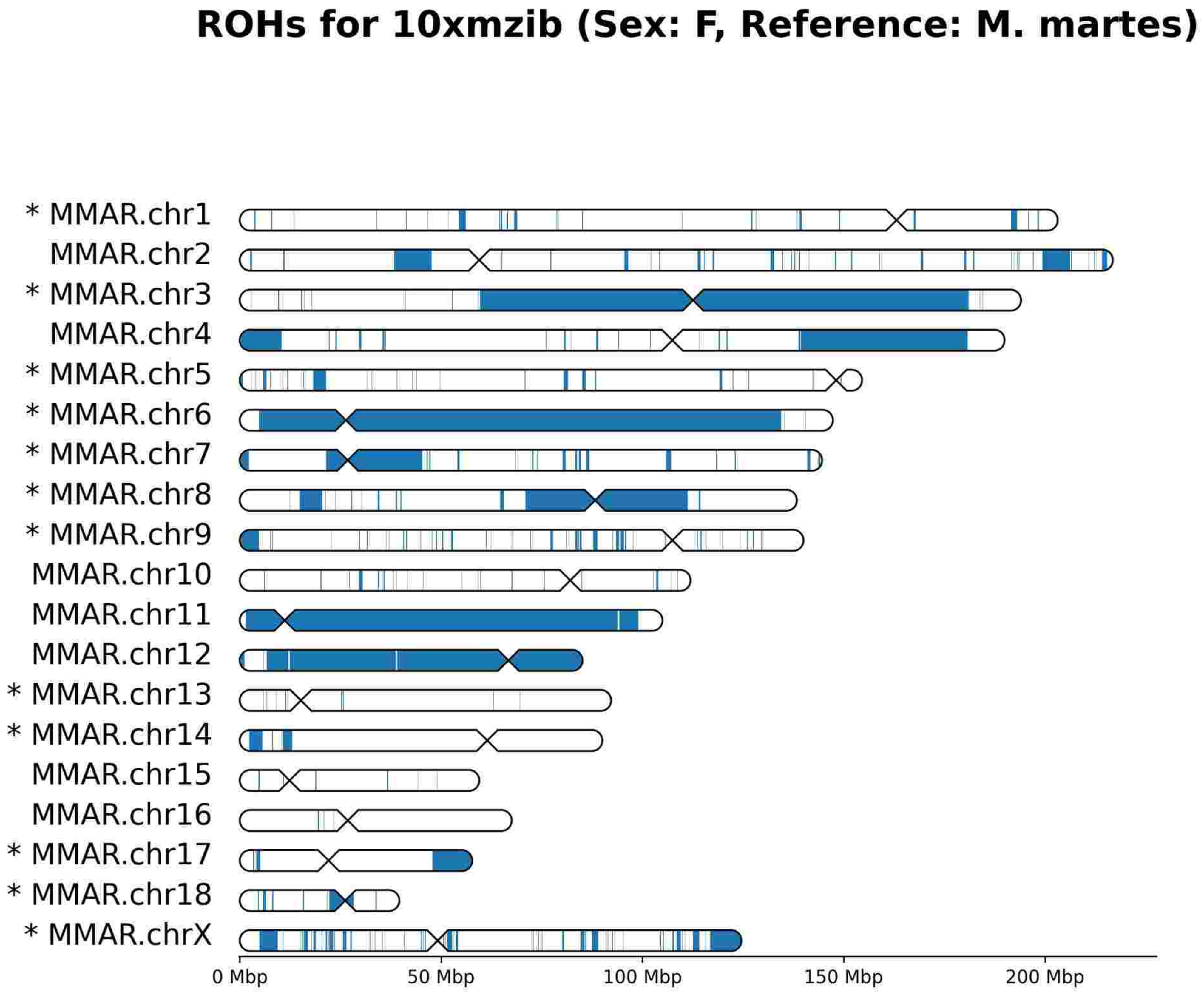

### CHN

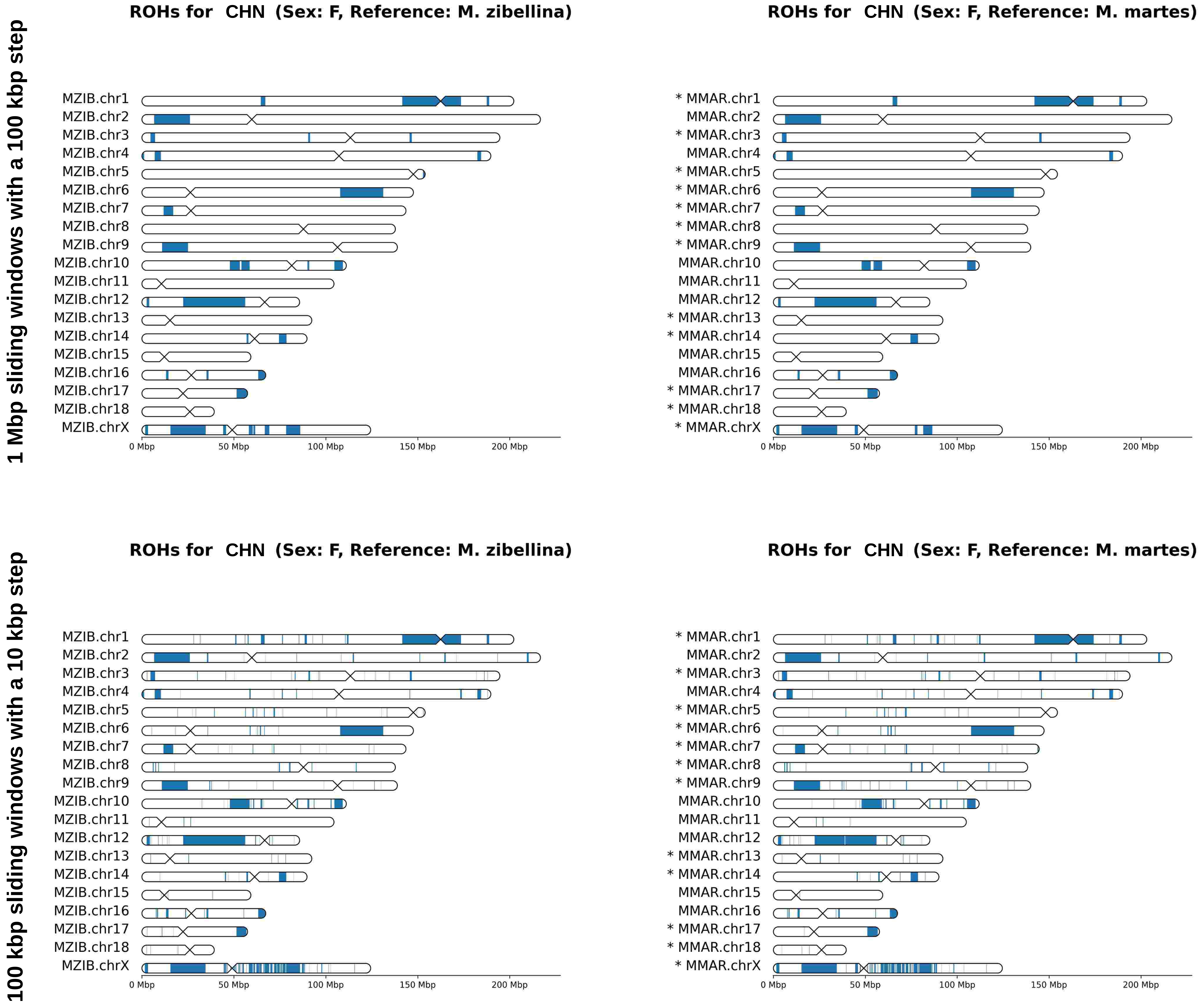
1 Mbp sliding windows with a 100 kbp step
100 kbp sliding windows with a 10 kbp step
ROHs for CHN (Sex: F, Reference: M. zibellina)
ROHs for CHN (Sex: F, Reference: M. martes)
MZIB.chr1
MZIB.chr2
MZIB.chr3
MZIB.chr4
MZIB.chr5
MZIB.chr6
MZIB.chr7
MZIB.chr8
MZIB.chr9
MZIB.chr10
MZIB.chr11
MZIB.chr12
MZIB.chr13
MZIB.chr14
MZIB.chr15
MZIB.chr16
MZIB.chr17
MZIB.chr18
MZIB.chrX
* MMAR.chr1
MMAR.chr2
* MMAR.chr3
MMAR.chr4
* MMAR.chr5
* MMAR.chr6
* MMAR.chr7
* MMAR.chr8
* MMAR.chr9
MMAR.chr10
MMAR.chr11
MMAR.chr12
* MMAR.chr13
* MMAR.chr14
MMAR.chr15
MMAR.chr16
* MMAR.chr17
* MMAR.chr18
* MMAR.chrX
0 Mbp
50 Mbp
100 Mbp
150 Mbp
200 Mbp

# S26

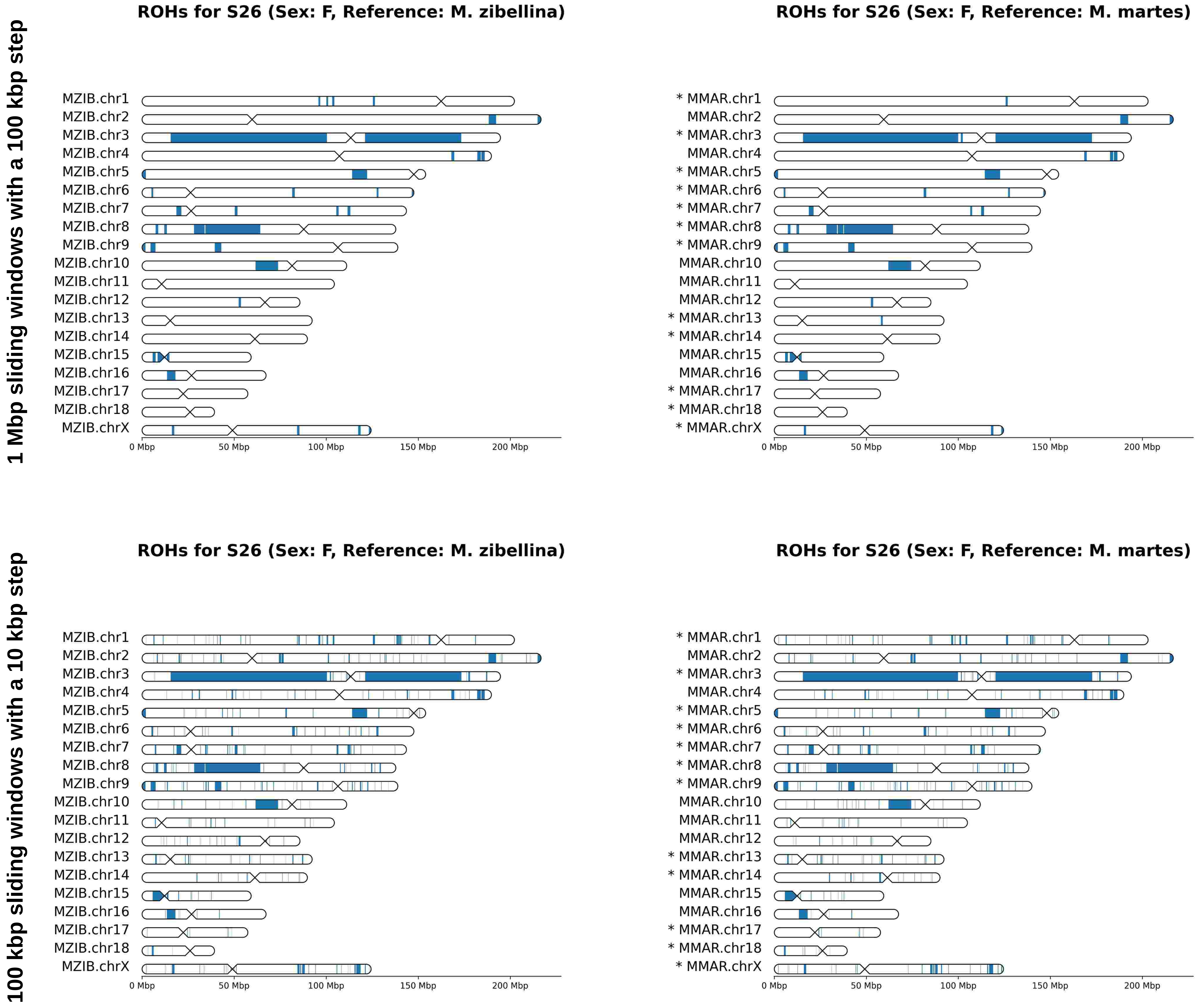
1 Mbp sliding windows with a 100 kbp step
100 kbp sliding windows with a 10 kbp step
ROHs for S26 (Sex: F, Reference: M. zibellina)
ROHs for S26 (Sex: F, Reference: M. martes)
MZIB.chr1
MZIB.chr2
MZIB.chr3
MZIB.chr4
MZIB.chr5
MZIB.chr6
MZIB.chr7
MZIB.chr8
MZIB.chr9
MZIB.chr10
MZIB.chr11
MZIB.chr12
MZIB.chr13
MZIB.chr14
MZIB.chr15
MZIB.chr16
MZIB.chr17
MZIB.chr18
MZIB.chrX
* MMAR.chr1
MMAR.chr2
* MMAR.chr3
MMAR.chr4
* MMAR.chr5
* MMAR.chr6
* MMAR.chr7
* MMAR.chr8
* MMAR.chr9
MMAR.chr10
MMAR.chr11
MMAR.chr12
* MMAR.chr13
* MMAR.chr14
MMAR.chr15
MMAR.chr16
* MMAR.chr17
* MMAR.chr18
* MMAR.chrX
0 Mbp
50 Mbp
100 Mbp
150 Mbp
200 Mbp

# S44

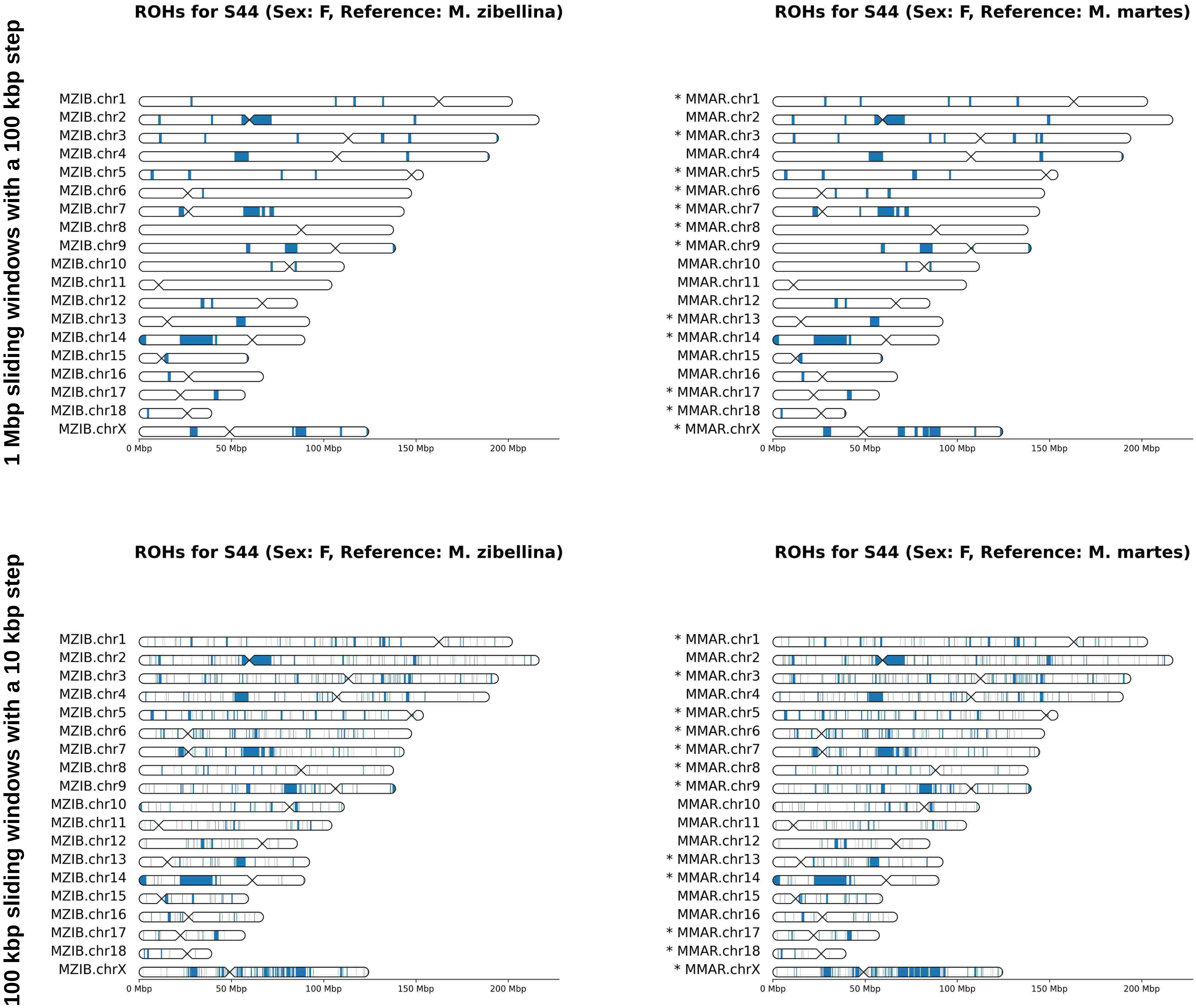
1 Mbp sliding windows with a 100 kbp step
100 kbp sliding windows with a 10 kbp step
ROHs for S44 (Sex: F, Reference: M. zibellina)
ROHs for S44 (Sex: F, Reference: M. martes)
MZIB.chr1
MZIB.chr2
MZIB.chr3
MZIB.chr4
MZIB.chr5
MZIB.chr6
MZIB.chr7
MZIB.chr8
MZIB.chr9
MZIB.chr10
MZIB.chr11
MZIB.chr12
MZIB.chr13
MZIB.chr14
MZIB.chr15
MZIB.chr16
MZIB.chr17
MZIB.chr18
MZIB.chrX
* MMAR.chr1
MMAR.chr2
* MMAR.chr3
MMAR.chr4
* MMAR.chr5
* MMAR.chr6
* MMAR.chr7
* MMAR.chr8
* MMAR.chr9
MMAR.chr10
MMAR.chr11
MMAR.chr12
* MMAR.chr13
* MMAR.chr14
MMAR.chr15
MMAR.chr16
* MMAR.chr17
* MMAR.chr18
* MMAR.chrX
0 Mbp
50 Mbp
100 Mbp
150 Mbp
200 Mbp

# S46

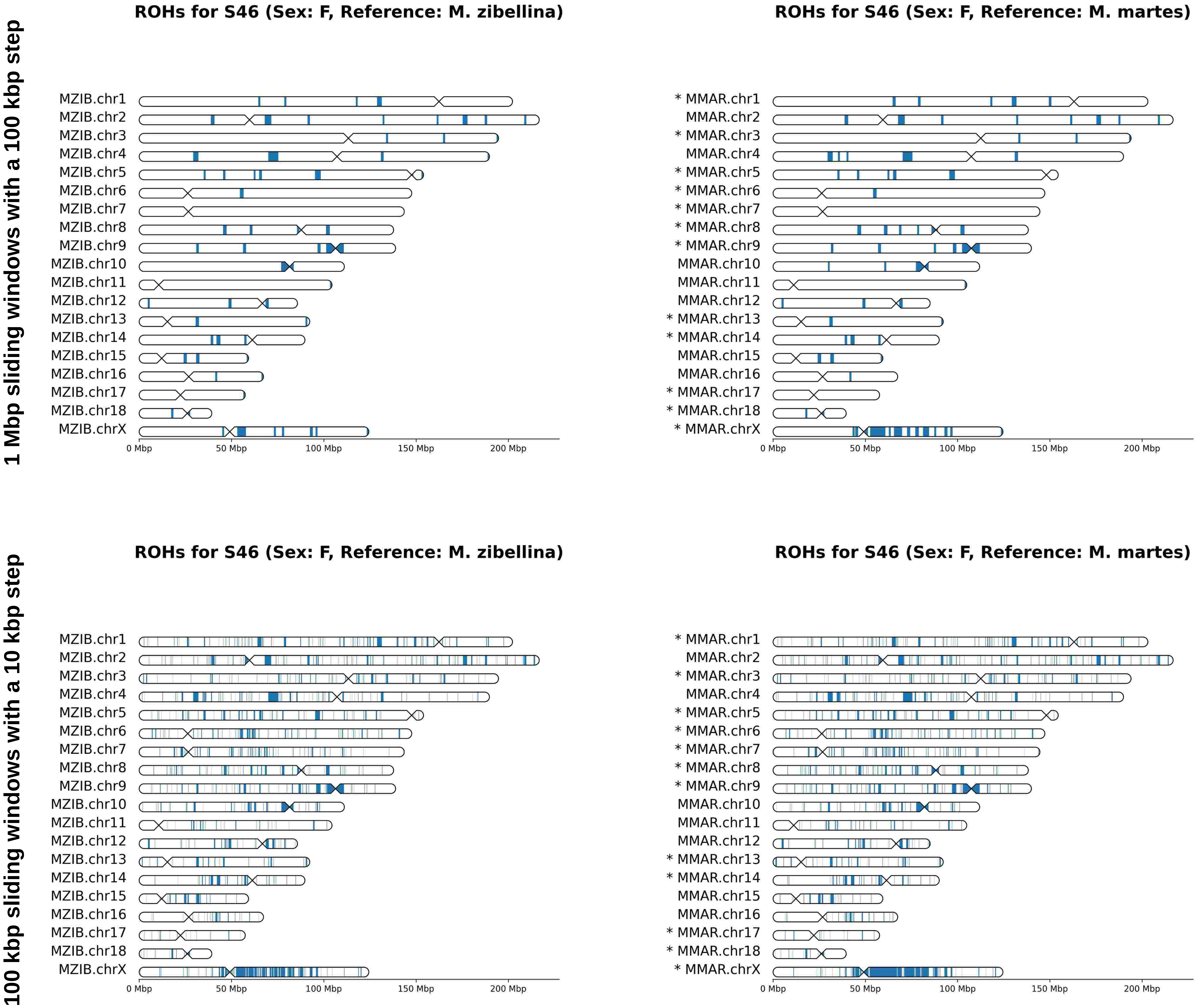
1 Mbp sliding windows with a 100 kbp step
100 kbp sliding windows with a 10 kbp step
ROHs for S46 (Sex: F, Reference: M. zibellina)
MZIB.chr1
MZIB.chr2
MZIB.chr3
MZIB.chr4
MZIB.chr5
MZIB.chr6
MZIB.chr7
MZIB.chr8
MZIB.chr9
MZIB.chr10
MZIB.chr11
MZIB.chr12
MZIB.chr13
MZIB.chr14
MZIB.chr15
MZIB.chr16
MZIB.chr17
MZIB.chr18
MZIB.chrX
0 Mbp
50 Mbp
100 Mbp
150 Mbp
200 Mbp
ROHs for S46 (Sex: F, Reference: M. martes)
* MMAR.chr1
MMAR.chr2
* MMAR.chr3
MMAR.chr4
* MMAR.chr5
* MMAR.chr6
* MMAR.chr7
* MMAR.chr8
* MMAR.chr9
MMAR.chr10
MMAR.chr11
MMAR.chr12
* MMAR.chr13
* MMAR.chr14
MMAR.chr15
MMAR.chr16
* MMAR.chr17
* MMAR.chr18
* MMAR.chrX
0 Mbp
50 Mbp
100 Mbp
150 Mbp
200 Mbp

# S49

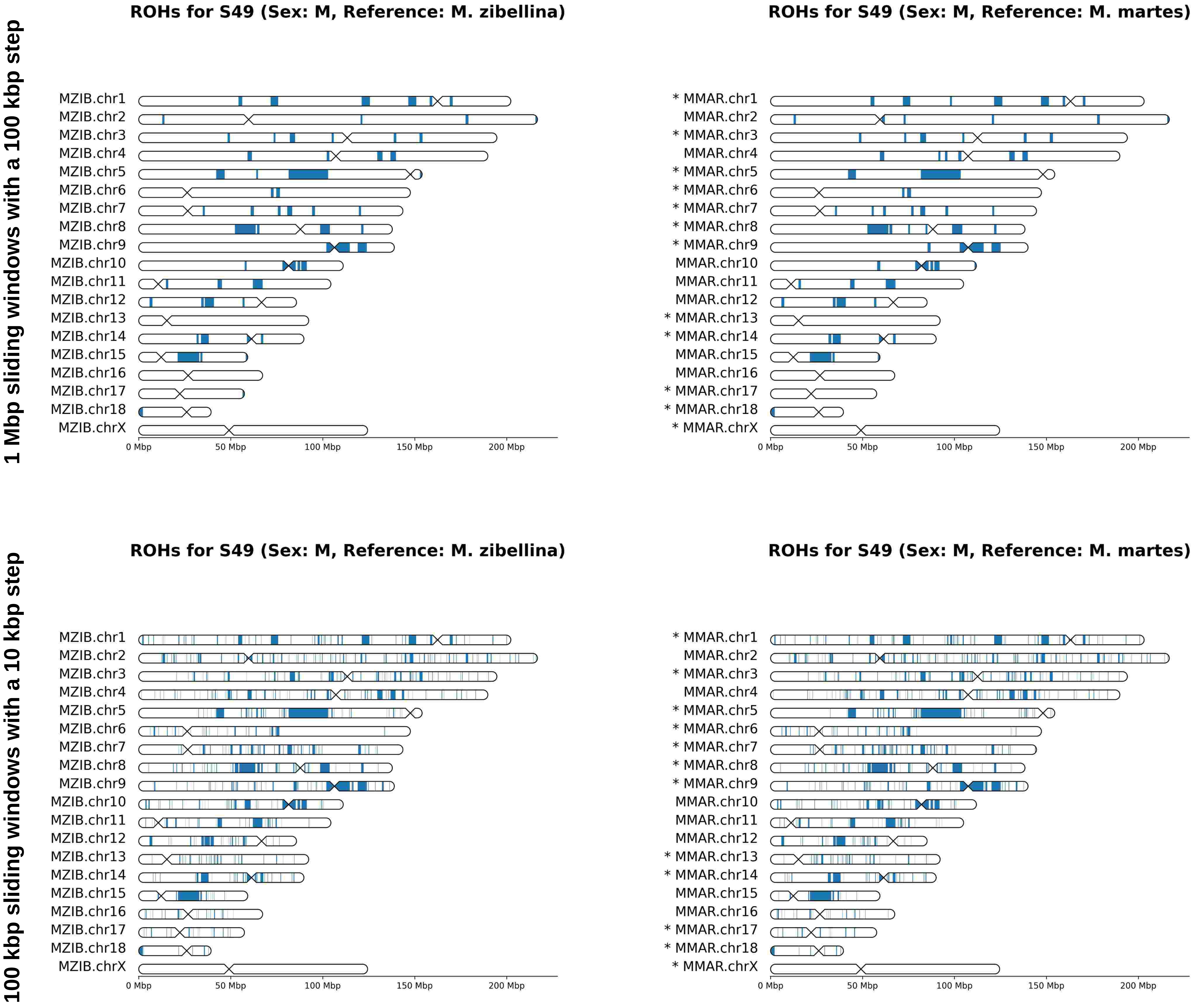
1 Mbp sliding windows with a 100 kbp step
100 kbp sliding windows with a 10 kbp step
ROHs for S49 (Sex: M, Reference: M. zibellina)
MZIB.chr1
MZIB.chr2
MZIB.chr3
MZIB.chr4
MZIB.chr5
MZIB.chr6
MZIB.chr7
MZIB.chr8
MZIB.chr9
MZIB.chr10
MZIB.chr11
MZIB.chr12
MZIB.chr13
MZIB.chr14
MZIB.chr15
MZIB.chr16
MZIB.chr17
MZIB.chr18
MZIB.chrX
0 Mbp
50 Mbp
100 Mbp
150 Mbp
200 Mbp
ROHs for S49 (Sex: M, Reference: M. martes)
* MMAR.chr1
MMAR.chr2
* MMAR.chr3
MMAR.chr4
* MMAR.chr5
* MMAR.chr6
* MMAR.chr7
* MMAR.chr8
* MMAR.chr9
MMAR.chr10
MMAR.chr11
MMAR.chr12
* MMAR.chr13
* MMAR.chr14
MMAR.chr15
MMAR.chr16
* MMAR.chr17
* MMAR.chr18
* MMAR.chrX
0 Mbp
50 Mbp
100 Mbp
150 Mbp
200 Mbp

# S50

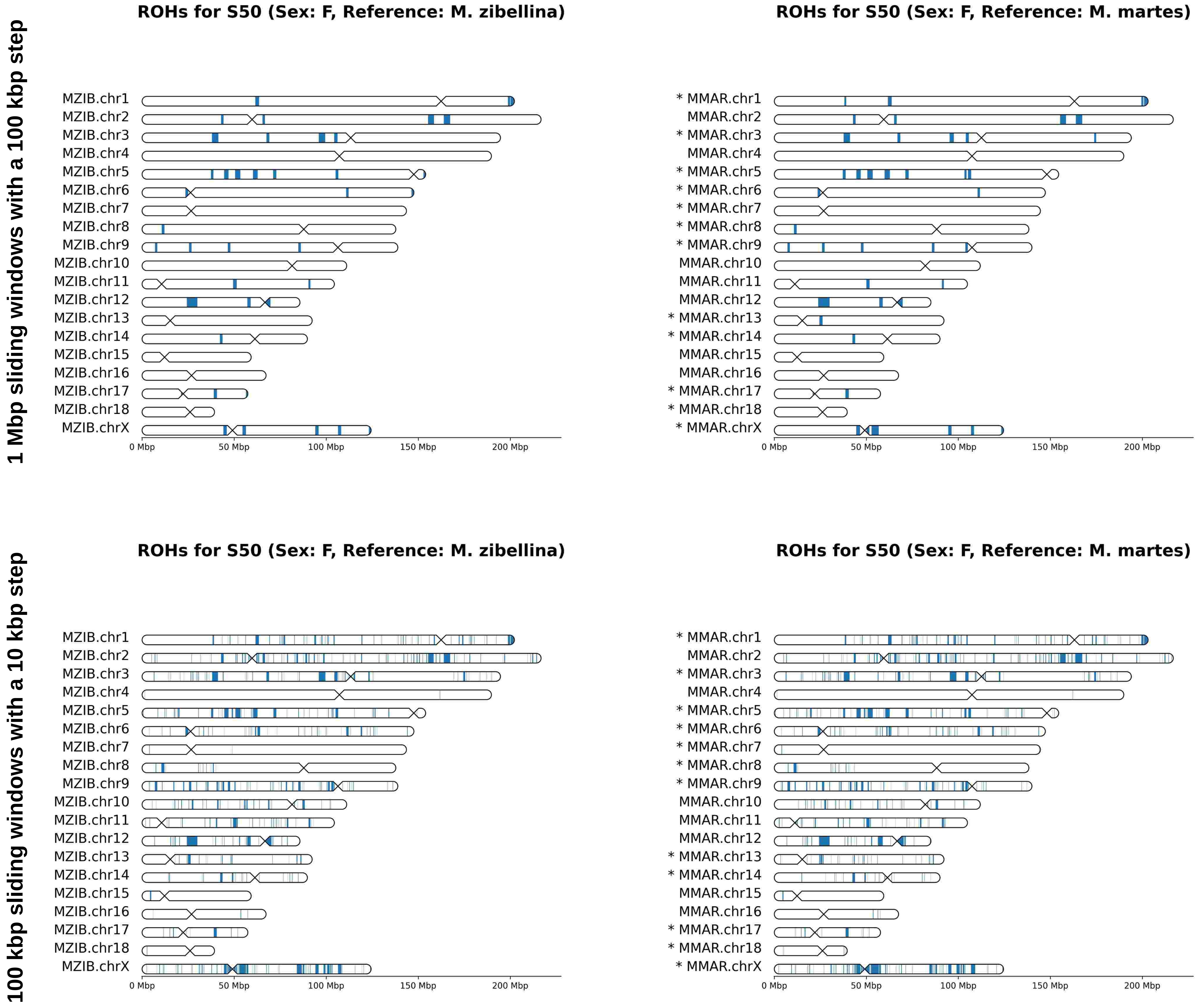
1 Mbp sliding windows with a 100 kbp step
100 kbp sliding windows with a 10 kbp step
ROHs for S50 (Sex: F, Reference: M. zibellina)
MZIB.chr1
MZIB.chr2
MZIB.chr3
MZIB.chr4
MZIB.chr5
MZIB.chr6
MZIB.chr7
MZIB.chr8
MZIB.chr9
MZIB.chr10
MZIB.chr11
MZIB.chr12
MZIB.chr13
MZIB.chr14
MZIB.chr15
MZIB.chr16
MZIB.chr17
MZIB.chr18
MZIB.chrX
0 Mbp
50 Mbp
100 Mbp
150 Mbp
200 Mbp
ROHs for S50 (Sex: F, Reference: M. martes)
* MMAR.chr1
MMAR.chr2
* MMAR.chr3
MMAR.chr4
* MMAR.chr5
* MMAR.chr6
* MMAR.chr7
* MMAR.chr8
* MMAR.chr9
MMAR.chr10
MMAR.chr11
MMAR.chr12
* MMAR.chr13
* MMAR.chr14
MMAR.chr15
MMAR.chr16
* MMAR.chr17
* MMAR.chr18
* MMAR.chrX
0 Mbp
50 Mbp
100 Mbp
150 Mbp
200 Mbp

# T8

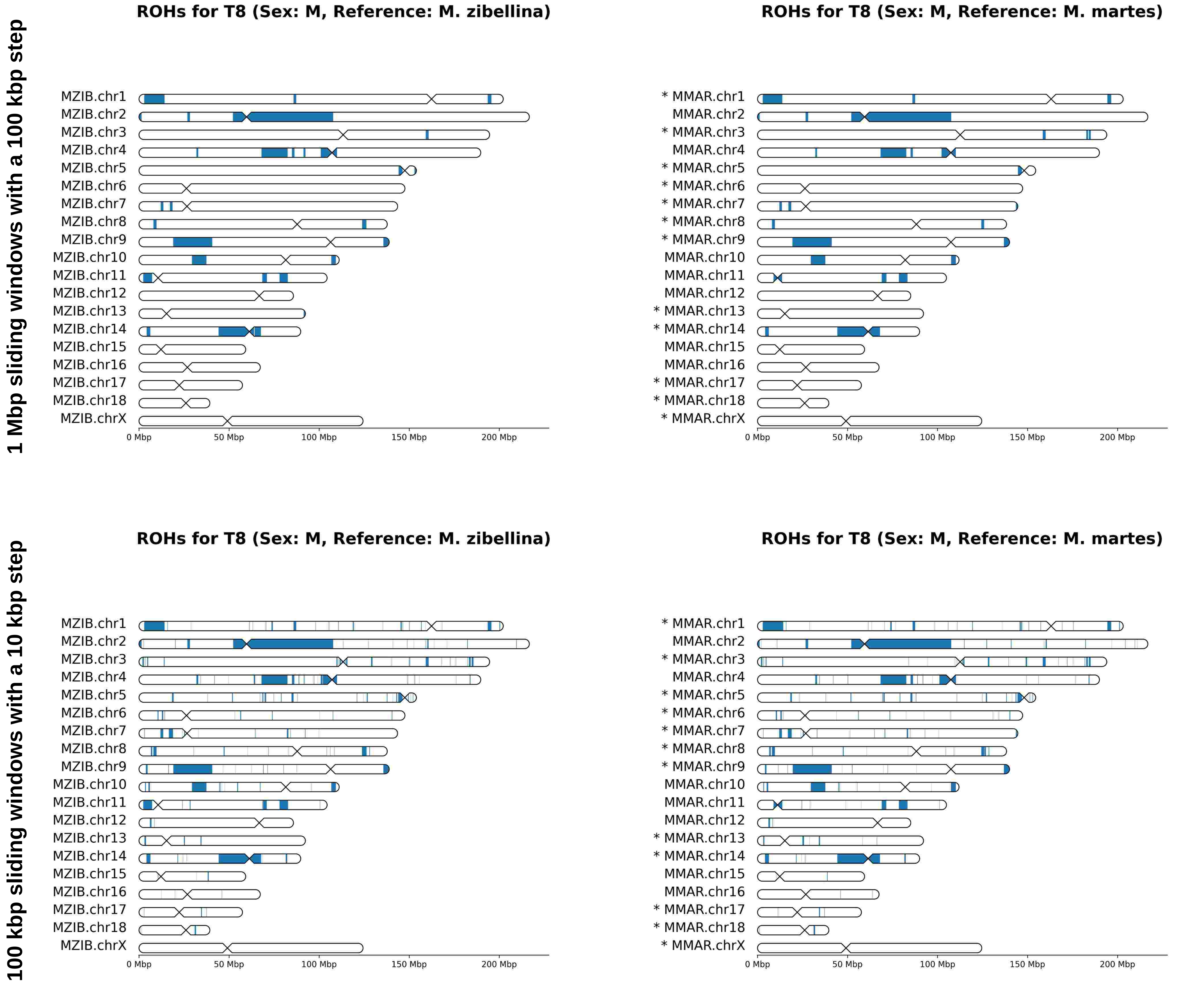

1 Mbp sliding windows with a 100 kbp step
100 kbp sliding windows with a 10 kbp step
ROHs for T8 (Sex: M, Reference: M. zibellina)
ROHs for T8 (Sex: M, Reference: M. martes)
MZIB.chr1
MZIB.chr2
MZIB.chr3
MZIB.chr4
MZIB.chr5
MZIB.chr6
MZIB.chr7
MZIB.chr8
MZIB.chr9
MZIB.chr10
MZIB.chr11
MZIB.chr12
MZIB.chr13
MZIB.chr14
MZIB.chr15
MZIB.chr16
MZIB.chr17
MZIB.chr18
MZIB.chrX
* MMAR.chr1
MMAR.chr2
* MMAR.chr3
MMAR.chr4
* MMAR.chr5
* MMAR.chr6
* MMAR.chr7
* MMAR.chr8
* MMAR.chr9
MMAR.chr10
MMAR.chr11
MMAR.chr12
* MMAR.chr13
* MMAR.chr14
MMAR.chr15
MMAR.chr16
* MMAR.chr17
* MMAR.chr18
* MMAR.chrX
0 Mbp
50 Mbp
100 Mbp
150 Mbp
200 Mbp

# T18

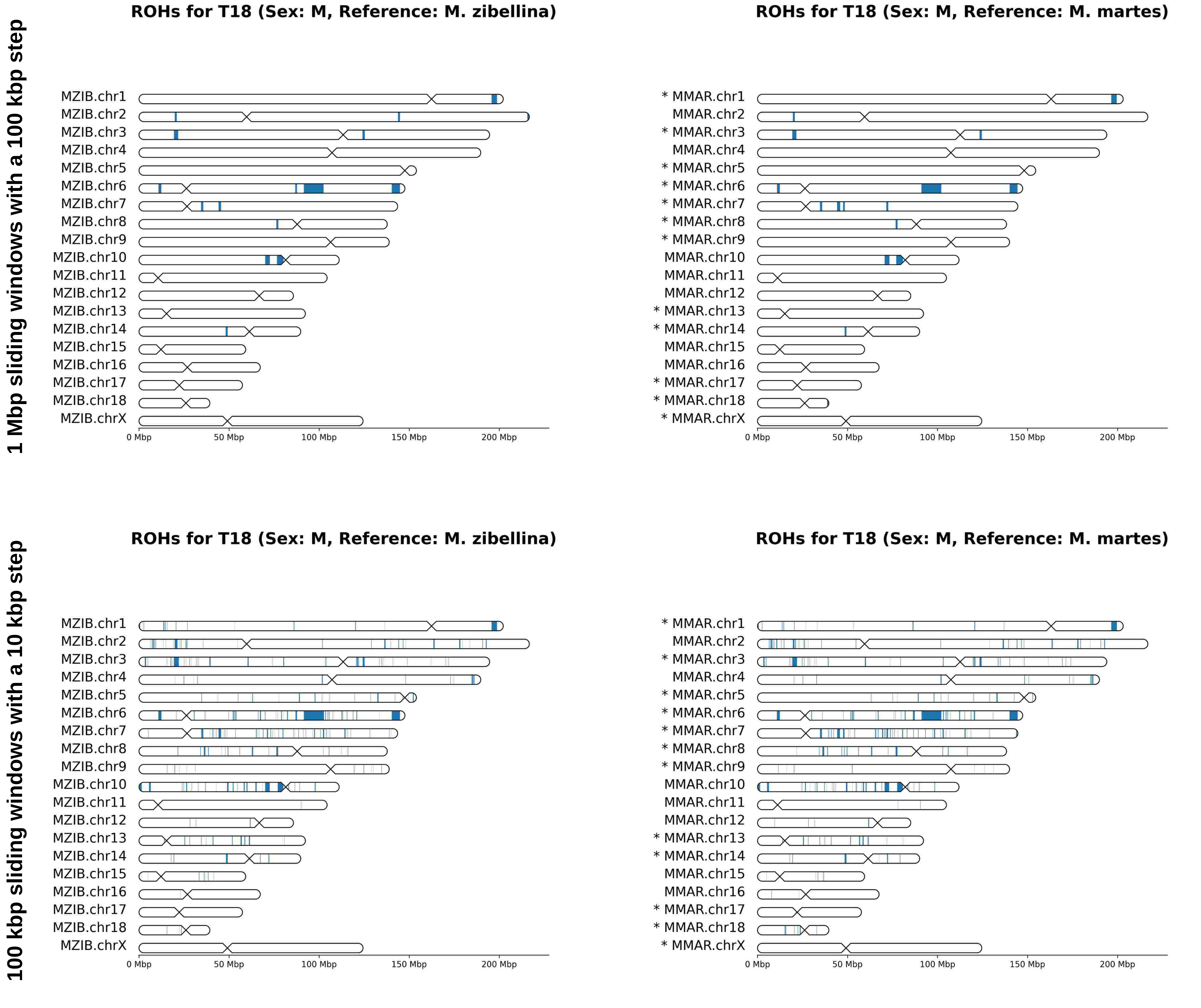
1 Mbp sliding windows with a 100 kbp step
100 kbp sliding windows with a 10 kbp step
ROHs for T18 (Sex: M, Reference: M. zibellina)
MZIB.chr1
MZIB.chr2
MZIB.chr3
MZIB.chr4
MZIB.chr5
MZIB.chr6
MZIB.chr7
MZIB.chr8
MZIB.chr9
MZIB.chr10
MZIB.chr11
MZIB.chr12
MZIB.chr13
MZIB.chr14
MZIB.chr15
MZIB.chr16
MZIB.chr17
MZIB.chr18
MZIB.chrX
0 Mbp
50 Mbp
100 Mbp
150 Mbp
200 Mbp
ROHs for T18 (Sex: M, Reference: M. martes)
* MMAR.chr1
MMAR.chr2
* MMAR.chr3
MMAR.chr4
* MMAR.chr5
* MMAR.chr6
* MMAR.chr7
* MMAR.chr8
* MMAR.chr9
MMAR.chr10
MMAR.chr11
MMAR.chr12
* MMAR.chr13
* MMAR.chr14
MMAR.chr15
MMAR.chr16
* MMAR.chr17
* MMAR.chr18
* MMAR.chrX
0 Mbp
50 Mbp
100 Mbp
150 Mbp
200 Mbp

# T24

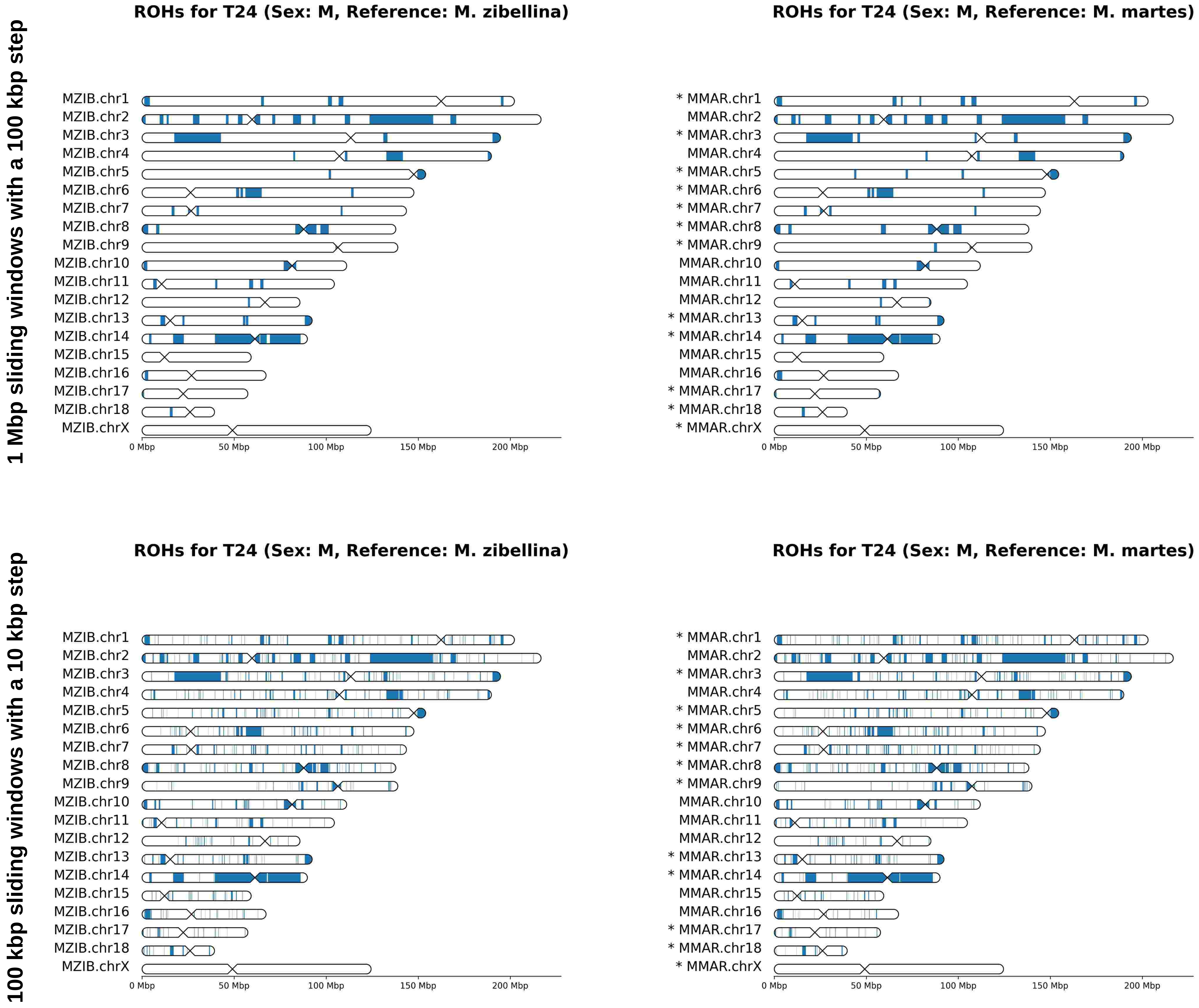
1 Mbp sliding windows with a 100 kbp step
100 kbp sliding windows with a 10 kbp step
ROHs for T24 (Sex: M, Reference: M. zibellina)
ROHs for T24 (Sex: M, Reference: M. martes)
MZIB.chr1
MZIB.chr2
MZIB.chr3
MZIB.chr4
MZIB.chr5
MZIB.chr6
MZIB.chr7
MZIB.chr8
MZIB.chr9
MZIB.chr10
MZIB.chr11
MZIB.chr12
MZIB.chr13
MZIB.chr14
MZIB.chr15
MZIB.chr16
MZIB.chr17
MZIB.chr18
MZIB.chrX
* MMAR.chr1
MMAR.chr2
* MMAR.chr3
MMAR.chr4
* MMAR.chr5
* MMAR.chr6
* MMAR.chr7
* MMAR.chr8
* MMAR.chr9
MMAR.chr10
MMAR.chr11
MMAR.chr12
* MMAR.chr13
* MMAR.chr14
MMAR.chr15
MMAR.chr16
* MMAR.chr17
* MMAR.chr18
* MMAR.chrX
0 Mbp
50 Mbp
100 Mbp
150 Mbp
200 Mbp

# T26

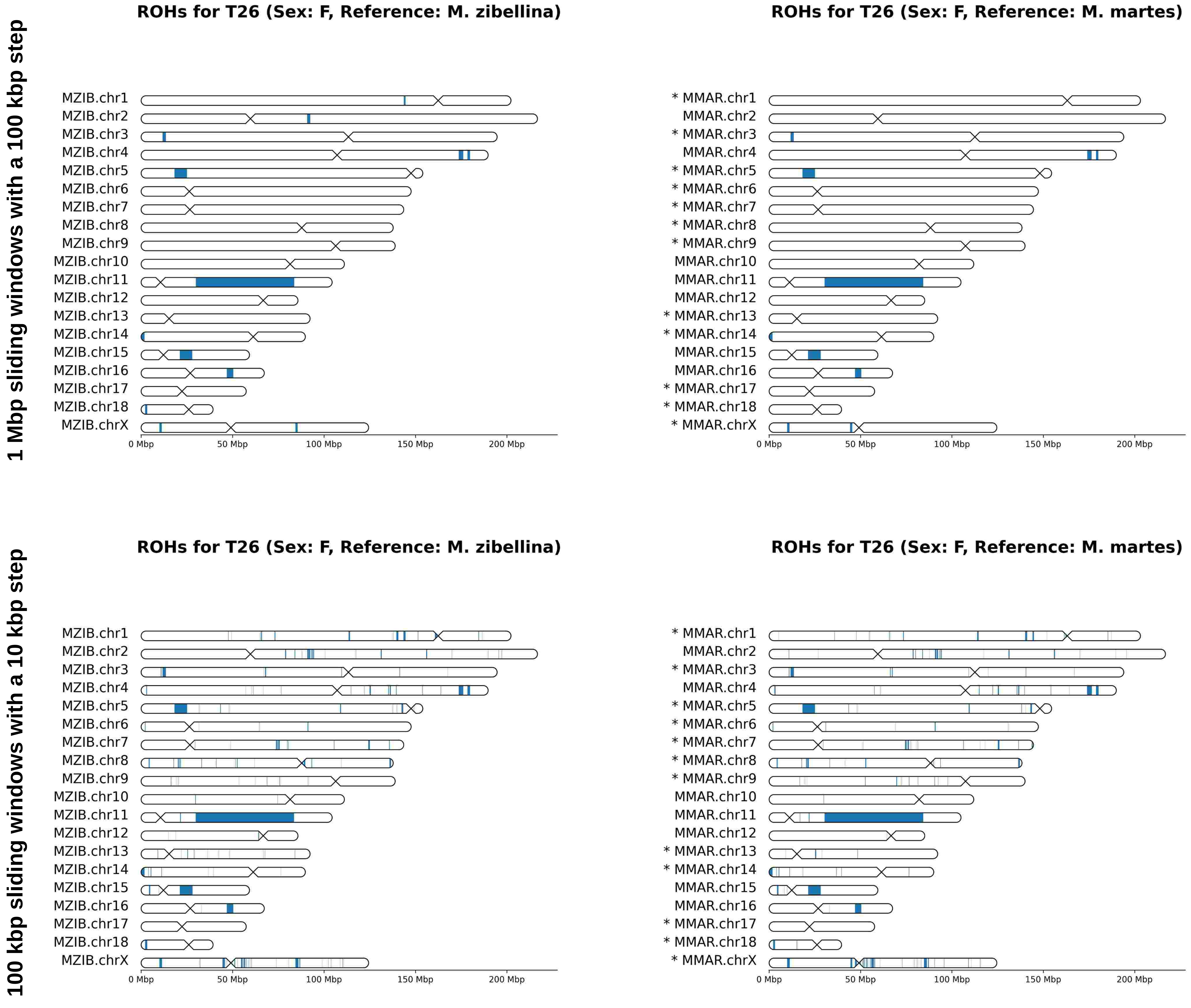
1 Mbp sliding windows with a 100 kbp step
100 kbp sliding windows with a 10 kbp step
ROHs for T26 (Sex: F, Reference: M. zibellina)
MZIB.chr1
MZIB.chr2
MZIB.chr3
MZIB.chr4
MZIB.chr5
MZIB.chr6
MZIB.chr7
MZIB.chr8
MZIB.chr9
MZIB.chr10
MZIB.chr11
MZIB.chr12
MZIB.chr13
MZIB.chr14
MZIB.chr15
MZIB.chr16
MZIB.chr17
MZIB.chr18
MZIB.chrX
0 Mbp
50 Mbp
100 Mbp
150 Mbp
200 Mbp
ROHs for T26 (Sex: F, Reference: M. martes)
* MMAR.chr1
MMAR.chr2
* MMAR.chr3
MMAR.chr4
* MMAR.chr5
* MMAR.chr6
* MMAR.chr7
* MMAR.chr8
* MMAR.chr9
MMAR.chr10
MMAR.chr11
MMAR.chr12
* MMAR.chr13
* MMAR.chr14
MMAR.chr15
MMAR.chr16
* MMAR.chr17
* MMAR.chr18
* MMAR.chrX

# T50

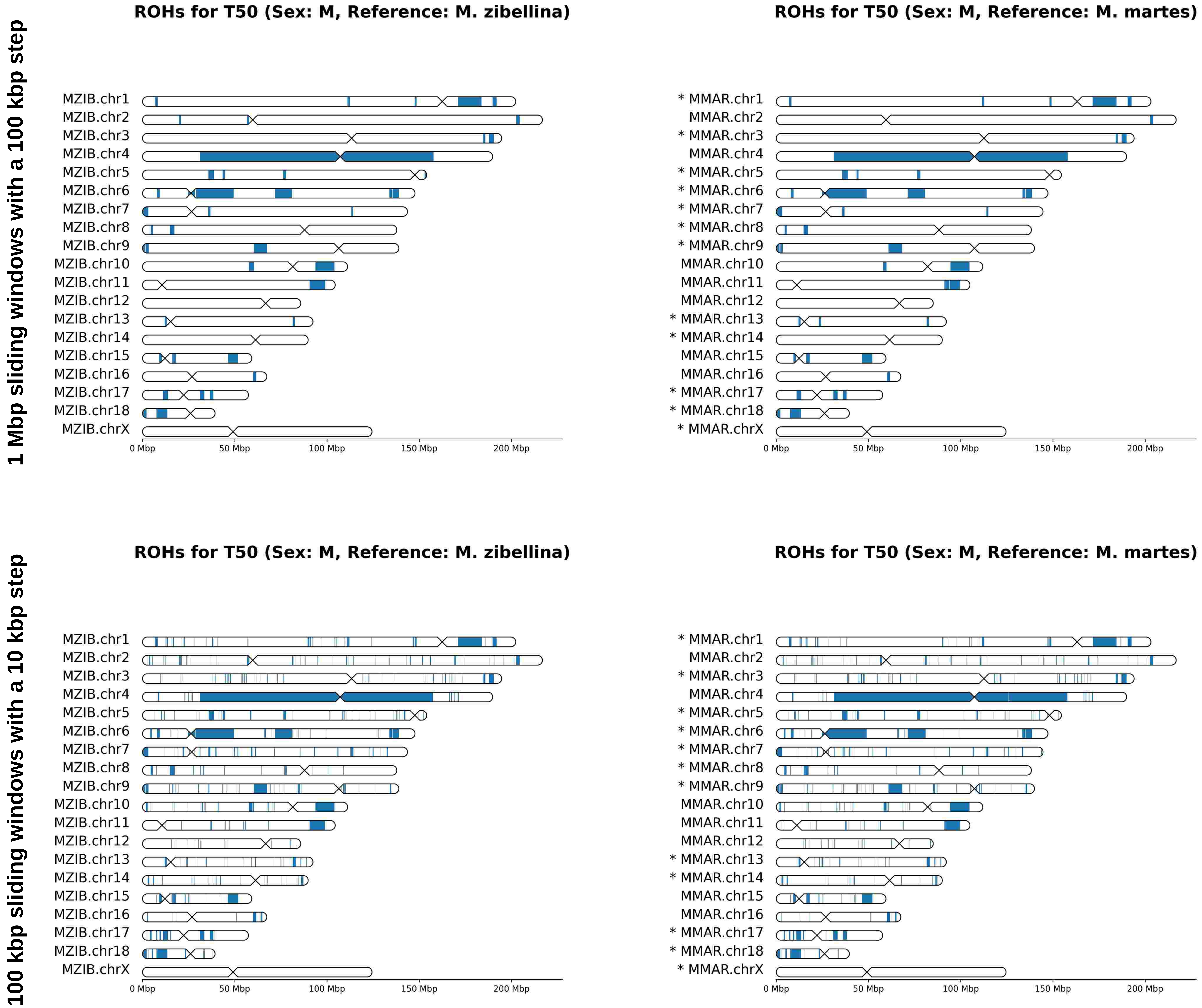
1 Mbp sliding windows with a 100 kbp step
100 kbp sliding windows with a 10 kbp step
ROHs for T50 (Sex: M, Reference: M. zibellina)
ROHs for T50 (Sex: M, Reference: M. martes)
MZIB.chr1
MZIB.chr2
MZIB.chr3
MZIB.chr4
MZIB.chr5
MZIB.chr6
MZIB.chr7
MZIB.chr8
MZIB.chr9
MZIB.chr10
MZIB.chr11
MZIB.chr12
MZIB.chr13
MZIB.chr14
MZIB.chr15
MZIB.chr16
MZIB.chr17
MZIB.chr18
MZIB.chrX
* MMAR.chr1
MMAR.chr2
* MMAR.chr3
MMAR.chr4
* MMAR.chr5
* MMAR.chr6
* MMAR.chr7
* MMAR.chr8
* MMAR.chr9
MMAR.chr10
MMAR.chr11
MMAR.chr12
* MMAR.chr13
* MMAR.chr14
MMAR.chr15
MMAR.chr16
* MMAR.chr17
* MMAR.chr18
* MMAR.chrX
0 Mbp
50 Mbp
100 Mbp
150 Mbp
200 Mbp

# T72

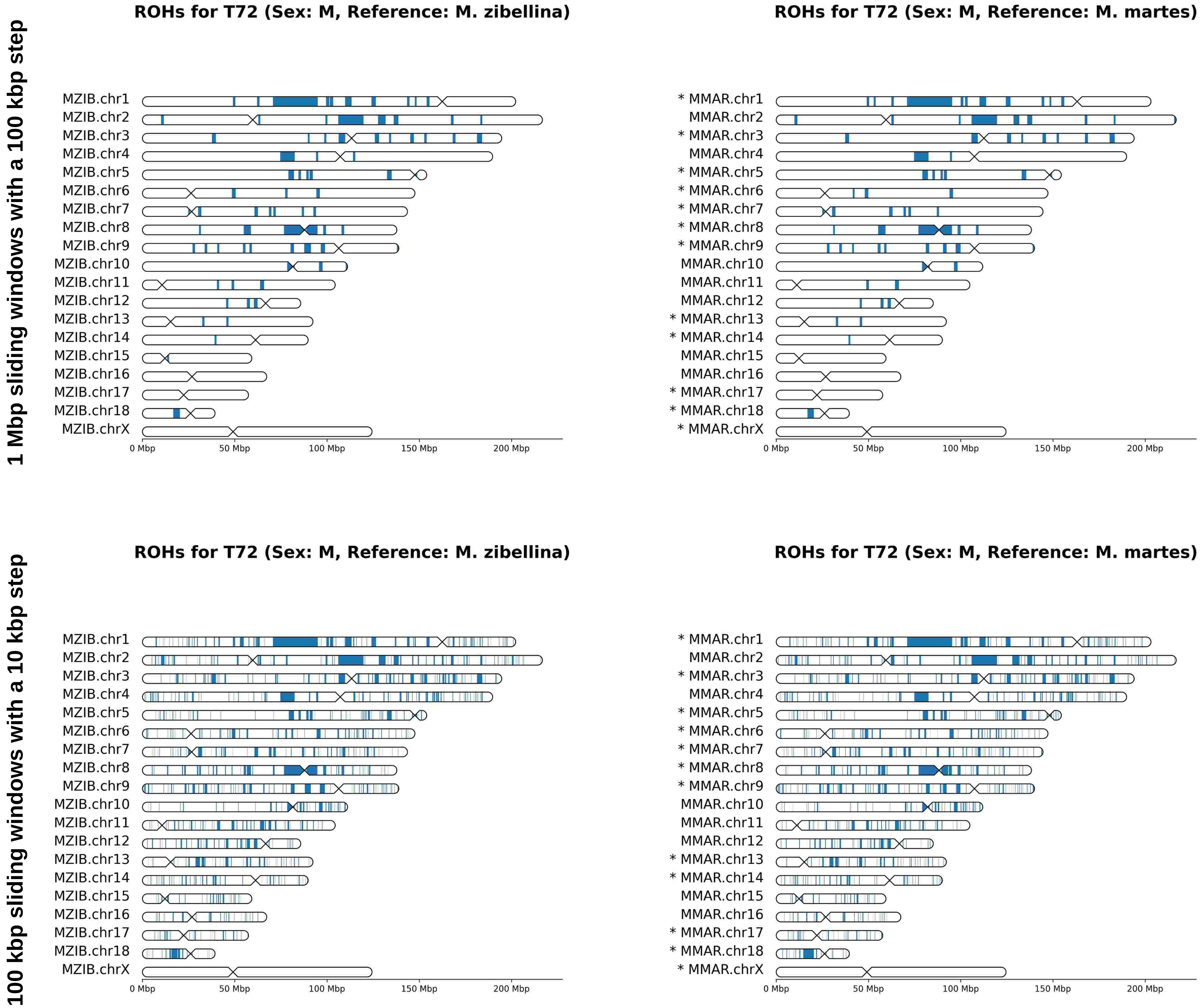
1 Mbp sliding windows with a 100 kbp step
100 kbp sliding windows with a 10 kbp step
ROHs for T72 (Sex: M, Reference: M. zibellina)
ROHs for T72 (Sex: M, Reference: M. martes)
MZIB.chr1
MZIB.chr2
MZIB.chr3
MZIB.chr4
MZIB.chr5
MZIB.chr6
MZIB.chr7
MZIB.chr8
MZIB.chr9
MZIB.chr10
MZIB.chr11
MZIB.chr12
MZIB.chr13
MZIB.chr14
MZIB.chr15
MZIB.chr16
MZIB.chr17
MZIB.chr18
MZIB.chrX
* MMAR.chr1
MMAR.chr2
* MMAR.chr3
MMAR.chr4
* MMAR.chr5
* MMAR.chr6
* MMAR.chr7
* MMAR.chr8
* MMAR.chr9
MMAR.chr10
MMAR.chr11
MMAR.chr12
* MMAR.chr13
* MMAR.chr14
MMAR.chr15
MMAR.chr16
* MMAR.chr17
* MMAR.chr18
* MMAR.chrX
0 Mbp
50 Mbp
100 Mbp
150 Mbp
200 Mbp

# T76

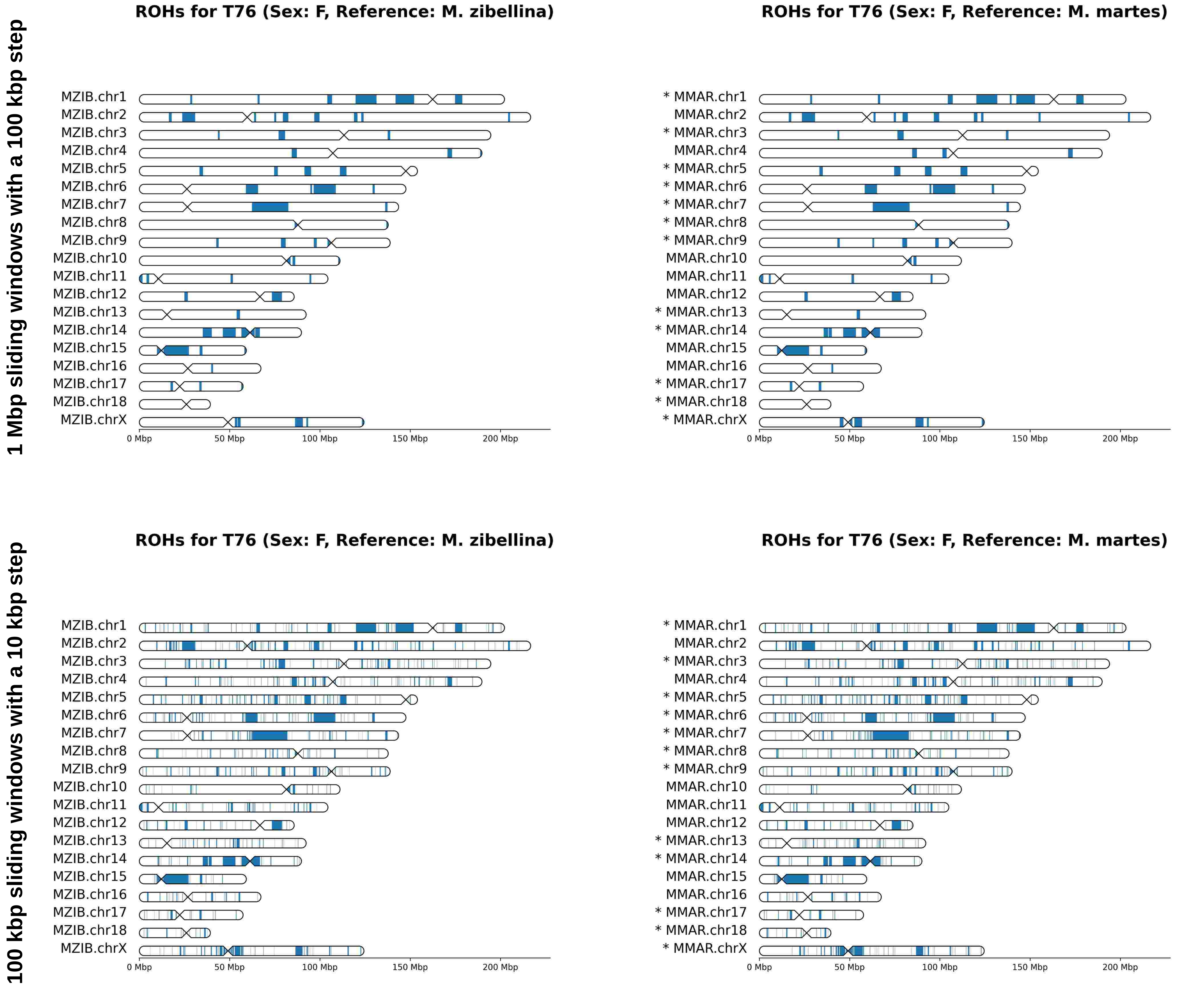
1 Mbp sliding windows with a 100 kbp step
100 kbp sliding windows with a 10 kbp step
ROHs for T76 (Sex: F, Reference: M. zibellina)
ROHs for T76 (Sex: F, Reference: M. martes)
MZIB.chr1
MZIB.chr2
MZIB.chr3
MZIB.chr4
MZIB.chr5
MZIB.chr6
MZIB.chr7
MZIB.chr8
MZIB.chr9
MZIB.chr10
MZIB.chr11
MZIB.chr12
MZIB.chr13
MZIB.chr14
MZIB.chr15
MZIB.chr16
MZIB.chr17
MZIB.chr18
MZIB.chrX
* MMAR.chr1
MMAR.chr2
* MMAR.chr3
MMAR.chr4
* MMAR.chr5
* MMAR.chr6
* MMAR.chr7
* MMAR.chr8
* MMAR.chr9
MMAR.chr10
MMAR.chr11
MMAR.chr12
* MMAR.chr13
* MMAR.chr14
MMAR.chr15
MMAR.chr16
* MMAR.chr17
* MMAR.chr18
* MMAR.chrX
0 Mbp
50 Mbp
100 Mbp
150 Mbp
200 Mbp

# T77

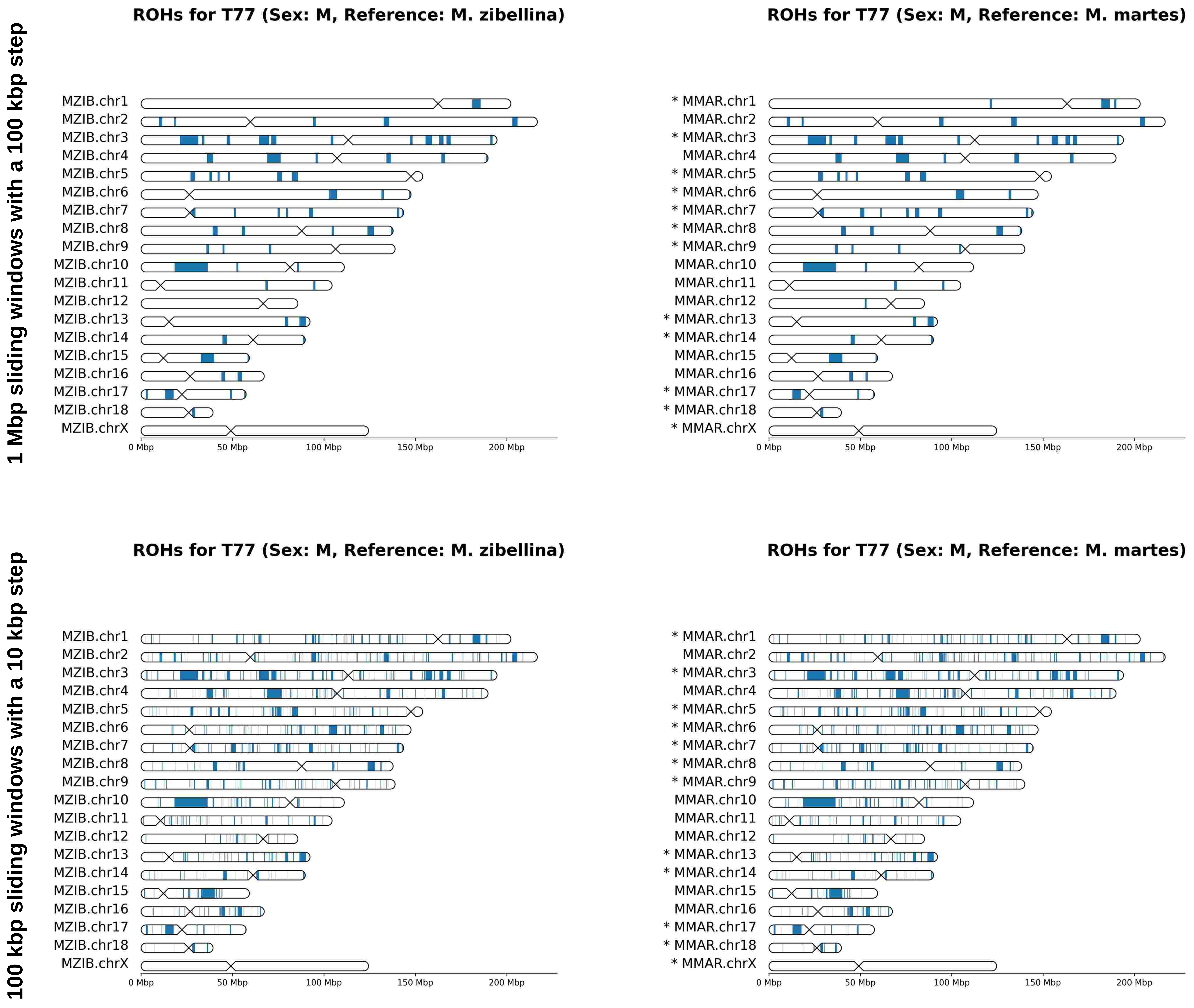
1 Mbp sliding windows with a 100 kbp step
100 kbp sliding windows with a 10 kbp step
ROHs for T77 (Sex: M, Reference: M. zibellina)
MZIB.chr1
MZIB.chr2
MZIB.chr3
MZIB.chr4
MZIB.chr5
MZIB.chr6
MZIB.chr7
MZIB.chr8
MZIB.chr9
MZIB.chr10
MZIB.chr11
MZIB.chr12
MZIB.chr13
MZIB.chr14
MZIB.chr15
MZIB.chr16
MZIB.chr17
MZIB.chr18
MZIB.chrX
0 Mbp
50 Mbp
100 Mbp
150 Mbp
200 Mbp
ROHs for T77 (Sex: M, Reference: M. martes)
* MMAR.chr1
MMAR.chr2
* MMAR.chr3
MMAR.chr4
* MMAR.chr5
* MMAR.chr6
* MMAR.chr7
* MMAR.chr8
* MMAR.chr9
MMAR.chr10
MMAR.chr11
MMAR.chr12
* MMAR.chr13
* MMAR.chr14
MMAR.chr15
MMAR.chr16
* MMAR.chr17
* MMAR.chr18
* MMAR.chrX
0 Mbp
50 Mbp
100 Mbp
150 Mbp
200 Mbp
ROHs for T77 (Sex: M, Reference: M. zibellina)
MZIB.chr1
MZIB.chr2
MZIB.chr3
MZIB.chr4
MZIB.chr5
MZIB.chr6
MZIB.chr7
MZIB.chr8
MZIB.chr9
MZIB.chr10
MZIB.chr11
MZIB.chr12
MZIB.chr13
MZIB.chr14
MZIB.chr15
MZIB.chr16
MZIB.chr17
MZIB.chr18
MZIB.chrX
0 Mbp
50 Mbp
100 Mbp
150 Mbp
200 Mbp
ROHs for T77 (Sex: M, Reference: M. martes)
* MMAR.chr1
MMAR.chr2
* MMAR.chr3
MMAR.chr4
* MMAR.chr5
* MMAR.chr6
* MMAR.chr7
* MMAR.chr8
* MMAR.chr9
MMAR.chr10
MMAR.chr11
MMAR.chr12
* MMAR.chr13
* MMAR.chr14
MMAR.chr15
MMAR.chr16
* MMAR.chr17
* MMAR.chr18
* MMAR.chrX
0 Mbp
50 Mbp
100 Mbp
150 Mbp
200 Mbp

# T78

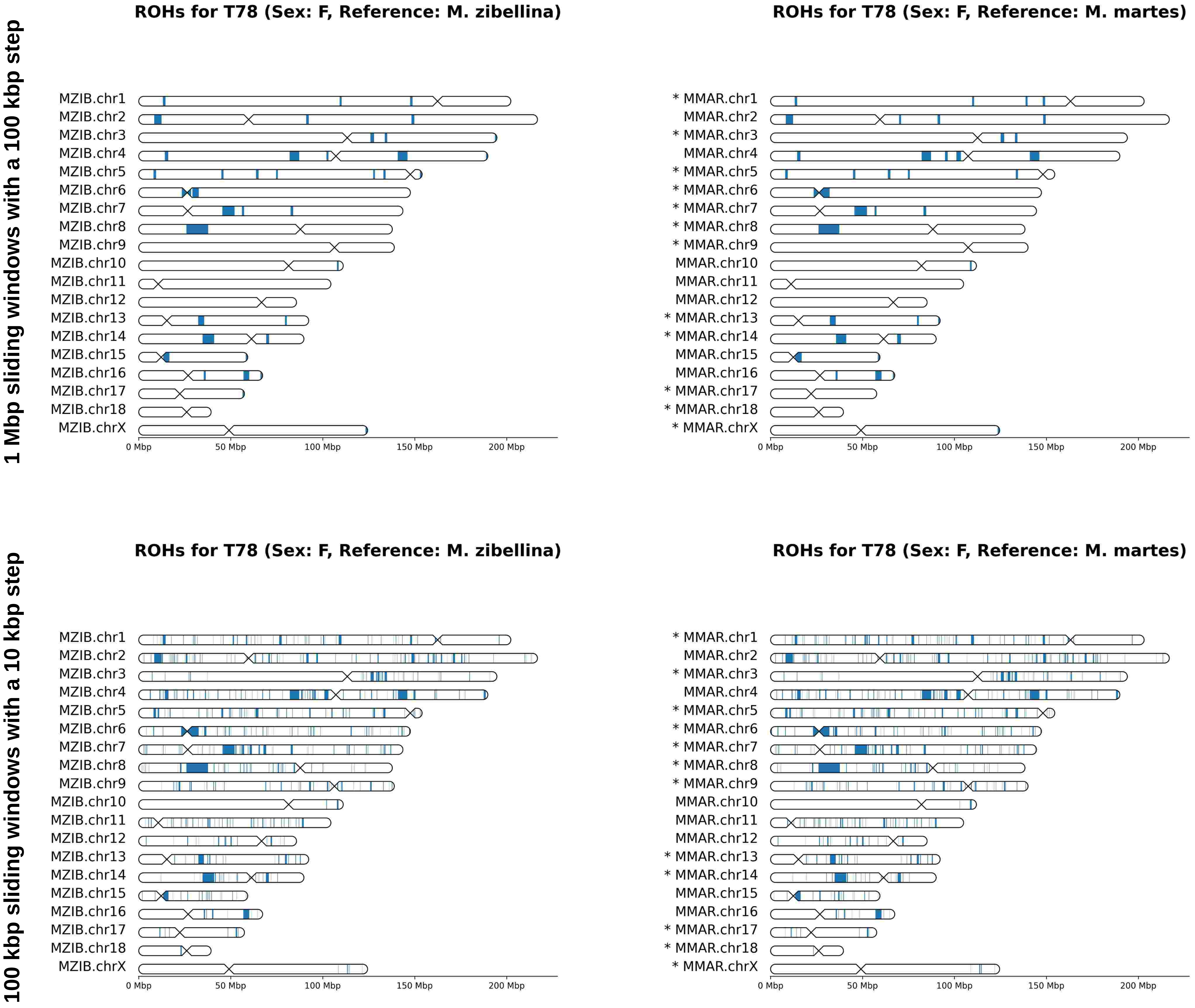
1 Mbp sliding windows with a 100 kbp step
100 kbp sliding windows with a 10 kbp step
ROHs for T78 (Sex: F, Reference: M. zibellina)
ROHs for T78 (Sex: F, Reference: M. martes)
MZIB.chr1
MZIB.chr2
MZIB.chr3
MZIB.chr4
MZIB.chr5
MZIB.chr6
MZIB.chr7
MZIB.chr8
MZIB.chr9
MZIB.chr10
MZIB.chr11
MZIB.chr12
MZIB.chr13
MZIB.chr14
MZIB.chr15
MZIB.chr16
MZIB.chr17
MZIB.chr18
MZIB.chrX
* MMAR.chr1
MMAR.chr2
* MMAR.chr3
MMAR.chr4
* MMAR.chr5
* MMAR.chr6
* MMAR.chr7
* MMAR.chr8
* MMAR.chr9
MMAR.chr10
MMAR.chr11
MMAR.chr12
* MMAR.chr13
* MMAR.chr14
MMAR.chr15
MMAR.chr16
* MMAR.chr17
* MMAR.chr18
* MMAR.chrX
0 Mbp
50 Mbp
100 Mbp
150 Mbp
200 Mbp

# T79

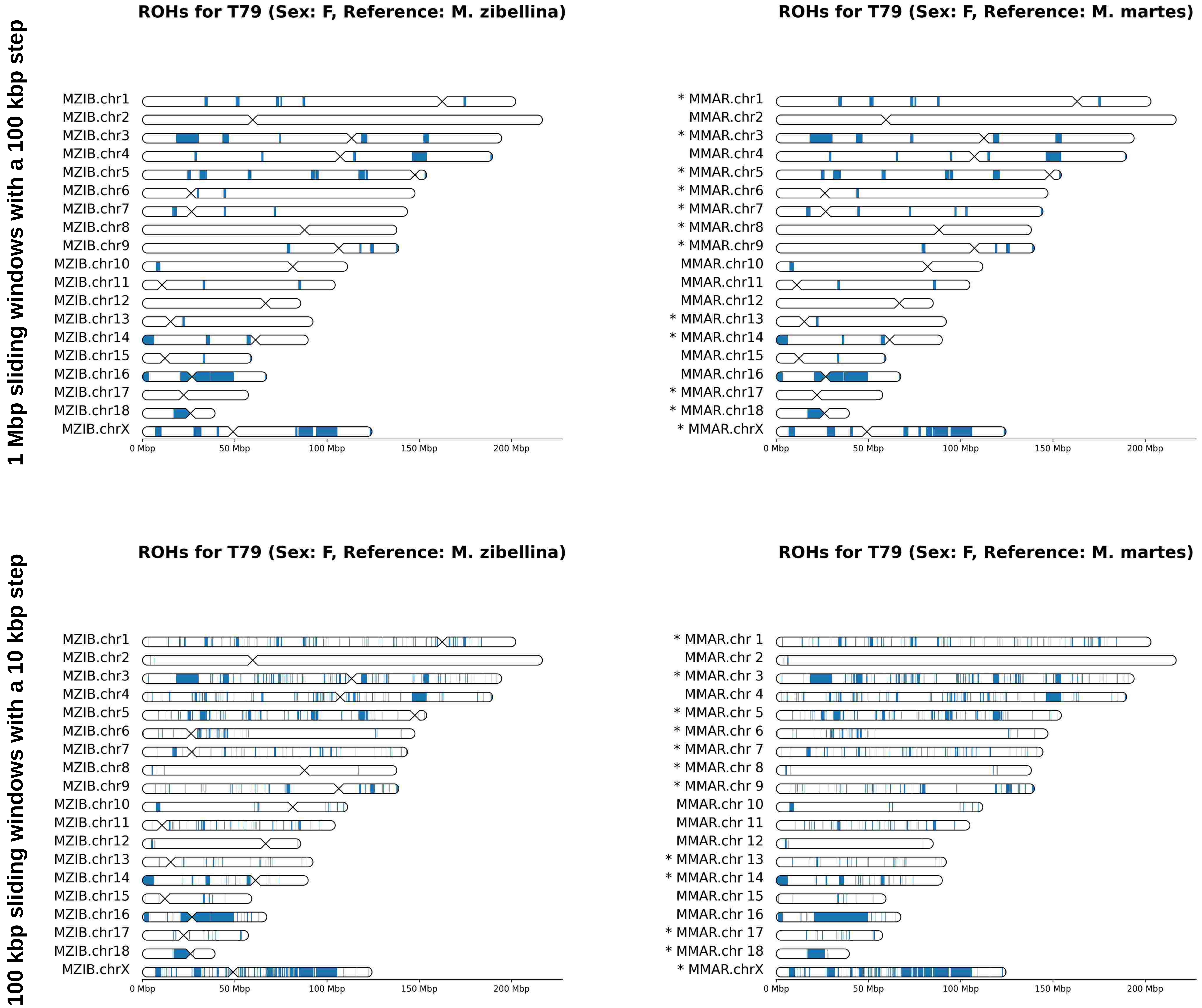
1 Mbp sliding windows with a 100 kbp step
100 kbp sliding windows with a 10 kbp step
ROHs for T79 (Sex: F, Reference: M. zibellina)
ROHs for T79 (Sex: F, Reference: M. martes)
MZIB.chr1
MZIB.chr2
MZIB.chr3
MZIB.chr4
MZIB.chr5
MZIB.chr6
MZIB.chr7
MZIB.chr8
MZIB.chr9
MZIB.chr10
MZIB.chr11
MZIB.chr12
MZIB.chr13
MZIB.chr14
MZIB.chr15
MZIB.chr16
MZIB.chr17
MZIB.chr18
MZIB.chrX
* MMAR.chr1
MMAR.chr2
* MMAR.chr3
MMAR.chr4
* MMAR.chr5
* MMAR.chr6
* MMAR.chr7
* MMAR.chr8
* MMAR.chr9
MMAR.chr10
MMAR.chr11
MMAR.chr12
* MMAR.chr13
* MMAR.chr14
MMAR.chr15
MMAR.chr16
* MMAR.chr17
* MMAR.chr18
* MMAR.chrX
* MMAR.chr 1
MMAR.chr 2
* MMAR.chr 3
MMAR.chr 4
* MMAR.chr 5
* MMAR.chr 6
* MMAR.chr 7
* MMAR.chr 8
* MMAR.chr 9
MMAR.chr 10
MMAR.chr 11
MMAR.chr 12
* MMAR.chr 13
* MMAR.chr 14
MMAR.chr 15
MMAR.chr 16
* MMAR.chr 17
* MMAR.chr 18
0 Mbp
50 Mbp
100 Mbp
150 Mbp
200 Mbp

# T81

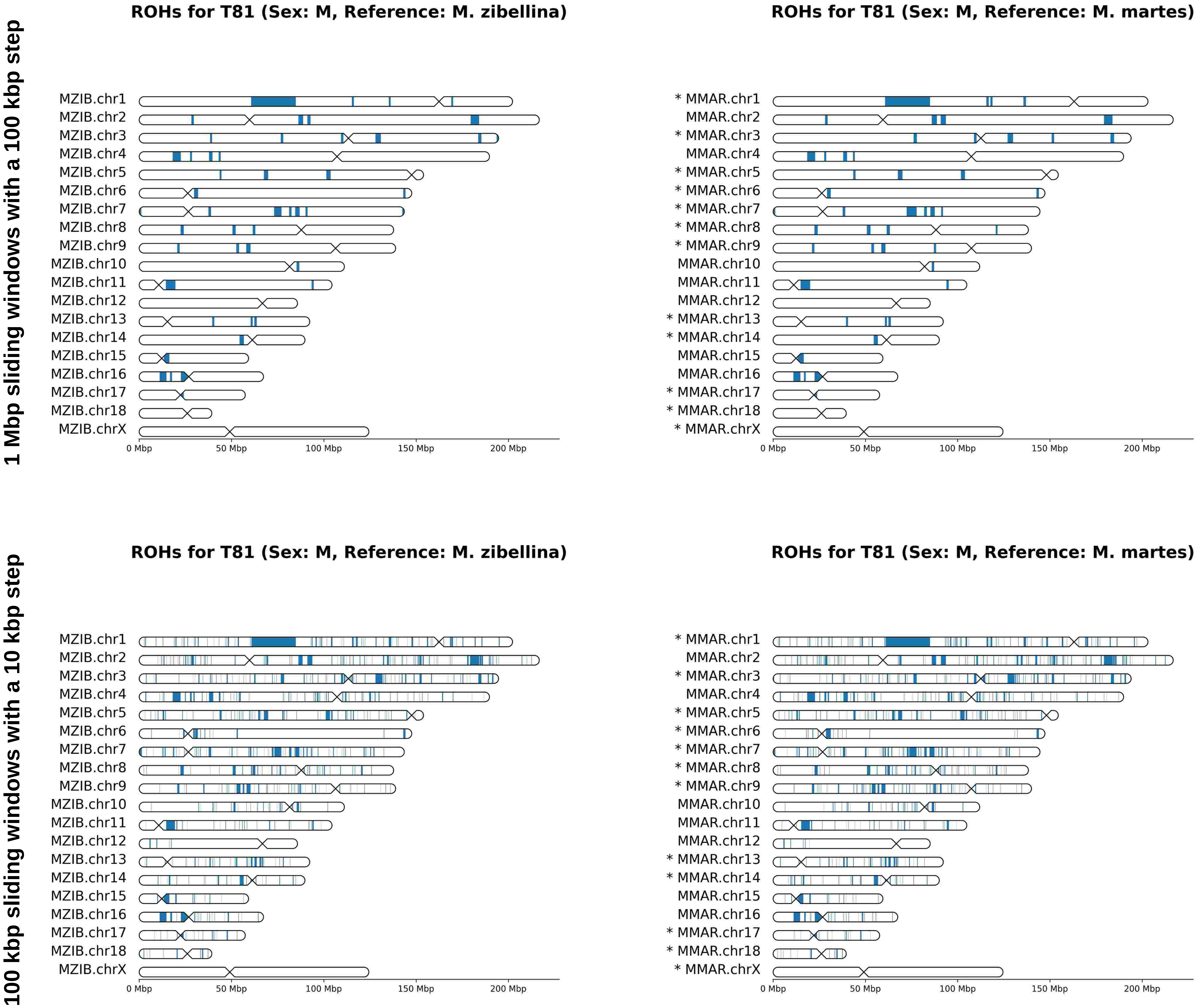
1 Mbp sliding windows with a 100 kbp step
100 kbp sliding windows with a 10 kbp step
ROHs for T81 (Sex: M, Reference: M. zibellina)
ROHs for T81 (Sex: M, Reference: M. martes)
MZIB.chr1
MZIB.chr2
MZIB.chr3
MZIB.chr4
MZIB.chr5
MZIB.chr6
MZIB.chr7
MZIB.chr8
MZIB.chr9
MZIB.chr10
MZIB.chr11
MZIB.chr12
MZIB.chr13
MZIB.chr14
MZIB.chr15
MZIB.chr16
MZIB.chr17
MZIB.chr18
MZIB.chrX
* MMAR.chr1
MMAR.chr2
* MMAR.chr3
MMAR.chr4
* MMAR.chr5
* MMAR.chr6
* MMAR.chr7
* MMAR.chr8
* MMAR.chr9
MMAR.chr10
MMAR.chr11
MMAR.chr12
* MMAR.chr13
* MMAR.chr14
MMAR.chr15
MMAR.chr16
* MMAR.chr17
* MMAR.chr18
* MMAR.chrX
0 Mbp
50 Mbp
100 Mbp
150 Mbp
200 Mbp

# T82

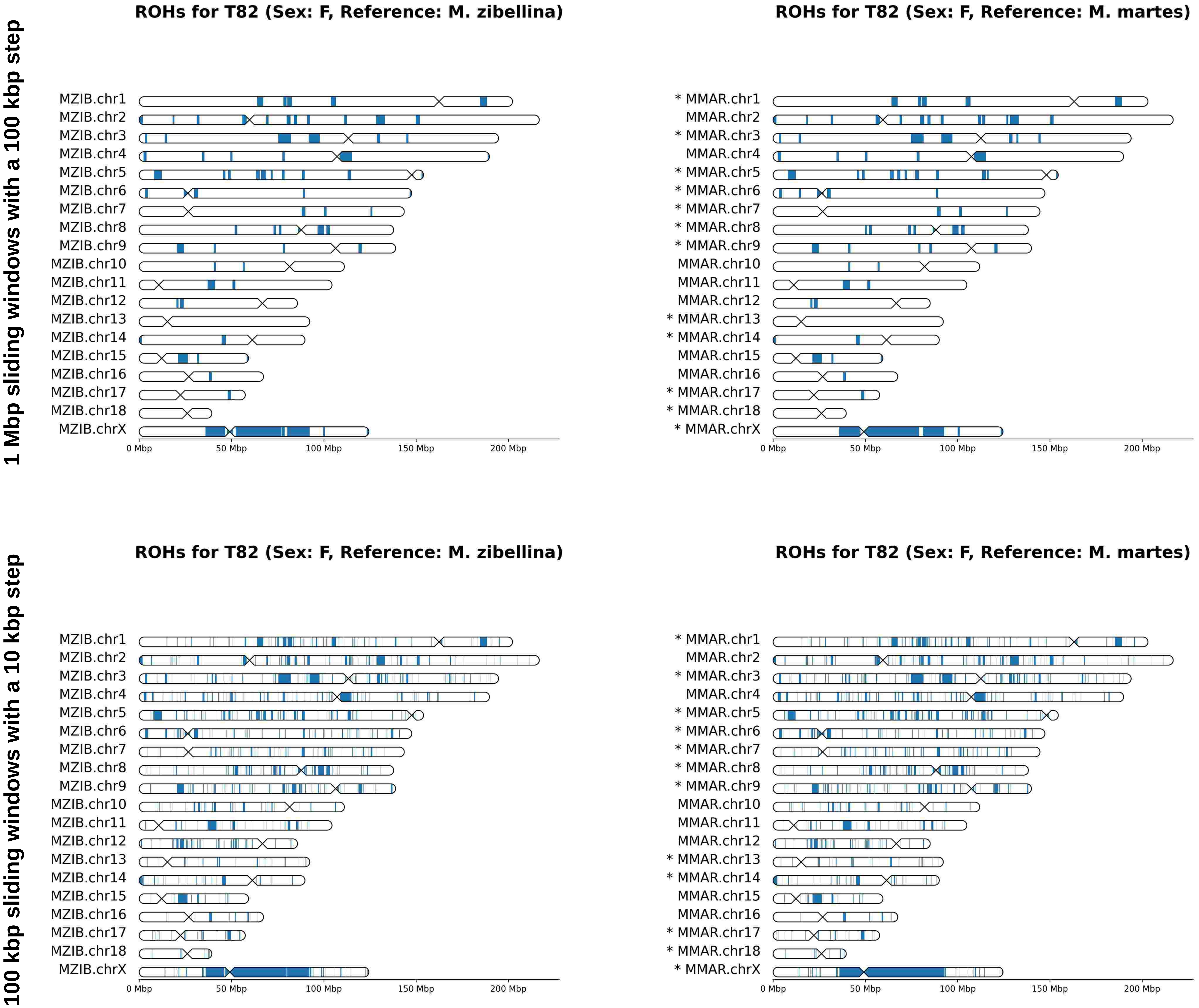
1 Mbp sliding windows with a 100 kbp step
100 kbp sliding windows with a 10 kbp step
ROHs for T82 (Sex: F, Reference: M. zibellina)
MZIB.chr1
MZIB.chr2
MZIB.chr3
MZIB.chr4
MZIB.chr5
MZIB.chr6
MZIB.chr7
MZIB.chr8
MZIB.chr9
MZIB.chr10
MZIB.chr11
MZIB.chr12
MZIB.chr13
MZIB.chr14
MZIB.chr15
MZIB.chr16
MZIB.chr17
MZIB.chr18
MZIB.chrX
ROHs for T82 (Sex: F, Reference: M. martes)
* MMAR.chr1
MMAR.chr2
* MMAR.chr3
MMAR.chr4
* MMAR.chr5
* MMAR.chr6
* MMAR.chr7
* MMAR.chr8
* MMAR.chr9
MMAR.chr10
MMAR.chr11
MMAR.chr12
* MMAR.chr13
* MMAR.chr14
MMAR.chr15
MMAR.chr16
* MMAR.chr17
* MMAR.chr18
* MMAR.chrX
0 Mbp
50 Mbp
100 Mbp
150 Mbp
200 Mbp

# T83

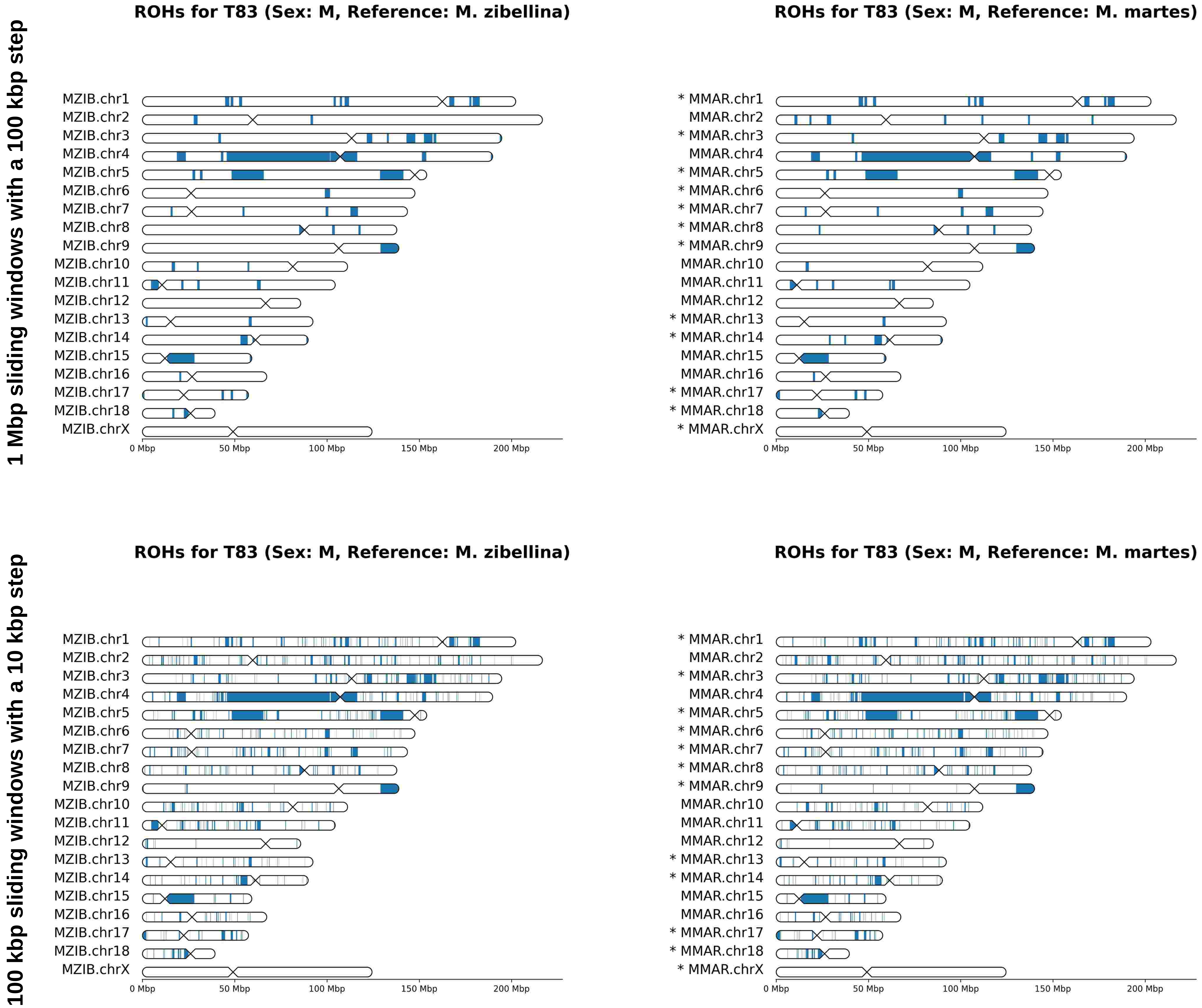
1 Mbp sliding windows with a 100 kbp step
100 kbp sliding windows with a 10 kbp step
ROHs for T83 (Sex: M, Reference: M. zibellina)
ROHs for T83 (Sex: M, Reference: M. martes)
MZIB.chr1
MZIB.chr2
MZIB.chr3
MZIB.chr4
MZIB.chr5
MZIB.chr6
MZIB.chr7
MZIB.chr8
MZIB.chr9
MZIB.chr10
MZIB.chr11
MZIB.chr12
MZIB.chr13
MZIB.chr14
MZIB.chr15
MZIB.chr16
MZIB.chr17
MZIB.chr18
MZIB.chrX
* MMAR.chr1
MMAR.chr2
* MMAR.chr3
MMAR.chr4
* MMAR.chr5
* MMAR.chr6
* MMAR.chr7
* MMAR.chr8
* MMAR.chr9
MMAR.chr10
MMAR.chr11
MMAR.chr12
* MMAR.chr13
* MMAR.chr14
MMAR.chr15
MMAR.chr16
* MMAR.chr17
* MMAR.chr18
* MMAR.chrX
0 Mbp
50 Mbp
100 Mbp
150 Mbp
200 Mbp

# T84

1 Mbp sliding windows with a 100 kbp step

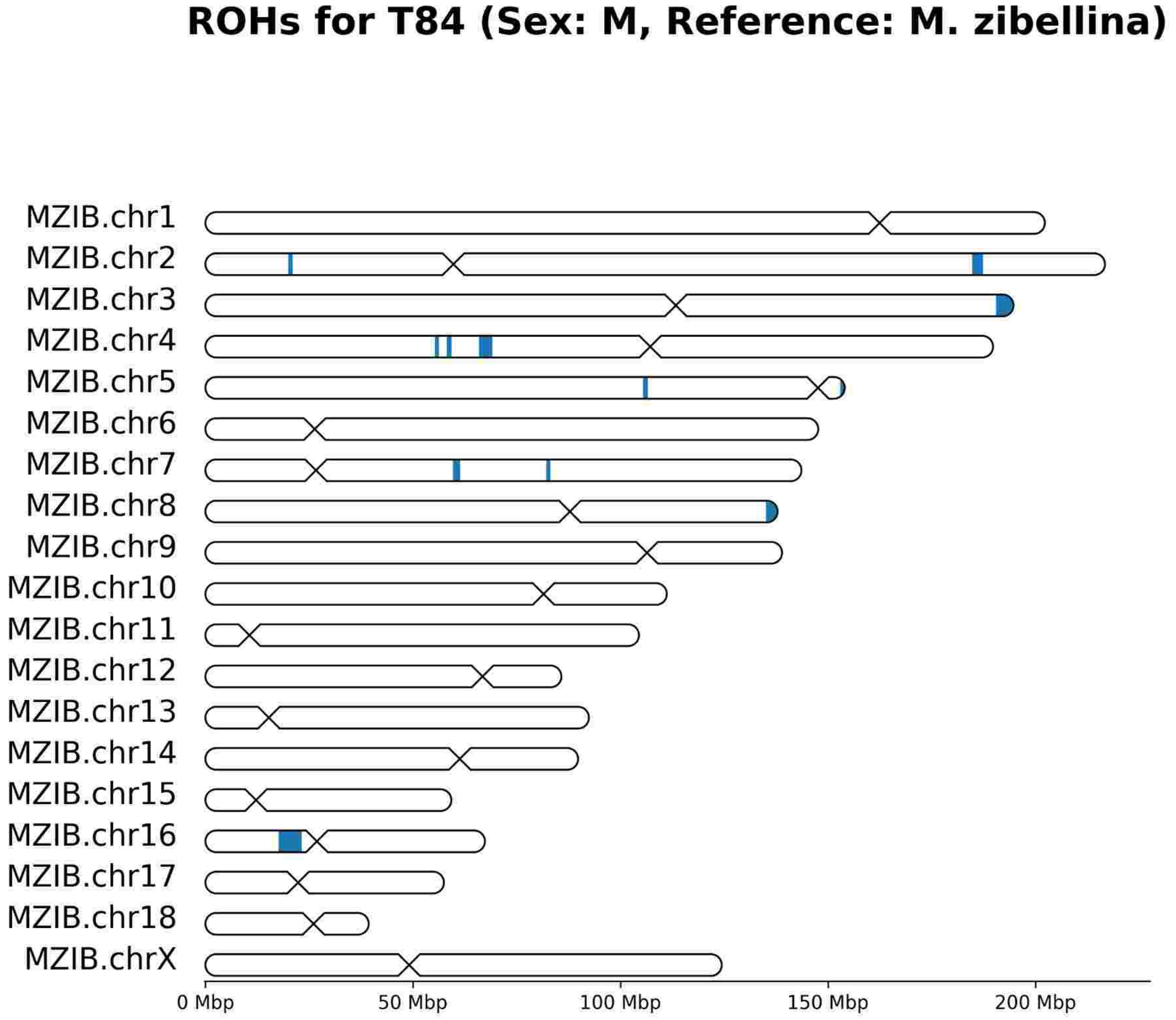


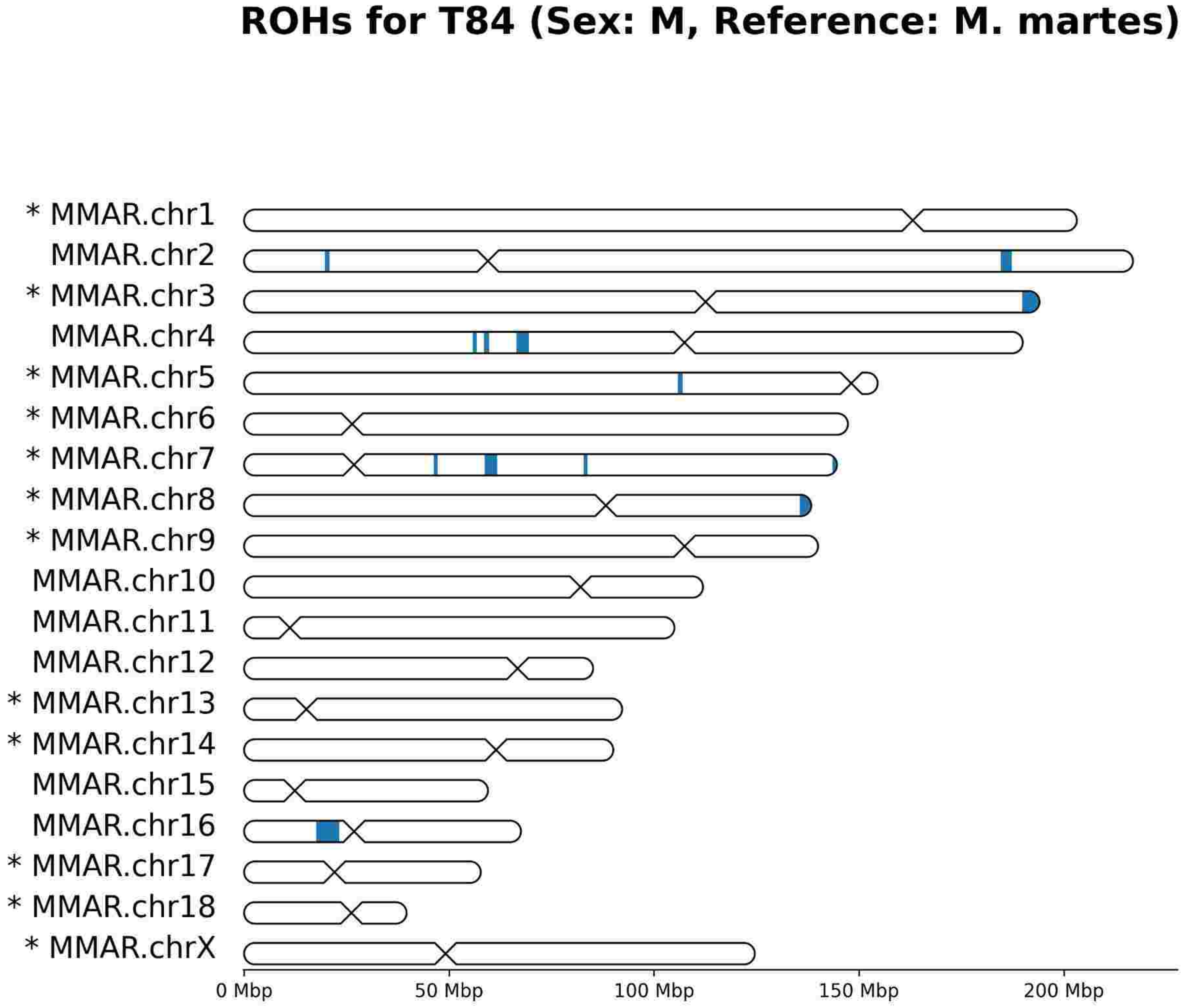


100 kbp sliding windows with a 10 kbp step

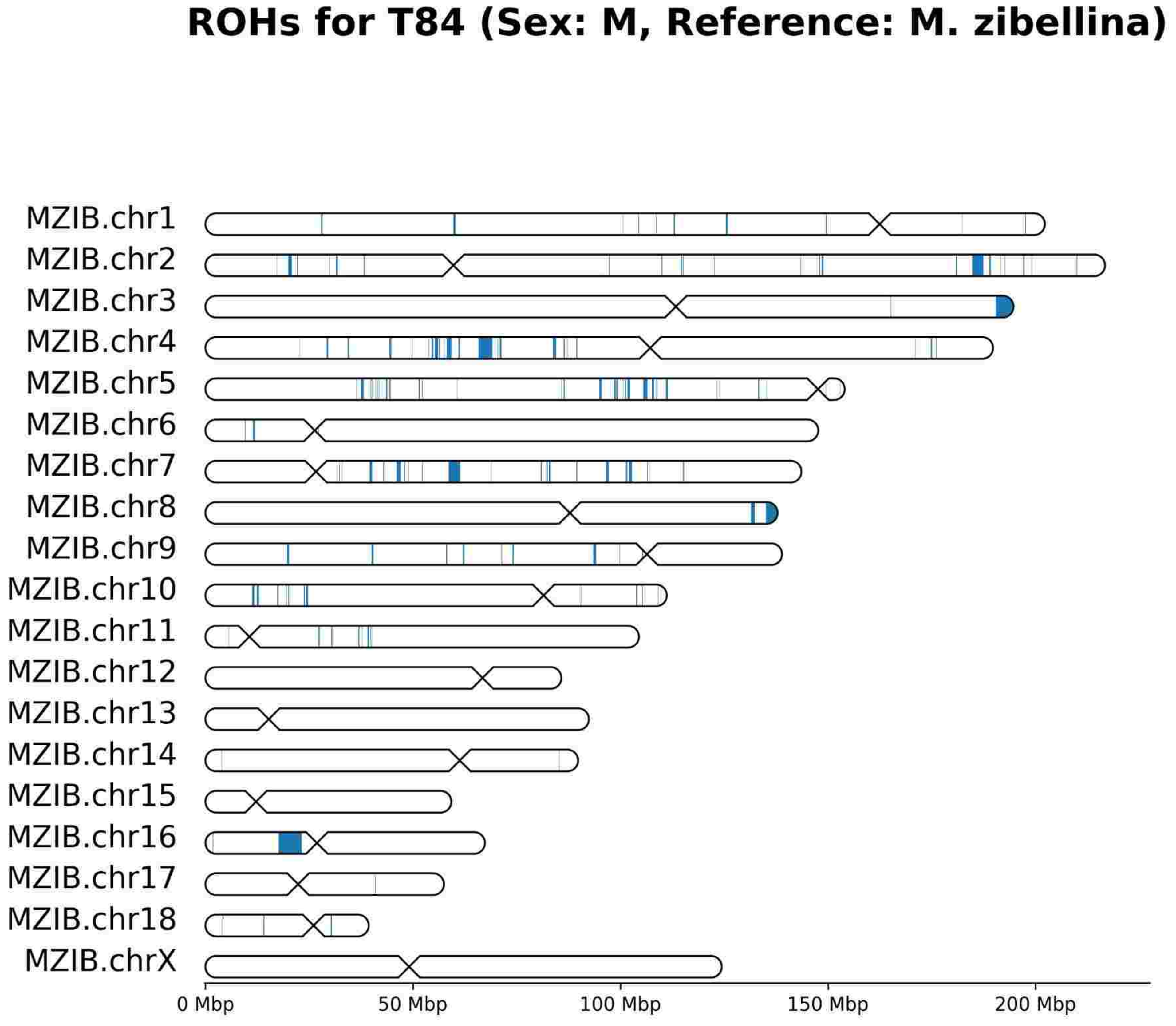


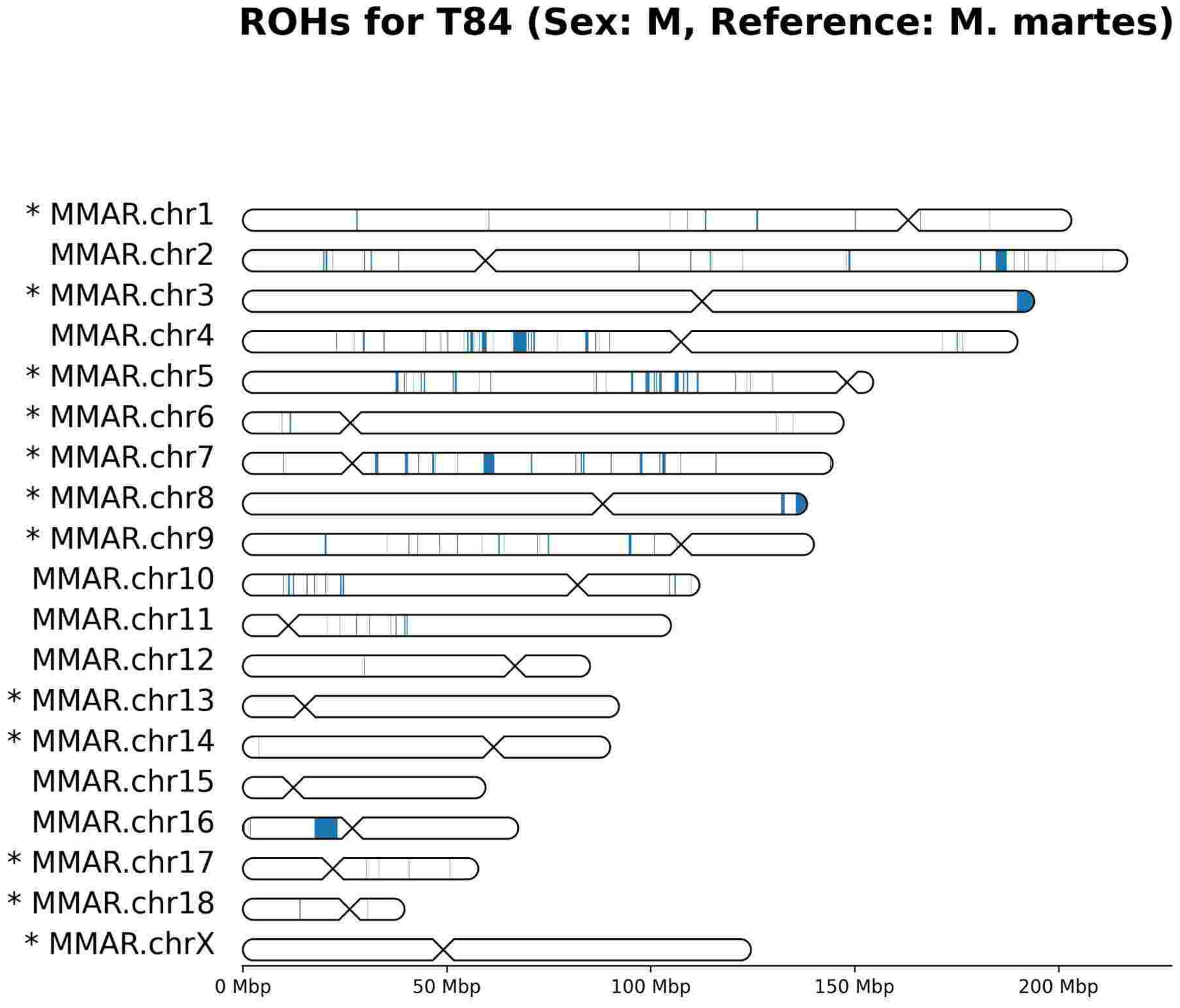

# T85

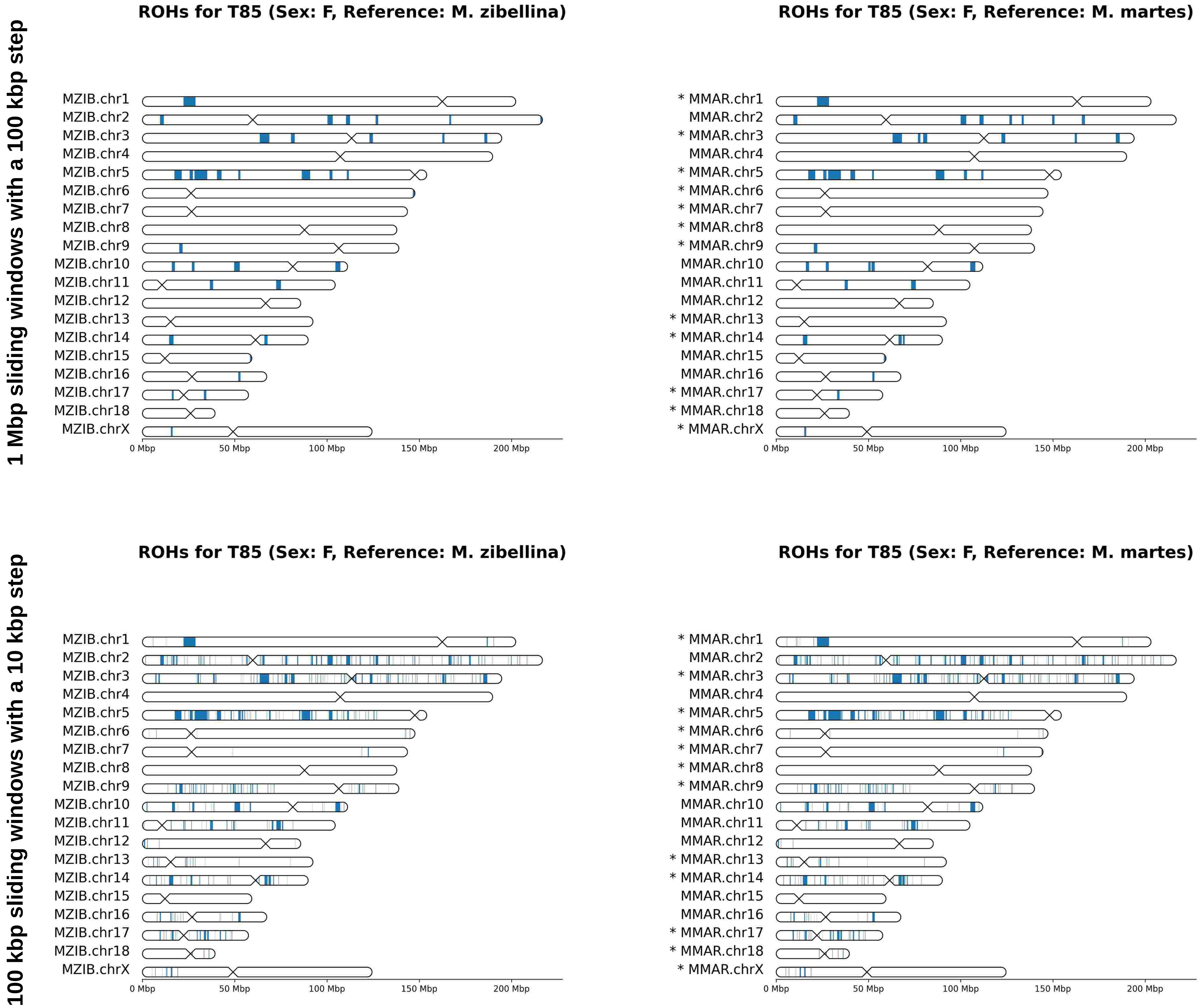
1 Mbp sliding windows with a 100 kbp step
100 kbp sliding windows with a 10 kbp step
ROHs for T85 (Sex: F, Reference: M. zibellina)
ROHs for T85 (Sex: F, Reference: M. martes)
MZIB.chr1
MZIB.chr2
MZIB.chr3
MZIB.chr4
MZIB.chr5
MZIB.chr6
MZIB.chr7
MZIB.chr8
MZIB.chr9
MZIB.chr10
MZIB.chr11
MZIB.chr12
MZIB.chr13
MZIB.chr14
MZIB.chr15
MZIB.chr16
MZIB.chr17
MZIB.chr18
MZIB.chrX
* MMAR.chr1
MMAR.chr2
* MMAR.chr3
MMAR.chr4
* MMAR.chr5
* MMAR.chr6
* MMAR.chr7
* MMAR.chr8
* MMAR.chr9
MMAR.chr10
MMAR.chr11
MMAR.chr12
* MMAR.chr13
* MMAR.chr14
MMAR.chr15
MMAR.chr16
* MMAR.chr17
* MMAR.chr18
* MMAR.chrX
0 Mbp
50 Mbp
100 Mbp
150 Mbp
200 Mbp

# T86

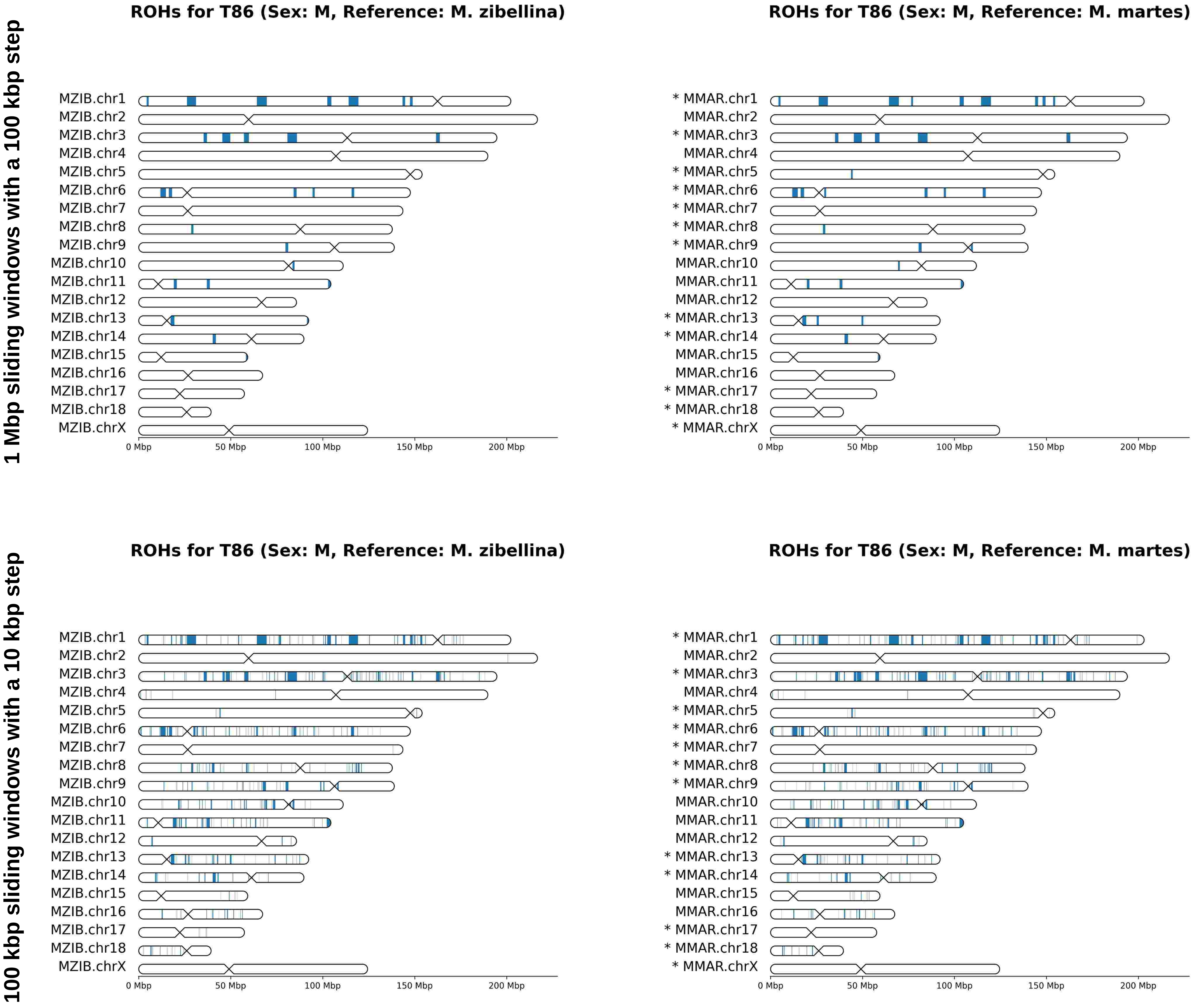

1 Mbp sliding windows with a 100 kbp step
ROHs for T86 (Sex: M, Reference: M. zibellina)
MZIB.chr1
MZIB.chr2
MZIB.chr3
MZIB.chr4
MZIB.chr5
MZIB.chr6
MZIB.chr7
MZIB.chr8
MZIB.chr9
MZIB.chr10
MZIB.chr11
MZIB.chr12
MZIB.chr13
MZIB.chr14
MZIB.chr15
MZIB.chr16
MZIB.chr17
MZIB.chr18
MZIB.chrX
0 Mbp
50 Mbp
100 Mbp
150 Mbp
200 Mbp
ROHs for T86 (Sex: M, Reference: M. martes)
* MMAR.chr1
MMAR.chr2
* MMAR.chr3
MMAR.chr4
* MMAR.chr5
* MMAR.chr6
* MMAR.chr7
* MMAR.chr8
* MMAR.chr9
MMAR.chr10
MMAR.chr11
MMAR.chr12
* MMAR.chr13
* MMAR.chr14
MMAR.chr15
MMAR.chr16
* MMAR.chr17
* MMAR.chr18
* MMAR.chrX
0 Mbp
50 Mbp
100 Mbp
150 Mbp
200 Mbp
100 kbp sliding windows with a 10 kbp step
ROHs for T86 (Sex: M, Reference: M. zibellina)
MZIB.chr1
MZIB.chr2
MZIB.chr3
MZIB.chr4
MZIB.chr5
MZIB.chr6
MZIB.chr7
MZIB.chr8
MZIB.chr9
MZIB.chr10
MZIB.chr11
MZIB.chr12
MZIB.chr13
MZIB.chr14
MZIB.chr15
MZIB.chr16
MZIB.chr17
MZIB.chr18
MZIB.chrX
0 Mbp
50 Mbp
100 Mbp
150 Mbp
200 Mbp
ROHs for T86 (Sex: M, Reference: M. martes)
* MMAR.chr1
MMAR.chr2
* MMAR.chr3
MMAR.chr4
* MMAR.chr5
* MMAR.chr6
* MMAR.chr7
* MMAR.chr8
* MMAR.chr9
MMAR.chr10
MMAR.chr11
MMAR.chr12
* MMAR.chr13
* MMAR.chr14
MMAR.chr15
MMAR.chr16
* MMAR.chr17
* MMAR.chr18
* MMAR.chrX
0 Mbp
50 Mbp
100 Mbp
150 Mbp
200 Mbp

# T87

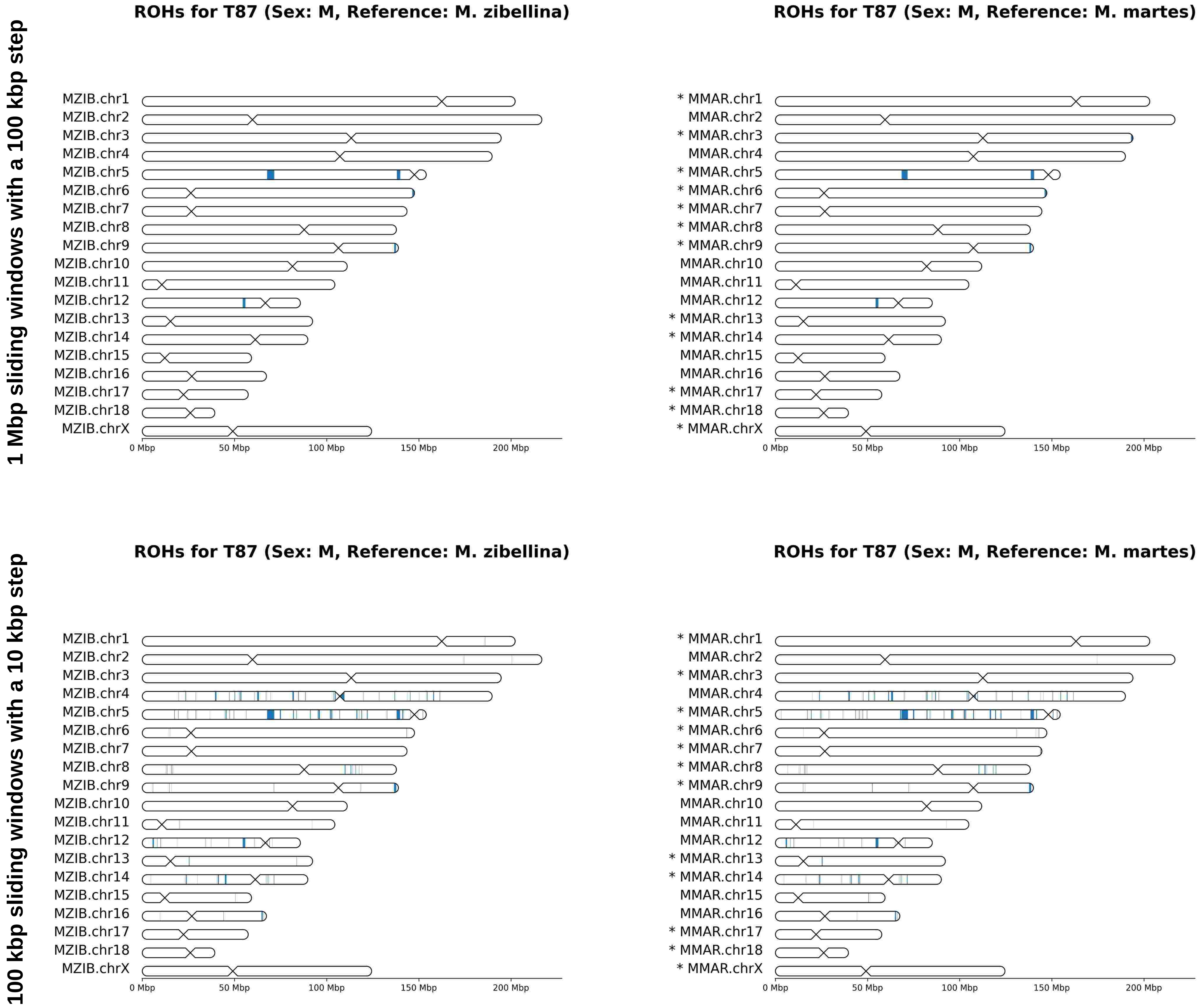
1 Mbp sliding windows with a 100 kbp step
100 kbp sliding windows with a 10 kbp step
ROHs for T87 (Sex: M, Reference: M. zibellina)
ROHs for T87 (Sex: M, Reference: M. martes)
MZIB.chr1
MZIB.chr2
MZIB.chr3
MZIB.chr4
MZIB.chr5
MZIB.chr6
MZIB.chr7
MZIB.chr8
MZIB.chr9
MZIB.chr10
MZIB.chr11
MZIB.chr12
MZIB.chr13
MZIB.chr14
MZIB.chr15
MZIB.chr16
MZIB.chr17
MZIB.chr18
MZIB.chrX
* MMAR.chr1
MMAR.chr2
* MMAR.chr3
MMAR.chr4
* MMAR.chr5
* MMAR.chr6
* MMAR.chr7
* MMAR.chr8
* MMAR.chr9
MMAR.chr10
MMAR.chr11
MMAR.chr12
* MMAR.chr13
* MMAR.chr14
MMAR.chr15
MMAR.chr16
* MMAR.chr17
* MMAR.chr18
* MMAR.chrX
0 Mbp
50 Mbp
100 Mbp
150 Mbp
200 Mbp

# T90

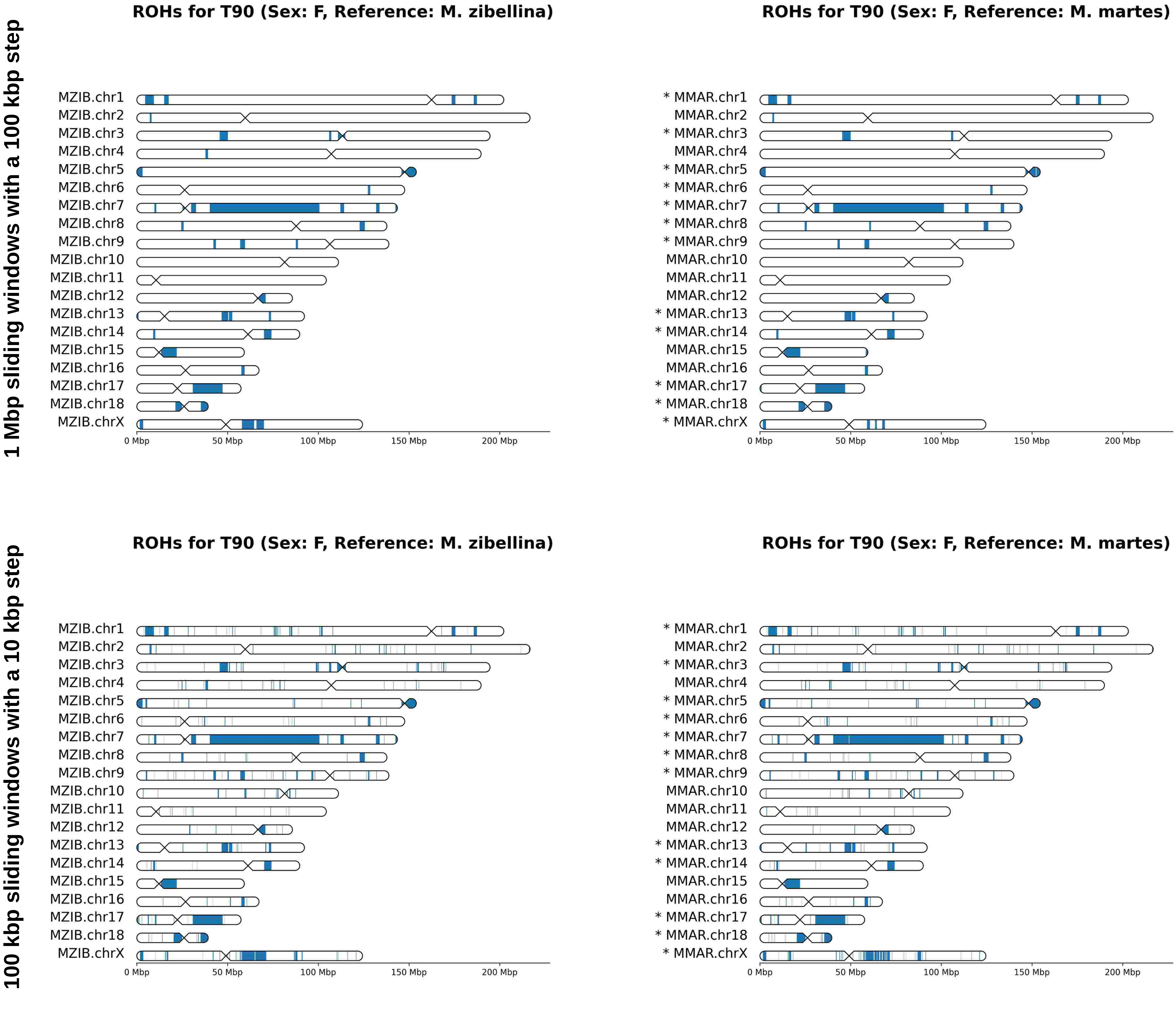
1 Mbp sliding windows with a 100 kbp step
100 kbp sliding windows with a 10 kbp step
ROHs for T90 (Sex: F, Reference: M. zibellina)
ROHs for T90 (Sex: F, Reference: M. martes)
MZIB.chr1
MZIB.chr2
MZIB.chr3
MZIB.chr4
MZIB.chr5
MZIB.chr6
MZIB.chr7
MZIB.chr8
MZIB.chr9
MZIB.chr10
MZIB.chr11
MZIB.chr12
MZIB.chr13
MZIB.chr14
MZIB.chr15
MZIB.chr16
MZIB.chr17
MZIB.chr18
MZIB.chrX
* MMAR.chr1
MMAR.chr2
* MMAR.chr3
MMAR.chr4
* MMAR.chr5
* MMAR.chr6
* MMAR.chr7
* MMAR.chr8
* MMAR.chr9
MMAR.chr10
MMAR.chr11
MMAR.chr12
* MMAR.chr13
* MMAR.chr14
MMAR.chr15
MMAR.chr16
* MMAR.chr17
* MMAR.chr18
* MMAR.chrX
0 Mbp
50 Mbp
100 Mbp
150 Mbp
200 Mbp

# T104

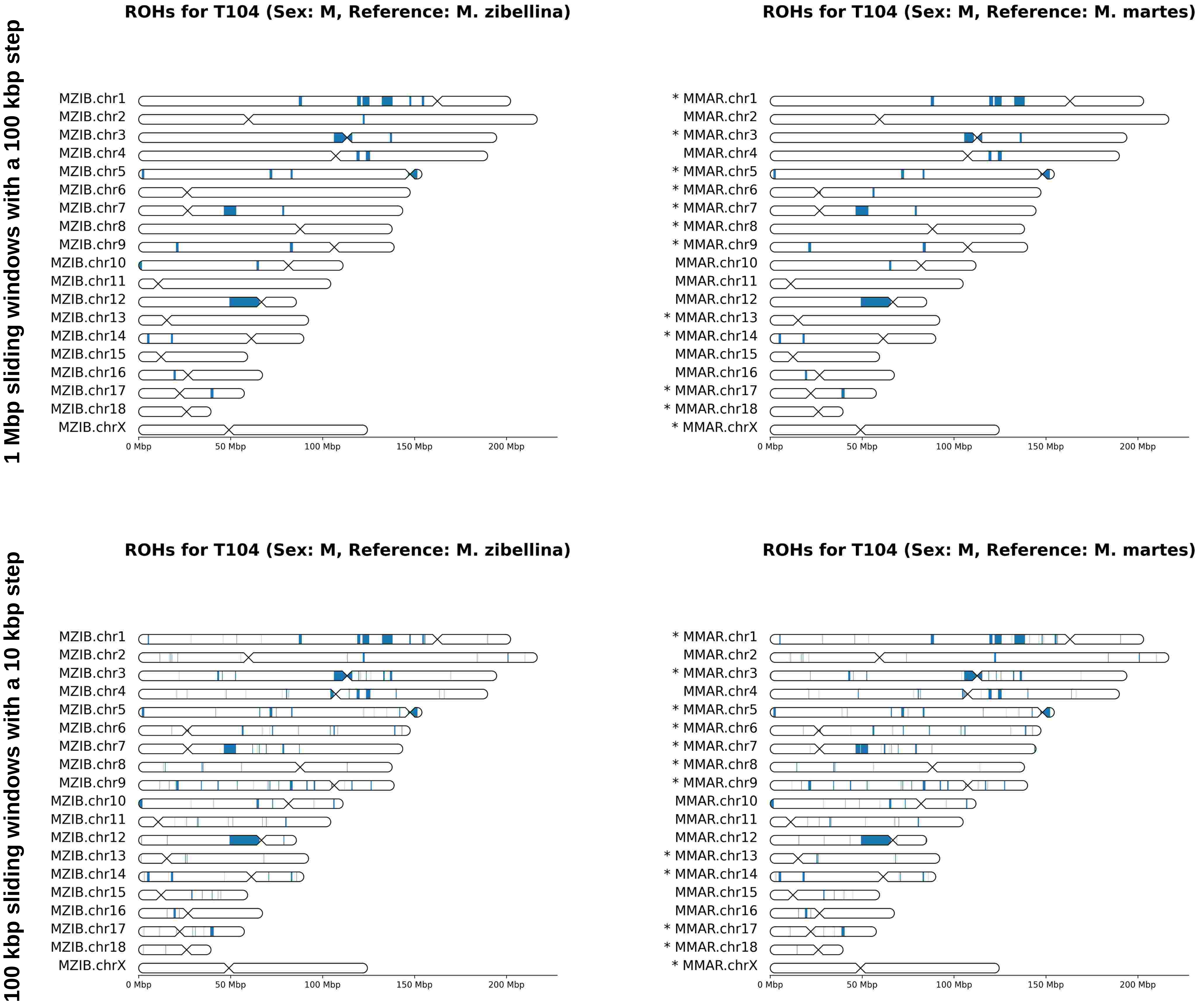
1 Mbp sliding windows with a 100 kbp step
ROHs for T104 (Sex: M, Reference: M. zibellina)
MZIB.chr1
MZIB.chr2
MZIB.chr3
MZIB.chr4
MZIB.chr5
MZIB.chr6
MZIB.chr7
MZIB.chr8
MZIB.chr9
MZIB.chr10
MZIB.chr11
MZIB.chr12
MZIB.chr13
MZIB.chr14
MZIB.chr15
MZIB.chr16
MZIB.chr17
MZIB.chr18
MZIB.chrX
0 Mbp
50 Mbp
100 Mbp
150 Mbp
200 Mbp
ROHs for T104 (Sex: M, Reference: M. martes)
* MMAR.chr1
MMAR.chr2
* MMAR.chr3
MMAR.chr4
* MMAR.chr5
* MMAR.chr6
* MMAR.chr7
* MMAR.chr8
* MMAR.chr9
MMAR.chr10
MMAR.chr11
MMAR.chr12
* MMAR.chr13
* MMAR.chr14
MMAR.chr15
MMAR.chr16
* MMAR.chr17
* MMAR.chr18
* MMAR.chrX
0 Mbp
50 Mbp
100 Mbp
150 Mbp
200 Mbp
100 kbp sliding windows with a 10 kbp step
ROHs for T104 (Sex: M, Reference: M. zibellina)
MZIB.chr1
MZIB.chr2
MZIB.chr3
MZIB.chr4
MZIB.chr5
MZIB.chr6
MZIB.chr7
MZIB.chr8
MZIB.chr9
MZIB.chr10
MZIB.chr11
MZIB.chr12
MZIB.chr13
MZIB.chr14
MZIB.chr15
MZIB.chr16
MZIB.chr17
MZIB.chr18
MZIB.chrX
0 Mbp
50 Mbp
100 Mbp
150 Mbp
200 Mbp
ROHs for T104 (Sex: M, Reference: M. martes)
* MMAR.chr1
MMAR.chr2
* MMAR.chr3
MMAR.chr4
* MMAR.chr5
* MMAR.chr6
* MMAR.chr7
* MMAR.chr8
* MMAR.chr9
MMAR.chr10
MMAR.chr11
MMAR.chr12
* MMAR.chr13
* MMAR.chr14
MMAR.chr15
MMAR.chr16
* MMAR.chr17
* MMAR.chr18
* MMAR.chrX
0 Mbp
50 Mbp
100 Mbp
150 Mbp
200 Mbp

# T118

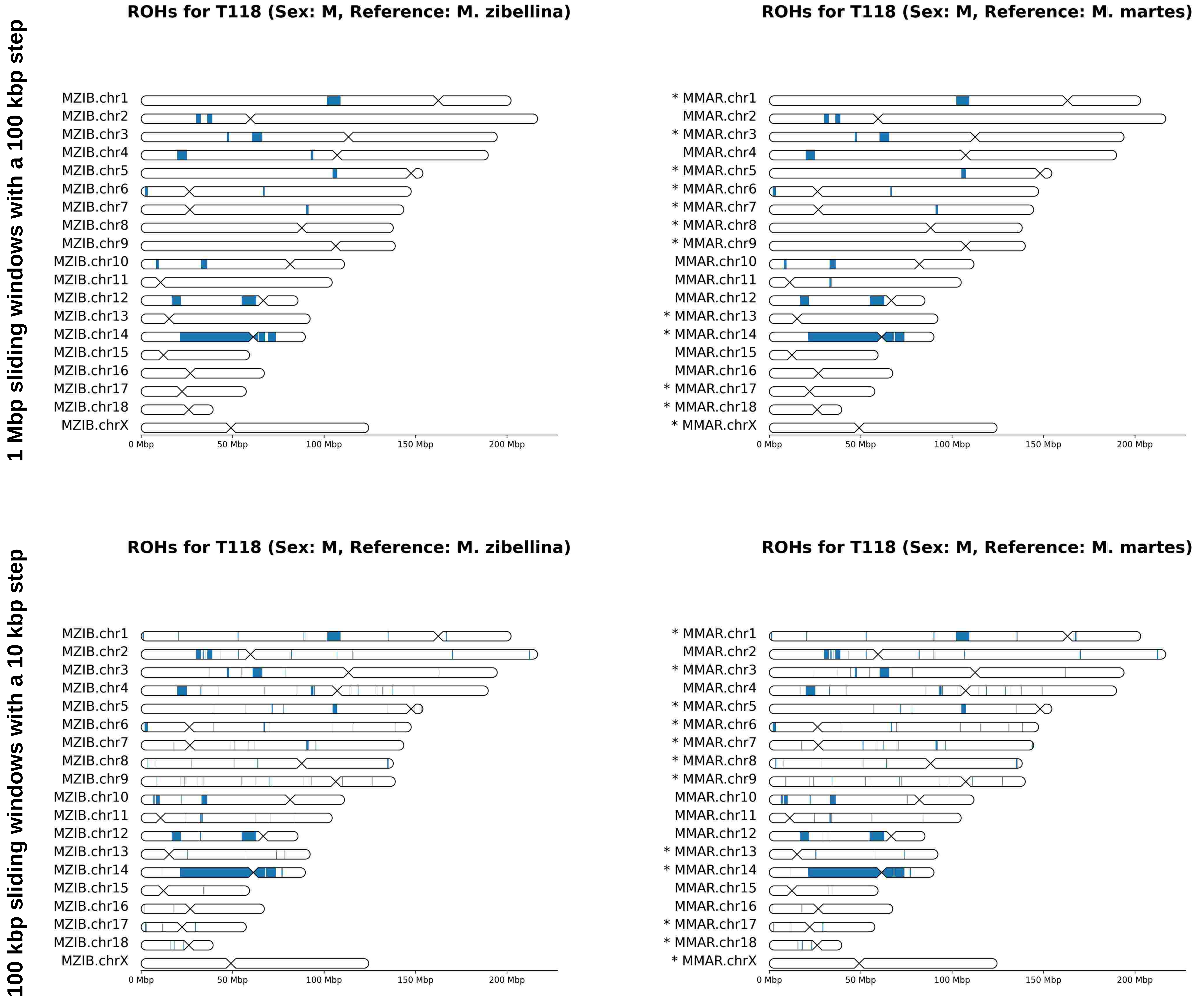
1 Mbp sliding windows with a 100 kbp step
ROHs for T118 (Sex: M, Reference: M. zibellina)
MZIB.chr1
MZIB.chr2
MZIB.chr3
MZIB.chr4
MZIB.chr5
MZIB.chr6
MZIB.chr7
MZIB.chr8
MZIB.chr9
MZIB.chr10
MZIB.chr11
MZIB.chr12
MZIB.chr13
MZIB.chr14
MZIB.chr15
MZIB.chr16
MZIB.chr17
MZIB.chr18
MZIB.chrX
0 Mbp
50 Mbp
100 Mbp
150 Mbp
200 Mbp
ROHs for T118 (Sex: M, Reference: M. martes)
* MMAR.chr1
MMAR.chr2
* MMAR.chr3
MMAR.chr4
* MMAR.chr5
* MMAR.chr6
* MMAR.chr7
* MMAR.chr8
* MMAR.chr9
MMAR.chr10
MMAR.chr11
MMAR.chr12
* MMAR.chr13
* MMAR.chr14
MMAR.chr15
MMAR.chr16
* MMAR.chr17
* MMAR.chr18
* MMAR.chrX
0 Mbp
50 Mbp
100 Mbp
150 Mbp
200 Mbp
100 kbp sliding windows with a 10 kbp step
ROHs for T118 (Sex: M, Reference: M. zibellina)
MZIB.chr1
MZIB.chr2
MZIB.chr3
MZIB.chr4
MZIB.chr5
MZIB.chr6
MZIB.chr7
MZIB.chr8
MZIB.chr9
MZIB.chr10
MZIB.chr11
MZIB.chr12
MZIB.chr13
MZIB.chr14
MZIB.chr15
MZIB.chr16
MZIB.chr17
MZIB.chr18
MZIB.chrX
0 Mbp
50 Mbp
100 Mbp
150 Mbp
200 Mbp
ROHs for T118 (Sex: M, Reference: M. martes)
* MMAR.chr1
MMAR.chr2
* MMAR.chr3
MMAR.chr4
* MMAR.chr5
* MMAR.chr6
* MMAR.chr7
* MMAR.chr8
* MMAR.chr9
MMAR.chr10
MMAR.chr11
MMAR.chr12
* MMAR.chr13
* MMAR.chr14
MMAR.chr15
MMAR.chr16
* MMAR.chr17
* MMAR.chr18
* MMAR.chrX
0 Mbp
50 Mbp
100 Mbp
150 Mbp
200 Mbp

# T148

1 Mbp sliding windows with a 100 kbp step

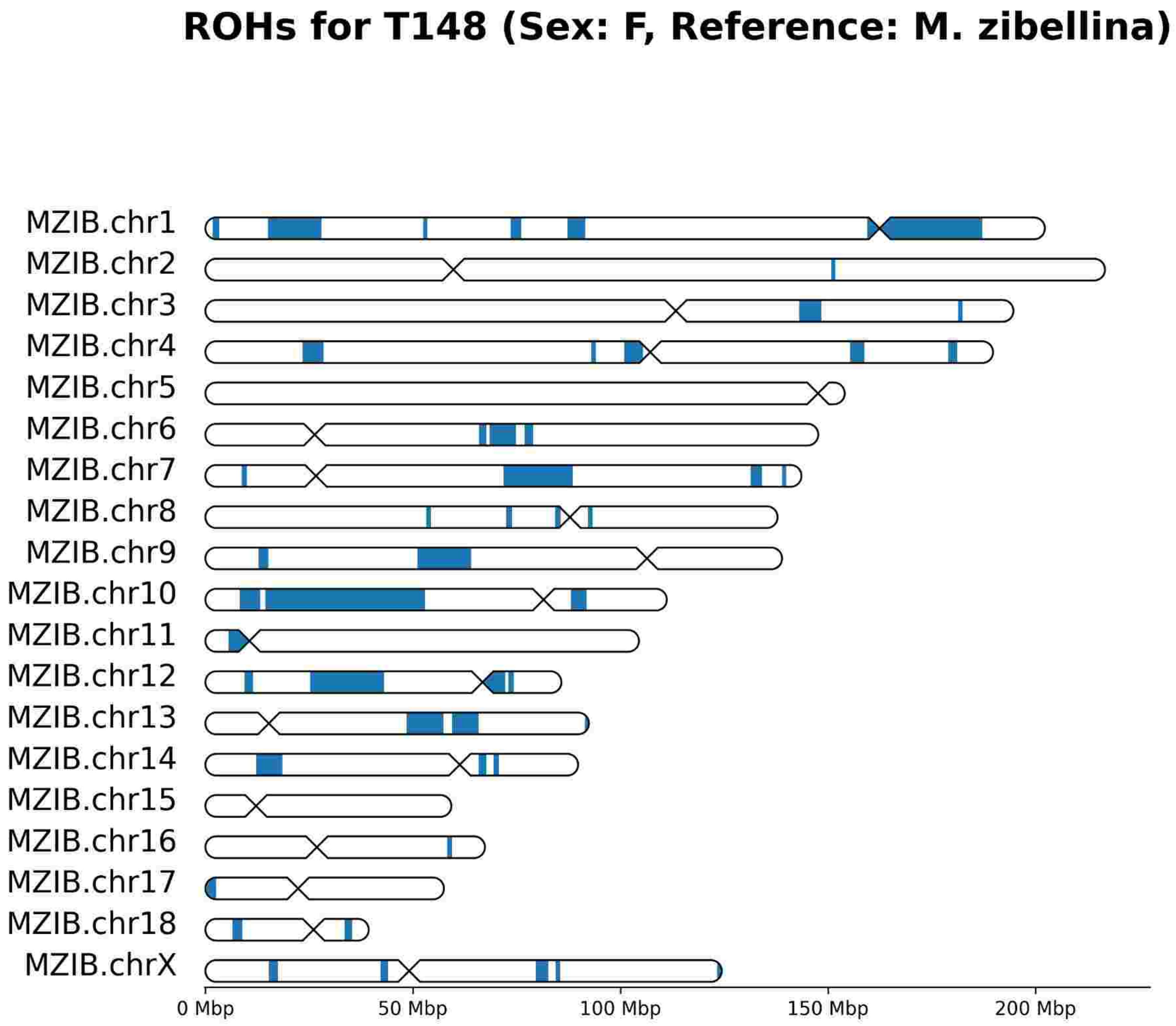


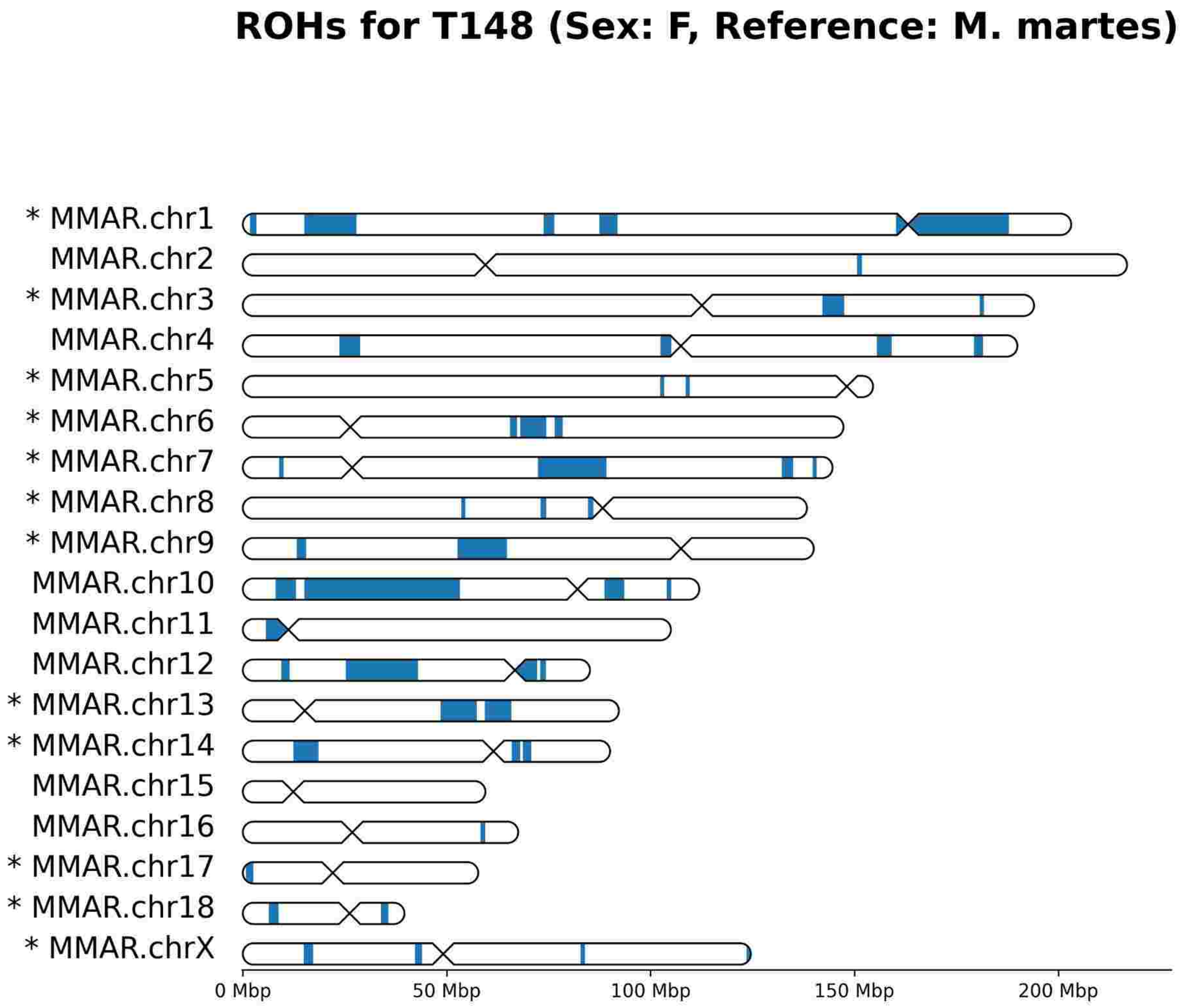


100 kbp sliding windows with a 10 kbp step

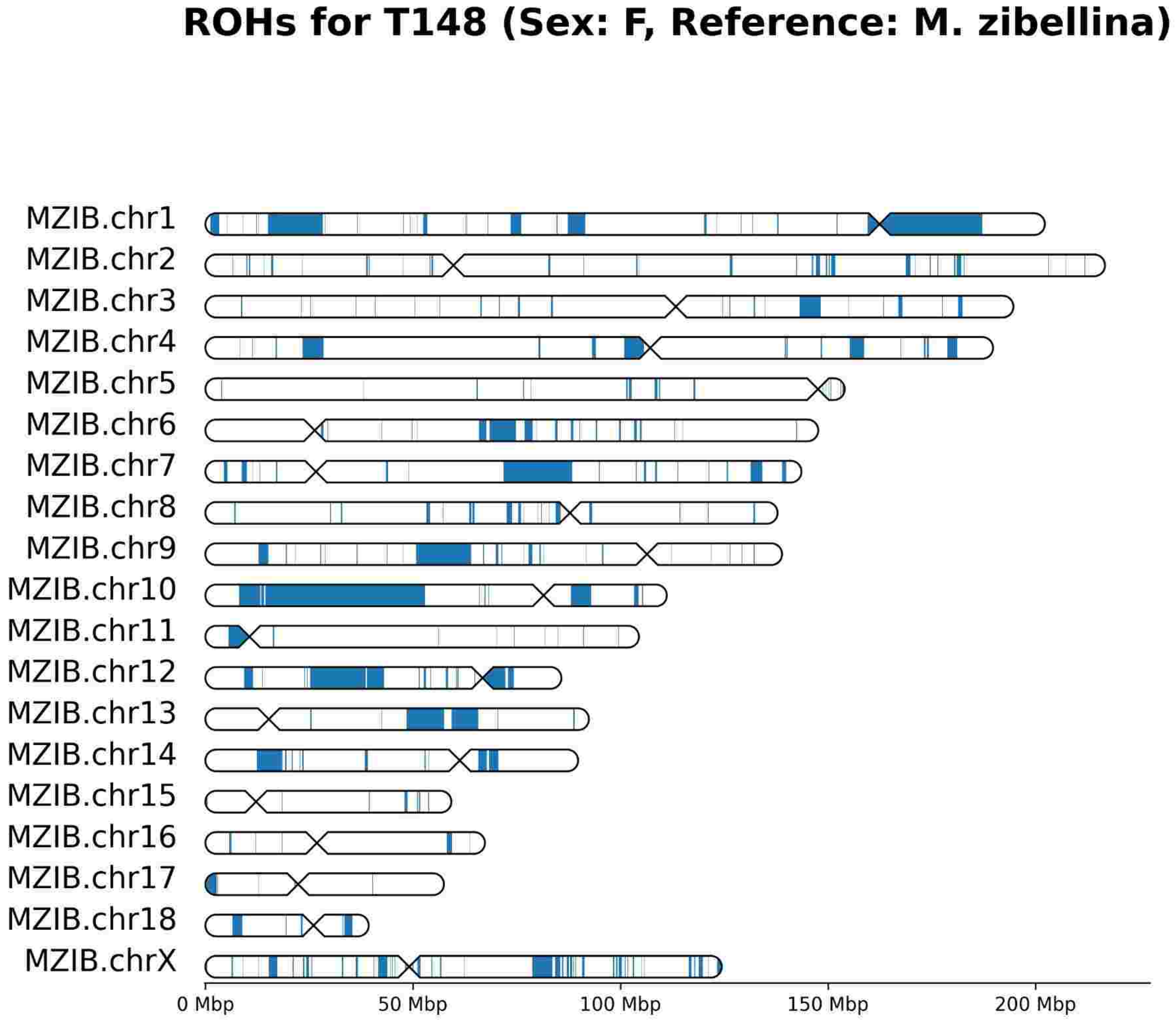


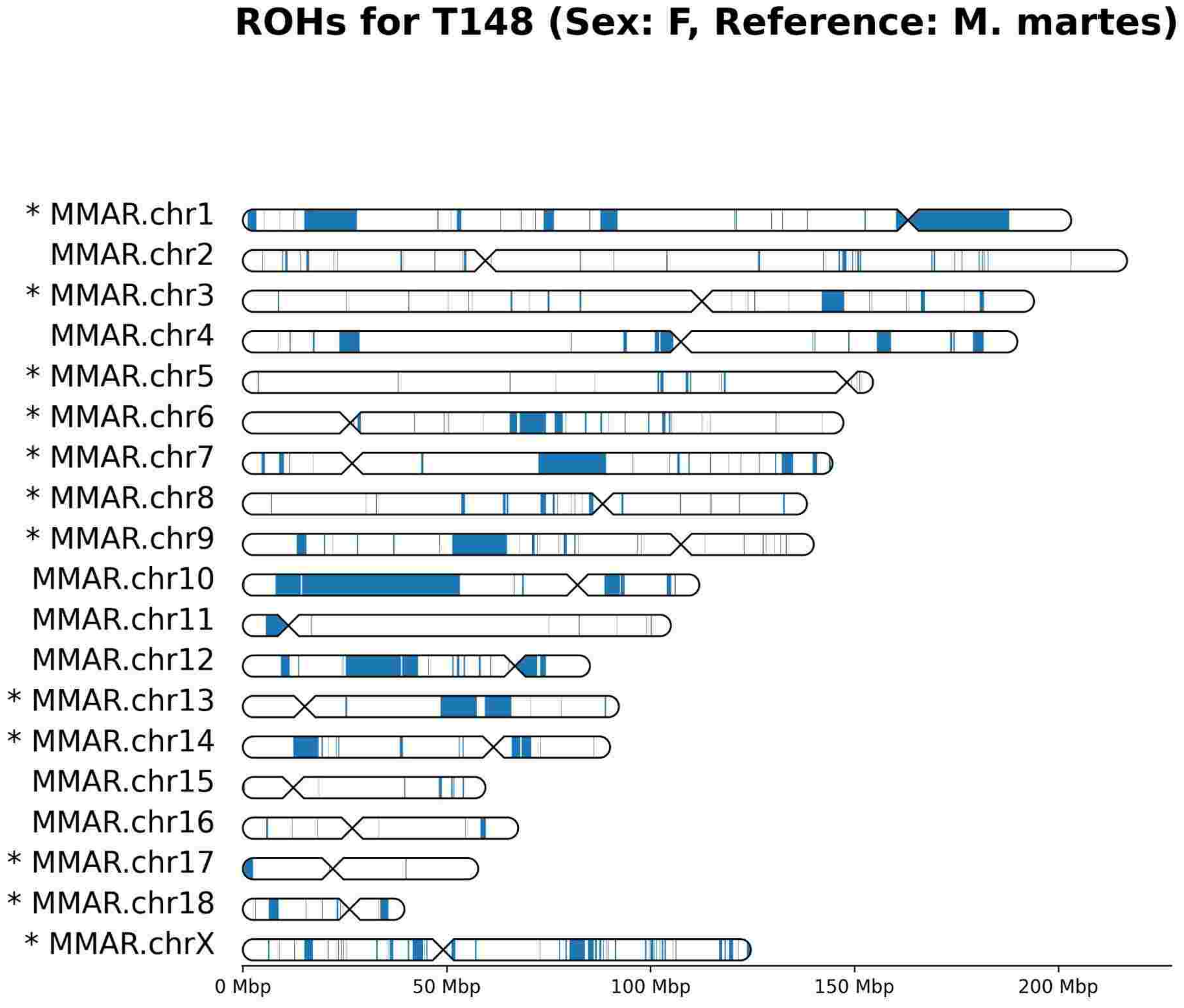

# T149

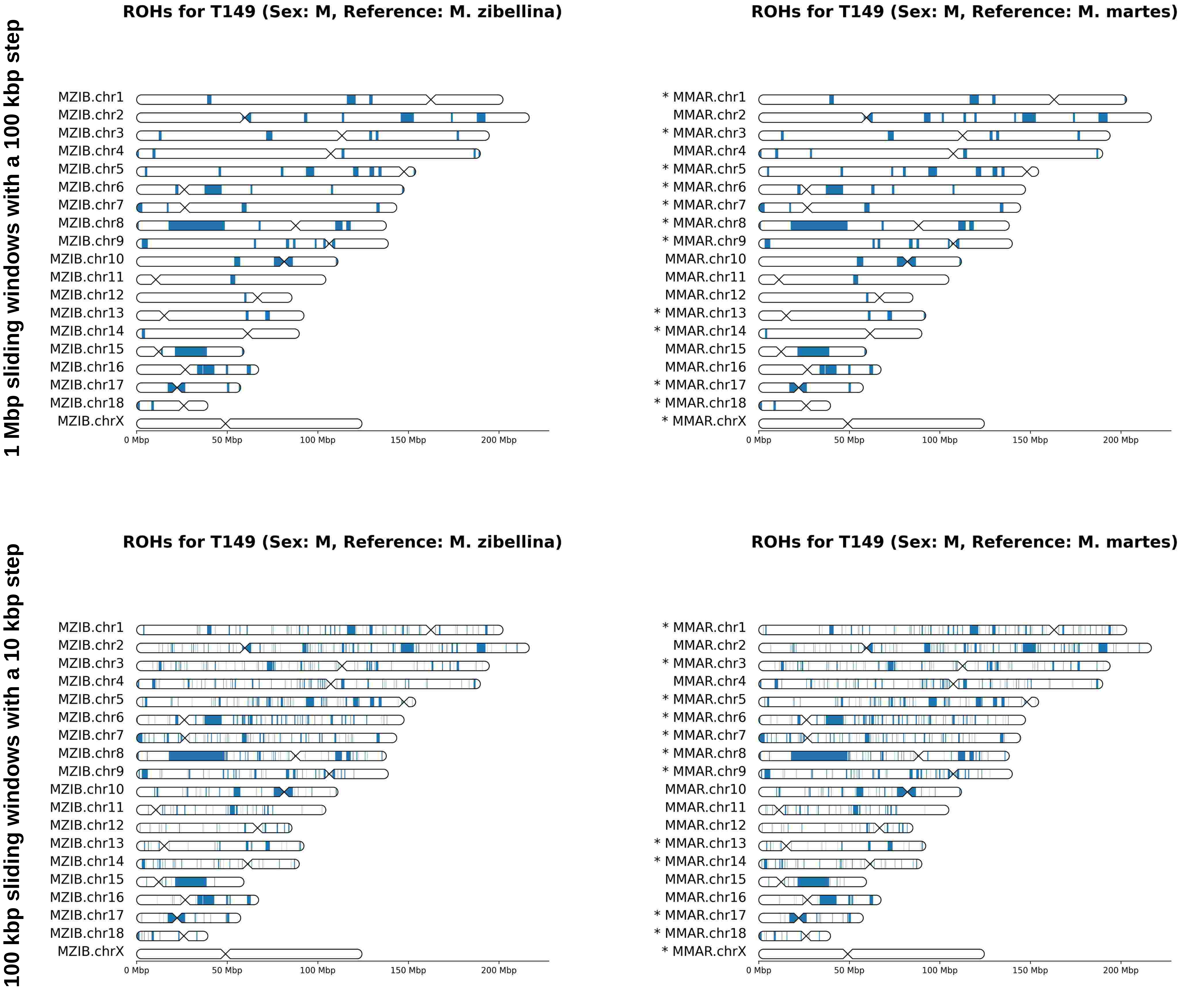

1 Mbp sliding windows with a 100 kbp step
100 kbp sliding windows with a 10 kbp step
ROHs for T149 (Sex: M, Reference: M. zibellina)
MZIB.chr1
MZIB.chr2
MZIB.chr3
MZIB.chr4
MZIB.chr5
MZIB.chr6
MZIB.chr7
MZIB.chr8
MZIB.chr9
MZIB.chr10
MZIB.chr11
MZIB.chr12
MZIB.chr13
MZIB.chr14
MZIB.chr15
MZIB.chr16
MZIB.chr17
MZIB.chr18
MZIB.chrX
0 Mbp
50 Mbp
100 Mbp
150 Mbp
200 Mbp
ROHs for T149 (Sex: M, Reference: M. martes)
* MMAR.chr1
MMAR.chr2
* MMAR.chr3
MMAR.chr4
* MMAR.chr5
* MMAR.chr6
* MMAR.chr7
* MMAR.chr8
* MMAR.chr9
MMAR.chr10
MMAR.chr11
MMAR.chr12
* MMAR.chr13
* MMAR.chr14
MMAR.chr15
MMAR.chr16
* MMAR.chr17
* MMAR.chr18
* MMAR.chrX
0 Mbp
50 Mbp
100 Mbp
150 Mbp
200 Mbp

# T150

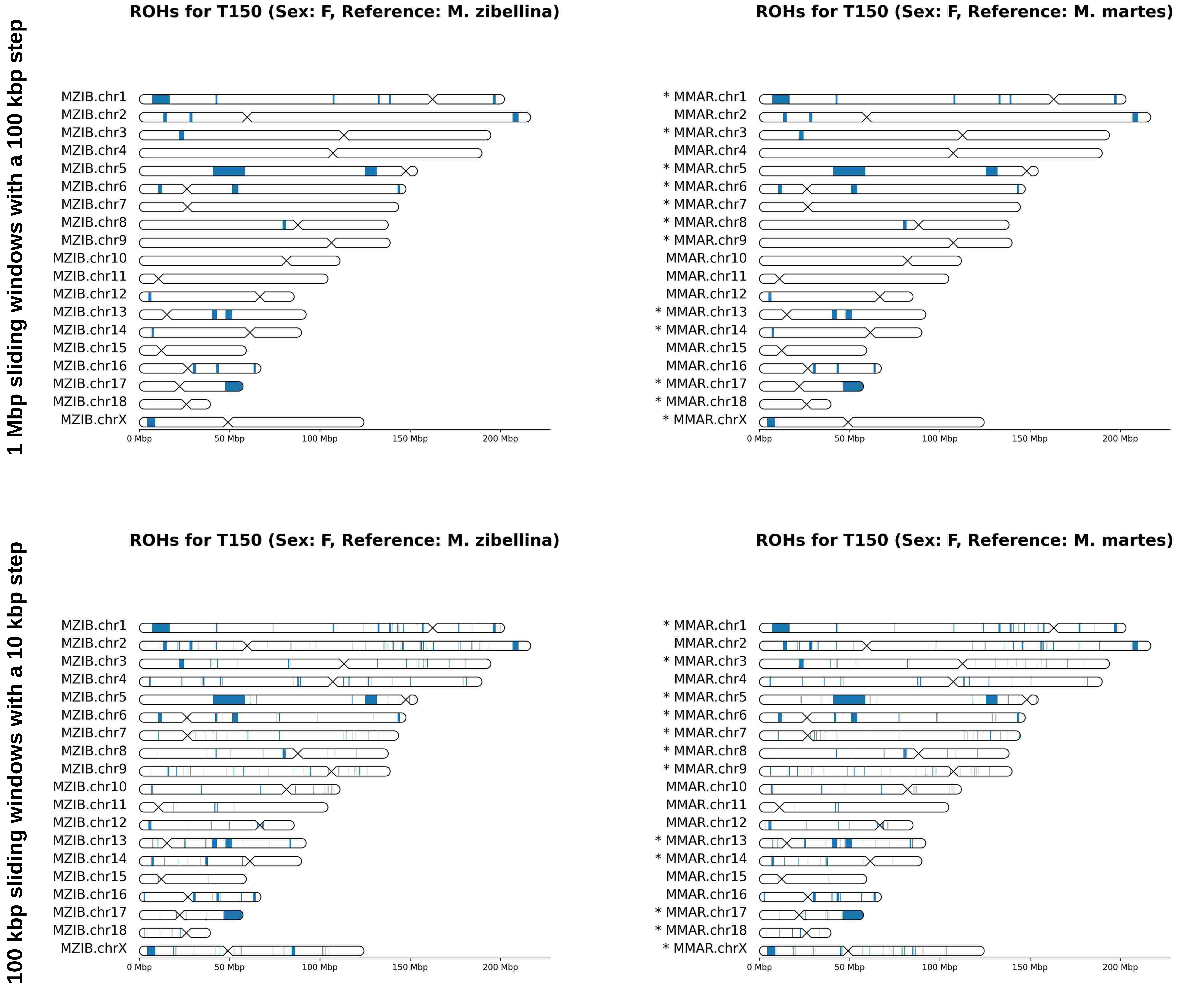
1 Mbp sliding windows with a 100 kbp step
ROHs for T150 (Sex: F, Reference: M. zibellina)
MZIB.chr1
MZIB.chr2
MZIB.chr3
MZIB.chr4
MZIB.chr5
MZIB.chr6
MZIB.chr7
MZIB.chr8
MZIB.chr9
MZIB.chr10
MZIB.chr11
MZIB.chr12
MZIB.chr13
MZIB.chr14
MZIB.chr15
MZIB.chr16
MZIB.chr17
MZIB.chr18
MZIB.chrX
0 Mbp
50 Mbp
100 Mbp
150 Mbp
200 Mbp
ROHs for T150 (Sex: F, Reference: M. martes)
* MMAR.chr1
MMAR.chr2
* MMAR.chr3
MMAR.chr4
* MMAR.chr5
* MMAR.chr6
* MMAR.chr7
* MMAR.chr8
* MMAR.chr9
MMAR.chr10
MMAR.chr11
MMAR.chr12
* MMAR.chr13
* MMAR.chr14
MMAR.chr15
MMAR.chr16
* MMAR.chr17
* MMAR.chr18
* MMAR.chrX
0 Mbp
50 Mbp
100 Mbp
150 Mbp
200 Mbp
100 kbp sliding windows with a 10 kbp step
ROHs for T150 (Sex: F, Reference: M. zibellina)
MZIB.chr1
MZIB.chr2
MZIB.chr3
MZIB.chr4
MZIB.chr5
MZIB.chr6
MZIB.chr7
MZIB.chr8
MZIB.chr9
MZIB.chr10
MZIB.chr11
MZIB.chr12
MZIB.chr13
MZIB.chr14
MZIB.chr15
MZIB.chr16
MZIB.chr17
MZIB.chr18
MZIB.chrX
0 Mbp
50 Mbp
100 Mbp
150 Mbp
200 Mbp
ROHs for T150 (Sex: F, Reference: M. martes)
* MMAR.chr1
MMAR.chr2
* MMAR.chr3
MMAR.chr4
* MMAR.chr5
* MMAR.chr6
* MMAR.chr7
* MMAR.chr8
* MMAR.chr9
MMAR.chr10
MMAR.chr11
MMAR.chr12
* MMAR.chr13
* MMAR.chr14
MMAR.chr15
MMAR.chr16
* MMAR.chr17
* MMAR.chr18
* MMAR.chrX
0 Mbp
50 Mbp
100 Mbp
150 Mbp
200 Mbp

# T151

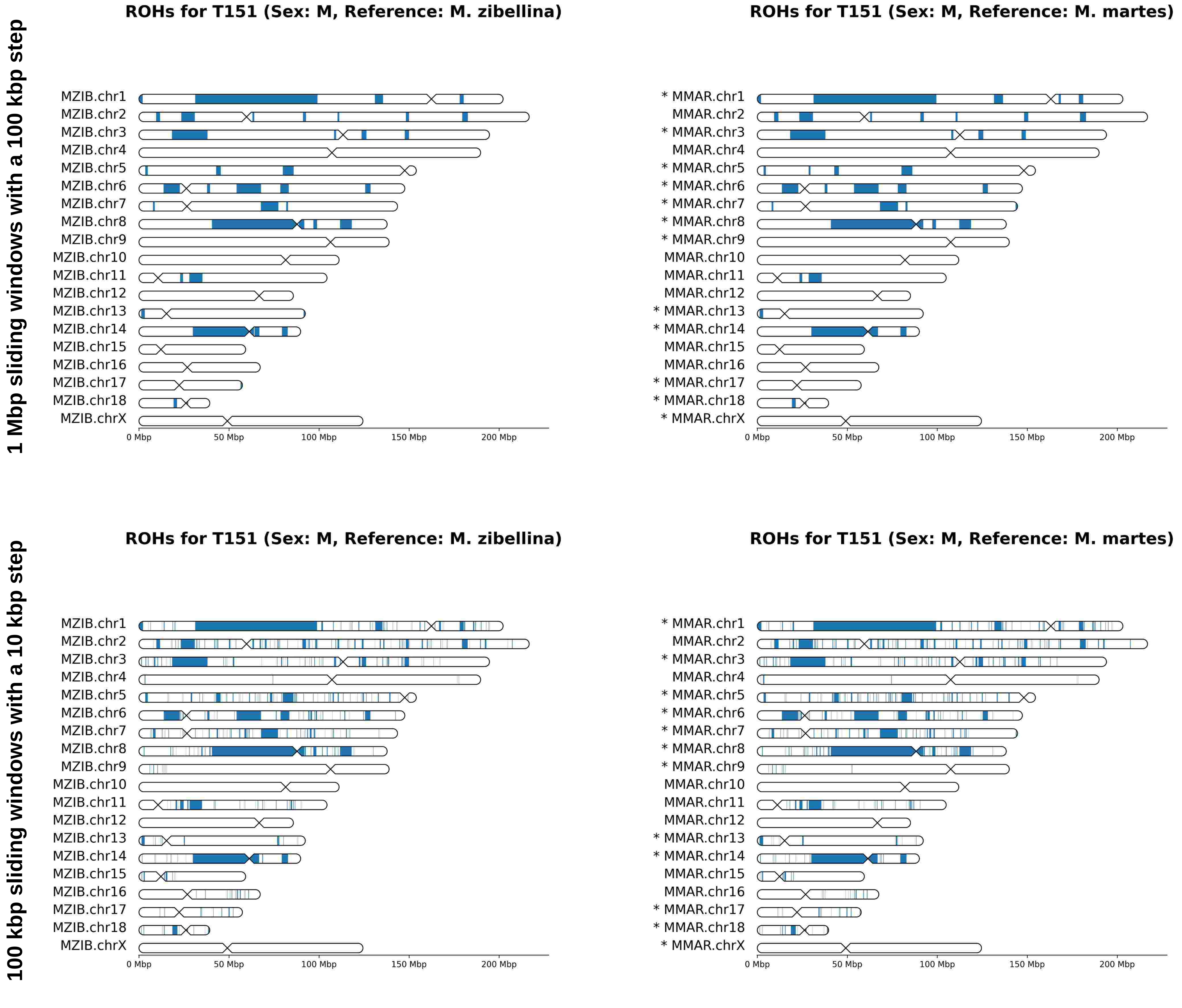

1 Mbp sliding windows with a 100 kbp step
ROHs for T151 (Sex: M, Reference: M. zibellina)
MZIB.chr1
MZIB.chr2
MZIB.chr3
MZIB.chr4
MZIB.chr5
MZIB.chr6
MZIB.chr7
MZIB.chr8
MZIB.chr9
MZIB.chr10
MZIB.chr11
MZIB.chr12
MZIB.chr13
MZIB.chr14
MZIB.chr15
MZIB.chr16
MZIB.chr17
MZIB.chr18
MZIB.chrX
0 Mbp
50 Mbp
100 Mbp
150 Mbp
200 Mbp
ROHs for T151 (Sex: M, Reference: M. martes)
* MMAR.chr1
MMAR.chr2
* MMAR.chr3
MMAR.chr4
* MMAR.chr5
* MMAR.chr6
* MMAR.chr7
* MMAR.chr8
* MMAR.chr9
MMAR.chr10
MMAR.chr11
MMAR.chr12
* MMAR.chr13
* MMAR.chr14
MMAR.chr15
MMAR.chr16
* MMAR.chr17
* MMAR.chr18
* MMAR.chrX
0 Mbp
50 Mbp
100 Mbp
150 Mbp
200 Mbp
100 kbp sliding windows with a 10 kbp step
ROHs for T151 (Sex: M, Reference: M. zibellina)
MZIB.chr1
MZIB.chr2
MZIB.chr3
MZIB.chr4
MZIB.chr5
MZIB.chr6
MZIB.chr7
MZIB.chr8
MZIB.chr9
MZIB.chr10
MZIB.chr11
MZIB.chr12
MZIB.chr13
MZIB.chr14
MZIB.chr15
MZIB.chr16
MZIB.chr17
MZIB.chr18
MZIB.chrX
0 Mbp
50 Mbp
100 Mbp
150 Mbp
200 Mbp
ROHs for T151 (Sex: M, Reference: M. martes)
* MMAR.chr1
MMAR.chr2
* MMAR.chr3
MMAR.chr4
* MMAR.chr5
* MMAR.chr6
* MMAR.chr7
* MMAR.chr8
* MMAR.chr9
MMAR.chr10
MMAR.chr11
MMAR.chr12
* MMAR.chr13
* MMAR.chr14
MMAR.chr15
MMAR.chr16
* MMAR.chr17
* MMAR.chr18
* MMAR.chrX
0 Mbp
50 Mbp
100 Mbp
150 Mbp
200 Mbp

# T194

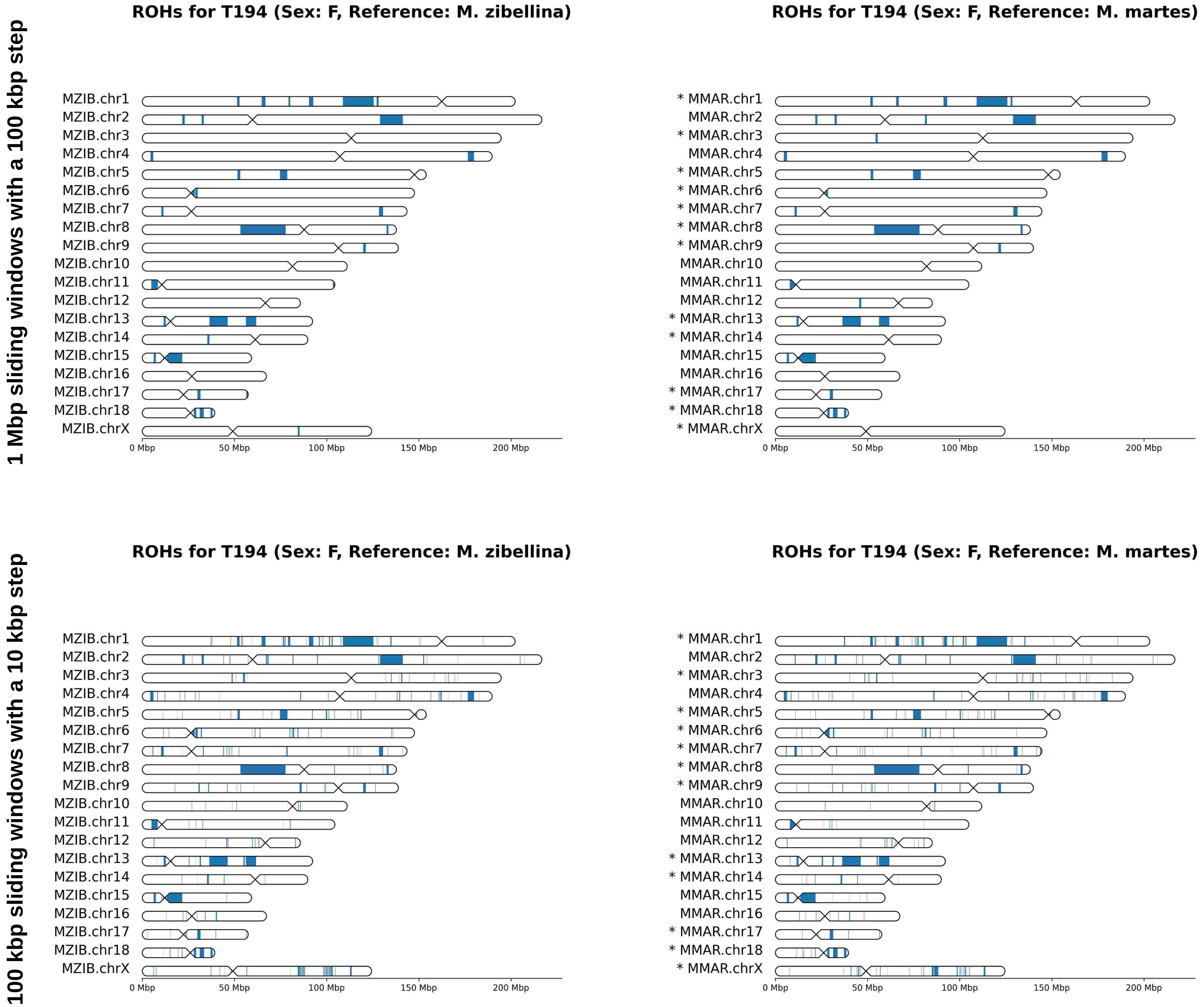
1 Mbp sliding windows with a 100 kbp step
100 kbp sliding windows with a 10 kbp step
ROHs for T194 (Sex: F, Reference: M. zibellina)
ROHs for T194 (Sex: F, Reference: M. martes)
MZIB.chr1
MZIB.chr2
MZIB.chr3
MZIB.chr4
MZIB.chr5
MZIB.chr6
MZIB.chr7
MZIB.chr8
MZIB.chr9
MZIB.chr10
MZIB.chr11
MZIB.chr12
MZIB.chr13
MZIB.chr14
MZIB.chr15
MZIB.chr16
MZIB.chr17
MZIB.chr18
MZIB.chrX
* MMAR.chr1
MMAR.chr2
* MMAR.chr3
MMAR.chr4
* MMAR.chr5
* MMAR.chr6
* MMAR.chr7
* MMAR.chr8
* MMAR.chr9
MMAR.chr10
MMAR.chr11
MMAR.chr12
* MMAR.chr13
* MMAR.chr14
MMAR.chr15
MMAR.chr16
* MMAR.chr17
* MMAR.chr18
* MMAR.chrX
0 Mbp
50 Mbp
100 Mbp
150 Mbp
200 Mbp
